# Predicting Composition of Genetic Circuits with Resource Competition: Demand and Sensitivity

**DOI:** 10.1101/2021.05.26.445862

**Authors:** Cameron D. McBride, Domitilla Del Vecchio

**Affiliations:** Department of Mechanical Engineering, MIT, 77 Massachusetts Ave, Cambridge, MA, USA

**Keywords:** Resource Competition, Modular Composition, Experiment Design

## Abstract

The design of genetic circuits typically relies on characterization of constituent modules in isolation to predict the behavior of modules’ composition. However, it has been shown that the behavior of a genetic module changes when other modules are in the cell due to competition for shared resources. In order to engineer multi-module circuits that behave as intended, it is thus necessary to predict changes in the behavior of a genetic module when other modules load cellular resources. Here, we introduce two characteristics of circuit modules: the demand for cellular resources and the sensitivity to resource loading. When both are known for every genetic module in a circuit library, they can be used to predict any module’s behavior upon addition of any other module to the cell. We develop an experimental approach to measure both characteristics for any circuit module using a resource sensor module. Using the measured resource demand and sensitivity for each module in a library, the outputs of the modules can be accurately predicted when they are inserted in the cell in arbitrary combinations. These resource competition characteristics may be used to inform the design of genetic circuits that perform as predicted despite resource competition.

## 1 Introduction

Genetic circuits are typically designed in a bottom-up fashion by combining previously characterized modules together to create new behaviors for particular applications (*1*). However, genetic modules often behave differently depending on the cellular context resulting in a lengthy design process and significant trial-and-error in designing genetic circuits (*2–4*). One major aspect of context dependence is resource competition between genetic circuit modules due to the fact that cellular resources such as RNAP and ribosomes are finite and shared among modules within a genetic circuit (*5–7*). It has previously been shown that resource competition between genes may result in more than a 60% decrease in expression levels (*6*), and that resource competition may cause significant changes in the qualitative behavior of genetic circuits (*7*). Resource competition has also been observed in cell-free systems (*8*), mammalian cells (*9*), and in computational models (*10–12*). Much work has gone into developing strategies to improve genetic circuits’ robustness to changes in cellular resources (*13–19*). In these strategies, resource competition is assumed to be an unknown disturbance and a controller (feedback or feedforward) is implemented to decrease the effects of resource competition on the system. However, characterization of genetic circuits with resource competition for prediction has remained largely unexplored.

Consider a situation where multiple genetic circuit modules are initially characterized in isolation. When the modules are combined to form a multi-module system, each module behaves differently since the modules must compete for resources with every other module, and, therefore, each module has access to a different amount of resources than when it was initially characterized. Therefore, to accurately design multi-module genetic circuits, the behavior of every module must be able to be predicted in the context of the final application despite resource competition effects. Additionally, it is highly desirable to characterize modules individually with a minimum number of experiments needed to characterize resource competition effects.

Here, we present an experimental method to estimate the effects of resource competition in genetic circuits, which can be used to predict circuit behavior in new contexts. We first present the method of estimating two key resource competition characteristics for a genetic circuit module of interest: resource demand and sensitivity. We define these quantities using a published mechanistic model of resource competition in genetic circuits (*6, 7*). These characteristics can be used to predict a module’s output with resource competition, which aids module design of genetic circuits. To experimentally measure the resource competition characteristics, we first characterize two resource sensors, which are then used to estimate the resource competition characteristics for any module of interest in a library. We then show that, using these characteristics, we can quantitatively predict the outputs of modules when they are inserted in the cell in any new combination. The predictions of module outputs are always more accurate than those obtained when neglecting resource competition.

## 2 Results

### 2.1 Using Resource Competition Properties to Predict Outputs

When genetic circuit modules are inserted into a cell, they use cellular resources for protein production and change the availability of cellular resources to other circuit modules in the cell. As shown in Figure 1a, the input-output response of a genetic circuit module of interest may be characterized in isolation. When the module of interest is in the same cell as other (perturbing) modules, the modules compete for cellular resources and, therefore, the output of the module of interest cannot be accurately predicted based only on its behavior in isolation (*6, 7*).

**Figure 1:**
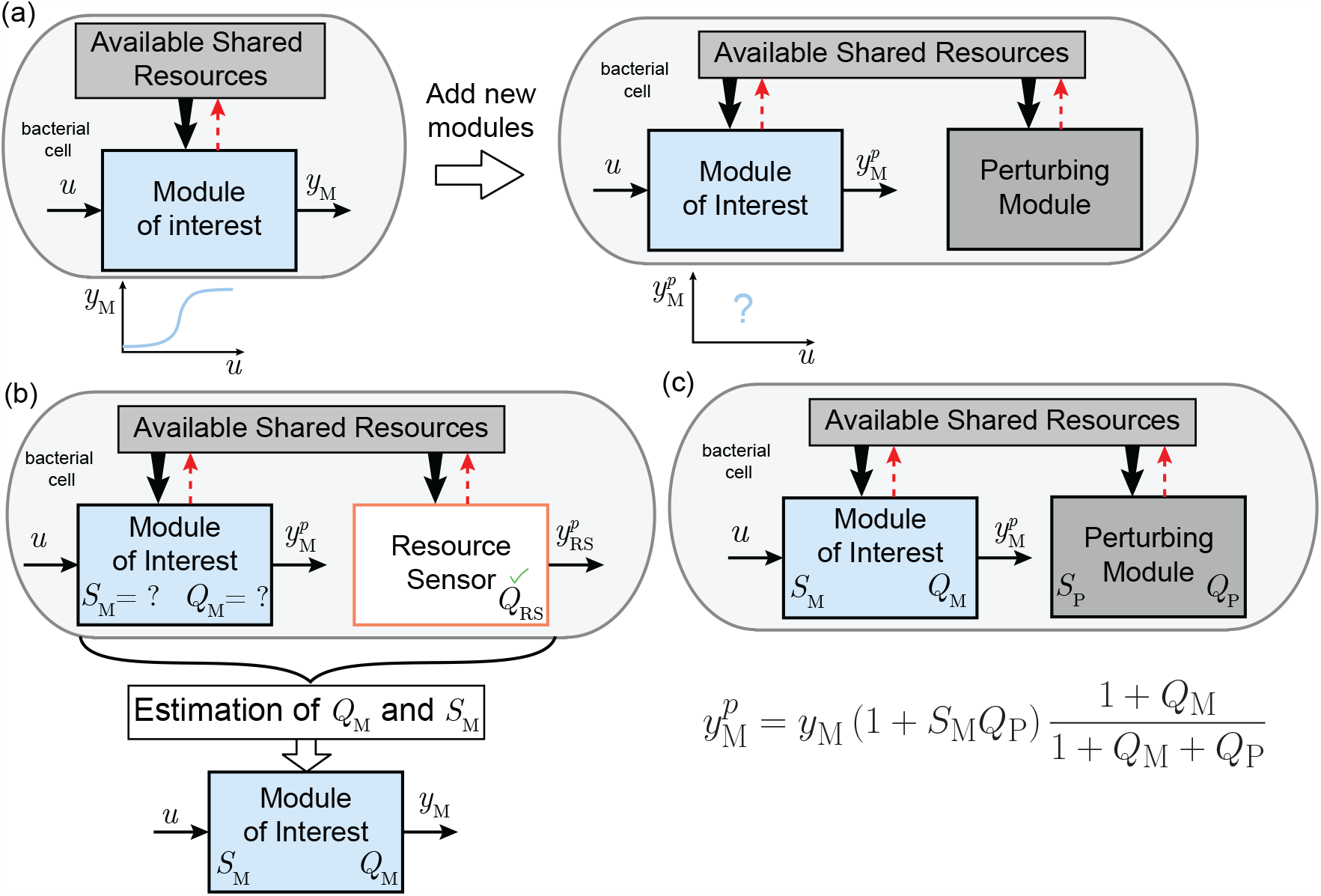
Resource competition characteristics allow prediction of modules’ output in new contexts. **(a)** The input-output response of a genetic circuit module of interest is characterized in isolation within the cell. When a new module is added to the cell, this module perturbs the pool of available cellular resources. This results in a change in the input-output response of the module of interest. **(b)** A resource sensor module enables the estimation of the resource competition characteristics *Q*_M_ (resource demand) and *S*_M_ (resource sensitivity) for a module of interest by measuring the input-output response of this module with the resource sensor in the cell. **(c)** Knowledge of the resource competition characteristics *Q*_M_ and *S*_M_ and of the *Q*_P_ for a perturbing module enables the prediction of the input-output response of the module of interest in new contexts such as with the perturbing module.

In order to predict the effect of resource competition on genetic circuit modules, we demonstrate a method to estimate two key *resource competition characteristics* of a genetic circuit module, which may be used to predict changes in the module’s input-output response due to resource competition in new situations.

For a module of interest M, the resource competition characteristics are the *resource demand Q*_M_ (“quantity” of resources used by the module) and *resource sensitivity S*_M_ (sensitivity of the module’s output to resource loads). As shown in Figure 1b, the output of the module of interest is predicted by a function of the module’s output when measured alone *y*_M_, the module of interest’s resource sensitivity *S*_M_, and the resource demands of both the module of interest *Q*_M_ and of the other, perturbing modules in the cell *Q*_P_. Thus, by estimating the resource demand and sensitivity for all modules, the changes in module outputs due to resource competition effects can be predicted as explained next.

The resource competition characteristics *Q*_M_ and *S*_M_ are defined based on a previously proposed model describing resource competition in bacterial cells, which we recall here (*6, 7*). We consider the production rate of a protein (which is related to the concentration of the protein at steady state; see the SI, Section Sl.2) as the output of a genetic circuit module of interest M, where the circuit module’s output is measured in the presence of another circuit module P. Based on this model, the production rate *y* of a module’s output protein is given as

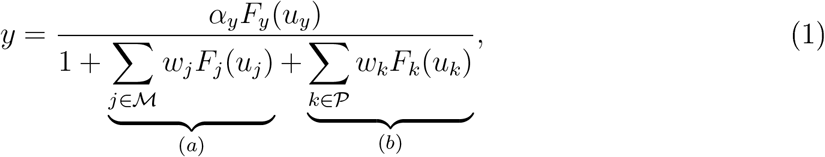

where *α*_*y*_ represents the maximum production rate of the output and is proportional to the total resources for transcription and translation, the functions *F*_*i*_(*u*_*i*_) represent the regulation function of the gene *i* by its input(s) *u*_*i*_ (which may be a Hill function (*20*)) and, for a single regulator is of the form 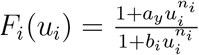 for some constants *a*_*i*_, *b*_*i*_, and *n*_*i*_. This model can be derived by applying the conservation law of gene expression resources during protein production and is derived in the SI, Section Sl.l as well as in (*7*). The quantity *w*_*j*_*F*_*j*_(*u*_*j*_) represents the resource demand of the individual gene *j* and is nondimensional (*7*) with *w*_*j*_ representing the resource demand coefficient of gene *j* which is a lumped parameter of biochemical binding constants that depends on the promoter strength, ribosome binding site (RBS) strength, and plasmid copy number. Furthermore, *γ* represents the protein degradation and dilution rate, and ℳ represents the set of genes in module M where the output *y* is in ℳ, while 𝒫 represents the set of genes in module P. Term (*a*) represents the resource demand of all the genes in module M and term (*b*) represents the resource demand of all genes in module P. This model assumes that total cellular resources are finite and constant.

Based on Eq. (1), the *resource demand* of the module of interest M, is defined as

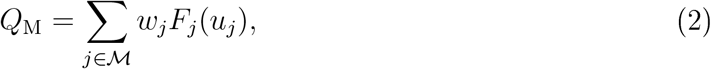

where ℳ is the set of all genes contained in the module of interest M. Note that the resource demand *Q*_M_ is the sum of the resource demands of all the constituent genes within the module.

The second resource competition characteristic is the sensitivity of the regulation function of the module’s output *F*_*y*_(*u*_*y*_) to changes in cellular resources, which we call the *resource sensitivity*, and is defined as

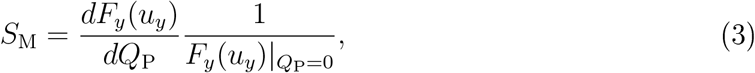

where *Q*_P_ is the resource demand of a module that is perturbing the resource pool, in other words, term (*b*) in Eq. (1). The resource sensitivity *S*_M_ determines how disturbances in the pool of cellular resources imparted by a perturbing module affect the regulation function of the module’s output. This change of the regulation function arises because, upon a resource perturbation, the expression of genes within the module of interest that regulate the modules’ output also change. Note that, for constitutive modules, *S*_M_ = 0 by definition since *F*_*y*_(*u*_*y*_) is constant in this case.

As shown in Figure 1b, in order to estimate the resource demand *Q*_M_ and resource sensitivity *S*_M_ of a general module, a special module called a *resource sensor* is used, where the resource demand *Q*_RS_ of the resource sensor was previously characterized. The output of the module of interest in new situations when the resource pool is perturbed by other modules can be predicted using the estimated 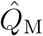 and *Ŝ*_M_ for all modules in the circuit, as shown in Figure 1c. Specifically, by combining the two resource competition characteristics *Q*_M_ and *S*_M_ based on their definitions, the output of a genetic circuit module can be predicted as other modules compete for resources through their own resource demand *Q*_P_ according to the equation

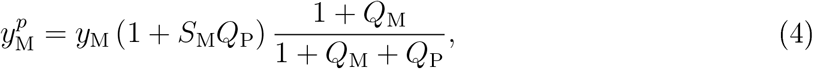

where *y*_M_ is the output of the module of interest in isolation and 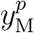 is the predicted output of the module when perturbed due to the depletion of the resource pool by the presence of the module P. For the derivation of Eq. (4), see the SI, Section Sl.

### 2.2 Characterization of a Resource Sensor

To estimate the resource competition characteristics *Q*_M_ and *S*_M_ for a generic module M, a special module called a *resource sensor* must be first characterized, which will be used to probe modules of interest. The resource sensor module consists of a single constitutively expressed fluorescent protein. Since the resource sensor is a single constitutive gene, *Q*_RS_ fully characterizes the resource competition characteristics of the resource sensor given that *F*_*y*_(*u*_*y*_) is a constant.

The resource demand of the resource sensor can be estimated from a procedure of three experiments as shown in Figure 2a. First, the outputs of two resource sensor modules are each measured in isolation. Then, both resource sensor modules are placed in the same cell, where they compete for resources, and their outputs are measured together. By comparing the change in output of each sensor under the resource perturbation by the other sensor, the resource demand *Q*_RS_ for both resource sensors l and 2 (RS1 and RS2) can be found according to the Resource Sensor Demand Estimation equations in Figure 2a.

**Figure 2:**
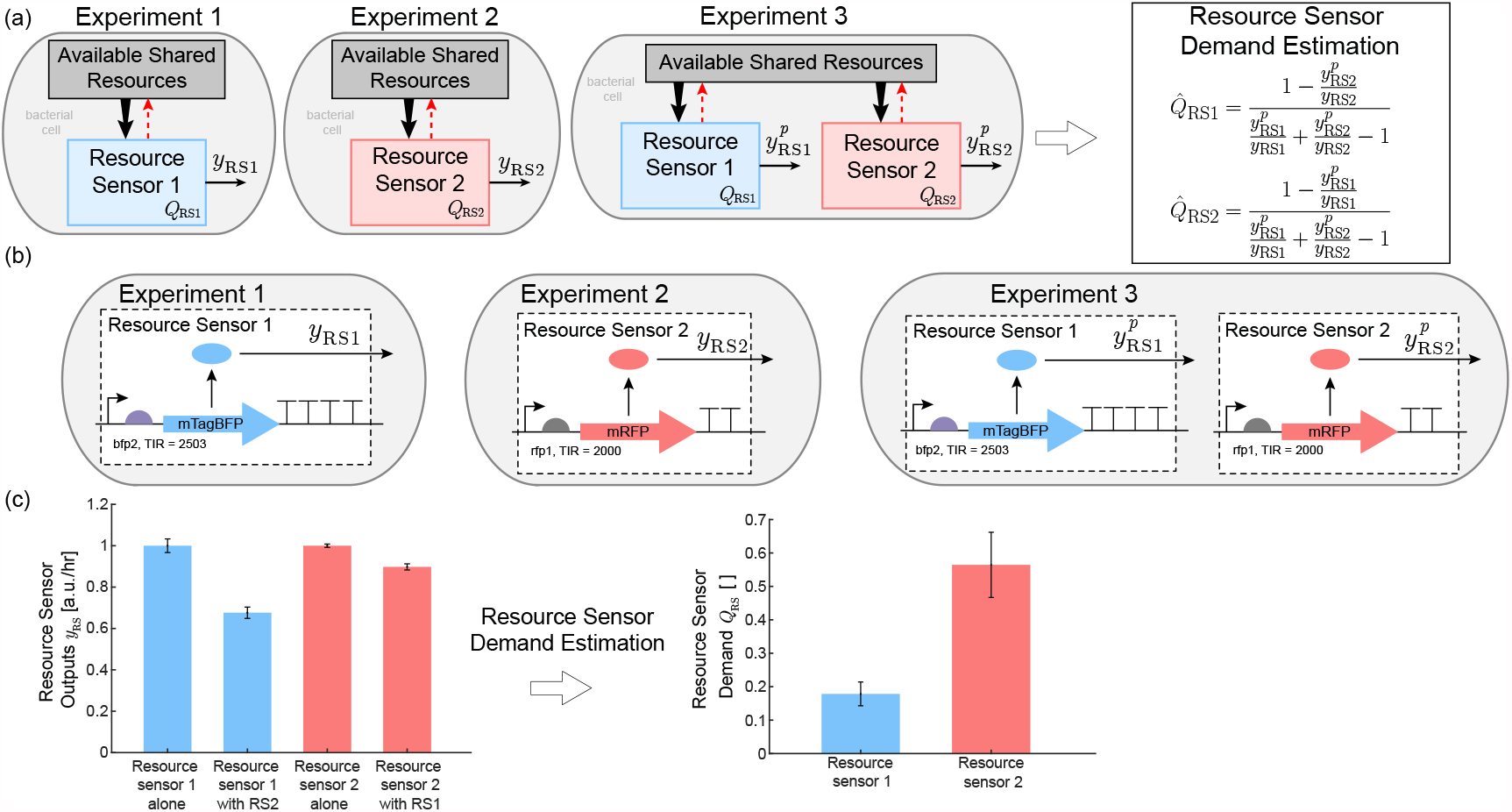
Experimental procedure to measure resource demand for a pair of resource sensors. **(a)** The outputs of two resource sensors, *y*_RS1_ and *y*_RS2_ when each sensor’s output is measured in isolation (Experiments 1 and 2) are compared to their respective outputs 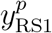 and 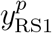 when the sensors are in the same cell together (Experiment 3). The superscript *y*^*p*^ represents the output of a sensor under a resource perturbation. **(b)** Genetic constructs used for implementing the resource sensors. See the SI, Section S4 for sequence details and Figure S14 for the relevant plasmid maps. The constitutive mTagBFP (BFP) gene with a weak RBS, bfp2, was chosen as resource sensor 1 (RS1), and a constitutive mRFP (RFP) gene with a strong RBS, rfp1, was chosen as resource sensor 2 (RS2). **(c)** From the measurements of the resource sensors outputs in isolation and together, the resource demands of both resource sensors can be estimated according to the Resource Sensor Demand Estimation Equations (Panel (a) or Eq. (5)). The fluorescent protein production rate per cell at steady state was used as the output of each resource sensor module. See Methods Section 4.4 for details on how production rate per cell was calculated. All measurements are the average of three technical replicates. Error bars represent one standard deviation from the mean and were propagated through the relevant equations using the standard error. Measurements were made using a microplate reader, see Methods Section 4.1 and Section 4.3 for details on cellular growth conditions and measurement conditions.

These equations were derived by expressing the output of each sensor using the model of Eq. (l) and using the fact that they are constitutive genes. See the SI, Section Sl.4 for the derivation of these equations. The system was set to its steady state, and solved for the resource demand characteristics *Q*_RS1_ and *Q*_RS2_ from the corresponding equations. Then, the estimators for the resource demand of the resource sensors are given by

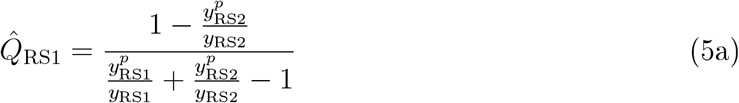

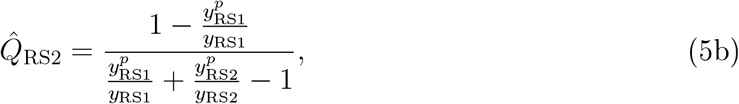

where *y*_RS1_ and *y*_RS2_ are the outputs of resource sensors 1 and 2 measured in isolation and 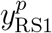 and 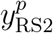 are the outputs of resource sensors 1 and 2 when they are measured together. In the experiments, the output fluorescent protein production rate per cell is used as the output of each module or resource sensor. The protein production rate is proportional to the steady state of the protein concentration per cell and independent of growth rate (see the SI, Section S1.2). For figures showing module and resource sensor outputs against time, see the SI, Section S5.3.

As in Figure 2b, the constitutive mTagBFP (BFP) gene with a weak ribosome binding site (RBS) was chosen as resource sensor 1, and a constitutive mRFP (RFP) gene with a strong RBS was chosen as resource sensor 2. The normalied output of resource sensors 1 and 2 are given in Figure 2c. These measurements are then used to calculate the resource demands *Q*_RS1_ and *Q*_RS2_ according to Eq. (5), with the resulting resource demands *Q*_RS1_ and *Q*_RS2_ for each resource sensor shown in Figure 2c. The resource demand of resource sensor 2 is significantly larger than the resource demand of resource sensor 1 since resource sensor 2 causes a larger decrease in the output of resource sensor 1 than resource sensor 1 causes in resource sensor 2. Detailed plasmid maps of all resource sensor constructs are shown in the SI, Figure S14. For additional experimental details on the design of the resource sensor modules and specific considerations for controlling for genetic context effects (*21*), see Methods Section 4.2.

### 2.3 Characterization of a Module

Using the estimated resource demand of the resource sensors 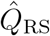, the resource competition characteristics *Q*_M_ and *S*_M_ of an arbitrary genetic module M may be estimated by following a similar set of experiments used to characterize the resource sensor. As shown in Figure 3a, the output of the module of interest is measured both with and without the presence of the resource sensor, and, similarly, the output of the resource sensor is measured both with and without the module of interest. These four measurements along with the measured resource demand 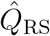 of the resource sensor are used to estimate the module’s resource competition characteristics 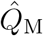 and *Ŝ*_M_.

**Figure 3:**
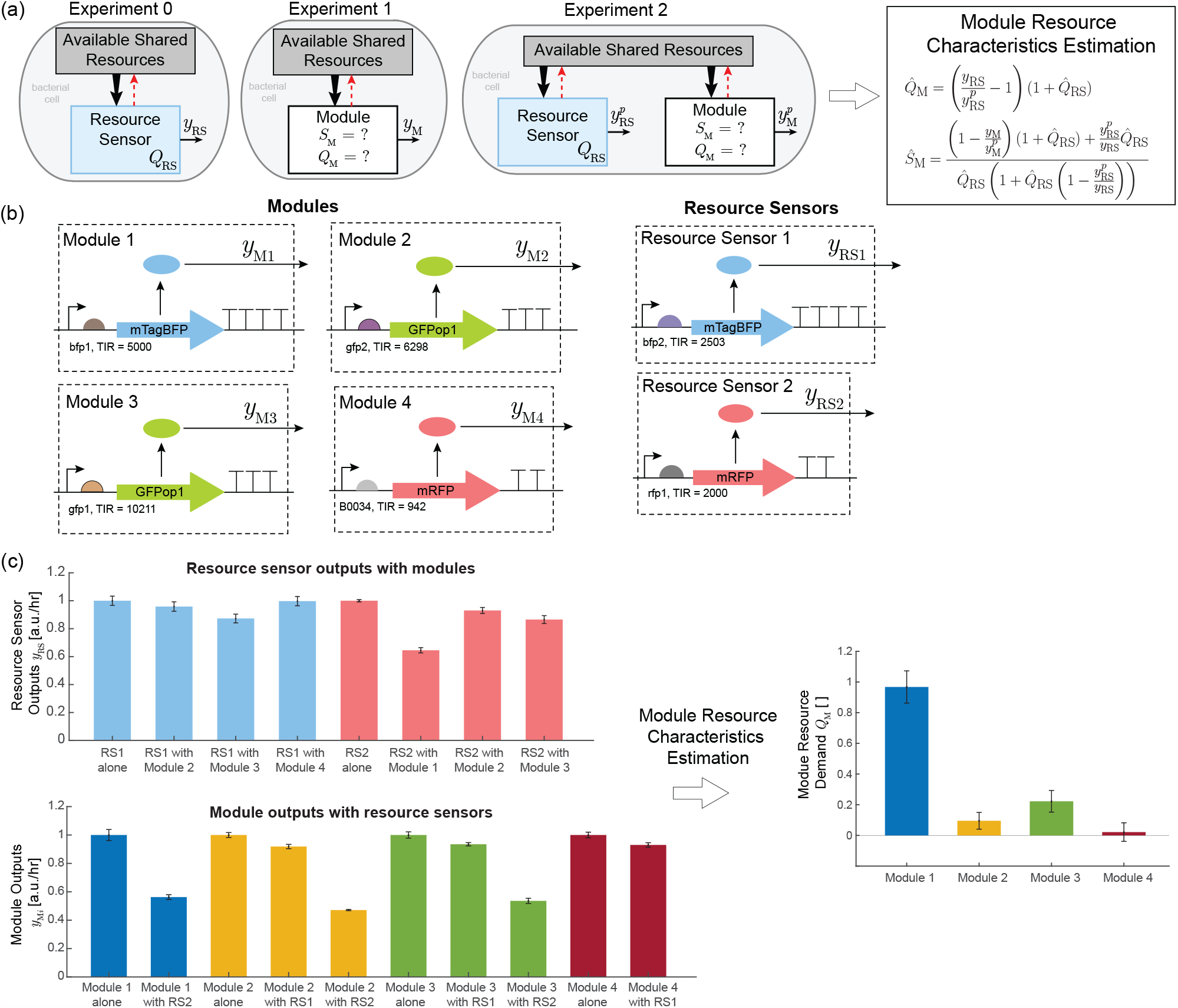
Estimation of the resource characteristics 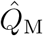 and *Ŝ*_M_ of a genetic circuit module. **(a)** Experiments to characterize the resource competition characteristics of the module of interest. The input-output responses of the resource sensor and of the module of interest are measured in isolation (Experiment 0 and 1). Then, the module is measured with the resource sensor (Experiment 2). **(b)** Genetic constructs of the circuit modules. See SI, Section S4 for sequence details and Figures S15 and S16 for plasmid maps. Module 1 consists of a constitutive mTagBFP (BFP) gene with a strong RBS, modules 2 and 3 consist of a constitutive GFPop1 (GFP) gene with a weak and strong RBS, respectively, and module 4 consists of a constitutive mRFP gene (RFP) with a weak RBS. The outputs of these modules were measured in isolation according to Experiment 1 and with resource sensors 1 and 2 according to Experiment 2, where the resource sensor demands 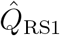 and 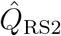 were previously estimated (Figure 2). **(c)** Measurements of the outputs of the resource sensors and modules according to Experiments 0, 1, and 2. Measurements are normalized such that the output of the module in isolation is set to 1. Superscripts ^*pi*^ indicate that the measurement is the perturbed output by module or resource sensor *i* for resource sensors *i* ∈ {1, 2} or modules *i* ∈ {1, 2, 3, 4}. The resource competition characteristics 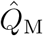 for each module are estimated according to the Module Resource Properties Estimation equations (Eq. (6)). Since the modules are constitutive the estimate for the sensitivity *Ŝ*_M_ was set to 0 for modules 1-4. The resource competition characteristics for module 1 was calculated using resource sensor 2 and the resource competition characteristics for module 4 were calculated using resource sensor 1. The resource demand and resource sensitivity for modules 2 and 3 are taken as the average of the respective resource demands and resource sensitivities found using both resource sensors 1 and 2. Non-averaged resource demands and sensitivities are shown in Figure S3. All measurements represent the average of three technical replicates. Error bars represent one standard deviation from the mean. Error estimates of 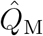 were obtained by propagating estimates for the error through Eq. (6) using the standard error. Measurements were made using a microplate reader, see Methods Section 4.1 and Section 4.3 for details on cellular growth conditions and measurement conditions.

As with the resource sensors, the resource demand characteristic *Q*_M_ of a module represents the resource competition term in Eq. (1) for all genes associated with the particular module. The resource competition characteristics are estimated by measuring the module output in isolation (Experiment 1) and the output perturbed by the resource sensor (Experiment 2) and using the resource competition model, Eq. (1), to solve for the desired quantities. The resource sensitivity *Ŝ*_M_ is solved for using an approximation for the derivative in the definition of *S*_M_, Eq. (3). The formulas of the estimates for the resource demand resource 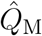 and resource sensitivity *Ŝ*_M_ of a module of interest M are given as

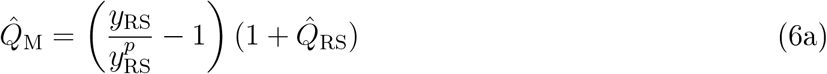

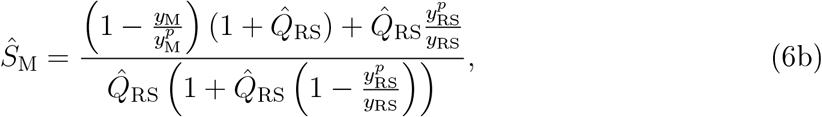

where *y*_RS_ is the output of the resource sensor measured in isolation, 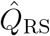 is the estimate for the resource demand of the resource sensor, 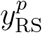 is the output of the resource sensor with the module of interest, *y*_M_ is the output of the module in isolation, and 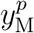 is the output of the module when measured with the resource sensor. For the derivation of these formulas, see SI, Section S1.5.

As shown in Figure 3b, four genetic circuit modules were characterized using both resource sensors. Each module consists of a constitutive gene with a fluorescent protein output, but could be a more complex module in practice (see Section 2.5). The output of each module was measured both in isolation and with the resource sensors. Resource sensors 1 and 2 were used depending on whether the resource sensor’s output is a different color from the module’s output. The outputs for each resource sensor alone and in the presence of the modules along with the outputs of the module alone and with the resource sensors are shown in Figure 3c. These data are then used to estimate the corresponding resource demand 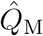 of the module of interest with the resource demand 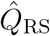 for the corresponding resource sensor following the Module Resource Estimation equations. Since modules 1-4 are constitutive, the sensitivity *Ŝ*_M_ was set to 0 by the definition of *S*_M_ (Eq. (3)). The estimated value of 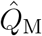 for each module is shown in Figure 3c. The sensitivity for an inducible module will be estimated from data in Section 2.5.

### 2.4 Prediction and Validation

Having estimated 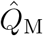 and *Ŝ*_M_ for each module, the module’s input-output response in new contexts accounting for resource competition can now be predicted. To demonstrate this, predictions of module outputs obtained from Eq. (4) are compared to measured outputs when pairs of modules are measured together in the cell. The four previously characterized modules were used, as shown in Figure 4a.

**Figure 4:**
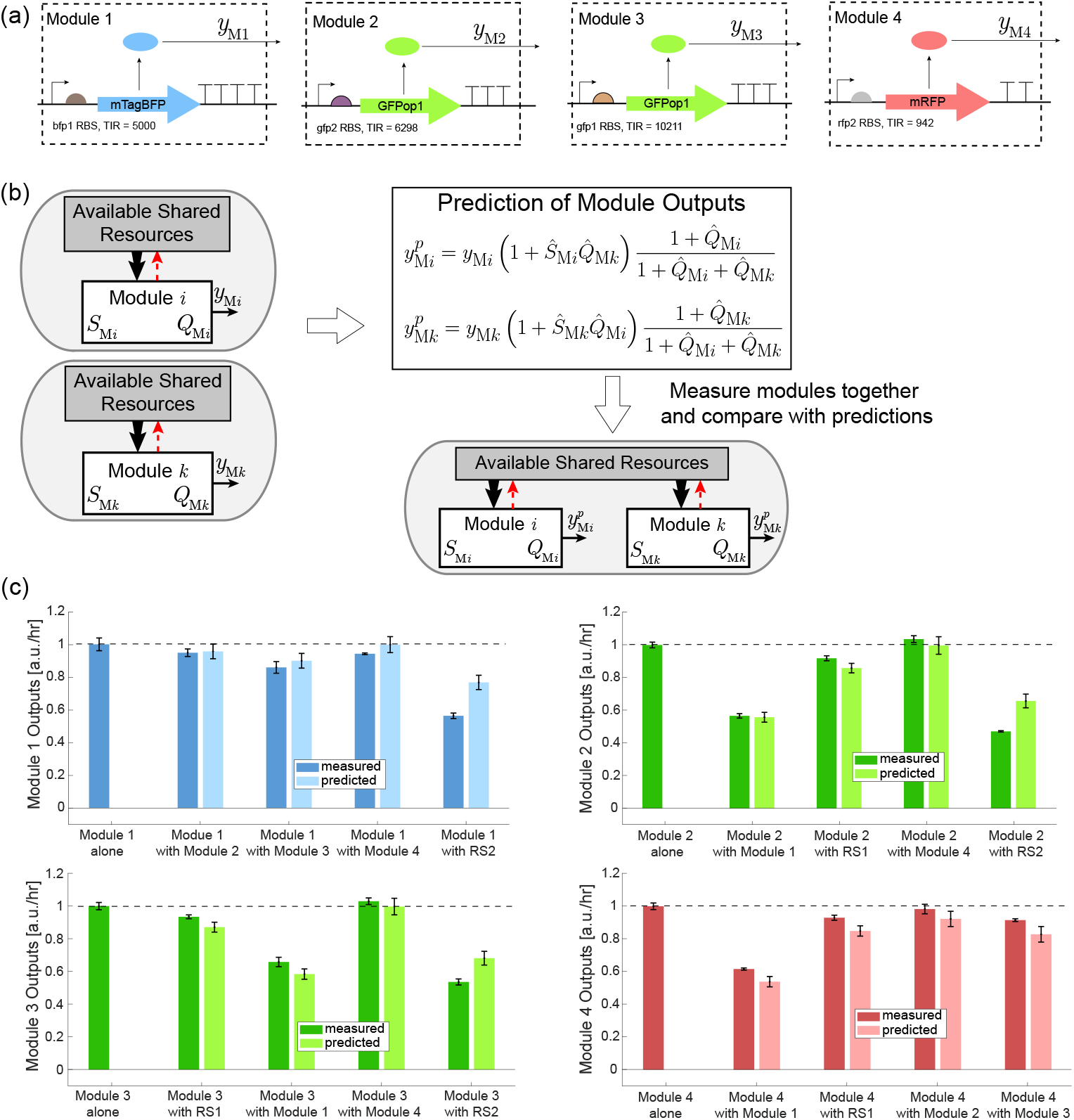
Validation of the method by prediction of module outputs in new contexts. **(a)** The four constitutive modules previously characterized in Figure 3 with known 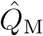 and *Ŝ*_M_ were used to validate the method. The sensitivity for all constitutive modules was set to 0. **(b)** Based on each module’s output in isolation and the estimated resource competition characteristics 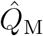 and *Ŝ*_M_, the outputs of the modules when measured together are predicted using the Prediction of Module Outputs equations (Eq. (7)). The superscript ^*p*^ indicates that the output is perturbed and the ^indicates that it is an estimated (or predicted) quantity. Additionally, the outputs of the modules are measured when they are together in pairs to compare with the predictions. See the SI, Section S4 for sequence details and Figure S17 for plasmid maps. **(c)** Comparison of the measured outputs of modules 1, 2, 3, and 4 with the predicted outputs based on 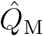 and *Ŝ*_M_ for each module when perturbed by another module or resource sensor. Measurement results are shown in darker colored bars and predictions shown with lighter colored bars. All measurements are the average of three technical replicates. Error bars represent one standard deviation from the mean. Error estimates of the predicted outputs were obtained by propagating the error estimates through Eq. (7) using the standard error. Measurements were made using a microplate reader, see Methods Sections 4.1 and 4.3 for details on cellular growth conditions and measurement conditions.

Each module was measured with every other module and resource sensor, resulting in a new cellular context due to the resource competition between the modules, as shown in Figure 4b. The output of each module when perturbed by another module was predicted using the estimated 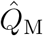 and *Ŝ*_M_ for each module along with its output in isolation *y*_M_. The predicted output of a module under resource competition from another, perturbing module P is given as

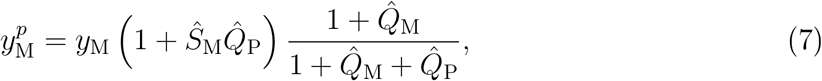

which is found by comparing the resource competition model, Eq. (1) with and without the perturbing module resource demand *Q*_P_ and combining with the definition of *S*_M_ (Eq. (3)). Further details on the derivation can be found in the SI, Section S1.3.

The measured outputs of the modules when measured with other modules were compared to the predictions using the resource competition characteristics of each module 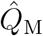 and *Ŝ*_M_, shown in Figure 4c. Predictions for the outputs of module 1 and module 2 are within one standard deviation from the measured values except when perturbed by resource sensor 2 and when module 2 was perturbed by resource sensor 1 the prediction was within two standard deviations. Predictions for modules 3 and 4 are within two standard deviations from the measured values except when module 3 is perturbed by resource sensor 2. The main reason for this discrepancy is that the growth rate changes significantly under the perturbation from resource sensor 2 (SI, Figures S6 and S7), resulting in effects that are unaccounted for in the model such as a change in total cellular resource concentration. More discussion on these discrepancies are presented in the Discussion, Section 3. Thus, the resource competition characteristics can be used to accurately predict the outputs of the constitutive modules in new contexts arising from resource competition between modules. The predictions using the resource competition characteristics are always a significant improvement compared to ignoring resource competition effects (modules alone). When the estimated sensitivities *Ŝ*_M_ of modules 1-4 are allowed to be non-,ero and estimated according to Eq. (6b), these predictions moderately improve in accuracy, see SI, Figure S2. This corresponds to allowing the gene regulation function *F*_*y*_(*u*_*y*_) in the resource competition model Eq. (1) to encompass regulatory functions of the whole cell, such as changes in gene expression due to a cellular stress response or changes in the total resource concentration in the cell.

### 2.5 Application to Inducible Genetic Modules

To verify that the new characterization method that we have proposed gives accurate predictions for more complex modules, we next considered inducible modules. Characterization of an inducible module is the same as that presented for the constitutive modules except that the resource competition characteristics *Q*_M_ and *S*_M_ must be estimated for each value of the input inducer concentration, *u*. We created module 5, consisting of an EYFP (YFP) gene and PhlF repressor, as shown in Figure 5a.

**Figure 5:**
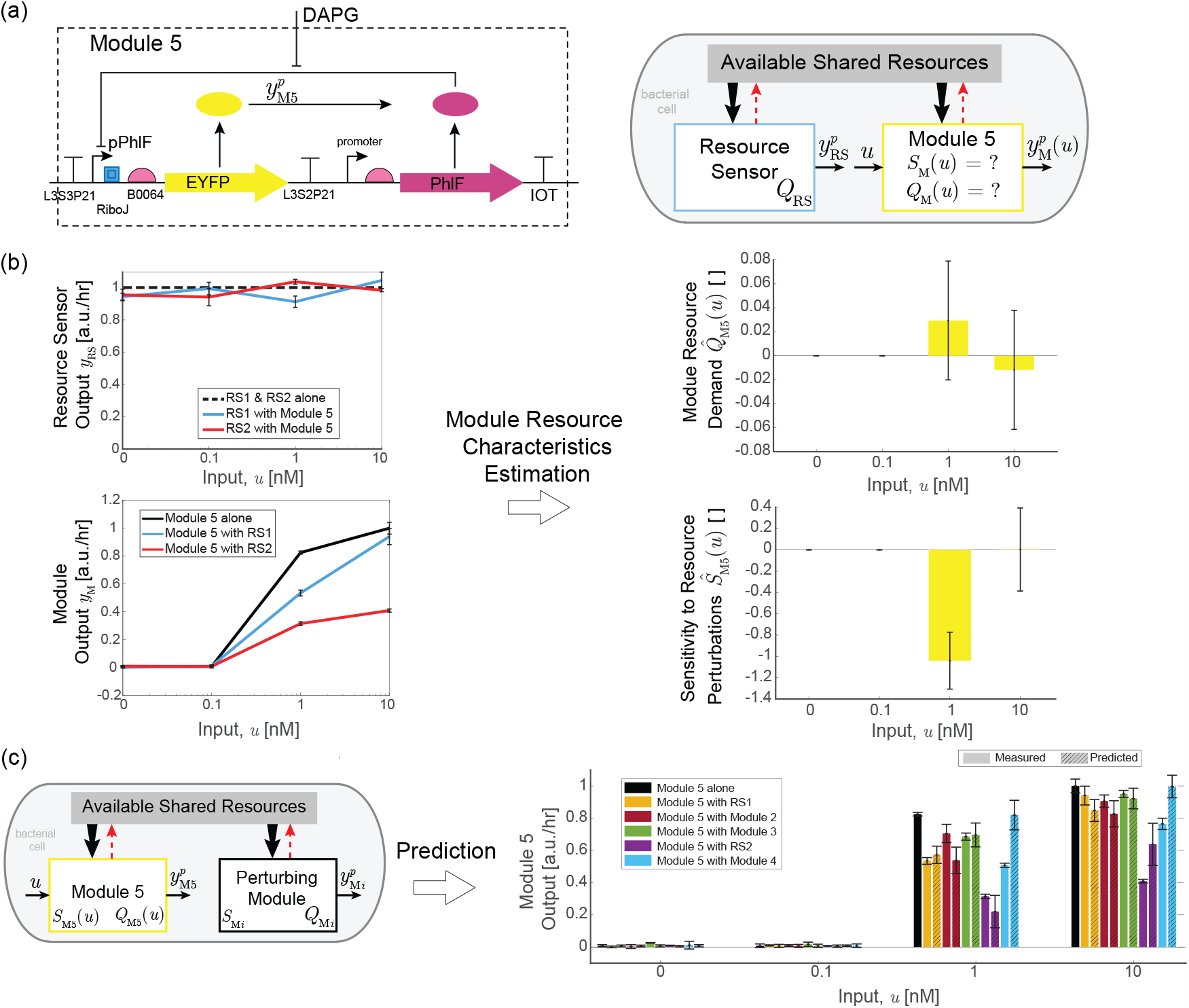
Application to predict an inducible module’s input-output response in new contexts. **(a)** Module 5 consists of a PhlF repressor and EYFP fluorescent protein output. The inducer 2,4-Diacetylphloroglucinol (DAPG) is used to vary the level of repression of the pPhlF promoter regulating EYFP by PhlF. See the SI, Section S4 for sequence details and Figure S18 for plasmid maps. **(b)** The outputs of module 5 and resource sensors 1 and 2 were measured for each module in isolation and when module 5 was measured with resource sensors 1 and 2 for each inducer concentration 0 nM, 0.1 nM, 1 nM, and 10 nM. The resource competition characteristics of the module were estimated for each value of the input using the Module Resource Properties Estimation equations (Eq. (5)). The resource demand and sensitivity were calculated using both resource sensor 1 and 2 and the average of each was taken for each input concentration. Non-averaged resource demand and sensitivity are shown in the SI, Figure S3. **(c)** Prediction of the behavior of module 5 when operating in new contexts with other modules based on its output in isolation, 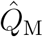, and *Ŝ*_M_. Solid bars represent measured data and dashed bars represent predictions made using Eq. (7). All measurements are the average of three technical replicates. Error bars represent one standard deviation from the mean and were propagated through the relevant equations using the standard error. Measurements were made using a microplate reader, see Methods Sections 4.1 and 4.3 for details on cellular growth conditions and measurement conditions.

After measuring module 5 alone and with the resource sensors, the resource sensor outputs and module outputs for each inducer concentration, shown in Figure 5b, were used to estimate the module’s resource demand 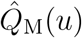 and resource sensitivity *Ŝ*_M_(*u*) for each inducer concentration. The input-output curves of the inducible module when paired in the cell with another characterized, perturbing module can be predicted for each input using these maps, as shown in Figure 5c. All predictions are within 1.5 standard deviations to the corresponding measurements except for the output of module 5 with module 4, where module 5 ’ s output is not predicted to significantly decrease under the perturbation, since 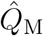 for module 4 is less than 0.1. This may be a result of genetic context effects, as explained further in the Discussion, Section 3.

## 3. Discussion

In this paper, the resource demand *Q*_M_ and sensitivity *S*_M_ were defined based on a mathematical model describing resource competition in genetic circuits. These quantities describe how modules behave in the presence of other modules that perturb the pool of cellular resources. Knowledge of *Q*_M_ and *S*_M_ for every module in a collection enables the quantitative prediction of changes in module outputs in new cellular context including different module combinations. Estimation of *Q*_M_ and *S*_M_ for a module of interest requires measuring the module of interest with a resource sensor module. This method was experimentally validated where predictions of module outputs using *Q*_M_ and *S*_M_ were compared with measurements of module outputs when pairs of modules were measured together. Predictions were within one standard deviation from the true measurements for the majority of the combinations tested, and were always strictly better than predictions ignoring resource competition (Figures 4 and 5).

Here, we discuss some considerations when implementing the method to estimate *Q*_M_ and *S*_M_ for a module and current limitations of the approach. The characteristics *Q*_M_ and *S*_M_ can predict large changes in module output due to resource competition effects, but care must be taken to detect or predict small changes in module behavior due to resource competition. The best resource sensors should have a large enough resource demand *Q*_RS_ to impose a measurable change in the output of the module of interest, but a small enough *Q*_RS_ to avoid creating a large stress response from the host cell. Based on our experimental results, it is recommended that the resource sensor demand *Q*_RS_ be between approximately 0.2 and 0.8. It should be noted that the resource competition characteristics are nondimensional quantities, so microplate readers do not need to use absolute units of fluorescence to obtain accurate estimates of the resource demand and resource sensitivity.

Differences in genetic context (*21*) were controlled for by including all coding sequences for the constitutive modules on the same plasmid and turning off module translation by using a very weak RBS (TIR approximately 1.0) for the corresponding coding sequence. It was verified that no significant fluorescence was observed from modules when they were turned off through the use of a very weak RBS, as shown in the SI, Section S3.1, Figures S8 S13. Resource competition has been shown to be mainly attributed to translational resource competition in exponentially growing bacteria (*6*), so transcriptional resource competition does not significantly effect our results. The resource competition characteristics due to transcriptional resources may also be estimated using a similar set of experiments, which have been shown to be important source of resource competition in mammalian cells (*17*).

One source of error in the validation of module 5 (Figure 5) is that the coding sequences for genes (mTagBFP, GFPop1, or mRFP) that were not relevant were not included on the plasmids with module 5, so genetic context was not held constant across all constructs in the same way as for the constitutive modules 1-4. Therefore, a possible reason for the difference between the predicted and measured outputs when modules 4 and 5 were measured together (Figure 5c) is that the spacer between the terminator for module 4 and the promoter of module 5 was only 19 bp, while for the other pairs of modules, the spacer was larger than 55 bp (see the SI Section S4 for details on the plasmid maps). This may have resulted in a change in the activity of the EYFP promoter due to transcriptional context differences such as supercoiling (*21*).

Theoretically, the estimated values of *Q*_M_ should be independent of the resource sensor used to perform the estimation however, when the resource sensor has a large demand 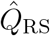, the estimated 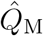 of the module of interest was found to be larger than when measured using a resource sensor with smaller 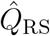, see the SI, Figure S5. This is likely due to the regulation of the cellular resources or stress response by the host cell (*12, 22*). In fact, the model on which the method is based assumes that total cellular resources are constant. However, if the cell undergoes a significant change in growth rate due to the presence of a genetic circuit module, the measurements of resource demand and sensitivity may not be accurate since total cellular resources is related to growth rate (*23, 24*). This is one possible source of error in the presented method and is especially likely to cause inaccuracies in the estimates for modules where the resource sensor demand *Q*_RS_ is large (greater than approximately 0.8).

Some approximations were necessary for estimating 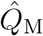 and *Ŝ* _M_ when the resource demand *Q*_M_ of a module of interest depends on unmeasured protein concentrations, such as in a more complex, inducible module (SI, Section S1.5). This explains why the predictions for the constitutive modules 1-4 (Figure 4) were closer to the measurements than for the inducible module 5 (Figure 5).

Growth rate is subject to change due to burden of the synthetic circuit on the host cell, which may cause additional unexpected changes in circuit behavior due to changes in dilution rate (*24* -*28*). In the SI, Section S2, we demonstrate that the cellular growth rate and the resource demand *Q*_M_ are closely related and show that estimation of the resource demand may be used to predict changes in growth rate due to metabolic burden on the host cell. This shows that the resource demand *Q*_M_ can be used to predict total cellular output and may be useful in metabolic engineering applications (*29* -*31*).

The resource competition characteristics may also be used to indicate whether resource competition effects can safely be ignored in genetic circuit design or need to be included for accurate circuit design. When the resource demands for all modules in a genetic circuit are small, e.g. 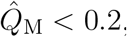, resource competition effects can safely be ignored as long as the number of modules is sufficiently small. When some module resource demands are moderate, e.g.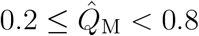, resource competition should be accounted for to make accurate predictions of module outputs and overall system behavior. When any module resource demand is large, e.g. 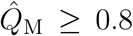, significant resource competition effects can be expected and the cellular growth rate will likely decrease due to the burden on the host cell. For an analysis of the experimental dependence of growth rate on the resource demand, see the SI, Section S2. Note that the resource demand estimation only depends on the resource sensor measurement (Eq. (6a)), so it can be estimated even for modules whose output is not measured.

When designing genetic circuits, it should be noted that, in general, the output of a module with a large resource demand *Q*_M_ will change less in response to the same resource perturbation than a similar module with a smaller resource demand, as can be seen from Eq. (7). However, modules with large resource demand will cause other modules to experience more significant changes in their behavior due to resource competition. Additionally, unexpected changes in behavior may occur due to resource competition, such as a change in the number of equilibria of a system (*32, 33*). The method assumes that disturbances due to resource competition are small enough such that these qualitative changes do not occur.

In order to ensure more accurate estimates of the sensitivity *S*_M_ of a module, the resource demand *Q*_RS_ of the resource sensor should not be too small as it should impart a clear change on the module’s output. As a consequence, chromosomally integrated resource sensors may not be well suited for this task (*25*). However, a chromosomally integrated resource sensor as in (*25*) would be very useful in measuring the resource demand *Q*_M_ in cases when the sensitivity *S*_M_ is not significant (e.g. for a constitutive module). This set up would minimize the number of experiments necessary by requiring only one experiment per module to find the module’s resource demand *Q*_M_. Since the resource sensor is chromosomally integrated, it will impart a much smaller resource demand compared to genes on the plasmid, giving *Q*_RS_ ≈ 0. Then, the resource demand for a module of interest, *Q*_M_, can be estimated by measuring the resource sensor output with and without the presence of the module of interest and using Eq. (6a) with *Q*_RS_ = 0.

Other recent work has attempted to better characterize genetic circuits for improved predictions. In (*34* -*36*), the authors used RNA sequencing and ribosome profiling to inform a mathematical model of a multi-module genetic circuit. However, the mathematical model does not include resource competition among the constituting genetic modules. Therefore, the model and its predictive power do not generalize to when the constituent modules are used in different combinations and contexts when the availability of cellular resources changes.

Future work may address how changes in the media or temperature change the resource demand *Q*_M_ and sensitivity *S*_M_. The resource competition characteristics *Q*_M_ and *S*_M_ may be applied to circuits with feedback or feedforward controllers implemented such as in (*14, 15*), so both improved robustness from the controller and improved prediction of module behavior can be obtained. Software that predicts the values of *Q*_M_ and *S*_M_ based on DNA sequences or the strength of promoters and ribosome binding sites may be designed as well. The values of *Q*_M_ and *S*_M_ may also be used to give conditions when it is advantageous to distribute genetic circuits across multiple cells within a consortium to reduce an excessive burden on the host cell such as in (*37*). Additionally, an extension of *Q*_M_ and *S*_M_ to situations where there are multiple pools of resources such as transcriptional resources in mammalian cells (*9*) or synthetic orthogonal ribosomes as in (*13*) would be valuable. Finally, *Q*_M_ and *S*_M_ may be used to investigate more complex models for resource competition such as when the pool of resources is regulated by the host cell.

## 4 Methods

### 4.1 Strain and Growth Medium

The Top10 strain of bacterium *E. coli* was used to perform all experiments. M9610 supplemented media and was used to test the constitutive plasmids was made by modifying M9 supplemented media (*14*). The recipe for 5x M9610 salt stock is 33.36 mM Na_2_HPO_4_ 7 H_2_O, 220.40 mM of KH_2_PO_4_, 8.56 mM of NaCl, and 18.69 mM of NH_4_Cl. The recipe for 50 mL of M9610 supplemented medium is 10 mL of 5x M9610 salt stock, 1 mL of 10% (w/v) casamino acid, 1 mL of 20% (w/v) glucose, 50 uL of 0.1 M CaCl_2_, 100 uL of 1M MgSO_4_, 333 uL of 50 g/L thiamine, 50 uL of 100x ampicillin, and the remainder of the 50 mL is sterile water.

### 4.2 Genetic Circuit Design and Construction

PCR, Gibson assembly and Golden Gate assembly were used to construct the plasmids. DNA fragments were amplified with PCR. Gibson assembly was performed using NEBuilder HiFi DNA Assembly Master Mix (NEB #E2621S) or Golden Gate master mix (NEB #E1601). Plasmids were incubated on ice for 30 min with competent cells which were prepared using CCMB80 buffer (TekNova, C3132)and were transformed by heat shocking competent cells at 42°C for 1 min. Plasmid DNA was prepared using the miniprep kit (Zymo Research, D4015), and sequencing was performed using the Quintarabio basic DNA sequencing service.

To control for transcriptional effects such as super-coiling (*21, 38*), transcription read-through, or promoter blocking by nearby terminators, all constitutive plasmids contained the coding sequences for all of BFP, GFP, and RFP proteins. Gene translation was turned off for the unwanted coding sequences by using extremely weak (TIR < 1.0) ribosome binding site strength (RBS) for the constitutive module experiments. This was verified by fluorescence measurements for the relevant genes and were observed to have no significant fluorescence. Translation initiation rates (TIR) values for each module were calculated using the RBS calculator 2.0 (*39*). Multiple strong terminators at the end of each coding sequence were used to minimize transcriptional read-through and spacers were introduced between terminators and promoters to minimize DNA-context-related effects between modules and resource sensors (see plasmid maps in the SI, Section S4, Figures S14-S18 and plasmid sequences in the SI, Table S10). Since ribosomes are the limiting resource in bacteria (*6, 10, 40*), and all promoters used for the constitutive modules were strong and of similar strength, varying the strength of the RBS gives a good estimation of the different resource competition effects between the modules. The strong constitutive promoters J23100, J23119, and J23104 were used for the constitutive modules and resource sensors to ensure resource competition effects would be significant enough to be easily measurable.

For the inducible EYFP module (module 5) experiments, only the coding sequence for the resource sensor/perturbing module of interest was included in the plasmids. It was found that there were only minor differences between the output of the module without a coding sequence of a fluorescent protein and one with the coding sequence of a fluorescent protein but a very weak RBS due to transcriptional context effects. The module 5 was taken from the pAJM.847 plasmid in (*41*) (Addgene #108524, additional details in the SI of (*41*), page 42). The molecule 2,4-diacetylphloroglucinol (DAPG) was used as the inducer for module 5.

Controlling for transcriptional context does not appear to be necessary in applications since module 5 predictions were still good even though the resource sensor or module on the plasmid with module 5 was different for different experiments. It is necessary to use multiple strong terminators to eliminate transcriptional read-through between the module and resource sensor and to use a long enough spacer (at least approximately 40 bp) between the resource sensor and the module of interest to ensure that terminators do not block nearby promoter activity.

### 4.3 Plate Reader

Cells with the relevant plasmids were streaked on LB agar plates with carbenicillin and grown in an incubator at 37°*C* for 20-24 hrs. Cells were then picked and inoculated into 800 μL of M9610 liquid media with ampicillin in a loose cap culture tube (VWR, 13 mL culture tube, cat #60818-667) and grown in an air shaker (New Brunswick Scientific Excella E24 M1352-0000) at 30°*C* with at 200 rpm for 7-9 hrs. Cells were then inoculated into a 96-well plate (Falcon #351172) and diluted to an initial optical density at 600 nm (OD _600_) reading of approximately 0.02 in 200 μL of M9610 liquid media. Cells were grown in a plate reader (Synergy MX, Biotek, Winooski, VT) at 30°*C* for 10 hrs with measurements every 5 mins. The plate was shaken for 3 sec before OD_600_ and fluorescence measurements. Excitation (EX) and emission (EM) wavelengths for the plate reader were chosen to maximi:ze difference between the different fluorescence channels while keeping a strong signal. The plate reader settings for each measurement are as follows. OD_600_ channel settings: absorbance at 600 nm; BFP channel settings: EX = 400 nm (bandwidth = 9 nm), EM = 460 nm (bandwidth = 9 nm), Sensitivity = 80; GFP channel settings: EX 485 nm (bandwidth = 9 nm), EM 513 nm (bandwidth = 9 nm), Sensitivity = 90; RFP channel settings: EX 584 nm (bandwidth = 9 nm), EM 619 nm (bandwidth = 13.5 nm), Sensitivity = 100; YFP channel settings: EX 520 nm (bandwidth = 9 nm), EM 545 nm (bandwidth = 13.5 nm), Sensitivity = 100. At least three wells filled with 200 uL M9610 liquid media were used to subtract background OD_600_ and fluorescence measurements. All experimental measurements consisted of three technical replicates. Measurements of the cells in the 96-well plate were spatially separated with at least one empty well between active wells to reduce cross-talk between neighboring wells (*42*).

### 4.4 Data Analysis

The protein production rate was used as the output of each circuit module, since it is independent of growth rate, see SI Section S1.2 for theoretical justification or (*25*). The protein production rate of a protein *X* per cell is given as

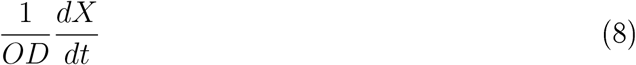

where *OD* is the optical density of the cells. Fluorescence measurements were taken every 5 minutes using the microplate reader. The numerical time derivative 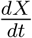 was found after sub-tracting background fluorescence and absorbance by using a 5-pt centered Savitzky-Golay filtered first derivative and then applying a (zero-phase) moving average filter with window size of 9 points. The steady state production rate was selected for each construct as a time window of at least 2 hours when the cells were in exponential phase where the protein production rate per cell was approximately constant and the fluorescent protein measurements were above the lower detection limit of the plate reader. This usually occurred between approximately 2 and 6 hours after the start of the experiment. Module output was calculated as the average production rate over the time window for each technical replicate and then averaged across all three replicates. Error bars on the module’s output represent one standard deviation in the difference of the means across the three replicates. Error estimates were propagated through all equations using the standard error estimates. Growth rate for each construct was determined using a robust nonlinear least squares logistic curve fit, utilizing the Levenberg-Marquardt algorithm. All computations were completed using custom scripts written in MATLAB R2020a, which are available in the Supporting Information. Corrections for fluorescence scattering due to cells and bleed-through across fluorescence channels were corrected for based off the raw fluorescence measurements. Graphs of OD _6_00 vs time are available in the SI, Figures S25-S27, and graphs of fluorescent protein production rate against time are available in the SI, Figures S28-S37.

### 4.5 Fluorescence Scattering Correction

As the optical density cells of the well change, the background fluorescence readings also change due to light scattering. To correct for this phenomena, correlations were measured for determining the relationship between background fluorescence measurements and OD _600_. Linear regression was applied between OD_600_ and the relevant raw fluorescence measurements for cells not expressing the relevant proteins. The correlation between BFP fluorescence and OD_600_ was *C*_*OD,B*_ = −2385*BFP/OD*(*a*.*u*.), and the correlation between GFP fluorescence and OD_600_ was *C*_*OD,G*_ = 1300*GFP/OD*(*a*.*u*.). These correlations are negligible for YFP and RFP measurement channels. Each was subtracted off of BFP and GFP fluorescence signals according to the equations *B*_*new*_ = *B*_*raw*_ − *C*_*OD,B*_*OD* and *G*_*new*_ = *G*_*raw*_ − *C*_*OD,G*_*OD*. The magnitude of this correction is less than 10% of the signal. Data used to estimate this effect and find correlations is shown in the SI, Figure S23.

Multiple scattering effects within the microplate reader become significant at larger optical densities (*43, 44*), so steady state data was selected for OD_600_ lower than this value before protein production rate was observed to drop, which normally occurred below OD _600_ *<* 0.05.

### 4.6 Bleed-Through Correction

Correction for fluorescence bleed between channels using linear unmixing (*45*) is applied to each fluorescence channel. The correlation between fluorescence intensities was found by applying linear regression between pairs of fluorescent measurement channels for a cell expressing only one protein to find the correction (correlation) matrix. Then, the measured fluorescent values may be expressed as a linear combinations of the true fluorescence values given by

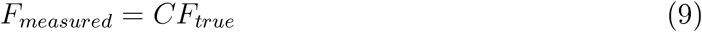

where *C* is the correlation matrix, which can be found by applying linear regression between measurement channels when only one type of fluorescent protein is expressed (I can’t find this treatment in the plate reader literature, only in the microscopy literature), and *F*_*measured*_ is the observed fluorescence intensity for each channel with cross-talk and *F*_*true*_ is the actual fluorescence intensity. For our data, the correlation matrix is

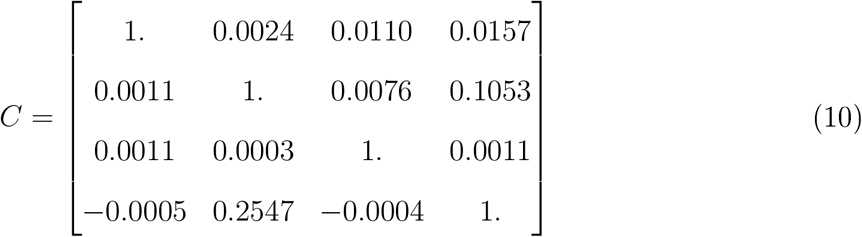

where first column encodes the correlation of the readings of the channels BFP, GFP, RFP, and YFP, respectively, when only mTagBFP2 is expressed and the correlation was found using linear regression. A similar procedure is used to populate the 2nd (GFP), 3rd (RFP), and 4th (YFP) columns with their respective proteins. Plots of pairs of fluorescence measurements against each other for purely expressed BFP, GFP, RFP, and YFP are given in the SI, Figures S19-S22. Entries in *C* less than 0.01 are set to 0. Note that the is no significant fluorescence bleed between channels except for between the YFP and GFP channels. The compensated fluorescence measurements are given as

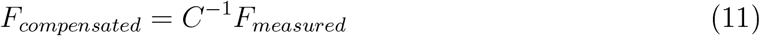

where *F*_*measured*_ is a column vector containing the fluorescent channels in the order BFP, GFP, RFP, and YFP. Eq. (11) is applied for each well at each time-step.

## Supporting information

Experimental data and Matlab code

## Acknowledgement

The authors thank Yili Qian for helpful discussions and Hsin-Ho Huang for help proofreading the manuscript. We also thank Massimo Bellato for his contributions to earlier versions of the constructs. This work was supported by NSF Expeditions in Computing Award #1521925.

## Author Contributions

All authors designed experiments and interpreted the data; CM preformed the research, cloned constructs, performed experiments, developed the mathematical models, wrote the paper; DDV designed the research and edited the paper.

## Supporting Information Available

The following files are freely available.

- Filename: Supporting Information
- Filename: MATLAB code and and experimental data zip folder

## TOC Graphic

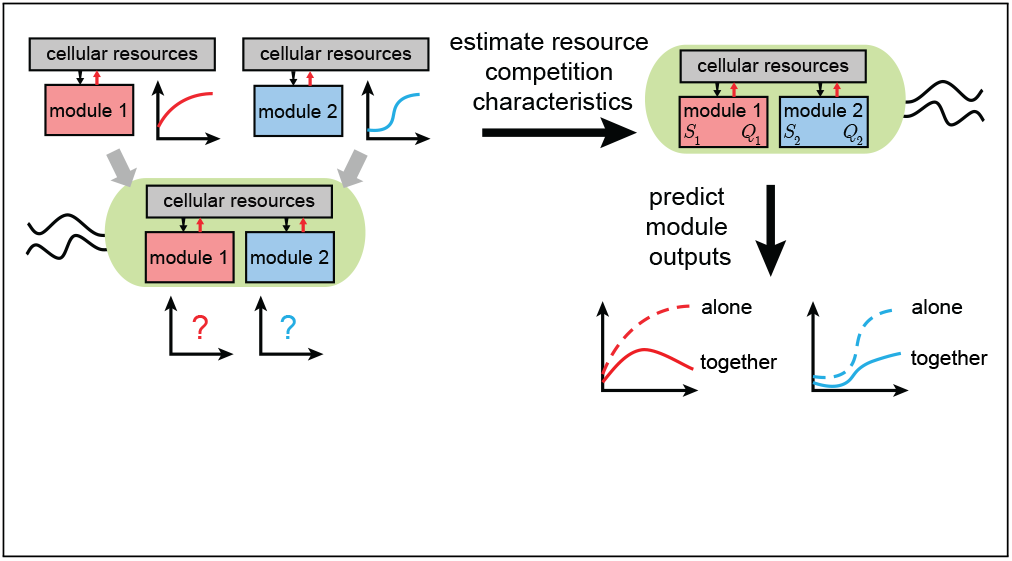

