## Supplementary material for "Predicting Composition of Genetic Circuits with Resource Competition: Demand and Sensitivity": Experimental data and Matlab code: metadataRS.m

```
%this file contains all metadata for loading in data, applying each
%exerimental condition to the relevant module and assigning properties for
%each module.

%filenames for loading in data for each dilution or each new experimental
%condition (according to indstruct)
filenamecell = {...
    'data\Exp1_3-02-21.txt','data\Exp2_3-02-21.txt',...
    'data\Exp1_3-03-21.txt','data\Exp2_3-03-21.txt',...
    'data\Exp1_3-04-21.txt','data\Exp2_3-04-21.txt',...
    'data\Exp1_3-05-21.txt','data\Exp2_3-05-21.txt',...
    'data\Exp1_3-10-21.txt','data\Exp2_3-10-21.txt',...
    'data\Exp1_3-12-21.txt','data\Exp2_3-12-21.txt',...
    'data\Exp1_3-18-21.txt','data\Exp2_3-18-21.txt',...
    'data\Exp1_3-24-21.txt','data\Exp2_3-24-21.txt',...
    'data\Exp3_3-24-21.txt'};
sheetcell = repmat({'none'},1,length(filenamecell));

%match wells on plates with experimental conditions
indstruct = cell(length(filenamecell),1);
indstruct{1}.RY192_0  = [1,1; 1,2; 1,3];
indstruct{1}.RY192_01 = [3,1; 3,2; 3,3];
%indstruct{1}.RY192_1  = [5,1; 5,2; 5,3];
%indstruct{1}.RY192_10 = [7,1; 7,2; 7,3];
%indstruct{1}.RY193_0  = [1,5; 1,6; 1,7];
%indstruct{1}.RY193_01 = [3,5; 3,6; 3,7];
%indstruct{1}.RY193_1  = [5,5; 5,6; 5,7];
%indstruct{1}.RY193_10 = [7,5; 7,6; 7,7];
indstruct{1}.GR221    = [1,9; 1,10; 1,11];
indstruct{1}.RY194_0  = [3,9; 3,10; 3,11];
indstruct{1}.RY194_01 = [4,9; 4,10; 4,11];
indstruct{1}.RY194_1  = [5,9; 5,10; 5,11];
indstruct{1}.RY194_10 = [7,9; 7,10; 7,11];
indstruct{1}.M9       = [2,4; 2,8; 4,4; 6,4; 6,8];
indstruct{2} = indstruct{1};

indstruct{3}.GY184_0  = [1,1; 1,2; 1,3];
indstruct{3}.GY184_01 = [3,1; 3,2; 3,3];
indstruct{3}.GY184_1  = [5,1; 5,2; 5,3];
indstruct{3}.GY184_10 = [7,1; 7,2; 7,3];
indstruct{3}.GY183_0  = [1,5; 1,6; 1,7];
indstruct{3}.GY183_01 = [3,5; 3,6; 3,7];
%indstruct{3}.GY183_1  = [5,5; 5,6; 5,7];
%indstruct{3}.GY183_10 = [7,5; 7,6; 7,7];
indstruct{3}.GY182_0  = [1,9; 1,10; 1,11];
indstruct{3}.GY182_01 = [3,9; 3,10; 3,11];
%indstruct{3}.GY182_1  = [5,9; 5,10; 5,11];
%indstruct{3}.GY182_10 = [7,9; 7,10; 7,11];
indstruct{3}.M9       = [2,4; 2,8; 4,4; 6,4; 6,8];
indstruct{4} = indstruct{3};

%indstruct{5}.BG200    = [1,1; 1,2; 1,3];
indstruct{5}.BG202    = [3,1; 3,2; 3,3];
indstruct{5}.BR211    = [5,1; 5,2; 5,3];
%indstruct{5}.GR220    = [7,1; 7,2; 7,3];
%indstruct{5}.BY172_0  = [1,5; 1,6; 1,7];
%indstruct{5}.BY172_01 = [3,5; 3,6; 3,7];
%indstruct{5}.BY172_1  = [5,5; 5,6; 5,7];
%indstruct{5}.BY172_10 = [7,5; 7,6; 7,7];
%indstruct{5}.BY173_0  = [1,9; 1,10; 1,11];
%indstruct{5}.BY173_01 = [3,9; 3,10; 3,11];
%indstruct{5}.BY173_1  = [5,9; 5,10; 5,11];
%indstruct{5}.BY173_10 = [7,9; 7,10; 7,11];
indstruct{5}.M9       = [2,4; 2,8; 4,4; 6,4; 6,8];
indstruct{6} = indstruct{5};

indstruct{7}.B171     = [1,1; 1,2; 1,3];
indstruct{7}.G181     = [3,1; 3,2; 3,3];
indstruct{7}.R190     = [5,1; 5,2; 5,3];
%indstruct{7}.BG201    = [7,1; 7,2; 7,3];
indstruct{7}.BY174_0  = [1,5; 1,6; 1,7];
indstruct{7}.BY174_01 = [3,5; 3,6; 3,7];
indstruct{7}.BY174_1  = [5,5; 5,6; 5,7];
indstruct{7}.BY174_10 = [7,5; 7,6; 7,7];
indstruct{7}.Y240_0   = [1,9; 1,10; 1,11];
indstruct{7}.Y240_01  = [3,9; 3,10; 3,11];
%indstruct{7}.Y240_1   = [5,9; 5,10; 5,11];
%indstruct{7}.Y240_10  = [7,9; 7,10; 7,11];
indstruct{7}.M9       = [2,4; 2,8; 4,4; 6,4; 6,8];
indstruct{8} = indstruct{7};

indstruct{9}.B170     = [1,1; 1,2; 1,3];
indstruct{9}.G180     = [3,1; 3,2; 3,3];
indstruct{9}.R191     = [5,1; 5,2; 5,3];
indstruct{9}.BG200    = [7,1; 7,2; 7,3];
%indstruct{9}.BG201    = [1,5; 1,6; 1,7];
indstruct{9}.BG203    = [3,5; 3,6; 3,7];
%indstruct{9}.BR210    = [5,5; 5,6; 5,7];
indstruct{9}.BR212    = [7,5; 7,6; 7,7];
indstruct{9}.BR213    = [1,9; 1,10; 1,11];
indstruct{9}.GR220    = [3,9; 3,10; 3,11];
indstruct{9}.GR222    = [5,9; 5,10; 5,11];
indstruct{9}.GR223    = [7,9; 7,10; 7,11];
indstruct{9}.M9       = [2,4; 2,8; 4,4; 6,4; 6,8];
indstruct{10} = indstruct{9};

indstruct{11}.BG201    = [1,1; 1,2; 1,3];
indstruct{11}.BR210    = [3,1; 3,2; 3,3];
%indstruct{11}.GY183_10 = [5,1; 5,2; 5,3];
%indstruct{11}.RY192_1  = [7,1; 7,2; 7,3];
%indstruct{11}.BY172_0  = [1,5; 1,6; 1,7];
%indstruct{11}.BY172_01 = [3,5; 3,6; 3,7];
%indstruct{11}.BY172_1  = [5,5; 5,6; 5,7];
%indstruct{11}.BY172_10 = [7,5; 7,6; 7,7];
indstruct{11}.BY173_0  = [1,9; 1,10; 1,11];
indstruct{11}.BY173_01 = [3,9; 3,10; 3,11];
%indstruct{11}.BY173_1  = [5,9; 5,10; 5,11];
%indstruct{11}.BY173_10 = [7,9; 7,10; 7,11];
indstruct{11}.M9       = [2,4; 2,8; 4,4; 6,4; 6,8];
indstruct{12} = indstruct{11};

indstruct{13}.BY172_0  = [1,1; 1,2; 1,3];
indstruct{13}.BY172_01 = [3,1; 3,2; 3,3];
indstruct{13}.BY172_1  = [5,1; 5,2; 5,3];
indstruct{13}.BY172_10 = [7,1; 7,2; 7,3];
indstruct{13}.RY193_0  = [1,5; 1,6; 1,7];
indstruct{13}.RY193_01 = [3,5; 3,6; 3,7];
%indstruct{13}.RY193_1  = [5,5; 5,6; 5,7];
%indstruct{13}.RY193_10 = [7,5; 7,6; 7,7];
indstruct{13}.BY174_1  = [1,9; 1,10; 1,11];
indstruct{13}.BY174_10 = [3,9; 3,10; 3,11];
%indstruct{13}.Y240_1   = [5,9; 5,10; 5,11];
%indstruct{13}.Y240_10  = [7,9; 7,10; 7,11];
indstruct{13}.M9       = [2,4; 2,8; 4,4; 6,4; 6,8];
indstruct{14}.BY172_0  = [7,1; 7,2; 7,3];
indstruct{14}.BY172_01 = [1,1; 1,2; 1,3];
indstruct{14}.BY172_1  = [5,1; 3,2; 3,3];
indstruct{14}.BY172_10 = [3,1; 5,2; 5,3];
indstruct{14}.RY193_0  = [7,5; 7,6; 7,7];
indstruct{14}.RY193_01 = [1,5; 1,6; 1,7];
%indstruct{14}.RY193_1  = [3,5; 3,6; 3,7];
%indstruct{14}.RY193_10 = [5,5; 5,6; 5,7];
indstruct{14}.BY174_1  = [1,9; 1,10; 1,11];
indstruct{14}.BY174_10 = [3,9; 3,10; 3,11];
%indstruct{14}.Y240_1   = [5,9; 5,10; 5,11];
%indstruct{14}.Y240_10  = [7,9; 7,10; 7,11];
indstruct{14}.M9       = [2,4; 2,8; 4,4; 6,4; 6,8];

indstruct{15}.BY173_1  = [1,1; 1,2; 1,3];
indstruct{15}.BY173_10 = [3,1; 3,2; 3,3];
indstruct{15}.GY182_1  = [5,1; 5,2; 5,3];
indstruct{15}.GY182_10 = [7,1; 7,2; 7,3];
indstruct{15}.GY183_1  = [1,5; 1,6; 1,7];
indstruct{15}.GY183_10 = [3,5; 3,6; 3,7];
indstruct{15}.RY192_1  = [5,5; 5,6; 5,7];
indstruct{15}.RY192_10 = [7,5; 7,6; 7,7];
indstruct{15}.RY193_1  = [1,9; 1,10; 1,11];
indstruct{15}.RY193_10 = [3,9; 3,10; 3,11];
indstruct{15}.Y240_1   = [5,9; 5,10; 5,11];
indstruct{15}.Y240_10  = [7,9; 7,10; 7,11];
indstruct{15}.M9       = [2,4; 2,8; 4,4; 6,4; 6,8];
indstruct{16} = indstruct{15};
indstruct{17} = indstruct{15};

%upper data bound for finding color correction matrix
maxstruct.BFP = 2e4;
maxstruct.GFP = 2.4e4;
maxstruct.RFP = 2e4;
maxstruct.YFP = 1e4;
maxstruct.OD  = 0.2;

%tell which fields should be zero to subtract background
%for background subtraction
mediastruct = struct;
mediastruct.OD  = 'M9';
mediastruct.BFP = 'M9';
mediastruct.GFP = 'M9';
mediastruct.RFP = 'M9';
mediastruct.YFP = 'M9';
ODcontrolstruct = [];

%calibration for adjusting changes in fluorescence due to changes in OD
ODFPstruct = struct;
ODFPstruct.BFP = {'R191','R190','G181','G180'}; %not BFP
ODFPstruct.GFP = {'R190','B171'}; %not GFP

[cellholder,basalstruct] = deal(cell(length(filenamecell),1));
%struct for compenstating for fluorescence bleed
clear C
FPbleedstruct = struct;
FPbleedstruct.BFP = {'BY172_0','BY173_0','B170','B171'};  %only BFP
FPbleedstruct.GFP = {'GY182_0','GY183_0','G181','G180'};
FPbleedstruct.RFP = {'RY192_0','RY193_0','R191','R190'};
FPbleedstruct.YFP = {'Y240_10','RY194_10','GY184_10'};

%sort module names for plotting
allnames = cellfun(@fieldnames,indstruct,'UniformOutput',false);
allnames = unique(vertcat(allnames{:}));
bluenames   = allnames(contains(allnames,'B'));
greennames  = allnames(contains(allnames,'G'));
rednames    = allnames(contains(allnames,'R'));
yellownames = allnames(contains(allnames,'Y'));
constnames  = allnames(~contains(allnames,'_'));
bluenames1   = allnames(contains(allnames,'B') & ~contains(allnames,'_'));
greennames1  = allnames(contains(allnames,'G') & ~contains(allnames,'_'));
rednames1    = allnames(contains(allnames,'R') & ~contains(allnames,'_'));
yellownames1 = allnames(contains(allnames,'Y') & ~contains(allnames,'4'));
bluenames2   = allnames(contains(allnames,'B') & contains(allnames,'_'));
greennames2  = allnames(contains(allnames,'G') & contains(allnames,'_'));
rednames2    = allnames(contains(allnames,'R') & contains(allnames,'_'));
yellownames2 = allnames(contains(allnames,'Y') & contains(allnames,'4'));
notbluenames   = allnames(~contains(allnames,'B') & ~contains(allnames,'_') & ~contains(allnames,'M9'));
notgreennames  = allnames(~contains(allnames,'G') & ~contains(allnames,'_') & ~contains(allnames,'M9'));
notrednames    = allnames(~contains(allnames,'R') & ~contains(allnames,'_') & ~contains(allnames,'M9'));
notyellownames = allnames(~contains(allnames,'Y') & ~contains(allnames,'_') & ~contains(allnames,'M9'));

%windows where production rate is in steady state for each experimental
%condition
ssstruct = struct;
ssstruct.B170  = [6.9300, 11.3300];
ssstruct.B171  = [7.0300, 9.5300];
ssstruct.G180  = [6.7300, 9.4300];
ssstruct.G181  = [7.3300, 10.8300];
ssstruct.R190  = [6.8300, 9.8300];
ssstruct.R191  = [5.7300, 10.2300];
ssstruct.BG200 = [6.6700, 11.8700];
ssstruct.BG201 = [7.2700, 11.6700];
ssstruct.BG202 = [7.1200, 10.4200];
ssstruct.BG203 = [5.6200, 10.2200];
ssstruct.BR210 = [7.7000, 12.5000];
ssstruct.BR211 = [6.6200, 10.2200];
ssstruct.BR212 = [6.4200, 11.2200];
ssstruct.BR213 = [7.9200, 12.9200];
ssstruct.GR220 = [7.6700, 9.6700];
ssstruct.GR221 = [7.6200, 10.8200];
ssstruct.GR222 = [7.5200, 11.6200];
ssstruct.GR223 = [6.8200, 11.9200];
ssstruct.BY172_0  = [7.4200, 12.1200];
ssstruct.BY172_01 = [7.8200, 12.4200];
ssstruct.BY172_1  = [7.7200, 12.1200];
ssstruct.BY172_10 = [6.6200, 11.5200];
ssstruct.BY173_0  = [8.1700, 10.4700];
ssstruct.BY173_01 = [8.4700, 10.1700];
ssstruct.BY173_1  = [11.2200, 14.2200];
ssstruct.BY173_10 = [10.6200, 13.9200];
ssstruct.BY174_0  = [2.0, 4.0];
ssstruct.BY174_01 = [2.0, 4.1];
ssstruct.BY174_1  = [1.8, 3.6];
ssstruct.BY174_10 = [1.9, 4.0];
ssstruct.GY182_0  = [7.3, 10.8];
ssstruct.GY182_01 = [7.6, 11.3];
ssstruct.GY182_1  = [13.9800, 11.2800];
ssstruct.GY182_10 = [11.6800, 14.7800];
ssstruct.GY183_0  = [7.6, 10.5];
ssstruct.GY183_01 = [7.2, 10.2];
ssstruct.GY183_1  = [10.9800, 13.0800];
ssstruct.GY183_10 = [11.3300, 12.9300];
ssstruct.GY184_0  = [7.0, 9.0];
ssstruct.GY184_01 = [7.0, 9.0];
ssstruct.GY184_1  = [6.7, 9.3];
ssstruct.GY184_10 = [7.0, 9.5];
ssstruct.RY192_0  = [6.7, 10.4];
ssstruct.RY192_01 = [7.7, 10.7];
ssstruct.RY192_1  = [5.2000, 7.8000];
ssstruct.RY192_10 = [5.4500, 7.7500];
ssstruct.RY193_0  = [5.9500, 9.7500];
ssstruct.RY193_01 = [5.5500, 9.4500];
ssstruct.RY193_1  = [10.3000, 13.5000];
ssstruct.RY193_10 = [10.1000, 12.8000];
ssstruct.RY194_0  = [6.0, 9.0];
ssstruct.RY194_01 = [6.0, 9.0];
ssstruct.RY194_1  = [6.8, 9.7];
ssstruct.RY194_10 = [7.2, 10.1];
ssstruct.Y240_0   = [6.9200, 10.1200];
ssstruct.Y240_01  = [6.3200, 9.3200];
ssstruct.Y240_1   = [10.1500, 12.5500];
ssstruct.Y240_10  = [11.0500, 13.0500];

%struct assigning induction levels for each module. If module is
%constitutive, input u = 0.
inductionmeta = struct;
inductionmeta.B170.mods  = {'B170'};
inductionmeta.B170.u     = 0;
inductionmeta.B171.mods  = {'B171'};
inductionmeta.B171.u     = 0;
inductionmeta.G180.mods  = {'G180'};
inductionmeta.G180.u     = 0;
inductionmeta.G181.mods  = {'G181'};
inductionmeta.G181.u     = 0;
inductionmeta.R190.mods  = {'R190'};
inductionmeta.R190.u     = 0;
inductionmeta.R191.mods  = {'R191'};
inductionmeta.R191.u     = 0;
inductionmeta.BG200.mods  = {'BG200'};
inductionmeta.BG200.u     = 0;
inductionmeta.BG201.mods  = {'BG201'};
inductionmeta.BG201.u     = 0;
inductionmeta.BG202.mods  = {'BG202'};
inductionmeta.BG202.u     = 0;
inductionmeta.BG203.mods  = {'BG203'};
inductionmeta.BG203.u     = 0;
inductionmeta.BR210.mods  = {'BR210'};
inductionmeta.BR210.u     = 0;
inductionmeta.BR211.mods  = {'BR211'};
inductionmeta.BR211.u     = 0;
inductionmeta.BR212.mods  = {'BR212'};
inductionmeta.BR212.u     = 0;
inductionmeta.BR213.mods  = {'BR213'};
inductionmeta.BR213.u     = 0;
inductionmeta.GR220.mods  = {'GR220'};
inductionmeta.GR220.u     = 0;
inductionmeta.GR221.mods  = {'GR221'};
inductionmeta.GR221.u     = 0;
inductionmeta.GR222.mods  = {'GR222'};
inductionmeta.GR222.u     = 0;
inductionmeta.GR223.mods  = {'GR223'};
inductionmeta.GR223.u     = 0;
inductionmeta.BY172.mods = {'BY172_0','BY172_01','BY172_1','BY172_10'};
inductionmeta.BY172.u    = [0, 0.1, 1, 10];
inductionmeta.BY173.mods = {'BY173_0','BY173_01','BY173_1','BY173_10'};
inductionmeta.BY173.u    = [0, 0.1, 1, 10];
inductionmeta.GY182.mods = {'GY182_0','GY182_01','GY182_1','GY182_10'};
inductionmeta.GY182.u    = [0, 0.1, 1, 10];
inductionmeta.GY183.mods = {'GY183_0','GY183_01','GY183_1','GY183_10'};
inductionmeta.GY183.u    = [0, 0.1, 1, 10];
inductionmeta.RY192.mods = {'RY192_0','RY192_01','RY192_1','RY192_10'};
inductionmeta.RY192.u    = [0, 0.1, 1, 10];
inductionmeta.RY193.mods = {'RY193_0','RY193_01','RY193_1','RY193_10'};
inductionmeta.RY193.u    = [0, 0.1, 1, 10];
inductionmeta.Y240.mods  = {'Y240_0','Y240_01','Y240_1','Y240_10'};
inductionmeta.Y240.u     = [0, 0.1, 1, 10];

%metadata struct for assigning properties to each module and describing
%which experimental data applies to which module
modulemetadata = struct;
modulemetadata.B170.isResourceSensor = false;
modulemetadata.B171.isResourceSensor = true;
modulemetadata.G180.isResourceSensor = false;
modulemetadata.G181.isResourceSensor = false;
modulemetadata.R190.isResourceSensor = false;
modulemetadata.R191.isResourceSensor = true;
modulemetadata.Y240.isResourceSensor = false;
modulemetadata.B170.containingmods = {'B170'};
modulemetadata.B171.containingmods = {'B171'};
modulemetadata.G180.containingmods = {'G180'};
modulemetadata.G181.containingmods = {'G181'};
modulemetadata.R190.containingmods = {'R190'};
modulemetadata.R191.containingmods = {'R191'};
modulemetadata.Y240.containingmods = {'Y240'};

%modules together
modulemetadata.BG200.containingmods = {'B170','G180'};
modulemetadata.BG200.isResourceSensor = false;
modulemetadata.BG201.containingmods = {'B171','G180'};
modulemetadata.BG201.isResourceSensor = false;
modulemetadata.BG202.containingmods = {'B171','G181'};
modulemetadata.BG202.isResourceSensor = false;
modulemetadata.BG203.containingmods = {'B170','G181'};
modulemetadata.BG203.isResourceSensor = false;
modulemetadata.BR210.containingmods = {'B170','R190'};
modulemetadata.BR210.isResourceSensor = false;
modulemetadata.BR211.containingmods = {'B171','R190'};
modulemetadata.BR211.isResourceSensor = false;
modulemetadata.BR212.containingmods = {'B171','R191'};
modulemetadata.BR212.isResourceSensor = true;
modulemetadata.BR213.containingmods = {'B170','R191'};
modulemetadata.BR213.isResourceSensor = false;
modulemetadata.GR220.containingmods = {'G180','R190'};
modulemetadata.GR220.isResourceSensor = false;
modulemetadata.GR221.containingmods = {'G181','R190'};
modulemetadata.GR221.isResourceSensor = false;
modulemetadata.GR222.containingmods = {'G180','R191'};
modulemetadata.GR222.isResourceSensor = false;
modulemetadata.GR223.containingmods = {'G181','R191'};
modulemetadata.GR223.isResourceSensor = false;
modulemetadata.BY172.containingmods = {'B170','Y240'};
modulemetadata.BY172.isResourceSensor = false;
modulemetadata.BY173.containingmods = {'B171','Y240'};
modulemetadata.BY173.isResourceSensor = false;
modulemetadata.GY182.containingmods = {'G181','Y240'};
modulemetadata.GY182.isResourceSensor = false;
modulemetadata.GY183.containingmods = {'G180','Y240'};
modulemetadata.GY183.isResourceSensor = false;
modulemetadata.RY192.containingmods = {'R191','Y240'};
modulemetadata.RY192.isResourceSensor = false;
modulemetadata.RY193.containingmods = {'R190','Y240'};
modulemetadata.RY193.isResourceSensor = false;

modulemetadata.B170.FPout = {'BFPdiffOD'};
modulemetadata.B171.FPout = {'BFPdiffOD'};
modulemetadata.G180.FPout = {'GFPdiffOD'};
modulemetadata.G181.FPout = {'GFPdiffOD'};
modulemetadata.R190.FPout = {'RFPdiffOD'};
modulemetadata.R191.FPout = {'RFPdiffOD'};
modulemetadata.Y240.FPout = {'YFPdiffOD'};
modulemetadata.BG200.FPout = {'BFPdiffOD','GFPdiffOD'};
modulemetadata.BG201.FPout = {'BFPdiffOD','GFPdiffOD'};
modulemetadata.BG202.FPout = {'BFPdiffOD','GFPdiffOD'};
modulemetadata.BG203.FPout = {'BFPdiffOD','GFPdiffOD'};
modulemetadata.BR210.FPout = {'BFPdiffOD','RFPdiffOD'};
modulemetadata.BR211.FPout = {'BFPdiffOD','RFPdiffOD'};
modulemetadata.BR212.FPout = {'BFPdiffOD','RFPdiffOD'};
modulemetadata.BR213.FPout = {'BFPdiffOD','RFPdiffOD'};
modulemetadata.GR220.FPout = {'GFPdiffOD','RFPdiffOD'};
modulemetadata.GR221.FPout = {'GFPdiffOD','RFPdiffOD'};
modulemetadata.GR222.FPout = {'GFPdiffOD','RFPdiffOD'};
modulemetadata.GR223.FPout = {'GFPdiffOD','RFPdiffOD'};
modulemetadata.BY172.FPout = {'BFPdiffOD','YFPdiffOD'};
modulemetadata.BY173.FPout = {'BFPdiffOD','YFPdiffOD'};
modulemetadata.GY182.FPout = {'GFPdiffOD','YFPdiffOD'};
modulemetadata.GY183.FPout = {'GFPdiffOD','YFPdiffOD'};
modulemetadata.RY192.FPout = {'RFPdiffOD','YFPdiffOD'};
modulemetadata.RY193.FPout = {'RFPdiffOD','YFPdiffOD'};

modulemetadata.B170.isalone = true;
modulemetadata.B171.isalone = true;
modulemetadata.G180.isalone = true;
modulemetadata.G181.isalone = true;
modulemetadata.R190.isalone = true;
modulemetadata.R191.isalone = true;
modulemetadata.Y240.isalone = true;
modulemetadata.BY172.isalone = false;
modulemetadata.BY173.isalone = false;
modulemetadata.GY182.isalone = false;
modulemetadata.GY183.isalone = false;
modulemetadata.RY192.isalone = false;
modulemetadata.RY193.isalone = false;
modulemetadata.BG200.isalone = false;
modulemetadata.BG201.isalone = false;
modulemetadata.BG202.isalone = false;
modulemetadata.BG203.isalone = false;
modulemetadata.BR210.isalone = false;
modulemetadata.BR211.isalone = false;
modulemetadata.BR212.isalone = false;
modulemetadata.BR213.isalone = false;
modulemetadata.GR220.isalone = false;
modulemetadata.GR221.isalone = false;
modulemetadata.GR222.isalone = false;
modulemetadata.GR223.isalone = false;
```
