## Supplementary material for "Predicting Composition of Genetic Circuits with Resource Competition: Demand and Sensitivity": Experimental data and Matlab code: README.txt

Matlab files for the paper:
C McBride and D Del Vecchio, "Predicting Composition of Genetic Circuits with Resource Competition: Demand and Sensitivity", 2021

metadatRS.m: describes how runplate_RS4.m should load and analyze data.
runplate_RS4.m: loads data, analyzes data, creates plots describing Q and S for modules and predictions based on the helper functions in the folder utility. Data is stored in the custom matlab classes RSexpdata and RSmodule.

These files were developed using Matlab R2020a on Windows 10.
runplate_RS4.m should be able to run as is.
