## Supplementary material for "Predicting Composition of Genetic Circuits with Resource Competition: Demand and Sensitivity": Experimental data and Matlab code: runplate_RS4.m

```
addpath utility

if 0
%load in metadata describing modules and steady state locations
metadataRS;

%-----------------------------------
%init
[cellholder,basalstruct] = deal(cell(length(filenamecell),1));
%load in fluorescence correction matrix and scattering correlations COD.
%this data can also be created by the commands ODFPcorrection() and
%fluorescencebleed2() on lines 50 and 52
load('C.mat')

%load in data and apply corrections
for ii = 1:length(filenamecell)
    %init data and load data from file
    platedata = RSexpdata;
    platedata = loaddata(platedata,filenamecell{ii},sheetcell{ii});
    platedata.FPfields = {'BFP','GFP','RFP','YFP'};
    platedata.samplename = filenamecell{ii};
    %repackage data into individual fields for each experimental condition
    cellstruct0 = bulkdata2cells2(platedata,indstruct{ii});
    %replace outliers in OD data
    cellstruct0 = replaceoutliers(cellstruct0,'OD');
    %subtract the basal fluorescence based on background media vs time
    [cellstruct0, basalstruct] = subtractbasal7(cellstruct0,mediastruct,ODcontrolstruct);
    
    if exist('C','var')
        %subtract off fluorescence due to OD on BFP and GFP channels
        [cellstruct0,COD] = ODFPcorrection(cellstruct0,ODFPstruct,maxstruct,COD);
        %subtract off fluorescence bleed
        cellstruct0 = applycolorcorrection(cellstruct0,C);
        %fit growthrate
        cellstruct0 = fitgrowthrate3(cellstruct0,false);
    end
    %take time derivative of raw fluorescent data
    cellstruct0 = diffcellfields(cellstruct0,{'BFP','GFP','RFP','YFP','OD'},'time');
    %store data from batch
    cellholder{ii} = cellstruct0;
end

%combine batches into one
cellstruct = combinedilutions(cellholder);

if ~exist('C','var')
    %find fluorescence corrections if not previously found
    %get correction factor for change in FP due to change in OD
    [cellstruct,COD] = ODFPcorrection(cellstruct,ODFPstruct,maxstruct,[],true,1);
    %find fluorescence bleed between channels
    C = fluorescencebleed2(cellstruct,FPbleedstruct,maxstruct,true,[6,7,8,9]);
    %need to re-run script to apply fluorescence corrections to data if
    %this code was excecuted
end

%check compensation--Cout should be close to identity
Cout = fluorescencebleed2(cellstruct,FPbleedstruct,maxstruct,false);
%divide raw fluorescence and time derivative of fluorescence by OD
cellstruct = normalizeFP3(cellstruct,{'BFP','GFP','RFP','YFP',...
    'BFPdiff','GFPdiff','RFPdiff','YFPdiff'});
else
    load(data2.mat)
end

%---------------------
%can skip previous code section and load('data2.mat') instead (need to
%addpath utility first)

%average over time windows to get steady state
modulestruct = findss4(cellstruct,ssstruct,{'BFPdiffOD','GFPdiffOD',...
    'RFPdiffOD','YFPdiffOD'},@mean,false);
%combine submodules for each dilution level
modulestruct2 = combineinductions(modulestruct,inductionmeta);
%add metadata to modules
modulestruct3 = addmetadata(modulestruct2,modulemetadata);
%calculate resource demand for resource sensor
RSmodules = calcRS3(modulestruct3,true,7,6);
%find Q and S for non-resource sensor modules
[modulesout,RSmodules2] = calcQS3(modulestruct3,RSmodules,true,8,9,10,11,12,13,14,15);
%combine modules together
modulesout2 = combinemodstructs(RSmodules2,modulesout,modulestruct3);
%predict outputs
modsout3 = predictRS6(modulesout2,true,[16,17,18,19,20,21,22],true,23);
%plot growth rate and resource demand
[modsout4,grfitobj,gof] = plotGRvsQ(modulesout2,true,24);
%plot growth rate bar graphs
plotGR(modulesout2)

return

%---------------------------
%plotting raw data vs time, etc
plotsubfield2(cellstruct,'time','ODdiff',2,allnames(51:end),ssstruct);
setallsubplots(2,'axis',{'ylim'},{[0,0.8]});

plotsubfield2(cellstruct,'time','OD',2,yellownames(1:12),ssstruct);
setallsubplots(2,'axis',{'ylim'},{[0,Inf]});
plotsubfield2(cellstruct,'time','YFPdiffOD',5,yellownames(1:12),ssstruct);
setallsubplots(5,'axis',{'ylim'},{[0,12e4]});
plotsubfield2(cellstruct,'time','BFPdiffOD',1,yellownames(1:12),ssstruct);
setallsubplots(1,'axis',{'ylim'},{[0,Inf]});

%plot steady state production rates
plotsubfield2(cellstruct,'time','OD',1,'BG201',ssstruct);
setallsubplots(1,'axis',{'yscale','ylim'},{'linear',[1e-3,0.5]});
plotsubfield2(cellstruct,'time','BFPdiffOD',2,bluenames(1:10),ssstruct);
setallsubplots(2,'axis',{'ylim'},{[0,12e4]});
plotsubfield2(cellstruct,'time','GFPdiffOD',3,greennames(1:10),ssstruct);
setallsubplots(3,'axis',{'ylim'},{[0,Inf]});
plotsubfield2(cellstruct,'time','RFPdiffOD',4,rednames(1:10),ssstruct);
setallsubplots(4,'axis',{'ylim'},{[0,Inf]});

%plot steady state production rates
plotsubfield2(cellstruct,'time','OD',1,constnames([1,2,11,12,18,19,3,4,5,6,7,8,9,10,13,14,15,16]),ssstruct,@(OD) (OD));
setallsubplots(1,'axis',{'yscale','ylim','xlim'},{'log',[1e-3,0.3],[0,15]});
plotsubfield2(cellstruct,'time','OD',2,yellownames([1:8,13:22]),ssstruct,@(OD) (OD));
setallsubplots(2,'axis',{'yscale','ylim','xlim'},{'log',[1e-3,0.3],[0,15]});
plotsubfield2(cellstruct,'time','OD',3,yellownames(23:40),ssstruct,@(OD) (OD));
setallsubplots(3,'axis',{'yscale','ylim','xlim'},{'log',[1e-3,0.3],[0,15]});

plotsubfield2(cellstruct,'time','BFPdiffOD',2,bluenames(11:18),ssstruct);
setallsubplots(2,'axis',{'ylim'},{[0,Inf]});
plotsubfield2(cellstruct,'time','GFPdiffOD',3,greennames(11:18),ssstruct);
setallsubplots(3,'axis',{'ylim'},{[0,Inf]});
plotsubfield2(cellstruct,'time','RFPdiffOD',4,rednames(11:18),ssstruct);
setallsubplots(4,'axis',{'ylim'},{[0,Inf]});

plotsubfield2(cellstruct,'time','YFPdiffOD',5,yellownames(1:8),ssstruct);
setallsubplots(5,'axis',{'ylim','xlim'},{[0,6e4],[0,15]});
plotsubfield2(cellstruct,'time','YFPdiffOD',6,yellownames(13:20),ssstruct);
setallsubplots(6,'axis',{'ylim','xlim'},{[0,6e4],[0,20]});
plotsubfield2(cellstruct,'time','YFPdiffOD',7,yellownames(25:32),ssstruct);
setallsubplots(7,'axis',{'ylim','xlim'},{[0,6e4],[0,15]});
plotsubfield2(cellstruct,'time','YFPdiffOD',8,yellownames(33:end),ssstruct);
setallsubplots(8,'axis',{'ylim','xlim'},{[0,6e4],[0,15]});

%plot steady state production rates
plotsubfield2(cellstruct,'time','YFPdiffOD',5,yellownames([23,24,35,36,39,40]),ssstruct);
setallsubplots(5,'axis',{'ylim'},{[0,4e4]});

plotsubfield2(cellstruct,'time','GFPdiffOD',5,{'BY172_10','BY173_10',...
    'GY182_10','GY183_10','RY192_1','RY193_10','Y240_10'},ssstruct);
setallsubplots(5,'axis',{'ylim'},{[0,1.5e4]});

plotsubfield2(cellstruct,'time','GFPOD',5);
setallsubplots(5,'axis',{'ylim'},{[0,2e4]});

%plotting for checking off genes don't fluoresce
plotsubfield2(cellstruct,'time','BFP',1,{'G180','G181','R190','R191','GR220','GR221','GR222','GR223',...
    'BY174_0','BY174_01','BY174_1','BY174_10'});
setallsubplots(1,'axis',{'xlim','ylim'},{[0,10],[-200,200]});
plotsubfield2(cellstruct,'time','GFP',2,{'B170','B171','R190','R191','BR210','BR211','BR212','BR213',...
    'GY184_0','GY184_01','GY184_1','GY184_10'});
setallsubplots(2,'axis',{'xlim','ylim'},{[0,10],[-80,80]});
plotsubfield2(cellstruct,'time','RFP',3,{'G180','G181','B170','B171','BG200','BG201','BG202','BG203',...
    'RY194_0','RY194_01','RY194_1','RY194_10'});
setallsubplots(3,'axis',{'xlim','ylim'},{[0,10],[-70,70]});
plotsubfield2(cellstruct,'time','YFP',4,{'B170','B171','G180','G181','R190','R191','BG200','BG201','BG202','BG203',...
    'BR210','BR211','BR212','BR213','GR220','GR221','GR222','GR223'});
setallsubplots(4,'axis',{'xlim','ylim'},{[0,10],[-100,100]});
plotsubfield2(cellstruct,'time','BFP',5,{'B170','B171',...
    'BG200','BG201','BG202','BG203','BR210','BR211','BR212','BR213',...
    'BY172_0','BY172_01','BY172_1','BY172_10','BY173_0','BY173_01','BY173_1','BY173_10'});
setallsubplots(5,'axis',{'xlim'},{[0,10]});
plotsubfield2(cellstruct,'time','GFP',6,{'G180','G181',...
    'BG200','BG201','BG202','BG203','GR220','GR221','GR222','GR223',...
    'GY182_0','GY182_01','GY182_1','GY182_10','GY183_0','GY183_01','GY183_1','GY183_10'});
setallsubplots(6,'axis',{'xlim'},{[0,10]});
plotsubfield2(cellstruct,'time','RFP',7,{'R190','R191',...
    'BR210','BR211','BR212','BR213','GR220','GR221','GR222','GR223',...
    'RY192_0','RY192_01','RY192_1','RY192_10','RY193_0','RY193_01','RY193_1','RY193_10'});
setallsubplots(7,'axis',{'xlim'},{[0,10]});
plotsubfield2(cellstruct,'time','YFP',8,{...
    'BY172_0','BY172_01','BY172_1','BY172_10','BY173_0','BY173_01','BY173_1','BY173_10',...
    'GY182_0','GY182_01','GY182_1','GY182_10','GY183_0','GY183_01','GY183_1','GY183_10',...
    'RY192_0','RY192_01','RY192_1','RY192_10','RY193_0','RY193_01','RY193_1','RY193_10'});
setallsubplots(8,'axis',{'xlim'},{[0,10]});
```
