## Supplementary material for "Predicting Composition of Genetic Circuits with Resource Competition: Demand and Sensitivity": Experimental data and Matlab code: bulkdata2cells2.m

```
function cellout = bulkdata2cells2(platedata, indstruct)
%convert raw data in an object to sorted by individual cells in an object

if nargin < 2
    indstruct = struct;
    indstruct.R = 1:3;
    indstruct.YG = 4:6;
    indstruct.G = 11:13;
    indstruct.GR = 14:16;
    indstruct.Y = 21:23;
    indstruct.YR = 24:26;
    indstruct.M9 = [7,17,27,34:37,41:44];
    indstruct.T10 = 31:33;
    indstruct.none = [8:10,18:20,28:30,38:40,45:60];
end

%init structs
cellout = struct;
cellnames = fieldnames(indstruct);
FPfields0 = platedata.FPfields;
%check to make sure FP fields aren't empty
inds = false(length(FPfields0),1);
for m = 1:length(FPfields0)
    inds(m) = ~isempty(platedata.(FPfields0{m}));
end
FPfields = FPfields0(inds);
ODfields = platedata.ODfield;
datafields = fieldnames(platedata);

%loop across cell names
for ii = 1:length(cellnames)
    celldata = RSexpdata;
    %loop through datafields
    for jj = 1:length(datafields)
        %reshape FP data and OD data
        if any(contains(FPfields,datafields{jj})) || any(contains(ODfields,datafields{jj}))
            numsamples = size(platedata.(datafields{jj}),3);
            tmp = permute(platedata.(datafields{jj}),[3,1,2]);   %put time field first
            data = zeros(numsamples,size(indstruct.(cellnames{ii}),1));
            %loop through each sample repeat
            for k = 1:size(indstruct.(cellnames{ii}),1)
                %add data into array
                data(:,k) = tmp(:,indstruct.(cellnames{ii})(k,1),indstruct.(cellnames{ii})(k,2));
            end
            %copy data into new object
            celldata.(datafields{jj}) = data;
        else
            %copy data into new object
            celldata.(datafields{jj}) = platedata.(datafields{jj});
        end
    end
    %output into struct of experimental conditions
    celldata.samplename = cellnames{ii};
    celldata.inducerconc = cellnames{ii};
    cellout.(cellnames{ii}) = celldata;
end
```
