## Supplementary material for "Predicting Composition of Genetic Circuits with Resource Competition: Demand and Sensitivity": Experimental data and Matlab code: filtplatedata3.m

```
function filtdata = filtplatedata3(platedata,wn,filtopts)
%filtdata = filtplatedata(datastruct,wn)
%run all data in datastruct through forward/backwards butterworth filter.
%If field has 'time' in it, the field is copied directly. wn is the filter
%cutoff frequency and is a value between 0 and 1 with 1 corresponding to
%the Nyquist frequency

if nargin < 3
    filtopts = 'low';
end
if nargin < 2
    wn = 0.04;
end
%input error checking
if isempty(wn) || wn >= 1 || wn <= 0
    %warning('data not filtered')
    filtdata = platedata;
    return
end

%init filter
filtdata = platedata;
[b,a] = butter(5,wn,filtopts);

%get fields to filter through
FPfields = platedata.FPfields;
ODfield = platedata.ODfield;
dataprops = [FPfields(:); ODfield(:)];

%filter data
for jj = 1:length(dataprops)
    %recursion for structs within structs
    if isstruct(platedata.(dataprops{jj}))
        filtdata.(dataprops{jj}) = filtplatedata2(platedata.(dataprops{jj}),wn,filtopts);
    else
        %filter along the 3rd dimension only
        datatemp = platedata.(dataprops{jj});
        for k = 1:size(datatemp,1)
            for ii = 1:size(datatemp,2)
                if all(isfinite(datatemp(k,ii,:)))
                    %filter data
                    filtdata.(dataprops{jj})(k,ii,:) = ...
                        filtfilt(b,a,squeeze(datatemp(k,ii,:)));
                else
                    %skip filtering if data has Inf's or NaN's
                    filtdata.(dataprops{jj})(k,ii,:) = datatemp(k,ii,:);
                end
            end
        end
    end
end
```
