## Supplementary material for "Predicting Composition of Genetic Circuits with Resource Competition: Demand and Sensitivity": Experimental data and Matlab code: RSexpdata.m

```
classdef RSexpdata
    properties
        samplename
        inducerconc
        time                                %units of hours
        BFP
        GFP
        RFP
        YFP
        OD
        timestep = 5/60        %measurements every 5 mins
        FPfields cell = {'BFP','GFP','RFP'}
        ODfield cell = {'OD'}
        growthrate
        growthratestd
        BFPdiff
        GFPdiff
        RFPdiff
        YFPdiff
        ODdiff
        BFPOD
        RFPOD
        GFPOD
        YFPOD
        BFPdiffOD
        GFPdiffOD
        RFPdiffOD
        YFPdiffOD
    end
    methods
        function obj = loaddata(obj,filename,sheet)
            if nargin < 3; sheet = 1; end
            if contains(filename,'.xls')
                obj0 = loadxlsdata(filename,sheet);
                if isempty(obj.time)
                    timevec = obj0.time(:);
                else
                    timeprev = obj.time(end);
                    timevec = [obj.time(:); obj0.time + timeprev + obj0.timestep];
                end
                obj = obj0;
                obj.time = timevec;
            elseif contains(filename,'.txt')
                platedata = importplate(filename);
                %load([filename,'.mat'],'platedata')
                obj.OD = platedata.OD600_600;
                obj.RFP = platedata.RFP1_584_619;
                if isfield(platedata,'GFP1_485_513')
                    obj.GFP = platedata.GFP1_485_513;
                elseif isfield(platedata,'GFP1_485_530')
                    obj.GFP = platedata.GFP1_485_530;
                end
                obj.BFP = platedata.GFP1_400_460;
                if isfield(platedata,'RFP1_520_550')
                    obj.YFP = platedata.RFP1_520_550;
                elseif isfield(platedata,'RFP1_510_541')
                    obj.YFP = platedata.RFP1_510_541;
                elseif isfield(platedata,'RFP1_520_545')
                    obj.YFP = platedata.RFP1_520_545;
                end
                %add time
                numsamples = max(structfun(@(x) size(x,3), platedata));
                timevec = obj.timestep*(0:(numsamples-1));
                if isempty(obj.time)
                    obj.time = timevec(:);
                else
                    timeprev = obj.time(end);
                    obj.time = [obj.time(:); timevec + timeprev + obj.timestep];
                end
            else
                error('need file extension')
            end
        end
    end
end
```
