## Supplementary material for "Predicting Composition of Genetic Circuits with Resource Competition: Demand and Sensitivity": Experimental data and Matlab code: RSmodule.m

```
classdef RSmodule
    properties
        u = []
        FPout cell
        containingmods cell
        BFPdiffOD
        GFPdiffOD
        RFPdiffOD
        YFPdiffOD
        BFPdiffODstd
        GFPdiffODstd
        RFPdiffODstd
        YFPdiffODstd
        y
        ystd
        perturbname
        perturby
        perturbystd
        predperturby
        predperturbystd
        Q
        Qstd
        S
        Sstd
        isResourceSensor logical
        isalone logical
        growthrate
        growthratestd
    end
    methods
        function plotmoduleIO(RSmodule,varargin)
            if length(RSmodule.u) > 1
                plot(RSmodule.u,RSmodule.y,varargin{:})
                if ~isempty(RSmodule.ystd)
                    hold on;
                    errorbar(RSmodule.u,RSmodule.ystd,'.k')
                end
            else
                bar(RSmodule.y,varargin{:})
                errorbar(RSmodule.ystd,'.k')
            end
        end
    end
end
```
