## Supplementary material for "Predicting Composition of Genetic Circuits with Resource Competition: Demand and Sensitivity": Experimental data and Matlab code: addmetadata.m

```
function outstruct = addmetadata(modulestruct,metadatastruct)
%add metadata to modules

cellnames = intersect(fieldnames(metadatastruct),fieldnames(modulestruct),'stable');
outstruct = modulestruct;

%loop through cell conditions
for ii = 1:length(cellnames)
    
    %unpack data and metadata
    metadata = metadatastruct.(cellnames{ii});
    moduledata = modulestruct.(cellnames{ii});
    metadatafields = fieldnames(metadata);
    
    %loop through metadata fields
    for jj = 1:length(metadatafields)
        
        if isprop(moduledata,metadatafields{jj}) || isfield(moduledata,metadatafields{jj})
            %add metadata to module data
            moduledata.(metadatafields{jj}) = metadata.(metadatafields{jj});
        end
    end
    outstruct.(cellnames{ii}) = moduledata;
end
```
