## Supplementary material for "Predicting Composition of Genetic Circuits with Resource Competition: Demand and Sensitivity": Experimental data and Matlab code: applycolorcorrection.m

```
function outstruct = applycolorcorrection(celldata,C,metastruct)


if nargin < 3 || isempty(metastruct)
    metastruct.BFP = {'B'};
    metastruct.GFP = {'G'};
    metastruct.RFP = {'R'};
    metastruct.YFP = {'Y'};
end

%ignore any values of C that are negative or insignificant
C2 = C;
C2(C2 < 0.01) = 0;
%chop off columns that are all 0's
emptyinds = all(C2 == 0);
C3 = C2(~emptyinds,~emptyinds);

%init
cellnames = fieldnames(celldata);
datafields = fieldnames(metastruct);
datafields2 = datafields(~emptyinds);
outstruct = celldata;

%loop through each experimental condition
for ii = 1:length(cellnames)
    %init
    [p,q] = size(celldata.(cellnames{ii}).(datafields2{1}));
    tmp = zeros(p,q,nnz(~emptyinds));
    %unpack data
    for jj = 1:length(datafields2)
        tmp(:,:,jj) = celldata.(cellnames{ii}).(datafields2{jj});
    end
    %reshape
    data2 = shiftdim(tmp,2);
    %apply fluorescence bleed conversion
    Y = applymat(data2,C3);
    %shape back
    Y2 = shiftdim(Y,1);
    %write output
    for k = 1:length(datafields2)
        if ndims(Y2) == 3
            outstruct.(cellnames{ii}).(datafields2{k}) = Y2(:,:,k);
        elseif ismatrix(Y2)
            outstruct.(cellnames{ii}).(datafields2{k}) = Y2(:,k);
        end
    end
end


function Y = applymat(X,C)

Y = zeros(size(X));
for ii = 1:size(X,3)
    %apply color correction to each set of measurements
    Y(:,:,ii) = C\X(:,:,ii);
end
```
