## Supplementary material for "Predicting Composition of Genetic Circuits with Resource Competition: Demand and Sensitivity": Experimental data and Matlab code: RSnetwork.m

```
classdef RSnetwork
    properties
        modulenames
        modules
        Qvec
        yvec
        y0vec
        u
    end
    methods
        function obj = predictoutputs(obj)
            obj.modules
        end
    end
end
```
