## Supplementary material for "Predicting Composition of Genetic Circuits with Resource Competition: Demand and Sensitivity": Experimental data and Matlab code: addtime.m

```
function [platedataout,timeprev] = addtime(platedata,timeprev)

platedataout = platedata;

%find the longest field
numsamples = max(structfun(@(x) size(x,3), platedata));
t = (1:5:5*numsamples)/60;     %time step in hours
%concatenate time data
if timeprev == 0
    timevec = t(:);
else
    timevec = t(:) + timeprev + 5/60;
end
platedataout.time = timevec;
timeprev = timevec(end);
```
