## Supplementary material for "Predicting Composition of Genetic Circuits with Resource Competition: Demand and Sensitivity": Experimental data and Matlab code: aveJ0time.m

```
function [yout,youtstd] = aveJ0time(time,y,ystd,tmin,tmax)
%[yout,youtstd] = aveJ0time(time,y,ystd,tmin,tmax)
%average y over time and calculate uncertianty. time must be a vector and
%tmax and tmin must be scalar values appearing between min(time) and 
%max(time)

if nargin < 5
    tmax = max(time);
    if nargin < 4
        tmin = min(time);
        if nargin < 2
            ystd = zeros(size(y));
        end
    end
end

%find indicies for y between tmin and tmax
[ymax,indmax] = min(abs(time-tmax));
[ymin,indmin] = min(abs(time-tmin));
tstep = time(2) - time(1);
if ymax > tstep || ymin > tstep
    warning('tmax or tmin not values in time vector')
end

%calculate y averaged between tmin and tmax
yout = mean(y(indmin:indmax,:));
if nargout > 1
    youtstd = sqrt(sum(ystd(indmin:indmax,:).^2))/size(ystd(indmin:indmax,:),1);
end
```
