## Supplementary material for "Predicting Composition of Genetic Circuits with Resource Competition: Demand and Sensitivity": Experimental data and Matlab code: aveJs.m

```
function Jstructout = aveJs(Jdatastruct,Jmetadata)

Jnames = fieldnames(Jdatastruct);

if nargin < 2
    Jmetadata.B = {'JB_G','JB_R','JB_l_s'};
    Jmetadata.G = {'JG_B','JG_R','JG_l_s'};
    Jmetadata.R = {'JR_B','JR_G','JR_l_s'};
end

modnames = fieldnames(Jmetadata);
Jstructout = Jdatastruct;

for jj = 1:length(modnames)
    
    Jmatches = Jmetadata.(modnames{jj});
    
    all(cellfun(@(x) any(strcmpi(x,Jnames)),Jmatches))
    
    %check to make sure names are valid
    if all(cellfun(@(x) any(strcmpi(x,Jnames)),Jmetadata.(modnames{jj})))
        
        %init
        [Jtmp,Jstdtmp] = deal(zeros(length(Jmatches),1));
        
        %loop through
        for ii = 1:length(Jmatches)
            
            Jtmp(ii) = Jdatastruct.(Jmatches{ii}).J0;
            Jstdtmp(ii) = Jdatastruct.(Jmatches{ii}).J0std;
        end
        %average J's
        Jstructout.(modnames{jj}).J0 = mean(Jtmp);
        Jstructout.(modnames{jj}).J0std = sqrt(sum(Jstdtmp.^2))./length(Jstdtmp);
        
    end
end
```
