## Supplementary material for "Predicting Composition of Genetic Circuits with Resource Competition: Demand and Sensitivity": Experimental data and Matlab code: avetimeARIMA.m

```
function yout = avetimeARIMA(time,y,ystd,tmin,tmax)


if nargin < 5
    tmax = max(time);
    if nargin < 4
        tmin = min(time);
        if nargin < 2
            ystd = zeros(size(y));
        end
    end
end

%find indicies for y between tmin and tmax
[ymax,indmax] = min(abs(time-tmax));
[ymin,indmin] = min(abs(time-tmin));
tstep = time(2) - time(1);
if ymax > tstep || ymin > tstep
    warning('tmax or tmin not values in time vector')
end

Mdl = arima(1,0,indmax-indmin)
[EstMdl,EstParamCov,logL,info] = estimate(Mdl,y)

%calculate y averaged between tmin and tmax
yout = mean(y(indmin:indmax,:));
```
