## Supplementary material for "Predicting Composition of Genetic Circuits with Resource Competition: Demand and Sensitivity": Experimental data and Matlab code: avetrials.m

```
function [outstruct,figh] = avetrials(cellstruct,datafields,timefld,ploton,figh,cellfields)
%outstruct = avetrials(cellstruct,datafields,timefld,ploton,figh,cellfields)
%average datafields across samples vs point in timefld

if nargin < 3 || isempty(timefld)
    timefld = 'time';
end
if nargin < 4 || isempty(ploton)
    ploton = false;
end
if ploton
    if nargin < 5 || isempty(figh)
        figh = figure;
    else
        if ishandle(figh); clf(figh);
        end
    end
    n = length(datafields);
    m = ceil(9/16*sqrt(n));
end

outstruct = cellstruct;
if nargin < 6 || isempty(cellfields)
    cellnames = fieldnames(cellstruct);
else
    cellnames = cellfields;
end

%get time vector
timedata = cellstruct.(cellnames{1}).(timefld);

for jj = 1:length(datafields)
    
    %init
    [ydataave,ydatastd] = deal(zeros(length(timedata),length(cellnames)));
    
    %loop through cell types
    for ii = 1:length(cellnames)
        
        %check if field matches desired fields
        if all(isfield(cellstruct.(cellnames{ii}),{timefld,datafields{jj}}))
            
            %get data
            ydata = cellstruct.(cellnames{ii}).(datafields{jj});
            %calculate mean at each time point
            ydataave(:,ii) = mean(ydata,2);
            %calculate std
            ydatastd(:,ii) = std(ydata,0,2);
            %output
            outstruct.(cellnames{ii}).([datafields{jj},'ave']) = ydataave(:,ii);
            outstruct.(cellnames{ii}).([datafields{jj},'std']) = ydatastd(:,ii);
        end
    end
    
    if ploton
        %plot average with error bars for intersample variance (std)
        figure(figh);
        subplot(m,ceil(n/m),jj); hold on;
        for k = 1:size(ydataave,2)
            hl(k) = subsampleerrorbar(timedata,ydataave(:,k),ydatastd(:,k),20);
        end
        set(gca,'fontsize',12)
        xlabel(timefld)
        ylabel(datafields{jj})
        legend(hl,cellnames,'location','best','fontsize',10)
        ylim([0,Inf])
    end
end
```
