## Supplementary material for "Predicting Composition of Genetic Circuits with Resource Competition: Demand and Sensitivity": Experimental data and Matlab code: bulkdata2cells.m

```
function cellstruct = bulkdata2cells(datastruct, indstruct, datafields)
%convert raw data in a struct to sorted by individual cells in a struct

if nargin < 3
    datafields = {'OD','GFP','RFP','YFP','time'};
    if nargin < 2
        indstruct = struct;
        indstruct.R = 1:3;
        indstruct.YG = 4:6;
        indstruct.G = 11:13;
        indstruct.GR = 14:16;
        indstruct.Y = 21:23;
        indstruct.YR = 24:26;
        indstruct.M9 = [7,17,27,34:37,41:44];
        indstruct.T10 = 31:33;
        indstruct.none = [8:10,18:20,28:30,38:40,45:60];
    end
end

%init structs
cellstruct = struct;
cellnames = fieldnames(indstruct);

%loop across cell names
for ii = 1:length(cellnames)
    %init individual cell struct
    cellstruct.(cellnames{ii}) = struct;
    
    %loop through datafields
    for jj = 1:length(datafields)
        %disp(datafields{jj})
        if ~contains(datafields{jj},'time')
            numsamples = size(datastruct.(datafields{jj}),3);
            tmp = permute(datastruct.(datafields{jj}),[3,1,2]);   %put time field first
            data = zeros(numsamples,size(indstruct.(cellnames{ii}),1));
            %loop through each sample repeat
            for k = 1:length(indstruct.(cellnames{ii}))
                %add data into array
                data(:,k) = tmp(:,indstruct.(cellnames{ii})(k,1),indstruct.(cellnames{ii})(k,2));
            end
            %copy into struct
            cellstruct.(cellnames{ii}).(datafields{jj}) = data;
        else
            %copy time into struct
            cellstruct.(cellnames{ii}).time = datastruct.(datafields{jj});
        end
    end
end
```
