## Supplementary material for "Predicting Composition of Genetic Circuits with Resource Competition: Demand and Sensitivity": Experimental data and Matlab code: calcFPhat2.m

```
function outstruct = calcFPhat2(instruct,FPfield,trng)


%[xhat, xhatstd] = calcFPhat(xalone,xtogether)

%calculate xhat0 for a pair of constitutive genes
xhat = mean(xtogether,2)./mean(xalone,2);

%estimate error bars assuming uncertianty is dominanted by variance across
%population samples and assuming var(GY), var(G) are uncorrelated
xhatstd = abs(xhat).*sqrt( (std(xtogether,0,2)./mean(xtogether,2)).^2 + ...
    (std(xalone,0,2)./mean(xalone,2)).^2);
```
