## Supplementary material for "Predicting Composition of Genetic Circuits with Resource Competition: Demand and Sensitivity": Experimental data and Matlab code: calcFPhat3.m

```
function [FPhat,FPhatstd] = calcFPhat3(FPalone,FPtogether,FPalonestd,FPtogetherstd)
%calculates ratios of fluorescent concentrations.

%calculate FPhat
FPhat = FPtogether./FPalone;

%estimate error bars assuming uncertianty is dominanted by variance across
%population samples and assuming var(FPtogether), var(FPalone) are uncorrelated
FPhatstd = abs(FPhat).*sqrt( (FPtogetherstd./FPtogether).^2 + ...
    (FPalonestd./FPalone).^2);
```
