## Supplementary material for "Predicting Composition of Genetic Circuits with Resource Competition: Demand and Sensitivity": Experimental data and Matlab code: calcJ0.m

```
function J0 = calcJ03(cellstruct)


[Jr,Jg] = calcJ0growthrate();
J0 = struct('G-R',Jg,'R-G',Jr);
J0 = struct('G-R',Jg,'R-G',Jr);


function [Jr,Jg] = calcJ0growthrate()
muGR = 1;
muG = 1;
muR = 1;

Ghat = 0.9;
Rhat = 0.9;

Jg = (1 - (muGR./muR)*Rhat)./((muGR./muG).*Ghat + (muGR./muR).*Rhat - 1);
Jr = (1 - (muGR./muG)*Ghat)./((muGR./muG).*Ghat + (muGR./muR).*Rhat - 1);

J0 = struct('G',Jg,'R',Jr)
```
