## Supplementary material for "Predicting Composition of Genetic Circuits with Resource Competition: Demand and Sensitivity": Experimental data and Matlab code: calcQS2.m

```
function [modsoutalone,RSmodsout] = calcQS2(modstruct,RSmodules,ploton,...
    fighy,fighyRS,fighQ,fighS,fighyi,fighyRSi,fighQi,fighSi)
%[modsout,RSmodsout] = calcQS2(modstruct,RSmodules,ploton,...
%    fighy,fighyRS,fighQ,fighS,fighyi,fighyRSi,fighQi,fighSi)
%calculate resource demand Q and sensitivity S for arbitrary modules under
%the perturbation from a resource sensor.

if nargin < 3
    ploton = false;
end
if ploton
    if nargin < 4 || isempty(fighy)
        fighy = figure;
    elseif ishandle(fighy); clf(fighy);
    end
    if nargin < 5 || isempty(fighy)
        fighyRS = figure;
    elseif ishandle(fighyRS); clf(fighyRS);
    end
    if nargin < 6 || isempty(fighQ)
        fighQ = figure;
    elseif ishandle(fighQ); clf(fighQ);
    end
    if nargin < 7 || isempty(fighS)
        fighS = figure;
    elseif ishandle(fighS); clf(fighS);
    end
end

%find modules that are perturbed by resource sensors
RSnames = fieldnames(RSmodules);
RSperturb = cell(length(RSnames),1);
for jj = 1:length(RSnames)
    RSperturb{jj} = findpropRS(modstruct,'containingmods',RSnames{jj})';
end

%find groups of non-RS modules that share properties
modsalone = findpropRS(modstruct,'isalone');
modstogether = findpropRS(modstruct,'isalone',false);
normalmods = findpropRS(modstruct,'isResourceSensor',false);  %not resource sensors

%experiments that are modules together and one module isn't a resource sensor
normalmodstogeth = intersect(normalmods,modstogether);
normalmodsalone = intersect(normalmods,modsalone);
normalRSmodstogeth = intersect(normalmodstogeth,unique([RSperturb{:}]'));

%loop through perturbed modules to calculate Q and S for each under the
%perturbation from the Resource Sensor
for ii = 1:length(normalRSmodstogeth)
    
    %need at least one resource sensor and one non resource sensor module
    pairmod = modstruct.(normalRSmodstogeth{ii});
    containedmods = pairmod.containingmods;
    modname = intersect(containedmods,normalmods);
    mod1 = modstruct.(modname{1});
    RSmodname = intersect(containedmods,RSnames);
    RSmod1 = RSmodules.(RSmodname{1});
    
    %get fluorescence channels corresponding to outputs
    mod_FPoutname = mod1.FPout;
    RS_FPoutname = RSmod1.FPout;
    
    %get module and resource sensor outputs
    modalone = abs(mod1.(mod_FPoutname{1}));
    modalonestd = mod1.([mod_FPoutname{1},'std']);
    modtogeth = abs(pairmod.(mod_FPoutname{1}));
    modtogethstd = pairmod.([mod_FPoutname{1},'std']);
    %for resource sensor
    RSalone = abs(RSmod1.(RS_FPoutname{1}));
    RSalonestd = RSmod1.([RS_FPoutname{1},'std']);
    RStogeth = abs(pairmod.(RS_FPoutname{1}));
    RStogethstd = pairmod.([RS_FPoutname{1},'std']);
    
    %get resource sensor Q
    QRS = RSmod1.Q;
    QRSstd = RSmod1.Qstd;
    
    %find Q and S for module from data
    [Q,S,Qstd,Sstd,S2,S2std,S3,S3std,S4,S4std,S9,S9std] = calcQS0(modalone,RSalone,...
        modtogeth,RStogeth,QRS,modalonestd,RSalonestd,modtogethstd,...
        RStogethstd,QRSstd);
    
    %theoretical prediction for constitutive nodes: S in CDC paper
    S5 = -1./(1 + Q + QRS);
    S5std = S5.^2.*sqrt(Qstd.^2 + QRSstd.^2);
    %S2 original S
    S6 = -(1 + Q)./(1 + Q + QRS).^2;
    S6std = sqrt((2*S6./(1+Q+QRS)).^2.*QRSstd.^2 + ((1+Q-QRS)./(1+Q+QRS).^3).^2.*Qstd.^2);
    %S3 = Fhat = 1
    S7 = ones(size(Q));
    S7std = zeros(size(Q));
    S8 = zeros(size(Q));
    S8std = zeros(size(Q));
    
    %output updated modules
    mod1.Q = [mod1.Q, Q];
    mod1.Qstd = [mod1.Qstd, Qstd];
    mod1.S = [mod1.S, S];
    mod1.S2 = [mod1.S2, S2];
    mod1.S3 = [mod1.S3, S3];
    mod1.S4 = [mod1.S4, S4];
    mod1.S5 = [mod1.S5, S5];
    mod1.S6 = [mod1.S6, S6];
    mod1.S7 = [mod1.S7, S7];
    mod1.S8 = [mod1.S8, S8];
    mod1.S9 = [mod1.S9, S9];
    %mod1.S3 = [mod1.S3, S3];
    mod1.Sstd = [mod1.Sstd, Sstd];
    mod1.S2std = [mod1.S2std, S2std];
    mod1.S3std = [mod1.S3std, S3std];
    mod1.S4std = [mod1.S4std, S4std];
    mod1.S5std = [mod1.S5std, S5std];
    mod1.S6std = [mod1.S6std, S6std];
    mod1.S7std = [mod1.S7std, S7std];
    mod1.S8std = [mod1.S8std, S8std];
    mod1.S9std = [mod1.S9std, S9std];
    %mod1.S3std = [mod1.S3std, S3std];
    mod1.y = modalone;
    mod1.ystd = modalonestd;
    mod1.perturbname = [mod1.perturbname, RSmodname(1)];
    mod1.perturby = [mod1.perturby, {modtogeth}];
    mod1.perturbystd = [mod1.perturbystd, {modtogethstd}];
    if ~isempty(mod1.predperturby)
        mod1.predperturby = [mod1.predperturby,{[]}];
        mod1.predperturbystd = [mod1.predperturbystd,{[]}];
    else
        mod1.predperturby = {[]};
        mod1.predperturbystd = {[]};
    end
    modstruct.(modname{1}) = mod1;
    
    %output for resource sensor
    %total output, keeping initial perturbation data from resource sensors
    RSmod1.perturbname = [RSmod1.perturbname, modname(1)];
    RSmod1.perturby = [RSmod1.perturby, {RStogeth}];
    RSmod1.perturbystd = [RSmod1.perturbystd, {RStogethstd}];
    if ~isempty(RSmod1.predperturby)
        RSmod1.predperturby = [RSmod1.predperturby,{[]}];
        RSmod1.predperturbystd = [RSmod1.predperturbystd,{[]}];
    else
        RSmod1.predperturby = {[]};
        RSmod1.predperturbystd = {[]};
    end
    modstruct.(RSmodname{1}) = RSmod1;
    RSmodules.(RSmodname{1}) = RSmod1;
end

%output data only for modules that are not resource sensors
modsoutalone = struct;
for k = 1:length(normalmodsalone)
    modsoutalone.(normalmodsalone{k}) = modstruct.(normalmodsalone{k});
end
%modsout = modstruct;
RSmodsout = RSmodules;

if ploton
    %split into constitutive and induced modules
    [constmods, inducedmods] = splitinducedmods(modsoutalone);
    
    %constitutive modules
    %plot mean and std of RS sensor outputs y and Q
    %note: modules are only perturbed by Resource sensors in this function
    [yvec,ystdvec,RSyvec,RSystdvec,Qvec,Qvecstd,Svec,Svecstd,S2vec,S2vecstd,...
        S3vec,S3vecstd,S4vec,S4vecstd,S5vec,S5vecstd,S6vec,S6vecstd,...
        S7vec,S7vecstd,S8vec,S8vecstd,S9vec,S9vecstd] = ...
        constplots(constmods,RSmodsout);
    
    constmodnames = fieldnames(constmods);
    RSmodnames = fieldnames(RSmodsout);
    inducednames = fieldnames(inducedmods);
    
    %plot module y outputs
    figure(fighy);
    plotRSbars(yvec,ystdvec);
    xticklabels(combinenames(constmodnames,RSmodnames));
    ylabel('$y$','interpreter','latex','fontsize',16)
    title('Module output with resource sensor','fontsize',16)
    
    %plot resource sensor y outputs
    figure(fighyRS);
    plotRSbars(RSyvec,RSystdvec);
    xticklabels(combinenames(RSmodnames,constmodnames));
    ylabel('$y$ Resource Sensor','interpreter','latex','fontsize',16)
    title('Resource sensor output with module','fontsize',16)
    
    %plot Q for module
    figure(fighQ);
    %plotRSbars2(Qvec,Qvecstd);
    plotRSbars(Qvec,Qvecstd,false);
    xticklabels(constmodnames);
    %xticklabels(combinenames(constmodnames,RSmodnames,false));
    ylabel('$Q$','Interpreter','Latex','fontsize',16);
    title('Module Q','fontsize',16)
    
    %plot S for modules
    figure(fighS);
    %subplot(3,1,[1,2]);
    plotRSbars(S4vec,S4vecstd,false);
    %plotRSbars([S4vec,S9vec],[S4vecstd,S9vecstd],true);
    %plotRSbars2([S4vec,S9vec],[S4vecstd,S9vecstd]);
    xticklabels(constmodnames);
    %plotRSbars([S4vec],[S4vecstd],false);
    %xticklabels(combinenames(constmodnames,RSmodnames,false));
    ylabel('$S$','Interpreter','Latex','fontsize',16);
    title('Module S','fontsize',16)
    
    if isempty(inducednames)
        return
    end
    
    %plot Q and S for inducible modules
    if nargin < 8 || isempty(fighyi)
        fighyi = figure;
    elseif ishandle(fighyi); clf(fighyi);
    end
    if nargin < 9 || isempty(fighyRSi)
        fighyRSi = figure;
    elseif ishandle(fighyRSi); clf(fighyRSi);
    end
    if nargin < 10 || isempty(fighQi)
        fighQi = figure;
    elseif ishandle(fighQi); clf(fighQi);
    end
    if nargin < 11 || isempty(fighSi)
        fighSi = figure;
    elseif ishandle(fighSi); clf(fighSi);
    end
    
    %----------
    %induced modules
    %plot curves of module properties vs induction
    %plot RS y
    [uRS,RSyperturb,RSyperturbstd,RSyalone,RSyalonestd] = ...
        deal(cell(length(RSmodnames),1));
    for n = 1:length(RSmodnames)
        RSmod2 = RSmodsout.(RSmodnames{n});
        RSyalone{n} = RSmod2.y;                %scalar
        RSyalonestd{n} = RSmod2.ystd;
        RSperturbname = RSmod2.perturbname;
        %get perturbations from induced modules only
        [inducedRSnames,ia,~] = intersect(RSperturbname,inducednames);
        if isempty(inducedRSnames)
            continue
        end
        [uRS0,RSyperturb0,RSyperturbstd0] = deal(cell(length(ia),1));
        for p = 1:length(ia)
            %look up input u
            uRS0{p} = inducedmods.(inducedRSnames{p}).u;
            %get RS perturbed output as a function of u
            RSyperturb0{p} = RSmod2.perturby{ia(p)};
            RSyperturbstd0{p} = RSmod2.perturbystd{ia(p)};
        end
        uRS{n} = uRS0;
        RSyperturb{n} = RSyperturb0;
        RSyperturbstd{n} = RSyperturbstd0;
    end
    
    %plot
    figure(fighyRSi);
    [lh,lh0] = plotRSycurves(uRS,RSyperturb,RSyperturbstd,RSyalone);
    lh2 = [lh{:}];
    xlabel('Input, u [nM]')
    ylabel('Resource Sensor y, AU')
    legend([lh0,lh2{:}],['alone',RSmodnames(:)'],'Location','best')  
    title('Resource sensor output')
    ylim([0, Inf])
    
    %plot module y
    [u,yvec,ystdvec,Qivec,Qivecstd,Sivec,Sivecstd,Sivec2,Sivec2std,Sivec3,...
        Sivec3std,Sivec4,Sivec4std,Sivec5,Sivec5std,Sivec6,Sivec6std,...
        Sivec7,Sivec7std,Sivec8,Sivec8std] = ...
        deal(cell(length(inducednames),1));
    for m = 1:length(inducednames)
        mod2 = inducedmods.(inducednames{m});
        u{m} = mod2.u;
        yalone = mod2.y;
        yalonestd = mod2.ystd;
        yperturb = mod2.perturby;
        yperturbstd = mod2.perturbystd;
        if ~isempty(yperturb)
        %store normalized output for plotting
            yvec{m} = [yalone, [yperturb{:}]]./max(yalone);
            ystdvec{m} = [yalonestd, [yperturbstd{:}]]./max(yalone);
            %store resource demand for plotting
            Qivec{m} = mod2.Q;   Qivecstd{m} = mod2.Qstd;
            Sivec{m} = mod2.S;   Sivecstd{m} = mod2.Sstd;
            Sivec2{m} = mod2.S2; Sivec2std{m} = mod2.S2std;
            Sivec3{m} = mod2.S3; Sivec3std{m} = mod2.S3std;
            Sivec4{m} = mod2.S4; Sivec4std{m} = mod2.S4std;
            Sivec5{m} = mod2.S5; Sivec5std{m} = mod2.S5std;
            Sivec6{m} = mod2.S6; Sivec6std{m} = mod2.S6std;
            Sivec7{m} = mod2.S7; Sivec7std{m} = mod2.S7std;
            Sivec8{m} = mod2.S8; Sivec8std{m} = mod2.S8std;
        else
            yvec{m} = NaN(length(u{m}),1);
            ystdvec{m} = NaN(length(u{m}),1);
        end
    end
    
    figure(fighyi);
    plotQScurves(u,yvec,ystdvec);
    xlabel('Input, u [nM]'); ylabel('Module y, AU')
    legend(['alone',combinenames(inducednames(:)',RSmodnames(:)',false)],...
        'Location','best')
    
    figure(fighQi);
    plotRSbars2(Qivec{1},Qivecstd{1},true);
    %qlh = plotQScurves(u,Qvec,Qvecstd,true);
    %set(qlh{1},'Color',[0.75,0.75,0.11])
    xticklabels(u{1})
    legend(inducednames,'Location','Best')
    xlabel('Input, u [nM]'); ylabel('Q [ ]')
    
    figure(fighSi);
    %plotRSbars()
    plotRSbars2(Sivec4{1},Sivec4std{1},true);
    %slh = plotQScurves(u,{[Sivec4{:}]},{[Sivec4std{:}]},true);
    %set(slh{1},'Color',[0.75,0.75,0.11])
    xticklabels(u{1})
    legend(inducednames,'Location','Best')
    xlabel('Input, u [nM]'); ylabel('S [ ]')
end


function [Q,S,Qstd,Sstd,S2,S2std,S3,S3std,S4,S4std,S5,S5std] = calcQS0(modalone,RSalone,...
    modtogeth,RStogeth,QRS,modalonestd,RSalonestd,modtogethstd,RStogethstd,QRSstd)
%calculate Q and S according to CDC paper

modalone2 = modalone;
modalone2(modalone < 0.05*max(modalone)) = 0;
modtogeth2 = modtogeth;
modtogeth2(modtogeth < 0.05*max(modtogeth)) = 0;

%module resource demand
Q = ((RSalone./RStogeth) - 1)*(1 + QRS);

%module sensitivity = S in CDC paper;
%S = -1/(1+Q+w0) for constitutive nodes
S = ((modtogeth2 - modalone2)./modtogeth2).*...
    ((RSalone + QRS*(RSalone - RStogeth))./(QRS*RSalone*(QRS + 1)));
%module sensitivity = dy/dw*(1/(y|w=0));
%S2 = -(1+Q)./(1+Q+w0)^2 for constitutive nodes
S2 = ((modtogeth2 - modalone2)./modalone2).*...
    ((RSalone + QRS*(RSalone - RStogeth))./(QRS*RSalone*(QRS + 1)));
%module sensitivity S3 = Fhat from CDC paper
%S3 = 1 for constitutive nodes
S3 = modtogeth.*RSalone*(1 + QRS)./(modalone.*(RSalone + QRS*(RSalone - RStogeth)));

%S4 = (Fhat - 1)/QRS
%S4 = 0 for constituitive nodes
S4 = (S3 - 1)./QRS;
S5 = (1 - 1./S3)./QRS;

%calculate partial derivatives for standard error calculation
dQdRSa = (1+QRS)./RStogeth;                     %dQ/dRSalone
dQdRSt = -(1+QRS)*(RSalone./(RStogeth.^2));     %dQ/dRStogether
dQdQRS = ((RSalone./RStogeth)-1);               %dQ/dQRS

%error when module sensitivity = S in CDC paper
dSdma = -(RSalone + QRS.*(RSalone-RStogeth))./...
    (modtogeth .* RSalone .* QRS .* (1 + QRS));             %dS/dmodalone
dSdmt = modalone .* (RSalone + QRS*(RSalone - RStogeth))./...
    (modtogeth.^2 .* QRS .* RSalone .* (QRS + 1));          %dS/dmodtogether
dSdRSa = ((modtogeth - modalone).*RStogeth)./...
    (modtogeth .* RSalone.^2 * (QRS+1));                    %dS/dRSalone
dSdRSt = (modalone - modtogeth)./...
    (modtogeth .* RSalone .* (QRS + 1));                    %dS/dRStogether
dSdQRS = -((modalone - modtogeth).*(RSalone-RStogeth + RSalone.*(2*QRS+1)))./...
    (modtogeth .* RSalone .* QRS.^2 .* (QRS+1)^2);          %dS/dQRS

%error when module sensitivity = dy/dw*(1/(y|w=0))
dS2dma = -modalone .* (RSalone + QRS*(RSalone - RStogeth))./...
    (modtogeth.^2 .* QRS .* RSalone .* (QRS + 1));          %dS2/dmodtogether
dS2dmt = (RSalone + QRS.*(RSalone-RStogeth))./...
    (modalone .* RSalone .* QRS .* (1 + QRS));              %dS2/dmodtogeth
dS2dRSa = ((modtogeth - modalone).*RStogeth)./...
    (modalone .* RSalone.^2 .* (QRS+1));                    %dS2/dRSalone
dS2dRSt = (modalone - modtogeth)./...
    (modalone .* RSalone .* (QRS + 1));                     %dS2/dRStogether
dS2dQRS = -((modalone - modtogeth).*(RSalone-RStogeth + RSalone.*(2.*QRS+1)))./...
    (modalone .* RSalone .* QRS.^2 .* (QRS+1).^2);          %dS2/dQRS

%error when module sensitivity = Fhat
dS3dma = -modtogeth.*RSalone*(1+QRS)./...
    (modalone.^2 .* (RSalone+QRS.*(RSalone-RStogeth)));    %dS3/dmodalone
dS3dmt = RSalone.*(1+QRS)./...
    (modalone.*(RSalone+QRS.*(RSalone-RStogeth)));         %dS3/dmodtogether
dS3dRSa = -modtogeth.*RStogeth.*QRS.*(1+QRS)./...
    (modalone.*(RSalone+QRS.*(RSalone-RStogeth)).^2);      %dS3/dRSalone
dS3dRSt = modtogeth.*RSalone.*QRS.*(1+QRS)./...
    (modalone.*(RSalone+QRS.*(RSalone-RStogeth)).^2);      %dS3/dRStogether
dS3dQRS = modtogeth.*RSalone.*RStogeth./...
    (modalone.*(RSalone+QRS.*(RSalone-RStogeth)).^2);      %dS3/dQRS

%calculate standard error
Qstd = sqrt(dQdRSa.^2.*RSalonestd.^2 + dQdRSt.^2.*RStogethstd.^2 + ...
    dQdQRS.^2.*QRSstd.^2);
Sstd = sqrt(dSdma.^2.*modalonestd.^2 + dSdmt.^2.*modtogethstd.^2 + ...
    dSdRSa.^2.*RSalonestd.^2 + dSdRSt.^2.*RStogethstd.^2 + dSdQRS.^2.*QRSstd.^2);
S2std = sqrt(dS2dma.^2.*modalonestd.^2 + dS2dmt.^2.*modtogethstd.^2 + ...
    dS2dRSa.^2.*RSalonestd.^2 + dS2dRSt.^2.*RStogethstd.^2 + dS2dQRS.^2.*QRSstd.^2);
S3std = sqrt(dS3dma.^2.*modalonestd.^2 + dS3dmt.^2.*modtogethstd.^2 + ...
    dS3dRSa.^2.*RSalonestd.^2 + dS3dRSt.^2.*RStogethstd.^2 + dS3dQRS.^2.*QRSstd.^2);

S4std = sqrt(S3std.^2./QRS.^2 + (S4./QRS).^2*QRSstd.^2);
S5std = sqrt(S3std.^2.*(1./(S3.*QRS)) + (S5./QRS).^2.*QRSstd.^2);

%set NaN entries to defaults when data is too noisy
emptyinds = isnan(S);
Q(emptyinds) = 0;
S(emptyinds) = 0;
S2(emptyinds) = 0;
S3(emptyinds) = 1;
S4(emptyinds) = 0;
S5(emptyinds) = 0;
Qstd(emptyinds) = 0;
Sstd(emptyinds) = 0;
S2std(emptyinds) = 0;
S3std(emptyinds) = 0;
S4std(emptyinds) = 0;
S5std(emptyinds) = 0;


function [constmods, inducedmods] = splitinducedmods(modstruct)
%split modules into induced and constitutive modules based off the length
%of the input vector u
modnames = fieldnames(modstruct);
inducedmods = struct;
constmods = struct;
for ii = 1:length(modnames)
    if length(modstruct.(modnames{ii}).u) > 1
        inducedmods.(modnames{ii}) = modstruct.(modnames{ii});
    else
        constmods.(modnames{ii}) = modstruct.(modnames{ii});
    end
end


function [y,ystd,RSy,RSystd,Q,Qstd,S,Sstd,S2,S2std,S3,S3std,S4,S4std,S5,S5std,...
    S6,S6std,S7,S7std,S8,S8std,S9,S9std] = constplots(constmods,RSmods)
%constitutive modules
%plot mean and std of RS sensor outputs y and Q
constmodnames = fieldnames(constmods);
RSmodnames = fieldnames(RSmods);
[y,ystd,Q,Qstd,S,Sstd,S2,S2std,S3,S3std,S4,S4std,S5,S5std,S6,S6std,S7,S7std,...
    S8,S8std,S9,S9std] = deal(cell(length(constmodnames),1));
%loop through resource sensors to unpack data for plotting
for m = 1:length(constmodnames)
    mod = constmods.(constmodnames{m});
    yalone = mod.y;
    yalonestd = mod.ystd;
    yperturb = mod.perturby;
    yperturbstd = mod.perturbystd;
    %store normalized output for plotting
    y{m} = [yalone; [yperturb{:}]']/yalone;
    ystd{m} = [yalonestd; [yperturbstd{:}]']/yalone;
    %store resource demand for plotting
    Q{m} = mod.Q;   Qstd{m} = mod.Qstd;
    S{m} = mod.S;   Sstd{m} = mod.Sstd;
    S2{m} = mod.S2; S2std{m} = mod.S2std;
    S3{m} = mod.S3; S3std{m} = mod.S3std;
    S4{m} = mod.S4; S4std{m} = mod.S4std;
    S5{m} = mod.S5; S5std{m} = mod.S5std;
    S6{m} = mod.S6; S6std{m} = mod.S6std;
    S7{m} = mod.S7; S7std{m} = mod.S7std;
    S8{m} = mod.S8; S8std{m} = mod.S8std;
    S9{m} = mod.S9; S9std{m} = mod.S9std;
end

%reshape data for resource sensor outputs when perturbed by modules for plotting
[RSy,RSystd] = deal(cell(length(RSmodnames),1));
for p = 1:length(RSmodnames)
    RSmod = RSmods.(RSmodnames{p});
    RSyalone = RSmod.y;                %scalar
    RSyalonestd = RSmod.ystd;
    RSperturbname = RSmod.perturbname;
    %ignore perturbations from other resource sensors or inducible modules
    [~,ia,~] = intersect(RSperturbname,constmodnames,'stable');
    RSyperturb = [RSmod.perturby{ia}];
    RSyperturbstd = [RSmod.perturbystd{ia}];
    RSy{p} = [RSyalone; RSyperturb(:)]./RSyalone;
    RSystd{p} = [RSyalonestd; RSyperturbstd(:)]./RSyalone;
end


function [bh,ebh] = plotRSbars(Yvec,Yvecstd,aveon)
%plot bars with error bars
if nargin < 3
    aveon = false;
end

if aveon
    Yvec = cellfun(@mean,Yvec,'UniformOutput', false);       %average within each cell
    Yvecstd = cellfun(@(x) sqrt(sum(x.^2)./length(x)),Yvecstd,'UniformOutput', false);
end

xprev = 0;
[bh,ebh] = deal(cell(length(Yvec),1));
for l = 1:length(Yvec)
    xvec = (1:length(Yvec{l})) + xprev;
    xprev = length(xvec) + xprev;
    %make sepperate bar objects for each resoruce sensor module to
    %change colors independently
    bh{l} = bar(xvec,Yvec{l}); hold on
    set(bh{l},'edgecolor','none','barwidth',0.6);
    ebh{l} = errorbar(xvec,Yvec{l},Yvecstd{l},'.k','linewidth',1.5);
    set(gca,'fontsize',12)
end
xticks(1:xprev);
xlim([0.3,xprev+0.8])
box off
set(gca,'linewidth',2);
hold off


function [bh,ebh] = plotRSbars2(Yvec,Yvecstd,aveon)
%plot bars with error bars
if nargin < 3; aveon = false; end

if aveon
    data = mean(Yvec,2);
    datastd = sqrt(sum(Yvecstd.^2,2)./size(Yvecstd,2));
else
    data = Yvec;
    datastd = Yvecstd;
end
%data = cellfun(@(x) mean(x,2),Yvec);       %average within each cell
%datastd = cellfun(@(x) sqrt(sum(x.^2,2)./length(x)),Yvecstd);
%make sepperate bar objects for each resoruce sensor module to
%change colors independently
nbars = size(data,2);
ngroups = size(data,1);
groupwidth = min(0.8, nbars/(nbars + 1.5));
x = zeros(nbars,ngroups);
for ii = 1:nbars
    % Calculate center of each bar
    x(ii,:) = (1:ngroups) - groupwidth/2 + (2*ii-1) * groupwidth / (2*nbars);
end

bh = bar(data); hold on
set(bh,'edgecolor','none','barwidth',0.6);
ebh = errorbar(x',data,datastd,'.k','linewidth',1.5);
set(gca,'fontsize',12)
xticks(1:ngroups);
box off
hold off


function [lh,lh0,ebh] = plotRSycurves(uvec,Yvec,Yvecstd,yalone)

%init
zerooffset = 1e-2;
inds0 = ~cellfun(@isempty,uvec);
[lh,ebh] = deal(cell(length(Yvec),1));
umax = max(cellfun(@max,[uvec{inds0}]));
umin = min(max(cellfun(@min,[uvec{inds0}]),zerooffset));

%plot module alone reference
lh0 = semilogx([umin,umax],ones(2,1),'k--','linewidth',1.2); hold on
%errorbar(umin,1,yalonestd./yalone,'.k','linewidth',1.5);
set(gca,'ColorOrderIndex',1)

%loop through each resource sensor
for jj = 1:length(Yvec)
    if isempty(Yvec{jj})
        continue
    end
    %loop through each perturbation conditions
    for ii = 1:length(Yvec{jj})
        %deal with the zero in u
        u = uvec{jj}{ii};
        if length(u) == 1
            u3 = [umin,umax];
        else
            %replace 0 with a small offset to plot on log scale
            u(u <= 0) = u(u <= 0) + zerooffset;
            u3 = [u(:,1),u(:,2:end)];
        end
        %cast size up if y is scalar
        if length(Yvec{jj}{ii}) == 1 && length(u3) > 1
            %plot horizontal line at constant level
            y = Yvec{jj}{ii}*ones(size(u3));
            ystd = Yvecstd{jj}{ii};
            %plot perturbed y
            lh{jj}{ii} = semilogx(u3,y./yalone{jj},'--','linewidth',1);
            ebh{jj}{ii} = errorbar(umin,y(1)./yalone{jj},ystd./yalone{jj},'.k',...
                'linewidth',1.5);
        else
            y = Yvec{jj}{ii};
            ystd = Yvecstd{jj}{ii};
            %plot perturbed y
            lh{jj}{ii} = semilogx(u3,y./yalone{jj},'linewidth',2);
            coi = get(gca,'ColorOrderIndex');
            ebh{jj}{ii} = errorbar(u3,y./yalone{jj},ystd./yalone{jj},'.k',...
                'linewidth',1.5);
            set(gca,'ColorOrderIndex',coi);
        end
        
        set(gca,'fontsize',14);
        %change zero tick mark
        if any(uvec{jj}{ii} == 0)
            xt = xticks;
            xticklabels([0,xt(2:end)])
        end
    end
end
xlim([umin,umax])
hold off;


function lh = plotQScurves(uvec,yvec,yvecstd,aveon)
if nargin < 4
    aveon = false;
end
if aveon
    yvec = cellfun(@(x)mean(x,2),yvec,'UniformOutput', false);       %average within each cell
    yvecstd = cellfun(@(x) sqrt(sum(x.^2,2)./size(x,2)),yvecstd,'UniformOutput', false);
end

%plot Q or S curves vs input u
zerooffset = 1e-2;
if iscell(uvec)
    lh = cell(length(uvec),1);
    umax = max(cellfun(@max,uvec));
    umin = min(max(cellfun(@min,uvec),zerooffset));
    for ii = 1:length(uvec)
        lh{ii} = semilogx(max(uvec{ii},zerooffset),yvec{ii},'linewidth',2); hold on
        for k = 1:size(yvec{ii},2)
            errorbar(max(uvec{ii},zerooffset),yvec{ii}(:,k),yvecstd{ii}(:,k),...
                '.k','linewidth',1.5);
        end
    end
else
    umax = max(uvec,[],'all');
    umin = max([min(uvec,[],'all'),zerooffset]);
    lh = semilogx(max(uvec,zerooffset),yvec,'linewidth',2); hold on
    for jj = 1:size(yvec,2)
        errorbar(max(uvec,zerooffset),yvec(:,jj),yvecstd(:,jj),'.k','linewidth',1.5);
    end
end
%change zero tick mark
set(gca,'fontsize',14);
if iscell(uvec) && any(cellfun(@(x) any(x == 0),uvec))
    xt = xticks;
    xticklabels([0,xt(2:end)])
elseif any(uvec == 0)
    xt = xticks;
    xticklabels([0,xt(2:end)])
end
xlim([umin,umax])
```
