## Supplementary material for "Predicting Composition of Genetic Circuits with Resource Competition: Demand and Sensitivity": Experimental data and Matlab code: calcQS3.m

```
function [modsoutalone,RSmodsout] = calcQS2(modstruct,RSmodules,ploton,fighy,fighyRS,fighQ,fighS)
%[modsout,RSmodsout] = calcQS(modstruct,RSmodules,ploton,fighy,fighyRS,fighQ,fighS)
%calculate resource demand Q and sensitivity S for arbitrary modules under
%the perturbation from a resource sensor.

%OLD version

if nargin < 3
    ploton = false;
end
if ploton
    if nargin < 4 || isempty(fighy)
        fighy = figure;
    elseif ishandle(fighy); clf(fighy);
    end
    if nargin < 5 || isempty(fighy)
        fighyRS = figure;
    elseif ishandle(fighyRS); clf(fighyRS);
    end
    if nargin < 6 || isempty(fighQ)
        fighQ = figure;
    elseif ishandle(fighQ); clf(fighQ);
    end
    if nargin < 7 || isempty(fighS)
        fighS = figure;
    elseif ishandle(fighS); clf(fighS);
    end
end

%find modules that are perturbed by resource sensors
RSnames = fieldnames(RSmodules);
RSperturb = cell(length(RSnames),1);
for jj = 1:length(RSnames)
    RSperturb{jj} = findpropRS(modstruct,'containingmods',RSnames{jj})';
end

%find groups of non-RS modules that share properties
modsalone = findpropRS(modstruct,'isalone');
modstogether = findpropRS(modstruct,'isalone',false);
normalmods = findpropRS(modstruct,'isResourceSensor',false);  %not resource sensors

%experiments that are modules together and one module isn't a resource sensor
normalmodstogeth = intersect(normalmods,modstogether);
normalmodsalone = intersect(normalmods,modsalone);
normalRSmodstogeth = intersect(normalmodstogeth,unique([RSperturb{:}]'));

%loop through perturbed modules to calculate Q and S for each under the
%perturbation from the Resource Sensor
for ii = 1:length(normalRSmodstogeth)
    
    %need at least one resource sensor and one non resource sensor module
    pairmod = modstruct.(normalRSmodstogeth{ii});
    containedmods = pairmod.containingmods;
    modname = intersect(containedmods,normalmods);
    mod1 = modstruct.(modname{1});
    RSmodname = intersect(containedmods,RSnames);
    RSmod1 = RSmodules.(RSmodname{1});
    
    %get fluorescence channels corresponding to outputs
    mod_FPoutname = mod1.FPout;
    RS_FPoutname = RSmod1.FPout;
    
    %get module and resource sensor outputs
    modalone = mod1.(mod_FPoutname{1});
    modalonestd = mod1.([mod_FPoutname{1},'std']);
    modtogeth = pairmod.(mod_FPoutname{1});
    modtogethstd = pairmod.([mod_FPoutname{1},'std']);
    %for resource sensor
    RSalone = RSmod1.(RS_FPoutname{1});
    RSalonestd = RSmod1.([RS_FPoutname{1},'std']);
    RStogeth = pairmod.(RS_FPoutname{1});
    RStogethstd = pairmod.([RS_FPoutname{1},'std']);
    
    %get resource sensor Q
    QRS = RSmod1.Q;
    QRSstd = RSmod1.Qstd;
    
    %find Q and S for module from data
    [Q,S,Qstd,Sstd,S2,S2std,S3,S3std] = calcQS0(modalone,RSalone,...
        modtogeth,RStogeth,QRS,modalonestd,RSalonestd,modtogethstd,...
        RStogethstd,QRSstd);
    
    %output updated modules
    mod1.Q = [mod1.Q, Q];
    mod1.Qstd = [mod1.Qstd, Qstd];
    mod1.S = [mod1.S, S];
    mod1.S2 = [mod1.S2, S2];
    mod1.S3 = [mod1.S3, S3];
    mod1.Sstd = [mod1.Sstd, Sstd];
    mod1.S2std = [mod1.S2std, S2std];
    mod1.S3std = [mod1.S3std, S3std];
    mod1.y = modalone;
    mod1.ystd = modalonestd;
    mod1.perturbname = [mod1.perturbname, {RSmodname{1}}];
    mod1.perturby = [mod1.perturby, modtogeth];
    mod1.perturbystd = [mod1.perturbystd, modtogethstd];
    modstruct.(modname{1}) = mod1;
    
    %output for resource sensor
    %total output, keeping initial perturbation data from resource sensors
    RSmod1.perturbname = [RSmod1.perturbname, {modname{1}}];
    RSmod1.perturby = [RSmod1.perturby, {RStogeth}];
    RSmod1.perturbystd = [RSmod1.perturbystd, {RStogethstd}];
    modstruct.(RSmodname{1}) = RSmod1;
    RSmodules.(RSmodname{1}) = RSmod1;
end

%output data only for modules that are not resource sensors
modsoutalone = struct;
for k = 1:length(normalmodsalone)
    modsoutalone.(normalmodsalone{k}) = modstruct.(normalmodsalone{k});
end
modsout = modstruct;
RSmodsout = RSmodules;

if ploton
    
    [constmods, inducedmods] = splitinducedmods(modsoutalone);
    
    [yvec,ystdvec,RSyvec,RSystdvec,Qvec,Qvecstd,Svec,Svecstd] = ...
        constplots(constmods,RSmodsout);
    %constitutive modules
    %plot mean and std of RS sensor outputs y and Q
    constmodnames = fieldnames(constmods);
    [yvec,ystdvec,Qvec,Qvecstd,Svec,Svecstd,Svec2,Svec2std,Svec3,Svec3std] = ...
        deal(cell(length(constmodnames),1));
    %loop through resource sensors to reshape data for plotting
    for m = 1:length(constmodnames)
        mod2 = constmods.(constmodnames{m})
        yalone = mod2.y;
        yalonestd = mod2.ystd;
        yperturb = mod2.perturby;
        yperturbstd = mod2.perturbystd;
        %store normalized output for plotting
        yvec{m} = [yalone; yperturb(:)]/yalone;
        ystdvec{m} = [yalonestd; yperturbstd(:)]/yalone;
        %store resource demand for plotting
        Qvec{m} = mod2.Q;
        Qvecstd{m} = mod2.Qstd;
        Svec{m} = mod2.S;
        Svecstd{m} = mod2.Sstd;
        Svec2{m} = mod2.S2;
        Svec2std{m} = mod2.S2std;
        Svec3{m} = mod2.S3;
        Svec3std{m} = mod2.S3std;
    end
    
    %reshape data for resource sensor outputs when perturbed by modules for
    %plotting
    RSmodnames = fieldnames(RSmodsout);
    [RSyvec,RSystdvec] = deal(cell(length(RSmodnames),1));
    for p = 1:length(RSmodnames)
        RSmod2 = RSmodsout.(RSmodnames{p});
        RSyalone = RSmod2.y;                %scalar
        RSyalonestd = RSmod2.ystd;
        RSyperturb = RSmod2.perturby;
        RSyperturbstd = RSmod2.perturbystd;
        RSyvec{p} = unique([RSyalone; [RSyperturb{~contains(...
            RSmod2.perturbname,RSmodnames)}]']./RSyalone,'stable');
        RSystdvec{p} = unique([RSyalonestd; [RSyperturbstd{~contains(...
            RSmod2.perturbname,RSmodnames)}]']./RSyalone,'stable');
    end
    
    %plot module y outputs
    figure(fighy);
    plotRSbars(yvec,ystdvec);
    xticklabels(combinenames(constmodnames,RSmodnames));
    ylabel('$y$','interpreter','latex','fontsize',16)
    title('Module output with resource sensor','fontsize',16)
    
    %plot resource sensor y outputs
    figure(fighyRS);
    plotRSbars(RSyvec,RSystdvec);
    xticklabels(combinenames(RSmodnames,constmodnames));
    ylabel('$y$ Resource Sensor','interpreter','latex','fontsize',16)
    title('Resource sensor output with module','fontsize',16)
    
    %plot Q for module
    figure(fighQ);
    plotRSbars(Qvec,Qvecstd);
    xticklabels(combinenames(constmodnames,RSmodnames,false));
    ylabel('$Q$','Interpreter','Latex','fontsize',16);
    title('Module Q','fontsize',16)
    
    %plot S for modules
    figure(fighS); %subplot(211);
    plotRSbars(Svec,Svecstd);
    xticklabels(combinenames(constmodnames,RSmodnames,false));
    ylabel('$S$','Interpreter','Latex','fontsize',16);
    title('Module S','fontsize',16)
    %subplot(212);
    %plotRSbars(Svec2,Svec2std);
    %xticklabels(combinenames(normmodnames,RSmodnames,false));
    %ylabel('$S_2$','Interpreter','Latex','fontsize',16);
    %title('Module S2','fontsize',16)
    %subplot(313);
    %plotRSbars(Svec3,Svec3std);
    %xticklabels(combinenames(normmodnames,RSmodnames,false));
    %ylabel('$S_3$','Interpreter','Latex','fontsize',16);
    %title('Module S3','fontsize',16)
    
    %----------
    %induced modules
    %plot curves of module properties vs induction
    figure(fighy); clf;
    figure(fighyRS); clf;
    figure(fighQ); clf;
    figure(fighS); clf;
    %plot RS y
    RSmodnames = fieldnames(RSmodsout);
    uRS = cell(length(RSmodnames),1);
    for n = 1:length(RSmodnames)
        RSmod2 = RSmodsout.(RSmodnames{n});
        RSyalone = RSmod2.y;                %scalar
        RSyalonestd = RSmod2.ystd;
        RSyperturb = RSmod2.perturby;
        RSyperturbstd = RSmod2.perturbystd;
        %get input u
        RSperturb = RSmod2.perturbname;
        for k = 1:length(RSperturb)
            if isfield(modsoutalone,RSperturb{k})
                uRS{k} = modsoutalone.(RSperturb{k}).u;
            else; uRS{k} = 0;
            end
        end
        
        %plot
        figure(fighyRS); subplot(2,1,n);
        plotRSycurves(uRS,RSyperturb,RSyperturbstd,RSyalone,RSyalonestd);
        xlabel('Input, u [nM]'); ylabel('Resource Sensor y, AU')
        legend(['alone'],'Location','best')
        title(RSmodnames{n}); ylim([0, Inf])
    end
    
    %plot module y
    constmodnames = fieldnames(modsoutalone);
    [u,yvec,ystdvec,Qvec,Qvecstd,Svec,Svecstd,Svec2,Svec2std,Svec3,Svec3std] = ...
        deal(cell(length(constmodnames),1));
    for m = 1:length(constmodnames)
        mod2 = modsoutalone.(constmodnames{m});
        u{m} = mod2.u;
        yalone = mod2.y;
        yalonestd = mod2.ystd;
        yperturb = mod2.perturby;
        yperturbstd = mod2.perturbystd;
        %store normalized output for plotting
        yvec{m} = [yalone, yperturb]./yalone;
        ystdvec{m} = [yalonestd, yperturbstd]./yalone;
        %store resource demand for plotting
        Qvec{m} = mod2.Q;
        Qvecstd{m} = mod2.Qstd;
        Svec{m} = mod2.S;
        Svecstd{m} = mod2.Sstd;
        Svec2{m} = mod2.S2;
        Svec2std{m} = mod2.S2std;
        Svec3{m} = mod2.S3;
        Svec3std{m} = mod2.S3std;
        
        figure(fighy); subplot(2,2,m);
        plotmodycurves(u{m},yalone,yperturb,yalonestd,yperturbstd)
        xlabel('Input, u [nM]'); ylabel('Module y, AU')
        legend(['alone'],'Location','best')
        title(constmodnames{m}); ylim([0, Inf])
        
        figure(fighQ); subplot(2,2,m);
        plotQScurves(u{m},Qvec{m},Qvecstd{m})
        title(constmodnames{m}); ylim([0,Inf])
        xlabel('Input, u [nM]'); ylabel('Q [ ]')
        
        figure(fighS); subplot(2,2,m);
        plotQScurves(u{m},Svec{m},Svecstd{m})
        title(constmodnames{m})
        xlabel('Input, u [nM]'); ylabel('S [ ]')
        %figure(fighS); %subplot(2,2,m);
        %plotQScurves(u{m},Svec2{m},Svec2std{m})
        %title(normmodnames{m}); %xlabel('Input, u [nM]'); %ylabel('S_2 [ ]')
    end
end


function [Q,S,Qstd,Sstd,S2,S2std,S3,S3std] = calcQS0(modalone,RSalone,...
    modtogeth,RStogeth,QRS,modalonestd,RSalonestd,modtogethstd,RStogethstd,QRSstd)
%calculate Q and S according to CDC paper

%module resource demand
Q = ((RSalone./RStogeth) - 1)*(1 + QRS);

%module sensitivity = S in CDC paper;
%S = -1/(1+Q+w0) for constitutive nodes
S = ((modtogeth - modalone)./modtogeth).*...
    ((RSalone + QRS*(RSalone - RStogeth))./(QRS*RSalone*(QRS + 1)));
%module sensitivity = dy/dw*(1/(y|w=0));
%S2 = -(1+Q)./(1+Q+w0)^2 for constitutive nodes
S2 = ((modtogeth - modalone)./modalone).*...
    ((RSalone + QRS*(RSalone - RStogeth))./(QRS*RSalone*(QRS + 1)));
%module sensitivity = Fhat; from CDC paper
%S3 = 1 for constitutive nodes
S3 = modtogeth.*RSalone*(1 + QRS)./...
    (modalone.*(RStogeth + QRS*(RSalone - RStogeth)));

%calculate partial derivatives for standard error calculation
dQdRSa = (1+QRS)./RStogeth;                 %dQ/dRSalone
dQdRSt = -(1+QRS)*(RSalone./(RStogeth.^2));   %dQ/dRStogether
dQdQRS = ((RSalone./RStogeth)-1);          %dQ/dQRS

%error when module sensitivity = S in CDC paper
dSdma = -(RSalone + QRS.*(RSalone-RStogeth))./...
    (modtogeth .* RSalone .* QRS .* (1 + QRS));             %dS/dmodalone
dSdmt = modalone .* (RSalone + QRS*(RSalone - RStogeth))./...
    (modtogeth.^2 .* QRS .* RSalone .* (QRS + 1));          %dS/dmodtogether
dSdRSa = ((modtogeth - modalone).*RStogeth)./...
    (modtogeth .* RSalone.^2 * (QRS+1));                    %dS/dRSalone
dSdRSt = (modalone - modtogeth)./...
    (modtogeth .* RSalone .* (QRS + 1));                    %dS/dRStogether
dSdQRS = -((modalone - modtogeth).*(RSalone-RStogeth + RSalone.*(2*QRS+1)))./...
    (modtogeth .* RSalone .* QRS.^2 .* (QRS+1)^2);          %dS/dQRS

%error when module sensitivity = dy/dw*(1/(y|w=0))
dS2dma = -modalone .* (RSalone + QRS*(RSalone - RStogeth))./...
    (modtogeth.^2 .* QRS .* RSalone .* (QRS + 1));          %dS/dmodtogether
dS2dmt = (RSalone + QRS.*(RSalone-RStogeth))./...
    (modalone .* RSalone .* QRS .* (1 + QRS));              %dS/dmodtogeth
dS2dRSa = ((modtogeth - modalone).*RStogeth)./...
    (modalone .* RSalone.^2 .* (QRS+1));                    %dS/dRSalone
dS2dRSt = (modalone - modtogeth)./...
    (modalone .* RSalone .* (QRS + 1));                     %dS/dRStogether
dS2dQRS = -((modalone - modtogeth).*(RSalone-RStogeth + RSalone.*(2.*QRS+1)))./...
    (modalone .* RSalone .* QRS.^2 .* (QRS+1).^2);          %dS/dQRS

%error when module sensitivity = Fhat
dS3dma = -modtogeth.*RSalone*(1+QRS)./...
    (modalone.^2 .* (RStogeth+QRS.*(RSalone-RStogeth)));    %dS3/dmodalone
dS3dmt = RSalone.*(1+QRS)./...
    (modalone.*(RStogeth+QRS.*(RSalone-RStogeth)));         %dS3/dmodtogether
dS3dRSa = -modtogeth.*RStogeth.*(1+QRS).*(QRS-1)./...
    (modalone.*(RStogeth+QRS.*(RSalone-RStogeth)).^2);      %dS3/dRSalone
dS3dRSt = modtogeth.*RSalone.*(1+QRS).*(QRS-1)./...
    (modalone.*(RStogeth+QRS.*(RSalone-RStogeth)).^2);      %dS3/dRStogether
dS3dQRS = modtogeth.*RSalone.*(RSalone-2*RStogeth)./...
    (modalone.*(RStogeth+QRS.*(RSalone-RStogeth)).^2);      %dS3/dQRS

%calculate standard error
Qstd = sqrt(dQdRSa.^2.*RSalonestd.^2 + dQdRSt.^2.*RStogethstd.^2 + ...
    dQdQRS.^2.*QRSstd.^2);
Sstd = sqrt(dSdma.^2.*modalonestd.^2 + dSdmt.^2.*modtogethstd.^2 + ...
    dSdRSa.^2.*RSalonestd.^2 + dSdRSt.^2.*RStogethstd.^2 + dSdQRS.^2.*QRSstd.^2);
S2std = sqrt(dS2dma.^2.*modalonestd.^2 + dS2dmt.^2.*modtogethstd.^2 + ...
    dS2dRSa.^2.*RSalonestd.^2 + dS2dRSt.^2.*RStogethstd.^2 + dS2dQRS.^2.*QRSstd.^2);
S3std = sqrt(dS3dma.^2.*modalonestd.^2 + dS3dmt.^2.*modtogethstd.^2 + ...
    dS3dRSa.^2.*RSalonestd.^2 + dS3dRSt.^2.*RStogethstd.^2 + dS3dQRS.^2.*QRSstd.^2);


function [constmods, inducedmods] = splitinducedmods(modstruct)
%split modules into induced and constitutive modules based off the length
%of the input vector u
modnames = fieldnames(modstruct);
for ii = 1:length(modnames)
    if length(modstruct.(modnames{ii}).u) > 1
        inducedmods.(modnames{ii}) = modstruct.(modnames{ii});
    else
        constmods.(modnames{ii}) = modstruct.(modnames{ii});
    end
end


function [yvec,ystdvec,RSyvec,RSystdvec,Qvec,Qvecstd,Svec,Svecstd] = ...
    constplots(constmods,RSmodsout)
%constitutive modules
%plot mean and std of RS sensor outputs y and Q
constmodnames = fieldnames(constmods);
[yvec,ystdvec,Qvec,Qvecstd,Svec,Svecstd,Svec2,Svec2std,Svec3,Svec3std] = ...
    deal(cell(length(constmodnames),1));
%loop through resource sensors to reshape data for plotting
for m = 1:length(constmodnames)
    mod = constmods.(constmodnames{m});
    yalone = mod.y;
    yalonestd = mod.ystd;
    yperturb = mod.perturby;
    yperturbstd = mod.perturbystd;
    %store normalized output for plotting
    yvec{m} = [yalone; yperturb(:)]/yalone;
    ystdvec{m} = [yalonestd; yperturbstd(:)]/yalone;
    %store resource demand for plotting
    Qvec{m} = mod.Q;
    Qvecstd{m} = mod.Qstd;
    Svec{m} = mod.S;
    Svecstd{m} = mod.Sstd;
    Svec2{m} = mod.S2;
    Svec2std{m} = mod.S2std;
    Svec3{m} = mod.S3;
    Svec3std{m} = mod.S3std;
end

%reshape data for resource sensor outputs when perturbed by modules for
%plotting
RSmodnames = fieldnames(RSmodsout);
[RSyvec,RSystdvec] = deal(cell(length(RSmodnames),1));
for p = 1:length(RSmodnames)
    RSmod2 = RSmodsout.(RSmodnames{p});
    RSyalone = RSmod2.y;                %scalar
    RSyalonestd = RSmod2.ystd;
    RSyperturb = RSmod2.perturby;
    RSyperturbstd = RSmod2.perturbystd;
    RSyvec{p} = unique([RSyalone; [RSyperturb{~contains(...
        RSmod2.perturbname,RSmodnames)}]']./RSyalone,'stable');
    RSystdvec{p} = unique([RSyalonestd; [RSyperturbstd{~contains(...
        RSmod2.perturbname,RSmodnames)}]']./RSyalone,'stable');
end


function [bh,ebh] = plotRSbars(Yvec,Yvecstd)
%plot bars with error bars

xprev = 0;
[bh,ebh] = deal(cell(length(Yvec),1));
for l = 1:length(Yvec)
    xvec = (1:length(Yvec{l})) + xprev;
    xprev = length(xvec) + xprev;
    %make sepperate bar objects for each resoruce sensor module to
    %change colors independently
    bh{l} = bar(xvec,Yvec{l}); hold on
    set(bh{l},'edgecolor','none','barwidth',0.6);
    ebh{l} = errorbar(xvec,Yvec{l},Yvecstd{l},'.k','linewidth',1.5);
    set(gca,'fontsize',12)
end
xticks(1:xprev);
xlim([0.3,xprev+0.8])
box off
set(gca,'linewidth',2);
hold off


function [lh,ebh] = plotRSycurves(uvec,Yvec,Yvecstd,yalone,yalonestd)

%init
zerooffset = 1e-4;
[lh,ebh] = deal(cell(length(Yvec),1));
umax = max(cellfun(@max,uvec));
umin = min(max(cellfun(@min,uvec),zerooffset));

%plot module alone reference
semilogx([umin,umax],ones(2,1),'--','linewidth',1.2); hold on
errorbar(umin,1,yalonestd./yalone,'.k','linewidth',1.5);

%loop through each perturbation condition
for l = 1:length(Yvec)
    
    %deal with the zero in u
    u = uvec{l};
    if length(u) == 1
        u3 = [umin,umax];
    else
        u(u <= 0) = u(u <= 0) + zerooffset;
        %u3 = [u(:,1),1.2*u(:,1),u(:,2:end)];
        u3 = [u(:,1),u(:,2:end)];
    end
    
    %cast size up if y is scalar
    if length(Yvec{l}) == 1 && length(u3) > 1
        %plot horizontal line at constant level
        y = Yvec{l}*ones(size(u3));
        ystd = Yvecstd{l};
        %plot perturbed y
        lh{l} = semilogx(u3,y./yalone,'--','linewidth',1);
        ebh{l} = errorbar(umin,y(1)./yalone,ystd./yalone,'.k','linewidth',1.5);
    else
        y = Yvec{l};
        ystd = Yvecstd{l};
        %plot perturbed y
        lh{l} = semilogx(u3,y./yalone,'linewidth',2);
        ebh{l} = errorbar(u3,y./yalone,ystd./yalone,'.k','linewidth',1.5);
    end
    
    set(gca,'fontsize',14);
    %change zero tick mark
    if any(uvec{l} <= 0)
        xt = xticks;
        xticklabels([0,xt(2:end)])
    end
end


function plotmodycurves(uvec,yalone,yperturb,yalonestd,yperturbstd)
%uvec and y should have the same number of rows. all inputs should be
%numeric arrays

zerooffset = 1e-4;
umax = max(uvec,[],'all');
umin = min(uvec,[],'all');
if length(uvec) == 1
    u3 = [umin,umax];
else
    uvec(uvec <= 0) = uvec(uvec <= 0) + zerooffset;
    u3 = [uvec(:,1),uvec(:,2:end)];
end

%normalize by max y value
ymax = max(yalone,[],'all');
%plot module alone
semilogx(u3,yalone./ymax,'linewidth',2); hold on
errorbar(u3,yalone./ymax,yalonestd./ymax,'.k','linewidth',1.5);
%plot module perturbed
semilogx(u3,yperturb./ymax,'linewidth',2);
for jj = 1:size(yperturb,2)
    errorbar(u3,yperturb(:,jj)./ymax,yperturbstd(:,jj)./ymax,'.k','linewidth',1.5);
end

set(gca,'fontsize',14);
%change zero tick mark
if any(uvec == 0)
    xt = xticks;
    xticklabels([0,xt(2:end)])
end


function plotQScurves(uvec,yvec,yvecstd)
%plot Q or S curves vs input u
zerooffset = 1e-4;
semilogx(max(uvec,zerooffset),yvec,'linewidth',2); hold on
for jj = 1:size(yvec,2)
    errorbar(max(uvec,zerooffset),yvec(:,jj),yvecstd(:,jj),'.k','linewidth',1.5);
end

%change zero tick mark
set(gca,'fontsize',14);
if any(uvec == 0)
    xt = xticks;
    xticklabels([0,xt(2:end)])
end
```
