## Supplementary material for "Predicting Composition of Genetic Circuits with Resource Competition: Demand and Sensitivity": Experimental data and Matlab code: calcRS2.m

```
function RSmodules = calcRS2(modulestruct,ploton,fighQ,fighy)
%RSmodules = calcRSall2(modulestruct,ploton,figh)
%calculate resource demand for Resource Sensor. Modulestruct should be a
%struct of all experimental conditions with approporate metadata about
%whether each experiment is for the Resource Sensor or not and whether the
%modules are alone or with other modules. Outputs a struct of resource
%sensor modules alone with appropriate Q and S data filled in.

%NOTE: I eleminated monte-carlo & least squares formulation functionality
%in this version. Use calcRSall.m for that (uses different input/output
%data formatting though).

if nargin < 2
    ploton = false;
end
if ploton
    if nargin < 3 || isempty(fighQ)
        fighQ = figure;
    else
        if ishandle(fighQ); clf(fighQ);
        end
    end
    if nargin < 4 || isempty(fighy)
        fighy = figure;
    else
        if ishandle(fighy); clf(fighy);
        end
    end
end

%calculate nominal resource demand RSD0 with standard error
modsalone = findpropRS(modulestruct,'isalone');
modstogether = findpropRS(modulestruct,'isalone',false);
RSmods = findpropRS(modulestruct,'isResourceSensor');

%experiments that are modules together and all modules are resource sensors
RSmodstogeth = intersect(RSmods,modstogether);
RSmodsalone = intersect(RSmods,modsalone);

%loop through pairs if more than 2 resource sensors present
for ii = 1:length(RSmodstogeth)
    %get metadata from the pair
    modtogeth = modulestruct.(RSmodstogeth{ii});
    alonemods = modtogeth.containingmods;
    FPfields = modtogeth.FPout;
    
    %get output data from modules when measured together
    ytogether1 = modtogeth.(FPfields{1});
    ytogether2 = modtogeth.(FPfields{2});
    ytogether1std = modtogeth.([FPfields{1},'std']);
    ytogether2std = modtogeth.([FPfields{2},'std']);
    
    %get data from cells alone
    alone1 = modulestruct.(alonemods{1});
    alone2 = modulestruct.(alonemods{2});
    yalone1 = alone1.(FPfields{1});
    yalone2 = alone2.(FPfields{2});
    yalone1std = alone1.([FPfields{1},'std']);
    yalone2std = alone2.([FPfields{2},'std']);
    
    %calculate resource demeand Q for each module alone pairwise
    %resource demand for module 1 pairwise
    [Q1,Q1std] = calcRSD0(yalone1,yalone2,ytogether1,ytogether2,...
        yalone1std,yalone2std,ytogether1std,ytogether2std);
    %resource demand for module 2 pairwise
    [Q2,Q2std] = calcRSD0(yalone2,yalone1,ytogether2,ytogether1,...
        yalone2std,yalone1std,ytogether2std,ytogether1std);
    
    %assign resource demand for resoruce sensor modules
    alone1.Q = [alone1.Q, Q1];
    alone1.Qstd = [alone1.Qstd, Q1std];
    alone2.Q = [alone2.Q, Q2];
    alone2.Qstd = [alone2.Qstd, Q2std];
    %assign resource sensitivity to resource sensor modules (known from model)
    alone1.S = -1/(1 + Q1 + Q2);
    alone1.Sstd = sqrt(Q1std^2/(1+Q1+Q2)^2 + Q2std^2/(1+Q1+Q2)^2);
    alone2.S = -1/(1 + Q1 + Q2);
    alone2.Sstd = sqrt(Q1std^2/(1+Q1+Q2)^2 + Q2std^2/(1+Q1+Q2)^2);
    alone1.S2 = -(1 + Q1)/(1 + Q1 + Q2).^2;
    alone1.S2std = sqrt(((1+Q1-Q2)/(1+Q1+Q2)^3)*Q1std^2 + (2*(1+Q1)/(1+Q1+Q2)^3)*Q2std^2);
    alone2.S2 = -(1 + Q2)/(1 + Q1 + Q2).^2;
    alone2.S2std = sqrt(((1+Q2-Q1)/(1+Q1+Q2)^3)*Q2std^2 + (2*(1+Q2)/(1+Q1+Q2)^3)*Q1std^2);
    alone1.S3 = 1;
    alone1.S3std = 0;
    alone2.S3 = 1;
    alone2.S3std = 0;
    alone1.S4 = 0;
    alone1.S4std = 0;
    alone2.S4 = 0;
    alone2.S4std = 0;
    alone1.S5 = alone1.S;
    alone1.S5std = alone1.Sstd;
    alone2.S5 = alone2.S;
    alone2.S5std = alone2.Sstd;
    alone1.S6 = alone1.S2;
    alone1.S6std = alone1.S2std;
    alone2.S6 = alone2.S2;
    alone2.S6std = alone2.S2std;
    alone1.S7 = 1;
    alone1.S7std = 0;
    alone2.S7 = 1;
    alone2.S7std = 0;
    alone1.S8 = 0;
    alone1.S8std = 0;
    alone2.S8 = 0;
    alone2.S8std = 0;
    alone1.S9 = 0;
    alone1.S9std = 0;
    alone2.S9 = 0;
    alone2.S9std = 0;
    %document which module perturbed the resource sensor and corresponding outputs
    alone1.y = yalone1;
    alone2.y = yalone2;
    alone1.ystd = yalone1std;
    alone2.ystd = yalone2std;
    alone1.perturbname = [alone1.perturbname, alonemods(2)];
    alone2.perturbname = [alone2.perturbname, alonemods(1)];
    alone1.perturby = [alone1.perturby, {ytogether1}];
    alone2.perturby = [alone2.perturby, {ytogether2}];
    alone1.perturbystd = [alone1.perturbystd, {ytogether1std}];
    alone2.perturbystd = [alone2.perturbystd, {ytogether2std}];
    
    %fill RS predictions with empty since don't need to predict pair for 2
    %resource sensors
    if ~isempty(alone1.predperturby)
        alone1.predperturby = [alone1.predperturby,{[]}];
        alone1.predperturbystd = [alone1.predperturbystd,{[]}];
    else
        alone1.predperturby = {[]};
        alone1.predperturbystd = {[]};
    end
    if ~isempty(alone2.predperturby)
        alone2.predperturby = [alone2.predperturby,{[]}];
        alone2.predperturbystd = [alone2.predperturbystd,{[]}];
    else
        alone2.predperturby = {[]};
        alone2.predperturbystd = {[]};
    end
    
    %save output
    modulestruct.(alonemods{1}) = alone1;
    modulestruct.(alonemods{2}) = alone2;
end

%output data for Resource Sensor modules alone
RSmodules = struct;
for jj = 1:length(RSmodsalone)
    RSmodules.(RSmodsalone{jj}) = modulestruct.(RSmodsalone{jj});
end


if ploton
    %plot mean and std of RS sensor outputs y and Q
    RSnames = fieldnames(RSmodules);
    [yvec,ystdvec,bh1,bh2,ebh2] = deal(cell(length(RSnames),1));
    [Qvec,Qvecstd] = deal(zeros(length(RSnames),1));
    %loop through resource sensors to reshape data for plotting
    for k = 1:length(RSnames)
        RSmod = RSmodules.(RSnames{k});
        yalone = RSmod.y;
        yperturb = RSmod.perturby;
        yalonestd = RSmod.ystd;
        yperturbstd = RSmod.perturbystd;
        %store normalized output for plotting
        yvec{k} = [yalone; [yperturb{:}]']./yalone;
        ystdvec{k} = [yalonestd; [yperturbstd{:}]']./yalone;
        %store resource demand for plotting
        Qvec(k) = RSmod.Q;
        Qvecstd(k) = RSmod.Qstd;
    end
    
    %plot module outputs
    figure(fighy);
    xprev = 0;
    for m = 1:length(RSnames)
        %get x coordinates to plot bars
        xvec2 = (1:length(yvec{m})) + xprev;
        xprev = length(yvec{m}) + xprev;
        %plot bars
        bh2{m} = bar(xvec2,yvec{m}); hold on
        set(bh2{m},'edgecolor','none','barwidth',0.6);
        ebh2{m} = errorbar(xvec2,yvec{m},ystdvec{m},'.k','linewidth',1.5);
        set(gca,'fontsize',14);
    end
    ylabel('$y$','interpreter','latex')
    xticks(1:xprev);
    xticklabels(combinenames(RSmodsalone,RSmodsalone))
    box off
    set(gca,'linewidth',2)
    xlim([0.3,xprev+0.8])
    title('RS module outputs alone v together')
    
    %plot Q for Resource Sensor
    figure(fighQ);
    xvec = 1:length(RSnames);
    for l = 1:length(RSnames)
        %make sepperate bar objects for each resoruce sensor module to
        %change colors independently
        bh1{l} = bar(xvec(l),Qvec(l)); hold on
        set(bh1{l},'edgecolor','none','barwidth',0.6);
    end
    errorbar(xvec,Qvec,Qvecstd,'.k','linewidth',1.5);
    xticks(1:length(RSnames));
    xticklabels(RSnames);
    ylabel('$Q$','Interpreter','Latex');
    title('Resource Sensor Q')
    set(gca,'fontsize',16)
    box off
    set(gca,'linewidth',2)
    xlim([0.3,length(xvec)+0.8])
end


function [Q,Qstd] = calcRSD0(y1alone,y2alone,y1togeth,y2togeth,...
    y1alonestd,y2alonestd,y1togethstd,y2togethstd)
%calculates Resource Demand of resource sensor and standard error of the
%first gene y1alone.

%compute errors comparing alone and perturbed conditions
e2 = y2alone - y2togeth;
%calculate resource demand for a pair of constitutive genes
num = y1alone.*e2;
den = y1alone.*y2togeth + y1togeth.*y2alone - y1alone.*y2alone;
Q = num./den;

%calculate standard error propagation for resource sensor demand
dfdF1alone = y2alone.*y1togeth.*e2./(den.^2);
dfdF1togeth = -y2alone.*y1alone.*e2./(den.^2);
dfdF2alone = y1alone.*y2togeth.*y1togeth./(den.^2);
dfdF2togeth = -y1alone.*y2alone.*y1togeth./(den.^2);
Qstd = sqrt(dfdF1alone.^2.*y1alonestd.^2 + dfdF1togeth.^2.*y1togethstd.^2 + ...
    dfdF2alone.^2.*y2alonestd.^2 + dfdF2togeth.^2.*y2togethstd.^2);
```
