## Supplementary material for "Predicting Composition of Genetic Circuits with Resource Competition: Demand and Sensitivity": Experimental data and Matlab code: calcRSall.m

```
function RSDstruct = calcRSall(cellstruct,FPstruct,nummods,nummctrials,ploton,figh)
%RSDstruct = calcRSall(cellstruct,FPstruct,nummods,ploton,figh)
%calculate resource demand by standard error and monte carlo simulation

if nargin < 4
    nummctrials = 10000;
end
if nargin < 5
    ploton = true;
end
if ploton
    if nargin < 6 || isempty(figh)
        figh = figure;
    else
        if ishandle(figh); clf(figh);
        end
    end
end

%calculate nominal resource demand RSD0 with standard error
RSDstruct = findRSD0(cellstruct,FPstruct,nummods);
%run Monte-Carlo simulation calculating resource demand RSD0
mcfun = @(x) RSDmcfun(cellstruct,FPstruct,nummods);
mcout = rsD0mc(mcfun,0,0,nummctrials);
%calculate stats from MC simulation
mcmean = mean(mcout,2);
mcstd = std(mcout,0,2);

%monte-carlo output for pairwise
mcpairs = reshape(mcmean(1:end-nummods),[],2);
mcpairsstd = reshape(mcstd(1:end-nummods),[],2);

FPnames = fieldnames(FPstruct);
cellaloneinds = cell(nummods,1);
for k = 1:length(FPnames)
    %get 
    cellalone1 = FPstruct.(FPnames{k}).alone1;
    cellalone2 = FPstruct.(FPnames{k}).alone2;
    %output back to RSDstruct
    RSDstruct.(cellalone1).([FPnames{k},'']) = mcpairs(k,1);
    RSDstruct.(cellalone2).([FPnames{k},'']) = mcpairs(k,2);
    RSDstruct.(cellalone1).([FPnames{k},'std']) = mcpairsstd(k,1);
    RSDstruct.(cellalone1).([FPnames{k},'std']) = mcpairsstd(k,2);
    %get map from indicies to cell alone names
    RSDinds = FPstruct.(FPnames{k}).RSDinds;
    cellaloneinds{RSDinds(1)} = cellalone1;
    cellaloneinds{RSDinds(2)} = cellalone2;
end

%monte-carlo output for least squares
mcls = mcmean(end-(nummods-1):end);
mclsstd = mcstd(end-(nummods-1):end);
for ii = 1:length(cellaloneinds)
    RSDstruct.(cellaloneinds{ii}).ls = mcls(ii);
    RSDstruct.(cellaloneinds{ii}).lsstd = mclsstd(ii);
end

if ploton
    %plot mean and std of FP hats and RSD0's
    figure(figh); clf;
    RSDnames = fieldnames(RSDstruct);
    for jj = 1:length(RSDnames)
        subplot(length(RSDnames),1,jj);
        bh = barsubfield(orderfields(RSDstruct.(RSDnames{jj})),'','std');
        ylabel(['w_',RSDnames{jj},' demand'])
        set(bh,'facecolor',[0.82,0.83,0.97]);
        set(bh,'barwidth',0.6);
    end
end


function [RSDstructout,RSDvec2] = findRSD0(instruct,FPstruct,nummods,MCsim)
%RSDstructout = findRSD04(instruct,FPstruct,nummods,MCsim)
%calculate RSD's for experiments by specifying how to combine data into
%fluorescence hat with FPHATSTRUCT. Fieldnames of FPHATSTRUCT should
%correspond to names of fluroescence hats and should contain the subfields:
%'alone', 'perturb', 'FPfield', and 'growthfield' which each hold a string
%corresponding to fields of INSTRUCT for 'alone' and 'perturb', and
%subfields for 'FPfield' and 'growthfield'. RSDSTRUCT specifies how to
%combine fluorescence hat data to obtain RSD0s. RSDSTRUCT should have the
%subfields 'hat1' and 'hat2', each holding a string specifiying a field in
%FPHATSTRUCT used to calculate RSD0 in each case.
%MCSIM is a bool whether to run the monte-carlo simulation

if nargin < 2
    %specify how to combine data to get resource demands for each experiment
    %blue & green
    FPstruct.BG.alone1 = 'B';    FPstruct.BG.FPfield1 = 'BFPdiffOD_ss';
    FPstruct.BG.alone2 = 'G';    FPstruct.BG.FPfield2 = 'GFPdiffOD_ss';
    FPstruct.BG.perturb = 'BG';
    FPstruct.BG.RSDinds = [1,2];             %for least squares
    %blue & red
    FPstruct.BR.alone1 = 'B';    FPstruct.BR.FPfield1 = 'BFPdiffOD_ss';
    FPstruct.BR.alone2 = 'G';    FPstruct.BR.FPfield2 = 'GFPdiffOD_ss';
    FPstruct.BR.perturb = 'BR';
    FPstruct.BR.RSDinds = [1,3];
    %green & red
    FPstruct.GR.alone1 = 'G';    FPstruct.GR.FPfield1 = 'GFPdiffOD_ss';
    FPstruct.GR.alone2 = 'R';    FPstruct.GR.FPfield2 = 'RFPdiffOD_ss';
    FPstruct.GR.perturb = 'GR';
    FPstruct.GR.RSDinds = [2,3];             %index of resource demand coefficient
end
if nargin < 3
    nummods = 3;
end
if nargin < 4 || isempty(MCsim)
    MCsim = false;
end

%init
RSDstructout = struct;
FPnames = fieldnames(FPstruct);
Y = zeros(2*length(FPnames),1);
X = zeros(2*length(FPnames),nummods);
covY = zeros(length(FPnames));
allRSDinds = zeros(2,length(FPnames));
allcellalone = cell(2,length(FPnames));
RSDvec = zeros(length(FPnames),2);      %MC simulation

%loop through fields in FP struct to calculate module resource demands
for ii = 1:length(FPnames)
    
    %get metadata
    cellalone1 = FPstruct.(FPnames{ii}).alone1;
    cellalone2 = FPstruct.(FPnames{ii}).alone2;
    FPfield1 = FPstruct.(FPnames{ii}).FPfield1;
    FPfield2 = FPstruct.(FPnames{ii}).FPfield2;
    cellperturb = FPstruct.(FPnames{ii}).perturb;
    RSDinds = FPstruct.(FPnames{ii}).RSDinds;
    
    %unpack steady state data with standard errors
    FP1alone = mean(instruct.(cellalone1).(FPfield1));
    FP1alonestd = instruct.(cellalone1).([FPfield1,'_std']);
    FP1togeth = mean(instruct.(cellperturb).(FPfield1));
    FP1togethstd = instruct.(cellperturb).([FPfield1,'_std']);
    FP2alone = mean(instruct.(cellalone2).(FPfield2));
    FP2alonestd = instruct.(cellalone2).([FPfield2,'_std']);
    FP2togeth = mean(instruct.(cellperturb).(FPfield2));
    FP2togethstd = instruct.(cellperturb).([FPfield2,'_std']);
    
    %add noise according to normal distribution with standard error std for
    %monte carlo simulation
    if MCsim == true
        FP1alone = addnoise(FP1alone,FP1alonestd);
        FP2alone = addnoise(FP2alone,FP2alonestd);
        FP1togeth = addnoise(FP1togeth,FP1togethstd);
        FP2togeth = addnoise(FP2togeth,FP2togethstd);
    end
    
    %calculate resource demand for module1 pairwise
    [RSD01,RSD01std] = calcRSD0(FP1alone,FP2alone,FP1togeth,FP2togeth,...
        FP1alonestd,FP2alonestd,FP1togethstd,FP2togethstd);
    %calculate resource demand for module2 pairwise
    [RSD02,RSD02std] = calcRSD0(FP2alone,FP1alone,FP2togeth,FP1togeth,...
        FP2alonestd,FP1alonestd,FP2togethstd,FP1togethstd);
    
    %output
    RSDvec(ii,:) = [RSD01,RSD02];
    RSDstructout.(cellalone1).(FPnames{ii}) = RSD01;
    RSDstructout.(cellalone2).(FPnames{ii}) = RSD02;
    RSDstructout.(cellalone1).([FPnames{ii},'std']) = RSD01std;
    RSDstructout.(cellalone2).([FPnames{ii},'std']) = RSD02std;
    
    %create data matricies for least squares regression
    Y(2*ii-1) = FP1alone - FP1togeth;
    X(2*ii-1,RSDinds(1)) = FP1alone - FP1togeth;
    X(2*ii-1,RSDinds(2)) = FP1togeth;
    Y(2*ii) = FP2alone - FP2togeth;
    X(2*ii,RSDinds(2)) = FP2alone - FP2togeth;
    X(2*ii,RSDinds(1)) = FP2togeth;
    
    %covariance of Y
    covY(2*ii-1,2*ii-1) = FP1alonestd.^2 + FP1togethstd.^2;
    covY(2*ii,2*ii) = FP2alonestd.^2 + FP2togethstd.^2;
    
    allRSDinds(:,ii) = RSDinds(:);
    allcellalone(:,ii) = [{cellalone1};{cellalone2}];
end

%want weights to be reciprocal of variance if measurements uncorrelated,
%otherwise W = inv(cov(y))
W = inv(covY);
%model: Y = X*RSDest;
RSDlsq = (X'*W*X)\X'*W*Y;
RSDvec2 = [RSDvec(:); RSDlsq];

%RSDlsq = (X'*X)\X'*Y; %this is biased due to correlation between X columns
%output least squares estimate
cellsalone = unique(allcellalone,'stable');
for k = 1:length(cellsalone)
    RSDstructout.(cellsalone{k}).ls = RSDlsq(k);
    RSDstructout.(cellsalone{k}).lsstd = 0; %calculate standard error from MC simulation
end


function [RSD0,RSD0std] = calcRSD0(FP1alone,FP2alone,FP1togeth,FP2togeth,...
    FP1astd,FP2astd,FP1tstd,FP2tstd)
%calculates Resource Demand  of constitutive node
%(resources used by the perturbation) assuming xhat2 is the
%perturbation and xhat1 is the basal protein. Also calculates the standard
%deviation of RSD0 if standard deviations of xhat1 and xhat2 are provided

%compute errors comparing alone and perturbed conditions
%e1 = FP1alone - FP1togeth;
e2 = FP2alone - FP2togeth;
%calculate resource demand for a pair of constitutive genes
num = FP1alone*e2;
den = FP1alone*FP2togeth + FP1togeth*FP2alone - FP1alone*FP2alone;
RSD0 = num./den;

%calculate standard error propagation for resource demand
dfdF1alone = FP2alone*FP1togeth*e2./(den.^2);
dfdF1togeth = -FP2alone*FP1alone*e2./(den.^2);
dfdF2alone = FP1alone*FP2togeth*FP1togeth./(den.^2);
dfdF2togeth = -FP1alone*FP2alone*FP1togeth./(den.^2);
RSD0std = sqrt(dfdF1alone^2*FP1astd^2 + dfdF1togeth^2*FP1tstd^2 + ...
    dfdF2alone^2*FP2astd^2 + dfdF2togeth^2*FP2tstd^2);


function y = addnoise(xmean,xstd)
%add normally distributed noise for MC simulation
y = xmean + randn(1)*xstd;


function out = rsD0mc(funh,nominputs,inputstd,numsamples)
%monte carlo for resoruce sensor

if nargin < 4
    numsamples = 100;
end

m = length(nominputs);              %size of inputs
outsize = size(funh(nominputs));     %size of outputs
%generate random input data according to a normal distribution with nominal
%values and prescribed std
data = nominputs(:) + inputstd(:).*randn([numsamples,m]);
%apply funh to data for each random sample
out = zeros(max(outsize),numsamples);
disp(['running monte-carlo: ',num2str(numsamples),' trials'])
for ii = 1:numsamples
    %evalulate funh at each point
    tmp = funh(data(ii,:));
    out(:,ii) = tmp(:);
end


function RSDvec = RSDmcfun(cellstruct,FPstruct,nummods)
%get 2nd output for running montecarlo
[~,RSDvec] = findRSD0(cellstruct,FPstruct,nummods,true);
```
