## Supplementary material for "Predicting Composition of Genetic Circuits with Resource Competition: Demand and Sensitivity": Experimental data and Matlab code: correctFP.m

```
function [outstruct,cvtstruct] = correctFP(instruct,cellcontrolfields,FPfields,ploton)
%[outstruct, cvtstruct] = correctFP(instruct, cellcontrolfields, FPfields)
%find fluorescence correction matrix 'C2' based on fluorescence controls 
%listed in the cell array CELLCONTROLFIELDS and apply to all fields listed 
%in the cell array FPFIELDS
%
%[outstruct, cvtstruct] = correctFP(instruct, correctionmatfilename, FPfields)
%if only two inputs are supplied, the second input is the path for a matlab 
%savefile containing the correction matrix 'C2'

if nargin < 3
    error('not enough inputs')
end
if nargin < 4
    ploton = false;
end

if iscellstr(cellcontrolfields)
    %find fluorescence correction based on fluorescence control for the
    %fields in the cell array FPfields
    
    %sensor limits for linear range
    GFPmax = 1e4;
    RFPmax = 5000;
    YFPmax = inf;
    BFPmax = inf;
    
    %error checking
    if ~all(isfield(instruct,cellcontrolfields))
        error('cellcontrolfields do not match with fields in cellstruct')
    end
    if length(FPfields) > 9
        error('too many FPfields')
    end
    if length(cellcontrolfields) ~= length(FPfields)
        error('number of cell control fields to not match the number of fluorescence channels')
    end
    
    %make sure order matches
    FPfields = sort(FPfields);
    cellcontrolfields = sort(cellcontrolfields);
    
    %init
    C = zeros(length(cellcontrolfields),length(FPfields));
    cvtstruct = struct;
    outstruct = instruct;
    
    %compensate for spectrial leakage in GFP/RFP/YFP signal
    disp('correcting for fluorescence leakage...')
    
    %loop through each control cell
    for ii = 1:length(cellcontrolfields)
        if ~all(isfield(instruct.(cellcontrolfields{ii}),FPfields))
            error('FPfields do not match with property fields in cellstruct')
        end
        disp(['finding effect of ',cellcontrolfields{ii},' on fluorescence'])
        
        %find all combinations of two FP channels
        %FPfieldpairs = nchoosek(FPfields,2);
        %loop through all pairs of FP channels
        for jj = 1:length(FPfields)
            
            %skip diagonal element
            if jj == ii
                continue
            end
            
            %get correct limits on fluorescence channels where linear range of
            %sensor is valid
            switch FPfields{ii}
                case 'RFP1_584_607'; xmax = RFPmax;
                case 'RFP'; xmax = RFPmax;
                case 'GFP1_465_498'; xmax = GFPmax;
                case 'GFP'; xmax = GFPmax;
                case 'GFP1_518_540'; xmax = YFPmax;
                case 'YFP'; xmax = YFPmax;
                case 'BFP'; xmax = BFPmax;
            end
            switch FPfields{jj}
                case 'RFP1_584_607'; ymax = RFPmax;
                case 'RFP'; ymax = RFPmax;
                case 'GFP1_465_498'; ymax = GFPmax;
                case 'GFP'; ymax = GFPmax;
                case 'GFP1_518_540'; ymax = YFPmax;
                case 'YFP'; ymax = YFPmax;
                case 'BFP'; ymax = BFPmax;
            end
            
            %fit line between all pairs of FPfields and take slope
            x = instruct.(cellcontrolfields{ii}).(FPfields{ii});
            y = instruct.(cellcontrolfields{ii}).(FPfields{jj});
            inds = (x < xmax) & (y < ymax);
            [p1,p2] = deal(zeros(size(inds,2),2));
            for k = 1:size(inds,2)
                %run fit for each column
                %dependence of GFP and RFP channels
                p1(k,:) = polyfit(x(inds(:,k),k),y(inds(:,k),k),1);
                %dependence of GFP and RFP channels
                p2(k,:) = polyfit(y(inds(:,k),k),x(inds(:,k),k),1);
            end
            %get average slopes over all trials
            m1 = mean(p1(:,1),1);
            m2 = mean(p2(:,1),1);
            %take smaller absolute value of slopes p1(1) and p(2) since
            %bleedover should never be larger than 1 (hopefully)
            foo = [m1,m2];
            [~,ind] = min(abs(foo));
            
            if ploton
                figure;
                fity = foo(ind)*x(:,1);
                for l = 1:size(inds,2)
                    plot(x(inds(:,l),l),y(inds(:,l),l),'linewidth',1.5); hold on
                end
                plot(x(:,1),fity,'k--','linewidth',1.5); drawnow;
                xlabel(strrep(FPfields{ii},'_','-'))
                ylabel(strrep(FPfields{jj},'_','-'))
                title(['slope: ',num2str(foo(ind))])
            end
            
            cvtstruct.(cellcontrolfields{ii}).(FPfields{jj}) = foo(ind);
            %cvtstruct.(cellcontrolfields{ii}).(FPfields{jj}) = foo(ind);
            C(jj,ii) = foo(ind);%max([0,foo(ind)]);
        end
    end
    
    %correction matrix
    C2 = C + eye(size(C));
    
    %save output
    info = struct('GFPexcitation','465nm pm 5','RFPexcitation','584nm pm 5',...
        'GFPemmission','498nm pm 20','RFPemmission','607nm pm 20');
    save('GFP-RFPcorrection.mat','C2','info');
else
    
    %load corretion matrix
    if ischar(cellcontrolfields) || isstring(cellcontrolfields)
        load(cellcontrolfields,'C2');
    else
        load('GFP_RFPcorrection.mat','C2');
    end
end

disp('Correction matrix:')
disp(FPfields')
disp(C2)
%apply correction matrix to fields

%C2 = [1,0,0; 0,1,0.2; 0,0,1]

cellnames = fieldnames(instruct);
%loop through all cells to correct fluorescence measurements
for k = 1:length(cellnames)
    
    %stack fields into vector for multiplying with correction matrix
    temp = [];
    for l = 1:length(FPfields)
        temp = [temp;instruct.(cellnames{k}).(FPfields{l})(:)'];
        fieldsize = size(instruct.(cellnames{k}).(FPfields{l}));
    end
    
    %apply fluorescence correction matrix
    outtemp = C2\temp;
    
    %unpack fields
    for m = 1:length(FPfields)
        outstruct.(cellnames{k}).(FPfields{m}) = reshape(outtemp(m,:),fieldsize);
    end
end
```
