## Supplementary material for "Predicting Composition of Genetic Circuits with Resource Competition: Demand and Sensitivity": Experimental data and Matlab code: diff5pt.m

```
function dxdt = diff5pt(f,h)
%run 5 pt stencil differentiation on x with respect to t
%f is vector of inputs, h is step size. h can be vector of evenly spaced
%points where f is sampled.

if length(h) > 1
    h2 = abs(mean(h(2:end) - h(1:end-1)));
else; h2 = h;
end

f2 = [zeros(2,1);f(:)];      %append 0's to beginning and end of x

dxdt = (f2(1:end-3) - 8*f2(2:end-2) + 8*f2(3:end-1) - f2(4:end))/(12*h2);

dxdt = [dxdt;0];    %pad with zero
```
