## Supplementary material for "Predicting Composition of Genetic Circuits with Resource Competition: Demand and Sensitivity": Experimental data and Matlab code: diffcellstruct.m

```
function outstruct = diffcellstruct(cellstruct,difffields,timefield)
%outstruct = diffcellstruct(cellstruct,difffields,timefield)
%apply smooth differentation to difffields in cellstruct with respect to
%timefield. difffields may be a cell array containing the fields to be
%differentaited. timefield must be a string.

cellnames = fieldnames(cellstruct);
outstruct = cellstruct;

%loop through cells
for ii = 1:length(cellnames)
    %loop through fields to be differentiated
    for jj = 1:length(difffields)
        %get fields
        FP = cellstruct.(cellnames{ii}).(difffields{jj});
        time = cellstruct.(cellnames{ii}).(timefield);
        %take numerical derivative with respect to time
        dFPdt0 = diff5pt(FP,time);
        %lowpass filter
        dFPdt = moveavefilt(dFPdt0,7);
        %dFPdt = lowpassfilt(dFPdt0,0.5);
        %output
        outstruct.(cellnames{ii}).([difffields{jj},'diff']) = dFPdt;
    end
end


function dfdh = diff5pt(f,h)
%run 5 pt stencil differentiation on x with respect to t
%f is vector of inputs, h is step size. h can be vector of evenly spaced
%points where f is sampled.

%find step size
if length(h) > 1
    h2 = abs(mean(h(2:end) - h(1:end-1)));
else; h2 = h;
end

% 9-pt Savitzky-Golay filtered first derivative (1st/2nd order polynomial fit)
a = 60*h2;
b = [-4,-3,-2,-1,0,1,2,3,4];
dfdh = filter(-b,a,f);


function y = moveavefilt(x,n)
%moving average filter with n points
a = 1;
b = 1/n*ones(1,n);
if nnz(isnan(x)) > 0
    y = x;
else
    y = filtfilt(b,a,x);
end


function y = lowpassfilt(x,wn)
%design 5th order Butterworth lowpass filter with cutoff freq wn
[b,a] = butter(5,wn,'low');
y = filtfilt(b,a,x);
```
