## Supplementary material for "Predicting Composition of Genetic Circuits with Resource Competition: Demand and Sensitivity": Experimental data and Matlab code: experimentsettings.m

```
function [filename,infostruct,indstct] = experimentsettings(expname)
%[filename,infostruct,indstct] = experimentsettings(expname)
%output structs corresponding to necessary information required to load
%data contained in the experiment data file. expname is a string
%corresponding to the filename of the data file the user wishes to load.

%input error checking
filelib = {'1st experiment to calibrate GYR',...
    '2nd experiment to calibrate GYR',...
    '3rd experiment to calibrate GYR',...
    '4th experiment to calibrate GYR',...
    '5th experiment to calibrate GYR',...
    '6th experiment to calibrate GYR',...
    'experiments5_6 to calibrate GYR',...
    'WS_WW GFP','SS_SW GFP 1-29','SS_SW GFP 2-4',...
    'ss_sw_yfp','ws_ww_yfp'};
if ~any(contains(filelib,expname))
    error('filename does not match a valid file')
end

%load file
if any(contains(expname,'1st experiment to calibrate GYR','IgnoreCase',true))
    %setup options
    filename = 'J0 characterization\1st experiment to calibrate GYR.xlsx';
    %struct indicating data ranges and labels in raw data excel file
    infostruct = struct();
    infostruct.OD = 'B92:BK591';
    infostruct.GFP = 'B596:BK1095';
    infostruct.RFP = 'B1100:BK1599';
    infostruct.YFP = 'B1604:BK2103';
    %struct of indicies corresponding to cells
    indstct = struct;
    indstct.R =     [1,11,21];
    indstct.Y =     [2,12,22];
    indstct.YG =    [3,13,23];
    indstct.G =     [4,14,24];
    indstct.nG =    [5,15,25];
    indstct.G3 =    [6,16,26];
    indstct.M9 =    [7,9:10,17,19:20,27,29:30,37:40,47:50,57:60];
    indstct.T10 =   [8,18,28];
    indstct.SuperR = [31,41,51];
    indstct.YR =    [32,42,52];
    indstct.BBa04450 = [33,43,53];
    indstct.GR =    [34,44,54];
    indstct.nGR =   [35,45,55];
    indstct.GR3 =   [36,46,56];
elseif any(contains(expname,'2nd experiment to calibrate GYR','IgnoreCase',true))
    %setup options
    filename = 'J0 characterization\2nd experiment to calibrate GYR - wide RFP bandwidth.xlsx';
    %struct indicating data ranges and labels in raw data excel file
    infostruct = struct();
    infostruct.OD = 'B90:BK489';
    infostruct.GFP = 'B494:BK893';
    infostruct.RFP = 'B898:BK1297';
    infostruct.YFP = 'B1302:BK2003';
    %struct of indicies corresponding to cells
    indstct = struct;
    indstct.R =     1:3;
    indstct.GY =    4:6;
    indstct.G =     11:13;
    indstct.GR =    14:16;
    indstct.Y =     21:23;
    indstct.RY =    24:26;
    indstct.M9 =    [7,17:20,27,31:37];
    indstct.T10 =   8:10;
    %indstct.none = [28:30,38:60];
elseif any(contains(expname,'3rd experiment to calibrate GYR','IgnoreCase',true))
    %some problems with RFP in the GFP channel
    %setup options
    filename = 'J0 characterization\3rd experiment to calibrate GYR - wide RFP bandwidth and shifted GFP.xlsx';
    %struct indicating data ranges and labels in raw data excel file
    infostruct = struct();
    infostruct.OD = 'B90:BK889';
    infostruct.GFP = 'B894:BK1693';
    infostruct.RFP = 'B1698:BK2497';
    infostruct.YFP = 'B2502:BK3301';
    %struct of indicies corresponding to cells
    indstct = struct;
    indstct.R =     1:3;
    indstct.GY =    4:6;
    indstct.G =     11:13;
    indstct.GR =    14:16;
    indstct.Y =     21:23;
    indstct.RY =    24:26;
    indstct.M9 =    [7,17:20,27,31:37];
    indstct.T10 =   8:10;
    %indstct.none = [28:30,38:60];
elseif any(contains(expname,'4th experiment to calibrate GYR','IgnoreCase',true))
    %YFP channel saturated on this run
    %setup options
    filename = 'J0 characterization\4th experiment to calibrate GYR.xlsx';
    %struct indicating data ranges and labels in raw data excel file
    infostruct = struct();
    infostruct.OD = 'B90:BK889';
    infostruct.GFP = 'B894:BK1693';
    infostruct.RFP = 'B1698:BK2497';
    infostruct.YFP = 'B2502:BK3301';
    %struct of indicies corresponding to cells
    indstct = struct;
    indstct.R =     1:3;
    indstct.GY =    4:6;
    indstct.G =     11:13;
    indstct.GR =    14:16;
    indstct.Y =     21:23;
    indstct.RY =    24:26;
    indstct.M9 =    [7,17,27,34:37,41:44];
    indstct.T10 =   31:33;
    %indstct.none = [8:10,18:20,28:30,38:40,45:60];
elseif any(contains(expname,'5th experiment to calibrate GYR','IgnoreCase',true))
    %setup options
    filename = 'J0 characterization\5th experiment to calibrate GYR.xlsx';
    %struct indicating data ranges and labels in raw data excel file
    infostruct = struct();
    infostruct.OD = 'B92:BK891';
    infostruct.GFP = 'B896:BK1695';
    infostruct.RFP = 'B1700:BK2499';
    infostruct.YFP = 'B2504:BK3303';
    %struct of indicies corresponding to cells
    indstct = struct;
    indstct.R =     1:3;
    indstct.GY =    4:6;
    indstct.G =     [11,12,13]; %12 shows low GFP levels
    indstct.GR =    14:15;      %16 didn't grow
    indstct.Y =     22:23;      %21 grew very slowly
    indstct.RY =    24:26;      %some GFP in 24
    indstct.M9 =    7;          %,17,27]; %,34:37,41:44];
    indstct.T10 =   31:33;
    %indstct.none = [8:10,18:20,28:30,38:40,45:60];
elseif any(contains(expname,'6th experiment to calibrate GYR','IgnoreCase',true))
    %setup options
    filename = 'J0 characterization\6th experiment to calibrate GYR.xls';
    %struct indicating data ranges and labels in raw data excel file
    infostruct = struct();
    infostruct.OD = 'B92:BK1090';
    infostruct.GFP = 'B1095:BK2093';
    infostruct.RFP = 'B2098:BK3096';
    infostruct.YFP = 'B3101:BK4099';
    %struct of indicies corresponding to cells
    indstct = struct;
    indstct.R =     [8,18,28];%[5,15,25];
    indstct.GY =    [6,16,26];
    indstct.G =     [1,11,21];
    indstct.GR =    [2,12];     %22 grew funny
    indstct.Y =     [3,13,23];
    indstct.RY =    [4,14,24];
    indstct.M9 =    [7,9,17,19,27,29];
    indstct.T10 =   [10,20,30];
elseif any(contains(expname,'experiments_all','IgnoreCase',true))
    %setup options
    filename = 'J0 characterization\experiments_all.xlsx';
    %struct indicating data ranges and labels in raw data excel file
    infostruct = struct();
    infostruct.OD = 'B92:GG891';
    infostruct.GFP = 'B896:GG1695';
    infostruct.RFP = 'B1700:GG2499';
    infostruct.YFP = 'B2504:GG3303';
    %struct of indicies corresponding to cells
    indstct = struct;
    indstct.R =     [1:3,[8,18,28]+63,[39,49,59]+126];         %[5,15,25]+63?;
    indstct.GY =    [4:6,[6,16,26]+63];
    indstct.G =     [11:13,[1,11,21]+63,[9,19,29]+126];       %12 shows low GFP levels
    indstct.GR =    [14:15,[2,12]+63];          %16 didn't grow,22+63 grew funny
    indstct.Y =     [22:23,[3,13,23]+63];       %21 grew very slowly
    indstct.RY =    [24:26,[4,14,24]+63];       %some GFP in 24
    indstct.M9 =    [7,[7,9,17,19,27,29]+63,[40,50,60]+126];   %,17,27]; %,34:37,41:44];
    indstct.T10 =   [31:33,[10,20,30]+63,[10,20,30]+126];
elseif any(contains(expname,'SS_SW GFP 1-29','IgnoreCase',true))
    filename = 'GFP module\SS_SW GFP 1-29.xlsx';
    %struct indicating data ranges and labels in raw data excel file
    infostruct = struct();
    infostruct.OD = 'B90:BK889';
    infostruct.GFP = 'B894:BK1693';
    infostruct.RFP = 'B1698:BK2497';
    infostruct.YFP = 'B2502:BK3301';
    %struct of indicies corresponding to cells
    indstct = struct;
    indstct.SS0G =   [1,11,21];
    indstct.SS1G =   [2,12,22];
    indstct.SS10G =  [3,13,23];
    indstct.SS100G = [4,14,24];
    indstct.SW0G =   [15,25];       %5 didn't grow
    indstct.SW1G =   [6,16,26];
    indstct.SW10G =  [7,17,27];
    indstct.SW100G = [8,18];        %28 didn't grow
    indstct.G =      [9,19,29];
    indstct.T10 =    [10,20,30];
    indstct.SS0R =   [31,41,51];
    indstct.SS1R =   [32,42,52];
    indstct.SS10R =  [33,43,53];
    indstct.SS100R = [34,44,54];
    indstct.SW0R =   [45,55];       %35 didn't grow
    indstct.SW1R =   [36,46,56];
    indstct.SW10R =  [37,47,57];
    indstct.SW100R = [38,48,58];
    indstct.R =      [39,49,59];
    indstct.M9 =     [40,50,60];
elseif any(contains(expname,'SS_SW GFP 2-4','IgnoreCase',true))
    filename = 'GFP module\SS_SW GFP 2-4.xlsx';
    %struct indicating data ranges and labels in raw data excel file
    disp('this only has RFP and YFP for some reason...')
    infostruct = struct();
    infostruct.OD = 'B92:BK1090';
    infostruct.GFP = 'B1095:BK2093';
    infostruct.RFP = 'B2098:BK3096';
    infostruct.YFP = 'B3101:BK4099';
    %struct of indicies corresponding to cells
    indstct = struct;
    indstct.SS0G =    [1,11,21];
    indstct.SS1G =    [2,12,22];
    indstct.SS10G =   [3,13,23];
    indstct.SS100G =  [4,14,24];
    indstct.SW0G =    [5,15,25];
    indstct.SW1G =    [6,16,26];
    indstct.SW10G =   [7,17,27];
    indstct.SW100G =  [8,18,28];
    indstct.G =       [9,19,29];
    indstct.T10 =     [10,20,30];
    indstct.SS0GR =   [31,41,51];
    indstct.SS1GR =   [32,42,52];
    indstct.SS10GR =  [33,43,53];
    indstct.SS100GR = [34,44,54];
    indstct.SW0GR =   [35,45,55];
    indstct.SW1GR =   [36,46,56];
    indstct.SW10GR =  [37,47,57];
    indstct.SW100GR = [38,48,58];
    indstct.R =       [39,49,59];
    indstct.M9 =      [40,50];
elseif any(contains(expname,'WS_WW GFP','IgnoreCase',true))
    filename = 'GFP module\WS_WW GFP 1-30.xlsx';
    %struct indicating data ranges and labels in raw data excel file
    infostruct = struct();
    infostruct.OD = 'B95:BK894';
    infostruct.GFP = 'B899:BK1698';
    infostruct.RFP = 'B1703:BK2502';
    infostruct.YFP = 'B2507:BK3306';
    %struct of indicies corresponding to cells
    indstct = struct;
    indstct.WS0G =    [1,11,21];
    indstct.WS1G =    [2,12,22];
    indstct.WS10G =   [3,13,23];
    indstct.WS100G =  [4,14,24];
    indstct.WW0G =    [5,15,25];
    indstct.WW1G =    [6,16,26];
    indstct.WW10G =   [7,17,27];
    indstct.WW100G =  [8,18,28];
    indstct.G =       [9,19,29];
    indstct.T10 =     [10,20,30];
    indstct.WS0GR =   [31,41,51];
    indstct.WS1GR =   [32,42,52];
    indstct.WS10GR =  [33,43,53];
    indstct.WS100GR = [34,44,54];
    indstct.WW0GR =   [35,45];      %55 grew too fast/too much red?
    indstct.WW1GR =   [36,46,56];
    indstct.WW10GR =  [37,47,57];
    indstct.WW100GR = [38,48,58];
    indstct.R =       [39,49,59];
    indstct.M9 =      [40,50,60];
elseif any(contains(expname,'ss_sw_yfp','IgnoreCase',true))
    filename = 'YFP module\ss_sw_yfp_2-5.xlsx';
    %struct indicating data ranges and labels in raw data excel file
    infostruct = struct();
    infostruct.OD = 'B92:BK1090';
    infostruct.GFP = 'B1095:BK2093';
    infostruct.RFP = 'B2098:BK3096';
    infostruct.YFP = 'B3101:BK4099';
    %struct of indicies corresponding to cells
    indstct = struct;
    indstct.SS0Y =    [1,11,21];
    indstct.SS1Y =    [2,12,22];
    indstct.SS10Y =   [3,13,23];
    indstct.SS100Y =  [4,14,24];
    indstct.SW0Y =    [5,15,25];
    indstct.SW1Y =    [6,16,26];
    indstct.SW10Y =   [7,17,27];
    indstct.SW100Y =  [8,18,28];
    indstct.Y =       [9,19,29];
    indstct.T10 =     [10,20,30];
    indstct.SS0RY =   [31,41,51];
    indstct.SS1RY =   [32,42,52];
    indstct.SS10RY =  [33,43,53];
    indstct.SS100RY = [34,44,54];
    indstct.SW0RY =   [35,45,55];
    indstct.SW1RY =   [36,46,56];
    indstct.SW10RY =  [37,47,57];
    indstct.SW100RY = [38,48,58];
    indstct.R =       [39,49,59];
    indstct.M9 =      [40,50,60];
elseif any(contains(expname,'ws_ww_yfp','IgnoreCase',true))
    filename = 'YFP module\ws_ww_yfp_2-6.xls';
    %struct indicating data ranges and labels in raw data excel file
    infostruct = struct();
    infostruct.OD = 'B92:BK1060';
    infostruct.GFP = 'B1095:BK2063';
    infostruct.RFP = 'B2098:BK3066';
    infostruct.YFP = 'B3101:BK4069';
    %struct of indicies corresponding to cells
    indstct = struct;
    indstct.WS0Y =   [1,11,21];
    indstct.WS1Y =   [2,12,22];
    indstct.WS10Y =  [3,13,23];
    indstct.WS100Y = [4,14,24];
    indstct.WW0Y =   [5,15,25];
    indstct.WW1Y =   [6,16,26];
    indstct.WW10Y =  [7,17,27];
    indstct.WW100Y = [8,18,28];
    indstct.Y =      [9,19,29];
    indstct.T10 =    [10,20,30];
    indstct.WS0R =   [31,41,51];
    indstct.WS1R =   [32,42,52];
    indstct.WS10R =  [33,43,53];
    indstct.WS100R = [34,44,54];
    indstct.WW0R =   [35,45,55];
    indstct.WW1R =   [36,46,56];
    indstct.WW10R =  [37,47,57];
    indstct.WW100R = [38,48,58];
    indstct.R =      [39,49,59];
    indstct.M9 =     [40,50,60];
end
```
