## Supplementary material for "Predicting Composition of Genetic Circuits with Resource Competition: Demand and Sensitivity": Experimental data and Matlab code: filtplatedata.m

```
function filtdata = filtplatedata(datastruct,wn,filtopts)
%filtdata = filtplatedata(datastruct,wn)
%run all data in datastruct through forward/backwards butterworth filter.
%If field has 'time' in it, the field is copied directly. wn is the filter
%cutoff frequency and is a value between 0 and 1 with 1 corresponding to
%the Nyquist frequency

if nargin < 3
    filtopts = 'low';
    if nargin < 2
        wn = 0.2;
    end
end
%input error checking
if wn > 1
    wn = 1;
elseif wn < 0
    wn = 0;
end

%init filter
filtdata = struct;
[b,a] = butter(5,wn,filtopts);
dataprops = fieldnames(datastruct);
%filter data
for jj = 1:length(dataprops)
    %recursion for structs within structs
    if isstruct(datastruct.(dataprops{jj}))
        filtdata.(dataprops{jj}) = filtplatedata(datastruct.(dataprops{jj}),wn,filtopts);
    else
        if ~contains(dataprops{jj},'time')
            %don't filter time fields
            filtdata.(dataprops{jj}) = filtfilt(b,a,datastruct.(dataprops{jj}));
        else
            %filter data
            filtdata.(dataprops{jj}) = datastruct.(dataprops{jj});
        end
    end
end
```
