## Supplementary material for "Predicting Composition of Genetic Circuits with Resource Competition: Demand and Sensitivity": Experimental data and Matlab code: findJ02.m

```
function [outstruct,celldata] = findJ02(celldata,trng,ploton)
%[outstruct,celldata] = findJ02(celldata,ploton)
%use data to calculate values of J0 for all pairs of fluorescent proteins:
%Green, Red, Yellow. celldata must have fields named G, R, Y, GR, GY, RY,
%and RG, each with subfields OD, GFP, RFP, and YFP. celldata should also
%have basal levels subtracted off. ploton is a bool of whether to show
%figures

if nargin < 2
    ploton = false;
end

if ~isempty(trng)
    inds = (celldata.G.time >= min(trng)) & (celldata.G.time <= max(trng));
else
    inds = 1:length(celldata.G.time);
end

%calculate Rhat0, Ghat0, Yhat0 for each pair of constitutive genes
[GhatR, GhatRstd] = calcFPhat(celldata.G.GFPOD(inds,:), celldata.GR.GFPOD(inds,:));
[GhatY, GhatYstd] = calcFPhat(celldata.G.GFPOD(inds,:), celldata.GY.GFPOD(inds,:));
[RhatG, RhatGstd] = calcFPhat(celldata.R.RFPOD(inds,:), celldata.GR.RFPOD(inds,:));
[RhatY, RhatYstd] = calcFPhat(celldata.R.RFPOD(inds,:), celldata.RY.RFPOD(inds,:));
% [YhatG, YhatGstd] = calcFPhat(celldata.Y.GFPOD(inds,:), celldata.GY.GFPOD(inds,:));
% [YhatR, YhatRstd] = calcFPhat(celldata.Y.GFPOD(inds,:), celldata.RY.GFPOD(inds,:));
[YhatG, YhatGstd] = calcFPhat(celldata.Y.YFPOD(inds,:), celldata.GY.YFPOD(inds,:));
[YhatR, YhatRstd] = calcFPhat(celldata.Y.YFPOD(inds,:), celldata.RY.YFPOD(inds,:));

%adjust FPhats by growthrate
%calculate Rhat0, Ghat0, Yhat0 for each pair of constitutive genes
% [GhatR, GhatRstd] = calcFPhat(celldata.G.dODdt(inds,:).*celldata.G.GFPOD(inds,:), ...
%     celldata.GR.dODdt(inds,:).*celldata.GR.GFPOD(inds,:));
% [GhatY, GhatYstd] = calcFPhat(celldata.G.dODdt(inds,:).*celldata.G.GFPOD(inds,:), ...
%     celldata.GY.dODdt(inds,:).*celldata.GY.GFPOD(inds,:));
% [RhatG, RhatGstd] = calcFPhat(celldata.R.dODdt(inds,:).*celldata.R.RFPOD(inds,:), ...
%     celldata.GR.dODdt(inds,:).*celldata.GR.RFPOD(inds,:));
% [RhatY, RhatYstd] = calcFPhat(celldata.R.dODdt(inds,:).*celldata.R.RFPOD(inds,:), ...
%     celldata.RY.dODdt(inds,:).*celldata.RY.RFPOD(inds,:));
% [YhatG, YhatGstd] = calcFPhat(celldata.Y.dODdt(inds,:).*celldata.Y.GFPOD(inds,:), ...
%     celldata.GY.dODdt(inds,:).*celldata.GY.GFPOD(inds,:));
% [YhatR, YhatRstd] = calcFPhat(celldata.Y.dODdt(inds,:).*celldata.Y.GFPOD(inds,:), ...
%     celldata.RY.dODdt(inds,:).*celldata.RY.GFPOD(inds,:));

% %adjust FPhats by growthrate
% %calculate Rhat0, Ghat0, Yhat0 for each pair of constitutive genes
% [GhatR, GhatRstd] = calcFPhat(mean(celldata.G.r(end,:)).*celldata.G.GFPOD(inds,:), ...
%     mean(celldata.GR.r(end,:)).*celldata.GR.GFPOD(inds,:));
% [GhatY, GhatYstd] = calcFPhat(mean(celldata.G.r(end,:)).*celldata.G.GFPOD(inds,:), ...
%     mean(celldata.GY.r(end,:)).*celldata.GY.GFPOD(inds,:));
% [RhatG, RhatGstd] = calcFPhat(mean(celldata.R.r(end,:)).*celldata.R.RFPOD(inds,:), ...
%     mean(celldata.GR.r(end,:)).*celldata.GR.RFPOD(inds,:));
% [RhatY, RhatYstd] = calcFPhat(mean(celldata.R.r(end,:)).*celldata.R.RFPOD(inds,:), ...
%     mean(celldata.RY.r(end,:)).*celldata.RY.RFPOD(inds,:));
% [YhatG, YhatGstd] = calcFPhat(mean(celldata.Y.r(end,:)).*celldata.Y.GFPOD(inds,:), ...
%     mean(celldata.GY.r(end,:)).*celldata.GY.GFPOD(inds,:));
% [YhatR, YhatRstd] = calcFPhat(mean(celldata.Y.r(end,:)).*celldata.Y.GFPOD(inds,:), ...
%     mean(celldata.RY.r(end,:)).*celldata.RY.GFPOD(inds,:));

%output to a struct
FPhatstruct = struct('GhatR',GhatR,'GhatY',GhatY,'RhatG',RhatG,'RhatY',RhatY,...
    'YhatG',YhatG,'YhatR',YhatR,'GhatRstd',GhatRstd,'GhatYstd',GhatYstd,...
    'RhatGstd',RhatGstd,'RhatYstd',RhatYstd,'YhatGstd',YhatGstd,'YhatRstd',YhatRstd);

%calculate J0 for each pair of constitutive genes
[JGR, JGRstd] = calcJ0(GhatR, RhatG, GhatRstd, RhatGstd);
[JGY, JGYstd] = calcJ0(GhatY, YhatG, GhatYstd, YhatGstd);
[JRG, JRGstd] = calcJ0(RhatG, GhatR, RhatGstd, GhatRstd);
[JRY, JRYstd] = calcJ0(RhatY, YhatR, RhatYstd, YhatRstd);
[JYG, JYGstd] = calcJ0(YhatG, GhatY, YhatGstd, GhatYstd);
[JYR, JYRstd] = calcJ0(YhatR, RhatY, YhatRstd, RhatYstd);

%output struct for J calculation and the errors/standard deviations
J0struct = struct('JGR',JGR,'JGY',JGY,'JRG',JRG,'JRY',JRY,'JYG',JYG,'JYR',JYR,...
    'JGRstd',JGRstd,'JGYstd',JGYstd,'JRGstd',JRGstd,'JRYstd',JRYstd,...
    'JYGstd',JYGstd,'JYRstd',JYRstd);

%output data
outstruct = struct();
outstruct.FPhat = FPhatstruct;
outstruct.J0 = J0struct;

if ploton
    tstart = 0;%.2e4;
    %plotting
    %plot Rhat0, Ghat0, Yhat0
    figure; clf;
    subplot(131); hold on
    hp1 = subsampleerrorbar(celldata.M9.time(inds),GhatR,GhatRstd);
    hp2 = subsampleerrorbar(celldata.M9.time(inds),GhatY,GhatYstd);
    ylabel('Ghat0')
    xlabel('time (s)')
    legend([hp1,hp2],{'with RFP','with YFP'},'Location','best')
    xlim([min(celldata.M9.time(inds)),max(celldata.M9.time(inds))])
    set(gca,'fontsize',14)
    
    subplot(132); hold on
    hp3 = subsampleerrorbar(celldata.M9.time(inds),RhatG,RhatGstd);
    hp4 = subsampleerrorbar(celldata.M9.time(inds),RhatY,RhatYstd);
    ylabel('Rhat0')
    xlabel('time (s)')
    legend([hp3,hp4],{'with GFP','with YFP'},'Location','best')
    xlim([min(celldata.M9.time(inds)),max(celldata.M9.time(inds))])
    set(gca,'fontsize',14)
    
    subplot(133); hold on
    hp5 = subsampleerrorbar(celldata.M9.time(inds),YhatG,YhatGstd);
    hp6 = subsampleerrorbar(celldata.M9.time(inds),YhatR,YhatRstd);
    ylabel('Yhat0')
    xlabel('time (s)')
    legend([hp5,hp6],{'with GFP','with RFP'},'Location','best')
    xlim([min(celldata.M9.time(inds)),max(celldata.M9.time(inds))])
    set(gca,'fontsize',14)
    
    %plot J's
    figure;
    subplot(131); hold on
    hp7 = subsampleerrorbar(celldata.M9.time(inds),JGR,JGRstd);
    hp8 = subsampleerrorbar(celldata.M9.time(inds),JGY,JGYstd);
    ylabel('JG')
    xlabel('time (s)')
    xlim([min(celldata.M9.time(inds)),max(celldata.M9.time(inds))])
    %ylim([-2,5])
    legend([hp7,hp8],{'with RFP','with YFP'},'Location','best')
    set(gca,'fontsize',14)
    
    subplot(132); hold on
    hp9 = subsampleerrorbar(celldata.M9.time(inds),JRG,JRGstd);
    hp10 = subsampleerrorbar(celldata.M9.time(inds),JRY,JRYstd);
    ylabel('JR')
    xlabel('time (s)')
    %ylim([-5,5])
    xlim([min(celldata.M9.time(inds)),max(celldata.M9.time(inds))])
    legend([hp9,hp10],{'with GFP','with YFP'},'Location','best')
    set(gca,'fontsize',14)
    
    subplot(133); hold on
    hp11 = subsampleerrorbar(celldata.M9.time(inds),JYG,JYGstd);
    hp12 = subsampleerrorbar(celldata.M9.time(inds),JYR,JYRstd);
    ylabel('JY')
    xlabel('time (s)')
    legend([hp11,hp12],{'with GFP','with RFP'},'Location','best')
    %ylim([-5,5])
    xlim([min(celldata.M9.time(inds)),max(celldata.M9.time(inds))])
    set(gca,'fontsize',14)
end


function [J0,J0std] = calcJ0(xhat1,xhat2,xhat1std,xhat2std)
%calculates J0 (resources used by the perturbation) assuming xhat2 is the
%perturbation and xhat1 is the basal protein. Also calculates the standard
%deviation of J0 if standard deviations of xhat1 and xhat2 are provided

if nargin < 4
    xhat2std = 0;
    if nargin < 3
        xhat1std = 0;
    end
end

%calculate J0 for a pair of constitutive genes
J0 = (1 - xhat2)./(xhat2 + xhat1 - 1);
%estimate error (1 standard deviation) in J's
J0std = sqrt(((1-xhat2).*xhat1std).^2 + (xhat1.*xhat2std).^2)./((xhat2 + xhat1 - 1).^2);
```
