## Supplementary material for "Predicting Composition of Genetic Circuits with Resource Competition: Demand and Sensitivity": Experimental data and Matlab code: findJ03.m

```
function [Jstructout,FPstructout] = findJ03(instruct,FPhatstruct,Jstruct)
%[Jstructout,FPstructout] = findJ03(instruct,FPhatstruct,Jstruct,lambda)
%calculate J's for experiments by specifying how to combine data into
%fluorescence hat with FPHATSTRUCT. Fieldnames of FPHATSTRUCT should
%correspond to names of fluroescence hats and should contain the subfields:
%'alone', 'perturb', 'FPfield', and 'growthfield' which each hold a string
%corresponding to fields of INSTRUCT for 'alone' and 'perturb', and
%subfields for 'FPfield' and 'growthfield'. JSTRUCT specifies how to
%combine fluorescence hat data to obtain J0s. JSTRUCT should have the
%subfields 'hat1' and 'hat2', each holding a string specifiying a field in
%FPHATSTRUCT used to calculate J0 in each case.
%LAMBDA is regularizer for calculating ratios of fluorsecence measurements

if nargin < 2
    %specify how to combine data to get Ghat0, Rhat0, Yhat0 for each
    %experiment
    FPhatstruct.G_R.alone = 'G';
    FPhatstruct.G_R.perturb = 'GR';
    FPhatstruct.G_R.FPfield = 'GFPOD_ss';
    FPhatstruct.G_R.growthfield = 'r_ss';
    FPhatstruct.G_Y.alone = 'G';
    FPhatstruct.G_Y.perturb = 'GY';
    FPhatstruct.G_Y.FPfield = 'GFPOD_ss';
    FPhatstruct.G_Y.growthfield = 'r_ss';
    FPhatstruct.R_G.alone = 'R';
    FPhatstruct.R_G.perturb = 'GR';
    FPhatstruct.R_G.FPfield = 'RFPOD_ss';
    FPhatstruct.R_G.growthfield = 'r_ss';
    FPhatstruct.R_Y.alone = 'R';
    FPhatstruct.R_Y.perturb = 'RY';
    FPhatstruct.R_Y.FPfield = 'RFPOD_ss';
    FPhatstruct.R_Y.growthfield = 'r_ss';
    FPhatstruct.Y_G.alone = 'Y';
    FPhatstruct.Y_G.perturb = 'GY';
    FPhatstruct.Y_G.FPfield = 'YFPOD_ss';
    FPhatstruct.Y_G.growthfield = 'r_ss';
    FPhatstruct.Y_R.alone = 'Y';
    FPhatstruct.Y_R.perturb = 'RY';
    FPhatstruct.Y_R.FPfield = 'YFPOD_ss';
    FPhatstruct.Y_R.growthfield = 'r_ss';
end
if nargin < 3
    %G and R
    Jstruct.JG_R.hat1 = 'G_R';
    Jstruct.JG_R.hat2 = 'R_G';
    Jstruct.JR_G.hat1 = 'R_G';
    Jstruct.JR_G.hat2 = 'G_R';
    %G and Y
    Jstruct.JG_Y.hat1 = 'G_Y';
    Jstruct.JG_Y.hat2 = 'Y_G';
    Jstruct.JY_G.hat1 = 'Y_G';
    Jstruct.JY_G.hat2 = 'G_Y';
    %R and Y
    Jstruct.JR_Y.hat1 = 'R_Y';
    Jstruct.JR_Y.hat2 = 'Y_R';
    Jstruct.JY_R.hat1 = 'Y_R';
    Jstruct.JY_R.hat2 = 'R_Y';
end

%copy info to output
FPstructout = FPhatstruct;
Jstructout = Jstruct;

%loop across FP hat fields
FPhatnames = fieldnames(FPhatstruct);
for ii = 1:length(FPhatnames)
    %locate data to use
    cellalone = FPhatstruct.(FPhatnames{ii}).alone;
    cellperturb = FPhatstruct.(FPhatnames{ii}).perturb;
    FPfield = FPhatstruct.(FPhatnames{ii}).FPfield;
    
    %unpack data
    FPalone = mean(instruct.(cellalone).(FPfield));
    FPalonestd = std(instruct.(cellalone).(FPfield));
    FPperturb = mean(instruct.(cellperturb).(FPfield));
    FPperturbstd = std(instruct.(cellperturb).(FPfield));
    
    %run method to find fluorescence ratios
    [FPhat,FPhatstd] = calcFPhat3(FPalone,FPperturb,FPalonestd,FPperturbstd);
    
    %output FP hats
    FPstructout.(FPhatnames{ii}).FPhatmean = FPhat;
    FPstructout.(FPhatnames{ii}).FPhatstd = FPhatstd;
    
    %with growth rate correction
    if all(isfield(FPhatstruct.(FPhatnames{ii}),{'growthfield'}))
        growthrate = FPhatstruct.(FPhatnames{ii}).growthfield;
        %unpack data with growth rate correction
        xaloneGR = instruct.(cellalone).(FPfield).*instruct.(cellalone).(growthrate);
        xperturbGR = instruct.(cellperturb).(FPfield).*instruct.(cellperturb).(growthrate);
        FPaloneGR = mean(xaloneGR);
        FPalonestdGR = std(xaloneGR);
        FPperturbGR = mean(xperturbGR);
        FPperturbstdGR = std(xperturbGR);
        %run method to find fluorescence ratios with growth rate correction
        [FPhatGR,FPhatstdGR] = calcFPhat3(FPaloneGR,FPperturbGR,FPalonestdGR,FPperturbstdGR);
        %otuput FP hat corrected for growth rate
        FPstructout.(FPhatnames{ii}).FPhatmeangrowth = FPhatGR;
        FPstructout.(FPhatnames{ii}).FPhatstdgrowth = FPhatstdGR;
    end
end

%least squares solution. FIX TO BE MORE ELEGANT?
Br = FPstructout.B_R.FPhatmean;
Rb = FPstructout.R_B.FPhatmean;
Gr = FPstructout.G_R.FPhatmean;
Rg = FPstructout.R_G.FPhatmean;
Bg = FPstructout.B_G.FPhatmean;
Gb = FPstructout.G_B.FPhatmean;

%data matrix
X = [Br-1,0,Br; Rb, 0 Rb-1; Gb, Gb-1, 0; Bg-1, Bg,0; 0, Gr-1, Gr; 0, Rg, Rg-1];
Y = -[Br; Rb; Gb; Bg; Gr; Rg] + 1;

%want weights to be reciprocal of variance if measurements uncorrelated,
%otherwise W = inv(cov(y))
%W = inv(cov(Y));

%model: Y = X*Jest;
%Jest = (X'*W*X)\X'*W*Y
Jlsq = (X'*X)\X'*Y; %this is biased due to correlation between X columns

%output least squares estimate
Jstructout.JB_l_s.J0 = Jlsq(1);
Jstructout.JB_l_s.J0std = 0;     %idk how to calculate variance from ols?
Jstructout.JG_l_s.J0 = Jlsq(2);
Jstructout.JG_l_s.J0std = 0;
Jstructout.JR_l_s.J0 = Jlsq(3);
Jstructout.JR_l_s.J0std = 0;

Jnames = fieldnames(Jstruct);
%loop across J fields
for jj = 1:length(Jnames)
    if all(isfield(Jstruct.(Jnames{jj}),{'hat1','hat2'}))
        %identify data to use
        hat1 = Jstruct.(Jnames{jj}).hat1;
        hat2 = Jstruct.(Jnames{jj}).hat2;
        %unpack data
        FPhat1 = FPstructout.(hat1).FPhatmean;
        FPhat1std = FPstructout.(hat1).FPhatstd;
        FPhat2 = FPstructout.(hat2).FPhatmean;
        FPhat2std = FPstructout.(hat2).FPhatstd;
        %run method
        [J0,J0std] = calcJ0(FPhat1,FPhat2,FPhat1std,FPhat2std);
        %output
        Jstructout.(Jnames{jj}).J0 = J0;
        Jstructout.(Jnames{jj}).J0std = J0std;
        
        %growth rate corrected
        if all(isfield(FPhatstruct.(FPhatnames{ii}),{'growthfield'}))
            %unpack data
            FPhat1GR = FPstructout.(hat1).FPhatmeangrowth;
            FPhat1GRstd = FPstructout.(hat1).FPhatstdgrowth;
            FPhat2GR = FPstructout.(hat2).FPhatmeangrowth;
            FPhat2GRstd = FPstructout.(hat2).FPhatstdgrowth;
            %run method
            [J0GR,J0GRstd] = calcJ0(FPhat1GR,FPhat2GR,FPhat1GRstd,FPhat2GRstd);
            %output
            Jstructout.(Jnames{jj}).J0GR = J0GR;
            Jstructout.(Jnames{jj}).J0GRstd = J0GRstd;
        end
    end
end


function [FPhat,FPhatstd] = calcFPhat3(FPalone,FPtogether,FPalonestd,FPtogetherstd)
%calculates ratios of fluorescent concentrations.
if nargin < 4
    FPalonestd = 0;
end
if nargin < 3
    FPtogetherstd = 0;
end
if nargin < 5
    lambda = 0;
end

%calculate FPhat
FPhat = (FPtogether)./(FPalone);
%estimate error bars assuming uncertianty is dominanted by variance across
%population samples and assuming var(FPtogether), var(FPalone) are uncorrelated
FPhatstd = abs(FPhat).*sqrt( (FPtogetherstd./FPtogether).^2 + ...
    (FPalonestd./FPalone).^2);


function [J0, J0std] = calcJ0(FPhat1,FPhat2,FPhat1std,FPhat2std)
%calculates J0 (resources used by the perturbation) assuming xhat2 is the
%perturbation and xhat1 is the basal protein. Also calculates the standard
%deviation of J0 if standard deviations of xhat1 and xhat2 are provided
if nargin < 4
    FPhat2std = 0;
end
if nargin < 3
    FPhat1std = 0;
end
%calculate J0 for a pair of constitutive genes
J0 = (1 - FPhat2)./(FPhat2 + FPhat1 - 1);
%estimate error (1 standard deviation) in J's
J0std = sqrt(((1-FPhat2).*FPhat1std).^2 + (FPhat1.*FPhat2std).^2)./((FPhat2 + FPhat1 - 1).^2);
```
