## Supplementary material for "Predicting Composition of Genetic Circuits with Resource Competition: Demand and Sensitivity": Experimental data and Matlab code: findJ04.m

```
function [Jstructout,Jvec] = findJ04(instruct,FPstruct,nummods,MCsim)
%Jstructout = findJ04(instruct,FPstruct,nummods,MCsim)
%calculate J's for experiments by specifying how to combine data into
%fluorescence hat with FPHATSTRUCT. Fieldnames of FPHATSTRUCT should
%correspond to names of fluroescence hats and should contain the subfields:
%'alone', 'perturb', 'FPfield', and 'growthfield' which each hold a string
%corresponding to fields of INSTRUCT for 'alone' and 'perturb', and
%subfields for 'FPfield' and 'growthfield'. JSTRUCT specifies how to
%combine fluorescence hat data to obtain J0s. JSTRUCT should have the
%subfields 'hat1' and 'hat2', each holding a string specifiying a field in
%FPHATSTRUCT used to calculate J0 in each case.
%MCSIM is a bool whether to run the monte-carlo simulation

if nargin < 2
    %specify how to combine data to get Ghat0, Rhat0, Yhat0 for each
    %experiment
    %BFP
    FPstruct.B_G.alone1 = 'B';
    FPstruct.B_G.FPfield1 = 'BFPdiffOD_ss';
    FPstruct.B_G.alone2 = 'G';
    FPstruct.B_G.FPfield2 = 'GFPdiffOD_ss';
    FPstruct.B_G.perturb = 'BG';
    FPstruct.B_G.aloneJind = 1;             %for least squares
    FPstruct.B_G.perturbJind = 2;
    FPstruct.B_R.alone1 = 'B';
    FPstruct.B_R.FPfield1 = 'BFPdiffOD_ss';
    FPstruct.B_R.alone2 = 'G';
    FPstruct.B_R.FPfield2 = 'GFPdiffOD_ss';
    FPstruct.B_R.perturb = 'BR';
    FPstruct.B_R.aloneJind = 1;
    FPstruct.B_R.perturbJind = 3;
    %GFP
    FPstruct.G_B.alone1 = 'G';
    FPstruct.G_B.FPfield1 = 'GFPdiffOD_ss';
    FPstruct.G_B.alone2 = 'G';
    FPstruct.G_B.FPfield2 = 'GFPdiffOD_ss';
    FPstruct.G_B.perturb = 'BG';
    FPstruct.G_B.aloneJind = 2;
    FPstruct.G_B.perturbJind = 1;
    FPstruct.G_R.alone1 = 'G';
    FPstruct.G_R.FPfield1 = 'GFPdiffOD_ss';
    FPstruct.G_R.alone2 = 'R';
    FPstruct.G_R.FPfield2 = 'RFPdiffOD_ss';
    FPstruct.G_R.perturb = 'GR';
    FPstruct.G_R.aloneJind = 2;             %index of resource demand coefficient
    FPstruct.G_R.perturbJind = 3;
    %RFP
    FPstruct.R_B.alone1 = 'R';
    FPstruct.R_B.FPfield1 = 'RFPdiffOD_ss';
    FPstruct.R_B.alone2 = 'G';
    FPstruct.R_B.FPfield2 = 'GFPdiffOD_ss';
    FPstruct.R_B.perturb = 'BR';
    FPstruct.R_B.aloneJind = 3;
    FPstruct.R_B.perturbJind = 1;
    FPstruct.R_G.alone1 = 'R';
    FPstruct.R_G.FPfield1 = 'RFPdiffOD_ss';
    FPstruct.R_G.alone2 = 'G';
    FPstruct.R_G.FPfield2 = 'GFPdiffOD_ss';
    FPstruct.R_G.perturb = 'GR';
    FPstruct.R_G.aloneJind = 3;
    FPstruct.R_G.perturbJind = 2;
end
if nargin < 3
    nummods = 3;
end
if nargin < 4 || isempty(MCsim)
    MCsim = false;
end

%init
Jstructout = struct;
FPnames = fieldnames(FPstruct);
Y = zeros(length(FPnames),1);
X = zeros(length(FPnames),nummods);
covY = zeros(length(FPnames));
Jvec = zeros(length(FPnames),1);      %MC simulation

%loop through fields in FP struct to calculate module resource demands
for ii = 1:length(FPnames)
    
    %get metadata
    cellalone1 = FPstruct.(FPnames{ii}).alone1;
    cellalone2 = FPstruct.(FPnames{ii}).alone2;
    FPfield1 = FPstruct.(FPnames{ii}).FPfield1;
    FPfield2 = FPstruct.(FPnames{ii}).FPfield2;
    cellperturb = FPstruct.(FPnames{ii}).perturb;
    aloneJind = FPstruct.(FPnames{ii}).aloneJind;
    perturbJind = FPstruct.(FPnames{ii}).perturbJind;
    
    %unpack data
    FP1alone = mean(instruct.(cellalone1).(FPfield1));
    FP1alonestd = std(instruct.(cellalone1).(FPfield1))./...
        sqrt(length(instruct.(cellalone1).(FPfield1)));     %standard error of mean of FP1alone
    FP1togeth = mean(instruct.(cellperturb).(FPfield1));
    FP1togethstd = std(instruct.(cellperturb).(FPfield1))./...
        sqrt(length(instruct.(cellperturb).(FPfield1)));   %standard error of mean of FP1perturb
    FP2alone = mean(instruct.(cellalone2).(FPfield2));
    FP2alonestd = std(instruct.(cellalone2).(FPfield2))./...
        sqrt(length(instruct.(cellalone2).(FPfield2)));     %standard error of mean of FP2alone
    FP2togeth = mean(instruct.(cellperturb).(FPfield2));
    FP2togethstd = std(instruct.(cellperturb).(FPfield2))./...
        sqrt(length(instruct.(cellperturb).(FPfield2)));    %standard error of mean of FP2perturb
    
    %add noise according to normal distribution with standard error std for
    %monte carlo simulation
    if MCsim == true
        FP1alone = addnoise(FP1alone,FP1alonestd);
        FP2alone = addnoise(FP2alone,FP2alonestd);
        FP1togeth = addnoise(FP1togeth,FP1togethstd);
        FP2togeth = addnoise(FP2togeth,FP2togethstd);
    end
    
    %calculate resource demand for each module pairwise
    [J0,J0std] = calcJ02(FP1alone,FP2alone,FP1togeth,FP2togeth,...
        FP1alonestd,FP2alonestd,FP1togethstd,FP2togethstd);
    %output
    Jvec(ii) = J0;
    Jstructout.(FPnames{ii}).J0 = J0;
    Jstructout.(FPnames{ii}).J0std = J0std;
    
    %create data matricies for least squares regression
    Y(ii) = FP1alone - FP1togeth;
    X(ii,aloneJind) = FP1alone - FP1togeth;
    X(ii,perturbJind) = FP1togeth;
    
    %covariance of Y
    covY(ii,ii) = FP1alonestd.^2 + FP1togethstd.^2;
    
end

%want weights to be reciprocal of variance if measurements uncorrelated,
%otherwise W = inv(cov(y))
W = inv(covY);
%model: Y = X*Jest;
Jlsq = (X'*W*X)\X'*W*Y;
Jvec = [Jvec; Jlsq];

%Jlsq = (X'*X)\X'*Y; %this is biased due to correlation between X columns
%output least squares estimate
Jstructout.B_l_s.J0 = Jlsq(1);
Jstructout.B_l_s.J0std = 0;     %calculate standard error from MC simulation
Jstructout.G_l_s.J0 = Jlsq(2);
Jstructout.G_l_s.J0std = 0;
Jstructout.R_l_s.J0 = Jlsq(3);
Jstructout.R_l_s.J0std = 0;
%Jstructout.Jvec = Jvec;


function [J0,J0std] = calcJ02(FP1alone,FP2alone,FP1togeth,FP2togeth,FP1astd,FP2astd,FP1tstd,FP2tstd)
%calculates J0 (resources used by the perturbation) assuming xhat2 is the
%perturbation and xhat1 is the basal protein. Also calculates the standard
%deviation of J0 if standard deviations of xhat1 and xhat2 are provided

%compute errors comparing alone and perturbed conditions
%e1 = FP1alone - FP1togeth;
e2 = FP2alone - FP2togeth;
%calculate J0 for a pair of constitutive genes
num = FP1alone*e2;
den = FP1alone*FP2togeth + FP1togeth*FP2alone - FP1alone*FP2alone;
J0 = num./den;

dfdF1alone = FP2alone*FP1togeth*e2./(den.^2);
dfdF1togeth = -FP2alone*FP1alone*e2./(den.^2);
dfdF2alone = FP1alone*FP2togeth*FP1togeth./(den.^2);
dfdF2togeth = -FP1alone*FP2alone*FP1togeth./(den.^2);

J0std = sqrt(dfdF1alone^2*FP1astd^2 + dfdF1togeth^2*FP1tstd^2 + ...
    dfdF2alone^2*FP2astd^2 + dfdF2togeth^2*FP2tstd^2);


%for MC simulation
function y = addnoise(xmean,xstd)
y = xmean + randn(1)*xstd;


%-------------------old------------------
function [FPhat,FPhatstd] = calcFPhat3(FPalone,FPtogether,FPalonestd,FPtogetherstd)
%calculates ratios of fluorescent concentrations.
if nargin < 4
    FPalonestd = 0;
end
if nargin < 3
    FPtogetherstd = 0;
end
if nargin < 5
    lambda = 0;
end

%calculate FPhat
FPhat = (FPtogether)./(FPalone);
%estimate error bars assuming uncertianty is dominanted by variance across
%population samples and assuming var(FPtogether), var(FPalone) are uncorrelated
FPhatstd = abs(FPhat).*sqrt( (FPtogetherstd./FPtogether).^2 + ...
    (FPalonestd./FPalone).^2);

function [J0, J0std] = calcJ0(FPhat1,FPhat2,FPhat1std,FPhat2std)
%calculates J0 (resources used by the perturbation) assuming xhat2 is the
%perturbation and xhat1 is the basal protein. Also calculates the standard
%deviation of J0 if standard deviations of xhat1 and xhat2 are provided
if nargin < 4
    FPhat2std = 0;
end
if nargin < 3
    FPhat1std = 0;
end
%calculate J0 for a pair of constitutive genes
J0 = (1 - FPhat2)./(FPhat2 + FPhat1 - 1);
%estimate error (1 standard deviation) in J's
J0std = sqrt(((1-FPhat2).*FPhat1std).^2 + (FPhat1.*FPhat2std).^2)./((FPhat2 + FPhat1 - 1).^2);
```
