## Supplementary material for "Predicting Composition of Genetic Circuits with Resource Competition: Demand and Sensitivity": Experimental data and Matlab code: findQ.m

```
function Qstruct = findQ(celldata,J0,J0std,measurename,perturbname,ploton,errscale)
%Qstruct = findQ(celldata,J0,J0std,measurename,perturbname,ploton,errscale)
%find Q and S and estimate uncertianty

if nargin < 7
    errscale = 1;
    if nargin < 6
        ploton = true;
        if nargin < 5
            perturbname = 'R';
            if nargin < 4
                measurename = 'G';
                if nargin < 3
                    J0std = 0;
                end
            end
        end
    end
end

%init
Qstruct = struct();
meashat = cell(0);
perturbhat = cell(0);
meashatstd = cell(0);
perturbhatstd = cell(0);
% Yhat = cell(0);
% Yhatstd = cell(0);
Q = cell(0);
S = cell(0);
Qstd = cell(0);
Sstd = cell(0);

%names of induction levels labels
cellnames = fieldnames(celldata);
inducedinds = contains(cellnames,'W') | contains(cellnames,'S');
namesmeasonly = cellnames(contains(cellnames,measurename) & ...
    ~(contains(cellnames,[measurename,perturbname]) | ...
    contains(cellnames,[perturbname,measurename])) & inducedinds);
namesboth = cellnames(contains(cellnames,perturbname) & inducedinds);

%check that inductions for RFP and GFP have same length
if length(namesmeasonly) ~= length(namesboth)
    error('length of inductions is not the same for GFP and RFP')
else
    %loop across all induction levels
    for ii = 1:length(namesmeasonly)
        %find Ghat, Rhat, Yhat and estimate uncertianty
        [meashat{ii}, meashatstd{ii}] = ...
            calcFPhat(celldata.(namesmeasonly{ii}).([measurename,'FPnorm']),...
            celldata.(namesboth{ii}).([measurename,'FPnorm']));
        [perturbhat{ii}, perturbhatstd{ii}] = ...
            calcFPhat(celldata.(perturbname).([perturbname,'FPnorm']),...
            celldata.(namesboth{ii}).([perturbname,'FPnorm']));
        
        %find Q and S and estimate uncertianty
        [Q{ii},Qstd{ii}] = calcQ(perturbhat{ii},J0,perturbhatstd{ii},J0std);
        [S{ii},Sstd{ii}] = calcS(meashat{ii},perturbhat{ii},J0,...
            meashatstd{ii},perturbhatstd{ii},J0std);
        
        %output to struct
        Qstruct.(namesmeasonly{ii}) = struct();
        Qstruct.(namesmeasonly{ii}).measurename = measurename;
        Qstruct.(namesmeasonly{ii}).perturbname = perturbname;
        Qstruct.(namesmeasonly{ii}).meashat = meashat{ii};
        Qstruct.(namesmeasonly{ii}).meashatstd = meashatstd{ii};
        Qstruct.(namesmeasonly{ii}).perturbhat = perturbhat{ii};
        Qstruct.(namesmeasonly{ii}).perturbhatstd = perturbhatstd{ii};
        %Qstruct.(namesmeasonly{ii}).Yhat = Yhat{ii};
        %Qstruct.(namesmeasonly{ii}).Yhatstd = Yhatstd{ii};
        Qstruct.(namesmeasonly{ii}).Q = Q{ii};
        Qstruct.(namesmeasonly{ii}).Qstd = Qstd{ii};
        Qstruct.(namesmeasonly{ii}).S = S{ii};
        Qstruct.(namesmeasonly{ii}).Sstd = Sstd{ii};
    end
end

if ploton
    tstart = 0;
    
    %plotting
    %plot Rhat0, Ghat0, Yhat0
    figure; clf;
    subplot(121); hold on
    for jj = 1:length(meashat)
        subsampleerrorbar(celldata.M9.time,meashat{jj},errscale*meashatstd{jj})
    end
    ylabel([measurename,'hat, measurement'])
    xlabel('time (s)')
    legend(namesmeasonly,'Location','best')
    xlim([tstart,max(celldata.M9.time)])
    
    subplot(122); hold on
    for jj = 1:length(perturbhat)
        subsampleerrorbar(celldata.M9.time,perturbhat{jj},errscale*perturbhatstd{jj})
    end
    ylabel([perturbname,'hat, perturbation'])
    xlabel('time (s)')
    legend(namesmeasonly,'Location','best')
    xlim([tstart,max(celldata.M9.time)])
    
%     subplot(133); hold on
%     for jj = 1:length(Yhat)
%         subsampleerrorbar(celldata.M9.time,Yhat{jj},errorscale*Yhatstd{jj})
%     end
%     ylabel('Yhat')
%     xlabel('time (s)')
%     legend(namesmeasonly,'Location','best')
%     xlim([tstart,max(celldata.M9.time)])
    
    %plot Q and S
    figure;
    subplot(121); hold on
    for jj = 1:length(Q)
        subsampleerrorbar(celldata.M9.time,Q{jj},errscale*Qstd{jj})
    end
    ylabel('Q')
    xlabel('time (s)')
    xlim([tstart,max(celldata.M9.time)])
    legend(namesmeasonly,'Location','best')
    
    subplot(122); hold on
    for jj = 1:length(S)
        subsampleerrorbar(celldata.M9.time,S{jj},errscale*Sstd{jj})
    end
    ylabel('S')
    xlabel('time (s)')
    xlim([tstart,max(celldata.M9.time)])
    legend(namesmeasonly,'Location','best')
end


function [Q,Qstd] = calcQ(Rhat,J0,Rhatstd,J0std)

%calculate Q and estimate the uncertainty
Q = (1./Rhat - 1).*(1+J0);
%not sure if this is the correct error estimate formulat
Qstd = sqrt(((1+J0)./(Rhat.^2).*Rhatstd).^2 + ((1./Rhat - 1).*J0std).^2);


function [S,Sstd] = calcS(Ghat,Rhat,J0,Ghatstd,Rhatstd,J0std)

%calculate S and estimate the uncertainty
S = ((Ghat-1).*(1+J0.*(1-Rhat)))./(Ghat.*(J0.*(1+J0)));
dSdG = (1+J0.*(1-Rhat))./(Ghat.^2.*(J0.*(1+J0)));
dSdR = (Ghat-1)./(Ghat.*(1+J0));
dSdJ = ((Ghat-1).*(1-Rhat))./(Ghat.*(J0.^2));
%estimate uncertainty
Sstd = sqrt((dSdG.*Ghatstd).^2 + (dSdR.*Rhatstd).^2 + (dSdJ.*J0std).^2);
```
