## Supplementary material for "Predicting Composition of Genetic Circuits with Resource Competition: Demand and Sensitivity": Experimental data and Matlab code: findss2.m

```
function outstruct = findss2(instruct,timestruct,datafields)
%average values over a user specificed interval

cellnames = fieldnames(instruct);
outstruct = instruct;

%loop through different cell types
for ii = 1:length(cellnames)
    
    %check that fields in timestruct match instruct celltypes
    if isfield(timestruct,cellnames{ii})
        
        %find time indicies of steady state
        time = instruct.(cellnames{ii}).time;
        tmin = min(timestruct.(cellnames{ii}));
        tmax = max(timestruct.(cellnames{ii}));
        [~,indmin] = min(abs(time - tmin));
        [~,indmax] = min(abs(time - tmax));
        
        %loop across datafields
        for jj = 1:length(datafields)
            if isfield(instruct.(cellnames{ii}),datafields{jj})
                data = instruct.(cellnames{ii}).(datafields{jj})(indmin:indmax,:);
                %ss = mean(data);
                ss = median(data);
                outstruct.(cellnames{ii}).([datafields{jj},'_ss']) = ss;
                disp([cellnames{ii},' ',datafields{jj}])
                
                if adtest(data(:)) == true
                    disp('fails AD test--samples NOT drawn from a Gaussian')
                else
                    disp('passes AD test--samples are drawn from a Gaussian')
                end
                
            end
        end
        %add growth rate at final time measurement
        if isfield(instruct.(cellnames{ii}),'r')
            rss = instruct.(cellnames{ii}).r;
            outstruct.(cellnames{ii}).r_ss = rss(end,:);
        end
    end
end
```
