## Supplementary material for "Predicting Composition of Genetic Circuits with Resource Competition: Demand and Sensitivity": Experimental data and Matlab code: findss3.m

```
function outstruct = findss3(instruct,timestruct,datafields,avefun,ploton,figh)
%outstruct = findss3(instruct,timestruct,datafields,avefun,ploton,figh)
%average values over a user specificed interval

if nargin < 5 || isempty(ploton)
    ploton = false;
end
if nargin < 4 || isempty(avefun)
    %function to apply to get steady state
    avefun = @mean;       %or @median
end

cellnames = fieldnames(timestruct);
outstruct = instruct;

%setup plot
if ploton
    if nargin < 5 || isempty(figh)
        figh = figure;
    end
    n = length(datafields);
    m = ceil(9/16*sqrt(n));
end

data = cell(length(datafields),1);
[outmat,outstdmat] = deal(zeros(length(datafields),length(cellnames)));

%loop through fields
for ii = 1:length(datafields)
    %loop through cells
    for jj = 1:length(cellnames)
        
        %check that fields in timestruct match instruct celltypes
        if isfield(instruct,cellnames{jj})
            
            %get indicies for range in steady state window
            time = instruct.(cellnames{jj}).time;
            tmin = min(timestruct.(cellnames{jj}));
            tmax = max(timestruct.(cellnames{jj}));
            [~,indmin] = min(abs(time - tmin));
            [~,indmax] = min(abs(time - tmax));
            
            %get data from time range
            tmp = instruct.(cellnames{jj}).(datafields{ii})(indmin:indmax,:);
            %apply average/median function to each sample
            ss = avefun(tmp);
            
            %store for histogram plots
            data{ii}{jj} = tmp(:);
            %store steady state data
            outstruct.(cellnames{jj}).([datafields{ii},'_ss']) = avefun(ss);
            %outstruct.(cellnames{jj}).([datafields{ii},'_ss_std']) = ...
            %   sqrt(sum(var(tmp))./numel(tmp));  %gives too small error
            %   bars due to correlations between consecutive samples
            outstruct.(cellnames{jj}).([datafields{ii},'_ss_std']) = std(ss)./numel(ss);
            
            outmat(ii,jj) = avefun(ss);
            outstdmat(ii,jj) = std(ss)./numel(ss);
            
            %test for normality
            %disp([cellnames{jj},' ',datafields{ii}])
            if 0 %numel(tmp(~isnan(tmp))) > 4
                if adtest(tmp(:)) == true
                    disp('fails AD test--samples NOT drawn from a Gaussian')
                else
                    disp('passes AD test--samples are drawn from a Gaussian')
                end
            end
        end
    end
    
    %plot
    if ploton
        %plot bar chart of steady state data
        if length(figh) == length(datafields)
            figure(figh(ii)); clf;
            set(gca,'fontsize',16)
        else
            figure(figh);
            subplot(m,ceil(n/m),ii); cla;
        end
        x = 1:size(outmat,2);
        bh = bar(x,outmat(ii,:)); hold on
        set(bh,'barwidth',0.6);
        if contains(datafields{ii},'BFP')
            set(bh,'FaceColor',[0.48,0.79,1])
        elseif contains(datafields{ii},'GFP')
            set(bh,'FaceColor',[0.7,1,0.49])
        elseif contains(datafields{ii},'RFP')
            set(bh,'FaceColor',[1,0.51,0.51])
        end
        xticks(x);
        xticklabels(cellnames);
        errorbar(outmat(ii,:),outstdmat(ii,:),'.k','linewidth',2);
        ylabel([datafields{ii},' steady state'])
        
        if 0
            %plot histograms of distributions
            figure(figh);
            ax = subplot(m,ceil(n/m),ii);
            multiplehist(data{ii},ax,30,'probability')
            xlabel(datafields{ii})
            legend(cellnames,'Location','Best','fontsize',10);
            ylabel('prob mass')
        end
    end
end
```
