## Supplementary material for "Predicting Composition of Genetic Circuits with Resource Competition: Demand and Sensitivity": Experimental data and Matlab code: findsteadystate.m

```
function [outstruct,ssoutstruct] = findsteadystate(cellstruct,ploton)
%[outstruct,youtstruct] = findsteadystate(cellstruct,fitparams,ploton)
%find steady state in plate reader data

if nargin < 3
    ploton = true;
end

%settings for determining "goodness" of steady state
slopetol = 100;
rsqtol = 0.8;
tmintime = 3;           %[hrs]

%init
outstruct = struct;
ssoutstruct = struct;
t = cellstruct.M9.time;
celltype = fieldnames(cellstruct);

%loop across cell types
for ii = 1:length(celltype)
    
    if strcmp(celltype{ii},'M9')
        %outstruct.(celltypes{ii}) = cellstruct.(celltypes{ii});
        continue
    end
    
    disp(celltype{ii})
    K = cellstruct.(celltype{ii}).K;
    r = cellstruct.(celltype{ii}).r;
    P0 = cellstruct.(celltype{ii}).P0;
    
    %find time window of good growthrate in exp phase
    tmax = 0.25*sqrt(K.^2./((K-P0).*r.^2.*P0));     %taken from quadratic term in Taylor expansion
    tmin = tmintime*ones(length(tmax),1);           %wait for transient
    
    if tmin > tmax
        warning('no steady state possible due to poor growthrate')
    end
    
    %find indicies corresponding to time window
    [~,minind] = min(abs(t(:)*ones(1,length(tmin)) - repmat(tmin(:)',[length(t),1])));
    [~,maxind] = min(abs(t(:)*ones(1,length(tmax)) - repmat(tmax(:)',[length(t),1])));
    
    dataflds = fieldnames(cellstruct.(celltype{ii}));
    if ploton
        figure;
        q = 1;
    end
    %loop across data fields
    for k = 1:length(dataflds)
        
        if contains(dataflds{k},'time') || (contains(dataflds{k},'OD') ...
                && ~contains(dataflds{k},'FP')) || contains(dataflds{k},...
                'growth') || (contains(dataflds{k},'FP') && endsWith(dataflds{k},'FP'))
            %outstruct.(celltypes{ii}).(datafields{k}) = cellstruct.(celltypes{ii}).(datafields{k});
            continue
        end
        
        %init
        [m1,m2,m3,b1,b2,b3,rsq1,rsq2,rsq3,rmse1,rmse2,rmse3,yout,ystdout] = ...
            deal(zeros(length(maxind),1));
        %loop across cell populations of same type
        for jj = 1:length(maxind)
            %data in desired time window
            y = cellstruct.(celltype{ii}).(dataflds{k})(minind(jj):maxind(jj),jj);
            y = y(:);
            x = t(minind(jj):maxind(jj));
            x = x(:);
            %linear fit first half, last half, whole dataset to investigate
            %trends
            [fitobj1,gof1] = fit(x(1:round(end/2)),y(1:round(end/2)),'poly1');
            [fitobj2,gof2] = fit(x(round(end/2):end),y(round(end/2):end),'poly1');
            [fitobj3,gof3] = fit(x,y,'poly1');
            %unpack fit output
            m1(jj) = fitobj1.p1;
            b1(jj) = fitobj1.p2;
            rsq1(jj) = gof1.rsquare;
            rmse1(jj) = gof1.rmse;
            m2(jj) = fitobj2.p1;
            b2(jj) = fitobj2.p2;
            rsq2(jj) = gof2.rsquare;
            rmse2(jj) = gof2.rmse;
            m3(jj) = fitobj3.p1;
            b3(jj) = fitobj3.p2;
            rsq3(jj) = gof3.rsquare;
            rmse3(jj) = gof3.rmse;
            
            %criterion for determining if the concentration reached steady state
            if all([m1(jj),m2(jj),m3(jj)] < 0)
                %take last half of data
                yout(jj) = max([0,mean(y(round(end/2):end))]);
                ystdout(jj) = std(y(round(end/2):end));
                disp('yout')
            elseif m1(jj) > slopetol && m2(jj) > slopetol && m3(jj) < slopetol
                %take last half of data
                yout(jj) = max([0,mean(y(round(end/2):end))]);
                ystdout(jj) = std(y(round(end/2):end));
                disp('yout')
            elseif all([m1(jj),m2(jj),m3(jj)] < slopetol) && any([m1(jj),m2(jj),m3(jj)] > 0)
                yout(jj) = max([0,mean(y(round(end/2):end))]);
                ystdout(jj) = std(y(round(end/2):end));
                disp('yout')
            else
                %no steady state
                yout(jj) = NaN;
                ystdout(jj) = NaN;
                disp(['No steady state',num2str(jj)])
            end
        end
        
        %erase datapoints that are not steady states
        yout(isnan(yout)) = [];
        ystdout(isnan(ystdout)) = [];
        
        %output data
        disp([celltype{ii},', ',dataflds{k}])
        disp([rsq1,rsq2,rsq3])
        disp([m1,m2,m3])
        disp([b1,b2,b3])
        outstruct.(celltype{ii}).tmax = tmax;
        outstruct.(celltype{ii}).tmin = tmin;
        outstruct.(celltype{ii}).(dataflds{k}).fit = {fitobj1,fitobj2,fitobj3};
        outstruct.(celltype{ii}).(dataflds{k}).gof = {gof1,gof2,gof3};
        outstruct.(celltype{ii}).(dataflds{k}).yout = yout;
        outstruct.(celltype{ii}).(dataflds{k}).m = [m1,m2,m3];
        outstruct.(celltype{ii}).(dataflds{k}).b = [b1,b2,b3];
        outstruct.(celltype{ii}).(dataflds{k}).rsq = [rsq1,rsq2,rsq3];
        outstruct.(celltype{ii}).(dataflds{k}).rmse = [rmse1,rmse2,rmse3];
        outstruct.(celltype{ii}).(dataflds{k}).ystdout = ystdout;
        
        %output just steady state
        ssoutstruct.([celltype{ii},'_',dataflds{k}]) = mean(yout);
        
        %plot data and fits to verify
        if ploton
            subplot(2,3,q);
            plot(t(min(minind):max(maxind)),...
                cellstruct.(celltype{ii}).(dataflds{k})(min(minind):max(maxind),:));
            hold on;
            for l = 1:length(m1)
                plot(t(min(minind):max(maxind)),saturate(t(min(minind):max(maxind))*m1(l)+b1(l)),'b-.')
                plot(t(min(minind):max(maxind)),saturate(t(min(minind):max(maxind))*m2(l)+b2(l)),'r-.')
                plot(t(min(minind):max(maxind)),saturate(t(min(minind):max(maxind))*m3(l)+b3(l)),'k--')
                if ~isempty(yout)
                    plot(linspace(min(tmin),max(tmax),5),yout*ones(1,5),'-ko')
                end
            end
            title([celltype{ii},', ',dataflds{k}])
            xlabel('time (hrs)')
            ylabel(dataflds{k})
            xlim([min(tmin),max(tmax)])
            q = q + 1;
            hold off;
        end
    end
end
```
