## Supplementary material for "Predicting Composition of Genetic Circuits with Resource Competition: Demand and Sensitivity": Experimental data and Matlab code: fitgrowthrate2.m

```
function cellstructout = fitgrowthrate2(cellstruct,growthrateopts,ODfld,ploton,verbose,figh)
%cellstructout = fitgrowthrate2(cellstruct,growthrateopts,ODfld,ploton,verbose)
%fit OD with either exponential, logisitic curve and take deriviative to
%find a smooth growthrate curve or take finite difference and filter to
%find growthrate. cellstruct must contain fields for each growth condition
%and each should be a struct containing the fields OD and time.
%growthrateopts is a string either 'expfit', 'logisticfit', or 'derivative'
%to decide on the fitting method. ploton is a bool whether to plot the OD
%fits and growthrates.

if nargin < 6 || isempty(figh)
    figh = figure;
end
if nargin < 5
    verbose = false;
end
if nargin < 4
    ploton = true;
end
if nargin < 3
    ODfld = 'OD';
end
if nargin < 2
    growthrateopts = 'logisticfit';  %expfit ,logisticfit, derivative
end
fntsze = 12;
%barplot colors
barcolor = [149,209,249]./255;      %light blue
% barcolor = [0,0.4470,0.7410];     %dark blue
strexp = '(\<[0-9]*\.)\w*(\s)?';    %regexp for formatting parameter strings in figure titles
%match string beginning with numeric characters [0-9] then string has a '.'
%then match rest of the word until an optional 1 trailing whitespace character

%init
cellstructout = cellstruct;
celltypes = fieldnames(cellstruct);

%loop across cell conditions
disp('fitting growth rate...')
if contains(growthrateopts,'exp','IgnoreCase',true)
    %fit with exponential curve
    %init
    [acell,bcell,astdcell,bstdcell] = deal(zeros(length(celltypes),1));
    for ii = 1:length(celltypes)
        %init
        ODfld = fieldnames(cellstruct.(celltypes{ii}));
        n = size(cellstruct.(celltypes{ii}).(ODfld),2);
        time = cellstruct.(celltypes{ii}).time;
        ODfit = zeros(size(cellstruct.(celltypes{ii}).(ODfld)));
        dODdt = zeros(size(cellstruct.(celltypes{ii}).(ODfld)));
        rsq = zeros(n,1);
        a = zeros(n,1);
        b = zeros(n,1);
        %loop through each cell type
        for jj = 1:n
            %run fitting on each column of data within the fields (celltypes)
            if iscell(cellstruct.(celltypes{ii}).(ODfld))
                [fitobj,gof] = fit(time,cellstruct.(celltypes{ii}).(ODfld){jj},'exp1');
            elseif isnumeric(cellstruct.(celltypes{ii}).(ODfld))
                [fitobj,gof] = fit(time,cellstruct.(celltypes{ii}).(ODfld)(:,jj),'exp1');
            end
            %output fitted OD and dOD/dt
            a(jj) = fitobj.a;
            b(jj) = fitobj.b;
            ODfit(:,jj) = a*exp(b.*time);
            dODdt(:,jj) = a*b*exp(b*time);
            rsq(jj) = gof.rsquare;
        end
        
        %output parameters
        cellstructout.(celltypes{ii}).a = a(:)';
        cellstructout.(celltypes{ii}).b = b(:)';
        cellstructout.(celltypes{ii}).rsquare = rsq(:)';
        cellstructout.(celltypes{ii}).fittype = growthrateopts;
        cellstructout.(celltypes{ii}).ODfit = ODfit;
        cellstructout.(celltypes{ii}).dODdt = dODdt;
        cellstructout.(celltypes{ii}).growthrate = dODdt./ODfit;
        
        %for plotting average parameters for each cell type
        acell(ii) = mean(a);
        astdcell(ii) = std(a);
        bcell(ii) = mean(b);
        bstdcell(ii) = std(b);
        
        %plotting
        if (ploton && verbose)% || any(rsq < 0.99) || any(rsq > 1)
            rsqstr = regexp(num2str(rsq(:)',3),strexp,'match');
            astr = regexp(num2str(a(:)',3),strexp,'match');
            bstr = regexp(num2str(b(:)',3),strexp,'match');
            
            %plot
            figure(figh); clf;
            subplot(211);
            plot(time,cellstruct.(celltypes{ii}).(ODfld),time,ODfit,'--','linewidth',2)
            title([celltypes{ii},': exp fit, R^2 = [',[rsqstr{:}],']'])
            xlabel('time (hrs)')
            ylabel('OD')
            set(gca,'fontsize',fntsze)
            subplot(212);
            plot(time,cellstructout.(celltypes{ii}).growthrate,'linewidth',2)
            title(['a = [',[astr{:}],'; b = [',[bstr{:}],']'])
            xlabel('time (hrs)')
            ylabel('growth rate (OD/s)')
            set(gca,'fontsize',fntsze)
        end
    end
elseif contains(growthrateopts,'logistic','IgnoreCase',true)
    %fit with logisitic curve
    %init
    [Kcell,Kstdcell,rcell,rstdcell,P0cell,P0stdcell] = deal(zeros(length(celltypes),1));
    for ii = 1:length(celltypes)
        %init
        n = size(cellstruct.(celltypes{ii}).(ODfld),2);
        time = cellstruct.(celltypes{ii}).time;
        ODfit = zeros(size(cellstruct.(celltypes{ii}).(ODfld)));
        dODdt = zeros(size(cellstruct.(celltypes{ii}).(ODfld)));
        rsq = zeros(n,1);
        K = zeros(n,1);
        P0 = zeros(n,1);
        r = zeros(n,1);
        %set up logistic fitting problem
        opts = fitoptions('method','NonlinearLeastSquares',...
            'StartPoint',[0.5,0.02,0.4],'lower',[0,0,0]);
        ft = fittype('K*a*exp(r*x)/(K+a*(exp(r*x)-1))',...
            'coeff',{'K','a','r'},'options',opts);
        %loop through each cell type
        for jj = 1:n
            %run fitting
            if iscell(cellstruct.(celltypes{ii}).(ODfld))
                [fitobj,gof] = fit(time,cellstruct.(celltypes{ii}).(ODfld){jj},ft);
            elseif isnumeric(cellstruct.(celltypes{ii}).(ODfld))
                [fitobj,gof] = fit(time,cellstruct.(celltypes{ii}).(ODfld)(:,jj),ft);
            end
            %output parameters from fit to OD, and dOD/dt as logistic curves
            K(jj) = fitobj.K;
            P0(jj) = fitobj.a;
            r(jj) = fitobj.r;
            ODfit(:,jj) = K(jj)*P0(jj)*exp(r(jj).*time)./(K(jj) + P0(jj)* ...
                (exp(r(jj).*time) - 1));
            dODdt(:,jj) = K(jj)*P0(jj)*r(jj)*(K(jj) - P0(jj))*exp(r(jj).*time)./...
                ((K(jj) + P0(jj)*(exp(r(jj).*time) - 1)).^2);
            rsq(jj) = gof.rsquare;
        end
        %write to output
        OD = cellstruct.(celltypes{ii}).(ODfld);
        cellstructout.(celltypes{ii}).ODfit = ODfit;
        cellstructout.(celltypes{ii}).dODdt = dODdt;
        cellstructout.(celltypes{ii}).growthrate = dODdt./ODfit;
        cellstructout.(celltypes{ii}).growthrateOD = dODdt./(OD).^2;
        %growthrate = r*(K-a)/(K + a*(exp(r*t)-1))
        
        %output parameters
        cellstructout.(celltypes{ii}).K = K(:)';
        cellstructout.(celltypes{ii}).r = r(:)';
        cellstructout.(celltypes{ii}).P0 = P0(:)';
        cellstructout.(celltypes{ii}).rsquare = rsq(:)';
        cellstructout.(celltypes{ii}).fittype = growthrateopts;
        
        %for plotting average parameters for each cell type
        Kcell(ii) = mean(K);
        Kstdcell(ii) = std(K);
        rcell(ii) = mean(r);
        rstdcell(ii) = std(r);
        P0cell(ii) = mean(P0);
        P0stdcell(ii) = std(P0);
        
        %plotting
        if ploton && verbose% || (any(rsq < 0.85) && ~strcmpi(celltypes{ii},'M9'))
            %format parameter strings for titles
            rsqstr = regexp(num2str(rsq(:)',3),strexp,'match');
            Kstr = regexp(num2str(K(:)',3),strexp,'match');
            P0str = regexp(num2str(P0(:)',3),strexp,'match');
            rstr = regexp(num2str(r(:)',3),strexp,'match');
            %plot
            figure;
            subplot(311);
            plot(time,cellstruct.(celltypes{ii}).(ODfld),'linewidth',2); hold on;
            set(gca,'colororderindex',1)
            plot(time,ODfit,'--','linewidth',2)
            title([celltypes{ii},': logistic fit R^2 = ',[rsqstr{:}]])
            ylabel('OD')
            xlim([min(time),max(time)])
            set(gca,'fontsize',fntsze)
            subplot(312);
            plot(time,cellstructout.(celltypes{ii}).growthrate,'linewidth',2)
            ylabel('d OD/dt')
            xlim([min(time),max(time)])
            title(['K = [',[Kstr{:}],']; P0 = [',[P0str{:}],']'])
            set(gca,'fontsize',fntsze)
            subplot(313);
            plot(time,dODdt./ODfit,'linewidth',2)
            xlabel('time (hrs)')
            ylabel('growthrate')
            title(['r = [',[rstr{:}],']'])
            xlim([min(time),max(time)])
            set(gca,'fontsize',fntsze)
        end
    end
else
    for ii = 1:length(celltypes)
        %take finite difference derivative and filter. Does not give good
        %results
        %init
        time = cellstruct.(celltypes{ii}).time;
        m = 5;
        wn = 0.01;
        %set up low pass filter
        [b,a] = butter(5,wn,'low');
        %take finite difference and filter to find dOD/dt
        dODdt = filtfilt(b,a,diff(cellstruct.(celltypes{ii}).(ODfld),m));
        %write to output
        cellstructout.(celltypes{ii}).growthrate = ...
            [zeros(m,size(cellstruct.(celltypes{ii}).(ODfld),2));dODdt];
        cellstructout.(celltypes{ii}).fittype = growthrateopts;
        
        if ploton && verbose
            %plot
            figure;
            subplot(211)
            plot(time, cellstruct.(celltypes{ii}).(ODfld),'linewidth',2)
            title([celltypes{ii},': finite difference'])
            xlabel('time (hrs)')
            ylabel('OD')
            set(gca,'fontsize',fntsze)
            subplot(212);
            plot(time,cellstructout.(celltypes{ii}).growthrate,'linewidth',2)
            xlabel('time (s)')
            ylabel('growth rate (OD/s)')
            set(gca,'fontsize',fntsze)
        end
    end
end

if ploton
    %plot bargraph of parameters for each cell type
    c = categorical(strrep(celltypes,'_','-'));
    names = strrep(celltypes,'_','-');
    if strcmp(growthrateopts,'logisticfit')
        %plot bargraph of K, r, P0
        figure(figh); clf;
        %plot P0
        subplot(311);
        bar(P0cell,'FaceColor',barcolor); hold on
        set(gca,'XTick',1:length(P0cell),'XTickLabel',[],'fontsize',fntsze,...
            'XLim',[0.5,length(rcell)+0.5]);
        errorbar(P0cell,P0stdcell,'.k','linewidth',2)
        ylabel('P0')
        %plot K
        %subplot(311);
        %bar(c,Kcell,'FaceColor',barcolor); hold on
        %errorbar(Kcell,Kstdcell,'.k','linewidth',2)
        %ylabel('K')
        %set(gca,'fontsize',fntsze)
        %plot r
        subplot(3,1,[2,3]);
        bar(rcell,'FaceColor',barcolor); hold on
        set(gca,'XTickLabel',names,'XTick',1:length(rcell),...
            'XTickLabelRotation',90,'XLim',[0.5,length(rcell)+0.5]);
        errorbar(rcell,rstdcell,'.k','linewidth',2)
        ylabel('r')
        set(gca,'fontsize',fntsze)
    elseif strcmp(growthrateopts,'expfit')
        %plot bargraph of fitted parameters
        figure(figh); clf;
        %plot a
        subplot(211);
        bar(c,acell,'FaceColor',barcolor); hold on
        errorbar(acell,astdcell,'.k','linewidth',2)
        ylabel('a')
        set(gca,'fontsize',fntsze)
        %plot b
        subplot(212);
        bar(c,bcell,'FaceColor',barcolor); hold on
        errorbar(bcell,bstdcell,'.k','linewidth',2)
        ylabel('b')
        set(gca,'fontsize',fntsze)
    end
end
```
