## Supplementary material for "Predicting Composition of Genetic Circuits with Resource Competition: Demand and Sensitivity": Experimental data and Matlab code: fitgrowthrate.m

```
function [cellstructout,params] = fitgrowthrate(cellstruct,growthrateopts,ploton)
%cellstructout = fitgrowthrate(cellstruct,growthrateopts,ploton)
%fit OD with either exponential, logisitic curve and take deriviative to
%find a smooth growthrate curve or take finite difference and filter to
%find growthrate. cellstruct must contain fields for each growth condition
%and each should be a struct containing the fields OD and time.
%growthrateopts is a string either 'expfit', 'logisticfit', or 'derivative'
%to decide on the fitting method. ploton is a bool whether to plot the OD
%fits and growthrates. Initial conditions may need to be adjusted for the
%logistic fit to ensure a good fit.

if nargin < 3
    ploton = false;
end
if nargin < 2 || isempty(growthrateopts)
    growthrateopts = 'logisticfit';  %expfit ,logisticfit, derivative
end

fntsze = 14;
cellstruct = orderfields(cellstruct);
params = struct;

cellstructout = cellstruct;
celltypes = fieldnames(cellstruct);
%loop across cell conditions
disp('fitting growth rate...')
for ii = 1:length(celltypes)
    n = size(cellstruct.(celltypes{ii}).OD,2);
    %init
    time = cellstruct.(celltypes{ii}).time;
    ODfit = zeros(size(cellstruct.(celltypes{ii}).OD));
    dODdt = zeros(size(cellstruct.(celltypes{ii}).OD));
    if strcmp(growthrateopts,'expfit')
        %fit with exponential curve
        rsq = zeros(n,1);
        a = zeros(n,1);
        b = zeros(n,1);
        for jj = 1:n
            [fitobj,gof] = fit(time,cellstruct.(celltypes{ii}).OD(:,jj),'exp1');
            a(jj) = fitobj.a;
            b(jj) = fitobj.b;
            ODfit(:,jj) = a*exp(b.*time);
            dODdt(:,jj) = a*b*exp(b*time);
            rsq(jj) = gof.rsquare;
        end
        
        %output parameters
        params.(celltypes{ii}).a = a;
        params.(celltypes{ii}).b = b;
        params.(celltypes{ii}).rsquare = rsq;
        params.(celltypes{ii}).fittype = growthrateopts;
        
        cellstructout.(celltypes{ii}).ODfit = ODfit;
        cellstructout.(celltypes{ii}).dODdt = dODdt;
        cellstructout.(celltypes{ii}).growthrate = dODdt./ODfit;
        
        %for plotting average parameters for each cell type
        acell(ii) = mean(a);
        astdcell(ii) = std(a);
        bcell(ii) = mean(b);
        bstdcell(ii) = std(b);
        
        %plotting
        if ploton || any(rsq < 0.99) || any(rsq > 1)
            figure;
            subplot(211)
            plot(time,cellstruct.(celltypes{ii}).OD,time,ODfit,'--')
            title([celltypes{ii},': exp fit, R^2 = ',num2str(rsq)])
            xlabel('time (hrs)')
            ylabel('OD')
            subplot(212)
            plot(time,cellstructout.(celltypes{ii}).growthrate)
            xlabel('time (hrs)')
            ylabel('growth rate (OD/s)')
        end
    elseif strcmp(growthrateopts,'logisticfit')
        %fit with logisitic curve
        opts = fitoptions('method','NonlinearLeastSquares',...
            'StartPoint',[0.5,0.02,0.4],'lower',[0,0,0]);
        ft = fittype('K*a*exp(r*x)/(K+a*(exp(r*x)-1))',...
            'coeff',{'K','a','r'},'options',opts);
        rsq = zeros(n,1);
        K = zeros(n,1);
        P0 = zeros(n,1);
        r = zeros(n,1);
        for jj = 1:n
            [fitobj,gof] = fit(time,cellstruct.(celltypes{ii}).OD(:,jj),ft);
            K(jj) = fitobj.K;
            P0(jj) = fitobj.a;
            r(jj) = fitobj.r;
            ODfit(:,jj) = K(jj)*P0(jj)*exp(r(jj).*time)./(K(jj) + P0(jj)* ...
                (exp(r(jj).*time) - 1));
            dODdt(:,jj) = K(jj)*P0(jj)*r(jj)*(K(jj) - P0(jj))*exp(r(jj).*time)./...
                ((K(jj) + P0(jj)*(exp(r(jj).*time) - 1)).^2);
            rsq(jj) = gof.rsquare;
        end
        cellstructout.(celltypes{ii}).ODfit = ODfit;
        cellstructout.(celltypes{ii}).dODdt = dODdt;
        cellstructout.(celltypes{ii}).growthrate = dODdt./ODfit;
        %growthrate = r*(K-a)/(K + a*(exp(r*t)-1))
        
        %output parameters
        params.(celltypes{ii}).K = K;
        params.(celltypes{ii}).r = r;
        params.(celltypes{ii}).P0 = P0;
        params.(celltypes{ii}).rsquare = rsq;
        params.(celltypes{ii}).fittype = growthrateopts;
        
        %for plotting average parameters for each cell type
        Kcell(ii) = mean(K);
        Kstdcell(ii) = std(K);
        rcell(ii) = mean(r);
        rstdcell(ii) = std(r);
        P0cell(ii) = mean(P0);
        P0stdcell(ii) = std(P0);
        
        %plotting
        if ploton || any(rsq < 0.99) || any(rsq > 1)
            figure;
            subplot(311);
            plot(time,cellstruct.(celltypes{ii}).OD,time,ODfit,'--','linewidth',2)
            title([celltypes{ii},': logistic fit R^2 = ',num2str(rsq(:)')])
            ylabel('OD')
            xlim([min(time),max(time)])
            set(gca,'fontsize',fntsze)
            
            subplot(312);
            plot(time,cellstructout.(celltypes{ii}).growthrate,'linewidth',2)
            ylabel('d OD/dt')
            xlim([min(time),max(time)])
            title(['K = ',num2str(K(:)'),'; P0 = ',num2str(P0(:)')])
            set(gca,'fontsize',fntsze)
            
            subplot(313);
            plot(time,dODdt./ODfit,'linewidth',2)
            xlabel('time (hrs)')
            ylabel('growthrate')
            title(['r = ',num2str(r(:)')])
            xlim([min(time),max(time)])
            set(gca,'fontsize',fntsze)
        end
    else
        %take finite difference derivative and filter. Does not give good
        %results
        m = 5;
        wn = 0.01;
        [b,a] = butter(5,wn,'low');
        dODdt = filtfilt(b,a,diff(cellstruct.(celltypes{ii}).OD,m));
        cellstructout.(celltypes{ii}).growthrate = ...
            [zeros(m,size(cellstruct.(celltypes{ii}).OD,2));dODdt];
        
        params.(celltypes{ii}).fittype = growthrateopts;
        
        if ploton
            figure;
            subplot(211)
            plot(time, cellstruct.(celltypes{ii}).OD)
            title([celltypes{ii},': finite difference'])
            xlabel('time (hrs)')
            ylabel('OD')
            subplot(212)
            plot(time,cellstructout.(celltypes{ii}).growthrate)
            xlabel('time (s)')
            ylabel('growth rate (OD/s)')
        end
    end
end

%plot bargraph of parameters for each cell type
c = categorical(celltypes);
if strcmp(growthrateopts,'logisticfit') && (nargin < 3 || ploton)
    %plot bargraph of K, r, P0
    figure;
    %plot K
    subplot(311);
    bar(c,Kcell,'FaceColor',[0,0.4470,0.7410]); hold on
    errorbar(Kcell,Kstdcell,'.k','linewidth',2)
    ylabel('K')
    set(gca,'fontsize',fntsze)
    %plot a
    subplot(312);
    bar(c,rcell,'FaceColor',[0,0.4470,0.7410]); hold on
    errorbar(rcell,rstdcell,'.k','linewidth',2)
    ylabel('r')
    set(gca,'fontsize',fntsze)
    %plot P0
    subplot(313);
    bar(c,P0cell,'FaceColor',[0,0.4470,0.7410]); hold on
    errorbar(P0cell,P0stdcell,'.k','linewidth',2)
    ylabel('P0')
    set(gca,'fontsize',fntsze)
    
elseif strcmp(growthrateopts,'expfit') && (nargin < 3 || ploton)
    %plot bargraph of fitted parameters
    figure;
    %plot a
    subplot(211);
    bar(c,acell,'FaceColor',[0,0.4470,0.7410]); hold on
    errorbar(acell,astdcell,'.k','linewidth',2)
    ylabel('a')
    set(gca,'fontsize',fntsze)
    %plot b
    subplot(212);
    bar(c,bcell,'FaceColor',[0,0.4470,0.7410]); hold on
    errorbar(bcell,bstdcell,'.k','linewidth',2)
    ylabel('b')
    set(gca,'fontsize',fntsze)
end
```
