## Supplementary material for "Predicting Composition of Genetic Circuits with Resource Competition: Demand and Sensitivity": Experimental data and Matlab code: fluorescencebleed.m

```
function C = fluorescencebleed(celldata,metastruct,ploton,maxstruct)
%compensate for fluorescence bleed between channels
%this should be done after background subtraction


if nargin < 2 || isempty(metastruct)
    metastruct.BFP = {'B'};
    metastruct.GFP = {'G'};
    metastruct.RFP = {'R'};
end
if nargin < 3 || isempty(ploton)
    ploton = false;
end
if nargin < 4 || isempty(maxstruct)
    maxstruct.BFP = 1e4;
    maxstruct.GFP = 1e4;
    maxstruct.RFP = 0.7e4;
end


FPfields = fieldnames(metastruct);
n = length(FPfields); %number of channels
C = zeros(n);

%find correlation between channels using cells expressing only BFP/GFP/RFP
%as a control
for ii = 1:length(FPfields)
    
    %get cells and channels
    celltype = metastruct.(FPfields{ii}){1};
    
    if ploton
        figure; clf;
    end
    
    for jj = 1:n
        %compute the correlation between fluorescence channels
        ydata = celldata.(celltype).(FPfields{ii});
        xdata = celldata.(celltype).(FPfields{jj});
        
        %get timepoints where all cell repeats are below thresholds
        inds = all(ydata<maxstruct.(FPfields{ii}) & xdata<maxstruct.(FPfields{jj}),2);
        
        %trim data
        ydata2 = ydata(inds,:);
        xdata2 = xdata(inds,:);
        
        %averaging in this way may cause some problems if cell repeats
        %arent lined up correctly against time
        corr = mean(xdata2,2)'/mean(ydata2,2)';
        C(jj,ii) = corr;
        
        if ploton
            %plot
            subplot(1,n,jj);
            plot(xdata,ydata,xdata,1/corr*xdata,'--',xdata2,ydata2,'linewidth',1.5)
            title(celltype)
            ylabel(FPfields{ii})
            xlabel(FPfields{jj})
            set(gca,'fontsize',14)
        end
    end
end
```
