## Supplementary material for "Predicting Composition of Genetic Circuits with Resource Competition: Demand and Sensitivity": Experimental data and Matlab code: formatplatedilutions.m

```
function datastruct = formatplatedilutions(OD_data,GFP_data,RFP_data,datastruct)
%datastruct = formatplatedilutions(datastruct,OD_data,GFP_data,RFP_data)
if nargin < 4 || ~all(isfield(datastruct,{'OD','GFP','RFP','time','inds'}))
    %init struct
    datastruct = struct('OD',[],'GFP',[],'RFP',[],'time',[],'inds',[]);
end

%shape raw data into (rows x col x time)
ODout = permute(reshape(OD_data',12,8,[]),[2,1,3]);
GFPout = permute(reshape(GFP_data',12,8,[]),[2,1,3]);
RFPout = permute(reshape(RFP_data',12,8,[]),[2,1,3]);

%concatenate rows and cols of the plate into one column and output (time x wells(96))
OD = reshape(ODout,96,[])';
GFP = reshape(GFPout,96,[])';
RFP = reshape(RFPout,96,[])';

time = (1:5:5*size(ODout,3))/60;    %time step in hours
inds = size(ODout,3);               %index of last row to take for each dilution

%output struct
datastruct.OD = cat(1,datastruct.OD, OD);
datastruct.GFP = cat(1,datastruct.GFP, GFP);
datastruct.RFP = cat(1,datastruct.RFP, RFP);
datastruct.time = cat(1,datastruct.time, time(:));
datastruct.inds = cat(1,datastruct.inds, inds);
```
