## Supplementary material for "Predicting Composition of Genetic Circuits with Resource Competition: Demand and Sensitivity": Experimental data and Matlab code: growthrate.m

```
function outstruct = growthrate(instruct,ODfld)

if nargin < 2 || isempty(ODfld)
    ODfld = 'OD';
end

%smallest valid value of OD measurement
thresh = 0.01;

ploton = false;

cellnames = fieldnames(instruct);
outstruct = instruct;

%loop through cells
for ii = 1:length(cellnames)
    
    if isfield(instruct.(cellnames{ii}),ODfld)
        %data
        data = instruct.(cellnames{ii}).(ODfld);
        %only take rows where smallest 
        minrow = min(data,[],2);
        
        %log transform
        X = log(data(minrow > thresh,:));
        tdata = instruct.(cellnames{ii}).time;
        t = tdata(minrow > thresh);
        dt = mean(diff(t));
        %calculate slope
        if size(X,1) > 10
            %max of numerical difference
            gr1 = max(diff5pt(X,dt));
            %robust fit
            gr2 = zeros(2,size(X,2));
            for jj = 1:size(X,2)
                gr2(:,jj) = robustfit(t,X(:,jj));
            end
            
            if ploton
                %predictions
                y2fit = gr2(1,:) + (t)*gr2(2,:);
                y1fit = X(1,:) + (t-t(1))*gr1;
                %plot
                figure;
                subplot(211);
                plot(t,X,t,y1fit,'--',t,y2fit,':');
                title(cellnames{ii})
                subplot(212);
                plot(t(1:end-4),diff5pt(X,dt),t(2:end),diff(X)/dt,'--')
            end
            
            %output data
            outstruct.(cellnames{ii}).gr1 = gr1;
            outstruct.(cellnames{ii}).gr2 = gr2(2,:);
        else
            outstruct.(cellnames{ii}).gr1 = zeros(1,size(X,2));
            outstruct.(cellnames{ii}).gr2 = zeros(1,size(X,2));
        end
    end
end


function y = diff5pt(X,dx)
%five point stencil derivative approximation
y = (X(1:end-4,:) - 8*X(2:end-3,:) + 8*X(4:end-1,:) - X(5:end,:))./(12*dx);
```
