## Supplementary material for "Predicting Composition of Genetic Circuits with Resource Competition: Demand and Sensitivity": Experimental data and Matlab code: importfile96well.m

```
function importfile96well(filename,numRow,numCol,outputname)
% PLEASE MAKE SURE DATA INPUT CONTAINS BLOCKS IN THE SEQUENCE OF RFP, GFP, OD
% importfile(filename,numRow,numCol,outputname)
% filename: "filename" file (.txt format) output from platereader
% numRow: number of rows (of wells) read by the plate reader
% numCol: number of columns (of wells) read by the plate reader
% outputname: string, outputname.mat saved in current folder
% outputname.mat contains GFPOut, RFPOut and ODOut, each is a matrix with
% size Tx(total number of samples), where total number of samples =
% numRow*numCol and T is the total number of time points collected. The
% columns are arranged such that the first row of wells are read from left
% to right and then the second row...until the last row.
% Yili 20180903
%% Initialize variables.
delimiter = '\t';

startRow = 3;
endRow = inf;

%% Read columns of data as strings:
% For more information, see the TEXTSCAN documentation.
formatSpec = '%q%q%q%q%q%q%q%q%q%q%q%q%q%q%[^\n\r]';

%% Open the text file.
fileID = fopen(filename,'r');

%% Read columns of data according to format string.
% This call is based on the structure of the file used to generate this
% code. If an error occurs for a different file, try regenerating the code
% from the Import Tool.
textscan(fileID, '%[^\n\r]', startRow(1)-1, 'WhiteSpace', '', 'ReturnOnError', false);
dataArray = textscan(fileID, formatSpec, endRow(1)-startRow(1)+1, 'Delimiter', delimiter, 'ReturnOnError', false);
for block=2:length(startRow)
    frewind(fileID);
    textscan(fileID, '%[^\n\r]', startRow(block)-1, 'WhiteSpace', '', 'ReturnOnError', false);
    dataArrayBlock = textscan(fileID, formatSpec, endRow(block)-startRow(block)+1, 'Delimiter', delimiter, 'ReturnOnError', false);
    for col=1:length(dataArray)
        dataArray{col} = [dataArray{col};dataArrayBlock{col}];
    end
end

%% Close the text file.
fclose(fileID);

%% Convert the contents of columns containing numeric strings to numbers.
% Replace non-numeric strings with NaN.
raw = repmat({''},length(dataArray{1}),length(dataArray)-1);
for col=1:length(dataArray)-1
    raw(1:length(dataArray{col}),col) = dataArray{col};
end
numericData = NaN(size(dataArray{1},1),size(dataArray,2));

for col=[1:1:14]
    % Converts strings in the input cell array to numbers. Replaced non-numeric
    % strings with NaN.
    rawData = dataArray{col};
    for row=1:size(rawData, 1);
        % Create a regular expression to detect and remove non-numeric prefixes and
        % suffixes.
        regexstr = '(?<prefix>.*?)(?<numbers>([-]*(\d+[\,]*)+[\.]{0,1}\d*[eEdD]{0,1}[-+]*\d*[i]{0,1})|([-]*(\d+[\,]*)*[\.]{1,1}\d+[eEdD]{0,1}[-+]*\d*[i]{0,1}))(?<suffix>.*)';
        try
            result = regexp(rawData{row}, regexstr, 'names');
            numbers = result.numbers;
            
            % Detected commas in non-thousand locations.
            invalidThousandsSeparator = false;
            if any(numbers==',');
                thousandsRegExp = '^\d+?(\,\d{3})*\.{0,1}\d*$';
                if isempty(regexp(thousandsRegExp, ',', 'once'))
                    numbers = NaN;
                    invalidThousandsSeparator = true;
                end
            end
            % Convert numeric strings to numbers.
            if ~invalidThousandsSeparator;
                numbers = textscan(strrep(numbers, ',', ''), '%f');
                numericData(row, col) = numbers{1};
                raw{row, col} = numbers{1};
            end
        catch me
        end
    end
end


%% Replace non-numeric cells with NaN
R = cellfun(@(x) ~isnumeric(x) && ~islogical(x),raw); % Find non-numeric cells
raw(R) = {NaN}; % Replace non-numeric cells

%% Create output variable
data = cell2mat(raw);

%% Above autogenerated by Matlab
% importfile(filename,numRow,plateCol,SampleLocations);

data(:,1) = [];
data(:,end) = [];

[~,W] = size(data);
% get rid of empty entries and rows that indicate column numbers on the
% plate

NaN_Inx = find(sum(isnan(data),2)==W);
Num_Inx = find(ismember(data,[1:1:12],'rows'));
data([NaN_Inx;Num_Inx],:) = [];

NaN_Inx_col = find(sum(isnan(data),1)==length(data));
data(:,NaN_Inx_col) = [];

[L,~] = size(data);

RFP_data = data(1:L/3,:);
GFP_data = data(L/3+1:L*2/3,:);
OD_data = data(L*2/3+1:end,:);

T_length = L/numRow/3;

% NumSamples = numRow*numCol;
% RFPOut = zeros(T_length,NumSamples);
% GFPOut = zeros(T_length,NumSamples);
% ODOut = zeros(T_length,NumSamples);

% for i = 1:1:T_length
%     for j = 1:1:NumSamples
%         SampleRow = floor((j-1)/numCol+1)
%         SampleCol = mod((j-1),numCol)+1
%         
%         RFPOut(i,j) = RFP_data((i-1)*numRow+SampleRow,SampleCol);
%         GFPOut(i,j) = GFP_data((i-1)*numRow+SampleRow,SampleCol);
%         ODOut(i,j) = OD_data((i-1)*numRow+SampleRow,SampleCol);
%     end
% end

save(outputname,'RFP_data','GFP_data','OD_data')
%save(outputname,'RFPOut','GFPOut','ODOut')
```
