## Supplementary material for "Predicting Composition of Genetic Circuits with Resource Competition: Demand and Sensitivity": Experimental data and Matlab code: importplate.m

```
function [platedata,platetrim] = importplate(filename,outputname,platesize)
% platedata = importplate(filename,outputname,platesize)
% filename: "filename" file (.txt format) output from platereader
% outputname: string, outputname.mat saved in current folder. Default is
% same as FILENAME
% platesize: array: [numrows, numcols]. Default is [8,12]
% outputname.mat contains a struct 'platedata' with fields corresponding to
% labels by the platereader. Each field has an 3 dimensional array 
% with [numrows,numcols,numsamples]. Also creates the struct 'platetrim'
% without empty wells (no NaN's)
% Yili 2018-09-03
% major edit by Cameron 7-13-2019

if nargin < 1
    filename = 'data\J0_Experiment_7-3-2019_4.txt';
end
if nargin < 2
    outputname = [filename(1:end-4),'.mat'];
end
if nargin < 3
    platesize = [8,12];
end
%% Initialize variables.
delimiter = '\t';

startRow = 1;
endRow = inf;

%% Read columns of data as strings:
% For more information, see the TEXTSCAN documentation.
formatSpec = '%s\t%[^\n\r]';

%% Open the text file.
fileID = fopen(filename,'r');

%% Read columns of data according to format string.
dataArray = textscan(fileID, formatSpec, endRow(1)-startRow(1)+1, 'Delimiter', delimiter, 'ReturnOnError', false);

%% Close the text file.
fclose(fileID);

%%
%first column
regexstr = '^\w*:';
[names,~,rowinds] = unique(dataArray{1}(~cellfun(@isempty,regexp(dataArray{1},regexstr))),'stable');
names2 = strrep(strrep(names,':','_'),',','_');

%indicies of row at start of each measurement
nameinds = find(~cellfun(@isempty,regexp(dataArray{1},'^\w*:')));

%split on tab character
platestrs = regexp(dataArray{2},delimiter,'split');

%init structs
platedata = struct;
platetrim = struct;

%loop through each measurement
for ii = 1:length(nameinds)
    
    %search for start index
    startind = nameinds(ii) + find(strncmp(dataArray{1}(nameinds(ii):nameinds(ii)+5),'A',1)) - 1;
    
    %get strings where data is
    plate0 = platestrs(startind:(startind+platesize(1)-1));
    plate1 = NaN(platesize);      %init; empty cells are NaN
    %loop through rows in the plate array
    for jj = 1:length(plate0)
        %find indicies where there is numeric data in each row
        inds = ~cellfun(@isempty,plate0{jj}(1:platesize(2)));
        %convert strings to numbers and put numeric data in array
        plate1(jj,inds) = cellfun(@str2double,plate0{jj}(inds));
    end
    %append into struct
    if ~all(isnan(plate1),'all')
        if isfield(platedata,names2{rowinds(ii)})
            platedata.(names2{rowinds(ii)}) = cat(3,platedata.(names2{rowinds(ii)}),plate1);
        else
            platedata.(names2{rowinds(ii)}) = plate1;
        end
    end
end

%trim off any wells that are always NaN/empty
datafields = fieldnames(platedata);
for k = 1:length(datafields)
    %get each field
    data = platedata.(datafields{k});
    
    %find active wells
    activewells = ~all(isnan(data),3);
    
    %find size of active well area
    numsamples = size(data,3);
    numrows = nnz(any(activewells,2));
    numcols = nnz(any(activewells,1));
    
    activewells2 = any(activewells,1) & any(activewells,2);
    
    %only keep data where wells are active
    activeinds = repmat(activewells2,[1,1,numsamples]);
    platetrim.(datafields{k}) = reshape(data(activeinds),[numrows,numcols,numsamples]);
end

%save as matlab data
if nargout == 0
    save(outputname,'platedata','platetrim')
end
```
