## Supplementary material for "Predicting Composition of Genetic Circuits with Resource Competition: Demand and Sensitivity": Experimental data and Matlab code: inducedmodplots.m

```
Yrfp = abs([modulestruct2.Y240.RFPdiffOD,modulestruct2.RY193.RFPdiffOD,...
    abs(modulestruct2.RY192.RFPdiffOD)]);
Yyfp = abs([modulestruct2.Y240.YFPdiffOD,modulestruct2.RY193.YFPdiffOD,...
    abs(modulestruct2.RY192.YFPdiffOD)]);
Erfp = [modulestruct2.Y240.RFPdiffODstd,modulestruct2.RY193.RFPdiffODstd,...
    abs(modulestruct2.RY192.RFPdiffODstd)];
Eyfp = [modulestruct2.Y240.YFPdiffODstd,modulestruct2.RY193.YFPdiffODstd,...
    abs(modulestruct2.RY192.YFPdiffODstd)];

figure(7); clf;
errorbar([1e-3, 0.3, 1, 10]',Yyfp(:,1),Eyfp(:,1),'--o','linewidth',1.7); hold on
errorbar([1e-3, 0.3, 1, 10]',Yyfp(:,2),Eyfp(:,2),'--o','linewidth',1.7); 
errorbar([1e-3, 1e-3, 1, 10]',Yyfp(:,3),Eyfp(:,3),'--o','linewidth',1.7)
set(gca,'yscale','log','xscale','log','fontsize',14)
xlabel('inducer [nM]')
ylabel('ss YFP production rate')

figure(8); clf;
errorbar([1e-3, 0.3, 1, 10]',Yrfp(:,1),Erfp(:,1),'--o','linewidth',1.7); hold on
errorbar([1e-3, 0.3, 1, 10]',Yrfp(:,2),Erfp(:,2),'--o','linewidth',1.7); 
errorbar([1e-3, 1e-3, 1, 10]',Yrfp(:,3),Erfp(:,3),'--o','linewidth',1.7)
set(gca,'yscale','log','xscale','log','fontsize',14)
ylim([1e4,1e5])
xlabel('inducer [nM]')
ylabel('ss RFP production rate')
```
