## Supplementary material for "Predicting Composition of Genetic Circuits with Resource Competition: Demand and Sensitivity": Experimental data and Matlab code: linearfitcellstruct.m

```
function fitstruct = linearfitcellstruct(cellstruct,xfield,yfield,xrng,cellfields,figh)
%fitstruct = linearfitcellstruct(cellstruct,xfield,yfield,xrng,cellfields,figh)
%fit a line to the fields XFIELD and YFIELD across the points within XRNG.
%CELLFIELDS is a cell array of cell names to plot with the respective fit
%lines and FIGH is the figure handle on which to plot. Returns FITSTRUCT
%which contains the subsubfields: p = fit parameters with p(1) = y intercept
%and p(2) = slope, x, and y.

ploton = true;

%get fields for each cell type
celltypes = fieldnames(cellstruct);
if nargin > 4 && ~isempty(cellfields)
    cellnames = intersect(cellfields,celltypes,'stable');
else; cellnames = celltypes; 
end

%loop through cell conditions
for ii = 1:length(cellnames)
    if all(isfield(cellstruct.(cellnames{ii}),{xfield,yfield}))
        %init
        fitstruct.(cellnames{ii}) = struct;
        fitparams = zeros(2,size(x0,2));
        [xfit,yfit] = deal(zeros(size(x0)));
        
        %get data
        x0 = cellstruct.(cellnames{ii}).(xfield);
        y0 = cellstruct.(cellnames{ii}).(yfield);
        
        %loop through trials
        for jj = 1:size(x0,2)
            %locate indicies 
            inds = x0(:,jj) > min(xrng) & x0(:,jj) < max(xrng);
        
            if nnz(inds) > 10
                %get data within range
                x2 = x0(inds,jj);
                y2 = y0(inds,jj);
                
                %run linear fit
                %fitparams(:,jj) = robustfit(x2,y2);
                fitparams(:,jj) = regress(y2,[ones(size(x2)),x2]);
                
                %calculate 
                xfit = x2;
                yfit = fitparams(1,jj) + x2*fitparams(2,jj);
            else
                fitparams(:,jj) = [];
            end
        end
        %output
        fitstruct.(cellnames{ii}).p = fitparams;
        fitstruct.(cellnames{ii}).x = xfit;
        fitstruct.(cellnames{ii}).y = yfit;
    else
        continue
    end
end

%plot
if ploton
    if nargin < 6 || isempty(figh)
        figh = figure;
    end
    figure(figh); clf;
    plotsubfield2(cellstruct,xfield,yfield,figh,cellfields);
    setallsubplots(figh,'axis','NextPlot','add');       %hold on for all subplots
    plotsubfield2(fitstruct,'x','y',figh,cellfields);
    %setallsubplots(figh,'line',{'LineStyle','LineColor'},{'--','k'});
end
```
