## Supplementary material for "Predicting Composition of Genetic Circuits with Resource Competition: Demand and Sensitivity": Experimental data and Matlab code: logsubtract.m

```
function outstruct = logsubtract(cellstruct,FPfields)

if nargin < 2
    FPfields = {'BFP','GFP','RFP'};
end

ODfield = 'OD';

cellfields = fieldnames(cellstruct);
outstruct = cellstruct;

for ii = 1:length(cellfields)
    
    log(cellstruct.(cellfields{ii}).OD);
    
    for jj = 1:length(FPfields)
        outstruct.(cellfields{ii}).([FPfields{jj},'logdiff']) = ...
            exp(log(cellstruct.(cellfields{ii}).(FPfields{jj}))...
            - log(cellstruct.(cellfields{ii}).(ODfield)));
    end
end
```
