## Supplementary material for "Predicting Composition of Genetic Circuits with Resource Competition: Demand and Sensitivity": Experimental data and Matlab code: makefigs_RS5.m

```
tic

foldername = 'data 2-27-21';
addpath utility
cd(foldername)
metadata_2_27_21;
cd ..

%resource sensor module pairs
RSmods0 = {'B170','G180';  'B170','G181';  'B170','R190';  'B170','R191';
    'B171','G180';  'B171','G181';  'B171','R190';  'B171','R191';
    'G180','R190';  'G180','R191';  'G181','R190';  'G181','R191'};
addpath utility

if 0
%-----------------------------------
[cellholder,basalstruct] = deal(cell(length(filenamecell),1));
load('C.mat')

%analyze data
for ii = 1:length(filenamecell)
    %init data and load data from file
    platedata = RSexpdata;
    platedata = loaddata(platedata,filenamecell{ii},sheetcell{ii});
    platedata.FPfields = {'BFP','GFP','RFP','YFP'};
    platedata.samplename = filenamecell{ii};
    %repackage data into individual fields for each experimental condition
    cellstruct0 = bulkdata2cells2(platedata,indstruct{ii});
    %replace outliers in OD data
    cellstruct0 = replaceoutliers(cellstruct0,'OD');
    %subtract the basal fluorescence based on background media vs time
    [cellstruct0, basalstruct] = subtractbasal7(cellstruct0,mediastruct,ODcontrolstruct);
    
    if exist('C','var')
        %subtract off fluorescence due to OD on BFP and GFP channels
        %[cellstruct0,COD] = ODFPcorrection(cellstruct0,ODFPstruct,maxstruct,COD);
        %subtract off fluorescence bleed
        cellstruct0 = applycolorcorrection(cellstruct0,C);
        %fit growthrate
        cellstruct0 = fitgrowthrate3(cellstruct0,false,true,4);
    end
    %correct for GFP fluorescence on neighboring wells?
    %neighborFP(cellstruct0,indstruct{ii},neighborsmeta)
    %take time derivative of raw fluorescent data
    cellstruct0 = diffcellfields(cellstruct0,{'BFP','GFP','RFP','YFP','OD'},'time');
    %combine batches
    cellholder{ii} = cellstruct0;
end

%combine batches into one
cellstruct = combinedilutions(cellholder);
%get correction factor for change in FP due to change in OD
%[cellstruct,COD] = ODFPcorrection(cellstruct,ODFPstruct,maxstruct,[],true,1);
%find fluorescence bleed between channels
%C = fluorescencebleed2(cellstruct,FPbleedstruct,maxstruct,false,[6,7,8,9]);
%check compensation
Cout = fluorescencebleed2(cellstruct,FPbleedstruct,maxstruct,false);
%make sure no offdiagonal entries are large
norm(Cout(~diag(ones(4,1))),'Inf')
%divide fluorescence and time derivative of fluorescence by OD
cellstruct = normalizeFP3(cellstruct,{'BFP','GFP','RFP','YFP',...
    'BFPdiff','GFPdiff','RFPdiff','YFPdiff'});

cd(foldername)
savefigpdf('GRstats',4,'png')
cd ..

end

%average over time intervals to get steady state
modulestruct = findss4(cellstruct,ssstruct,{'BFPdiffOD','GFPdiffOD',...
    'RFPdiffOD','YFPdiffOD'},@mean,false);
%combine submodules for each dilution level
modulestruct2 = combineinductions(modulestruct,inductionmeta);

%loop through pairs of resource sensors
for k = 1:size(RSmods0,1)
    
    RSmods = RSmods0(k,:)
    
    %turn on RS experiment with both of them together
    cellnames = fieldnames(modulemetadata);
    for jj = 1:length(cellnames)
        foo = modulemetadata.(cellnames{jj}).containingmods;
        if all(strcmpi(foo,RSmods))
            modulemetadata.(cellnames{jj}).isResourceSensor = true;
        else
            modulemetadata.(cellnames{jj}).isResourceSensor = false;
        end
    end
    %turn on desired resource sensor pair
    for ii = 1:length(RSmods)
        modulemetadata.(RSmods{ii}).isResourceSensor = true;
    end
    
    %add metadata to modules
    modulestruct3 = addmetadata(modulestruct2,modulemetadata);
    %calculate resource demand for resource sensor
    RSmodules = calcRS2(modulestruct3,true,7,6);
    %find Q and S for non-resource sensor modules
    [modulesout,RSmodules2] = calcQS2(modulestruct3,RSmodules,true,8,9,10,11,12,13,14,15);
    %combine modules together
    modulesout2 = combinemodstructs(RSmodules2,modulesout,modulestruct3);
    %predict outputs
    modsout3 = predictRS5(modulesout2,true,[16,17,18,19,20,21,22]);
    %plot growth rate and resource demand
    [modsout4,grfitobj] = plotGRvsQ(modulesout2,true,23);
    
    drawnow;
    
    %save figure outputs
    cd(foldername)
    savefigpdf(['06RSy_',RSmods{1},'_',RSmods{2}],6,'png')
    savefigpdf([RSmods{1},'_',RSmods{2},'_06RSy'],6,'png')
    savefigpdf([RSmods{1},'_',RSmods{2},'_06RSy'],6,'pdf')
    savefigpdf(['07RSQ_',RSmods{1},'_',RSmods{2}],7,'png')
    savefigpdf([RSmods{1},'_',RSmods{2},'_07RSQ'],7,'png')
    savefigpdf([RSmods{1},'_',RSmods{2},'_07RSQ'],7,'pdf')
    savefigpdf(['08mody+RSy_',RSmods{1},'_',RSmods{2}],8,'png')
    savefigpdf([RSmods{1},'_',RSmods{2},'_08mody+RSy'],8,'png')
    savefigpdf([RSmods{1},'_',RSmods{2},'_08mody+RSy'],8,'pdf')
    savefigpdf(['09RSy+mody_',RSmods{1},'_',RSmods{2}],9,'png')
    savefigpdf([RSmods{1},'_',RSmods{2},'_09RSy+mody'],9,'png')
    savefigpdf([RSmods{1},'_',RSmods{2},'_09RSy+mody'],9,'pdf')
    savefigpdf(['10modQ_',RSmods{1},'_',RSmods{2}],10,'png')
    savefigpdf([RSmods{1},'_',RSmods{2},'_10modQ'],10,'png')
    savefigpdf([RSmods{1},'_',RSmods{2},'_10modQ'],10,'pdf')
    savefigpdf(['11modS_',RSmods{1},'_',RSmods{2}],11,'png')
    savefigpdf([RSmods{1},'_',RSmods{2},'_11modS'],11,'png')
    savefigpdf([RSmods{1},'_',RSmods{2},'_11modS'],11,'pdf')
    savefigpdf(['12modyind+RSy_',RSmods{1},'_',RSmods{2}],12,'png')
    savefigpdf([RSmods{1},'_',RSmods{2},'_12modyind+RSy'],12,'png')
    savefigpdf([RSmods{1},'_',RSmods{2},'_12modyind+RSy'],12,'pdf')
    savefigpdf(['13RSy+modyind_',RSmods{1},'_',RSmods{2}],13,'png')
    savefigpdf([RSmods{1},'_',RSmods{2},'_13RSy+modyind'],13,'png')
    savefigpdf([RSmods{1},'_',RSmods{2},'_13RSy+modyind'],13,'pdf')
    savefigpdf(['14modQind_',RSmods{1},'_',RSmods{2}],14,'png')
    savefigpdf([RSmods{1},'_',RSmods{2},'_14modQind'],14,'png')
    savefigpdf([RSmods{1},'_',RSmods{2},'_14modQind'],14,'pdf')
    savefigpdf(['15modSind_',RSmods{1},'_',RSmods{2}],15,'png')
    savefigpdf([RSmods{1},'_',RSmods{2},'_15modSind'],15,'png')
    savefigpdf([RSmods{1},'_',RSmods{2},'_15modSind'],15,'pdf')
    savefigpdf(['16predB170_',RSmods{1},'_',RSmods{2}],16,'png')
    savefigpdf([RSmods{1},'_',RSmods{2},'_16predB170-all'],16,'png')
    savefigpdf([RSmods{1},'_',RSmods{2},'_16predB170-all'],16,'pdf')
    savefigpdf(['17predG180_',RSmods{1},'_',RSmods{2}],17,'png')
    savefigpdf([RSmods{1},'_',RSmods{2},'_17predG180-all'],17,'png')
    savefigpdf([RSmods{1},'_',RSmods{2},'_17predG180-all'],17,'pdf')
    savefigpdf(['18predB171_',RSmods{1},'_',RSmods{2}],18,'png')
    savefigpdf([RSmods{1},'_',RSmods{2},'_18predB171-all'],18,'png')
    savefigpdf([RSmods{1},'_',RSmods{2},'_18predB171-all'],18,'pdf')
    savefigpdf(['19predG181_',RSmods{1},'_',RSmods{2}],19,'png')
    savefigpdf([RSmods{1},'_',RSmods{2},'_19predG181-all'],19,'png')
    savefigpdf([RSmods{1},'_',RSmods{2},'_19predG181-all'],19,'pdf')
    savefigpdf(['20predR190_',RSmods{1},'_',RSmods{2}],20,'png')
    savefigpdf([RSmods{1},'_',RSmods{2},'_20predR190-all'],20,'png')
    savefigpdf([RSmods{1},'_',RSmods{2},'_20predR190-all'],20,'pdf')
    savefigpdf(['21predR191_',RSmods{1},'_',RSmods{2}],21,'png')
    savefigpdf([RSmods{1},'_',RSmods{2},'_21predR191-all'],21,'png')
    savefigpdf([RSmods{1},'_',RSmods{2},'_21predR191-all'],21,'pdf')
    savefigpdf(['22predY240_',RSmods{1},'_',RSmods{2}],22,'png')
    savefigpdf([RSmods{1},'_',RSmods{2},'_22predY240-4'],22,'png')
    savefigpdf([RSmods{1},'_',RSmods{2},'_22predY240-4'],22,'pdf')
    savefigpdf(['23GRvsQ_',RSmods{1},'_',RSmods{2}],23,'png')
    savefigpdf([RSmods{1},'_',RSmods{2},'_23GRvsQ'],23,'png')
    savefigpdf([RSmods{1},'_',RSmods{2},'_23GRvsQ'],23,'pdf')
    
    cd ..
end

toc
```
