## Supplementary material for "Predicting Composition of Genetic Circuits with Resource Competition: Demand and Sensitivity": Experimental data and Matlab code: makepred0.m

```
function makepred0(modulesout, RSmodules)

%GFP with BFP estimate from RFP measurements
[ypred(1),ypredstd(1)] = calcypred(modulesout.B.y,modulesout.B.S(2),RSmodules.G.Q,...
    modulesout.B.ystd,modulesout.B.Sstd(2),RSmodules.G.Qstd);

%GFP with RFP estimate from BFP measurements
[ypred(2),ypredstd(2)] = calcypred(modulesout.B.y,modulesout.B.S(1),RSmodules.R.Q,...
    modulesout.B.ystd,modulesout.B.Sstd(1),RSmodules.R.Qstd);

data = [modulesout.B.y, modulesout.B.perturby;
    NaN, ypred(:)'];
datastd = [modulesout.B.ystd, modulesout.B.perturbystd;
    NaN, ypredstd(:)'];

figure;
plotbarpredict(data,datastd)


function [ypred,ypredstd] = calcypred(yalone,S,Q,ystd,Sstd,Qstd)

%calculate perturbed module estimate
ypred = yalone./(1 - S*Q);

%calculate output standard error
dypdy = 1./(1-S*Q);
dypdS = yalone*Q./(1 - S*Q)^2;
dypdQ = yalone*S./(1 - S*Q)^2;
ypredstd = sqrt(dypdy^2*ystd^2 + dypdS^2*Sstd^2 + dypdQ^2*Qstd^2);


function plotbarpredict(data,datastd)

nbars = size(data,1);
ngroups = size(data,2);

groupwidth = min(0.8, nbars/(nbars + 1.5));
% Set the position of each error bar in the centre of the main bar
% Based on barweb.m by Bolu Ajiboye from MATLAB File Exchange
x = zeros(nbars,ngroups);
for ii = 1:nbars
    % Calculate center of each bar
    x(ii,:) = (1:ngroups) - groupwidth/2 + (2*ii-1) * groupwidth / (2*nbars);
end

bh = bar(data'); hold on;
set(bh,'edgecolor','none');
errorbar(x,data,datastd,'.k','linestyle','none','Linewidth',1.5);
```
