## Supplementary material for "Predicting Composition of Genetic Circuits with Resource Competition: Demand and Sensitivity": Experimental data and Matlab code: neighborFP.m

```
function outstruct = neighborFP(datastruct, locstruct, metastruct)
%correct for fluorescence bleed due to neighboring wells

%metastruct contains
if nargin < 3
    metastruct = struct;
    metastruct.GFP = {'B170','B171','BY172_0','BY173_0'}; %wells without GFP
end

FPfields = fieldnames(metastruct);
[p,p2] = deal(cell(length(FPfields),1));
cellnames = fieldnames(datastruct);
outstruct = datastruct;

for ii = 1:length(FPfields)
    FPfields{ii}
    count = 1;
    FPneighs = []; FP = [];
    %wells used as controls
    controlfields = intersect(cellnames,metastruct.(FPfields{ii}));
    for jj = 1:length(controlfields)
        %controlfields{jj}
        %find well locations of controls
        controlinds = locstruct.(controlfields{jj});
        %find well locations of neighboring wells
        neighinds = getneighbors(controlinds);
        for k = 1:length(neighinds)
            %look up wells
            neighlocs = structindfind(locstruct,neighinds{k});
            controlloc = structindfind(locstruct,controlinds(k,:));
            %get FP signal from control well
            FP(:,count) = datastruct.(controlfields{jj}).(FPfields{ii})(:,controlloc{2});
            n = size(FP,1);
            %get FP signals from neighboring wells and sum together
            z = [];
            for m = 1:size(neighlocs,1)
                %neighlocs(m,:)
                x = datastruct.(neighlocs{m,1}).(FPfields{ii})(:,neighlocs{m,2});
                if size(x,1) >= n
                    z = [z,x(1:n,:)];
                end
            end
            FPneighs(:,count) = sum(z,2);
            count = count+1;
        end
    end
    
    %figure; plot(FPneighs, FP)
    
    %use linear regression
    p{ii}  = polyfit(FPneighs,FP,1);
    p2{ii} = 1./polyfit(FP,FPneighs,1);
    slopevec = [p{ii}(1),p2{ii}(1)];
    [~,pind] = min(abs(slopevec));
    C(ii,:) = slopevec;
end
C

%apply correction to all wells on plate?


function neighinds = getneighbors(inds)
%get all neighbors within bounds of indicies. Inds is a n x 2 array with
%x-coordinates as the first column and y-coordinates as the 2nd column
xmin = 2;
ymin = 2;
xmax = 11;
ymax = 11;

neighinds = cell(size(inds,1),1);
for ii = 1:size(inds,1)
    ind1 = inds(ii,1);
    ind2 = inds(ii,2);
    %get neighboring indicies in the 4 cardinal directions
    neighinds{ii} = [ind1-1, ind2; ind1, ind2-1; ind1+1, ind2; ind1,ind2+1];
    %delete neighbor indicies that are out of bounds or matching an input
    xminind = neighinds{ii}(:,1) < xmin;
    yminind = neighinds{ii}(:,2) < ymin;
    xmaxind = neighinds{ii}(:,1) > xmax;
    ymaxind = neighinds{ii}(:,2) > ymax;
    inputinds = ismember(neighinds{ii},inds,'rows');
    badinds = xminind | yminind | xmaxind | ymaxind | inputinds;
    neighinds{ii}(badinds,:) = [];
end


function loc = structindfind(S,neighinds)
%find field names and indicies of neighinds matching in 2-layer struct S
%first column of the output is the fieldname, 2nd column is the index in the
%data corresponding to the column

names = fieldnames(S);
loc = cell(0);%size(neighinds,1),2);
count = 1;
for jj = 1:size(neighinds,1)
    for ii = 1:length(names)
        inds1 = find(S.(names{ii})(:,1) == neighinds(jj,1));
        inds2 = find(S.(names{ii})(:,2) == neighinds(jj,2));
        if ~isempty(inds1) && ~isempty(inds2)
            loc{count,1} = names{ii};              %cell name of the neighbor
            loc{count,2} = find(inds2 == inds1);   %index of the neighbor
        end
    end
    count = count + 1;
end


function neighinds = getneighborsold(inds)
%get all neighbors within bounds of indicies. Inds is a n x 2 array with
%x-coordinates as the first column and y-coordinates as the 2nd column
xmin = 2;
ymin = 2;
xmax = 11;
ymax = 11;

neighinds0 = [];
for ii = 1:size(inds,1)
    ind1 = inds(ii,1);
    ind2 = inds(ii,2);
    %get neighboring indicies
    neighinds0 = [neighinds0; ind1-1, ind2; ind1, ind2-1; ...
        ind1+1, ind2; ind1,ind2+1];
end
neighinds = unique(neighinds0,'rows','stable');

%remove rows out of range
xminind = neighinds(:,1) < xmin;
yminind = neighinds(:,2) < ymin;
xmaxind = neighinds(:,1) > xmax;
ymaxind = neighinds(:,2) > ymax;

%remove rows matching inputs
inputinds = ismember(neighinds,inds,'rows');
badinds = xminind | yminind | xmaxind | ymaxind | inputinds;
neighinds(badinds,:) = [];
```
