## Supplementary material for "Predicting Composition of Genetic Circuits with Resource Competition: Demand and Sensitivity": Experimental data and Matlab code: normalizeFP2.m

```
function outstruct = normalizeFP2(instruct, ploton)
%outstruct = normalizeFP2(instruct, ploton)
%normalize fluorescence data by OD or growthrate

if nargin < 2
    ploton = false;
end

names = fieldnames(instruct);
% dataprops = fieldnames(instruct.(names{1}));
% if ~any(contains(dataprops,{'ODfit','dODdt'}))
%     disp('fitting growth rate in normalizeFP2...')
%     instruct = fitgrowthrate2(instruct);
% end

%copy to output
outstruct = instruct;

%OD field
if isfield(instruct.(names{1}),'OD'); ODfld = 'OD';
elseif isfield(instruct.(names{1}),'OD600_600'); ODfld = 'OD600_600';
else; error('OD field does not match');
end

%normalize fluorescence data based on growth rate or by number of cells (OD)
names = fieldnames(instruct);
for k = 1:length(names)
    %loop through cell names
    props = fieldnames(instruct.(names{k}));
    for ii = 1:length(props)
        %loop through properties of each cell
        if any(contains(props{ii},'FP','IgnoreCase',true))
            %if fluorescence data, normalize by growth rate and OD
            %normalize FP per cell based on OD
            outstruct.(names{k}).([props{ii},'OD']) = instruct.(names{k}).(props{ii}) ./...
                instruct.(names{k}).(ODfld);
            %divide by ODfit
            if isfield(instruct.(names{k}),'ODfit')
                outstruct.(names{k}).([props{ii},'ODfit']) = instruct.(names{k}).(props{ii}) ./...
                    instruct.(names{k}).ODfit;
            end
            %multiply OD by gr2 to get get ODdil
            if isfield(instruct.(names{k}),'gr2')
                outstruct.(names{k}).([props{ii},'ODdil']) = ...
                    outstruct.(names{k}).([props{ii},'OD']) .*instruct.(names{k}).gr2;
            end
            %normalize based on growth rate
            %y = FP/(OD * d/dt ln(a*OD)) = a*FP/(d(OD)/dt)
            %outstruct.(names{k}).([props{ii},'mu']) = instruct.(names{k}).(props{ii}) ./...
            %    instruct.(names{k}).dODdt;
            %y = (FP/OD)*growth rate = FP*(dODdt/OD)/(OD)
            %outstruct.(names{k}).([props{ii},'mu2']) = instruct.(names{k}).(props{ii}) .*...
            %    instruct.(names{k}).dODdt./(instruct.(names{k}).OD.^2);
        end
    end
end


if ploton
    tstart = 0.2e4;
    %plot normalized data
    if any(strcmp(names,'G')) && any(strcmp(names,'GR')) && ...
            any(strcmp(names,'GY')) && any(strcmp(names,'R')) && ...
            any(strcmp(names,'RY')) && any(strcmp(names,'Y'))
        %plot timecourses of normalized fluorescent proteins
        figure;
        subplot(131); hold on
        plot(outstruct.G.time,outstruct.G.GFPmu);
        plot(outstruct.GR.time,outstruct.GR.GFPmu,'--');
        plot(outstruct.GY.time,outstruct.GY.GFPmu,'-.');
        legend('G','G','G','GR','GR','GR','GY','GY','GY')
        %set(gca,'yscale','log')
        ylabel('GFP/(dOD/dt)')
        xlabel('time (s)')
        xlim([tstart,max(outstruct.Y.time)]);
        
        subplot(132); hold on
        plot(outstruct.M9.time, outstruct.R.RFPmu);
        plot(outstruct.GR.time,outstruct.GR.RFPmu,'--');
        plot(outstruct.RY.time,outstruct.RY.RFPmu,'-.');
        legend('R','R','R','RG','RG','RG','RY','RY','RY')
        %set(gca,'yscale','log')
        ylabel('RFP/(dOD/dt)')
        xlabel('time (s)')
        xlim([tstart,max(outstruct.Y.time)]);
        
        subplot(133); hold on
        plot(outstruct.Y.time,outstruct.Y.YFPmu);
        plot(outstruct.GY.time,outstruct.GY.YFPmu,'--');
        plot(outstruct.RY.time,outstruct.RY.YFPmu,'-.');
        %set(gca,'yscale','log')
        legend('Y','Y','Y','YG','YG','YG','YR','YR','YR')
        ylabel('YFP/(dOD/dt)')
        xlabel('time (s)')
        xlim([tstart,max(outstruct.Y.time)]);
        
        %plot timecourses of normalized fluorescent proteins on OD
        figure;
        subplot(131); hold on
        plot(outstruct.G.time,outstruct.G.GFPOD);
        plot(outstruct.GR.time,outstruct.GR.GFPOD,'--');
        plot(outstruct.GY.time,outstruct.GY.GFPOD,'-.');
        legend('G','G','G','GR','GR','GR','GY','GY','GY')
        %set(gca,'yscale','log')
        ylabel('GFP/OD')
        xlabel('time (s)')
        xlim([tstart,max(outstruct.Y.time)]);
        
        subplot(132); hold on
        plot(outstruct.M9.time, outstruct.R.RFPOD);
        plot(outstruct.GR.time,outstruct.GR.RFPOD,'--');
        plot(outstruct.RY.time,outstruct.RY.RFPOD,'-.');
        legend('R','R','R','RG','RG','RG','RY','RY','RY')
        %set(gca,'yscale','log')
        ylabel('RFP/OD')
        xlabel('time (s)')
        xlim([tstart,max(outstruct.Y.time)]);
        
        subplot(133); hold on
        plot(outstruct.Y.time,outstruct.Y.YFPOD);
        plot(outstruct.GY.time,outstruct.GY.YFPOD,'--');
        plot(outstruct.RY.time,outstruct.RY.YFPOD,'-.');
        %set(gca,'yscale','log')
        legend('Y','Y','Y','YG','YG','YG','YR','YR','YR')
        ylabel('YFP/OD')
        xlabel('time (s)')
        xlim([tstart,max(outstruct.Y.time)]);
    else
        %plot normalized fluorescence fields
        plotsubfield(outstruct,'GFPmu')
        plotsubfield(outstruct,'RFPmu')
        plotsubfield(outstruct,'YFPmu')
        plotsubfield(outstruct,'GFPOD')
        plotsubfield(outstruct,'RFPOD')
        plotsubfield(outstruct,'YFPOD')
    end
end
```
