## Supplementary material for "Predicting Composition of Genetic Circuits with Resource Competition: Demand and Sensitivity": Experimental data and Matlab code: normalizeFP.m

```
function [celldataout, cnvstct] = normalizeFP(celldata, ploton)
%[celldataout, cnvstct] = normalizeFP(celldata, ploton)
%normalize fluorescence data by OD or growthrate and compensate for
%fluorescence leakage

if nargin < 2
    ploton = false;
end

celldataout = celldata;
cellnames = fieldnames(celldata);
dataprops = fieldnames(celldata.(cellnames{1}));
if ~any(strcmp(dataprops,'GFP')) || ~any(contains(dataprops,'RFP')) ||...
        ~any(contains(dataprops,'YFP'))
    error('input struct does not have the required fields')
end

%compensate for spectrial leakage in GFP/YFP signal
if any(strcmp(cellnames,'G')) && any(strcmp(cellnames,'Y')) && any(strcmp(cellnames,'R'))
    disp('correcting for fluorescence leakage...')
    GYcnvt = mean(celldata.G.YFP./celldata.G.GFP,2);     %green in the yellow channel
    GRcnvt = mean(celldata.G.RFP./celldata.G.GFP,2);     %green in the red channel
    YGcnvt = mean(celldata.Y.GFP./celldata.Y.YFP,2);     %yellow in the green channel
    YRcnvt = mean(celldata.Y.RFP./celldata.Y.YFP,2);     %yellow in the red channel
    RGcnvt = mean(celldata.R.GFP./celldata.R.RFP,2);     %red in the green channel
    RYcnvt = mean(celldata.R.YFP./celldata.R.RFP,2);     %red in the yellow channel
    %output conversions to a struct
    cnvstct = struct();
    cnvstct.GYconvert = mean(GYcnvt(end-20:end));
    cnvstct.GRconvert = mean(GRcnvt(end-20:end));
    cnvstct.YGconvert = mean(YGcnvt(end-20:end));
    cnvstct.YRconvert = mean(YRcnvt(end-20:end));
    cnvstct.RGconvert = mean(RGcnvt(end-20:end));
    cnvstct.RYconvert = mean(RYcnvt(end-20:end));
    %cnvstct = filtplatedata(cnvstct,0.2);
    %compensate for fluorescence leakage between GFP and YFP assuming a linear
    %dependence of leakage and normal fluorescence
    celldataout.GY.YFP = (celldataout.GY.YFP - cnvstct.YGconvert.*celldataout.GY.YFP)./...
        (1 - cnvstct.GYconvert.*cnvstct.YGconvert);
    celldataout.GY.GFP = (celldataout.GY.GFP - cnvstct.GYconvert.*celldataout.GY.GFP)./...
        (1 - cnvstct.GYconvert.*cnvstct.YGconvert);
else
    cnvstct = struct();
end

%normalize fluorescence data based on growth rate or by number of cells (OD)
cellnames = fieldnames(celldataout);
for k = 1:length(cellnames)
    %normalize based on growth rate
    %y = FP/(OD * d/dt ln(a*OD)) = a*FP/(d(OD)/dt)
    celldataout.(cellnames{k}).GFPmu = celldataout.(cellnames{k}).GFP./...
        celldataout.(cellnames{k}).dODdt;
    celldataout.(cellnames{k}).RFPmu = celldataout.(cellnames{k}).RFP./...
        celldataout.(cellnames{k}).dODdt;
    celldataout.(cellnames{k}).YFPmu = celldataout.(cellnames{k}).YFP./...
        celldataout.(cellnames{k}).dODdt;
    %normalize FP per cell based on OD
    celldataout.(cellnames{k}).GFPOD = celldataout.(cellnames{k}).GFP./...
        celldataout.(cellnames{k}).ODfit;
    celldataout.(cellnames{k}).RFPOD = celldataout.(cellnames{k}).RFP./...
        celldataout.(cellnames{k}).ODfit;
    celldataout.(cellnames{k}).YFPOD = celldataout.(cellnames{k}).YFP./...
        celldataout.(cellnames{k}).ODfit;
end

if ploton
    tstart = 0.2e4;
    if any(strcmp(cellnames,'G')) && any(strcmp(cellnames,'Y')) && any(strcmp(cellnames,'R'))
        %plot spectral leakage conversion factors
        figure;
        plot(celldataout.M9.time,[GYcnvt,GRcnvt,YGcnvt,YRcnvt,RGcnvt,RYcnvt])
        legend({'Y in G channel','R in G channel','G in Y channel','R in Y channel',...
            'G in R channel','Y in R channel'},'Location','best')
        ylabel('spectral conversion')
        xlabel('time (s)')
        xlim([tstart,max(celldataout.M9.time)])
    end
    
    %plot normalize data
    if any(strcmp(cellnames,'G')) && any(strcmp(cellnames,'GR')) && ...
            any(strcmp(cellnames,'GY')) && any(strcmp(cellnames,'R')) && ...
            any(strcmp(cellnames,'RY')) && any(strcmp(cellnames,'Y'))
        %plot timecourses of normalized fluorescent proteins
        figure;
        subplot(131); hold on
        plot(celldataout.G.time,celldataout.G.GFPmu);
        plot(celldataout.GR.time,celldataout.GR.GFPmu,'--');
        plot(celldataout.GY.time,celldataout.GY.GFPmu,'-.');
        legend('G','G','G','GR','GR','GR','GY','GY','GY')
        %set(gca,'yscale','log')
        ylabel('GFP/(dOD/dt)')
        xlabel('time (s)')
        xlim([tstart,max(celldataout.Y.time)]);
        
        subplot(132); hold on
        plot(celldataout.M9.time, celldataout.R.RFPmu);
        plot(celldataout.GR.time,celldataout.GR.RFPmu,'--');
        plot(celldataout.RY.time,celldataout.RY.RFPmu,'-.');
        legend('R','R','R','RG','RG','RG','RY','RY','RY')
        %set(gca,'yscale','log')
        ylabel('RFP/(dOD/dt)')
        xlabel('time (s)')
        xlim([tstart,max(celldataout.Y.time)]);
        
        subplot(133); hold on
        plot(celldataout.Y.time,celldataout.Y.YFPmu);
        plot(celldataout.GY.time,celldataout.GY.YFPmu,'--');
        plot(celldataout.RY.time,celldataout.RY.YFPmu,'-.');
        %set(gca,'yscale','log')
        legend('Y','Y','Y','YG','YG','YG','YR','YR','YR')
        ylabel('YFP/(dOD/dt)')
        xlabel('time (s)')
        xlim([tstart,max(celldataout.Y.time)]);
        
        %plot timecourses of normalized fluorescent proteins on OD
        figure;
        subplot(131); hold on
        plot(celldataout.G.time,celldataout.G.GFPOD);
        plot(celldataout.GR.time,celldataout.GR.GFPOD,'--');
        plot(celldataout.GY.time,celldataout.GY.GFPOD,'-.');
        legend('G','G','G','GR','GR','GR','GY','GY','GY')
        %set(gca,'yscale','log')
        ylabel('GFP/OD')
        xlabel('time (s)')
        xlim([tstart,max(celldataout.Y.time)]);
        
        subplot(132); hold on
        plot(celldataout.M9.time, celldataout.R.RFPOD);
        plot(celldataout.GR.time,celldataout.GR.RFPOD,'--');
        plot(celldataout.RY.time,celldataout.RY.RFPOD,'-.');
        legend('R','R','R','RG','RG','RG','RY','RY','RY')
        %set(gca,'yscale','log')
        ylabel('RFP/OD')
        xlabel('time (s)')
        xlim([tstart,max(celldataout.Y.time)]);
        
        subplot(133); hold on
        plot(celldataout.Y.time,celldataout.Y.YFPOD);
        plot(celldataout.GY.time,celldataout.GY.YFPOD,'--');
        plot(celldataout.RY.time,celldataout.RY.YFPOD,'-.');
        %set(gca,'yscale','log')
        legend('Y','Y','Y','YG','YG','YG','YR','YR','YR')
        ylabel('YFP/OD')
        xlabel('time (s)')
        xlim([tstart,max(celldataout.Y.time)]);
    else
        %plot normalized fluorescence fields
        plotsubfield(celldataout,'GFPmu')
        plotsubfield(celldataout,'RFPmu')
        plotsubfield(celldataout,'YFPmu')
        plotsubfield(celldataout,'GFPOD')
        plotsubfield(celldataout,'RFPOD')
        plotsubfield(celldataout,'YFPOD')
    end
end
```
