## Supplementary material for "Predicting Composition of Genetic Circuits with Resource Competition: Demand and Sensitivity": Experimental data and Matlab code: platewrapper.m

```
function cellstructout = platewrapper(filenamecell,indstruct,celldatafields,nonefield)
%cellstruct2 = platewrapper(filenamecell,indstruct,nonefield,celldatafields)

wn = 0.5;               %low pass filter cutoff frequency
plotgrowthon = true;    %plot bar graph of growth rates
ODfitploton = false;    %plot fit curves for OD

if nargin < 1
    filenamecell = {'Exp1_9-26-2019.mat','Exp2_9-26-2019.mat',...
   'Exp3_9-26-2019.mat','Exp4_9-26-2019.mat'};
end
if nargin < 2
    indstruct{1}.atc0ahl0 = [1,1; 1,2; 1,3; 1,7; 1,8; 1,9];
    %aTc = 0; titrate AHL \in [0,0.1,1,3,10,30,100,300]
    indstruct{1}.atc0ahl1 = [2,1; 2,2; 2,3];
    indstruct{1}.atc0ahl2 = [3,1; 3,2; 3,3];
    indstruct{1}.atc0ahl3 = [4,1; 4,2; 4,3];
    indstruct{1}.atc0ahl4 = [5,1; 5,2; 5,3];
    indstruct{1}.atc0ahl5 = [6,1; 6,2; 6,3];
    indstruct{1}.atc0ahl6 = [7,1; 7,2; 7,3];
    indstruct{1}.atc0ahl7 = [8,1; 8,2; 8,3];
    
    %aTc = 10; titrate AHL \in [0,0.1,1,3,10,30,100,300]
    indstruct{1}.atc10ahl1 = [2,4; 2,5; 2,6];
    indstruct{1}.atc10ahl2 = [3,4; 3,5; 3,6];
    indstruct{1}.atc10ahl3 = [4,4; 4,5; 4,6];
    indstruct{1}.atc10ahl4 = [5,4; 5,5; 5,6];
    indstruct{1}.atc10ahl5 = [6,4; 6,5; 6,6];
    indstruct{1}.atc10ahl6 = [7,4; 7,5; 7,6];
    indstruct{1}.atc10ahl7 = [8,4; 8,5; 8,6];
    
    %AHL = 0; titrate aTc \in [0,0.01,0.1,0.3,1,3,10,30]
    indstruct{1}.ahl0atc1 = [2,7; 2,8; 2,9];
    indstruct{1}.ahl0atc2 = [3,7; 3,8; 3,9];
    indstruct{1}.ahl0atc3 = [4,7; 4,8; 4,9];
    indstruct{1}.ahl0atc4 = [5,7; 5,8; 5,9];
    indstruct{1}.ahl0atc5 = [6,7; 6,8; 6,9];
    indstruct{1}.ahl0atc6 = [7,7; 7,8; 7,9];
    indstruct{1}.ahl0atc7 = [8,7; 8,8; 8,9];
    
    %AHL = 100; titrate aTc \in [0,0.01,0.1,0.3,1,3,10,30]
    indstruct{1}.ahl100atc1 = [2,10; 2,11; 2,12];
    indstruct{1}.ahl100atc2 = [3,10; 3,11; 3,12];
    indstruct{1}.ahl100atc3 = [4,10; 4,11; 4,12];
    indstruct{1}.ahl100atc4 = [5,10; 5,11; 5,12];
    indstruct{1}.ahl100atc5 = [6,10; 6,11; 6,12];
    indstruct{1}.ahl100atc6 = [7,10; 7,11; 7,12];
    indstruct{1}.ahl100atc7 = [8,10; 8,11; 8,12];
end
if nargin < 3
    celldatafields = {'GFP','RFP','OD','time'};
end
if nargin < 4
    nonefield = '';
end

%init
[cellstruct,basalstruct] = deal(cell(length(filenamecell),1));
timeprev = 0;

%loop through each dilution
for ii = 1:length(filenamecell)
    
    %load data
    if exist(filenamecell{ii},'file')
        load(filenamecell{ii})
    else
        platedata = importplate([filenamecell{ii}(1:end-4),'.txt']);
    end
    
    %add time
    [platedata,timeprev] = addtime(platedata,timeprev);
    
    %rename fields
    datafields = fieldnames(platedata);
    for jj = 1:length(celldatafields)
        fieldinds = contains(datafields,celldatafields{jj});
        if nnz(fieldinds) > 1
            disp(datafields)
            error(['multiple fields match with ',celldatafields{jj}])
        end
        if nnz(fieldinds) < 1
            disp(datafields)
            error(['no match for field with ',celldatafields{jj}])
        end
        platedata.(celldatafields{jj}) = platedata.(datafields{fieldinds});
    end
    
    %filter and format into substructs
    platedata = filtplatedata2(platedata,wn,'low');
    cellstruct{ii} = bulkdata2cells(platedata,indstruct{ii},celldatafields);
    
    %subtract background fluorescence and OD levels
    if ~isempty(nonefield)
        [cellstruct{ii}, basalstruct{ii}] = subtractbasal3(cellstruct{ii},...
            nonefield,nonefield);
    else
        [cellstruct{ii}, basalstruct{ii}] = subtractbasalnomedia(cellstruct{ii});
    end
    
    %find smoothed growthrate for cells
    cellstruct{ii} = fitgrowthrate2(cellstruct{ii},'logisticfit','OD',plotgrowthon,ODfitploton,ii);
end

cellstructout = combinecellrepeats(cellstruct);

% %correct for fluorescence bleed between channels
% cellstruct2 = correctFP(cellstruct2,{'G','R','Y'},{'GFP','RFP','YFP'});

%divide fluorescence by OD and growthrate
cellstructout = normalizeFP2(cellstructout);
```
