## Supplementary material for "Predicting Composition of Genetic Circuits with Resource Competition: Demand and Sensitivity": Experimental data and Matlab code: plot3subfield.m

```
function [figh,axh] = plot3subfield(cellstruct,xfield,yfield,zfield,figh,cellfields)
%[figh,axh] = plotsubfield2(cellstruct, xfield, yfield,figh, cellfields)
%plot one subfield for each field in a struct. datafield must match to a
%valid field for all substructs contined in cellstruct

%input verification
if nargin < 4 || isempty(figh)
    figh = figure;
end

fntsze = 10;

%get fields for each cell type
celltypes = fieldnames(cellstruct);
if nargin > 5 && ~isempty(cellfields)
    cellnames = intersect(cellfields,celltypes,'stable');
else; cellnames = celltypes;
end

n = length(cellnames);                  %number of subplots (fields) to make
m = ceil(9/16*sqrt(n));                 %number of rows of subplots

%plot each struct's subfield on one plot
figure(figh);
axh = cell(n,1);
for jj = 1:n
    %make subplot
    axh{jj} = subplot(m,ceil(n/m),jj);
    %keep hold on if already set
    holdon = ishold;
    if holdon; hold on; end
    %plot
    xdata = cellstruct.(cellnames{jj}).(xfield);
    ydata = cellstruct.(cellnames{jj}).(yfield);
    zdata = cellstruct.(cellnames{jj}).(zfield);
    plot3(xdata,ydata,zdata,'linewidth',2);
    
    title(strrep(cellnames{jj},'_','-'))
    %put ylabels only on farthest left plots
    %if mod(jj,ceil(n/m)) == 1; ylabel(strrep(yfield,'_',' ')); end
    %put xlabels only on bottom plots
    %if n - jj < ceil(n/m); xlabel(strrep(xfield,'_',' ')); end
    xlabel(strrep(xfield,'_',' '));
    ylabel(strrep(yfield,'_',' '));
    zlabel(strrep(zfield,'_',' '));
    set(gca,'fontsize',fntsze)
    hold off
end
```
