## Supplementary material for "Predicting Composition of Genetic Circuits with Resource Competition: Demand and Sensitivity": Experimental data and Matlab code: plot3testing.m

```
figure(6);
plot3(cellstruct.G180.BFP,cellstruct.G180.GFP,cellstruct.G180.time); hold on;
plot3(cellstruct.G181.BFP,cellstruct.G181.GFP,cellstruct.G181.time)
title('G180, G181')
xlabel('BFP'); ylabel('GFP'); zlabel('time')
hold off;

figure(7); clf;
plot3(cellstruct.R190.time,cellstruct.R190.GFP,cellstruct.R190.RFP); hold on;
plot3(cellstruct.R191.time,cellstruct.R191.GFP,cellstruct.R191.RFP)
title('R190, R191')
xlabel('time'); ylabel('GFP'); zlabel('RFP')
hold off;

figure(8); clf;
plot3(cellstruct.B170.BFP,cellstruct.B170.GFP,cellstruct.B170.time); hold on;
plot3(cellstruct.B171.BFP,cellstruct.B171.GFP,cellstruct.B171.time)
title({'B170, B171'})
xlabel('BFP'); ylabel('GFP'); zlabel('time')
hold off;

figure(9); clf;
plot3(cellstruct.B171.BFP,cellstruct.B171.GFP,cellstruct.R191.time); hold on;
plot3(cellstruct.G181.BFP,cellstruct.G181.GFP,cellstruct.R191.time);
plot3(cellstruct.R191.BFP,cellstruct.R191.GFP,cellstruct.R191.time);
xlabel('BFP'); ylabel('GFP'); zlabel('time')

%leakage of GFP onto BFP and RFP due to neighboring wells
time = cellstruct.R191.time;
figure(10); clf;
plot3(cellstruct.G181.GFP,cellstruct.B171.GFP,cellstruct.B171.BFP,'linewidth',1.2); hold on
plot3(cellstruct.G181.GFP,cellstruct.R191.GFP,cellstruct.R191.RFP,'linewidth',1.2)
xlabel('G181 GFP'); ylabel('B171/R191 GFP'); zlabel('BFP/RFP')
set(gca,'fontsize',14)

inds = all(~isnan(cellstruct.G181.GFP),2);

p = polyfit(cellstruct.G181.GFP(inds,:),[cellstruct.B171.GFP(inds,:),cellstruct.R191.GFP(inds,:)],1)

%leakage of BFP onto GFP and RFP due to neighboring wells
figure(11); clf;
plot3(cellstruct.B170.BFP,cellstruct.G180.BFP,time); hold on
plot3(cellstruct.B170.BFP,cellstruct.R190.BFP,time)
xlabel('B170 BFP'); ylabel('G180/R190 BFP'); zlabel('time')

%leakage of RFP onto G180 and M9 due to neighboring wells
figure(12); clf;
plot3(cellstruct.R190.RFP,cellstruct.M9.RFP(:,1:3),time); hold on
plot3(cellstruct.R190.RFP,cellstruct.G180.RFP,time)
xlabel('R191 RFP'); ylabel('M9/G181 RFP'); zlabel('time')

figure(13); clf;
plot3(cellstruct.M9.BFP,cellstruct.M9.GFP,cellstruct.M9.RFP); hold on;
%plot3(cellstruct.R190.BFP,cellstruct.R190.GFP,cellstruct.R190.RFP);
%plot3(cellstruct.R191.BFP,cellstruct.R191.GFP,cellstruct.R191.RFP);
%plot3(cellstruct.BR210.BFP,cellstruct.BR210.GFP,cellstruct.BR210.RFP);
%plot3(cellstruct.BR211.BFP,cellstruct.BR211.GFP,cellstruct.BR211.RFP);
```
