## Supplementary material for "Predicting Composition of Genetic Circuits with Resource Competition: Demand and Sensitivity": Experimental data and Matlab code: plotcellstruct.m

```
function plotcellstruct(cellstruct,datatag,xfield)
%plotcellstruct(cellstruct,datatag,xfield)
%plot timeseries for each dataset contained labeled by the fields in 
%cellstruct
%cellstruct cannot be a nested struct (it is not recursive)
%datatag is a string for the title of the graph
%xfield is a string corresponding to the name of the field used for the
%independent variable on the plot

if nargin < 3
    xfield = 'time';
    if nargin < 2
        datatag = '';
    end
end

cellfields = fieldnames(cellstruct);
cellfields2 = cellfields(~contains(cellfields,xfield));
n = sum(~contains(cellfields,xfield));  %number of fields to plot not continaing xfield

%missing explicit time vector
if ~any(strcmp(cellfields,xfield))
    %if time is in the name of some fields
    if any(contains(cellfields,xfield))
        %average all timeseries at each point
        cellxfield = cellfields(contains(cellfields,xfield));
        z = zeros(length(cellstruct.(cellxfield{1})),length(cellxfield));
        for jj = 1:length(cellxfield)
             z(:,jj) = cellstruct.(cellxfield{jj})(:);
             cellstruct.(xfield) = mean(z,2);
        end
    else
        %if no time dependence, just plot data points
        cellstruct.(xfield) = 1:length(cellstruct.(cellfields2{1}));
    end
end

figure;
for ii = 1:n
    subplot(2,ceil(n/2),ii);
    if contains(cellfields2{ii},'OD')
        plot(cellstruct.(xfield),cellstruct.(cellfields2{ii}))
    else
        plot(cellstruct.(xfield),cellstruct.(cellfields2{ii}))
    end
    if strcmp(xfield,'time')
        xlabel([xfield,' (hrs)'])
    else
        xlabel(xfield)
    end
    ylabel(cellfields2{ii})
    title(datatag)
    xlim([min(cellstruct.time),max(cellstruct.time)])
end
```
