## Supplementary material for "Predicting Composition of Genetic Circuits with Resource Competition: Demand and Sensitivity": Experimental data and Matlab code: plotFPandOD.m

```
function plotFPandOD(cellstruct,datatag,tmin,tmax)
%plotFPandOD(cellstruct,datatag,tmin,tmax)
%plot fluorescence fields vs time along with growthrate vs time

celltype = fieldnames(cellstruct);
%check if recursion is necessary
if isstruct(cellstruct.(celltype{1}))
    for m = 1:length(celltype)
        if nargin == 4
            plotFPandOD(cellstruct.(celltype{m}),celltype{m},tmin,tmax)
        elseif nargin == 3
            plotFPandOD(cellstruct.(celltype{m}),celltype{m},tmin)
        elseif nargin == 2
            plotFPandOD(cellstruct.(celltype{m}),celltype{m})
        elseif nargin == 1
            plotFPandOD(cellstruct.(celltype{m}),celltype{m})
        end
    end
else
    %check inputs
    if nargin < 4
        tmax = max(cellstruct.time);
        if nargin < 3
            tmin = 0;
            if nargin < 2
                datatag = '';
            end
        end
    end
    
    n = sum(contains(celltype,'FP'));
    %plot fluorescence
    fig = figure;
    jj = 1;
    for ii = 1:length(celltype)
        if any(contains(celltype{ii},'FP'))
            subplot(ceil(n/3)+1,3,jj)
            plot(cellstruct.time,cellstruct.(celltype{ii}))
            ylabel(celltype{ii})
            xlim([min([tmin,tmax]),max([tmin,tmax])])
            if jj < 4
                title(datatag)
            end
            jj = jj + 1;
        end
    end
    
    %plot OD or growthrate
    for k = 1:3
        subplot(ceil(n/3)+1,3,n+k)
        semilogy(cellstruct.time,cellstruct.dODdt)
        xlim([min([tmin,tmax]),max([tmin,tmax])])
        ylim([min(min(cellstruct.dODdt)),max(max(cellstruct.dODdt))])
        ylabel('dOD/dt')
        xlabel('time (hrs)')
    end
    %stretch vertically
    pos = get(fig,'Position');
    pos(2) = 50;              %bottom attribute
    pos(4) = 600;             %height attribute
    set(fig,'Position',pos)
end
```
