## Supplementary material for "Predicting Composition of Genetic Circuits with Resource Competition: Demand and Sensitivity": Experimental data and Matlab code: plotplatestitch.m

```
function plotplatestitch(ycell, indscell, xcell, axh)
%plotplatestitch(ycell,indscell,xcell,axh)
%plot multiple platereader fluorescence experiments on the same figure.
%YCELL is a cell array containing flourescence data
%INDS is a cell array of n x 2 integers where row ii contains the 
%corresponding indicies for the desired entry in FP{ii}
%XCELL is a cell array of time vectors. The cell array must have the same
%size as FP and each cell must contain a vector the same size as the 3rd
%dimension of each cell of FP.
%AXH is the axis handle.

if nargin < 4 || isempty(axh)
    axh = gca; cla;
end

axes(axh); cla;
for ii = 1:length(xcell)
    set(gca,'ColorOrderIndex',1);
    hold on;
    for jj = 1:length(indscell)
        plot(xcell{ii},squeeze(ycell{ii}(indscell{jj}(ii,1),indscell{jj}(ii,2),:)))
        hold on;
    end
end
```
