## Supplementary material for "Predicting Composition of Genetic Circuits with Resource Competition: Demand and Sensitivity": Experimental data and Matlab code: plotsubfield.m

```
function plotsubfield(cellstruct, xfield, yfield, figh, tmin, tmax, nonefield)
%plotsubfield(cellstruct, xfield, yfield, tmin, tmax)
%plot one subfield for each field in a struct. datafield must match to a
%valid field for all substructs contined in cellstruct

%input verification
if nargin < 7 || isempty(nonefield)
    nonefield = 'M9';
end
if nargin < 6 || isempty(tmin)
    if isfield(cellstruct,nonefield)
        tmin = min(cellstruct.(nonefield).(xfield));
    else
        tmin = [];
    end
end
if nargin < 5 || isempty(tmax)
    if isfield(cellstruct,nonefield)
        tmax = max(cellstruct.(nonefield).(xfield));
    else
        tmax = [];
    end
end
if nargin < 4 || isempty(figh)
    figh = figure;
end
if nargin < 2 || isempty(xfield)
    xfield = 'time';
end
if nargin < 3 || isempty(yfield)
    yfield = 'OD';
end
if isfield(cellstruct,nonefield)
    if tmin < min(cellstruct.(nonefield).(xfield))
        tmin = min(cellstruct.(nonefield).(xfield));
    end
    if tmax > max(cellstruct.(nonefield).(xfield))
        tmax = max(cellstruct.(nonefield).(xfield));
    end
end

fntsze = 12;

%get fields for each cell type
celltypes = fieldnames(cellstruct);
n = length(celltypes);                  %number of subplots (fields) to make
m = ceil(9/16*sqrt(n));                 %number of rows of subplots

%input error checking
if ~ischar(yfield)
    error('datafield not string')
end

%check that each cell type has a time field
cellprops = cell(n,1);
for ii = 1:n
    cellprops{ii} = fieldnames(cellstruct.(celltypes{ii}));
    %missing explicit time vector
    if ~any(strcmp(cellprops{ii},xfield))
        %if time is in the name of some fields
        if any(contains(cellprops{ii},xfield))
            %average all timeseries at each point
            cellxfield = cellprops{ii}(contains(cellprops{ii},xfield));
            cellprops2 = zeros(length(cellstruct.(cellxfield{ii})),length(cellxfield));
            for k = 1:length(cellxfield)
                cellprops2(:,k) = cellstruct.(celltypes{ii}).(cellxfield{k})(:);
                cellstruct.(celltypes{ii}).(xfield) = mean(cellprops2,2);
            end
        else
            %if no time dependence, just plot data points
            cellstruct.(celltypes{ii}).(xfield) = 1:length(cellstruct.(celltypes{ii}));
        end
    end
end

%plot each struct's subfield on one plot
figure(figh);
for jj = 1:n
    subplot(m,ceil(n/m),jj);
    if ishold; hold on; end
    plot(cellstruct.(celltypes{jj}).(xfield), cellstruct.(celltypes{jj}).(yfield),'linewidth',2)
    title(strrep(celltypes{jj},'_','-'))
    if mod(jj,ceil(n/m)) == 1
        ylabel(strrep(yfield,'_',' '))
    end
    if n - jj < ceil(n/m)
        xlabel(strrep(xfield,'_',' '))
    end
    if contains(xfield,'time','ignorecase',true)
        if ~isempty(tmin) && ~isempty(tmax)
            xlim([tmin,tmax])
        end
    end
    set(gca,'fontsize',fntsze)
    hold off
end
```
