## Supplementary material for "Predicting Composition of Genetic Circuits with Resource Competition: Demand and Sensitivity": Experimental data and Matlab code: predictRS2.m

```
function outstruct = predictRS2(datastruct,FPstruct,RSDstruct,ploton,figh)
%predict outputs

if nargin < 4 || isempty(ploton)
    ploton = true;
end
if ploton && nargin < 5
    figh = 4;
end

%init
outstruct = datastruct;
FPnames = fieldnames(FPstruct);
modsalone = fieldnames(RSDstruct);
names = cell(length(FPnames),1);
[datamat,datamatstd] = deal(cell(length(modsalone),1));
mask = 1:length(modsalone);

%loop through perturbed modules and make predictions
for ii = 1:length(FPnames)
    %get names of modules alone and corresponding measurement fields
    alone1 = FPstruct.(FPnames{ii}).alone1;
    alone2 = FPstruct.(FPnames{ii}).alone2;
    FPfield1 = FPstruct.(FPnames{ii}).FPfield1;
    FPfield2 = FPstruct.(FPnames{ii}).FPfield2;
    
    %look in RSDstruct for corresponding resource demands
    cellpairs1 = fieldnames(RSDstruct.(alone1));
    cellpairs2 = fieldnames(RSDstruct.(alone2));
    stdfields1 = cellpairs1(contains(cellpairs1,'std'));
    stdfields2 = cellpairs2(contains(cellpairs2,'std'));
    
    %exclude current measurement, least squares, and std fields from prediction
    predfield1 = setxor(cellpairs1,['ls';FPnames{ii};stdfields1(:)]);
    predfield2 = setxor(cellpairs2,['ls';FPnames{ii};stdfields2(:)]);
    
    %get correpsonding resource demand
    [RSD1,RSD1std] = deal(zeros(length(predfield1),1));
    for jj = 1:length(predfield1)
        RSD1(jj) = RSDstruct.(alone1).(predfield1{jj});
        RSD1std(jj) = RSDstruct.(alone1).([predfield1{jj},'std']);
    end
    %get correpsonding resource demand
    [RSD2,RSD2std] = deal(zeros(length(predfield2),1));
    for k = 1:length(predfield2)
        RSD2(k) = RSDstruct.(alone2).(predfield2{k});
        RSD2std(k) = RSDstruct.(alone2).([predfield2{k},'std']);
    end
    
    %make prediction for data1
    data1 = datastruct.(alone1).(FPfield1);
    data1std = datastruct.(alone1).([FPfield1,'_std']);
    [pred1,pred1std] = applyRSpred(data1,mean(RSD1),mean(RSD2),...
        data1std,mean(RSD1std),mean(RSD2std));
    
    %get prediction for data2
    data2 = datastruct.(alone2).(FPfield2);
    data2std = datastruct.(alone2).([FPfield2,'_std']);
    [pred2,pred2std] = applyRSpred(data2,mean(RSD2),mean(RSD1),...
        data2std,mean(RSD2std),mean(RSD1std));
    
    %get measured perturbed output
    meas1 = datastruct.(FPnames{ii}).(FPfield1);
    meas1std = datastruct.(FPnames{ii}).([FPfield1,'_std']);
    meas2 = datastruct.(FPnames{ii}).(FPfield2);
    meas2std = datastruct.(FPnames{ii}).([FPfield2,'_std']);
    
    %package output data for plotting on bar chart
    modind1 = mask(strcmp(modsalone,alone1));
    modind2 = mask(strcmp(modsalone,alone2));
    if isempty(datamat{modind1})
        datamat{modind1} = [datastruct.(alone1).(FPfield1),NaN];
        datamatstd{modind1} = [datastruct.(alone1).([FPfield1,'_std']),NaN];
        names{modind1} = {alone1};
    end
    if isempty(datamat{modind2})
        datamat{modind2} = [datastruct.(alone2).(FPfield2),NaN];
        datamatstd{modind2} = [datastruct.(alone2).([FPfield2,'_std']),NaN];
        names{modind2} = {alone2};
    end 
    datamat{modind1} = [datamat{modind1};[meas1,pred1]];
    datamat{modind2} = [datamat{modind2};[meas2,pred2]];
    datamatstd{modind1} = [datamatstd{modind1};[meas1std,pred1std]];
    datamatstd{modind2} = [datamatstd{modind2};[meas2std,pred2std]];
    names{modind1} = [names{modind1}(:);FPnames{ii}];
    names{modind2} = [names{modind2}(:);FPnames{ii}];
    
    %write output
    %predstruct.(FPnames{ii}).(alone1).([FPfield1,'pred']) = pred1;
    %predstruct.(FPnames{ii}).(alone2).([FPfield1,'pred']) = pred1;
    
end

if ploton
    %plot bar chart of measured and predicted values with error bars
    figure(figh); clf;
    numalone = numel(modsalone);
    for k = 1:numalone
        %make subplot for each module alone
        subplot(1,numalone,k);
        bar(datamat{k},'grouped'); hold on
        set(gca, 'XTickLabel', names{k});
        
        %plot error bars on top of bars
        ngroups = size(datamat{k},1);
        nbars = size(datamat{k},2);
        % Calculating the width for each bar group
        groupwidth = min(0.8, nbars/(nbars + 1.5));
        
        % Set the position of each error bar in the centre of the main bar
        % Based on barweb.m by Bolu Ajiboye from MATLAB File Exchange
        x = zeros(nbars,ngroups);
        for ii = 1:nbars
            % Calculate center of each bar
            x(ii,:) = (1:ngroups) - groupwidth/2 + (2*ii-1) * groupwidth / (2*nbars);
        end
        errorbar(x', datamat{k}, datamatstd{k}, '.k', 'linestyle', 'none','Linewidth',1.5);
        legend('measured','predicted','Location','Best')
        set(gca,'fontsize',12)
        ylabel(modsalone{k})
    end
end


function [pred,predstd] = applyRSpred(data,RSD1,RSD2,datastd,RSD1std,RSD2std)

%use resource demand to predict circuit expression
pred = data*(1+RSD1)./(1+RSD1+RSD2);

if nargin < 4 || any([isempty(datastd),isempty(RSD1std),isempty(RSD2std)])
    predstd = [];
else
    dfddata = (1+RSD1)./(1+RSD1+RSD2);
    dfdRSD1 = data.*RSD2./(1+RSD1+RSD2).^2;
    dfdRSD2 = -data.*(1+RSD1)./(1+RSD1+RSD2).^2;
    %calculate standard error on prediction
    predstd = sqrt(dfddata.^2*datastd.^2 + dfdRSD1.^2*RSD1std.^2 + dfdRSD2.^2*RSD2std.^2);
end
```
