## Supplementary material for "Predicting Composition of Genetic Circuits with Resource Competition: Demand and Sensitivity": Experimental data and Matlab code: predictRS4.m

```
function dataoutstruct = predictRS4(modstruct,modulesout,ploton,fighvec)
%predict outputs

if nargin < 3 || isempty(ploton)
    ploton = true;
end
if ploton && nargin < 4
    fighvec = 1:6;
end

%find groups of non-RS modules that share properties
% modsalone = findpropRS(modstruct,'isalone');
modstogether = findpropRS(modstruct,'isalone',false);
normalmods = findpropRS(modstruct,'isResourceSensor',false);  %not resource sensors
% RSmods = findpropRS(modstruct,'isResourceSensor',true);

%experiments that are modules together and one module isn't a resource sensor
normalmodstogeth = intersect(normalmods,modstogether);
% normalmodsalone = intersect(normalmods,modsalone);
% RSmodsalone = intersect(RSmods,modsalone);

%[constmods, inducedmods] = splitinducedmods(modsoutalone);

dataoutstruct = struct;

for k = 1:length(normalmodstogeth)
    
    submodnames = modstruct.(normalmodstogeth{k}).containingmods;
    FPfields = modstruct.(normalmodstogeth{k}).FPout;
    
    %initialize output structk
    for ii = 1:length(submodnames)
        if ~isfield(dataoutstruct,submodnames{ii})
            tmpstruct = struct('data',[],'datastd',[],'perturb',[],'FPfield',[]);
            dataoutstruct.(submodnames{ii}) = tmpstruct;
        end
    end
    
    %get measured module outputs alone and with perturbation
    y1 = modulesout.(submodnames{1}).y;
    y1std = modulesout.(submodnames{1}).ystd;
    y1p = modstruct.(normalmodstogeth{k}).(FPfields{1});
    y1pstd = modstruct.(normalmodstogeth{k}).([FPfields{1},'std']);
    y2p = modstruct.(normalmodstogeth{k}).(FPfields{2});
    y2pstd = modstruct.(normalmodstogeth{k}).([FPfields{2},'std']);
    
    %resource properties of module 1
    if length(modulesout.(submodnames{1}).Q) > 1
        %get the index \in [1,2] of perturbing Q that doesn't overlap with
        %the perturbing module
        ind1 = find(~strcmpi(modulesout.(submodnames{1}).perturbname,submodnames{2}),1);
    else; ind1 = 1;
    end
    Q1 = modulesout.(submodnames{1}).Q(ind1);
    Q1std = modulesout.(submodnames{1}).Qstd(ind1);
    S11 = modulesout.(submodnames{1}).S(ind1);
    S11std = modulesout.(submodnames{1}).Sstd(ind1);
    S12 = modulesout.(submodnames{1}).S2(ind1);
    S12std = modulesout.(submodnames{1}).S2std(ind1);
    S13 = modulesout.(submodnames{1}).S3(ind1);
    S13std = modulesout.(submodnames{1}).S3std(ind1);
    
    %resource properties of module 2
    if length(modulesout.(submodnames{2}).perturbname) > 1
        %get the index \in [1,2] of perturbing Q that doesn't overlap with
        %the perturbing module
        ind2 = find(~strcmpi(modulesout.(submodnames{2}).perturbname,submodnames{1}),1);
    else; ind2 = 1;
    end
    Q2 = modulesout.(submodnames{2}).Q(:,ind2);
    Q2std = modulesout.(submodnames{2}).Qstd(:,ind2);
    S21 = modulesout.(submodnames{2}).S(:,ind2);
    S21std = modulesout.(submodnames{2}).Sstd(:,ind2);
    S22 = modulesout.(submodnames{2}).S2(:,ind2);
    S22std = modulesout.(submodnames{2}).S2std(:,ind2);
    S23 = modulesout.(submodnames{2}).S3(:,ind2);
    S23std = modulesout.(submodnames{2}).S3std(:,ind2);
    y2 = modulesout.(submodnames{2}).y;
    y2std = modulesout.(submodnames{2}).ystd;
    
    submodnames
    %NOTE: need to update for inducible modules
    
    %make predictions
    [y11pred,y11predstd] = calcypred(y1,S11,Q2,y1std,S11std,Q2std);
    [y21pred,y21predstd] = calcypred(y2,S21,Q1,y2std,S21std,Q1std);
    [y12pred,y12predstd] = calcypred2(y1,S12,Q2,y1std,S12std,Q2std); %std = 0?
    [y22pred,y22predstd] = calcypred2(y2,S22,Q1,y2std,S22std,Q1std);
    [y13pred,y13predstd] = calcypred3(y1,S13,Q1,Q2,y1std,S13std,Q1std,Q2std);
    [y23pred,y23predstd] = calcypred3(y2,S23,Q2,Q1,y2std,S23std,Q2std,Q1std);
    
    %format output for bar plots
    if length(y1) == 1 && length(y2) == 1
        %[alone, perturbed measured, predicted1, ...]
        dataout1 = [y1,NaN,NaN,NaN; y1p,y11pred,y12pred,y13pred];
        dataout1std = [y1std,NaN,NaN,NaN; y1pstd,y11predstd,y12predstd,y13predstd];
        dataout2 = [y2,NaN,NaN,NaN; y2p,y21pred,y22pred,y23pred];
        dataout2std = [y2std,NaN,NaN,NaN; y2pstd,y21predstd,y22predstd,y23predstd];
    else
        warning(['skipped data: ',normalmodstogeth{k}]);
        %shape outputs for plotting (induced modules). Predictions are
        %column vectors. Use cell arrays
        dataout1 = {y1,y1p,y11pred,y12pred,y13pred};
        dataout1std = {y1std,y1pstd,y11predstd,y12predstd,y13predstd};
        dataout2 = {y2,y2p,y21pred,y22pred,y23pred};
        dataout2std = {y2std,y2pstd,y21predstd,y22predstd,y23predstd};
        %{alone, perturbed measured, predicted1, ...}
    end
    
    %output
    olddata1 = dataoutstruct.(submodnames{1}).data;
    olddata2 = dataoutstruct.(submodnames{2}).data;
    olddata1std = dataoutstruct.(submodnames{1}).datastd;
    olddata2std = dataoutstruct.(submodnames{2}).datastd;
    oldperturbnames1 = dataoutstruct.(submodnames{1}).perturb;
    oldperturbnames2 = dataoutstruct.(submodnames{2}).perturb;
    oldFPfields1 = dataoutstruct.(submodnames{1}).FPfield;
    oldFPfields2 = dataoutstruct.(submodnames{2}).FPfield;
    
    dataoutstruct.(submodnames{1}).data = [olddata1; dataout1];
    dataoutstruct.(submodnames{2}).data = [olddata2; dataout2];
    dataoutstruct.(submodnames{1}).datastd = [olddata1std; dataout1std];
    dataoutstruct.(submodnames{2}).datastd = [olddata2std; dataout2std];
    dataoutstruct.(submodnames{1}).perturb = [oldperturbnames1(:)',submodnames(2)];
    dataoutstruct.(submodnames{2}).perturb = [oldperturbnames2(:)',submodnames(1)];
    dataoutstruct.(submodnames{1}).FPfield = unique([oldFPfields1(:)',FPfields(1)]);
    dataoutstruct.(submodnames{2}).FPfield = unique([oldFPfields2(:)',FPfields(2)]);
end

if ploton
    %plot
    outnames = fieldnames(dataoutstruct);
    if length(fighvec) < length(outnames)
        warning('fighvec length less than the number of modules')
        fighvec = fighvec(1):(length(outnames)+fighvec(1)-1);
    end
    for jj = 1:length(outnames)
        data = dataoutstruct.(outnames{jj}).data;
        datastd = dataoutstruct.(outnames{jj}).datastd;
        %determine which predictions to plot:
        %1st column = measured; 2nd = predicted with S1, 3rd = with S2, 4th = with S3
        pinds = 1:2;
        
        %plot
        figure(fighvec(jj)); clf;
        bh = plotbarpredict([data(1,pinds);data(2:2:end,pinds)]'/data(1,1),...
            [datastd(1,pinds);datastd(2:2:end,pinds)]'/data(1,1));
        ylabel(dataoutstruct.(outnames{jj}).FPfield)
        names = combinenames({outnames{jj}},dataoutstruct.(outnames{jj}).perturb,true);
        xticklabels(names)
        set(gca,'fontsize',14)
        %legend('measured','predicted_1','predicted_2','location','best')
        legend('measured','predicted','location','best')
        ylim([0,1.2])
        
        if contains(dataoutstruct.(outnames{jj}).FPfield,'BFP')
            %make colors match
            blue1 = [91,155,213]/255;       %dark blue
            blue2 = [125,195,251]/255;      %medium blue
            blue3 = [173,222,255]/255;      %light blue
            set(bh(1),'FaceColor',blue1)    %measured
            set(bh(2),'FaceColor',blue3)    %predicted1
            %set(bh(3),'FaceColor',blue3)    %predicted2
        elseif contains(dataoutstruct.(outnames{jj}).FPfield,'GFP')
            green1 = [120,171,48]/255;      %dark green
            green2 = [45,190,6]/255;        %emerald green
            green3 = [163,252,67]/255;      %light green
            set(bh(1),'FaceColor',green2)   %measured
            set(bh(2),'FaceColor',green3)   %predicted1
            %set(bh(3),'FaceColor',green3)   %predicted2
        elseif contains(dataoutstruct.(outnames{jj}).FPfield,'RFP')
            red1 = [207,81,81]/255;         %dark red
            red2 = [240,122,122]/255;       %light red
            red3 = [255,168,168]/255;       %lighter red
            set(bh(1),'FaceColor',red1)     %measured
            set(bh(2),'FaceColor',red3)     %predicted1
            %set(bh(3),'FaceColor',red3)     %predicted2
        end
    end
end


function [ypred,ypredstd] = calcypred(yalone,S,Q,ystd,Sstd,Qstd)

%calculate perturbed module estimate
ypred = yalone./(1 - S.*Q);

%calculate output standard error
dypdy = 1./(1-S.*Q);
dypdS = yalone.*Q./(1 - S.*Q).^2;
dypdQ = yalone.*S./(1 - S.*Q).^2;
ypredstd = sqrt(dypdy.^2.*ystd.^2 + dypdS.^2.*Sstd.^2 + dypdQ.^2.*Qstd.^2);


function [ypred,ypredstd] = calcypred2(yalone,S2,Q1,ystd,Sstd,Qstd)

%calculate perturbed module estimate
ypred = yalone.*(1 + S2.*Q1);

%calculate output standard error
dypdy = (1 + S2.*Q1);
dypdS = yalone.*Q1;
dypdQ = yalone.*S2;
ypredstd = sqrt(dypdy.^2.*ystd.^2 + dypdS.^2.*Sstd.^2 + dypdQ.^2.*Qstd.^2);


function [pred,predstd] = calcypred3(y1,S2,Q1,Q2,ystd,Sstd,Q1std,Q2std)

%use resource demand to predict circuit expression
pred = y1.*S2.*(1+Q1)./(1+Q1+Q2);

%calculate standard error on predicted output
dfdy = S2.*(1+Q1)./(1+Q1+Q2);
dfdS = y1.*(1+Q1)./(1+Q1+Q2);
dfdQ1 = y1.*S2*Q2./(1+Q1+Q2).^2;
dfdQ2 = -y1.*S2.*(1+Q1)./(1+Q1+Q2).^2;
predstd = sqrt(dfdy.^2.*ystd.^2 + dfdS.^2.*Sstd.^2 + dfdQ1.^2.*Q1std.^2 + ...
    dfdQ2.^2.*Q2std.^2);


function bh = plotbarpredict(data,datastd)

nbars = size(data,1);
ngroups = size(data,2);

groupwidth = min(0.8, nbars/(nbars + 1.5));
% Set the position of each error bar in the centre of the main bar
% Based on barweb.m by Bolu Ajiboye from MATLAB File Exchange
x = zeros(nbars,ngroups);
for ii = 1:nbars
    % Calculate center of each bar
    x(ii,:) = (1:ngroups) - groupwidth/2 + (2*ii-1) * groupwidth / (2*nbars);
end

bh = bar(data'); hold on;
set(bh,'edgecolor','none');
errorbar(x,data,datastd,'.k','linestyle','none','Linewidth',1.5);


function [constmods, inducedmods] = splitinducedmods(modstruct)
%split modules into induced and constitutive modules based off the length
%of the input vector u
modnames = fieldnames(modstruct);
for ii = 1:length(modnames)
    if length(modstruct.(modnames{ii}).u) > 1
        inducedmods.(modnames{ii}) = modstruct.(modnames{ii});
    else
        constmods.(modnames{ii}) = modstruct.(modnames{ii});
    end
end


function plotQScurves(uvec,yvec,yvecstd)
%plot Q or S curves vs input u
zerooffset = 1e-3;
if iscell(uvec)
    umax = max(cellfun(@max,uvec));
    umin = min(max(cellfun(@min,uvec),zerooffset));
    for ii = 1:length(uvec)
        semilogx(max(uvec{ii},zerooffset),yvec{ii},'--','linewidth',2); hold on
        for k = 1:size(yvec{ii},2)
            errorbar(max(uvec{ii},zerooffset),yvec{ii}(:,k),yvecstd{ii}(:,k),'.k','linewidth',1.5);
        end
    end
else
    umax = max(uvec,[],'all');
    umin = max([min(uvec,[],'all'),zerooffset]);
    semilogx(max(uvec,zerooffset),yvec,'linewidth',2); hold on
    for jj = 1:size(yvec,2)
        errorbar(max(uvec,zerooffset),yvec(:,jj),yvecstd(:,jj),'.k','linewidth',1.5);
    end
end
%change zero tick mark
set(gca,'fontsize',14);
if iscell(uvec) && cellfun(@(x) any(x == 0),uvec)
    xt = xticks;
    xticklabels([0,xt(2:end)])
elseif any(uvec == 0)
    xt = xticks;
    xticklabels([0,xt(2:end)])
end
xlim([umin,umax])
```
