## Supplementary material for "Predicting Composition of Genetic Circuits with Resource Competition: Demand and Sensitivity": Experimental data and Matlab code: predictRS5.m

```
function dataout = predictRS5(moduledata,ploton,fighvec,constpinds,indpind,includeinduced)
%predict outputs

if nargin < 3 || isempty(ploton)
    ploton = false;
end
if ploton && nargin < 3
    fighvec = 1:6;
end
if ploton && nargin < 4
    %which predicitons to plot for constitutive modules
    %constpinds = [1,[1,5,4,8,11] + 1]; %[1,3+1];%,1,4,2,5,7,8] + 1];
    %constpinds = [1,[4,11,8]+1];
    constpinds = [1,[4]+1];
end
if ploton && nargin < 5
    %which S to use for inducible module predictions?
    indpind = 4;
    %indpind = 11;
end
if ploton && nargin < 6
    %include induced mod perturbations in constitutive mod predictions
    includeinduced = false;
end

%find groups of non-RS modules that share properties
modsalone = findpropRS(moduledata,'isalone');
modstogether = findpropRS(moduledata,'isalone',false);
normalmods = findpropRS(moduledata,'isResourceSensor',false);  %not resource sensors
%experiments that are modules together and one module isn't a resource sensor
normalmodstogeth = intersect(normalmods,modstogether);

dataout = moduledata;

for k = 1:length(normalmodstogeth)
    
    %unpackage data about modules in pairs
    modnames = moduledata.(normalmodstogeth{k}).containingmods;
    FPfields = moduledata.(normalmodstogeth{k}).FPout;
    
    %get measured module outputs alone and with perturbation
    y1 = moduledata.(modnames{1}).y;
    y1std = moduledata.(modnames{1}).ystd;
    y2 = moduledata.(modnames{2}).y;
    y2std = moduledata.(modnames{2}).ystd;
    y1p = moduledata.(normalmodstogeth{k}).(FPfields{1});
    y1pstd = moduledata.(normalmodstogeth{k}).([FPfields{1},'std']);
    y2p = moduledata.(normalmodstogeth{k}).(FPfields{2});
    y2pstd = moduledata.(normalmodstogeth{k}).([FPfields{2},'std']);
    
    %resource properties of module 1
    %average together if more than one Q measured
    Q1 = mean(moduledata.(modnames{1}).Q,2);
    Q1std = mean(moduledata.(modnames{1}).Qstd,2);
    S11 = mean(moduledata.(modnames{1}).S,2);
    S11std = mean(moduledata.(modnames{1}).Sstd,2);
    S11theory = mean(moduledata.(modnames{1}).S5,2);
    S11stdtheory = mean(moduledata.(modnames{1}).S5std,2);
    S12 = mean(moduledata.(modnames{1}).S2,2);
    S12std = mean(moduledata.(modnames{1}).S2std,2);
    S12theory = mean(moduledata.(modnames{1}).S6,2);
    S12stdtheory = mean(moduledata.(modnames{1}).S6std,2);
    S13 = mean(moduledata.(modnames{1}).S3,2);
    S13std = mean(moduledata.(modnames{1}).S3std,2);
    S13theory = mean(moduledata.(modnames{1}).S7,2);
    S13stdtheory = mean(moduledata.(modnames{1}).S7std,2);
    S14 = mean(moduledata.(modnames{1}).S4,2);
    S14std = mean(moduledata.(modnames{1}).S4std,2);
    S14theory = mean(moduledata.(modnames{1}).S8,2);
    S14stdtheory = mean(moduledata.(modnames{1}).S8std,2);
    S15 = mean(moduledata.(modnames{1}).S9,2);
    S15std = mean(moduledata.(modnames{1}).S9std,2);
    
    %resource properties of module 2
    %average together if more than one Q measured
    Q2 = mean(moduledata.(modnames{2}).Q,2);
    Q2std = mean(moduledata.(modnames{2}).Qstd,2);
    S21 = mean(moduledata.(modnames{2}).S,2);
    S21std = mean(moduledata.(modnames{2}).Sstd,2);
    S21theory = mean(moduledata.(modnames{2}).S5,2);
    S21stdtheory = mean(moduledata.(modnames{2}).S5std,2);
    S22 = mean(moduledata.(modnames{2}).S2,2);
    S22std = mean(moduledata.(modnames{2}).S2std,2);
    S22theory = mean(moduledata.(modnames{2}).S6,2);
    S22stdtheory = mean(moduledata.(modnames{2}).S6std,2);
    S23 = mean(moduledata.(modnames{2}).S3,2);
    S23std = mean(moduledata.(modnames{2}).S3std,2);
    S23theory = mean(moduledata.(modnames{2}).S7,2);
    S23stdtheory = mean(moduledata.(modnames{2}).S7std,2);
    S24 = mean(moduledata.(modnames{2}).S4,2);
    S24std = mean(moduledata.(modnames{2}).S4std,2);
    S24theory = mean(moduledata.(modnames{2}).S8,2);
    S24stdtheory = mean(moduledata.(modnames{2}).S8std,2);
    S25 = mean(moduledata.(modnames{2}).S9,2);
    S25std = mean(moduledata.(modnames{2}).S9std,2);
    
    %make predictions
    [y11pred,y11predstd] = calcypred(y1,S11, Q2,y1std,S11std,Q2std);
    [y21pred,y21predstd] = calcypred(y2,S21, Q1,y2std,S21std,Q1std);
    [y12pred,y12predstd] = calcypred2(y1,S12,Q2,y1std,S12std,Q2std); %std = 0?
    [y22pred,y22predstd] = calcypred2(y2,S22,Q1,y2std,S22std,Q1std);
    [y13pred,y13predstd] = calcypred3(y1,S13,Q1,Q2,y1std,S13std,Q1std,Q2std);
    [y23pred,y23predstd] = calcypred3(y2,S23,Q2,Q1,y2std,S23std,Q2std,Q1std);
    [y14pred,y14predstd] = calcypred5(y1,S14,Q1,Q2,y1std,S14std,Q1std,Q2std);
    [y24pred,y24predstd] = calcypred5(y2,S24,Q2,Q1,y2std,S24std,Q2std,Q1std);
    [y111pred,y111predstd] = calcypred6(y1,S15,Q1,Q2,y1std,S15std,Q1std,Q2std);
    [y211pred,y211predstd] = calcypred6(y2,S25,Q2,Q1,y2std,S25std,Q2std,Q1std);
    [y19pred,y19predstd] = calcypred4(y1,S11,Q2,y1std,S11std,Q2std);
    [y29pred,y29predstd] = calcypred4(y2,S21,Q1,y2std,S21std,Q1std);
    
    %based off of Q with module structure theory
    %make predictions
    [y15pred,y15predstd] = calcypred(y1, S11theory,Q2,y1std,S11stdtheory,Q2std);
    [y25pred,y25predstd] = calcypred(y2, S21theory,Q1,y2std,S21stdtheory,Q1std);
    [y16pred,y16predstd] = calcypred2(y1,S12theory,Q2,y1std,S12stdtheory,Q2std); %std = 0?
    [y26pred,y26predstd] = calcypred2(y2,S22theory,Q1,y2std,S22stdtheory,Q1std);
    [y17pred,y17predstd] = calcypred3(y1,S13theory,Q1,Q2,y1std,S13stdtheory,Q1std,Q2std);
    [y27pred,y27predstd] = calcypred3(y2,S23theory,Q2,Q1,y2std,S23stdtheory,Q2std,Q1std);
    [y18pred,y18predstd] = calcypred5(y1,S14theory,Q1,Q2,y1std,S14stdtheory,Q1std,Q2std);
    [y28pred,y28predstd] = calcypred5(y2,S24theory,Q2,Q1,y2std,S24stdtheory,Q2std,Q1std);
    [y110pred,y110predstd] = calcypred4(y1,S11theory,Q2,y1std,S11stdtheory,Q2std);
    [y210pred,y210predstd] = calcypred4(y2,S21theory,Q1,y2std,S21stdtheory,Q1std);
    
    %save perturbations
    prevpredy1 = dataout.(modnames{1}).predperturby;
    prevpredy1std = dataout.(modnames{1}).predperturbystd;
    prevperturby1 = dataout.(modnames{1}).perturby;
    prevperturby1std = dataout.(modnames{1}).perturbystd;
    prevperturbname1 = dataout.(modnames{1}).perturbname;
    prevpredy2 = dataout.(modnames{2}).predperturby;
    prevpredy2std = dataout.(modnames{2}).predperturbystd;
    prevperturby2 = dataout.(modnames{2}).perturby;
    prevperturby2std = dataout.(modnames{2}).perturbystd;
    prevperturbname2 = dataout.(modnames{2}).perturbname;
    
    %append predictions and perturbed outputs into output data object
    %for module 1
    if ~isempty(prevpredy1)
        dataout.(modnames{1}).predperturby    = [prevpredy1,...
            {[y11pred,y12pred,y13pred,y14pred,y15pred,y16pred,y17pred,...
            y18pred,y19pred,y110pred,y111pred]}];
        dataout.(modnames{1}).predperturbystd = [prevpredy1std,...
            {[y11predstd,y12predstd,y13predstd,y14predstd,y15predstd,...
            y16predstd,y17predstd,y18predstd,y19predstd,y110predstd,y111predstd]}];
    else
        dataout.(modnames{1}).predperturby    = {[y11pred,y12pred,y13pred,...
            y14pred,y15pred,y16pred,y17pred,y18pred,y19pred,y110pred,y111pred]};
        dataout.(modnames{1}).predperturbystd = {[y11predstd,y12predstd,...
            y13predstd,y14predstd,y15predstd,y16predstd,y17predstd,y18predstd,...
            y19predstd,y110predstd,y111predstd]};
    end
    dataout.(modnames{1}).perturbname = [prevperturbname1,modnames{2}];
    dataout.(modnames{1}).perturby    = [prevperturby1,y1p];
    dataout.(modnames{1}).perturbystd = [prevperturby1std,y1pstd];
    
    %for module 2
    if ~isempty(prevpredy2)
        dataout.(modnames{2}).predperturby    = [prevpredy2,...
            {[y21pred,y22pred,y23pred,y24pred,y25pred,y26pred,y27pred,...
            y28pred,y29pred,y210pred,y211pred]}];
        dataout.(modnames{2}).predperturbystd = [prevpredy2std,...
            {[y21predstd,y22predstd,y23predstd,y24predstd,y25predstd,...
            y26predstd,y27predstd,y28predstd,y29predstd,y210predstd,y211predstd]}];
    else
        dataout.(modnames{2}).predperturby    = {[y21pred,y22pred,y23pred,...
            y24pred,y25pred,y26pred,y27pred,y28pred,y29pred,y210pred,y211pred]};
        dataout.(modnames{2}).predperturbystd = {[y21predstd,y22predstd,...
            y23predstd,y24predstd,y25predstd,y26predstd,y27predstd,y28predstd,...
            y29predstd,y210predstd,y211predstd]};
    end
    dataout.(modnames{2}).perturbname = [prevperturbname2,modnames{1}];
    dataout.(modnames{2}).perturby    = [prevperturby2,y2p];
    dataout.(modnames{2}).perturbystd = [prevperturby2std,y2pstd];
end

%plot
if ploton
    if length(fighvec) < length(modsalone)
        fighvec = fighvec(1):(fighvec(1)+length(modsalone)-1);
        warning(['fighvec length less than the number of modules. Using figures',...
            num2str(fighvec)])
    end
    for jj = 1:length(modsalone)
        modplt = dataout.(modsalone{jj});
        %assuming all mods alone are modules contained in a pair
        if length(modplt.u) > 1
            %inducible module prediction plot
            uvec = modplt.u;
            inds = cellfun(@(x) ~isempty(x),modplt.predperturby);
            %get data
            y = abs(modplt.y);
            ystd = modplt.ystd./max(y);
            yp = abs([modplt.perturby{inds}]./max(y));
            ypstd = [modplt.perturbystd{inds}]./max(y);
            pred = abs([modplt.predperturby{inds}]./max(y));
            predstd = [modplt.predperturbystd{inds}]./max(y);
            
            %select predictions and group with the appropriate measured
            %perturbed data
            pred2 = pred(:,indpind:11:end);
            pred2std = predstd(:,indpind:11:end);
            
            %skip B170 perturbation?
            %yp = yp(:,2:end);
            %ypstd = ypstd(:,2:end);
            %pred2 = pred2(:,2:end);
            %pred2std = pred2std(:,2:end);
            
            [yppred,yppredstd] = deal(zeros(size(yp,1),2*size(yp,2)));
            for k = 1:size(yp,2)
                yppred(:,2*k-1) = yp(:,k);
                yppred(:,2*k) = pred2(:,k);
                yppredstd(:,2*k-1) = ypstd(:,k);
                yppredstd(:,2*k) = pred2std(:,k);
            end
            %note all predictions are stacked together
            dataind = [y./max(y),yppred];
            dataindstd = [ystd,yppredstd];
            
            figure(fighvec(jj)); clf;
            %plot bar chart
            bh = plotbarpredict(dataind',dataindstd');
            %make predictions and measurements same color
            for l = 1:size(yp,2)
                set(bh(2*l),'FaceColor',get(bh(1+2*l),'FaceColor'))
            end
            %ph = plotinducedmod(uvec,y./max(y),ystd./max(y),yp,ypstd,...
            %    pred(:,indpind:11:end),predstd(:,indpind:11:end));
            set(gca,'yscale','linear','xticklabels',uvec,'fontsize',14)
            ylim([-0.1,1.1])
            ylabel(modplt.FPout)
            xlabel('intput, u [nM]')
            legend([bh(1),bh(2:2:end)],['alone',modplt.perturbname(3:end)],'Location','Best')
        else
            %constitutive module predictions
            y = modplt.y;
            ystd = modplt.ystd;
            %indicies where module is perturbed by a non-resource sensor
            %constitutive module indicies
            indsC = cellfun(@(x) ~isempty(x) & size(x,1) == 1,modplt.predperturby);
            %indicies where module is perturbed by non-RS inducible module
            indsI = cellfun(@(x) ~isempty(x) & size(x,1) > 1,modplt.predperturby);
            %shape data and predictions into matrix for plotting
            yp = vertcat(modplt.perturby{indsC});
            datamat0 = vertcat(modplt.predperturby{indsC});
            %1st column = measured; 2nd = predicted with S1, 3rd = with S2, 4th = with S3
            datamat  = [y,NaN(1,size(datamat0,2)); [yp,datamat0]]./y;
            ypstd = vertcat(modplt.perturbystd{indsC});
            datamat0std = vertcat(modplt.predperturbystd{indsC});
            datamatstd  = [ystd,NaN(1,size(datamat0std,2)); [ypstd,datamat0std]]./y;
            
            %plot constitutive module predictions
            figure(fighvec(jj)); clf;
            if any(indsI) && includeinduced; subplot(1,3,[1,2]); end
            bh = plotbarpredict(datamat(:,constpinds)',datamatstd(:,constpinds)');
            ylabel(modplt.FPout)
            names = combinenames(modsalone(jj),modplt.perturbname(indsC),true);
            xticklabels(names)
            set(gca,'fontsize',14)
            legend('measured','predicted','location','best')
            ylim([-0.1,1.2])
            if contains(modplt.FPout,'BFP')
                %make colors match
                blue1 = [91,155,213]/255;       %dark blue
                blue2 = [125,195,251]/255;      %medium blue
                blue3 = [173,222,255]/255;      %light blue
                set(bh(1),'FaceColor',blue1)    %measured
                set(bh(2),'FaceColor',blue3)    %predicted1
                if length(constpinds) >= 3
                    set(bh(3),'FaceColor',blue2)    %predicted2
                end
            elseif contains(modplt.FPout,'GFP')
                green1 = [120,171,48]/255;      %dark green
                green2 = [45,190,6]/255;        %emerald green
                green3 = [163,252,67]/255;      %light green
                set(bh(1),'FaceColor',green2)   %measured
                set(bh(2),'FaceColor',green3)   %predicted1
                if length(constpinds) >= 3
                    set(bh(3),'FaceColor',green1)   %predicted2
                end
            elseif contains(modplt.FPout,'RFP')
                red1 = [207,81,81]/255;         %dark red
                red2 = [240,122,122]/255;       %light red
                red3 = [255,168,168]/255;       %lighter red
                set(bh(1),'FaceColor',red1)     %measured
                set(bh(2),'FaceColor',red3)     %predicted1
                if length(constpinds) >= 3
                    set(bh(3),'FaceColor',red2)     %predicted2
                end
            elseif contains(modplt.FPout,'YFP')
                yellow1 = [237,177,32]/255;
                yellow2 = [255,255,0]/255;
                yellow3 = [196,200,21]/255;
                set(bh(1),'FaceColor',yellow1)     %measured
                set(bh(2),'FaceColor',yellow2)     %predicted1
                if length(constpinds) >= 3
                    set(bh(3),'FaceColor',yellow3)     %predicted2
                end
            end
            
            if any(indsI) && includeinduced
                %plot curves for induced module perturbing constitutive one
                subplot(1,3,3);
                %loop through induced perturbing modules
                indsI2 = find(indsI);
                for k = indsI2
                    %get u vec from perturbing module
                    perturbname = modplt.perturbname{k};
                    uvec = dataout.(perturbname).u;
                    %get data
                    y = modplt.y;
                    y2 = modplt.y*ones(size(uvec))/y;           %cast up to correct size
                    y2std = modplt.ystd*ones(size(uvec))/y;     %cast up to correct size
                    yp = modplt.perturby{k}/y;
                    ypstd = modplt.perturbystd{k}/y;
                    pred = modplt.predperturby{k}/y;
                    predstd = modplt.predperturbystd{k}/y;
                    %ph2 = plotinducedmod(uvec,y2,y2std,yp,ypstd,pred(:,indpind),predstd(:,indpind));
                    
                    dataind = [y2',yp,pred(:,indpind)];
                    dataindstd = [y2',yp,pred(:,indpind)];
                    bh = plotbarpredict(dataind,dataindstd);
                    %(uvec,y2,y2std,yp,ypstd,pred(:,indpind),predstd(:,indpind));
                    ylim([0,1.2])
                    %legend([ph2{1};ph2{2}],{'measured','pred_1','pred_2','pred_3'},...
                    %    'Location','Best')
                end
            end
        end
    end
end


function [ypred,ypredstd] = calcypred(yalone,S,Q,ystd,Sstd,Qstd)
%calculate perturbed module estimate
ypred = yalone./(1 - S.*Q);

%calculate output standard error
dypdy = 1./(1-S.*Q);
dypdS = yalone.*Q./(1 - S.*Q).^2;
dypdQ = yalone.*S./(1 - S.*Q).^2;
ypredstd = sqrt(dypdy.^2.*ystd.^2 + dypdS.^2.*Sstd.^2 + dypdQ.^2.*Qstd.^2);


function [ypred,ypredstd] = calcypred2(yalone,S,Qp,ystd,Sstd,Qpstd)
%calculate perturbed module estimate
ypred = yalone.*(1 + S.*Qp + 2*(S*Qp).^2/2 + 6*(S*Qp).^3/6 + 24*(S*Qp).^4/24);

%calculate output standard error
dypdy = (1 + S.*Qp);
dypdS = yalone.*Qp;
dypdQ = yalone.*S;
ypredstd = sqrt(dypdy.^2.*ystd.^2 + dypdS.^2.*Sstd.^2 + dypdQ.^2.*Qpstd.^2);


function [pred,predstd] = calcypred3(y1,S1,Q1,Q2,ystd,Sstd,Q1std,Q2std)
%use resource demand to predict circuit expression
pred = y1.*S1.*(1+Q1)./(1+Q1+Q2);

%calculate standard error on predicted output
dfdy = S1.*(1+Q1)./(1+Q1+Q2);
dfdS = y1.*(1+Q1)./(1+Q1+Q2);
dfdQ1 = y1.*S1*Q2./(1+Q1+Q2).^2;
dfdQ2 = -y1.*S1.*(1+Q1)./(1+Q1+Q2).^2;
predstd = sqrt(dfdy.^2.*ystd.^2 + dfdS.^2.*Sstd.^2 + dfdQ1.^2.*Q1std.^2 + ...
    dfdQ2.^2.*Q2std.^2);


function [ypred,ypredstd] = calcypred4(yalone,S1,Q2,ystd,S1std,Q2std)
%calculate perturbed module estimate S1 = self S; Q2 = perturbing Q
ypred = yalone./(1 - S1.*Q2 + 2*S1.^2*Q2.^2/2 - 6*S1.^3*Q2.^3/6 + 24*S1.^4*Q2.^4/24);
%include higher order terms for consitutive module
%NOTE: error bars not correct for all new terms

%calculate output standard error
dypdy = 1./(1 - S1.*Q2 + 2*S1.^2*Q2.^2/2 - 6*S1.^3*Q2.^3/6 + 24*S1.^4*Q2.^4/24);
dypdS = yalone.*Q2./(1 - S1.*Q2).^2;
dypdQ = yalone.*S1./(1 - S1.*Q2).^2;
ypredstd = sqrt(dypdy.^2.*ystd.^2 + dypdS.^2.*S1std.^2 + dypdQ.^2.*Q2std.^2);


function [pred,predstd] = calcypred5(y1,S1,Q1,Q2,ystd,Sstd,Q1std,Q2std)
%use resource demand to predict circuit expression
pred = y1.*(1 + S1.*Q2).*(1+Q1)./(1+Q1+Q2);

%calculate standard error on predicted output
dfdy = (1+S1*Q2).*(1+Q1)./(1+Q1+Q2);
dfdS = y1*Q2.*(1+Q1)./(1+Q1+Q2);
dfdQ1 = y1.*(1+S1.*Q2).*Q2./(1+Q1+Q2).^2;
dfdQ2 = -y1.*(S1.*(1+Q1)./(1+Q1+Q2) - (1+S1.*Q2).*(1+Q1)./(1+Q1+Q2).^2);
predstd = sqrt(dfdy.^2.*ystd.^2 + dfdS.^2.*Sstd.^2 + dfdQ1.^2.*Q1std.^2 + ...
    dfdQ2.^2.*Q2std.^2);


function [pred,predstd] = calcypred6(y1,S1,Q1,Q2,ystd,Sstd,Q1std,Q2std)
%use resource demand to predict circuit expression
pred = y1.*(1./(1 - S1.*Q2)).*(1+Q1)./(1+Q1+Q2);

%calculate standard error on predicted output
dfdy = (1./(1 - S1.*Q2)).*(1+Q1)./(1+Q1+Q2);
dfdS = y1*Q2.*(1+Q1)./((1 - S1.*Q2).^2.*(1+Q1+Q2));
dfdQ1 = y1.*Q2./((1 - S1.*Q2).*(1+Q1+Q2).^2);
dfdQ2 = -y1.*(1+Q1).*(S1.*(Q1 + 2*Q2 + 1) - 1)./((1 - S1.*Q2).^2.*(1+Q1+Q2).^2);
predstd = sqrt(dfdy.^2.*ystd.^2 + dfdS.^2.*Sstd.^2 + dfdQ1.^2.*Q1std.^2 + ...
    dfdQ2.^2.*Q2std.^2);


function bh = plotbarpredict(data,datastd)
nbars = size(data,1);
ngroups = size(data,2);
groupwidth = min(0.8, nbars/(nbars + 1.5));
% Set the position of each error bar in the centre of the main bar
% Based on barweb.m by Bolu Ajiboye from MATLAB File Exchange
x = zeros(nbars,ngroups);
for ii = 1:nbars
    % Calculate center of each bar
    x(ii,:) = (1:ngroups) - groupwidth/2 + (2*ii-1) * groupwidth / (2*nbars);
end

bh = bar(data'); hold on;
set(bh,'edgecolor','none');
errorbar(x,data,datastd,'.k','linestyle','none','Linewidth',1.5);


function ph = plotinducedmod(uvec,yalone,yalonestd,yp,ypstd,ypred,ypredstd)
%plot Q or S curves vs input u
zerooffset = 1e-2;
ax = gca;
ph = cell(2,1);
umax = max(uvec,[],'all');
umin = max([min(uvec,[],'all'),zerooffset]);

%plot y alone
semilogx(max(uvec,zerooffset),yalone,'-k','linewidth',2); hold on
errorbar(max(uvec,zerooffset),yalone,yalonestd,'.k','linewidth',0.5);
set(ax,'ColorOrderIndex',1)
%plot actual y perturb
coi = get(ax,'colororderindex');
ph{1} = semilogx(max(uvec,zerooffset),yp,'linewidth',2); hold on
for jj = 1:size(yp,2)
    x = max(uvec,zerooffset).*(1 + 0*rand(size(uvec)));
    errorbar(x,yp(:,jj),ypstd(:,jj),'.k','linewidth',0.5);
end
%plot predictions
set(ax,'colororderindex',coi);
ph{2} = semilogx(max(uvec,zerooffset),ypred,'--','linewidth',2); hold on
for ii = 1:size(ypred,2)
    x = max(uvec,zerooffset).*(1 + 0*rand(size(uvec)));
    errorbar(x,ypred(:,ii),ypredstd(:,ii),'.k','linewidth',0.5);
end

%change zero tick mark
set(gca,'fontsize',14);
if iscell(uvec) && cellfun(@(x) any(x == 0),uvec)
    xt = xticks;
    xticklabels([0,xt(2:end)])
elseif any(uvec == 0)
    xt = xticks;
    xticklabels([0,xt(2:end)])
end
xlim([umin,umax])


function colorplots(name,ph1,ph2,bluename,greenname,redname,yellowname)

if isempty(bluename); bluename = 'B'; end
if isempty(greenname); greenname = 'G'; end
if isempty(redname); redname = 'R'; end
if isempty(yellowname); yellowname = 'Y'; end

if contains(name,bluename)
    %make colors match
    blue1 = [91,155,213]/255;       %dark blue
    %blue2 = [125,195,251]/255;      %medium blue
    blue3 = [173,222,255]/255;      %light blue
    set(ph1,'Color',blue1)          %measured
    set(ph2,'Color',blue3)          %predicted1
elseif contains(name,greenname)
    %green1 = [120,171,48]/255;     %dark green
    green2 = [45,190,6]/255;        %emerald green
    green3 = [163,252,67]/255;      %light green
    set(ph1,'Color',green2)         %measured
    set(ph2,'Color',green3)         %predicted1
elseif contains(name,redname)
    red1 = [207,81,81]/255;         %dark red
    %red2 = [240,122,122]/255;      %light red
    red3 = [255,168,168]/255;       %lighter red
    set(ph1,'Color',red1)           %measured
    set(ph2,'Color',red3)           %predicted1
elseif contains(name,yellowname)
    yellow1 = [237,177,32]/255;
    yellow2 = [255,255,0]/255;
    %yellow3 = [196,200,21]/255;
    set(ph1,'Color',yellow1)        %measured
    set(ph2,'Color',yellow2)        %predicted1
end
```
