## Supplementary material for "Predicting Composition of Genetic Circuits with Resource Competition: Demand and Sensitivity": Experimental data and Matlab code: predictRS.m

```
function outstruct = predictRS(datastruct,FPstruct,Jstruct,FPfields,ploton,figharray)

if nargin < 5 || isempty(ploton)
    ploton = true;
end
if ploton && nargin < 6
    figharray = 4:6;    %list of figure handles to plot to, must be same length as FPfields
end

outstruct = datastruct;
FPnames = fieldnames(FPstruct);
names = cell(length(FPnames),3);
datamat = zeros(length(FPnames),3);

for ii = 1:length(FPnames)
    
    %get metadata from FPstruct about pairwise experiments
    cellalone1 = FPstruct.(FPnames{ii}).alone1;
    cellalone2 = FPstruct.(FPnames{ii}).alone2;
    cellperturb = FPstruct.(FPnames{ii}).perturb;
    FPfield1 = FPstruct.(FPnames{ii}).FPfield1;
    FPfield2 = FPstruct.(FPnames{ii}).FPfield2;
    
    %get J's for predictions using least squares estimates
    J1 = Jstruct.([cellalone1,'_l_s']).J0mc;
    J2 = Jstruct.([cellalone2,'_l_s']).J0mc;
    
    time = datastruct.(cellalone1).time;
    
    %get data
    %constant
    dataalone1c = mean(datastruct.(cellalone1).(FPfield1))*ones(size(time));
    predalone1c = dataalone1c;
    predperturb1c = applyRSpred(dataalone1c,J1,J2);
    %constant
    dataalone2c = mean(datastruct.(cellalone2).(FPfield2))*ones(size(time));
    predalone2c = dataalone2c;
    predperturb2c = applyRSpred(dataalone2c,J2,J1);
    
    %measured perturbation
    mean(datastruct.(cellperturb).(FPfield1));
    mean(datastruct.(cellperturb).(FPfield2));
    
    %datamat2(ii,:) = []
    datamat(ii,:) = [dataalone1c,dataalone2c,predperturb2c];
    names(ii,:) = {cellalone1,cellalone2,cellperturb};
    
    %write output
    outstruct.(cellalone1).([FPfield1,'predc']) = predalone1c;
    outstruct.(cellalone2).([FPfield2,'predc']) = predalone2c;
    outstruct.(cellperturb).([FPfield1,'predc']) = predperturb1c;
    outstruct.(cellperturb).([FPfield2,'predc']) = predperturb2c;
    
    [FPnames{ii},'; ',FPfield1,'; ',FPfield2];
    [J1,J2,applyRSpred(1,J1,J2),applyRSpred(1,J2,J1)];
end

if ploton
    for k = 1:length(FPfields)
        if contains(FPfields{k},'BFP')
            celllist = {'B','BG','BR'};
        elseif contains(FPfields{k},'GFP')
            celllist = {'G','BG','GR'};
        elseif contains(FPfields{k},'RFP')
            celllist = {'R','BR','GR'};
        end
        datafieldpred = [FPfields{k},'_sspredc'];
        %plot data
        figh = plotsubfield2(outstruct,'time',FPfields{k},figharray(k),celllist);
        setallsubplots(figh,'axis','nextplot','add');
        %plot predicted
        plotsubfield2(outstruct,'time',datafieldpred,figh,celllist);
    end
end


function pred = applyRSpred(data,J1,J2)
pred = data*(1+J1)./(1+J1+J2);
```
