## Supplementary material for "Predicting Composition of Genetic Circuits with Resource Competition: Demand and Sensitivity": Experimental data and Matlab code: predODvstime.m

```
function predODvstime(modstruct,cellstruct,GRfitparams,ploton)


modnames = fieldnames(modstruct);

for ii = 1:length(modnames)
    
    Q = mean(modstruct.(modnames{ii}).Q);
    
    predalpha = mean(modstruct.(modnames{ii}).y,2);
    predGR = GRfitmodel(Q,GRfitparams);
    
    
    
    predFPOD = (predalpha./predGR)*(1 - exp(-predGR*t));
    
end


function gr = GRfitmodel(Q,GRfitparams)

gr = 0.7./(0.2*(1+Q) + 1);
```
