## Supplementary material for "Predicting Composition of Genetic Circuits with Resource Competition: Demand and Sensitivity": Experimental data and Matlab code: readplate2.m

```
function datastruct = readplate2(filename, infostruct)
%datastruct = readplate2(filename, infostruct)
%read in data from platereader and seperate into fields. Infostruct should
%have field names corresponding to properties measured by flow cytometer.
%Each field should correspond to a range in the excel file containing the
%corresponding data formatted as a string.

if nargin < 2
    infostruct = struct();
    infostruct.OD = 'B92:BK891';
    infostruct.GFP = 'B896:BK1695';
    infostruct.RFP = 'B1700:BK2499';
    infostruct.YFP = 'B2504:BK3303';
    if nargin < 1
        filename = 'Calibration - J0 characterization\5th experiment to calibrate GYR_new.xlsx';
    end
end

datastruct = struct;

root = pwd;
cd('..\experiments')
disp('loading data...')
if contains(filename,filesep)
    disp(filename(strfind(filename,filesep)+1:end))
else
    disp(filename)
end
datameasured = fieldnames(infostruct);
for ii = 1:length(datameasured)
    data = xlsread(filename,1,infostruct.(datameasured{ii}));
    if sum(sum(isnan(data))) > 0
        warning('dataset has NaN''s')
    end
    datastruct.(datameasured{ii}) = data(:,3:end);
    datastruct.([datameasured{ii},'time']) = data(:,1)/3600;
    %datastruct.([datameasured{ii},'temp']) = data(:,2);
end
cd(root);
pause(.1);
```
