## Supplementary material for "Predicting Composition of Genetic Circuits with Resource Competition: Demand and Sensitivity": Experimental data and Matlab code: runplate_RS3.m

```
%could write a class for this??
addpath utility

%experiment settings
%match wells on plate with experimental conditions
%change from .txt files to .mat files
importplate('J0_exp3_9-25-2020.txt')
filenamecell = {'J0_exp3_9-25-2020.mat'};
% %match wells on plate with experimental conditions
indstruct = cell(length(filenamecell),1);
indstruct{1}.B = [2,2; 2,3; 2,4];
indstruct{1}.G = [3,2; 3,3; 3,4];
indstruct{1}.R = [4,2; 4,3; 4,4];
indstruct{1}.BG = [5,2; 5,3; 5,4];
indstruct{1}.BR = [6,2; 6,3; 6,4];
indstruct{1}.GR = [7,2; 7,3; 7,4];
indstruct{1}.M9 = [2,5; 3,5; 4,5];
%windows where each cell type are in steady state
ssfun = @mean;
ssfields = {'BFPdiffOD','GFPdiffOD','RFPdiffOD'};
% ssstruct.B = [6.1,7.8];
% ssstruct.G = [4.0,5.8];
% ssstruct.R = [6.0,7.1];
% ssstruct.BG = [6.5,8.0];
% ssstruct.BR = [5.1,7.8];
% ssstruct.GR = [4.8,6.0];
ssstruct.B =  [4.5,5.3];
ssstruct.G =  [1.6,4.5];
ssstruct.R =  [1.5,4.6];
ssstruct.BG = [2.5,6.0];
ssstruct.BR = [5.1,7.8];
ssstruct.GR = [1.9,4.7];

%% shouldn't need to change these much
%tell which fields should be zero to subtract background
nonestruct = struct;
nonestruct.OD = {'M9'};
nonestruct.BFP = {'G','R','GR'};
nonestruct.GFP = {'B'};%,'R','BR'};
nonestruct.RFP = {'B','G','BG'};

%specify how to combine steady state fluorescence measurements to find RSD0s
%!for Resource sensor modules only
nummods = 2;            %number of modules for least squares
FPstruct = struct;
FPstruct.BG.alone1 = 'B';      FPstruct.BG.FPfield1 = 'BFPdiffOD_ss';
FPstruct.BG.alone2 = 'G';      FPstruct.BG.FPfield2 = 'GFPdiffOD_ss';
FPstruct.BG.perturb = 'BG';
FPstruct.BG.RSDinds = [1,2];             %for least squares
% FPstruct.BR.alone1 = 'B';      FPstruct.BR.FPfield1 = 'BFPdiffOD_ss';
% FPstruct.BR.alone2 = 'R';      FPstruct.BR.FPfield2 = 'RFPdiffOD_ss';
% FPstruct.BR.perturb = 'BR';
% FPstruct.BR.RSDinds = [1,3];
% FPstruct.GR.alone1 = 'G';      FPstruct.GR.FPfield1 = 'GFPdiffOD_ss';
% FPstruct.GR.alone2 = 'R';      FPstruct.GR.FPfield2 = 'RFPdiffOD_ss';
% FPstruct.GR.perturb = 'GR';
% FPstruct.GR.RSDinds = [2,3];             %index of resource demand coefficient

%need to get steady state data from module alone, module with RS, RS with module, RS alone

%plotting limits
tlims = [4,9];
ODlims = [0,0.1];
BFPlims = [0,7e3];
GFPlims = [0,4e4];
RFPlims = [0,7e3];
BFPODlims = [0,5e6];
GFPODlims = [0,4e5];
RFPODlims = [0,1e5];
BFPdiffODlims = [0,12e5];
GFPdiffODlims = [0,2e5];
RFPdiffODlims = [0,4e4];

%% analyze data
%init
[datastruct,cellstruct,basalstruct] = deal(cell(length(filenamecell),1));
timeprev = 0;

%loop through each dilution
for ii = 1:length(filenamecell)
    %load data
    load(filenamecell{ii})
    %rename fields
    platedata.OD = platedata.OD600_600;
    platedata.RFP = platedata.RFP1_584_619; %RFP1_584_607;
    platedata.GFP = platedata.GFP1_485_530; %GFP1_465_498;
    platedata.BFP = platedata.GFP1_400_460; %GFP1_518_540;
    %format into struct
    [datastruct{ii},timeprev] = addtime(platedata,timeprev);
    %low pass filter data
    datastruct{ii} = filtplatedata2(datastruct{ii},0.8);
    %format into substructs
    cellstruct{ii} = bulkdata2cells(datastruct{ii},indstruct{ii},{'GFP','RFP','BFP','OD','time'});
    %subtract background fluorescence and OD levels
    %NOTE: background fluoresence changes as cells use up metabolites/sugars
    %[cellstruct{ii}, basalstruct{ii}] = subtractbasal3(cellstruct{ii},...
    %   nonefield,nonefield,ODeps,FPeps);
    [cellstruct{ii}, basalstruct{ii}] = subtractbasal4(cellstruct{ii},nonestruct);
    %correct for fluorescence bleed between channels
    %cellstruct{ii} = correctFP(cellstruct{ii},{'G','R','B'},{'GFP','RFP','BFP'});
    %find smoothed growthrate for cells
    %cellstruct{ii} = fitgrowthrate2(cellstruct{ii},'logisticfit','OD',true,false,1);
    %cellstruct{ii} = growthrate(cellstruct{ii},'OD');
    %differentiate raw fluorescence with respect to time
    cellstruct{ii} = diffcellstruct(cellstruct{ii},{'BFP','RFP','GFP'},'time');
end

%%
%combine batches into one
%cellstruct2 = combinecellrepeats(cellstruct);
cellstruct2 = cellstruct{1};
%divide fluorescence by OD and growthrate
cellstruct2 = normalizeFP2(cellstruct2);
%average over time intervals to get steady state
cellstruct2 = findss3(cellstruct2,ssstruct,ssfields,ssfun,true,[2,3,4]);
%calculate resource demand for resource sensor
RSstruct = calcRSall(cellstruct2,FPstruct,nummods,10000,true,5);

%make predictions of combos from measurements alone
out = predictRS2(cellstruct2,FPstruct,RSstruct,true,6);

return

%average across all replicates
[~,figh] = avetrials(cellstruct2,{'BFPdiffOD','GFPdiffOD','RFPdiffOD'},...
    'time',true,1,{'B','G','R','BG','BR','GR'});
setallsubplots(1,'axis','YLim',[0,max([BFPdiffODlims;GFPdiffODlims;RFPdiffODlims],[],'all')]);
setallsubplots(1,'axis','XLim',tlims);

% %color correction
% plotsubfield2(cellstruct2,'GFP','BFP',19);
% plotsubfield2(cellstruct2,'GFP','RFP',20);
% plotsubfield2(cellstruct2,'RFP','BFP',21);

%%

%time vs OD
plotsubfield2(cellstruct2,'time','OD',4,{'B','G','R','BG','BR','GR'});
setallsubplots([],'axis',{'Xlim','YLim'},{tlims,ODlims});
% %time vs FP/OD
plotsubfield2(cellstruct2,'time','BFPOD',7,{'B','BG','BR'});
setallsubplots(7,'axis',{'Xlim','YLim'},{tlims,BFPODlims});
plotsubfield2(cellstruct2,'time','GFPOD',8,{'G','BG','GR'});
setallsubplots([],'axis',{'Xlim','YLim'},{tlims,GFPODlims});
plotsubfield2(cellstruct2,'time','RFPOD',9,{'R','BR','GR'});
setallsubplots([],'axis',{'Xlim','YLim'},{tlims,RFPODlims});

%fluorescence bleed
plotsubfield2(cellstruct2,'OD','BFP',8,{'G','R','GR'});
%setallsubplots([],'axis',{'Xlim','YLim'},{tlims});
plotsubfield2(cellstruct2,'OD','GFP',9,{'B','R','BR'});
%setallsubplots([],'axis',{'Xlim','YLim'},{tlims});
plotsubfield2(cellstruct2,'OD','RFP',10,{'B','G','BG'});
%setallsubplots([],'axis',{'Xlim','YLim'},{tlims});

%time vs dot{FP}/OD
plotsubfield2(cellstruct2,'time','BFPdiffOD',11,{'B','BG','BR'});
setallsubplots([],'axis','nextplot','add')
plotsubfield2(cellstruct2,'time','BFPdiffODpred',11,{'B','BG','BR'});
%setallsubplots([],'line',{'LineStyle','Marker'},{'none','.'});
setallsubplots([],'line',{'Linewidth'},{1});
setallsubplots([],'axis',{'Ylim','Xlim'},{BFPdiffODlims,tlims});
plotsubfield2(cellstruct2,'time','GFPdiffOD',12,{'G','BG','GR'});
%setallsubplots([],'line',{'LineStyle','Marker'},{'none','.'});
setallsubplots([],'line',{'Linewidth'},{1});
setallsubplots([],'axis',{'Ylim','Xlim'},{GFPdiffODlims,tlims});
plotsubfield2(cellstruct2,'time','RFPdiffOD',13,{'R','BR','GR'});
%setallsubplots([],'line',{'LineStyle','Marker'},{'none','.'});
setallsubplots([],'line',{'Linewidth'},{1});
setallsubplots([],'axis',{'Ylim','Xlim'},{RFPdiffODlims,tlims});

%% notes:
%protease degradation?
%lack of linear binding assumption causing extra correlations?

%%model assumptions:
%shared resources
%no post transcriptional/translational resource modifications
%cells in exponential growth
%total resources constant


%% archive
%change from .txt files to .mat files
% importplate('data 2-4-20\J0_exp_2-4-2020_1.txt')
% filenamecell = {'data 2-4-20\J0_exp_2-4-2020_1.mat'};
% % %match wells on plate with experimental conditions
% indstruct = cell(length(filenamecell),1);
% indstruct{1}.B = [2,2; 2,3; 2,4];
% indstruct{1}.G = [3,2; 3,3; 3,4];
% indstruct{1}.R = [4,2; 4,3; 4,4];
% indstruct{1}.BG = [5,2; 5,3; 5,4];
% indstruct{1}.BR = [6,2; 6,3; 6,4];
% indstruct{1}.GR = [7,2; 7,3; 7,4];
% indstruct{1}.M9 = [2,5; 3,5; 4,5];
% %steady state
% ssstruct.B = [6.1,7.8];
% ssstruct.G = [4.0,5.8];
% ssstruct.R = [6.0,7.1];
% ssstruct.BG = [6.5,8.0];
% ssstruct.BR = [5.1,7.8];
% ssstruct.GR = [4.8,6.0];

% %experiment settings
% importplate('data 2-15-20\J0_exp_2-15-2020_2_short.txt')
% filenamecell = {'data 2-15-20\J0_exp_2-15-2020_2.mat'};
% %match wells on plate with experimental conditions
% indstruct = cell(length(filenamecell),1);
% indstruct{1}.B = [2,4; 2,5; 2,6; 2,7; 2,8];
% indstruct{1}.G = [3,4; 3,5; 3,6; 3,7; 3,8];
% indstruct{1}.R = [4,4; 4,5; 4,6; 4,8];      %4,7
% indstruct{1}.BG = [5,4; 5,5; 5,6; 5,7; 5,8];
% indstruct{1}.BR = [6,4; 6,5; 6,6; 6,7; 6,8];
% indstruct{1}.GR = [7,4; 7,5; 7,6; 7,7; 7,8];
% indstruct{1}.M9 = [2,3; 3,3; 4,3; 2,9; 3,9; 4,9];
% %steady state time ranges
% ssstruct.B = [4.4,6.0];     %6.3?
% ssstruct.G = [2.8,4.0];
% ssstruct.R = [3.5,4.6];
% ssstruct.BG = [3.5,6.0];
% ssstruct.BR = [4.5,6.0];    %7.0?
% ssstruct.GR = [3.1,5.0];
```
