## Supplementary material for "Predicting Composition of Genetic Circuits with Resource Competition: Demand and Sensitivity": Experimental data and Matlab code: smooth_diff.m

```
function h=smooth_diff(n)
% A smoothed differentiation filter (digital differentiator). 
%
% Such a filter has the following advantages:
% 
% First, the filter involves both the smoothing operation and differentation operation. 
% It can be regarded as a low-pass differention filter (digital differentiator). 
% It is well known that the common differentiation operation amplifies the high-frequency noises.
% Therefore, the smoothded differentiation filter would be valuable in experimental (noisy) data processing. 
% 
% Secondly, the filter coefficients are all convenient integers (simple units) except for an integer scaling factor,
% as may be especially significant in some applications such as those in some single-chip microcomputers
% or digital signal processors. 
% 
% Usage:
% h=smooth_diff(n)
% n: filter length (positive integer larger no less than 2)
% h: filter coefficients (anti-symmetry)
%
% Examples:
% smooth_demo
%
% Author:
% Jianwen Luo <> 2004-11-02
% Department of Biomedical Engineering, Department of Electrical Engineering
% Tsinghua University, Beijing 100084, P. R. China  
% 
% References:
% Usui, S.; Amidror, I., 
% Digital Low-Pass Differentiation for Biological Signal-Processing. 
% IEEE Transactions on Biomedical Engineering 1982, 29, (10), 686-693.
% Luo, J. W.; Bai, J.; He, P.; Ying, K., 
% Axial strain calculation using a low-pass digital differentiator in ultrasound elastography. 
% IEEE Transactions on Ultrasonics Ferroelectrics and Frequency Control 2004, 51, (9), 1119-1127.

if n>=2 && floor(n)==ceil(n)
    if mod(n,2)==1%is odd
        m=fix((n-1)/2);
        h=[-ones(1,m) 0 ones(1,m)]/m/(m+1);
    else%is even
        m=fix(n/2);
        h=[-ones(1,m) ones(1,m)]/m^2;
    end
else    
    error('The input parameter (n) should be a positive integer larger no less than 2.');
end
```
