## Supplementary material for "Predicting Composition of Genetic Circuits with Resource Competition: Demand and Sensitivity": Experimental data and Matlab code: smooth_diff_demo.m

```
k=1/(2*sqrt(log(2)));
x=0:30/1999:30; 
a1=1.0; t1=5; w1=1.3;
y1=a1*exp(-((x-t1)/(w1*k)).^2);
a2=1.0; t2=15.0; w2=1.3;
y2=a2*(1+4*((x-t2)/w2).^2).^(-1);
a3=1.0; t3=25;
y3=a3*ones(size(x))./(1+exp(-3.0*(x-t3)));
y=y1+y2+y3;
noise=rand(size(y));
y=y+(noise-0.5)*0.01;

figure
subplot(3,1,1)
plot(x,y);
title('noisy data');

subplot(3,1,2)
plot(x(1:end-1)+(x(2)-x(1)),diff(y)/(x(2)-x(1)));
title('diff function');

subplot(3,1,3)
plot(x,filter(-smooth_diff(10),1,y)/(x(2)-x(1)));
title('smoothed differentiation');
```
