## Supplementary material for "Predicting Composition of Genetic Circuits with Resource Competition: Demand and Sensitivity": Experimental data and Matlab code: smoothdiff.m

```
function [dfdh,dfdh2,dfdh3] = smoothdiff(f,h)
%dfdh2 = smoothdiff(f,h)
%take smooth derivative of f with respect to h

%find step size
if length(h) > 1
    h2 = abs(mean(h(2:end) - h(1:end-1)));
else; h2 = h;
end

%Y = filter(B,A,X)
%a(1)*y(n) = b(1)*x(n) + b(2)*x(n-1) + ... + b(nb+1)*x(n-nb)
%                - a(2)*y(n-1) - ... - a(na+1)*y(n-na)
% 9-pt Savitzky-Golay filtered first derivative (1st/2nd order polynomial fit)
a = 60*h2;
b = [-4,-3,-2,-1,0,1,2,3,4];
dfdh = filter(-b,a,f);
%this gives best performance for y = 3*sin(x) + 0.1*randn(size(x));
%OR y = K*P0*exp(r*x)./(K+P0*(exp(r*x)-1)) + 0.01*randn(size(x));
%OR y = 3*x + 0.1*randn(size(x));

% 9-pt Savitzky-Golay filtered first derivative (3rd/4th order polynomial fit)
a2 = 1188*h2;
b2 = [86,-142,-193,-126,0,126,193,142,-86];
dfdh2 = filter(-b2,a2,f);

% %Savitzky-Golay filtered first derivative (equialent to the 5 pt stencil)
% %window size: 5 points, cubic polynomial
% f2 = [f(4,:); f(3,:); f(2,:); f; f(end-1,:); f(end-2,:); f(end-3,:)];
% %dfdh = (f2(1:end-4,:) - 8*f2(2:end-3,:) + 8*f2(4:end-1,:) - f2(5:end,:))/(12*h2);
% %window size: 7 pts, cubic polynomial
% dfdh = (22*f2(1:end-6,:) - 67*f2(2:end-5,:) - 58*f2(3:end-4,:) ...
%     + 58*f2(5:end-2,:) + 67*f2(6:end-1,:) - 22*f2(7:end,:))/(252*h2);


% moving average differentiator
dfdh3 = filter(-smooth_diff(9),1,f)./h2;


% %five point stencil
% %append 0's to beginning of f so output is same size as input
% f2 = [f(3,:); f(2,:); f; f(end-1,:); f(end-2,:)];
% %5 point stencil differentiation
% dxdt = (f2(1:end-4,:) - 8*f2(2:end-3,:) + 8*f2(4:end-1,:) - f2(5:end,:))/(12*h2);
% %pad with zero so output is same size as input
% dxdt = [dxdt;zeros(1,size(f,2))];
```
