## Supplementary material for "Predicting Composition of Genetic Circuits with Resource Competition: Demand and Sensitivity": Experimental data and Matlab code: structfun2.m

```
function structout = structfun2(funh, structin, datafields)
%structout = structfun2(funh, structin, datafields)
%apply FUNH to each field of STRUCTIN and returns a structure with the same
%fields. If DATAFIELDS is given, apply FUNH only to fields matching the
%strings in the cell array DATAFIELDS.

fields = fieldnames(structin);
structout = struct;


for ii = 1:length(fields)
    if isstruct(structin.(fields{ii}))
        structout.(fields{ii}) = structfun2(funh, structin.(fields{ii}));
    else
        if nargin < 3 || isempty(datafields)
            structout.(fields{ii}) = funh(structin.(fields{ii}));
        else
            %only apply funh to desired bottom-level fields in datafields
            if any(strcmpi(fields{ii},datafields))
                structout.(fields{ii}) = funh(structin.(fields{ii}));
            end
        end
    end
end
```
