## Supplementary material for "Predicting Composition of Genetic Circuits with Resource Competition: Demand and Sensitivity": Experimental data and Matlab code: subsampleerrorbar.m

```
function [hp,he] = subsampleerrorbar(x,y,yerr,maxerrorbars)

if nargin < 4
    maxerrorbars = 100;
end

%if data is too large only plot MAXERRORBARS number of error bars
if length(yerr) > maxerrorbars
    n = round(length(yerr)/maxerrorbars);
    yerr2 = yerr(1:n:end);
    y2 = y(1:n:end);
    x2 = x(1:n:end);
else
    y2 = y;
    x2 = x;
    yerr2 = yerr;
end

%plot error bars
coi = get(gca,'colororderindex');
hp = plot(x,y,'linewidth',2); hold on
set(gca,'colororderindex',coi);
he = errorbar(x2,y2,yerr2,'.','linewidth',0.5);
```
