## Supplementary material for "Predicting Composition of Genetic Circuits with Resource Competition: Demand and Sensitivity": Experimental data and Matlab code: subtractbasal2.m

```
function [newstruct,basalstruct] = subtractbasal2(cellstruct,ODeps,FPeps,nonefld)
%[newstruct,basalstruct] = subtractbasal2(cellstct,ODeps,FPeps,nonefld)
%find basal level for each property by assuming initial conditition should
%be 0 for all properties

if nargin < 4
    nonefld = 'M9';
    if nargin < 3
        FPeps = 0;
        if nargin < 2
            ODeps = 0;
        end
    end
end

%init
newstruct = struct;
cellnames = fieldnames(cellstruct);
%find basal levels
basalstruct = findbasallevel2(cellstruct,ODeps,FPeps,20);

for ii = 1:length(cellnames)
    dataprops = fieldnames(cellstruct.(cellnames{ii}));
    %loop across each cell type
    for jj = 1:length(dataprops)
        %loop across each data property and subtract off basal level
        if strcmp(dataprops{jj},'OD')
            newstruct.(cellnames{ii}).(dataprops{jj}) = ...
                cellstruct.(cellnames{ii}).(dataprops{jj}) - ...
                mean(basalstruct.(nonefld).(dataprops{jj}));
        else
            newstruct.(cellnames{ii}).(dataprops{jj}) = ...
                cellstruct.(cellnames{ii}).(dataprops{jj}) - ...
                basalstruct.(cellnames{ii}).(dataprops{jj});
        end
    end
end


function basalstruct = findbasallevel(nonestruct,ODeps,FPeps)
%find basal levels of each property based on media measurements

basalstruct = struct;
dataprops = fieldnames(nonestruct);

for ii = 1:length(dataprops)
    if contains(dataprops{ii},'OD')
        basalstruct.(dataprops{ii}) = ...
            mean(mean(nonestruct.(dataprops{ii})(1:100,:)),2) - ODeps;
    elseif contains(dataprops{ii},'FP')
        basalstruct.(dataprops{ii}) = ...
            mean(nonestruct.(dataprops{ii}),2) - FPeps;
    elseif contains(dataprops{ii},'time') || contains(dataprops{ii},'temp')
        basalstruct.(dataprops{ii}) = 0;
    else
        basalstruct.(dataprops{ii}) = ...
            mean(nonestruct.(dataprops{ii}),2);
    end
end


function basalstruct = findbasallevel2(cellstruct,ODeps,FPeps,n)
%find basal levels of each property based on media measurements

basalstruct = struct;
dprops = fieldnames(cellstruct);

for ii = 1:length(dprops)
    if isstruct(cellstruct.(dprops{ii}))
        basalstruct.(dprops{ii}) = findbasallevel2(cellstruct.(dprops{ii}),ODeps,FPeps,n);
    else
        if contains(dprops{ii},'OD')
            basalstruct.(dprops{ii}) = mean(cellstruct.(dprops{ii})(1:n,:),1) - ODeps;
        elseif contains(dprops{ii},'FP')
            basalstruct.(dprops{ii}) = mean(cellstruct.(dprops{ii})(1:n,:),1) - FPeps;
        elseif contains(dprops{ii},'time') || contains(dprops{ii},'temp')
            basalstruct.(dprops{ii}) = 0;
        else
            basalstruct.(dprops{ii}) = mean(cellstruct.(dprops{ii})(1:n,:),1);
        end
    end
end
```
