## Supplementary material for "Predicting Composition of Genetic Circuits with Resource Competition: Demand and Sensitivity": Experimental data and Matlab code: subtractbasal3.m

```
function [newstruct,basalstruct] = subtractbasal3(cellstruct,nonefld,mediafld,ODeps,FPeps)
%[newstruct,basalstruct] = subtractbasal3(cellstct,nonefld,mediafld,ODeps,FPeps)
%find basal level for each property by assuming initial conditition should
%be 0 for all properties

if nargin < 5
    ODeps = 0;
    if nargin < 4
        FPeps = 0;
        if nargin < 3
            mediafld = 'M9';
            if nargin < 2
                nonefld = 'T10';
            end
        end
    end
end

%init
newstruct = struct;
cellnames = fieldnames(cellstruct);

%find basal levels across time
if all(isfield(cellstruct,{nonefield,mediafld}))
    basalstruct = findbasallevel3(cellstruct.(nonefld),cellstruct.(mediafld),ODeps,FPeps);
elseif isfield(cellstruct,nonefield)
    basalstruct = findbasallevel3(cellstruct.(nonefld),cellstruct.(nonefld),ODeps,FPeps);
elseif isfield(cellstruct,mediafld)
    basalstruct = findbasallevel3(cellstruct.(mediafld),cellstruct.(mediafld),ODeps,FPeps);
else
    basalstruct = findbasalnomedia(cellstruct,ODeps,FPeps);
end

%subtract basal levels off data
for ii = 1:length(cellnames)
    dataprops = fieldnames(cellstruct.(cellnames{ii}));
    %loop across each cell type
    for jj = 1:length(dataprops)
        %loop across each data property and subtract off basal level
        if iscell(cellstruct.(cellnames{ii}).(dataprops{jj}))
            newstruct.(cellnames{ii}).(dataprops{jj}) = ...
                cellfun(@(x,y) x - y,cellstruct.(cellnames{ii}).(dataprops{jj}),...
                basalstruct.(dataprops{jj}),'UniformOutput',false);
        else
            newstruct.(cellnames{ii}).(dataprops{jj}) = ...
                cellstruct.(cellnames{ii}).(dataprops{jj}) - ...
                basalstruct.(dataprops{jj});
        end
    end
end

function basalstruct = findbasallevel3(nonestruct,mediastruct,ODeps,FPeps)
%find basal levels of each property based on media measurements

basalstruct = struct;
dprops = fieldnames(nonestruct);

for ii = 1:length(dprops)
    if isstruct(nonestruct.(dprops{ii}))
        basalstruct.(dprops{ii}) = findbasallevel3(nonestruct.(dprops{ii}),ODeps,FPeps);
    else
        %find average values for each property taking the average of each
        %column or each cell
        if contains(dprops{ii},'OD')
            %if property is OD
            if iscell(mediastruct.(dprops{ii}))
                basalstruct.(dprops{ii}) = cellfun(@(x) mean(x,2)-ODeps,...
                    mediastruct.(dprops{ii}),'UniformOutput',false);
            else
                basalstruct.(dprops{ii}) = mean(mediastruct.(dprops{ii}),2) - ODeps;
            end
        elseif contains(dprops{ii},'FP')
            %if property is fluorescence
            if iscell(nonestruct.(dprops{ii}))
                basalstruct.(dprops{ii}) = cellfun(@(x) mean(x,2)-FPeps,...
                    nonestruct.(dprops{ii}),'UniformOutput',false);
            else
                basalstruct.(dprops{ii}) = mean(nonestruct.(dprops{ii}),2) - FPeps;
            end
        elseif contains(dprops{ii},'time') || contains(dprops{ii},'temp')
            %if property is time
            if iscell(nonestruct.(dprops{ii}))
                basalstruct.(dprops{ii}) = repmat({0},1,size(nonestruct.(dprops{ii}),2));
            else
                basalstruct.(dprops{ii})  = 0;
            end
        else
            %if property is something else
            if iscell(nonestruct.(dprops{ii}))
                basalstruct.(dprops{ii}) = cellfun(@(x) mean(x,2),...
                    nonestruct.(dprops{ii}),'UniformOutput',false);
            else
                basalstruct.(dprops{ii}) = mean(nonestruct.(dprops{ii}),2);
            end
        end
    end
end


function basalstruct = findbasalnomedia(cellstruct,ODeps,FPeps)
%find basal levels of each property based on average of first n samples

n = 5;

basalstruct = struct;
cellnames = fieldnames(cellstruct);

for jj = 1:length(cellnames)
    dprops = fieldnames(cellstruct.(cellnames{jj}));
    for ii = 1:length(dprops)
        %basalstruct.(cellnames{jj}).(dprops{ii}) = findbasallevel3(nonestruct.(dprops{ii}),ODeps,FPeps,n);
        %find average values for each property taking the average of each
        %column or each cell
        if contains(dprops{ii},'OD')
            %if property is OD
            if iscell(cellstruct.(cellnames{jj}).(dprops{ii}))
                basalstruct.(dprops{ii}) = cellfun(@(x) mean(x,2)-ODeps,...
                    mediastruct.(dprops{ii}),'UniformOutput',false);
            else
                basalstruct.(dprops{ii}) = mean(mediastruct.(dprops{ii}),2) - ODeps;
            end
        elseif contains(dprops{ii},'FP')
            %if property is fluorescence
            if iscell(nonestruct.(dprops{ii}))
                basalstruct.(dprops{ii}) = cellfun(@(x) mean(x,2)-FPeps,...
                    nonestruct.(dprops{ii}),'UniformOutput',false);
            else
                basalstruct.(dprops{ii}) = mean(nonestruct.(dprops{ii}),2) - FPeps;
            end
        elseif contains(dprops{ii},'time') || contains(dprops{ii},'temp')
            %if property is time
            if iscell(nonestruct.(dprops{ii}))
                basalstruct.(dprops{ii}) = repmat({0},1,size(nonestruct.(dprops{ii}),2));
            else
                basalstruct.(dprops{ii})  = 0;
            end
        else
            %if property is something else
            if iscell(nonestruct.(dprops{ii}))
                basalstruct.(dprops{ii}) = cellfun(@(x) mean(x,2),...
                    nonestruct.(dprops{ii}),'UniformOutput',false);
            else
                basalstruct.(dprops{ii}) = mean(nonestruct.(dprops{ii}),2);
            end
        end
    end
end


%---------------------------------------------------
%not used
function basalstruct = findbasallevel(cellstruct,ODeps,FPeps)
%find basal levels of each property based on media measurements

basalstruct = struct;
dprops = fieldnames(cellstruct);

for ii = 1:length(dprops)
    if isstruct(cellstruct.(dprops{ii}))
        basalstruct.(dprops{ii}) = findbasallevel(cellstruct.(dprops{ii}),ODeps,FPeps);
    else
        if contains(dprops{ii},'OD')
            basalstruct.(dprops{ii}) = ...
                mean(cellstruct.(dprops{ii}),2) - ODeps;
        elseif contains(dprops{ii},'FP')
            basalstruct.(dprops{ii}) = ...
                mean(cellstruct.(dprops{ii}),2) - FPeps;
        elseif contains(dprops{ii},'time') || contains(dprops{ii},'temp')
            basalstruct.(dprops{ii}) = 0;
        else
            basalstruct.(dprops{ii}) = ...
                mean(cellstruct.(dprops{ii}),2);
        end
    end
end


function basalstruct = findbasallevel2(cellstruct,ODeps,FPeps,n)
%find basal levels of each property based on media measurements

basalstruct = struct;
dprops = fieldnames(cellstruct);

for ii = 1:length(dprops)
    if isstruct(cellstruct.(dprops{ii}))
        basalstruct.(dprops{ii}) = findbasallevel2(cellstruct.(dprops{ii}),ODeps,FPeps,n);
    else
        if contains(dprops{ii},'OD')
            basalstruct.(dprops{ii}) = mean(cellstruct.(dprops{ii})(1:n,:),1) - ODeps;
        elseif contains(dprops{ii},'FP')
            basalstruct.(dprops{ii}) = mean(cellstruct.(dprops{ii})(1:n,:),1) - FPeps;
        elseif contains(dprops{ii},'time') || contains(dprops{ii},'temp')
            basalstruct.(dprops{ii}) = 0;
        else
            basalstruct.(dprops{ii}) = mean(cellstruct.(dprops{ii})(1:n,:),1);
        end
    end
end
```
