## Supplementary material for "Predicting Composition of Genetic Circuits with Resource Competition: Demand and Sensitivity": Experimental data and Matlab code: subtractbasal4.m

```
function [outstruct,basalstruct] = subtractbasal4(cellstruct,nonestruct,ODfld,subfields)
%[newstruct,basalstruct] = subtractbasal4(cellstct,nonefld,mediafld,ODeps,FPeps)
%find basal level for each property by assuming initial conditition should
%be 0 for all properties

%make curves of basal fluorescence vs OD for each channel
%average across replicates
%subtract off average basal levels from raw data

if nargin < 3 || isempty(ODfld)
    %field for OD
    ODfld = 'OD';
end
if nargin < 2
    %tells the cells/wells where there should be no expression
    %for each datafield
    nonestruct = struct;
    nonestruct.OD = {'M9'};
    nonestruct.BFP = {'G','R','GR'};
    nonestruct.GFP = {'B','R','BR'};
    nonestruct.RFP = {'B','G','BG'};
end
if nargin < 4 || isempty(subfields)
    subfields = fieldnames(nonestruct);
end

%init
outstruct = struct;
cellnames = fieldnames(cellstruct);

%find basal levels as a function of OD for each channel
basalstruct = findbasallevel4(cellstruct,nonestruct,ODfld);
% basalstruct = findbasalnomedia(cellstruct,ODeps,FPeps);

%subtract basal levels off data
for ii = 1:length(cellnames)
    
    data = cellstruct.(cellnames{ii});
    %loop across each cell type
    dprops = fieldnames(data);
    
    %get OD curve for specific measurement
    ODdata = data.(ODfld);
    
    for jj = 1:length(dprops)
        %if matches desired fields, subtract fields
        if any(strcmpi(dprops{jj},subfields))
            if strcmp(dprops{jj},ODfld)
                %if OD, subtract off basal level from media
                outobj.(dprops{jj}) = data.(dprops{jj}) - basalstruct.(dprops{jj});
            else
                %if FP, loop across each data property and subtract off basal
                %level, dependent on OD
                FPODfun = basalstruct.(dprops{jj});
                outobj.(dprops{jj}) = data.(dprops{jj}) - FPODfun(ODdata);
            end
        else
            %copy remaining data
            outobj.(dprops{jj}) = data.(dprops{jj});
        end
    end
    outstruct.(cellnames{ii}) = outobj;
end


function basalstruct = findbasallevel4(cellstruct,nonestruct,ODfld)
%find basal levels of each property based on media measurements

basalstruct = struct;
dprops = fieldnames(nonestruct);

%loop across properties
for ii = 1:length(dprops)
    %split if FP
    if ~strcmpi(dprops{ii},ODfld)
        %make curve of basal FP vs OD
        cellnamesnone = nonestruct.(dprops{ii});
        FPtmp = [];
        ODtmp = [];
        for jj = 1:length(cellnamesnone)
            %get FP data
            FPtmp0 = cellstruct.(cellnamesnone{jj}).(dprops{ii});
            FPtmp = [FPtmp,FPtmp0];     %append FP data together
            %get OD data
            ODtmp0 = cellstruct.(cellnamesnone{jj}).(ODfld);
            ODtmp = [ODtmp,ODtmp0];     %append OD data together
        end
        %average fluorescence fields against OD
        [FPout,ODout] = makeFPbasalOD(FPtmp,ODtmp);
        %function that takes OD and ouputs basal fluorescence for the channel
        basalFP = @(OD) interp1(ODout,FPout,OD,'nearest','extrap');
        basalstruct.(dprops{ii}) = basalFP;
    else
        %If OD, take average of all media fields
        mediaflds = nonestruct.(dprops{ii});
        tmp = [];
        for jj = 1:length(mediaflds)
            tmp0 = cellstruct.(mediaflds{jj}).(dprops{ii});
            tmp = [tmp,tmp0];
        end
        basalstruct.(dprops{ii}) = mean(mean(tmp,2));
    end
end


function [FPout,ODout] = makeFPbasalOD(FP,OD)
%make single OD vs FP curve for each channel by combining all measurements

%make a standard OD curve
minOD = min(OD,[],'all');
maxOD = max(OD,[],'all');
ODout = linspace(minOD,maxOD,200)';     %standardized OD curve

FPnew = zeros(length(ODout),size(FP,2));
%loop across measurements
for ii = 1:size(FP,2)
    %line up fluorescence measurements with the same ODs
    [ODtmp,~,idx] = unique(OD(:,ii),'stable');
    FPtmp = accumarray(idx,FP(:,ii),[],@mean);
    %interpolate to get standard fluorescence curve
    FPnew(:,ii) = interp1(ODtmp,FPtmp,ODout,'nearest','extrap');
end
%take the average over all measurements
FPout = mean(FPnew,2);
```
