## Supplementary material for "Predicting Composition of Genetic Circuits with Resource Competition: Demand and Sensitivity": Experimental data and Matlab code: subtractbasal6.m

```
function [outstruct,basalstruct] = subtractbasal6(cellstruct,mediastruct,subfields)
%[newstruct,basalstruct] = subtractbasal5(cellstct,nonefld,mediafld,ODeps,FPeps)
%find basal level for each property by assuming only media contributes to
%background (ignores fluorescence of cells).
%subtract off average background fluorescence and OD levels from raw data

if nargin < 2 || isempty(mediastruct)
    %struct to determine background media contribution for each datafield
    mediastruct = struct;
    mediastruct.OD = {'M9'};
    mediastruct.BFP = {'M9'};
    mediastruct.GFP = {'M9'};
    mediastruct.RFP = {'M9'};
end
if nargin < 3 || isempty(subfields)
    subfields = fieldnames(mediastruct);
end

%init
outstruct = struct;
cellnames = fieldnames(cellstruct);

%find basal levels from media for each channel
basalstruct = findbasallevel5(cellstruct,mediastruct);

%subtract basal levels off data
for ii = 1:length(cellnames)
    
    outobj = RSexpdata;
    dataobj = cellstruct.(cellnames{ii});
    %loop across each cell type
    dprops = fieldnames(dataobj);
    for jj = 1:length(dprops)
        %if matches desired fields, subtract fields
        if any(strcmpi(dprops{jj},subfields))
            outobj.(dprops{jj}) = dataobj.(dprops{jj}) - basalstruct.(dprops{jj});
        else
            %copy remaining data
            outobj.(dprops{jj}) = dataobj.(dprops{jj});
        end
    end
    %output
    outstruct.(cellnames{ii}) = outobj;
end


function basalstruct = findbasallevel5(cellstruct,nonestruct)
%find basal levels of each property based on media measurements

basalstruct = struct;
dprops = fieldnames(nonestruct);

%loop across properties
for ii = 1:length(dprops)
    %take average of all media fields
    mediaflds = nonestruct.(dprops{ii});
    tmp = [];
    for jj = 1:length(mediaflds)
        tmp0 = cellstruct.(mediaflds{jj}).(dprops{ii});
        tmp = [tmp,tmp0(:)];
    end
    basalstruct.(dprops{ii}) = mean(tmp,'all');
end


function [FPout,ODout] = makeFPbasalOD(FP,OD)
%make single OD vs FP curve for each channel by combining all measurements

%make a standard OD curve
minOD = min(OD,[],'all');
maxOD = max(OD,[],'all');
ODout = linspace(minOD,maxOD,200)';     %standardized OD curve

FPnew = zeros(length(ODout),size(FP,2));
%loop across measurements
for ii = 1:size(FP,2)
    %line up fluorescence measurements with the same ODs
    [ODtmp,~,idx] = unique(OD(:,ii),'stable');
    FPtmp = accumarray(idx,FP(:,ii),[],@mean);
    %interpolate to get standard fluorescence curve
    FPnew(:,ii) = interp1(ODtmp,FPtmp,ODout,'nearest','extrap');
end
%take the average over all measurements
FPout = mean(FPnew,2);
```
