## Supplementary material for "Predicting Composition of Genetic Circuits with Resource Competition: Demand and Sensitivity": Experimental data and Matlab code: subtractbasal.m

```
function [newstruct,basalstruct] = subtractbasal(cellstruct,ODeps,FPeps,nonecell)
%find basal level for each property by comparing with M9 media and subtract
%it off for each field for each cell

if nargin < 4
    nonecell = 'T10';
end

%init
newstruct = struct;
cellnames = fieldnames(cellstruct);
%find basal levels
basalstruct = findbasallevel(cellstruct.(nonecell),ODeps,FPeps);
dataprops = fieldnames(basalstruct);

for ii = 1:length(cellnames)
    %loop across each cell type
    for jj = 1:length(dataprops)
        %loop across each data property and subtract off basal level
        newstruct.(cellnames{ii}).(dataprops{jj}) = ...
            cellstruct.(cellnames{ii}).(dataprops{jj}) - ...
            repmat(basalstruct.(dataprops{jj}),[1,...
            size(cellstruct.(cellnames{ii}).(dataprops{jj}),2)]);
    end
end


function basalstruct = findbasallevel(nonestruct,ODeps,FPeps)
%find basal levels of each property based on media measurements

basalstruct = struct;
dataprops = fieldnames(nonestruct);

for ii = 1:length(dataprops)
    if contains(dataprops{ii},'OD')
        basalstruct.(dataprops{ii}) = ...
            mean(mean(nonestruct.(dataprops{ii})(1:100,:)),2) - ODeps;
    elseif contains(dataprops{ii},'FP')
        basalstruct.(dataprops{ii}) = ...
            mean(nonestruct.(dataprops{ii}),2) - FPeps;
    elseif contains(dataprops{ii},'time') || contains(dataprops{ii},'temp')
        basalstruct.(dataprops{ii}) = 0;
    else
        basalstruct.(dataprops{ii}) = ...
            mean(nonestruct.(dataprops{ii}),2);
    end
end
```
