## Supplementary material for "Predicting Composition of Genetic Circuits with Resource Competition: Demand and Sensitivity": Experimental data and Matlab code: subtractbasalnomedia.m

```
function [newstruct,basalstruct] = subtractbasalnomedia(cellstruct,ODeps,FPeps,n)
%[newstruct,basalstruct] = subtractbasal3(cellstct,nonefld,mediafld,ODeps,FPeps)
%find basal level for each property by assuming initial conditition should
%be 0 for all properties

if nargin < 2
    ODeps = 0;
end
if nargin < 3
    FPeps = 0;
end
if nargin < 4
    n = 5;
end

%init
newstruct = struct;
cellnames = fieldnames(cellstruct);

%find basal levels
basalstruct = findbasalnomedia(cellstruct,ODeps,FPeps,n);

%subtract basal levels off data
for ii = 1:length(cellnames)
    dataprops = fieldnames(cellstruct.(cellnames{ii}));
    %loop across each cell type
    for jj = 1:length(dataprops)
        newstruct.(cellnames{ii}).(dataprops{jj}) = ...
            cellstruct.(cellnames{ii}).(dataprops{jj}) - ...
            basalstruct.(cellnames{ii}).(dataprops{jj});
    end
end


function basalstruct = findbasalnomedia(cellstruct,ODeps,FPeps,n)
%find basal levels of each property based on average of first n samples

if nargin < 4
    n = 5;
end

basalstruct = struct;
cellnames = fieldnames(cellstruct);

for jj = 1:length(cellnames)
    dprops = fieldnames(cellstruct.(cellnames{jj}));
    for ii = 1:length(dprops)
        %find average values for each property taking the average of each
        %column or each cell
        if contains(dprops{ii},'OD')
            %if property is OD
            if iscell(cellstruct.(cellnames{jj}).(dprops{ii}))
                basalstruct.(cellnames{jj}).(dprops{ii}) = ...
                    cellfun(@(x) mean(x(1:n,:),1)-ODeps,...
                    mediastruct.(dprops{ii}),'UniformOutput',false);
            else
                basalstruct.(cellnames{jj}).(dprops{ii}) = ...
                    mean(cellstruct.(cellnames{jj}).(dprops{ii})(1:n,:),1) - ODeps;
            end
        elseif contains(dprops{ii},'FP')
            %if property is fluorescence
            if iscell(cellstruct.(cellnames{jj}).(dprops{ii}))
                basalstruct.(cellnames{jj}).(dprops{ii}) = ...
                    cellfun(@(x) mean(x(:,1:n),2)-FPeps,...
                    cellstruct.(cellnames{jj}).(dprops{ii}),'UniformOutput',false);
            else
                basalstruct.(cellnames{jj}).(dprops{ii}) = ...
                    mean(cellstruct.(cellnames{jj}).(dprops{ii})(1:n,:),1) - FPeps;
            end
        elseif contains(dprops{ii},'time') || contains(dprops{ii},'temp')
            %if property is time
            basalstruct.(cellnames{jj}).(dprops{ii})  = 0;
        else
            %if property is something else
            if iscell(cellstruct.(cellnames{jj}).(dprops{ii}))
                basalstruct.(cellnames{jj}).(dprops{ii}) = cellfun(@(x) mean(x(1:n,:),1),...
                    cellstruct.(cellnames{jj}).(dprops{ii}),'UniformOutput',false);
            else
                basalstruct.(cellnames{jj}).(dprops{ii}) = ...
                    mean(cellstruct.(cellnames{jj}).(dprops{ii})(1:n,:),1);
            end
        end
    end
end
```
