## Supplementary material for "Predicting Composition of Genetic Circuits with Resource Competition: Demand and Sensitivity": Experimental data and Matlab code: testinducplot.m

```
figure(1); clf;

%plot RS y
u = cell(length(RSmodnames),1);
RSmodnames = fieldnames(RSmodsout);
for n = 1:length(RSmodnames)
    RSmod2 = RSmodsout.(RSmodnames{n});
    RSyalone = RSmod2.y;                %scalar
    RSyalonestd = RSmod2.ystd;
    RSyperturb = RSmod2.perturby;
    RSyperturbstd = RSmod2.perturbystd;
    
    RSperturb = RSmod2.perturbname;
    for k = 1:length(RSperturb)
        if isfield(modsout,RSperturb{k})
            u{k} = modsout.(RSperturb{k}).u;
        else
            u{k} = 0;
        end
    end
    
    %plot
    subplot(2,1,n)
    plotRSycurves(u,RSyperturb,RSyperturbstd,RSyalone,RSyalonestd);
    xlabel('Input, u [nM]')
    ylabel('Resource Sensor y, AU')
    legend(['alone'],'Location','best')
    title(RSmodnames{n})
    ylim([0, Inf])
end

figure(2); clf;
figure(3); clf;
figure(4); clf;

%plot module y
normmodnames = fieldnames(modsout);
[u,yvec,ystdvec,Qvec,Qvecstd,Svec,Svecstd,Svec2,Svec2std,Svec3,Svec3std] = ...
    deal(cell(length(normmodnames),1));
for m = 1:length(normmodnames)
    mod2 = modsout.(normmodnames{m});
    u{m} = mod2.u;
    yalone = mod2.y;
    yalonestd = mod2.ystd;
    yperturb = mod2.perturby;
    yperturbstd = mod2.perturbystd;
    %store normalized output for plotting
    yvec{m} = [yalone, yperturb]./yalone;
    ystdvec{m} = [yalonestd, yperturbstd]./yalone;
    %store resource demand for plotting
    Qvec{m} = mod2.Q;
    Qvecstd{m} = mod2.Qstd;
    Svec{m} = mod2.S;
    Svecstd{m} = mod2.Sstd;
    Svec2{m} = mod2.S2;
    Svec2std{m} = mod2.S2std;
    Svec3{m} = mod2.S3;
    Svec3std{m} = mod2.S3std;
    
    figure(2);
    subplot(2,2,m);
    plotQScurves(u{m},Qvec{m},Qvecstd{m})
    title(normmodnames{m})
    ylim([0,Inf])
    xlabel('Input, u [nM]')
    ylabel('Q [ ]')
    
    figure(3);
    subplot(2,2,m);
    plotQScurves(u{m},Svec{m},Svecstd{m})
    title(normmodnames{m})
    xlabel('Input, u [nM]')
    ylabel('S [ ]')
    
    figure(4);
    subplot(2,2,m);
    plotQScurves(u{m},Svec2{m},Svec2std{m})
    title(normmodnames{m})
    xlabel('Input, u [nM]')
    ylabel('S_2 [ ]')
end


function [lh,ebh] = plotRSycurves(uvec,Yvec,Yvecstd,yalone,yalonestd)

%init
zerooffset = 1e-4;
[lh,ebh] = deal(cell(length(Yvec),1));
umax = max(cellfun(@max,uvec));
umin = min(max(cellfun(@min,uvec),zerooffset));

%plot module alone reference
semilogx([umin,umax],ones(2,1),'--','linewidth',1.2); hold on
errorbar(umin,1,yalonestd./yalone,'.k','linewidth',1.5);

%loop through each perturbation condition
for l = 1:length(Yvec)
    
    %deal with the zero in u
    u = uvec{l};
    if length(u) == 1
        u3 = [umin,umax];
    else
        u(u <= 0) = u(u <= 0) + zerooffset;
        %u3 = [u(:,1),1.2*u(:,1),u(:,2:end)];
        u3 = [u(:,1),u(:,2:end)];
    end
    
    %cast size up if y is scalar
    if length(Yvec{l}) == 1 && length(u3) > 1
        %plot horizontal line at constant level
        y = Yvec{l}*ones(size(u3));
        ystd = Yvecstd{l};
        %plot perturbed y
        lh{l} = semilogx(u3,y./yalone,'--','linewidth',1);
        ebh{l} = errorbar(umin,y(1)./yalone,ystd./yalone,'.k','linewidth',1.5);
    else
        y = Yvec{l};
        ystd = Yvecstd{l};
        %plot perturbed y
        lh{l} = semilogx(u3,y./yalone,'linewidth',2);
        ebh{l} = errorbar(u3,y./yalone,ystd./yalone,'.k','linewidth',1.5);
    end
    
    set(gca,'fontsize',14);
    %change zero tick mark
    if any(uvec{l} <= 0)
        xt = xticks;
        xticklabels([0,xt(2:end)])
    end
end
end


function plotQScurves(uvec,yvec,yvecstd)

zerooffset = 1e-4;
%plot Q or S curves vs input u
semilogx(max(uvec,zerooffset),yvec,'linewidth',2); hold on
for jj = 1:size(yvec,2)
    errorbar(max(uvec,zerooffset),yvec(:,jj),yvecstd(:,jj),'.k','linewidth',1.5);
end

%change zero tick mark
set(gca,'fontsize',14);
if any(uvec <= 0)
    xt = xticks;
    xticklabels([0,xt(2:end)])
end

end
```
