## Supplementary material for "Predicting Composition of Genetic Circuits with Resource Competition: Demand and Sensitivity": Experimental data and Matlab code: threshdata.m

```
function outstruct = threshdata(cellstruct,threshfield,thresh,datafields)
%outstruct = threshdata(cellstruct,threshfield,thresh,datafields)
%threshold all subfields given by DATAFIELDS in CELLSTRUCT based on the
%threshold in THRESH where THRESH = [THRESHMIN,THRESHMAX] and is in the
%subfield THRESHFIELD.

outstruct = struct;
cellnames = fieldnames(cellstruct);

%loop through cells
for ii = 1:length(cellnames)
    %find indicies where to place threshold
    threshdata = cellstruct.(cellnames{ii}).(threshfield);
    [~,indmin] = min(abs(threshdata - min(thresh)));
    [~,indmax] = min(abs(threshdata - max(thresh)));
    
    %loop through datafields
    for jj = 1:length(datafields)
        
        data = cellstruct.(cellnames{ii}).(datafields{jj});
        data2 = data(indmin:indmax,:);
        
        outstruct.(cellnames{ii}).(datafields{jj}) = data2;
    end
    outstruct.(cellnames{ii}).(threshfield) = threshdata(indmin:indmax,:);
end
```
