## Supplementary material for "Predicting Composition of Genetic Circuits with Resource Competition: Demand and Sensitivity": Experimental data and Matlab code: zipsubstruct.m

```
function outstruct = zipsubstruct(cellstruct,mediafield)
%outstruct = zipsubstruct(cellstruct)

outstruct = struct;
cellnames = fieldnames(cellstruct);

for ii = 1:length(cellnames)
    if strcmpi(cellnames{ii},mediafield)
        continue
    end
    datafields = fieldnames(cellstruct.(cellnames{ii}));
    for jj = 1:length(datafields)
        disp([cellnames{ii},' ',datafields{jj}])
        %append data to output struct
        z = cellstruct.(cellnames{ii}).(datafields{jj});
        if isfield(outstruct,datafields{jj})
            x = outstruct.(datafields{jj});
            outstruct.(datafields{jj}) = [x,z(:)];
        else
            outstruct.(datafields{jj}) = z(:);
        end
    end
end
```
