## Supplementary material for "Predicting Composition of Genetic Circuits with Resource Competition: Demand and Sensitivity": Experimental data and Matlab code: barsubfield.m

```
function [hbar,hebar] = barsubfield(instruct,datafield,datafieldstd,axh)
%barsubfield(cellstruct,xfield,figh)
%make a bar chart for each field in INSTRUCT

%input verification
if nargin < 4 || isempty(axh)
    axh = gca;
end
if nargin < 3 || isempty(datafieldstd) || strcmpi(datafieldstd,'std')
    datafield0 = fieldnames(instruct);
    datafieldstd = datafield0(contains(datafield0,'std'));
end
if nargin < 2 || isempty(datafield)
    datafield0 = fieldnames(instruct);
    datafield = setxor(datafield0,datafieldstd);
end
if ~(ischar(datafield) || iscell(datafield))
    error('datafield not string')
end
if ~(ischar(datafieldstd) || isnumeric(datafieldstd) || iscell(datafieldstd))
    error('datafieldstd not string or number')
end

fntsze = 12;
barcolor = [149,209,249]./255;      %light blue
% barcolor = [0,0.4470,0.7410];     %dark blue

%get fields for each cell type
names = fieldnames(instruct);

if ~iscell(datafield)
    %get unique fields
    [~,ia,~] = unique(names,'stable');
    x2 = 1:length(names);
    x2 = x2(ia);
    [data,datastd] = deal(zeros(length(names),1));
    %loop through fields
    for ii = 1:length(names)
        %put data into vector
        if isfield(instruct.(names{ii}),datafield)
            data(ii) = instruct.(names{ii}).(datafield);
        end
        %put datastd into vector
        if ischar(datafieldstd)
            if isfield(instruct.(names{ii}),datafieldstd)
                datastd(ii) = instruct.(names{ii}).(datafieldstd);
            end
        elseif isnumeric(datafieldstd)
            if length(datafieldstd) >= length(names)
                datastd(ii) = datafieldstd(ii);
            else
                datastd(ii) = datafieldstd(1);
            end
        end
    end
else
    [x2,data,datastd] = deal(zeros(length(datafield),1));
    for ii = 1:length(datafield)
        x2(ii) = ii;
        data(ii) = instruct.(datafield{ii});
        datastd(ii) = instruct.(datafieldstd{ii});
    end
end

axes(axh);
hbar = bar(x2,data,'FaceColor',barcolor); hold on
xticks(x2);
if ~iscell(datafield); xticklabels(names); ylabel(datafield)
else; xticklabels(datafield);
end
hebar = errorbar(x2,data,datastd,'.k','linewidth',2);
set(gca,'fontsize',fntsze)
```
