## Supplementary material for "Predicting Composition of Genetic Circuits with Resource Competition: Demand and Sensitivity": Experimental data and Matlab code: combinedilutions.m

```
function outstruct = combinedilutions(datastruct)
%HELP need to fix for RSexpdata objects

outstruct = struct;

if iscell(datastruct)
    %loop across each element in the cell array
    for ii = 1:length(datastruct)
        %concatenate fields of each cell in cellstruct
        outstruct = catstructs(datastruct{ii},outstruct);
    end
else
    outstruct = datastruct;
end


function outstruct = catstructs(instruct,outstruct)
%outstruct = catstructs(instruct,outstruct)
%concatenate fields within the two structures instruct and outstuct

if isa(instruct,'RSexpdata')
    if ~isempty(outstruct)
        %concatenate objects
        outstruct = catRSexp(outstruct,instruct);
    else
        %copy object
        outstruct = instruct;
    end
elseif isstruct(instruct)
    names = fieldnames(instruct);
    %loop across subfields
    for ii = 1:length(names)
        %recursion
        if isfield(outstruct,names{ii})
            outstruct.(names{ii}) = catstructs(instruct.(names{ii}),outstruct.(names{ii}));
        else
            outstruct.(names{ii}) = catstructs(instruct.(names{ii}),[]);
        end
    end
elseif ischar(instruct)
    %append character arrays into cell array
    outstruct = [outstruct;{instruct}];
elseif isnumeric(instruct)
    %concatenate numeric data along the 1st dimension
    outstruct = cat(1,outstruct,instruct);
else
    outstruct = instruct;
end


function outobj = catRSexp(obj1,obj2)

names1 = fieldnames(obj1);
names2 = fieldnames(obj2);
names = union(names1,names2);
outobj = RSexpdata;

for ii = 1:length(names)
    if strcmp(names{ii},'time')
        prevtime = obj1.(names{ii})(end) + obj1.timestep;
        outobj.(names{ii}) = cat(1,obj1.(names{ii}),obj2.(names{ii}) + prevtime);
        continue
    end
    
    if isprop(obj1,names{ii}) && isprop(obj2,names{ii})
        if ~iscell(obj1.(names{ii}))
            if all(size(obj1.(names{ii})) == size(obj2.(names{ii})),'all') && ...
                all(obj1.(names{ii}) == obj2.(names{ii}),'all')
                %copy if arrays are exactly the same
                outobj.(names{ii}) = obj1.(names{ii});
            elseif all(size(obj1.(names{ii}))  == [1,1]) || ...
                    all(size(obj2.(names{ii}))  == [1,1])
                %scalar data
                outobj.(names{ii}) = cat(2,obj1.(names{ii}), obj2.(names{ii}));
            else
                %concatenate otherwise and pad with NaNs
                %numsamples1 = size(obj1.(names{ii}),2);
                %numsamples2 = size(obj2.(names{ii}),2);
                numsamplesdiff = size(obj1.(names{ii}),2) - size(obj2.(names{ii}),2);
                numtime1 = size(obj1.(names{ii}),1);
                numtime2 = size(obj2.(names{ii}),1);
                if numsamplesdiff > 0
                    %pad with NaNs
                    NaNpad = NaN(numtime2, numsamplesdiff);
                    outobj.(names{ii}) = cat(1,obj1.(names{ii}),[obj2.(names{ii}),NaNpad]);
                elseif numsamplesdiff < 0
                    %pad with NaNs
                    NaNpad = NaN(numtime1, -numsamplesdiff);
                    outobj.(names{ii}) = cat(1,[obj1.(names{ii}),NaNpad],obj2.(names{ii}));
                else
                    outobj.(names{ii}) = cat(1,obj1.(names{ii}),obj2.(names{ii}));
                end
            end
        else
            outobj.(names{ii}) = obj1.(names{ii});
        end
    elseif isprop(obj1,names{ii}) && ~isprop(obj2,names{ii})
        outobj.(names{ii}) = obj1.(names{ii});
    elseif ~isprop(obj1,names{ii}) && isprop(obj2,names{ii})
        outobj.(names{ii}) = obj2.(names{ii});
    end
end
```
