## Supplementary material for "Predicting Composition of Genetic Circuits with Resource Competition: Demand and Sensitivity": Experimental data and Matlab code: combineinductions.m

```
function modsout = combineinductions(modulesin, metadatain)
%combine module

%stack steady state outputs into col vector
specialfields = {'FPout','containingmods','isResourceSensor','isalone'};
modsout = struct;

%get field names
datafields = fieldnames(RSmodule);
modnames = fieldnames(metadatain);
cellnames = fieldnames(modulesin);

%loop over parent modules
for ii = 1:length(modnames)
    submodnames = intersect(metadatain.(modnames{ii}).mods,cellnames,'stable');
    if isempty(submodnames)
        continue
    end
    %check if output is initalized
    if ~isfield(modsout,modnames{ii})
        modsout.(modnames{ii}) = RSmodule;
    end
    %loop over each data field
    for k = 1:length(datafields)
        if ~any(strcmp(datafields{k},specialfields))
            %loop over submodules
            for jj = 1:length(submodnames)
                %append data to columns
                if isfield(modsout,modnames{ii}) && isprop(modsout.(modnames{ii}),datafields{k}) ...
                        && ~isempty(modsout.(modnames{ii}).(datafields{k}))
                    olddata = modsout.(modnames{ii}).(datafields{k});
                else; olddata = [];
                end
                modsout.(modnames{ii}).(datafields{k}) = ...
                    [olddata; modulesin.(submodnames{jj}).(datafields{k})];
            end
        else
            %just copy data over for special fields
            modsout.(modnames{ii}).(datafields{k}) = ...
                modulesin.(submodnames{1}).(datafields{k});
        end
    end
    %write inducer concentration
    modsout.(modnames{ii}).u = metadatain.(modnames{ii}).u;
end

%copy over any experiments that don't have inducers or were missed
modnames2 = [];
for p = 1:length(modnames)
    modnames2 = [modnames2,metadatain.(modnames{p}).mods];
end
noninducedmods = setdiff(cellnames,modnames2);

for m = 1:length(noninducedmods)
    modsout.(noninducedmods{m}) = modulesin.(noninducedmods{m});
    modsout.(noninducedmods{m}).u = 0;
end
    
    
```
