## Supplementary material for "Predicting Composition of Genetic Circuits with Resource Competition: Demand and Sensitivity": Experimental data and Matlab code: combinemodstructs.m

```
function modstructout = combinemodstructs(modstruct1,modstruct2,varargin)
%modstructout = combinemodstructs(modstruct1,modstruct2,varargin)
%combine structs containing the RSmodule objects. Appends fields together.

if ~isempty(varargin)
    modstructout = combinemodstructs(modstruct1,modstruct2);
    for jj = 1:length(varargin)
        modstructout = combinemodstructs(modstructout,varargin{jj});
    end
else
    %init
    modstructout = modstruct1;
    cellnames2 = fieldnames(modstruct2);
    
    %append data from modstruct2 onto modstruct1.
    for ii = 1:length(cellnames2)
        if ~isfield(modstructout,cellnames2{ii})
            modstructout.(cellnames2{ii}) = modstruct2.(cellnames2{ii});
        else
            %append objects
            %warning(['module names repeat, appending for cell ',cellnames2{ii}])
            mod1 = modstructout.(cellnames2{ii});
            mod2 = modstruct2.(cellnames2{ii});
            modstructout.(cellnames2{ii}) = appendRSmods(mod1,mod2);
        end
    end
end


function modout = appendRSmods(mod1,mod2)
%append RSmodule objects

modout = mod1;
metafields = {'u','FPout','containingmods','isResourceSensor','isalone'};
datanames = fieldnames(mod2);

%loop through fields
for ii = 1:length(datanames)
    if any(strcmp(metafields,datanames{ii}))
        %just copy fields
        if iscell(modout.(datanames{ii}))
            if ~isempty(setdiff(modout.(datanames{ii}),mod2.(datanames{ii})))
                warning(['metafield ',datanames{ii},' does not match; appending fields'])
                modout.(datanames{ii}) = [modout.(datanames{ii}),mod2.(datanames{ii})];
            end
        else
            if all(size(modout.(datanames{ii})) ~= size(mod2.(datanames{ii}))) || ...
                ~all(modout.(datanames{ii}) == mod2.(datanames{ii}))
                warning(['metafield ',datanames{ii},' does not match; appending fields'])
                modout.(datanames{ii}) = {modout.(datanames{ii}),mod2.(datanames{ii})};
            end
        end
    else
        if iscell(modout.(datanames{ii}))
            %append fields
            modout.(datanames{ii}) = [modout.(datanames{ii}), mod2.(datanames{ii})];
        else
            %append fields
            if all(size(modout.(datanames{ii})) ~= size(mod2.(datanames{ii}))) || ...
                all(modout.(datanames{ii}) ~= mod2.(datanames{ii}),[1,2])
                modout.(datanames{ii}) = [modout.(datanames{ii}), mod2.(datanames{ii})];
            end
        end
    end
end
```
