## Supplementary material for "Predicting Composition of Genetic Circuits with Resource Competition: Demand and Sensitivity": Experimental data and Matlab code: combinenames.m

```
function namesout = combinenames(measurednames,perturbnames,selfon)
%make strings for axis labeling with underscores of the perturbed
%conditions
if nargin < 3
    selfon = true;
end

n = 1;
namesout = cell(0);
for ii = 1:length(measurednames)
    if selfon
        %append alone condition first
        namesout{n} = measurednames{ii};
        n = n+1;
    end
    perturbname2 = setdiff(perturbnames,measurednames{ii},'stable');
    for jj = 1:length(perturbname2)
        colorm = regexp(measurednames{ii},'^\D+','match');
        colorp = regexp(perturbname2{jj},'^\D+','match');
        if strcmpi(colorm{1},colorp{1})
            %exclude pairs that have the same color
            continue
        end
        %append all perturbed conditions
        namesout{n} = [measurednames{ii},'+',perturbname2{jj}];
        n = n+1;
    end
end
```
