## Supplementary material for "Predicting Composition of Genetic Circuits with Resource Competition: Demand and Sensitivity": Experimental data and Matlab code: diffcellfields.m

```
function outstruct = diffcellfields(cellstruct,difffields,timefield,n)
%outstruct = diffcellstruct(cellstruct,difffields,timefield)
%apply smooth differentation to difffields in cellstruct with respect to
%timefield. difffields may be a cell array containing the fields to be
%differentaited. timefield must be a string.

cellnames = fieldnames(cellstruct);
outstruct = cellstruct;

if nargin < 4 || isempty(n)
    n = 9;          %moving average window size
end

%loop through cells
for ii = 1:length(cellnames)
    %loop through fields to be differentiated
    for jj = 1:length(difffields)
        if ~isempty(cellstruct.(cellnames{ii}).(difffields{jj}))
            %get fields
            FP = cellstruct.(cellnames{ii}).(difffields{jj});
            time = cellstruct.(cellnames{ii}).(timefield);
            %take natural log if OD is to be differentiated
            if strcmpi(difffields{jj},'OD')
                FP = log(FP);
            end
            %take filtered numerical derivative with respect to time
            dFPdt = moveavefilt(mydiff(FP,time),n);
            %output
            outstruct.(cellnames{ii}).([difffields{jj},'diff']) = dFPdt;
        end
    end
end


function dfdh = mydiff(f,h)
%run filtered numerical differentiation on x with respect to t
%f is vector of inputs, h is step size. h can be vector of evenly spaced
%points where f is sampled.

%find step size
if length(h) > 1
    h2 = abs(mean(h(2:end) - h(1:end-1)));
else; h2 = h;
end

% 5-pt Savitzky-Golay filtered first derivative (1st/2nd order polynomial fit)
%5 pt window
f0 = [repmat(f(1,:),2,1); f; repmat(f(end,:),2,1)];
dfdh = (-2*f0(1:end-4,:) - f0(2:end-3,:) + f0(4:end-1,:) + 2*f0(5:end,:))./(10*h2);


function y = moveavefilt(x,n)
%run zero phase moving average filter with window of n points
%a = 1;
%b = 1/n*ones(1,n);
if nnz(isnan(x)) > 0 || n < 2
    y = x;
else
    %zero phase moving average filter
    y = movmean(x,n);
end
```
