## Supplementary material for "Predicting Composition of Genetic Circuits with Resource Competition: Demand and Sensitivity": Experimental data and Matlab code: filtplatedata2.m

```
function filtdata = filtplatedata2(datastruct,wn,filtopts)
%filtdata = filtplatedata(datastruct,wn)
%run all data in datastruct through forward/backwards butterworth filter.
%If field has 'time' in it, the field is copied directly. wn is the filter
%cutoff frequency and is a value between 0 and 1 with 1 corresponding to
%the Nyquist frequency

if nargin < 3
    filtopts = 'low';
end
if nargin < 2
    wn = 0.04;
end
%input error checking
if wn >= 1 || wn <= 0
    warning('data not filtered')
    filtdata = datastruct;
    return
end

%init filter
filtdata = struct;
[b,a] = butter(5,wn,filtopts);
dataprops = fieldnames(datastruct);
%filter data
for jj = 1:length(dataprops)
    %recursion for structs within structs
    if isstruct(datastruct.(dataprops{jj}))
        filtdata.(dataprops{jj}) = filtplatedata2(datastruct.(dataprops{jj}),wn,filtopts);
    else
        %don't filter time fields
        if ~contains(dataprops{jj},'time')
            %filter along the 3rd dimension only
            datatemp = datastruct.(dataprops{jj});
            for k = 1:size(datatemp,1)
                for ii = 1:size(datatemp,2)
                    if all(isfinite(datatemp(k,ii,:)))
                        %filter data
                        filtdata.(dataprops{jj})(k,ii,:) = ...
                            filtfilt(b,a,squeeze(datatemp(k,ii,:)));
                    else
                        %skip filtering if data has Inf's or NaN's
                        filtdata.(dataprops{jj})(k,ii,:) = datatemp(k,ii,:);
                    end
                end
            end
        else
            %pass time data unfiltered
            filtdata.(dataprops{jj}) = datastruct.(dataprops{jj});
        end
    end
end
```
