## Supplementary material for "Predicting Composition of Genetic Circuits with Resource Competition: Demand and Sensitivity": Experimental data and Matlab code: findpropRS.m

```
function namesout = findpropRS(modules,propname,propvalue)
%namesout = findprop(modules,propname,propvalue)
%find cell conditions where the data in propname matches propvalue
if nargin < 3
    propvalue = true;
end
cellnames = fieldnames(modules);
indvec = zeros(length(cellnames),1);

%loop across data finding properties that match the desired value
for ii = 1:length(cellnames)
    if isprop(modules.(cellnames{ii}),propname) || ...
            isfield(modules.(cellnames{ii}),propname)
        if iscell(modules.(cellnames{ii}).(propname))
            if any(contains(modules.(cellnames{ii}).(propname),propvalue))
                indvec(ii) = true;
            else
                indvec(ii) = false;
            end
        else
            if modules.(cellnames{ii}).(propname) == propvalue
                indvec(ii) = true;
            else
                indvec(ii) = false;
            end
        end
    else
        indvec(ii) = false;
    end
end
namesout = cellnames(logical(indvec));
```
