## Supplementary material for "Predicting Composition of Genetic Circuits with Resource Competition: Demand and Sensitivity": Experimental data and Matlab code: findss4.m

```
function outstruct = findss4(instruct,steadystatestruct,datafields,avefun,ploton,figh)
%outstruct = findss3(instruct,timestruct,datafields,avefun,ploton,figh)
%average values over a user specificed interval

if nargin < 5 || isempty(ploton)
    ploton = false;
end
if nargin < 4 || isempty(avefun)
    %function to apply to get steady state
    avefun = @mean;       %or @median
end

cellnames = intersect(fieldnames(steadystatestruct),fieldnames(instruct),'stable');

%setup plot
if ploton
    if nargin < 5 || isempty(figh)
        figh = figure;
    end
    n = length(datafields);
    m = ceil(9/16*sqrt(n));
end

%init for plotting bar charts
data = cell(length(datafields),1);
[outmat,outstdmat] = deal(zeros(length(datafields),length(cellnames)));

%loop through cell conditions
for jj = 1:length(cellnames)
    
    %init steady state data output holder
    moduledata = RSmodule;
    %loop through fields
    for ii = 1:length(datafields)
        %check that fields in timestruct match instruct celltypes
        if isfield(instruct,cellnames{jj})
            %get indicies for range in steady state window
            time = instruct.(cellnames{jj}).time;
            tmin = min(steadystatestruct.(cellnames{jj}));
            tmax = max(steadystatestruct.(cellnames{jj}));
            [~,indmin] = min(abs(time - tmin));
            [~,indmax] = min(abs(time - tmax));
            
            %get data from time range
            tmp = instruct.(cellnames{jj}).(datafields{ii})(indmin:indmax,:);
            %apply average/median function to each sample
            ss = mean(tmp);
            %avevar = var(tmp,0,1);
            %samplestd = std(ss)./sqrt(numel(ss))
            %z = sqrt(sum(avevar))/numel(ss)
            
            %store steady state data in module
            moduledata.(datafields{ii}) = avefun(ss,'omitnan');
            %sqrt(sum(avevar))/numel(ss);
            moduledata.([datafields{ii},'std']) = sqrt(var(ss,'omitnan')./...
                nnz(~isnan(ss)));
            %moduledata.(datafields{ii}) = ss;
            %moduledata.([datafields{ii},'std']) = 0;%sqrt(var(ss)./numel(ss));
            
            
            %store data for histogram plots
            data{ii}{jj} = tmp(:);
            %output for bar chart
            outmat(ii,jj) = avefun(ss);
            outstdmat(ii,jj) = std(ss)./numel(ss);
        end
    end
    %output
    outstruct.(cellnames{jj}) = moduledata;
    %copy over growth rate
    outstruct.(cellnames{jj}).growthrate = instruct.(cellnames{jj}).growthrate;
    outstruct.(cellnames{jj}).growthratestd = instruct.(cellnames{jj}).growthratestd;
end

%plot
if ploton
    for k = 1:length(datafields)
        %plot bar chart of steady state data
        if length(figh) == length(datafields)
            figure(figh(k)); clf;
            set(gca,'fontsize',16)
        else
            figure(figh);
            subplot(m,ceil(n/m),k); cla;
        end
        x = 1:size(outmat,2);
        bh = bar(x,outmat(k,:)); hold on
        set(bh,'barwidth',0.6);
        if contains(datafields{k},'BFP')
            set(bh,'FaceColor',[0.48,0.79,1])
        elseif contains(datafields{k},'GFP')
            set(bh,'FaceColor',[0.7,1,0.49])
        elseif contains(datafields{k},'RFP')
            set(bh,'FaceColor',[1,0.51,0.51])
        end
        xticks(x);
        xticklabels(cellnames);
        errorbar(outmat(k,:),outstdmat(k,:),'.k','linewidth',2);
        ylabel([datafields{k},' steady state'])
        
        if 0
            %plot histograms of distributions
            figure(figh);
            ax = subplot(m,ceil(n/m),k);
            multiplehist(data{k},ax,30,'probability')
            xlabel(datafields{k})
            legend(cellnames,'Location','Best','fontsize',10);
            ylabel('prob mass')
        end
    end
end
```
