## Supplementary material for "Predicting Composition of Genetic Circuits with Resource Competition: Demand and Sensitivity": Experimental data and Matlab code: fitgrowthrate3.m

```
function cellstructout = fitgrowthrate3(cellstruct,ploton,verbose,figh)
%cellstructout = fitgrowthrate3(cellstruct,ploton,verbose,figh)
%fit OD with logisitic curve to find growthrate. cellstruct must contain 
%fields for each growth condition with an RSexpdata object in each.
%ploton is a bool whether to plot the OD
%fits and growthrates.

if nargin < 3
    verbose = false;
end
if nargin < 2
    ploton = true;
end
if (nargin < 4 || isempty(figh)) && (ploton)
    figh = figure;
end
tmax = 10;                          %[hrs] max time to fit
tmin = 2;                           %[hrs] chop off less than this
fntsze = 12;
%barplot colors
barcolor = [149,209,249]./255;      %light blue
% barcolor = [0,0.4470,0.7410];     %dark blue
strexp = '(\<[0-9]*\.)\w*(\s)?';    %regexp for formatting parameter strings in figure titles
%match string beginning with numeric characters [0-9] then string has a '.'
%then match rest of the word until an optional 1 trailing whitespace character

%init
cellstructout = cellstruct;
celltypes = fieldnames(cellstruct);
[Kcell,Kstdcell,rcell,rstdcell,P0cell,P0stdcell] = deal(zeros(length(celltypes),1));

disp('fitting growth rate...')

%loop across cell conditions
for ii = 1:length(celltypes)
    
    %init
    data = cellstruct.(celltypes{ii});
    ODfld = data.ODfield{1};
    n = size(data.(ODfld),2);
    time = data.time;
    
    if ~(max(time,[],'all') < 6 || strcmpi(celltypes{ii},'M9'))
        %skip fitting otherwise
        
        [ODfit,dODdt] = deal(zeros(size(data.(ODfld))));
        [rsq,K,P0,r,sigmaK,sigmaP0,sigmar] = deal(zeros(n,1));
        %only take time up to tmax hrs
        inds = time <= tmax & time >= tmin;
        %set up logistic fitting problem
        opts = fitoptions('method','NonlinearLeastSquares','Robust','on',...
            'StartPoint',[0.9,0.02,0.6],'lower',[0,0,0]);
        ft = fittype('K*a*exp(r*x)/(K+a*(exp(r*x)-1))',...
            'coeff',{'K','a','r'},'options',opts);
        %loop through each cell type
        for jj = 1:n
            %run fitting
            if iscell(data.(ODfld))
                [fitobj,gof] = fit(time(inds),data.(ODfld){jj}(inds,:),ft);
            elseif isnumeric(data.(ODfld))
                [fitobj,gof] = fit(time(inds),data.(ODfld)(inds,jj),ft);
            end
            %output parameters from fit to OD, and dOD/dt as logistic curves
            K(jj) = fitobj.K;
            P0(jj) = fitobj.a;
            r(jj) = fitobj.r;
            paracint = confint(fitobj);     %parameter confidence intervals
            sigmaK(jj) = max(abs(paracint(:,1) - fitobj.K));
            sigmaP0(jj) = max(abs(paracint(:,2) - fitobj.a));
            sigmar(jj) = max(abs(paracint(:,3) - fitobj.r));
            ODfit(:,jj) = K(jj)*P0(jj)*exp(r(jj).*time)./(K(jj) + P0(jj)* ...
                (exp(r(jj).*time) - 1));
            dODdt(:,jj) = K(jj)*P0(jj)*r(jj)*(K(jj) - P0(jj))*exp(r(jj).*time)./...
                ((K(jj) + P0(jj)*(exp(r(jj).*time) - 1)).^2);
            rsq(jj) = gof.rsquare;
        end
        %output parameters
        cellstructout.(celltypes{ii}).growthrate = {mean(r)};
        cellstructout.(celltypes{ii}).growthratestd = {sqrt(sum(sigmar.^2)/length(sigmar))};
        
        %for plotting average parameters for each cell type
        Kcell(ii) = mean(K);
        Kstdcell(ii) = sqrt(sum(sigmaK.^2)/length(sigmaK)); %std(K);
        rcell(ii) = mean(r);
        rstdcell(ii) = sqrt(sum(sigmar.^2)/length(sigmar)); %std(r);
        P0cell(ii) = mean(P0);
        P0stdcell(ii) = sqrt(sum(sigmaP0.^2)/length(sigmaP0)); %std(P0);
        
        %plotting
        if ploton && verbose% || (any(rsq < 0.85) && ~strcmpi(celltypes{ii},'M9'))
            %format parameter strings for titles
            rsqstr = regexp(num2str(rsq(:)',3),strexp,'match');
            Kstr = regexp(num2str(K(:)',3),strexp,'match');
            P0str = regexp(num2str(P0(:)',3),strexp,'match');
            rstr = regexp(num2str(r(:)',3),strexp,'match');
            %plot
            figure;
            subplot(311);
            semilogy(time,data.(ODfld),'linewidth',2); hold on;
            set(gca,'colororderindex',1)
            semilogy(time,ODfit,'--','linewidth',2)
            title([celltypes{ii},': logistic fit R^2 = ',[rsqstr{:}]])
            ylabel('OD')
            xlim([min(time),max(time)])
            set(gca,'fontsize',fntsze)
            subplot(312);
            plot(time,dODdt,'linewidth',2)
            ylabel('d OD/dt')
            xlim([min(time),max(time)])
            title(['K = [',[Kstr{:}],']; P0 = [',[P0str{:}],']'])
            set(gca,'fontsize',fntsze)
            subplot(313);
            plot(time,dODdt./ODfit,'linewidth',2)
            xlabel('time (hrs)')
            ylabel('growthrate')
            title(['r = [',[rstr{:}],']'])
            xlim([min(time),max(time)])
            set(gca,'fontsize',fntsze)
        end
    else
        
    end
end

if ploton && (nnz(rcell) > 1)
    %plot bargraph of parameters for each cell type
    names = strrep(celltypes,'_','-');
    %plot bargraph of K, r, P0
    figure(figh); clf;
    %plot P0
    subplot(311);
    bar(P0cell,'FaceColor',barcolor); hold on
    set(gca,'XTick',1:length(P0cell),'XTickLabel',[],'fontsize',fntsze,...
        'XLim',[0.5,length(P0cell)+0.5]);
    errorbar(P0cell,P0stdcell,'.k','linewidth',2)
    ylabel('P0')
    ylim([0,1.1*max(P0cell)])
    %plot K
    subplot(312);
    bar(Kcell,'FaceColor',barcolor); hold on
    set(gca,'XTick',1:length(Kcell),'XTickLabel',[],'fontsize',fntsze,...
        'XLim',[0.5,length(Kcell)+0.5]);
    errorbar(Kcell,Kstdcell,'.k','linewidth',2)
    ylabel('K')
    ylim([0,1.1*max(Kcell)])
    %plot r
    subplot(3,1,3);
    bar(rcell,'FaceColor',barcolor); hold on
    set(gca,'XTick',1:length(rcell),'XTickLabel',names,'XTickLabelRotation',90,...
        'XLim',[0.5,length(rcell)+0.5],'fontsize',fntsze);
    errorbar(rcell,rstdcell,'.k','linewidth',2)
    ylabel('r')
    ylim([0,1.1*max(rcell)])
end
```
