## Supplementary material for "Predicting Composition of Genetic Circuits with Resource Competition: Demand and Sensitivity": Experimental data and Matlab code: fluorescencebleed2.m

```
function C = fluorescencebleed2(celldata,metastruct,maxstruct,ploton,fighs)
%compensate for fluorescence bleed between channels
%this should be done after background subtraction, also include OD as a
%fluorescent contanimant

if nargin < 2 || isempty(metastruct)
    metastruct = struct;
    metastruct.BFP = {'BY172_0','BY173_0','B170','B171'};  %only BFP
    metastruct.GFP = {'GY182_0','GY183_0','G181','G180'};
    metastruct.RFP = {'RY192_0','RY193_0','R191','R190'};
    metastruct.YFP = {'Y240_10','RY194_10','GY184_10','BY174_10'};
end
if nargin < 3 || isempty(ploton)
    ploton = false;
end
if nargin < 4 || isempty(maxstruct)
    %upper data bound for finding color correction matrix
    maxstruct.BFP = 2e4;
    maxstruct.GFP = 2.4e4;
    maxstruct.RFP = 2e4;
    maxstruct.YFP = 1e4;
    maxstruct.OD  = 0.2;
end
if nargin < 5 || isempty(fighs)
    fighs = 6:9;
end

metastruct = orderfields(metastruct);
FPfields = fieldnames(metastruct);
cellnames = fieldnames(celldata);
n = length(FPfields); %number of channels
C = zeros(n);
C2 = zeros(n);

%find correlation between channels using cells expressing only BFP/GFP/RFP
%as a control
for ii = 1:length(FPfields)
    
    %get cells and channels
    celltypes = intersect(metastruct.(FPfields{ii}),cellnames,'stable');
    
    if ploton; figure(fighs(ii)); clf; end
    
    for jj = 1:n
        xdata = [];
        ydata = [];
        for k = 1:length(celltypes)
            if ~isempty(ydata)
                ysamples = min([size(ydata,1), size(celldata.(celltypes{k}).(FPfields{ii}),1)]);
                xsamples = min([size(xdata,1), size(celldata.(celltypes{k}).(FPfields{jj}),1)]);
                %compute the correlation between fluorescence channels
                ydata = [ydata(1:ysamples,:), celldata.(celltypes{k}).(FPfields{ii})(1:ysamples,:)];
                xdata = [xdata(1:xsamples,:), celldata.(celltypes{k}).(FPfields{jj})(1:xsamples,:)];
            else
                ydata = celldata.(celltypes{k}).(FPfields{ii});
                xdata = celldata.(celltypes{k}).(FPfields{jj});
            end
        end
        
        %get timepoints where all cell repeats are below thresholds
        %inds = all(ydata<maxstruct.(FPfields{ii}) & xdata<maxstruct.(FPfields{jj}),2);
        inds2 = ydata<maxstruct.(FPfields{ii}) & xdata<maxstruct.(FPfields{jj});
        
        %trim data
        %ydata2 = ydata(inds,:);
        %xdata2 = xdata(inds,:);
        ydata2 = ydata(inds2);
        xdata2 = xdata(inds2);
        
        %averaging in this way may cause some problems if cell repeats
        %arent lined up correctly against time
        %corr = mean(xdata2,2)'/mean(ydata2,2)';
        
        %b = robustfit(mean(xdata2,2)',mean(ydata2,2)');
        %b2 = robustfit(mean(ydata2,2)',mean(xdata2,2)');
        
        %fo = fit(mean(xdata2,2)',mean(ydata,2)','poly1');
        %fo2 = fit(mean(ydata2,2)',mean(xdata,2)','poly1');
        
        if ~isempty(xdata2)
            p = polyfit(mean(xdata2,2)',mean(ydata2,2)',1);
            p2 = polyfit(mean(ydata2,2)',mean(xdata2,2)',1);
        else
            [p,p2] = deal(zeros(2,1));
        end
        
        corr2 = 1./p(1);
        corr = p2(1);
        C(jj,ii) = corr;
        C2(jj,ii) = corr2;
        
        if ploton && ~isempty(xdata2)
            xplt = linspace(min(xdata2,[],'all'),max(xdata2,[],'all'),200)';
            %plot
            subplot(2,2,jj);
            %plot(xdata,ydata,xdata,1/corr*xdata,'--',xdata2,ydata2,'linewidth',1.5)
            plot(xdata,ydata,'.',xplt,1/corr*xplt + p(2),'k--','linewidth',1.5)
            ylim([min(ydata,[],'all'),max(ydata,[],'all')])
            if jj == 1; title(strjoin(strrep(celltypes,'_','-'),', '));
            end
            ylabel(['Measured ',FPfields{ii}])
            xlabel(['Measured ',FPfields{jj}])
            set(gca,'fontsize',12)
        end
    end
    %[C,C2]
end
```
