## Supplementary material for "Predicting Composition of Genetic Circuits with Resource Competition: Demand and Sensitivity": Experimental data and Matlab code: loadxlsdata.m

```
function dataout = loadxlsdata(filename, sheet, platesize)
%read in data from xls file from Tecan plate reader and output as RSexpdata
%object using a 96 well plate

if nargin < 3 || isempty(platesize)
    platesize = [8,12];
end

%init
dataout = RSexpdata;

%read in data
disp('reading in xlsx data...')
[numdata,txtdata,~] = xlsread(filename,sheet);

%find data headers
dataheaders = txtdata(find(contains(txtdata(:,1),'Cycle Nr.')) - 1,1);
if any(~strcmp(dataheaders,'OD') & contains(dataheaders,'OD'))
    dataheaders(~strcmp(dataheaders,'OD') & contains(dataheaders,'OD')) = {'OD'};
end

%find number of rows and columns
txtstartinds = find(contains(txtdata(:,1),'Cycle Nr.'));
wellnames = txtdata(txtstartinds(1)+3:txtstartinds(2)-3,1);

%find first row of data collection:
firstrowind = find(~any(isnan(numdata),2),1,'first');
numdatared = numdata(firstrowind:end,:);
numrows = size(numdatared,1);
numsamples = size(numdatared,2);

%find size and location of data blocks
gapinds = find(all(isnan(numdatared),2));
datarows = find(~all(isnan(numdatared),2));
datablocksize = [gapinds; numrows+1] - [0; gapinds] - 1;
datablocksize = [0; datablocksize(datablocksize > 0)];

%get indicies of start of each block
prevind = 1;
blockstartinds = ones(length(datablocksize)-1,1);
for ii = 2:length(datablocksize)-1
    blockstartinds(ii) = datarows(find(datarows > datablocksize(ii)+prevind,1));
    prevind = blockstartinds(ii);
end

%get each data block and format
[time,temp] = deal(zeros(length(blockstartinds),numsamples));
plate = nan(platesize(1),platesize(2),numsamples);
for jj = 1:length(blockstartinds)
    datablock = numdatared(blockstartinds(jj):blockstartinds(jj)+datablocksize(jj+1)-1,:);
    time(jj,:) = datablock(2,:);
    temp(jj,:) = datablock(3,:);
    datablockred = datablock(4:end,:);
    
    %put into plate array with correct well locations
    for k = 1:size(datablockred,1)
        loc = wellnames2loc(wellnames{k});
        plate(loc(1),loc(2),:) = datablockred(k,:);
    end
    dataout.(dataheaders{jj}) = plate;
end
dataout.time = time(1,:)/3600;
dataout.timestep = mean(diff(time(1,:)/3600));


function loc = wellnames2loc(namestr)
%convert string position info to numerical locations

loc = zeros(2,1);
%row location
loc(1) = double(upper(namestr(1))) - 64;
%column location
loc(2) = str2double(namestr(2:end));
```
