## Supplementary material for "Predicting Composition of Genetic Circuits with Resource Competition: Demand and Sensitivity": Experimental data and Matlab code: normalizeFP3.m

```
function outstruct = normalizeFP3(instruct,normfields)
%outstruct = normalizeFP3(instruct,normfields)
%normalize fluorescence data by OD in the fields contained in the cell
%array normfields.

if nargin < 2
    %fields to divide by OD
    normfields = {'BFP','GFP','RFP','BFPdiff','GFPdiff','RFPdiff'};
end

%normalize fluorescence data based on growth rate or by number of cells (OD)
cellnames = fieldnames(instruct);

outstruct = instruct;
%loop through cell names
for k = 1:length(cellnames)
    
    data = instruct.(cellnames{k});
    ODfield = data.ODfield{1};
    ODfilt = movmean(data.(ODfield),7);
    
    %props = fieldnames(instruct.(cellnames{k}));
    for ii = 1:length(normfields)
        %loop through properties of each cell
        if isprop(data,normfields{ii}) && ~isempty(data.(normfields{ii}))
            %normalize per cell based on OD
            outstruct.(cellnames{k}).([normfields{ii},'OD']) = ...
                data.(normfields{ii})./ODfilt;
        end
    end
end
```
