## Supplementary material for "Predicting Composition of Genetic Circuits with Resource Competition: Demand and Sensitivity": Experimental data and Matlab code: ODFPcorrection.m

```
function [outstruct,C] = ODFPcorrection(instruct,FPstruct,maxstruct,C,ploton,figh)
%fix correlation between fluorescence and OD (on BFP)

if nargin < 2
    %fluorescence fields to correct for against OD and the corresponding
    %calibration experiments to use
    FPstruct = struct;
    %something without BFP
    FPstruct.BFP = {'RY192_0','RY192_01','RY192_1','RY192_10',...
                    'RY193_0','RY193_01','RY193_1','RY193_10'};
    %FPstruct.GFP = {'BY172_0','BY172_0','Y240_0'};
    %FPstruct.RFP = {'BY172_0','BY173_0','Y240_0'};
    %FPstruct.YFP = {'BY173_0','BY172_0','Y240_0','RY192_0','RY193_0'};
end
if nargin < 3
    %max values where the linear range is
    %maxstruct.BFP = 1e5;
    maxstruct.OD = 0.2;
end
if nargin < 5
    ploton = false;
end
if ploton && (isempty(figh) || nargin < 6)
    figh = figure;
end

cellnames = fieldnames(instruct);
FPfields = fieldnames(FPstruct);
ODfield = 'OD';
outstruct = instruct;

%find OD and FP relationship if not given an input
if nargin < 4 || isempty(C)
    %get correlation between OD and FP
    [p,p2,C] = deal(cell(length(FPfields),1));
    for ii = 1:length(FPfields)
        
        %FPfields{ii};
        %cells used to calibrate
        calibcells = intersect(cellnames,FPstruct.(FPfields{ii}));
        
        %append OD and FP fields from all calibration experiments together
        OD = [];
        FP = [];
        for jj = 1:length(calibcells)
            OD = [OD(:); instruct.(calibcells{jj}).(ODfield)(:); NaN];
            FP = [FP(:); instruct.(calibcells{jj}).(FPfields{ii})(:); NaN];
        end
        
        %take only the datapoints in the linear region
        ODinds = OD < maxstruct.(ODfield);
        if isfield(maxstruct,FPfields{ii})
            FPinds = FP < maxstruct.(FPfields{ii});
        else; FPinds = true(size(FP));
        end
        inds = ODinds & FPinds;
        
        %get correlation between FP and OD by linear fit
        p{ii} = polyfit(OD(inds),FP(inds),1);
        p2{ii} = 1./(polyfit(FP(inds),OD(inds),1));
        slopevec = [p{ii}(1),p2{ii}(1)];
        [~,pind] = min(abs(slopevec));
        C{ii} = slopevec(pind);
        
        if ploton
            %plot linear fit and data
            xfit = linspace(min(OD(inds)),max(OD(inds)),200)';
            yfit = xfit*C{ii};
            
            figure(figh);
            subplot(length(FPfields),1,ii);
            plot(OD(inds),FP(inds),'.',xfit,yfit,'k--','linewidth',2)
            xlabel('OD')
            ylabel(FPfields{ii})
            set(gca,'fontsize',14)
            titlenames = strcat(calibcells,',   ');
            title([titlenames{:}])
        end
    end
end

%apply correction to all experiments
for k = 1:length(cellnames)
    for m = 1:length(FPfields)
        OD0 = instruct.(cellnames{k}).(ODfield);
        FP0 = instruct.(cellnames{k}).(FPfields{m});
        FPnew = FP0 - C{m}*OD0;
        outstruct.(cellnames{k}).(FPfields{m}) = FPnew;
    end
end
```
