## Supplementary material for "Predicting Composition of Genetic Circuits with Resource Competition: Demand and Sensitivity": Experimental data and Matlab code: plotGR.m

```
function plotGR(modstruct)

%init
modnames = fieldnames(modstruct);
[grmat,grmatstd,grmat2,grmatstd2] = deal([]);
[namesout,namesout2] = deal({});

%loop through modules
for ii = 1:length(modnames)
    mod1 = modstruct.(modnames{ii});
    
    %copy growth rate info
    if iscell(mod1.growthrate)
        gr0 = cell2mat(mod1.growthrate);
        gr0std = cell2mat(mod1.growthratestd);
    else
        gr0 = mod1.growthrate;
        gr0std = mod1.growthratestd;
    end
    gr1 = mean(gr0,2);
    gr1std = sqrt(var(gr0,0,2) + mean(gr0std,2).^2);
    
    if length(gr1) == 1
        grmat = [grmat; gr1];
        grmatstd = [grmatstd; gr1std];
        namesout = [namesout,modnames(ii)];
    else
        grmat2 = [grmat2; gr1];
        grmatstd2 = [grmatstd2; gr1std];
        namesout2 = [namesout2,repmat(modnames(ii),length(gr1),1)];
    end
end

blue1 = [91,155,213]/255;       %dark blue
blue2 = [125,195,251]/255;      %medium blue
blue3 = [173,222,255]/255;      %light blue
green1 = [120,171,48]/255;      %dark green
green2 = [45,190,6]/255;        %emerald green
green3 = [163,252,67]/255;      %light green
red1 = [207,81,81]/255;         %dark red
red2 = [240,122,122]/255;       %light red
red3 = [255,168,168]/255;       %lighter red
yellow1 = [237,177,32]/255;
yellow2 = [255,255,0]/255;
yellow3 = [196,200,21]/255;

%B171 plots
indsB171 = find(contains(namesout,{'B171','BG201','BG202','BR211','BR212'}));
indsB170 = find(contains(namesout,{'B170','BG200','BG203','BR210','BR213'}));
indsG180 = find(contains(namesout,{'G180','BG200','BG201','GR220','GR222'}));
indsG181 = find(contains(namesout,{'G181','BG202','BG203','GR221','GR223'}));
indsR191 = find(contains(namesout,{'R191','BR212','BR213','GR222','GR223'}));
indsR190 = find(contains(namesout,{'R190','BR210','BR211','GR220','GR221'}));
indsY240 = find(contains(namesout2,{'Y240'}));
indsBY172 = find(contains(namesout2,{'BY173'}));
indsBY173 = find(contains(namesout2,{'BY172'}));
indsGY182 = find(contains(namesout2,{'GY182'}));
indsGY183 = find(contains(namesout2,{'GY183'}));
indsRY192 = find(contains(namesout2,{'RY192'}));
indsRY193 = find(contains(namesout2,{'RY193'}));

inds1 = 1:18;

%constitutive modules
figure(1);
bh1 = plotbars(grmat(inds1)',grmatstd(inds1)');
set(bh1,'FaceColor',blue3)
xticks(1:length(inds1))
xticklabels(namesout(inds1));
ylabel('growth rate [1/hr]')
set(gca,'fontsize',14)
ylim([0,0.6])

%inducible module with inputs 0, 0.1, 1, and 10 nM per construct
figure(2); clf;
bh7 = plotbars(grmat2',grmatstd2');
set(bh7,'FaceColor',red1)
xticks(1:length(grmat2))
xticklabels(namesout2);
ylabel('growth rate [1/hr]')
set(gca,'fontsize',14)
ylim([0,0.6])
return
% 
% %module 1
% figure(2);
% bh2 = plotbars(grmat(indsB170)',grmatstd(indsB170)');
% set(bh2,'FaceColor',blue1)
% xticks(1:length(indsB170))
% xticklabels(namesout(indsB170));
% ylabel('growth rate [1/hr]')
% set(gca,'fontsize',14)
% ylim([0,0.6])
% 
% %module 2
% figure(3); clf;
% bh3 = plotbars(grmat(indsG180)',grmatstd(indsG180)');
% set(bh3,'FaceColor',green3)
% xticks(1:length(indsG180))
% xticklabels(namesout(indsG180));
% ylabel('growth rate [1/hr]')
% set(gca,'fontsize',14)
% ylim([0,0.6])
% 
% %module 3
% figure(4); clf;
% bh4 = plotbars(grmat(indsG181)',grmatstd(indsG181)');
% set(bh4,'FaceColor',green2)
% xticks(1:length(indsG181))
% xticklabels(namesout(indsG181));
% ylabel('growth rate [1/hr]')
% set(gca,'fontsize',14)
% ylim([0,0.6])
% 
% %module 4
% figure(5); clf;
% bh5 = plotbars(grmat(indsR190)',grmatstd(indsR190)');
% set(bh5,'FaceColor',red3)
% xticks(1:length(indsR190))
% xticklabels(namesout(indsR190));
% ylabel('growth rate [1/hr]')
% set(gca,'fontsize',14)
% ylim([0,0.6])
% 
% %resource sensor 2
% figure(6); clf;
% bh6 = plotbars(grmat(indsR191)',grmatstd(indsR191)');
% set(bh6,'FaceColor',red1)
% xticks(1:length(indsR191))
% xticklabels(namesout(indsR191));
% ylabel('growth rate [1/hr]')
% set(gca,'fontsize',14)
% ylim([0,0.6])

figure(7); clf;
bh7 = plotbars(grmat2(indsY240)',grmatstd2(indsY240)');
set(bh7,'FaceColor',red1)
xticks(1:length(indsY240))
xticklabels(namesout2(indsY240));
ylabel('growth rate [1/hr]')
set(gca,'fontsize',14)
ylim([0,0.6])

figure(8); clf;
bh8 = plotbars(grmat2(indsBY172)',grmatstd2(indsBY172)');
set(bh8,'FaceColor',red1)
xticks(1:length(indsBY172))
xticklabels(namesout2(indsBY172));
ylabel('growth rate [1/hr]')
set(gca,'fontsize',14)
ylim([0,0.6])

figure(9); clf;
bh9 = plotbars(grmat2(indsBY173)',grmatstd2(indsBY173)');
set(bh9,'FaceColor',red1)
xticks(1:length(indsBY173))
xticklabels(namesout2(indsBY173));
ylabel('growth rate [1/hr]')
set(gca,'fontsize',14)
ylim([0,0.6])

figure(10); clf;
bh10 = plotbars(grmat2(indsGY182)',grmatstd2(indsGY182)');
set(bh10,'FaceColor',red1)
xticks(1:length(indsGY182))
xticklabels(namesout2(indsGY182));
ylabel('growth rate [1/hr]')
set(gca,'fontsize',14)
ylim([0,0.6])

figure(11); clf;
bh11 = plotbars(grmat2(indsGY183)',grmatstd2(indsGY183)');
set(bh11,'FaceColor',red1)
xticks(1:length(indsGY183))
xticklabels(namesout2(indsGY183));
ylabel('growth rate [1/hr]')
set(gca,'fontsize',14)
ylim([0,0.6])

figure(12); clf;
bh12 = plotbars(grmat2(indsRY192)',grmatstd2(indsRY192)');
set(bh12,'FaceColor',red1)
xticks(1:length(indsRY192))
xticklabels(namesout2(indsRY192));
ylabel('growth rate [1/hr]')
set(gca,'fontsize',14)
ylim([0,0.6])

figure(13); clf;
bh13 = plotbars(grmat2(indsRY193)',grmatstd2(indsRY193)');
set(bh13,'FaceColor',red1)
xticks(1:length(indsRY193))
xticklabels(namesout2(indsRY193));
ylabel('growth rate [1/hr]')
set(gca,'fontsize',14)
ylim([0,0.6])


function bh = plotbars(data,datastd)
nbars = size(data,1);
ngroups = size(data,2);
groupwidth = min(0.8, nbars/(nbars + 1.5));
% Set the position of each error bar in the centre of the main bar
% Based on barweb.m by Bolu Ajiboye from MATLAB File Exchange
x = zeros(nbars,ngroups);
for ii = 1:nbars
    % Calculate center of each bar
    x(ii,:) = (1:ngroups) - groupwidth/2 + (2*ii-1) * groupwidth / (2*nbars);
end
bh = bar(data'); hold on;
set(bh,'edgecolor','none');
errorbar(x,data,datastd,'.k','linestyle','none','Linewidth',1.5);
```
