## Supplementary material for "Predicting Composition of Genetic Circuits with Resource Competition: Demand and Sensitivity": Experimental data and Matlab code: plotGRvsQ.m

```
function [outstruct,grfitobj,gof] = plotGRvsQ(modstruct,ploton,figh)
%outstruct = plotGRvsQ(modstruct,expdatastruct,ploton,figh)
%get growth rate and total resource demand for all modules in modstruct

if nargin < 2
    ploton = true;
end
if nargin < 3 || isempty(figh)
    figh = figure;
end

%init
modnames = fieldnames(modstruct);
outstruct = struct;

%loop through modules
for ii = 1:length(modnames)
    %modout = RSmodule;
    mod1 = modstruct.(modnames{ii});
    modout = mod1;
    
    %get total resource demand as sum of individual resource demands
    if ~mod1.isalone
        modsinside = mod1.containingmods;
        Qtot = 0;
        Qtotvar = 0;
        for jj = 1:length(modsinside)
            %get Q for submodules
            Qtot = Qtot + mean(modstruct.(modsinside{jj}).Q,2);
            Qtotvar = Qtotvar^2 + sum(modstruct.(modsinside{jj}).Qstd,2).^2/...
                size(modstruct.(modsinside{jj}).Qstd,2);
        end
        modout.Q = Qtot;
        modout.Qstd = sqrt(Qtotvar);
    else
        modout.Q = mean(mod1.Q,2);
        modout.Qstd = sqrt(sum(mod1.Qstd.^2,2)./size(mod1.Qstd,2));
    end
    
    %copy growth rate info
    if iscell(mod1.growthrate)
        gr0 = cell2mat(mod1.growthrate);
        gr0std = cell2mat(modout.growthratestd);
    else
        gr0 = mod1.growthrate;  %expdatastruct.(modnames{ii}).growthrate;
        gr0std = modout.growthratestd;
    end
    modout.growthrate = mean(gr0,2);
    modout.growthratestd = sqrt(var(gr0,0,2) + gr0std.^2);
    %expdatastruct.(modnames{ii}).growthratestd.^2);
    
    %output
    outstruct.(modnames{ii}) = modout;
end


if ploton
    [gr0,gr0std,Q0,Q0std] = deal(cell(length(modnames),1));
    for k = 1:length(modnames)
        %get data
        mod2 = outstruct.(modnames{k});
        if ~isempty(mod2.Q)
            gr0{k} = mean(mod2.growthrate,2);
            gr0std{k} = sqrt(sum(mod2.growthratestd.^2,2))./size(mod2.growthratestd,2);
            Q0{k} = mod2.Q;
            Q0std{k} = mod2.Qstd;
        end
    end
    
    Q = vertcat(Q0{:});
    Qstd = vertcat(Q0std{:});
    gr = vertcat(gr0{:});
    grstd = vertcat(gr0std{:});
    
    %set up logistic fitting problem
    opts = fitoptions('method','NonlinearLeastSquares','Robust','on',...
        'StartPoint',[0.6,1],'lower',[0,0],'upper',[1,Inf]);
    ft = fittype('a/(K*(1+x)^1+1)','coeff',{'a','K'},'options',opts);
    %opts = fitoptions('method','NonlinearLeastSquares','Robust','on',...
    %    'StartPoint',[0.6,1,2],'lower',[0,0,0],'upper',[1,Inf,5]);
    %ft = fittype('a/(K*(1+x)^n+1)','coeff',{'a','K','n'},'options',opts);
    [grfitobj,gof] = fit(Q,gr,ft);
    a = grfitobj.a
    K = grfitobj.K
    %b = grfitobj.b
    %n = grfitobj.n
    n = 1;
    
    Qfit = linspace(min(Q),max(Q),300)';
    grfit = a./(K*(1 + Qfit).^n + 1);
    %grfit = a*K./((1 + Qfit).^n + K);
    
    %plot
    figure(figh); clf;
    %plot(gr,Q,'o','linewidth',2); hold on;
    %errorbar(gr,Q,Qstd,'.k','vertical','linewidth',1.0)
    %errorbar(gr,Q,grstd,'.k','horizontal','linewidth',1.0)
    %ylabel('Resource Demand, Q [ ]')
	%plot(gr,Q./(1+Q),'o','linewidth',2); hold on;
    %errorbar(gr,Q./(1+Q),Qstd./(1+Q).^2,'.k','vertical','linewidth',1.0)
    %errorbar(gr,Q./(1+Q),grstd,'.k','horizontal','linewidth',1.0)
    %xlabel('Growthrate [1/hr]')
    %ylabel('Fraction of total resources bound [ ]')
    
    plot(Q,gr,'o','linewidth',2); hold on;
    errorbar(Q,gr,grstd,'.k','vertical','linewidth',1.0)
    errorbar(Q,gr,Qstd,'.k','horizontal','linewidth',1.0)
    plot(Qfit,grfit,'--','linewidth',1.2);
    ylabel('Growthrate [1/hr]')
    xlabel('Q [ ]')
    set(gca,'fontsize',14,'linewidth',2)
    %xlim([0,inf])
    %ylim([0,inf])
    box off
else
    grfitobj = [];
    gof = [];
end
```
