## Supplementary material for "Predicting Composition of Genetic Circuits with Resource Competition: Demand and Sensitivity": Experimental data and Matlab code: plotstruct.m

```
function plotstruct(cellstruct,xfield,yfield,figh,cellfields)
%plotstruct(cellstruct,xfield,yfield,figh,cellfields)
%plot subfields from a struct for all fields
%xfield and yfield may be cell arrays. The plot uses subplot to plot each
%xfield and yfield against each other.

if nargin < 4 || isempty(figh)
    figh = figure;
end
if nargin < 5 || isempty(cellfields)
    cellnames = fieldnames(cellstruct);
else
    cellnames = cellfields;
end

linewdth = 1.5;

figure(figh); clf;
if iscell(xfield) && length(xfield) > 1 && iscell(yfield) && length(yfield) > 1
    for k = 1:length(yfield)
        for jj = 1:length(xfield)
            subplot(length(yfield),length(xfield),jj + (k-1)*length(xfield));
            for ii = 1:length(cellnames)
                %make plots
                plot(cellstruct.(cellnames{ii}).(xfield{jj}),...
                    cellstruct.(cellnames{ii}).(yfield{k}),'linewidth',linewdth);
                hold on
            end
            legend(cellnames,'Location','Best')
            hold off
            xlabel(xfield{jj})
            ylabel(yfield{k})
            set(gca,'fontsize',12)
        end
    end
elseif iscell(xfield) && length(xfield) > 1
    for jj = 1:length(xfield)
        subplot(1,length(xfield),jj);
        for ii = 1:length(cellnames)
            %make plots
            plot(cellstruct.(cellnames{ii}).(xfield{jj}),...
                cellstruct.(cellnames{ii}).(yfield),'linewidth',linewdth);
            hold on
        end
        legend(cellnames,'Location','Best')
        hold off
        ylabel(yfield)
        xlabel(xfield{jj})
        set(gca,'fontsize',12)
    end
    
elseif iscell(yfield) && length(yfield) > 1
    for jj = 1:length(yfield)
        subplot(length(yfield),1,jj);
        for ii = 1:length(cellnames)
            %make plots
            plot(cellstruct.(cellnames{ii}).(xfield),...
                cellstruct.(cellnames{ii}).(yfield{jj}),'linewidth',linewdth);
            hold on
        end
        legend(cellnames,'Location','Best')
        hold off
        xlabel(xfield)
        ylabel(yfield{jj})
        set(gca,'fontsize',12)
    end
else
    for ii = 1:length(cellnames)
        %make plots
        plot(cellstruct.(cellnames{ii}).(xfield),...
            cellstruct.(cellnames{ii}).(yfield),'linewidth',2);
        hold on
    end
    legend(cellnames,'Location','Best')
    hold off
    xlabel(xfield)
    ylabel(yfield)
    set(gca,'fontsize',12)
end
```
