## Supplementary material for "Predicting Composition of Genetic Circuits with Resource Competition: Demand and Sensitivity": Experimental data and Matlab code: plotsubfield2.m

```
function [figh,axh] = plotsubfield2(cellstruct,xfield,yfield,figh,cellfields,ssstruct,yfun,fntsze)
%[figh,axh] = plotsubfield2(cellstruct, xfield, yfield,figh, cellfields)
%plot one subfield for each field in a struct. datafield must match to a
%valid field for all substructs contined in cellstruct

%input verification
if nargin < 4 || isempty(figh)
    figh = figure;
end
if nargin < 7 || isempty(yfun)
    yfun = @(x) x;
end

%labeling
if strcmp(xfield,'time')
    xfieldlabel = 'time [hrs]';
else
    xfieldlabel = xfield;
end
if any(contains(yfield,'diffOD'))
    yfieldlabel = [yfield(1:3),' production rate [a.u.]'];
else
    yfieldlabel = yfield;
end

%get fields for each cell type
celltypes = fieldnames(cellstruct);
if nargin > 4 && ~isempty(cellfields)
    cellnames = intersect(cellfields,celltypes,'stable');
else; cellnames = celltypes;
end

n = length(cellnames);                  %number of subplots (fields) to make
m = ceil(9/16*sqrt(n));                 %number of rows of subplots

if nargin < 8 || isempty(fntsze)
    if n*m > 20; fntsze = 10;
    else; fntsze = 12;
    end
end

%plot each struct's subfield on one plot
figure(figh);
axh = cell(n,1);
for jj = 1:n
    %make subplot
    axh{jj} = subplot(m,ceil(n/m),jj);
    %keep hold on if already set
    holdon = ishold;
    if holdon; hold on; end
    %plot
    xdata = cellstruct.(cellnames{jj}).(xfield);
    ydata = cellstruct.(cellnames{jj}).(yfield);
    if ~isempty(ydata)
        plot(xdata,yfun(ydata),'linewidth',1);
    else
        cla;
        title(strrep(cellnames{jj},'_','-'))
        continue
    end
    
    %put marks where steady state was chosen
    if nargin >= 6 && ~isempty(ssstruct) && isfield(ssstruct,cellnames{jj}) ...
            && ~isempty(ssstruct.(cellnames{jj}))...
            && max(ssstruct.(cellnames{jj}),[],'all') <= max(xdata,[],'all') ...
            && min(ssstruct.(cellnames{jj}),[],'all') >= min(xdata,[],'all')
        hold on;
        ssdata = ssstruct.(cellnames{jj});
        inds = xdata > min(ssdata) & xdata < max(ssdata);
        yssmax = max(yfun(ydata(inds,:)),[],'all');
        yssmin = min(yfun(ydata(inds,:)),[],'all');
        plot([ssdata(1),ssdata(1)],[0.9*yssmin,1.1*yssmax],'-k',...
            [ssdata(2),ssdata(2)],[0.9*yssmin,1.1*yssmax],'-k','linewidth',0.8)
        if ~holdon; hold off; end
    end
    
    title(strrep(cellnames{jj},'_','-'))
    %put ylabels only on farthest left plots
    if mod(jj,ceil(n/m)) == 1; ylabel(strrep(yfieldlabel,'_',' ')); end
    %put xlabels only on bottom plots
    if n - jj < ceil(n/m); xlabel(strrep(xfieldlabel,'_',' ')); end
    set(gca,'fontsize',fntsze)
    hold off
end
```
