## Supplementary material for "Predicting Composition of Genetic Circuits with Resource Competition: Demand and Sensitivity": Experimental data and Matlab code: predictRS6.m

```
function dataout = predictRS6(moduledata,ploton,fighvec,includeinduced,fighindpert)
%predict outputs

if nargin < 3 || isempty(ploton)
    ploton = false;
end
if ploton && nargin < 3
    fighvec = 1:6;
end
if ploton && nargin < 4
    %include induced mod perturbations in constitutive mod predictions
    includeinduced = true;
end
if ploton && nargin < 5
    fighindpert = max(fighvec) + 1;
end

%find groups of non-RS modules that share properties
modsalone = findpropRS(moduledata,'isalone');
modstogether = findpropRS(moduledata,'isalone',false);
normalmods = findpropRS(moduledata,'isResourceSensor',false);  %not resource sensors
%experiments that are modules together and one module isn't a resource sensor
normalmodstogeth = intersect(normalmods,modstogether);

dataout = moduledata;

for k = 1:length(normalmodstogeth)
    
    %unpackage data about modules in pairs
    modnames = moduledata.(normalmodstogeth{k}).containingmods;
    FPfields = moduledata.(normalmodstogeth{k}).FPout;
    
    %get measured module outputs alone and with perturbation
    y1 = moduledata.(modnames{1}).y;
    y1std = moduledata.(modnames{1}).ystd;
    y2 = moduledata.(modnames{2}).y;
    y2std = moduledata.(modnames{2}).ystd;
    y1p = moduledata.(normalmodstogeth{k}).(FPfields{1});
    y1pstd = moduledata.(normalmodstogeth{k}).([FPfields{1},'std']);
    y2p = moduledata.(normalmodstogeth{k}).(FPfields{2});
    y2pstd = moduledata.(normalmodstogeth{k}).([FPfields{2},'std']);
    
    %resource properties of module 1
    %average together if more than one Q or S measured
    Q1 = mean(moduledata.(modnames{1}).Q,2);
    Q1std = mean(moduledata.(modnames{1}).Qstd,2);
    S1 = mean(moduledata.(modnames{1}).S,2);
    S1std = mean(moduledata.(modnames{1}).Sstd,2);
    %resource properties of module 2
    %average together if more than one Q or S measured
    Q2 = mean(moduledata.(modnames{2}).Q,2);
    Q2std = mean(moduledata.(modnames{2}).Qstd,2);
    S2 = mean(moduledata.(modnames{2}).S,2);
    S2std = mean(moduledata.(modnames{2}).Sstd,2);
    
    if length(moduledata.(modnames{1}).u) > 1
        %make predictions (constitutive modules, S = 0)
        [y1pred,y1predstd] = calcypred5(y1,0,Q1,Q2,y1std,0,Q1std,Q2std);
    else
        %make predictions (non-constitutive modules, S neq 0)
        [y1pred,y1predstd] = calcypred5(y1,S1,Q1,Q2,y1std,S1std,Q1std,Q2std);
    end
    if length(moduledata.(modnames{2}).u) > 1
        %make predictions (constitutive modules, S = 0)
        [y2pred,y2predstd] = calcypred5(y2,0,Q2,Q1,y2std,0,Q2std,Q1std);
    else
        %make predictions (non-constitutive modules, S neq 0)
        [y2pred,y2predstd] = calcypred5(y2,S2,Q2,Q1,y2std,S2std,Q2std,Q1std);
    end
    
    %save perturbations
    prevpredy1 = dataout.(modnames{1}).predperturby;
    prevpredy2 = dataout.(modnames{2}).predperturby;
    prevpredy1std = dataout.(modnames{1}).predperturbystd;
    prevpredy2std = dataout.(modnames{2}).predperturbystd;
    prevperturby1 = dataout.(modnames{1}).perturby;
    prevperturby2 = dataout.(modnames{2}).perturby;
    prevperturby1std = dataout.(modnames{1}).perturbystd;
    prevperturby2std = dataout.(modnames{2}).perturbystd;
    prevperturbname1 = dataout.(modnames{1}).perturbname;
    prevperturbname2 = dataout.(modnames{2}).perturbname;
    
    %append predictions and perturbed outputs into output data object
    %for module 1
    if ~isempty(prevpredy1)
        dataout.(modnames{1}).predperturby    = [prevpredy1,{y1pred}];
        dataout.(modnames{1}).predperturbystd = [prevpredy1std,{y1predstd}];
    else
        dataout.(modnames{1}).predperturby    = {y1pred};
        dataout.(modnames{1}).predperturbystd = {y1predstd};
    end
    dataout.(modnames{1}).perturbname = [prevperturbname1,modnames{2}];
    dataout.(modnames{1}).perturby    = [prevperturby1,y1p];
    dataout.(modnames{1}).perturbystd = [prevperturby1std,y1pstd];
    
    %for module 2
    if ~isempty(prevpredy2)
        dataout.(modnames{2}).predperturby    = [prevpredy2,{y2pred}];
        dataout.(modnames{2}).predperturbystd = [prevpredy2std,{y2predstd}];
    else
        dataout.(modnames{2}).predperturby    = {y2pred};
        dataout.(modnames{2}).predperturbystd = {y2predstd};
    end
    dataout.(modnames{2}).perturbname = [prevperturbname2,modnames{1}];
    dataout.(modnames{2}).perturby    = [prevperturby2,y2p];
    dataout.(modnames{2}).perturbystd = [prevperturby2std,y2pstd];
end

%plot
if ploton
    if length(fighvec) < length(modsalone)
        fighvec = fighvec(1):(fighvec(1)+length(modsalone)-1);
        warning(['fighvec length less than the number of modules. Using figures',...
            num2str(fighvec)])
    end
    for jj = 1:length(modsalone)
        modplt = dataout.(modsalone{jj});
        %assuming all mods alone are modules contained in a pair
        if length(modplt.u) > 1
            %inducible module prediction plot
            uvec = modplt.u;
            inds = cellfun(@(x) ~isempty(x),modplt.predperturby);
            %get data
            y = abs(modplt.y);
            ystd = modplt.ystd./max(y);
            yp = abs([modplt.perturby{inds}]./max(y));
            ypstd = [modplt.perturbystd{inds}]./max(y);
            pred = abs([modplt.predperturby{inds}]./max(y));
            predstd = [modplt.predperturbystd{inds}]./max(y);
            
            %reshape to make bar plots
            [yppred,yppredstd] = deal(zeros(size(yp,1),2*size(yp,2)));
            for k = 1:size(yp,2)
                yppred(:,2*k-1) = yp(:,k);
                yppred(:,2*k) = pred(:,k);
                yppredstd(:,2*k-1) = ypstd(:,k);
                yppredstd(:,2*k) = predstd(:,k);
            end
            %note all predictions are stacked together
            dataind = [y./max(y),yppred];
            dataindstd = [ystd,yppredstd];
            
            figure(fighvec(jj)); clf;
            %plot bar chart
            bh = plotbarpredict(dataind',dataindstd');
            %make predictions and measurements same color
            for l = 1:size(yp,2)
                set(bh(2*l),'FaceColor',min(get(bh(1+2*l),'FaceColor')*1.4,1))
            end
            set(gca,'yscale','linear','xticklabels',uvec,'fontsize',14)
            ylim([-0.1,1.2])
            ylabel(modplt.FPout)
            xlabel('intput, u [nM]')
            legend([bh(2:2:end),bh(3:2:end)],[...
                combinenames(modplt.perturbname(3:end),{'measured'},false),...
                combinenames(modplt.perturbname(3:end),{'predicted'},false)],'Location','Best')
        else
            %constitutive module predictions
            y = modplt.y;
            ystd = modplt.ystd;
            %indicies where module is perturbed by a non-resource sensor
            %constitutive module indicies
            indsC = cellfun(@(x) ~isempty(x) & size(x,1) == 1,modplt.predperturby);
            %indicies where module is perturbed by non-RS inducible module
            indsI = cellfun(@(x) ~isempty(x) & size(x,1) > 1,modplt.predperturby);
            %shape data and predictions into matrix for plotting
            yp = vertcat(modplt.perturby{indsC});
            datamat0 = vertcat(modplt.predperturby{indsC});
            %1st column = measured; 2nd = predicted with S1, 3rd = with S2, 4th = with S3
            datamat  = [y,NaN(1,size(datamat0,2)); [yp,datamat0]]./y;
            ypstd = vertcat(modplt.perturbystd{indsC});
            datamat0std = vertcat(modplt.predperturbystd{indsC});
            datamatstd  = [ystd,NaN(1,size(datamat0std,2)); [ypstd,datamat0std]]./y;
            
            %plot constitutive module predictions
            figure(fighvec(jj)); clf;
            bh = plotbarpredict(datamat',datamatstd');
            ylabel(modplt.FPout)
            names = combinenames(modsalone(jj),modplt.perturbname(indsC),true);
            xticklabels(names)
            set(gca,'fontsize',14)
            ylim([-0.1,1.2])
            colorplots(modplt.FPout,bh)
            legend('measured','predicted','location','best')
            
            %outputs of constitutive modules perturbed by induced module
            if any(indsI) && includeinduced
                %plot curves for induced module perturbing constitutive one
                figure(fighindpert);
                if jj == 1; clf; end
                subplot(2,3,jj);
                %loop through induced perturbing modules
                indsI2 = find(indsI);
                for k = indsI2
                    %get u vec from perturbing module
                    perturbname = modplt.perturbname{k};
                    uvec = dataout.(perturbname).u;
                    %get data
                    y = modplt.y;
                    y2 = modplt.y*ones(size(uvec))/y;           %cast up to correct size
                    y2std = modplt.ystd*ones(size(uvec))/y;     %cast up to correct size
                    yp = modplt.perturby{k}/y;
                    ypstd = modplt.perturbystd{k}/y;
                    pred = modplt.predperturby{k}/y;
                    predstd = modplt.predperturbystd{k}/y;
                    
                    %make bar chart
                    dataind = [y2',yp,pred];
                    dataindstd = [y2std',ypstd,predstd];
                    bh = plotbarpredict(dataind',dataindstd');
                    
                    ylim([0,1.2])
                    xticklabels(uvec)
                    set(gca,'fontsize',14)
                    ylabel(modplt.FPout)
                    title([modplt.containingmods{1},'+',perturbname])
                    xlabel('input, u [nM]')
                    colorplots(modplt.FPout,bh)
                    legend('alone','measured','predicted','location','best')
                end
            end
        end
    end
end

function [pred,predstd] = calcypred5(y1,S1,Q1,Q2,ystd,Sstd,Q1std,Q2std)
%use resource demand to predict circuit expression
pred = y1.*(1 + S1.*Q2).*(1+Q1)./(1+Q1+Q2);

%calculate standard error on predicted output
dfdy = (1+S1*Q2).*(1+Q1)./(1+Q1+Q2);
dfdS = y1*Q2.*(1+Q1)./(1+Q1+Q2);
dfdQ1 = y1.*(1+S1.*Q2).*Q2./(1+Q1+Q2).^2;
dfdQ2 = -y1.*(S1.*(1+Q1)./(1+Q1+Q2) - (1+S1.*Q2).*(1+Q1)./(1+Q1+Q2).^2);
predstd = sqrt(dfdy.^2.*ystd.^2 + dfdS.^2.*Sstd.^2 + dfdQ1.^2.*Q1std.^2 + ...
    dfdQ2.^2.*Q2std.^2);


function bh = plotbarpredict(data,datastd)
nbars = size(data,1);
ngroups = size(data,2);
groupwidth = min(0.8, nbars/(nbars + 1.5));
% Set the position of each error bar in the centre of the main bar
% Based on barweb.m by Bolu Ajiboye from MATLAB File Exchange
x = zeros(nbars,ngroups);
for ii = 1:nbars
    % Calculate center of each bar
    x(ii,:) = (1:ngroups) - groupwidth/2 + (2*ii-1) * groupwidth / (2*nbars);
end
bh = bar(data'); hold on;
set(bh,'edgecolor','none');
errorbar(x,data,datastd,'.k','linestyle','none','Linewidth',1.5);


function ph = plotinducedmod(uvec,yalone,yalonestd,yp,ypstd,ypred,ypredstd)
%plot Q or S curves vs input u
zerooffset = 1e-2;
ax = gca;
ph = cell(2,1);
umax = max(uvec,[],'all');
umin = max([min(uvec,[],'all'),zerooffset]);

%plot y alone
semilogx(max(uvec,zerooffset),yalone,'-k','linewidth',2); hold on
errorbar(max(uvec,zerooffset),yalone,yalonestd,'.k','linewidth',0.5);
set(ax,'ColorOrderIndex',1)
%plot actual y perturb
coi = get(ax,'colororderindex');
ph{1} = semilogx(max(uvec,zerooffset),yp,'linewidth',2); hold on
for jj = 1:size(yp,2)
    x = max(uvec,zerooffset).*(1 + 0*rand(size(uvec)));
    errorbar(x,yp(:,jj),ypstd(:,jj),'.k','linewidth',0.5);
end
%plot predictions
set(ax,'colororderindex',coi);
ph{2} = semilogx(max(uvec,zerooffset),ypred,'--','linewidth',2); hold on
for ii = 1:size(ypred,2)
    x = max(uvec,zerooffset).*(1 + 0*rand(size(uvec)));
    errorbar(x,ypred(:,ii),ypredstd(:,ii),'.k','linewidth',0.5);
end
%change zero tick mark
set(gca,'fontsize',14);
if iscell(uvec) && cellfun(@(x) any(x == 0),uvec)
    xt = xticks;
    xticklabels([0,xt(2:end)])
elseif any(uvec == 0)
    xt = xticks;
    xticklabels([0,xt(2:end)])
end
xlim([umin,umax])


function colorplots(name,bh)

if contains(name,'BFP')
    %make colors match
    blue1 = [91,155,213]/255;       %dark blue
    blue2 = [125,195,251]/255;      %medium blue
    blue3 = [173,222,255]/255;      %light blue
    set(bh(1),'FaceColor',blue1)    %measured
    set(bh(2),'FaceColor',blue3)    %predicted1
    if length(bh) >= 3
        set(bh(3),'FaceColor',blue2)    %predicted2
    end
elseif contains(name,'GFP')
    green1 = [120,171,48]/255;      %dark green
    green2 = [45,190,6]/255;        %emerald green
    green3 = [163,252,67]/255;      %light green
    set(bh(1),'FaceColor',green2)   %measured
    set(bh(2),'FaceColor',green3)   %predicted1
    if length(bh) >= 3
        set(bh(3),'FaceColor',green1)   %predicted2
    end
elseif contains(name,'RFP')
    red1 = [207,81,81]/255;         %dark red
    red2 = [240,122,122]/255;       %light red
    red3 = [255,168,168]/255;       %lighter red
    set(bh(1),'FaceColor',red1)     %measured
    set(bh(2),'FaceColor',red3)     %predicted1
    if length(bh) >= 3
        set(bh(3),'FaceColor',red2)     %predicted2
    end
elseif contains(name,'YFP')
    yellow1 = [237,177,32]/255;
    yellow2 = [255,255,0]/255;
    yellow3 = [196,200,21]/255;
    set(bh(1),'FaceColor',yellow1)     %measured
    set(bh(2),'FaceColor',yellow2)     %predicted1
    if length(bh) >= 3
        set(bh(3),'FaceColor',yellow3)     %predicted2
    end
end
```
