## Supplementary material for "Predicting Composition of Genetic Circuits with Resource Competition: Demand and Sensitivity": Experimental data and Matlab code: replaceoutliers.m

```
function outstruct = replaceoutliers(instruct,datafield)


cellnames = fieldnames(instruct);
outstruct = instruct;

%loop through experiments
for ii = 1:length(cellnames)
    
    datanames0 = fieldnames(instruct.(cellnames{ii}));
    datanames = intersect(datanames0,datafield);
    
    %loop through data fields to smooth
    for jj = 1:length(datanames)
        
        indata = instruct.(cellnames{ii}).(datanames{jj});
        outdata = filloutliers(indata,'linear','movmean',5);
        outstruct.(cellnames{ii}).(datanames{jj}) = outdata;
    end
end
```
