## Supplementary material for "Predicting Composition of Genetic Circuits with Resource Competition: Demand and Sensitivity": Experimental data and Matlab code: savefigpdf.m

```
function savefigpdf(filename,figh,figformat,fntsze)
%savefigpdf(filename,figh,figformat,fntsze)
%save figure defined by FIGH to pdf with desired settings. FILENAME is the
%name of the file you want to output (not including .pdf). Saves the figure
%to the current folder

%filename
if nargin < 1
    filename = 'matlabfig';
end
%choose options
if nargin < 2 || isempty(figh)
    figh = gcf;
end
%output format
if nargin < 3
    figformat = 'pdf';
end
%apply settings to figure
if isnumeric(figh); fig = figure(figh);
elseif ishandle(figh); fig = figh;
else; fig = gcf;
end

if nargin > 3 && ~isempty(fntsze)
    %apply for all subplots
    for ii = 1:length(fig.Children)
        set(fig.Children(ii),'FontSize',fntsze)
    end
end

%remove extension
if any(contains(filename,{'.pdf','.fig'}))
    filename = filename(1:end-4);
end
minfntsze = 10;

%output style
myStyle = hgexport('factorystyle');
myStyle.Format = figformat;
myStyle.Resolution = 300;
myStyle.Units = 'inch';
% myStyle.FixedFontSize = fntsze;
myStyle.FontSizeMin = minfntsze;
myStyle.FontName = 'Helvetica';
myStyle.Bounds = 'tight';
myStyle.LineWidthMin = 1.5;
% myStyle.LockAxesTicks = 'on';         %makes factors of 10 above axes disappear
% set(fig,'PaperUnits','inches','PaperSize',[6,3.75]/2)
pos = get(fig,'PaperPosition');
set(fig,'PaperSize',[pos(3)*1.05,pos(4)*1.05]);

%output figure as png and matlab figure
hgexport(fig,[filename,'.',figformat],myStyle,'Format',figformat)
savefig(fig,[filename,'.fig']);
```
