## Supplementary material for "Predicting Composition of Genetic Circuits with Resource Competition: Demand and Sensitivity": Experimental data and Matlab code: setallsubplots.m

```
function obj = setallsubplots(figh, proptype, namearray, valuearray)
%obj = setallsubplots(FIGH, PROPTYPE, NAMEARRAY, VALUEARRAY)
%Change the properties of all children in the figure. May apply for any
%figure properties, axis properties, or line properties.
%Valid options for PROPTYPE are 'figure', 'axis', 'line', or 'legend'.
%NAMEARRAY may be any array of valid property fields for the correpsonding 
%figure, axis, or line properties. VALUEARRAY is the value to set the 
%desired property for all subplots in FIGH. OBJ returns a cell array 
%containing the handles for the objects that were set

if nargin < 4
    valuearray = 'log';
    if nargin < 3
        namearray = 'yscale';
        if nargin < 2
            proptype = 'axis';
            if nargin < 1
                figh = [];
            end
        end
    end
end

%input error checking
if isnumeric(figh) && ~isempty(figh)
    %if scalar, apply to one figure
    if length(figh) == 1
        figh = figure(figh);
    else
        %if vector, apply to all figures in vector
        for k = 1:length(figh)
            setallsubplots(figh(k), proptype, namearray, valuearray)
        end
        return
    end
elseif isgraphics(figh)
    %get figure handle
    figh = figure(figh);
elseif ischar(figh) && strcmp(figh,'all')
    %set properties for all active figures
    fighandles = get(groot,'Children');
    for k = 1:length(fighandles)
        setallsubplots(fighandles(k), proptype, namearray, valuearray);
    end
    return
else
    figh = gcf;
end

%apply property to class
if strcmpi(proptype,'line')
    obj = cell(length(figh.Children),1);
    for ii = 1:length(figh.Children)
        if strcmp(figh.Children(ii).Type,'axes')
            %only go into axes objects
            lineobj = cell(length(figh.Children(ii).Children),1);
            for jj = 1:length(figh.Children(ii).Children)
                %set line properties
                set(figh.Children(ii).Children(jj),namearray,valuearray)
                lineobj{jj} = figh.Children(ii).Children(jj);
            end
        else; lineobj = [];    
        end
        obj{ii} = lineobj;
    end
elseif strcmpi(proptype,'Legend')
    obj = cell(length(figh.Children),1);
    %loop through children only taken 'Legend' children
    for ii = 1:length(figh.Children)
        %set properties for Legends
        if strcmp(figh.Children(ii).Type,'Legend')
            set(figh.Children(ii),namearray,valuearray)
            obj{ii} = figh.Children(ii);
        end
    end
elseif strcmpi(proptype,'axis')
    %loop throuh children
    obj = cell(length(figh.Children),1);
    for ii = 1:length(figh.Children)
        %set only axis properties
        if strcmp(figh.Children(ii).Type,'axes')
            set(figh.Children(ii),namearray,valuearray)
            obj{ii} = figh.Children(ii);
        end
    end
elseif strcmpi(proptype,'figure')
    %set figure properties
    set(figh,namearray,valuearray)
    obj = figh;
end
```
