## Supplementary material for "Predicting Composition of Genetic Circuits with Resource Competition: Demand and Sensitivity": Experimental data and Matlab code: subtractbasal7.m

```
function [outstruct2,basalstruct] = subtractbasal7(cellstruct,mediastruct,ODcontrolstruct,subfields)
%[outstruct,basalstruct] = subtractbasal7(cellstruct,mediastruct,cellcontrol,subfields)
%find basal level for each property by assuming only media contributes to
%background (ignores fluorescence of cells).
%subtract off average background fluorescence and OD levels from raw data

if nargin < 2 || isempty(mediastruct)
    %struct to determine background media contribution for each datafield
    mediastruct = struct;
    mediastruct.OD  = 'M9';
    mediastruct.BFP = 'M9';
    mediastruct.GFP = 'M9';
    mediastruct.RFP = 'M9';
end
if nargin < 3 || isempty(ODcontrolstruct)
    ODcontrolstruct = [];
end
if nargin < 4 || isempty(subfields)
    subfields = fieldnames(mediastruct);
end

%init
outstruct = struct;
cellnames = fieldnames(cellstruct);

%find basal levels from media for each channel
basalstruct = findbasalvstime(cellstruct,mediastruct);

%subtract basal levels off data
for ii = 1:length(cellnames)
    outobj = RSexpdata;
    celldata = cellstruct.(cellnames{ii});
    %loop across each cell type
    dprops = fieldnames(celldata);
    for jj = 1:length(dprops)
        %if matches desired fields, subtract fields
        if any(strcmpi(dprops{jj},subfields))
            outobj.(dprops{jj}) = celldata.(dprops{jj}) - basalstruct.(dprops{jj});
        else
            %copy remaining data
            outobj.(dprops{jj}) = celldata.(dprops{jj});
        end
    end
    %output
    outstruct.(cellnames{ii}) = outobj;
end

if isempty(ODcontrolstruct)
    %if no cell OD control, return
    outstruct2 = outstruct;
    return
end

%subtract basal fluorescent levels based on control cells as a function of OD
outstruct2 = struct;
ODctrl = makeFPbasalOD(outstruct,ODcontrolstruct);
%apply to each cell type
for ii = 1:length(cellnames)
    outobj2 = RSexpdata;
    celldata2 = outstruct.(cellnames{ii});
    %loop across each cell type
    dprops2 = fieldnames(celldata2);
    OD = celldata2.OD;
    for jj = 1:length(dprops2)
        %if matches desired fields excluding OD, subtract fields
        if any(strcmpi(dprops2{jj},setdiff(subfields,'OD')))
            outobj2.(dprops2{jj}) = celldata2.(dprops2{jj}) - ODctrl.(dprops2{jj})(OD);
        else
            %copy remaining data
            outobj2.(dprops2{jj}) = celldata2.(dprops2{jj});
        end
    end
    %output
    outstruct2.(cellnames{ii}) = outobj2;
end


function basalstruct = findbasalvstime(cellstruct,mediastruct)
%find basal levels of each property based on media measurements

basalstruct = struct;
dprops = fieldnames(mediastruct);

%loop across properties
for ii = 1:length(dprops)
    %take average of all media fields
    mediafld = mediastruct.(dprops{ii});
    if iscell(mediafld) && length(mediafld) == 1
        mediafld = mediafld{1};
    elseif iscell(mediafld) && length(mediafld) > 1
        error('only 1 media field allowed and it must be a string')
    end
    tmp = cellstruct.(mediafld).(dprops{ii});
    %filter out high frequency noise
    %[b,a] = butter(3,0.3,'low');
    %filtfilt(b,a,mean(tmp,2));
    %tmp2 = mean(tmp,[],'all');
    basalFP = mean(tmp,'all');
    %col vector with same length as 'time'
    basalstruct.(dprops{ii}) = basalFP*ones(size(tmp,1),1);
end


function [out,ODout] = makeFPbasalOD(cellstruct,ODcontrolstruct)
%make single OD vs FP curve for each channel by combining all measurements

controlOD = cellstruct.(ODcontrolstruct.OD).OD;

%make a standard OD curve
minOD = min(controlOD,[],'all');
maxOD = max(controlOD,[],'all');
ODout = linspace(minOD,maxOD,200)';     %standardized OD curve

dprops = fieldnames(ODcontrolstruct);

out = struct;

for jj = 1:length(dprops)
    nonefld = ODcontrolstruct.(dprops{jj});
    FP = cellstruct.(nonefld).(dprops{jj});
    FPnew = zeros(length(ODout),size(FP,2));
    %loop across measurements
    for ii = 1:size(FP,2)
        %line up fluorescence measurements with the same ODs and average
        [ODtmp,~,idx] = unique(controlOD(:,ii),'stable');
        FPtmp = accumarray(idx,FP(:,ii),[],@mean);
        %interpolate to get standard fluorescence curve
        FPnew(:,ii) = interp1(ODtmp,FPtmp,ODout,'nearest','extrap');
    end
    %take the average over all measurements with low pass filter
    [b,a] = butter(3,0.2,'low');
    FPout = filtfilt(b,a,mean(FPnew,2));
    out.(dprops{jj}) = @(OD) interp1(ODout,FPout,OD,'linear');
end
```
