## Supplementary figures and images for "Predicting Composition of Genetic Circuits with Resource Competition: Demand and Sensitivity"

### smooth_diff.jpg

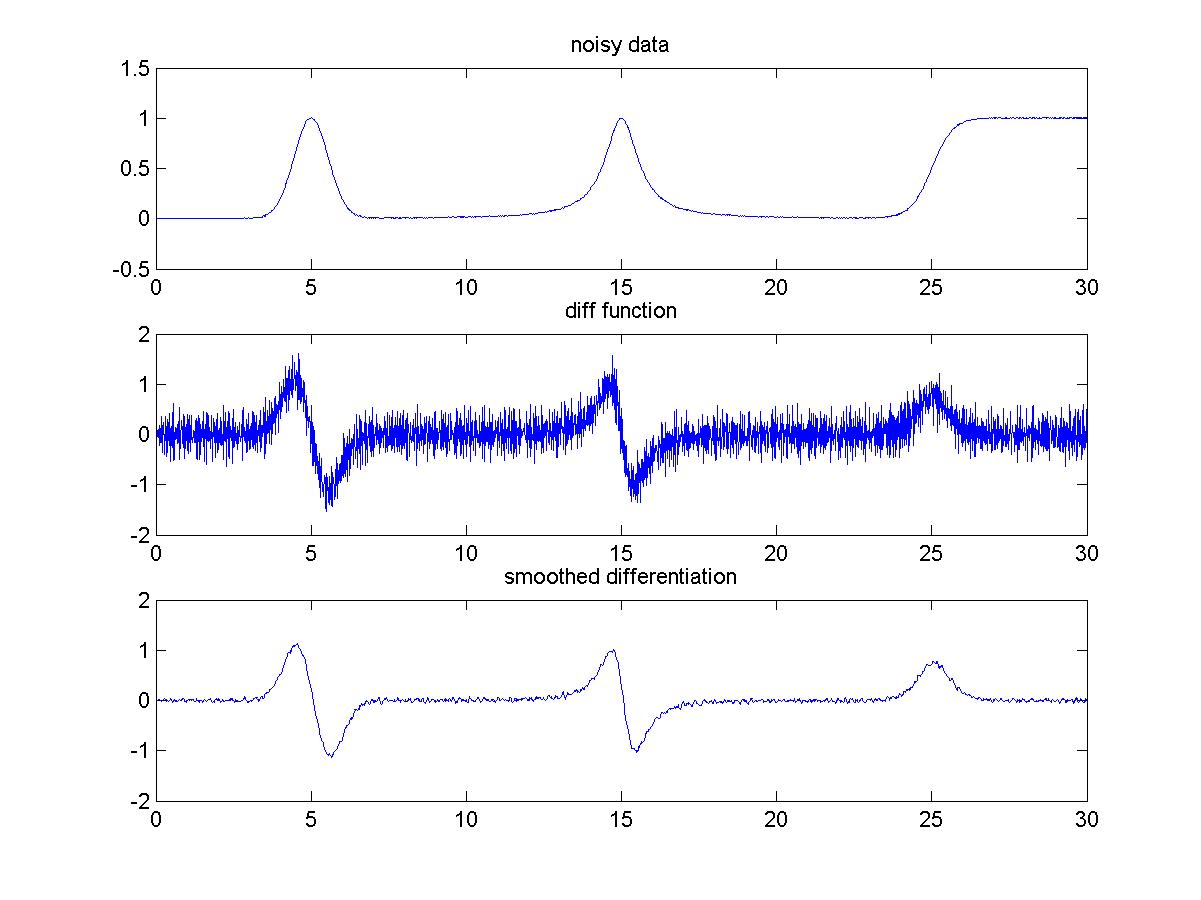
