## Supplementary Information pdf for "Predicting Composition of Genetic Circuits with Resource Competition: Demand and Sensitivity"

### S1 Mathematical Analysis

#### S1.1 Derivation of the Resource Competition Model

Here, we derive the model describing resource competition in genetic circuits beginning from the chemical reactions describing protein production from DNA utilizing a shared resource. Protein production is modeled as a two-step process taking DNA to mRNA (transcription) then to protein (translation). More details can be found in (*1*, *2*). The chemical reactions describing the process of protein production for gene $i$ in a genetic circuit are

$$\mathrm{D}_i + \mathrm{P} \underset{d_{1i}}{\overset{a_{1i}F_i(u)}{\rightleftharpoons}} \mathrm{C}_{1i} \xrightarrow{k_{1i}} \mathrm{P} + \mathrm{D}_i + \mathrm{m}_i \tag{S1a}$$

$$\mathrm{m}_i + \mathrm{R} \underset{d_{2i}}{\overset{a_{2i}}{\rightleftharpoons}} \mathrm{C}_{2i} \xrightarrow{k_{2i}} \mathrm{R} + \mathrm{m}_i + \mathrm{x}_i \tag{S1b}$$

$$\mathrm{m}_i \xrightarrow{\delta} \emptyset \qquad \mathrm{x}_i \xrightarrow{\gamma} \emptyset, \tag{S1c}$$

where $\mathrm{D}_i$ represents the DNA for gene $i$ (copy number), P represents the RNA polymerase (RNAP), $\mathrm{C}_{1i}$ represents the complex of DNA bound to the RNAP, $\mathrm{m}_i$ represents the mRNA for gene $i$, R represents the ribosome, $\mathrm{C}_{2i}$ represents the complex of mRNA bound to the ribosome, and $\mathrm{x}_i$ represents the protein output of gene $i$. Additionally $a_{1i}$ and $d_{1i}$ are the association and dissociation constants of DNA with RNAP, respectively, $k_{1i}$ is the catalytic rate of DNA transcription, $a_{2i}$ and $d_{2i}$ are the association and dissociation constants of mRNA with ribosomes, $k_{2i}$ is the catalytic rate of mRNA translation, $\delta$ is the dilution and degradation rate of mRNA, and $\gamma$ is the protein decay rate, which we assume is completely due to dilution. The association of DNA with RNAP depends on the concentration of some set of inputs, $u_i$, which regulates gene $i$ according to the function $F_i(u_i)$ and is a sigmoidal function (*2*). Then using the law of mass action, the dynamics of the system are given as

$$\frac{dC_{1i}}{dt} = a_{1i}F_i(u_i)D_iP - (d_{1i} + k_{1i})C_{1i} \tag{S2}$$

$$\frac{dC_{2i}}{dt} = a_{2i}m_iR - (d_{2i} + k_{2i})C_{2i} \tag{S3}$$

$$\frac{dm_i}{dt} = k_{1i}C_{1i} - a_{2i}m_iR + (d_{2i} + k_{2i})C_{2i} - \delta m_i \tag{S4}$$

$$\frac{dx_i}{dt} = k_{2i}C_{2i} - \gamma x_i, \tag{S5}$$

where the DNA and cellular resources obey the conservation laws

$$D_{i_{tot}} = D_i + C_{1i} \tag{S6}$$

$$P_{tot} = P + \sum_{i=1}^{n} C_{1i} \tag{S7}$$

$$R_{tot} = R + \sum_{i=1}^{n} C_{2i}. \tag{S8}$$

Here, $D_{i_{tot}}$ is the total concentration of DNA for gene $i$, $P_{tot}$ is the total concentration of RNAP, and $R_{tot}$ is the total concentration of ribosomes serving genes 1 through $n$. We assume that binding and unbinding reactions occur much faster than protein degradation and dilution and that mRNA degradation occurs much faster than protein degradation. This allows us to set dynamics of the complexes $C_{1i}$ and $C_{2i}$ as well as the mRNA dynamics $m_i$ to their quasi-steady states (QSS). Then, solving for the QSS of $C_{1i}$, $C_{2i}$, and $m_i$ in Eqs. (S2), (S3), and (S4), we have that the quasi-steady state concentrations are

$$\overline{C}_{1i} = \frac{D_iPF_i(u_i)}{K_{1i}} \tag{S9a}$$

$$\overline{C}_{2i} = \frac{m_iR}{K_{2i}} \tag{S9b}$$

$$\overline{m}_i = \frac{k_{1i}}{\delta}\overline{C}_{1i}, \tag{S9c}$$

where $K_{1i} = \frac{d_{1i}+k_{1i}}{a_{1i}}$ and $K_{2i} = \frac{d_{2i}+k_{2i}}{a_{2i}}$ are the Michaelis-Menten binding constants for transcription and translation, respectively. Next, we solve for the free resources $P$ and $R$, and the free DNA $D_i$ by substituting Eqs. (S9) into Eqs. (S6), (S7), and (S8), which gives

$$D_i = \frac{D_{i_{tot}}}{1 + \frac{PF_i(u_i)}{K_{1i}}} \tag{S10a}$$

$$P = \frac{P_{tot}}{1 + \sum_{j=1}^{n} \frac{D_j F_j(u_j)}{K_{1j}}} \tag{S10b}$$

$$R = \frac{R_{tot}}{1 + \sum_{j=1}^{n} \frac{m_j}{K_{2j}}}. \tag{S10c}$$

To simplify analysis, we assume that binding between the RNAP and every gene is weak, $P \ll K_{1i}$, and binding between the ribosome and mRNA is also weak, $R \ll K_{2i}$, as in (*1*, *2*). Then, applying this assumption to Eqs. (S10) and substituting Eqs. (S10) into Eqs. (S9), the QSS concentration of the complexes are given as

$$\overline{C}_{1i} = \frac{D_{i_{tot}} P_{tot} F_i(u_i)}{K_{1i}\left(1 + \sum_{j=1}^{n} \frac{D_{i_{tot}} F_i(u_i)}{K_{1i}}\right)} \tag{S11a}$$

$$\overline{C}_{2i} = \frac{k_{1i} D_{i_{tot}} P_{tot} R_{tot} F_i(u_i)}{K_{1i} K_{2i} \delta \left(1 + \sum_{j=1}^{n} \frac{D_{j_{tot}} F_j(u_j)}{K_{1j}}\right)\left(1 + \sum_{j=1}^{n} \frac{k_{1j}}{K_{2j}\delta} \overline{C}_{1j}\right)}. \tag{S11b}$$

After substituting Eq. (S11a) into Eq. (S11b), simplifying, and substituting into Eq. (S5), we find the dynamics of the protein $\mathrm{x}_i$ to be

$$\dot{x}_i = \underbrace{\frac{\alpha_i F_i(u_i)}{1 + \sum_{j=1}^{n} w_j F_j(u_j)}}_{(a)} - \gamma x_i, \tag{S12}$$

where $\alpha_i = \frac{k_{1i}k_{2i}P_{tot}R_{tot}D_{i_{tot}}}{K_{1i}K_{2i}\delta}$ represents the maximum rate of production and $w_j = \frac{D_{j_{tot}}}{K_{1j}}\left(1 + \frac{k_{1j}P_{tot}}{K_{2j}\delta}\right)$ represents the resource demand coefficient of gene $j$ and depends on the DNA copy number, $D_{j_{tot}}$, the promoter strength, determined by $K_{1j}$, the RBS strength, determined by $K_{2j}$, the total RNAP concentration, $P_{tot}$, and the transcription rate, $k_{1j}$. In genetic circuits, the regulation function $F_i(u_i)$ typically has a sigmoidal function such as a Hill function (*2–4*). Then, the resource competition model in the main text, Eq. (1), is found after partitioning the resource competition term in the denominator of term $(a)$ in Eq. (S12) into genes inside and outside a module of interest. In the following analysis and in the main text, we study the production rate of the protein $\mathrm{x}_i$, given by term $(a)$ in Eq. (S12).

#### S1.2 Production Rate

Here we derive how the protein production rate can be found from experimental measurements based on a logistic growth model and the protein concentration model. We consider a control volume consisting of all cells and media in a well. The number of cells $n$ follows the logistic growth model

$$\dot{n} = \mu n \underbrace{\left(1 - \frac{n}{K}\right)}_{(a)}, \tag{S13}$$

where $\mu$ is the specific growth rate and $K$ is the population carrying capacity. The dynamics governing the total protein concentration of a protein F in a microplate well based on the previously derived model, Eq. (S12), is given as

$$\dot{F} = \beta n \tag{S14}$$

where $\beta$ represents the protein production rate per cell, i.e. term $(a)$ in Eq. (S12). It is assumed that protein F is not actively degraded within the cells, which is a good assumption for fluorescent proteins since they are stable and not targeted by proteases (*5*). It is assumed that cells are only measured in exponential phase, i.e. the number of cells is much lower than the carrying capacity, $n \ll K$, so the term $(a)$ in Eq. (S13) is negligible. Then, the protein production rate $\beta$, which is affected by resource competition and is of interest in this paper, can be found experimentally from the quantity

$$\beta = \frac{1}{n}\frac{dF}{dt}. \tag{S15}$$

In the experiments, the quantity $y = \frac{1}{\mathrm{OD}_{600}}\frac{dF}{dt}$ is used for the protein production rate per cell since it is proportional to $\beta$, where $F$ is the absolute fluorescence from each channel after subtracting the background and correcting for scattering effects (see Methods, Section 4.3). For details on the computation of the numerical derivative in Eq. (S15) from data, see Methods Section 4.4. The protein production rate of the model term $(a)$ in Eq. (S12) is used as the starting point for the derivation of the expressions for estimating the resource competition characteristics $Q_{\mathrm{M}}$ and $S_{\mathrm{M}}$. For data corresponding to the production rate per cell for all constructs in this paper, see SI, Section S5.3.

Note that the dynamics of protein per cell $\frac{F}{n}$ follow the standard protein production model in genetic circuits

$$\frac{d}{dt}\left(\frac{F}{n}\right) = \beta - \mu\left(\frac{F}{n}\right), \tag{S16}$$

which can be found by applying the time derivative to $\frac{F}{n}$ and substituting Eq. (S13) and Eq. (S14). Note that the steady state of the protein concentration $\frac{F}{n}$ in Eq. (S16) is given as $\frac{\beta}{\mu}$, which is proportional to the production rate. When the growth rate (equivalent to the dilution rate) is equal for all experimental conditions, then the steady state production rate and steady state concentration are equivalent. However, the growth rate does change across different experiments depending on the resource load (Figures S6 and S7), so the steady state concentration, which depends on the dilution rate, is not useful for estimating the resource competition characteristics. Therefore, the protein production rate is used in the main text for all module outputs and analysis.

#### S1.3 Derivation of Predictions Using Resource Demand and Sensitivity

We now consider the situation where a module of interest M is perturbed by a module with known resource demand $Q_{\mathrm{P}}$. We will derive the estimate for the output of Module M in this new context. We assume all measurements are made at steady state. First, we compare the expressions for the perturbed and unperturbed outputs of the module based on the production rate of the resource competition model, Eq. (S12). Let the output of a module $y_{\mathrm{M}}$ be equal to the production rate (term $(a)$) at steady state in the resource competition model Eq. (S12), which is found experimentally using the expression for the production rate e.g. $\beta$ from Eq. (S15).

$$y_{\mathrm{M}} = \frac{\alpha_{\mathrm{M}} F_y(u_y)}{1 + Q_{\mathrm{M}}} \tag{S17}$$

$$y_{\mathrm{M}}^p = \frac{\alpha_{\mathrm{M}} F_y(u_y^p)}{1 + Q_{\mathrm{M}} + Q_{\mathrm{P}}}. \tag{S18}$$

Here, $\alpha_{\mathrm{M}}$ represents the maximum rate of protein production, $F_y(\cdot)$ is the regulation function, and $Q_{\mathrm{M}}$ and $Q_{\mathrm{P}}$ are the resource demands for modules M and P, according to Eq. (S12). We use the protein production rate, so the dilution term in the model describing protein concentration (Eq. S12) does not appear, as explained in Section S1.2. Next, we use the Taylor series expansion for the regulation function $F_y(u_y)$ under the resource perturbation $Q_{\mathrm{P}}$, given as

$$F_y(u_y^p) = F_y(u_y) + \underbrace{\frac{dF_y(u_y)}{dQ_P}}_{F_y(u_y)S_{\mathrm{M}}} Q_{\mathrm{P}} + \mathcal{O}\left(\frac{d^2F_y(u_y)}{dQ_P^2} Q_{\mathrm{P}}^2\right). \tag{S19}$$

By the definition of $S_{\mathrm{M}}$ in Eq. (3) in the main text, $\frac{dF_y(u_y)}{dQ_P} = F_y(u_y)S_{\mathrm{M}}$. We assume that the higher order terms in Eq. (S19) are insignificant, which is a good assumption when the resource perturbation is small or the cooperativity of the regulation function $F_y(u_y)$ is not large. We can then express the perturbed output $y_{\mathrm{M}}^p$ in terms of the isolated output $y_{\mathrm{M}}$,

$Q_{\mathrm{M}}$, $Q_{\mathrm{P}}$ and $S_{\mathrm{M}}$ by substituting and rearranging Eq. (S17), Eq. (S18), and Eq. (S19). This results in the equation for the prediction $\hat{y}^p_{\mathrm{M}}$ as

$$\hat{y}^p_{\mathrm{M}} = y_{\mathrm{M}} \left(1 + \hat{S}_{\mathrm{M}}\hat{Q}_{\mathrm{P}}\right) \frac{1 + \hat{Q}_{\mathrm{M}}}{1 + \hat{Q}_{\mathrm{M}} + \hat{Q}_{\mathrm{P}}}, \tag{S20}$$

where we also substituted the estimators $\hat{Q}_{\mathrm{M}}$, $\hat{Q}_{\mathrm{P}}$, and $\hat{S}_{\mathrm{M}}$ for the true parameters $Q_{\mathrm{M}}$, $Q_{\mathrm{P}}$ and $S_{\mathrm{M}}$, respectively. Additionally, it should be noted that the formulas for the estimates for $\hat{Q}_{\mathrm{M}}$, $\hat{S}_{\mathrm{M}}$, and $\hat{y}^p_{\mathrm{M}}$ are exact for constitutive modules since $F_y(u_y)$ is a constant, giving $S_{\mathrm{M}} = 0$, and therefore no Taylor series approximations are necessary.

#### S1.4 Derivation of Resource Demand Estimation for a Resource Sensor

According to the model of a circuit with resource competition Eq. (1) for a constitutive gene using the protein production rate, the outputs of the two resource sensors when measured without other modules are given by

$$y_{\mathrm{RS1}} = \frac{\alpha_{\mathrm{RS1}}}{1 + w_{\mathrm{RS1}}} \tag{S21a}$$

$$y_{\mathrm{RS2}} = \frac{\alpha_{\mathrm{RS2}}}{1 + w_{\mathrm{RS2}}} \tag{S21b}$$

where $y_{\mathrm{RS1}}$ and $y_{\mathrm{RS2}}$ are the measured outputs of resource sensors 1 and 2, respectively. Here, $\alpha_{RS1}$ and $\alpha_{RS2}$ are the maximum production rates of $y_{\mathrm{RS1}}$ and $y_{\mathrm{RS2}}$, with $w_{\mathrm{RS1}}$ and $w_{\mathrm{RS2}}$ representing the corresponding resource demand parameters of the single genes. The resource demand parameters of resource sensors 1 and 2 are $Q_{\mathrm{RS1}} = w_{\mathrm{RS1}}$ and $Q_{\mathrm{RS2}} = w_{\mathrm{RS2}}$. By measuring the two resource sensors in the same cell, their outputs are given as

$$y^p_{\mathrm{RS1}} = \frac{\alpha_{\mathrm{RS1}}}{1 + w_{\mathrm{RS1}} + w_{\mathrm{RS2}}} \tag{S22a}$$

$$y^p_{\mathrm{RS2}} = \frac{\alpha_{\mathrm{RS2}}}{1 + w_{\mathrm{RS1}} + w_{\mathrm{RS2}}}, \tag{S22b}$$

where $y^p_{\mathrm{RS1}}$ and $y^p_{\mathrm{RS2}}$ are the outputs of resource sensors 1 and 2, respectively, when they are competing for resources. Using Eq. (S21) and Eq. (S22) and solving for the resource demand parameters for both resource sensors and simplifying, we find

$$\hat{Q}_{\mathrm{RS1}} = \frac{y_{\mathrm{RS2}}(y_{\mathrm{RS1}} - y^p_{\mathrm{RS1}})}{y^p_{\mathrm{RS1}} y_{\mathrm{RS2}} + y_{\mathrm{RS1}} y^p_{\mathrm{RS2}} - y_{\mathrm{RS1}} y_{\mathrm{RS2}}} \tag{S23a}$$

$$\hat{Q}_{\mathrm{RS2}} = \frac{y_{\mathrm{RS1}}(y_{\mathrm{RS2}} - y^p_{\mathrm{RS2}})}{y^p_{\mathrm{RS1}} y_{\mathrm{RS2}} + y_{\mathrm{RS1}} y^p_{\mathrm{RS2}} - y_{\mathrm{RS1}} y_{\mathrm{RS2}}}, \tag{S23b}$$

which can be rearranged to give the expressions in Eq. (5) in the main text and in Figure 2a.

#### S1.5 Derivation of Resource Demand and Sensitivity for a Module

We now consider a genetic circuit module M with a fluorescent protein output. The fluorescent protein production rate output according to the general model Eq. (1) is given as

$$y_{\mathrm{M}} = \frac{\alpha_{\mathrm{M}} F_y(u_y)}{1 + \sum\limits_{j \in \mathcal{M}} w_j F_j(u_j)}, \tag{S24}$$

where $F_y(u_y)$ is the regulation function of the output protein of the module, $\alpha_{\mathrm{M}}$ is the maximum production rate, $\mathcal{M}$ is the set of genes in the module, and $w_j F_j(u_j)$ represents the resource demand of the $j$th gene in the module. Next, module and resource sensor are measured together in the same cell. The outputs of the module and resource sensor are given as

$$y^p_{\mathrm{RS}} = \frac{\alpha_{\mathrm{RS}}}{1 + \sum\limits_{j \in \mathcal{M}} w_j F_j(u^p_j) + Q_{\mathrm{RS}}} \tag{S25a}$$

$$y^p_{\mathrm{M}} = \frac{\alpha_{\mathrm{M}} F_y(u^p_y)}{1 + \sum\limits_{j \in \mathcal{M}} w_j F_j(u^p_j) + Q_{\mathrm{RS}}}, \tag{S25b}$$

where $F_y(u^p_y)$ and $F_j(u^p_j)$ represent the perturbed regulation functions of the output gene and the $j$th gene, respectively, which may be slightly different than the unperturbed regulation functions due to changes in other genes that regulate the module output, $u_j$.

We assume that change in the module's resource demand due to the resource sensor perturbation is much smaller than the module's resource demand itself, specifically $\sum_{j\in\mathcal{M}} w_j \frac{dF_j(u_j)}{dQ_{\mathrm{RS}}} Q_{\mathrm{RS}} \ll \sum_{j\in\mathcal{M}} w_j F_j(u_j)$, which is valid since all $w_j$ and $Q_{\mathrm{RS}}$ are smaller than 1 and the sensitivity $S_{\mathrm{M}}$ is less than 1 for all modules measured (Figures 3 and 5 in the main text). Then, by applying a Taylor series expansion of the perturbed resource demand under the perturbation $Q_{\mathrm{RS}}$, we can express the perturbed resource demand as

$$\sum_{j\in\mathcal{M}} w_j F_j(u_j^p) = \sum_{j\in\mathcal{M}} w_j F_j(u_j) + \mathcal{O}\left(\sum_{j\in\mathcal{M}} w_j \frac{dF_j(u_j)}{dQ_{\mathrm{RS}}} Q_{\mathrm{RS}}\right). \tag{S26}$$

Then, the unperturbed resource demand is approximately equal to the perturbed resource demand $\sum_{j\in\mathcal{M}} w_j F_j(u_j) \approx \sum_{j\in\mathcal{M}} w_j F_j(u_j^p)$ when $\frac{dF_j(u_j)}{dQ_{\mathrm{RS}}} Q_{\mathrm{RS}}$ is small corresponding to when the resource sensor $Q_{\mathrm{RS}}$ is small or the cooperativity of all regulation functions $F_j(u_j)$ inside the module $\mathcal{M}$ are not large.

Then, combining Eq. (S24) with Eq. (S25) along with the resource sensor measurements alone Eq. (S21), the module's resource demand parameter $Q_{\mathrm{M}}$ is solved for as

$$\hat{Q}_{\mathrm{M}} = \left(\frac{y_{\mathrm{RS}}}{y_{\mathrm{RS}}^p} - 1\right)(1 + \hat{Q}_{\mathrm{RS}}). \tag{S27}$$

The module's sensitivity $S_{\mathrm{M}}$ can be approximated using a Taylor series for the perturbed regulation function $F_y(u_y)$,

$$F_y(u_y^p) = F_y(u_y) + \underbrace{\frac{dF_y(u_y)}{dQ_P}}_{F_y(u_y)S_{\mathrm{M}}} Q_{\mathrm{P}} + \mathcal{O}\left(\frac{d^2F_y(u_y)}{dQ_P^2} Q_{\mathrm{P}}^2\right). \tag{S28}$$

We neglect the quadratic term in Eq. (S28), which is a good approximation when resource demand $Q_{\mathrm{P}}$ is small or the cooperativity of the regulation function $F_y(u_y)$ is not large. Then, rearranging Eq. (S28) and combining with the definition for $S_{\mathrm{M}}$ in Eq. (3), gives the estimator

$$\hat{S}_{\mathrm{M}} = \frac{1}{\hat{Q}_{\mathrm{RS}}} \left( \frac{F_y(u_y^p)}{F_y(u_y)} - 1 \right). \tag{S29}$$

The term $\frac{F_y(u_y)^p}{F_y(u_y)}$ may be solved for by combining Eq. (S24) with Eq. (S25) and Eq. (S21) to find

$$\frac{F_y(u_y^p)}{F_y(u_y)} = \frac{y_{\mathrm{M}}^p y_{\mathrm{RS}}(1+\hat{Q}_{\mathrm{RS}})}{y_{\mathrm{M}}(y_{\mathrm{RS}} + \hat{Q}_{\mathrm{RS}}(y_{\mathrm{RS}} - y_{\mathrm{RS}}^p))}. \tag{S30}$$

Then, substituting Eq. (S30) into Eq. (S29) and simplifying, the estimator for $S_{\mathrm{M}}$ in terms of the measured values is

$$\hat{S}_{\mathrm{M}} = \frac{\left(1 - \frac{y_{\mathrm{M}}}{y_{\mathrm{M}}^p}\right)(1+\hat{Q}_{\mathrm{RS}}) + \hat{Q}_{\mathrm{RS}} \frac{y_{\mathrm{RS}}^p}{y_{\mathrm{RS}}}}{\hat{Q}_{\mathrm{RS}}(1 + \hat{Q}_{\mathrm{RS}}(1 - \frac{y_{\mathrm{RS}}^p}{y_{\mathrm{RS}}}))}, \tag{S31}$$

which matches the equations given in the main text in Eq. (6).

### S2 Growth Rate and Resource Demand

We wish to determine the effect of resource loading on cellular growth rate. The resource demand parameter $Q_{\mathrm{M}}$ for a module M is related to the availability of free cellular resources (*6*). The relationship between ribosome availability and bacterial growth rate can be modeled as a Hill function (*6*), with the form

$$\mu(R_{free}) = \frac{\beta R_{free}}{K + R_{free}} + \beta_0 \tag{S32}$$

where $\mu$ represents the specific growth rate, $R_{free}$ is the concentration of available cellular resources, and $\beta$, $\beta_0$, and $K$ are constants. Using the model of protein production with resource competition, Eq. (1), the available resources $R_{free}$ is related to the resource demand

$Q_M$ as

$$R_{free} = \frac{R_{tot}}{1 + Q_M}, \tag{S33}$$

where $R_{tot}$ represents the total cellular resources. Therefore, the dependence of cellular growth rate on the resource demand $Q_M$ can be found by combining Eq. (S32) and Eq. (S33) and simplifying, which gives

$$\mu(Q_M) = \frac{\beta}{\frac{K}{R_{tot}}(1 + Q_M) + 1} + \beta_0. \tag{S34}$$

This model was fit to the experimentally estimated $\hat{Q}_M$ and measured growth rates for each previously run experiment as shown in Figure S1. The fitted parameters in Figure S1 are $\beta = 0.86 \pm 0.08$, $\beta_0 = 0.$, and $\frac{K}{R_{tot}} = 0.53 \pm 0.12$. This assumes that the genetic circuit does not cause any toxicity effects beyond depleting the pool of cellular resources.

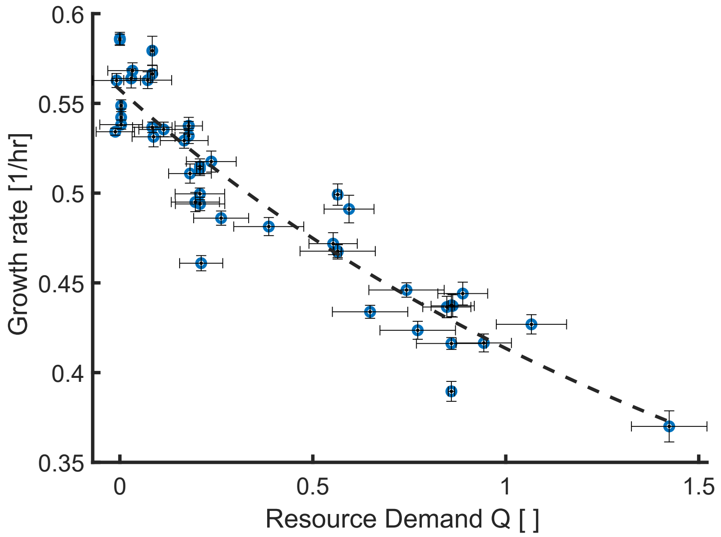


Figure S1: Relation between measured specific growth rate and resource demand for all experiments. The black dashed line represents the least-squares fit of the model Eq. (S34) to the experimentally measured specific growth rate and estimated resource demand $\hat{Q}_M$. The fit has an adjusted $R^2 = 0.864$.

### S3 Additional Data for Measurements of Resource Sensors and Modules

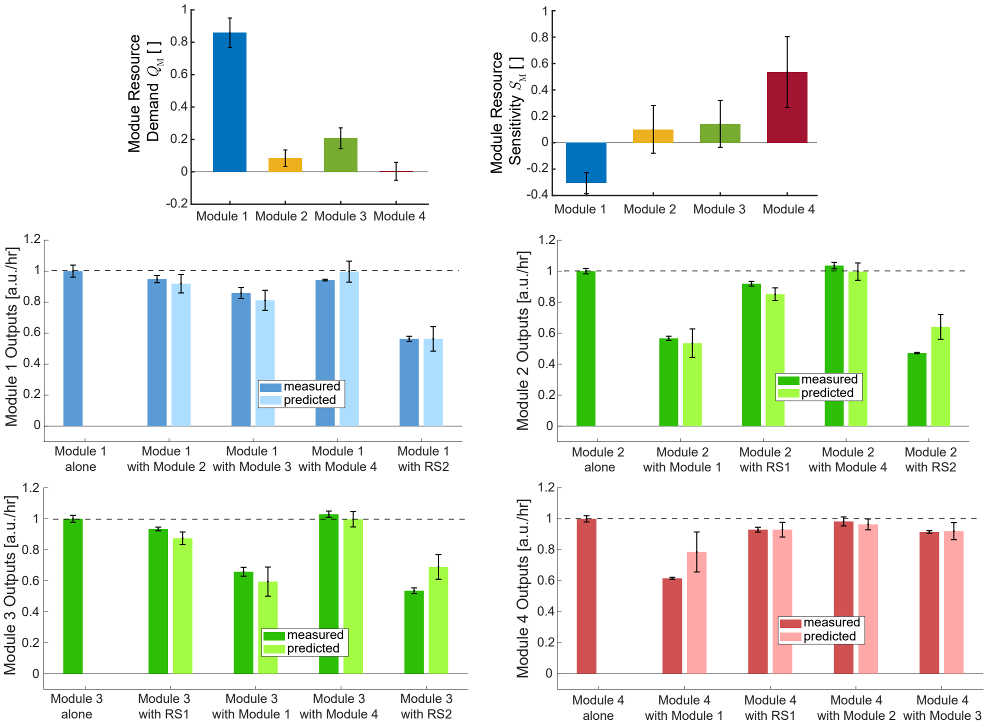


Figure S2: Resource demand and sensitivity for the constitutive modules when the sensitivity $S_{\mathrm{M}}$ is allowed to be nonzero, which may capture unmodeled effects in the regulation function of the gene $F_y(u_y)$ in Eq. (3) in the main text. Predictions follow according to the same expression for prediction with $Q$ and $S$ Eq. (7) in the main text. Allowing $S_{\mathrm{M}}$ to be non-zero for the constitutive modules enables better predictions than in the main text where $S_{\mathrm{M}} = 0$ for constitutive modules.

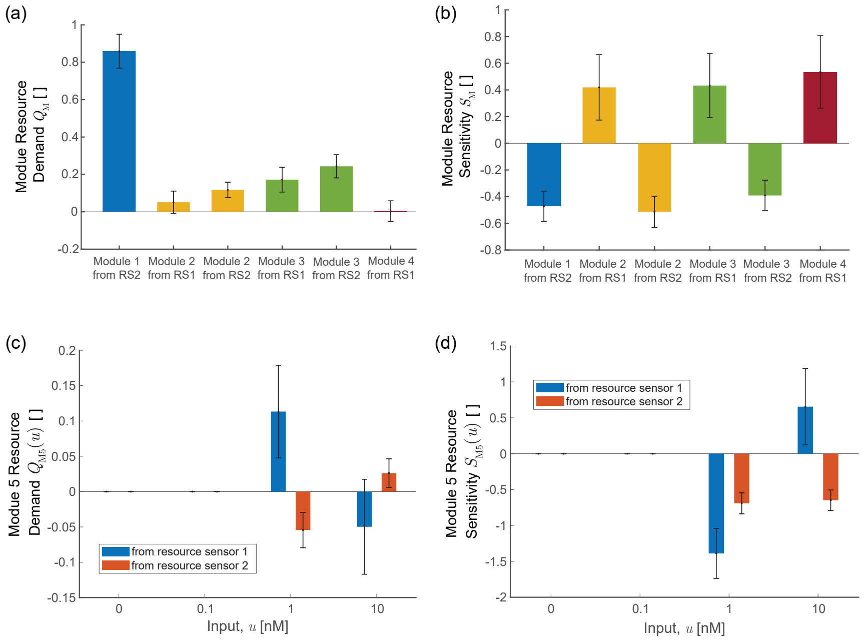


Figure S3: **(a)** Estimated module resource demands for modules 1-4 (constitutive) calculated using measurements of the modules with either resource sensor 1 (RS1) or with resource sensor 2 (RS2). Resource demands in Figures 3 and 4 are the average of the resource demands measured from both resource sensors for modules 2 and 3 with appropriate error bar propagation. **(b)** Estimated module resource sensitivities for modules 1-4 (constitutive) calculated using measurements of the modules with either resource sensor 1 (RS1) or with resource sensor 2 (RS2). Resource sensitivities in Figures 3 and 4 are the average of the resource sensitivities measured from both resource sensors for modules 2 and 3 with appropriate error bar propagation. **(c)** Estimated module resource demands for module 5 (inducible) calculated using measurements of the module with either resource sensor 1 (RS1) or with resource sensor 2 (RS2). Resource demands in Figure 5 are the average of the resource demands for module 5 measured from both resource sensors with appropriate error bar propagation. **(d)** Estimated module resource sensitivities for module 5 (inducible) calculated using measurements of the module with either resource sensor 1 (RS1) or with resource sensor 2 (RS2). Resource demands in Figure 5 are the average of the resource sensitivities for module 5 measured from both resource sensors with appropriate error bar propagation.

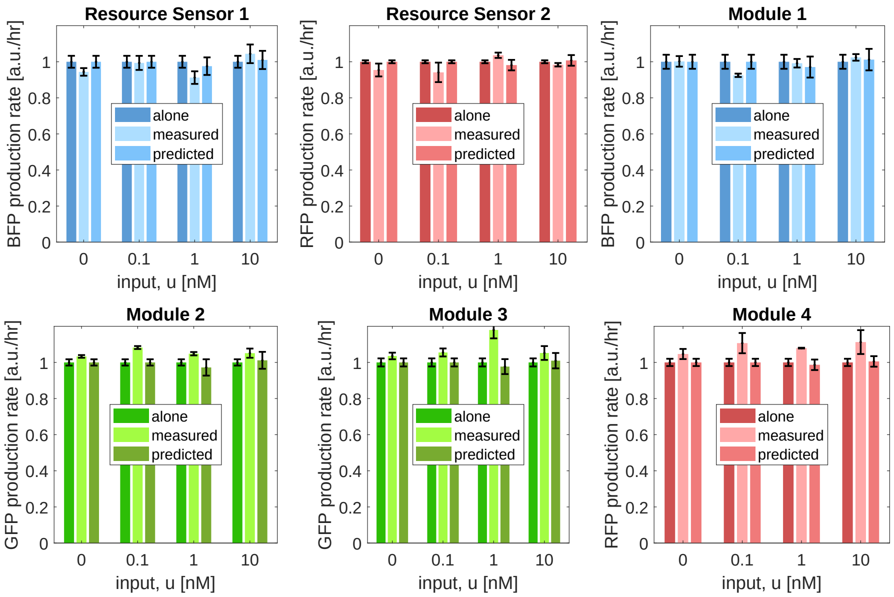


Figure S4: Outputs of Resource Sensors 1 and 2 and Modules 1, 2, 3, and 4 when measured with Module 5. Predictions for outputs are made using $\hat{Q}_{\mathrm{M}}$ and $\hat{S}_{\mathrm{M}}$ for each module in Eq. (7).

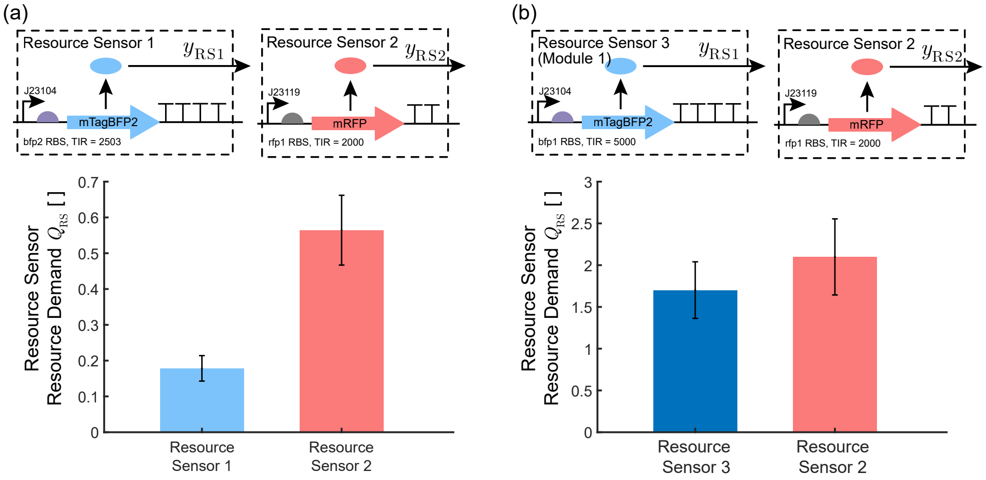


Figure S5: The estimated value of the resource demand for the resource sensors may change due to overloading of the cell. **(a)** Measurement of the resource demand of resource sensors 1 and 2 as in the main text. **(b)** Measurement of the resource demand according to the resource sensor equations in Figure 2 (Eq. (5)) using module 1 from the main text as resource sensor 3. The resource demand estimate for resource sensor 2 is significantly larger than in panel (a) due to the regulation of cellular resources and stress response as discussed in the Discussion Section 3.

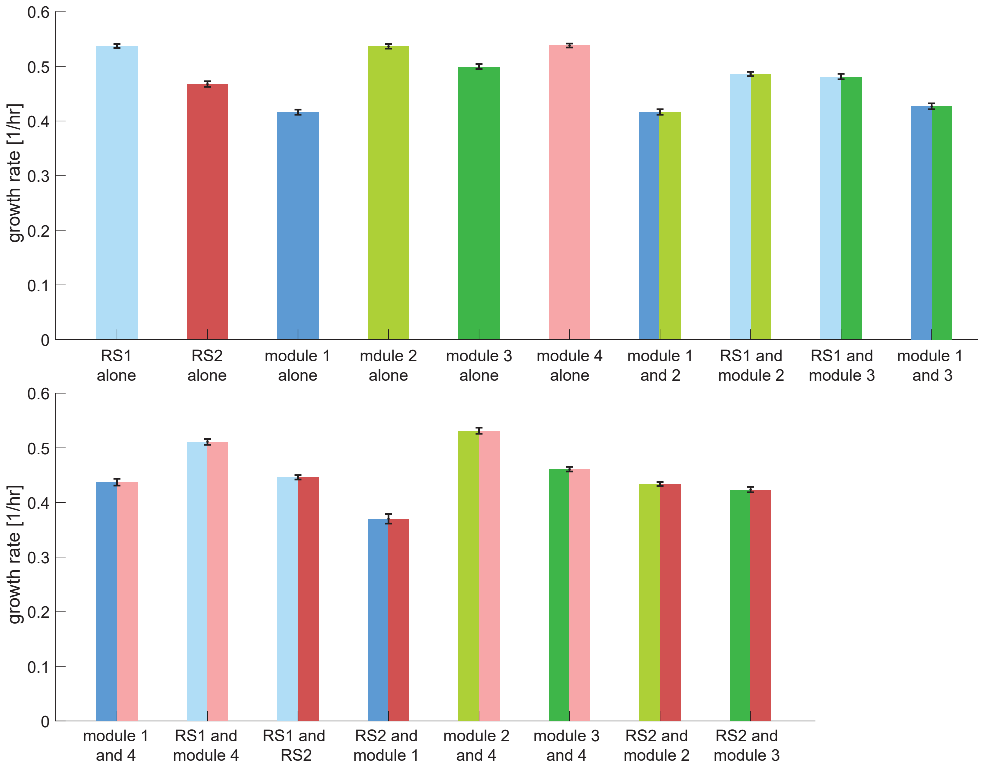


Figure S6: Growth rate of constitutive constructs.

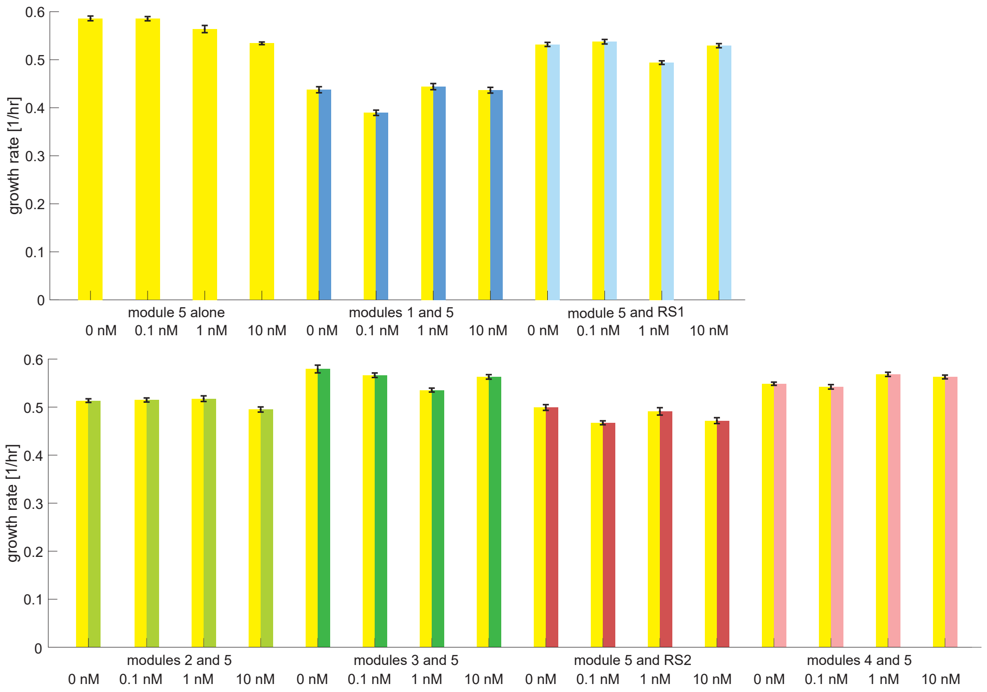


Figure S7: Growth rate of inducible constructs for different levels of induction.

#### S3.1 Verification of Fluorescence parts

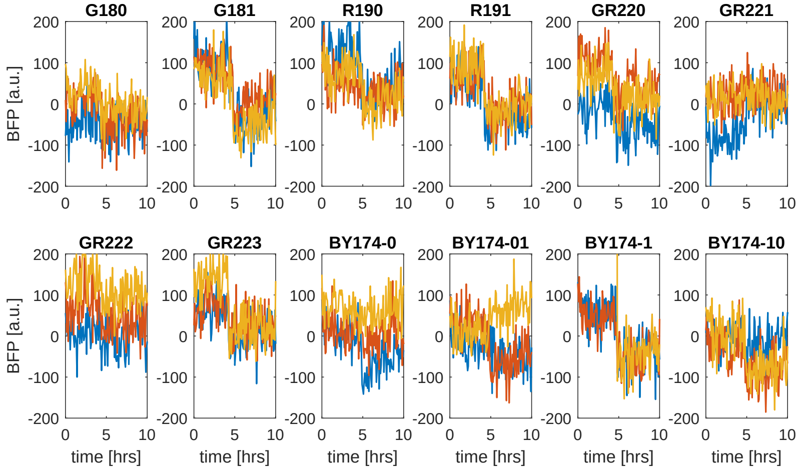


Figure S8: Blue fluorescence readings after background subtraction for constructs that contain a mTagBFP gene with the bfp3 RBS (TIR = 0.9). Blue fluorescence readings are not significantly larger than the measurement noise with this RBS. Compare with Figure S9 where the RBS for the blue fluorescent gene is stronger.

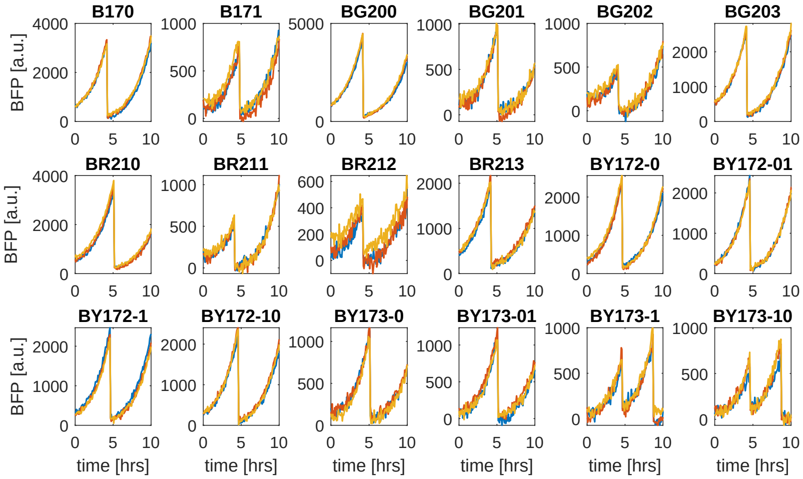


Figure S9: Blue fluorescence readings after background subtraction for constructs that contain a mTagBFP gene and have either the bfp1 (TIR = 5000) or the bfp2 (TIR = 2503) RBS. Blue fluorescence is significantly larger than the measurement noise with these RBSs. Compare with Figure S8.

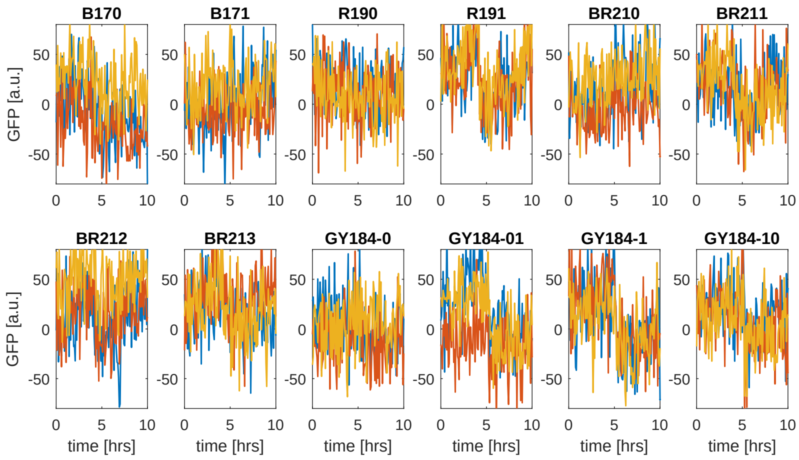


Figure S10: Green fluorescence readings after background subtraction for constructs that contain a GFPop1 gene with the gfp3 RBS (TIR = 1.1). No significant GFPop1 fluorescence is observed. Compare with Figure S11 where the RBS for the green fluorescent gene is stronger.

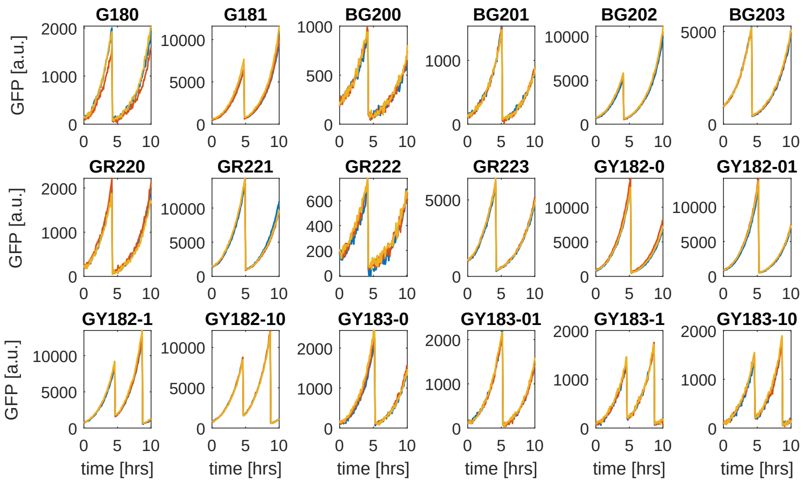


Figure S11: Green fluorescence readings after background subtraction for constructs that contain a GFPop1 gene and have either the gfp1 (TIR = 10211) or the gfp2 (TIR = 6298) RBS. Green fluorescence is significantly larger than the measurement noise with these RBSs. Compare with Figure S10.

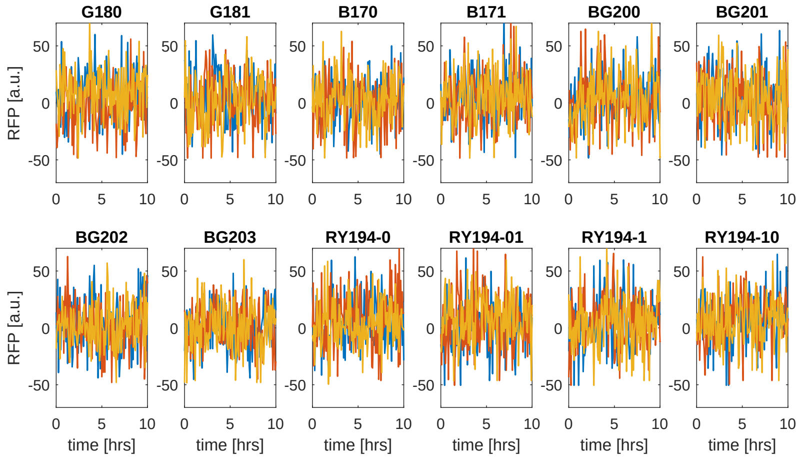


Figure S12: Red fluorescence readings after background subtraction for constructs that contain a mRFP gene with the rfp3 RBS (TIR = 1.0). No significant mRFP fluorescence is observed. Compare with Figure S13 where the RBS for the red fluorescent gene is stronger.

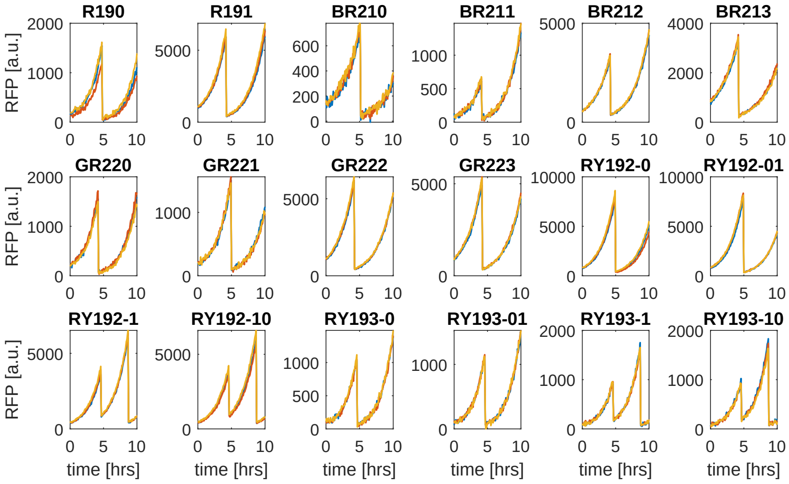


Figure S13: Red fluorescence readings after background subtraction for constructs that contain a mRFP gene and have either the rfp1 (TIR = 2000) or the rfp2 (TIR = 996) RBS. Red fluorescence is significantly larger than the measurement noise with these RBSs. Compare with Figure S12.

### S4 Plasmid Maps and DNA Sequences

Plasmid DNA maps for all constructs used are given in Figures S14, S15, S16, S17, and S18 with the sequences of the corresponding parts given in Tables S1, S2, S3, S4, S5, S6, S7, and S8. A p15a origin of replication and AmpR resistance selection marker were used for all constructs. For the sequences for all plasmids, see SI Table S10

Table S1: List of promoter sequences.

| **Promoter name** | **sequence** | **source** |
|---|---|---|
| J23104 | GGCGCGCCTTGACAGCTAGCTCAGTCCTAGGTATTGTGCTAGCTTACG | iGEM |
| J23100 | GGCGCGCCTTGACGGCTAGCTCAGTCCTAGGTACAGTGCTAGCTTAAT | iGEM |
| J23119 | ttgacagctagctcagtcctaggtataatgctagc | iGEM |
| PlacIQ | gcggcgcgccatcgaatggtgcaaaacctttcgcggtatggcatgatagcgccc | (*7*) |
| Pphlf | cgacgtacggtggaatctgattcgttaccaattgacatgatacgaaacgtaccgtatcgttaaggt | (*7*) |

Table S2: List of ribosome binding site sequences.

| **coding sequence** | **RBS name** | **TIR** | **RBS sequence** | **source** |
|---|---|---|---|---|
| mTagBFP | bfp1 | 5000 | tcaacccaccctaggagaggataag | RBS calculator (*8*) |
| | bfp2 | 2503 | CCGATTCTGCCGGGGGGAATTATA | RBS calculator (*8*) |
| | bfp3 | 0.9 | AGCGATGGGCTCACTGTTGAGCCTCA | RBS calculator (*8*) |
| GFPop1 (*9*) | gfp1 | 10211 | TTCTAATTAGGATAAGGACACCAAG | RBS calculator (*8*) |
| | gfp2 | 6298 | GCGAGAGGCTGGTTAAGTGGGTGTTG | RBS calculator (*8*) |
| | gfp3 | 1.1 | CTTGTCAAACAGTGGCTACGTGAACAG | RBS calculator (*8*) |
| mRFP | rfp1 | 2000 | CTAATTCTACATAGAGTGAGGTAATCTA | RBS calculator (*8*) |
| | B0034 | 942 | TCTAGAGAAAGAGGAGAAATACTAG | iGEM B0034 |
| | rfp3 | 1.0 | AAATATGTTGTAGAAACAATTGCACGA | RBS calculator (*8*) |
| EYFP | B0064 | | tactagagaaagaggggaaatactag | iGEM B0064 |
| PhlF | phl2 | | ggaagagagtcaattcatgggggtgaat | (*7*) |

Table S3: List of protein coding sequences.

| coding sequence | sequence | source |
|---|---|---|
| mTagBFP | atgagcgaactgatcaaagagaacatgcacatgaagctgtacatggagggtaccgtggataatcaccactttaagtgtacttctgagggcgagg<br>gtaagccgtatgaagggactcaaacgatgcgtattaaagtagtggagggtggcccactgccgtttgctttcgatattctggcgacgagctttct<br>gtatggtagcaaaacgtttataaaccacactcagggcattccggatttctttaaacaaagctttccggaaggttttacctgggagcgtgtgact<br>acgtatgaagatggtggtgtgttgactgctactcaagatacttcactgcaggacggctgtctgatctataacgtgaagattcgtggcgtgaact<br>ttacgagcaatgggccggtaatgcaaaaaaaaaccctgggttgggaagcgttcacggaaactctgtatccggctgacggcggcctggagggccg<br>taacgatatggcactgaagctggttggtggcagccacctgatcgcgaatatcaaaacgacttatcgctctaaaaaaccggcgaaaaatctgaag<br>atgccgggtgtttattatgttgactatcgtctggaacgcattaaagaagcgaataatgaaacttacgtggagcaacacgaggttgcagtggcgc<br>gctattgcgacttgccttcaaagctgggtcacaaactgaattaataa | (*10*) |
| GFPop1 | atgcgtaaaggagaagaactgttcactggagttgtcccaattcttgttgaattagatggtgatgttaatgggcacaaattttctgtgagtggag<br>agggtgaaggtgatgcgacatacggcaaactgacccttaaatttatttgtactaccggcaagctacctgttccttggccgacacttgtcactac<br>tttcggttatggtgttcaatgttttgcgagatacccagatcatatgaaacagcatgactttttcaaaagtgccatgcccgaaggttatgtacag<br>gaaagaactatatttttcaaagatgacgggaactacaaaacacgtgctgaagttaaatttgagggtgatactctggtaaatagaatcgaattaa<br>aaggtattgattttaaagaagatggcaacattcttggtcacaagctggaatacaactacaactcacacaatgtttacatcatggcagacaaaca<br>aaaaaatggcatcaaagttaacttcaaaattcgacacaacattgaagatggaagcgttcaactggcagaccattatcaacagaatactccgatt<br>ggcgatggccctgtactcttaccagacaaccattacctgtccacacaatctgccctttcgaaagatccgaacgaaaagagagatcacatggtcc<br>tgcttgagtttgtaacagctgctgggattacacatggtatggatgagctatacaaataataa | (*9*) |
| mRFP | atggcttcctccgaagacgttatcaaagagttcatgcgtttcaaagttcgtatggaaggttccgttaacggtcacgagttcgaaatcgaaggtg<br>aaggtgaaggtcgtccgtacgaaggtacccagaccgctaaactgaaagttaccaaaggtggtccgctgccgttcgcttgggacatcctgtcccc<br>gcagttccagtacggttccaaagcttacgttaaacacccggctgacatcccggactacctgaaactgtccttcccggaaggtttcaaatgggaa<br>cgtgttatgaacttcgaagacggtggtgttgttaccgttacccaggactcctccctgcaagacggtgagttcatctacaaagttaaactgcgtg<br>gtaccaacttcccgtccgacggtccggttatgcagaaaaaaaccatgggttgggaagcttccaccgaacgtatgtacccggaagacggtgctct<br>gaaaggtgaaatcaaaatgcgtctgaaactgaaagacggtggtcactacgacgctgaagttaaaaccacctacatggctaaaaaaccggttcag<br>ctgccgggtgcttacaaaaccgacatcaaactggacatcacctcccacaacgaagactacaccatcgttgaacagtacgaacgtgctgaaggtc<br>gtcactccaccggtgcttaataa | (*9*) |
| EYFP | atggtgagcaagggcgaggagctgttcaccggggtggtgcccatcctggtcgagctggacggcgacgtaaacggccacaagttcagcgtgtccg<br>gcgagggcgagggcgatgccacctacggcaagctgaccctgaagttcatctgcaccacaggcaagctgcccgtgccctggcccaccctcgtgac<br>caccttcggctacggcctgcaatgcttcgcccgctaccccgaccacatgaagctgcacgacttcttcaagtccgccatgcccgaaggctacgtc<br>caggagcgcaccatcttcttcaaggacgacggcaactacaagacccgcgccgaggtgaagttcgagggcgacaccctggtgaaccgcatcgagc<br>tgaagggcatcgacttcaaggaggacggcaacatcctggggcacaagctggagtacaactacaacagccacaacgtctatatcatggccgacaa<br>gcagaagaacggcatcaaggtgaacttcaagatccgccacaacatcgaggacggcagcgtgcagctcgccgaccactaccagcagaacaccccca<br>atcggcgacggccccgtgctgctgcccgacaaccactaccttagctaccagtccgccctgagcaaagaccccaacgagaagcgcgatcacatgg<br>tcctgctggagttcgtgaccgccgccgggatcactctcggcatggacgagctgtacaagtaa | (*7*) |
| PhlF | atggcacgtaccccgagccgtagcagcattggtagcctgcgtagtccgcatacccataaagcaattctgaccagcaccattgaaatcctgaaag<br>aatgtggttatagcggtctgagcattgaaagcgtggcacgtcgcgccggtgcaggcaaaccgaccatttatcgttggtggaccaacaaagcagc<br>actgattgccgaagtgtatgaaaatgaaatcgaacaggtacgtaaatttccggatttgggtagctttaaagccgatctggattttctgctgcat<br>aatctgtggaaagtttggcgtgaaaccatttgtggtgaagcatttcgttgtgttattgcagaagcacagttggaccctgtaaccctgacccaac<br>tgaaagatcagtttatggaacgtcgtcgtgagataccgaaaaaactggttgaagatgccattagcaatggtgaactgccgaaagatatcaatcg<br>tgaactgctgctggatatgatttttggtttttgttggtatcgcctgctgaccgaacagttgaccgttgaacaggatattgaagaatttaccttc<br>ctgctgattaatggtgtttgtccgggtacacagtgttga | (*7*) |

Table S4: RiboJ sequence (*7*).

| name | sequence |
|---|---|
| RiboJ | agctgtcaccggatgtgctttccggtctgatgagtccgtgaggacgaaacagcctctacaaataattttgtttaa |

Table S5: List of Terminator sequences (*11*).

| **Terminator** | **sequence** |
|---|---|
| ECK120033737 | ggaaacacagAAAAAAGCCCGCACCTGACAGTGCGGGCTTTTTTTTTcgaccaaagg |
| L3S1P22 | GACGAACAATAAGGCCGCAAATCGCGGCCTTcgTTATTGATAACAgcA |
| ECK120029600 | TTCAGCCAAAAAACTTAAGACCGCCGGTCTTGTCCACTACCTTGCAGTAATGCGGTGGACAGGATCGGCGGTTTTCTTTTCTCTTCTCAA |
| L3S3P21 | CCAATTATTGAAGGCCTCCCTAACGGGGGGCCTTTTTTTGTTTCTGGTCTCCC |
| terminator1 | ACAAAACTGCTGACGGCAGAAGCGTCTCGGTACCAAATTCCAGAAAAGAGACGCTTTCGAGCGTCTTTTTTCGTTTTGGTCCACGCTGATGAATATTCTACG<br>GAATTGCCATAGGCGTTGAACGCTACACGGACGATACGAATTTATGTATAGAGCG |
| Terminator_B0010 | ccaggcatcaaataaaacgaaaggctcagtcgaaagactgggcctttcgttttatctgttgtttgtcggtgaacgctctc |
| Terminator_B0012 | tcacactggctcaccttcgggtgggcctttctgcgtttata |
| terminator2 | tccttagcgaaagctaaggattttttttatctg |
| lambda tI terminator | ctgtaacagagcattagcgcaaggtgatttttgtcttcttgcgctaatttttt |

Table S6: Spacer sequences

| **Spacer** | **sequence** |
|---|---|
| spacer1 | TTCGCGACCCAGGATGAGGTCGCCAAAACTACTAGATTTG |
| spacer2 | GGaatCGTACGGTCGATCCATCTGACTATCGCCTACTTCA |
| spacer3 | gcttaacgatcgttggctg |

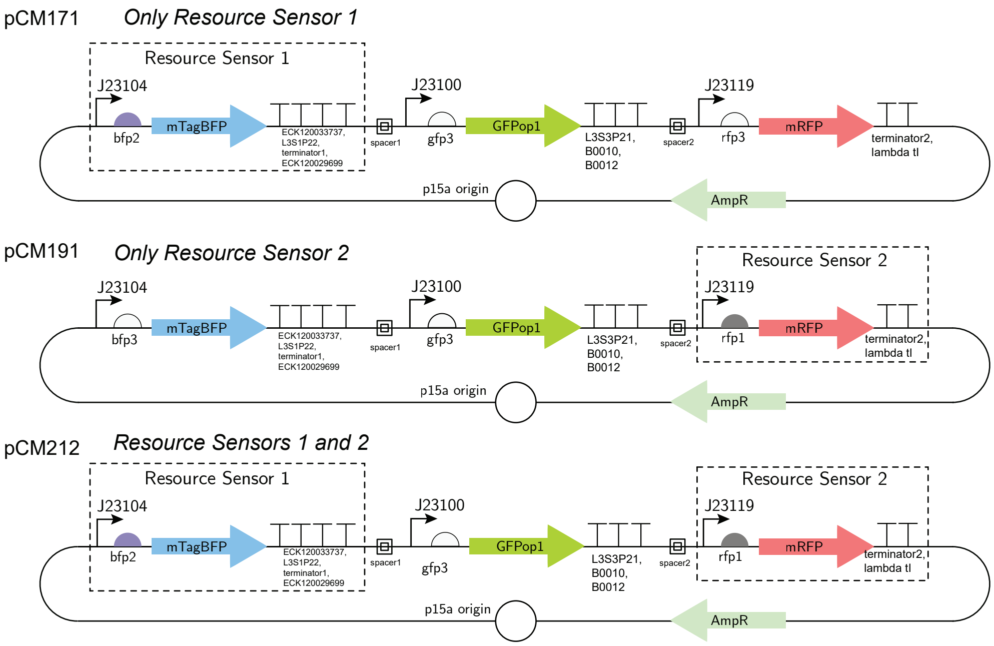


Figure S14: Resource sensors with each other.

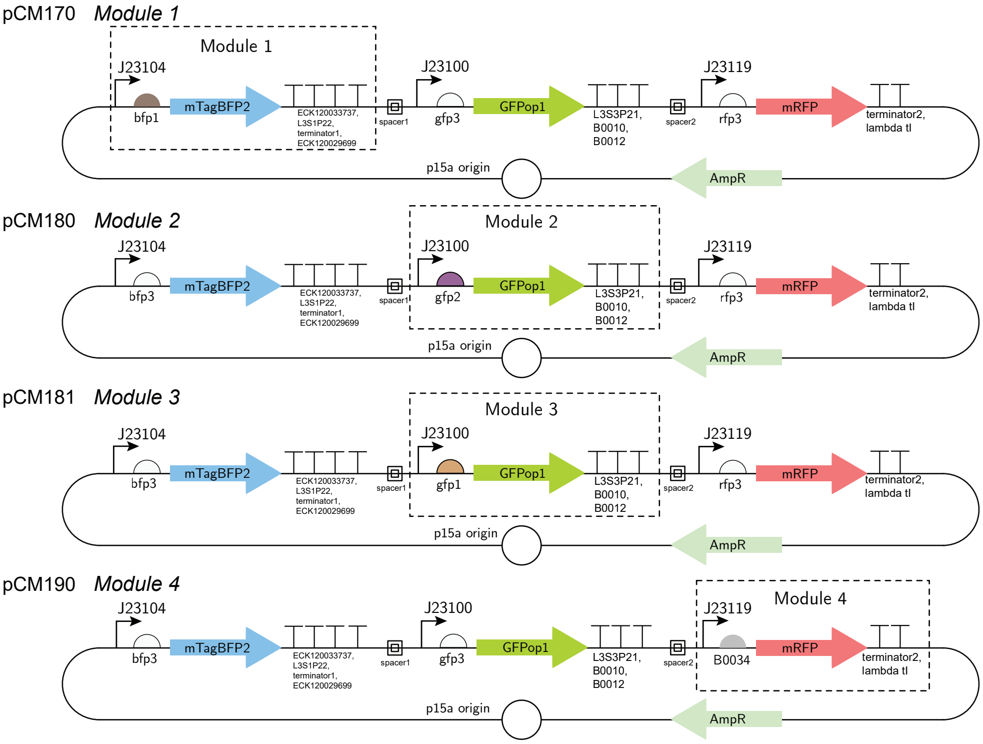


Figure S15: Plasmid maps of modules 1-4 alone.

Table S7: Description of ribosome binding site strengths for all constitutive constructs.

| **Plasmid** | **Description** | **BFP RBS** | **GFP RBS** | **RFP RBS** |
|---|---|---|---|---|
| pCM170 | Module 1 | bfp1 (TIR = 5000) | gfp3 (TIR = 1.1) | rfp3 (TIR = 1.0) |
| pCM171 | Resource sensor 1 | bfp2 (TIR = 2503) | gfp3 (TIR = 1.1) | rfp3 (TIR = 1.0) |
| pCM180 | Module 2 | bfp3 (TIR = 0.9) | gfp2 (TIR = 6298) | rfp3 (TIR = 1.0) |
| pCM181 | Module 3 | bfp3 (TIR = 0.9) | gfp1 (TIR = 10211) | rfp3 (TIR = 1.0) |
| pCM190 | Module 4 | bfp3 (TIR = 0.9) | gfp3 (TIR = 1.1) | rfp2 (TIR = 942) |
| pCM191 | Resource Sensor 2 | bfp3 (TIR = 0.9) | gfp3 (TIR = 1.1) | rfp1 (TIR = 2000) |
| pCM200 | Modules 1 and 2 | bfp1 (TIR = 5000) | gfp2 (TIR = 6298) | rfp3 (TIR = 1.0) |
| pCM201 | RS1 and Module 2 | bfp2 (TIR = 2503) | gfp2 (TIR = 6298) | rfp3 (TIR = 1.0) |
| pCM202 | RS1 and Module 3 | bfp2 (TIR = 2503) | gfp1 (TIR = 10211) | rfp3 (TIR = 1.0) |
| pCM203 | Modules 1 and 3 | bfp1 (TIR = 5000) | gfp1 (TIR = 10211) | rfp3 (TIR = 1.0) |
| pCM210 | Module 1 and 4 | bfp1 (TIR = 5000) | gfp3 (TIR = 1.1) | rfp2 (TIR = 942) |
| pCM211 | RS1 and Module 4 | bfp2 (TIR = 2503) | gfp3 (TIR = 1.1) | rfp2 (TIR = 942) |
| pCM212 | RS1 and RS2 | bfp2 (TIR = 2503) | gfp3 (TIR = 1.1) | rfp1 (TIR = 2000) |
| pCM213 | Module 1 and RS2 | bfp1 (TIR = 5000) | gfp3 (TIR = 1.1) | rfp1 (TIR = 2000) |
| pCM220 | Module 2 and 4 | bfp3 (TIR = 0.9) | gfp2 (TIR = 6298) | rfp2 (TIR = 942) |
| pCM221 | Module 3 and 4 | bfp3 (TIR = 0.9) | gfp1 (TIR = 10211) | rfp2 (TIR = 942) |
| pCM222 | Module 2 and RS2 | bfp3 (TIR = 0.9) | gfp2 (TIR = 6298) | rfp1 (TIR = 2000) |
| pCM223 | Module 3 and RS2 | bfp3 (TIR = 0.9) | gfp1 (TIR = 10211) | rfp1 (TIR = 2000) |

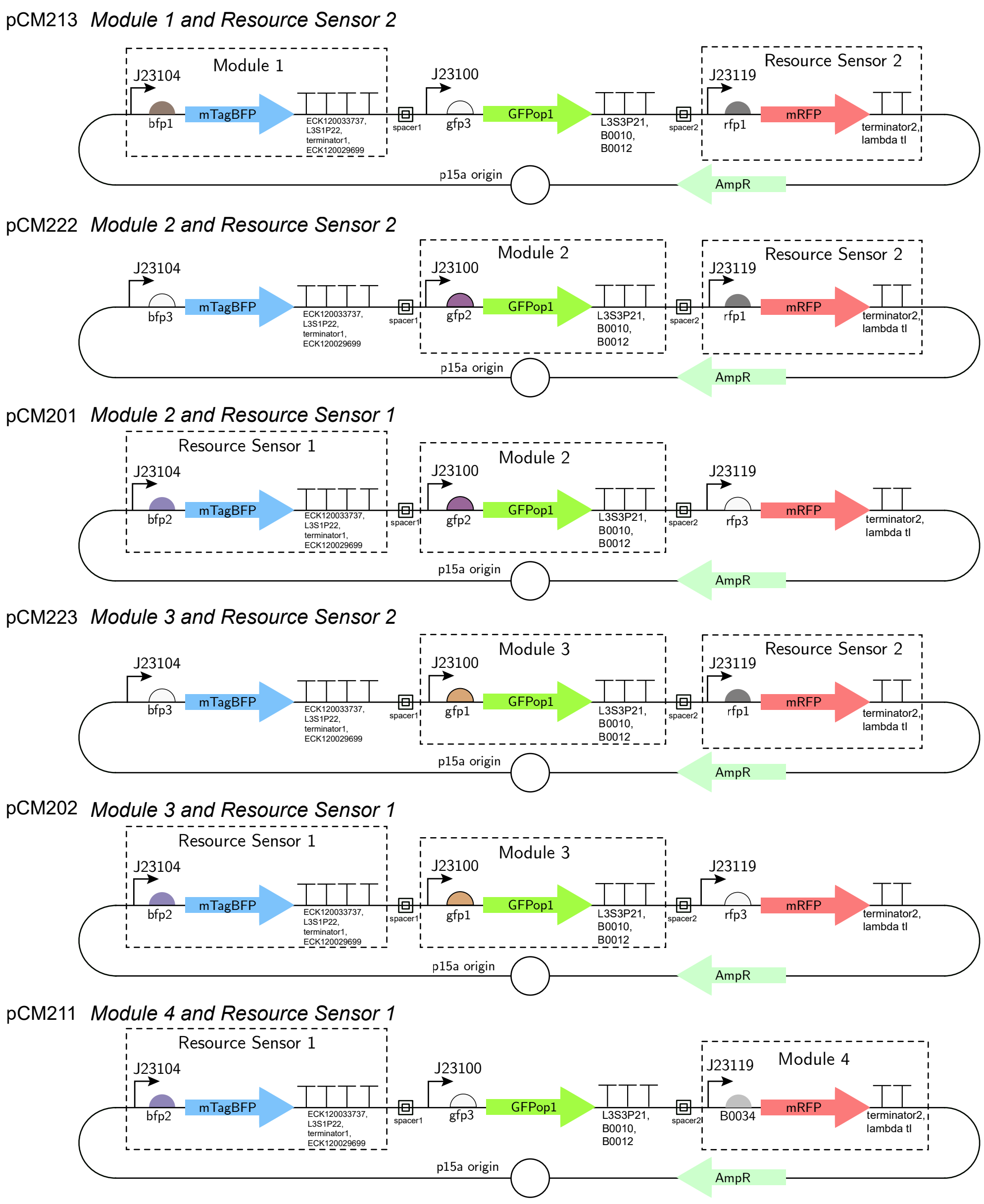


Figure S16: Plasmid maps of modules 1-4 with resource sensors.

Table S8: Description of parts for constitutive constructs (Modules 1-4 and resource sensors 1 and 2).

| Plasmid name | promoter | RBS | gene sequence | terminators | spacer |
|---|---|---|---|---|---|
| BFP module | J23104 | bfp1, bfp2, bfp3 | mTagBFP | ECK120033737, L3S1P22, terminator1, ECK120029600 | spacer1 |
| GFP module | J23100 | gfp1, gfp2, gfp3 | GFPop1 | L3S3P21, B0010, B0012 | spacer2 |
| RFP module | J23119 | rfp1, rfp2, rfp3 | mRFP | terminator2, lambda tI terminator | none |

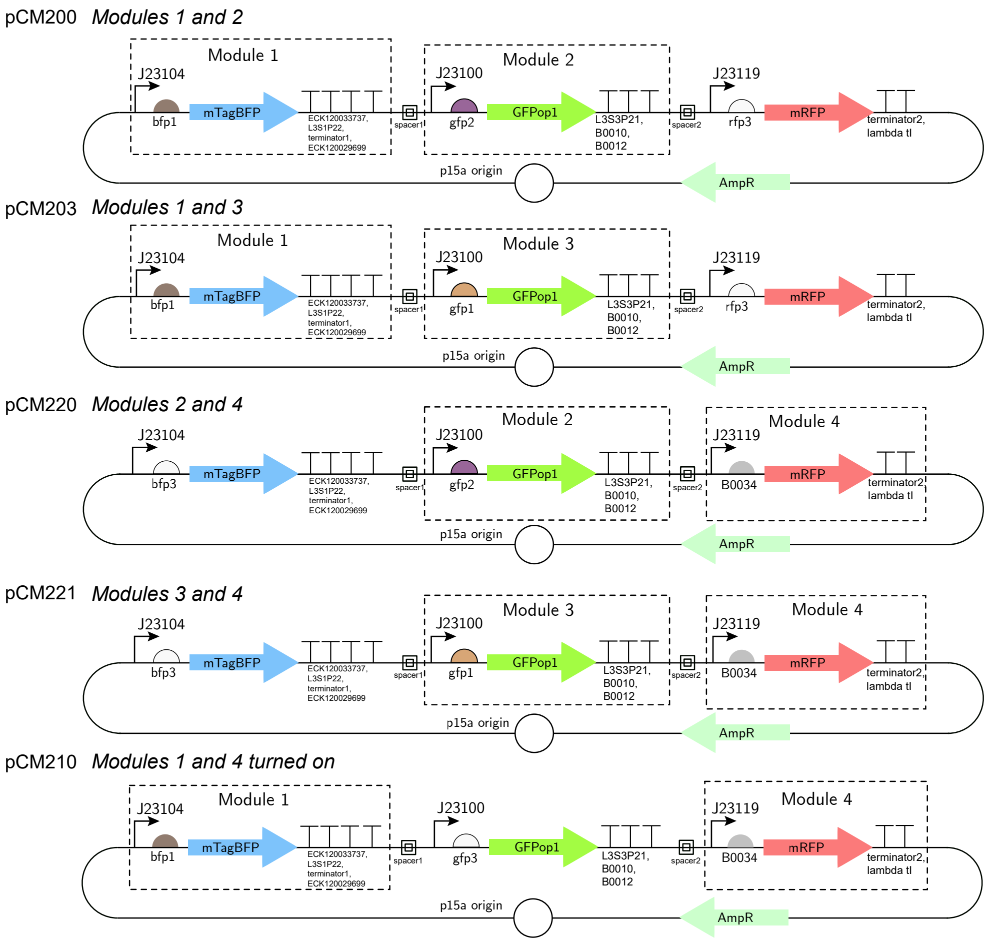


Figure S17: Plasmid maps of modules 1-4 perturbed by other modules.

Table S9: Description of ribosome binding site strengths for the inducible constructs.

| Plasmid | Description | RBS 1 | RBS 2 | RBS 3 |
|---|---|---|---|---|
| pCM172 | Module 1 and 5 | bfp1 (TIR = 5000) | B0064 | phl2 |
| pCM173 | Module 5 and RS1 | bfp2 (TIR = 2503) | B0064 | phl2 |
| pCM182 | Module 3 and 5 | gfp1 (TIR = 10211) | B0064 | phl2 |
| pCM183 | Module 2 and 5 | gfp2 (TIR = 6298) | B0064 | phl2 |
| pCM184 | Module 5 | gfp3 (TIR = 1.1) | B0064 | phl2 |
| pCM192 | Module 5 and RS2 | rfp1 (TIR = 2000) | B0064 | phl2 |
| pCM193 | Module 4 and 5 | rfp2 (TIR = 942) | B0064 | phl2 |
| pCM194 | Module 5 | rfp3 (TIR = 1.0) | B0064 | phl2 |
| pCM240 | Module 5 | none | B0064 | phl2 |

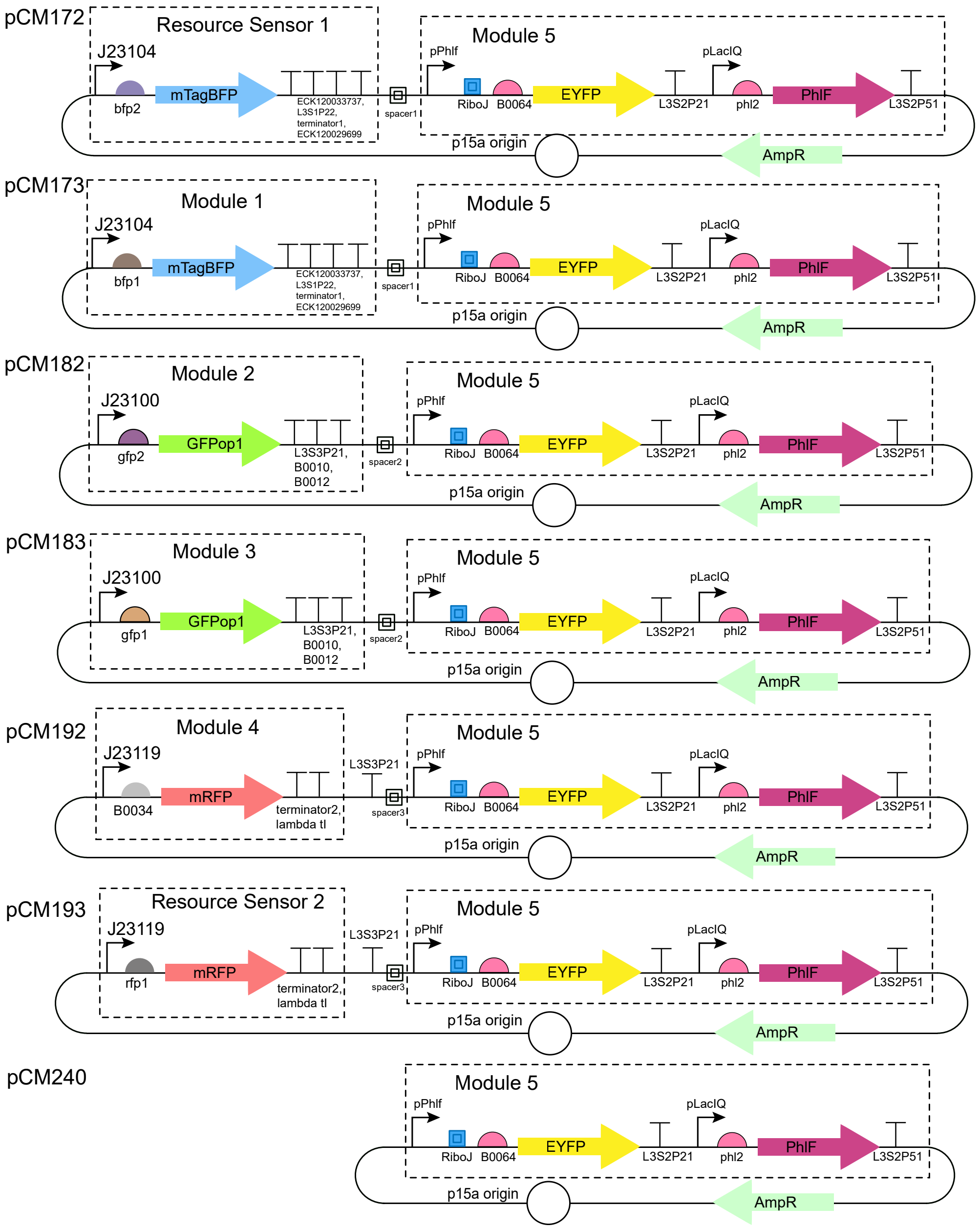


Figure S18: Plasmid maps of the inducible module with resource sensors and with other modules.

### S5 Plate Reader Data

#### S5.1 Correction graphs

Correlations were found to correct for fluorescence bleed between fluorescence measurement channels to apply linear unmixing techniques (*12*) as described in Methods, Section 4.6.

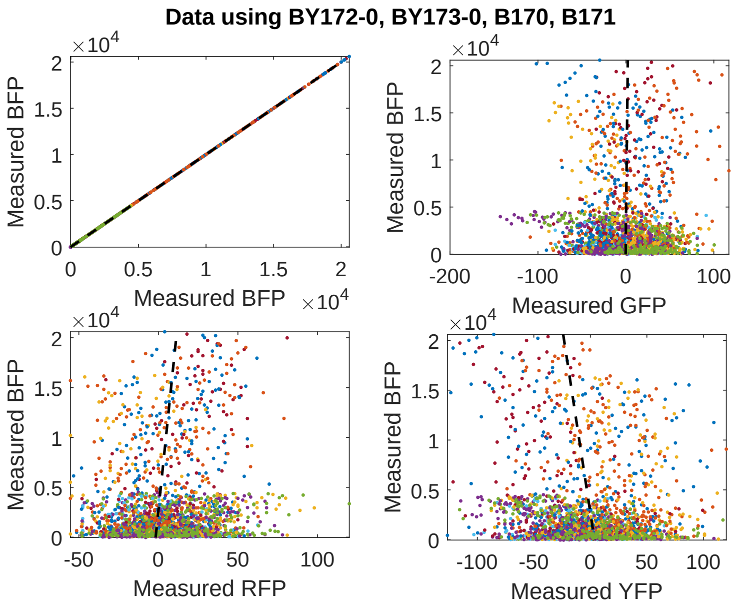


Figure S19: Correction of fluorescence bleed between fluorescence measurement channels for BFP proteins to populate the 1st column in the correction matrix $C$ in Section 4.6. Constructs that only translate mTagBFP proteins were used: pCM170, pCM171, pCM172 (0 nM induction), pCM173 (0 nM induction). Correlation between channels was used to measure the level of cross-talk between channels when GFP is present. Data were where the GFP measurement was greater than 20000 was not used as nonlinear effects become significant for large fluorescence values.

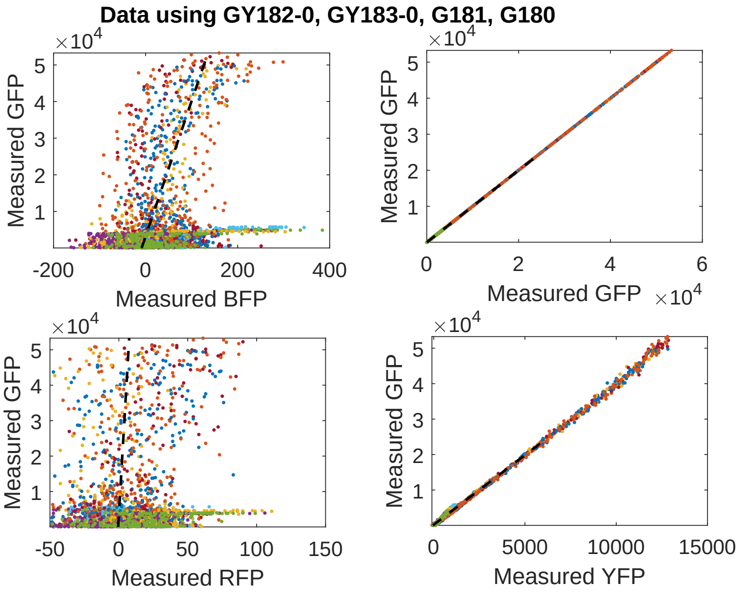


Figure S20: Correction of fluorescence bleed between fluorescence measurement channels for GFP proteins to populate the 2nd column in the correction matrix $C$ in Section 4.6. Constructs that only translate GFPop1 proteins were used: pCM180, pCM181, pCM182 (0 nM induction), pCM183 (0 nM induction). Correlation between channels was used to measure the level of cross-talk between channels when GFP is present. Data with GFP measurement values greater than 24000 were not used as nonlinear effects become significant for large fluorescence values.

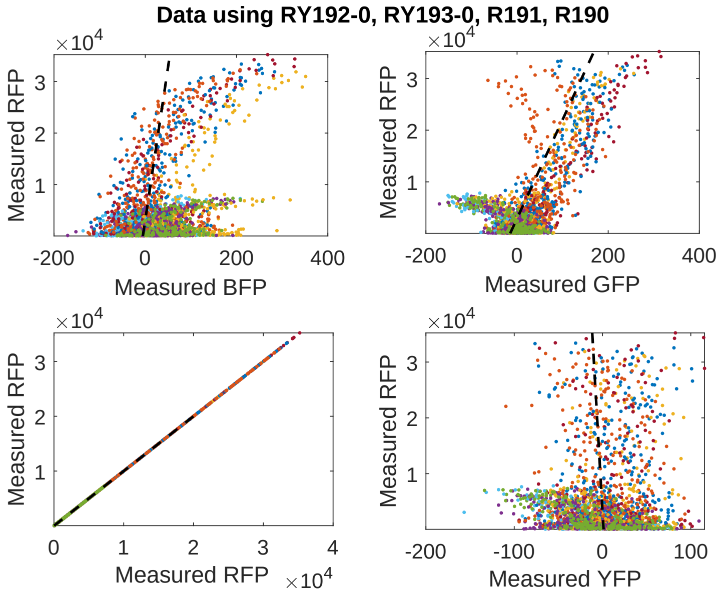


Figure S21: Correction of fluorescence bleed between fluorescence measurement channels for mRFP proteins to populate the 3rd column in the correction matrix $C$ in Section 4.6. Constructs that only translate mRFP proteins were used: pCM192 (0 nM induction), pCM193 (0 nM induction), pCM190, and pCM191. Correlation between channels was used to measure the level of cross-talk between channels when RFP is present. Data with RFP measurement values greater than 20000 were not used as nonlinear effects become significant for large fluorescence values.

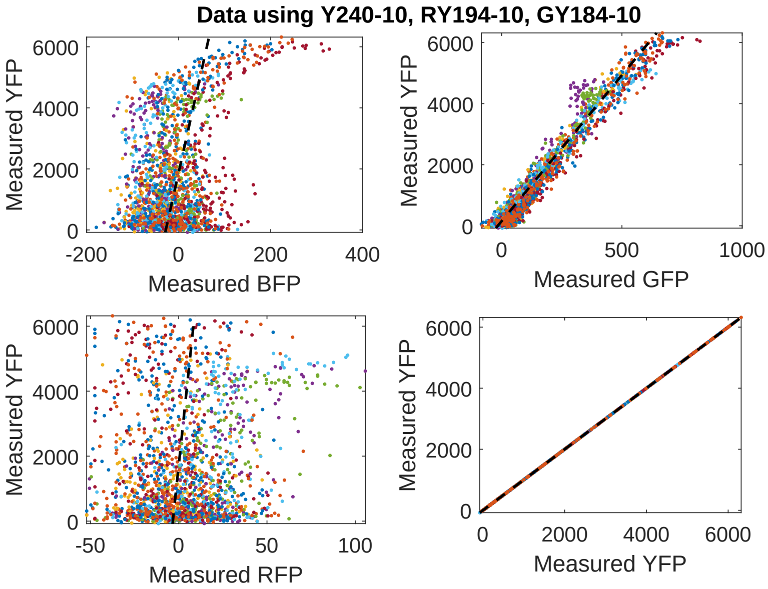


Figure S22: Correction of fluorescence bleed between fluorescence measurement channels for YFP proteins to populate the 4th column in the correction matrix $C$ in Section 4.6. Constructs that only translate EYFP proteins were used: pCM184 (10 nM induction), pCM194 (10 nM induction), and pCM240 (10 nM induction). Correlation between channels was used to measure the level of cross-talk between channels when YFP is present. Data with YFP measurement values greater than 10000 were not used as nonlinear effects become significant for large fluorescence values.

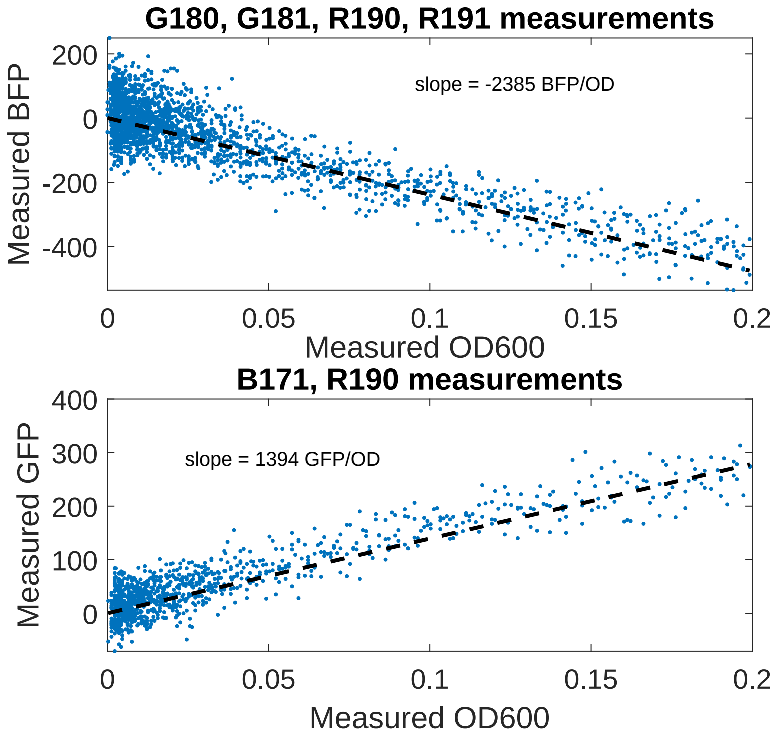


Figure S23: Correction of scattering effects on the BFP and GFP channels as a function of $OD_{600}$ measurements, see Section 4.5. Constructs that do not translate mTagBFP proteins were used to calibrate the BFP channel (top): pCM180, pCM181, pCM190, and pCM191. Constructs that do not translate GFPop1 proteins were used to calibrate the GFP channel (bottom): pCM171 and pCM190. Data with $OD_{600}$ measurement values larger than 0.2 were not used as nonlinear effects become significant for large $OD_{600}$ values.

#### S5.2 Transcriptional Context Comparison

There are some differences in module behavior due to transcriptional context the constructs used to test the module 5. As shown in Figure S24, the production rate of the module without other genes on its plasmid (other than the origin and selection marker) is larger than the production rate of the module when other genes are present but their ribosome binding site is turned off. This is likely due to transcriptional blocking effects of module 5's promoter from neighboring terminators.

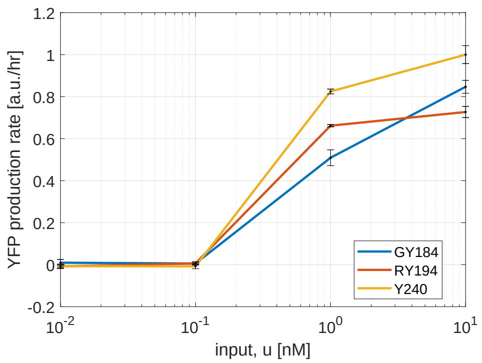


Figure S24: Comparison of output of YFP modules with different transcriptional context. The construct pCM184 has a GFP gene upstream of the YFP gene with the RBS turned off separated by a 40 bp spacer. The construct pCM184 has the RFP module upstream of the YFP gene with the RBS turned off. The construct pCM240 does not have another gene upstream of it and is used as the inducible module (module 5) alone in the main text. The YFP production rate of the pCM240 construct without an upstream module is larger than that of the other constructs due to transcriptional context.

#### S5.3 Production Rate Graphs

Time-course of fluorescent protein production rate per cell and measurements of $OD_{600}$ for all constructs. Fluorescent protein production rate per cell was used for the output of all modules, as described in SI Section S1.2.

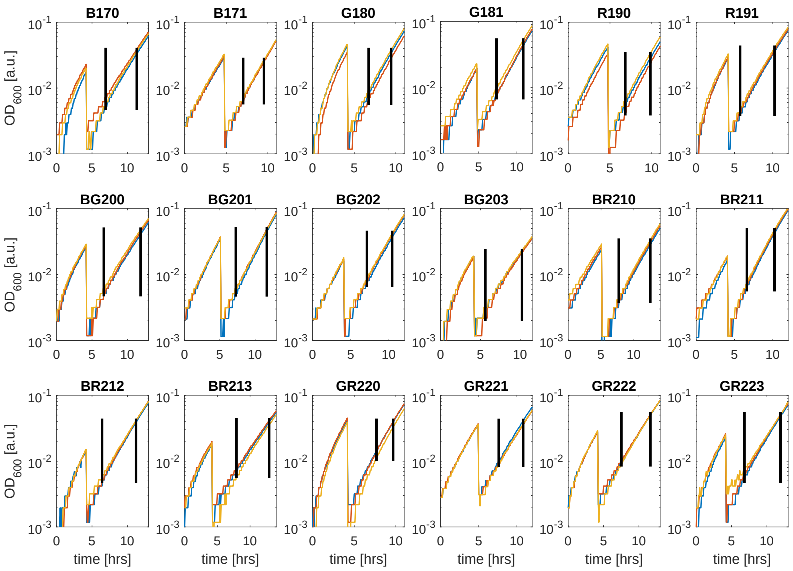


Figure S25: $OD_{600}$ vs time for plasmids pCM170, pCM171, pCM180, pCM181, pCM190, pCM191, pCM200, pCM201, pCM202, pCM203, pCM210, pCM211, pCM212, pCM213, pCM220, pCM221, pCM222, and pCM223. Three technical replicates of each construct are shown (blue, red, and yellow lines) and correspond to the fluorescent protein production rate data shown in Figures S28, S30, and S32. Steady states were selected in the time window between the vertical black lines. Note that the y-axis has a logarithmic scale.

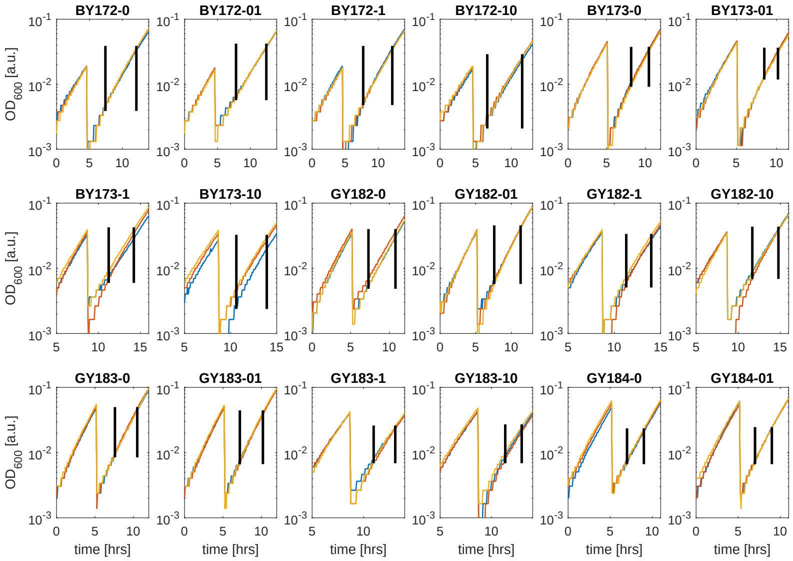


Figure S26: $OD_{600}$ vs time for plasmids pCM172, pCM173, pCM182, pCM183 and pCM14 for induction levels 0 nM, 0.1 nM, 1 nM, and 10 nM of DAPG. Three technical replicates of each construct are shown (blue, red, and yellow lines) and correspond to the fluorescent protein production rate data shown in Figures S29, S31, S34, and S35. Steady states were selected in the time window between the vertical black lines. Note that the y-axis has a logarithmic scale.

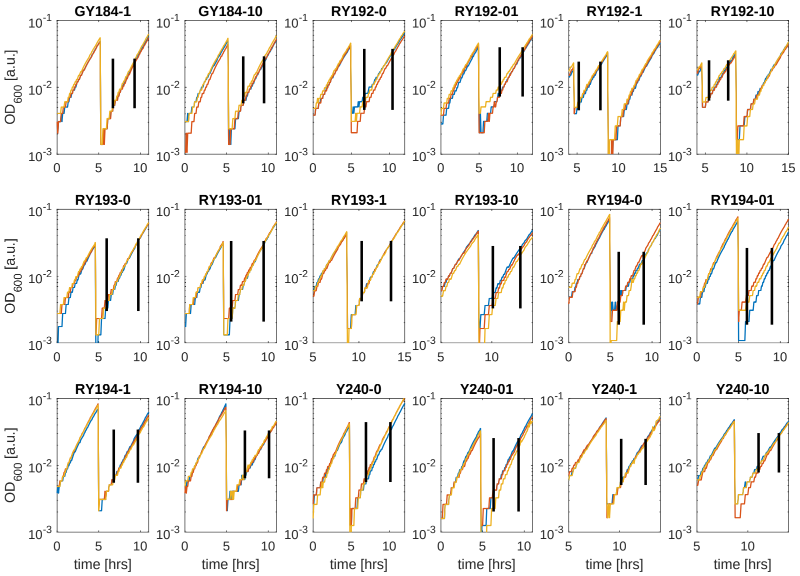


Figure S27: $OD_{600}$ vs time for plasmids pCM184, pCM192, pCM193, pCM194, and pCM240 for induction levels 0 nM, 0.1 nM, 1 nM, and 10 nM of DAPG. Three technical replicates of each construct are shown (blue, red, and yellow lines) and correspond to the fluorescent protein production rate data shown in Figures S31, S33, S36, and S37. Steady states were selected in the time window between the vertical black lines. Note that the y-axis has a logarithmic scale.

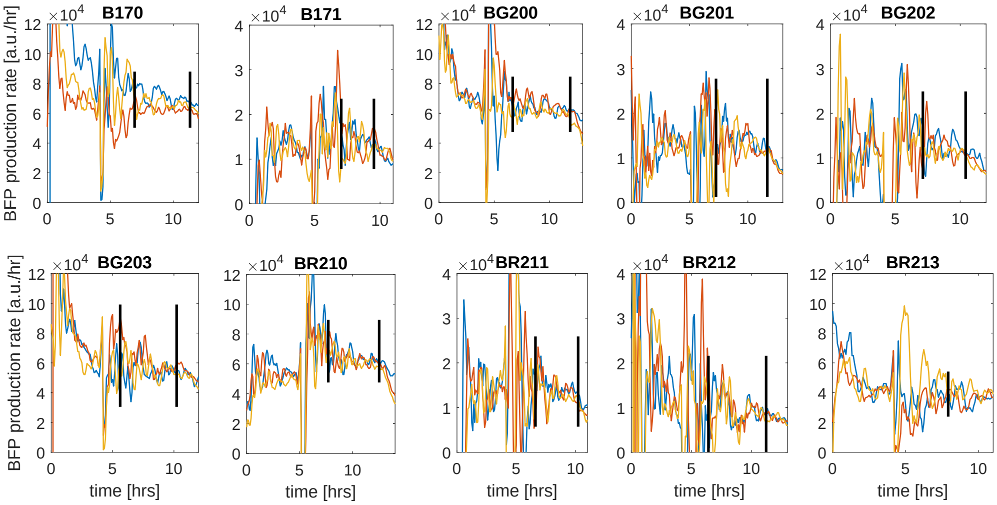


Figure S28: Production rate of BFP per cell for the constitutive plasmids pCM170, pCM171, pCM200, pCM201, pCM202, pCM203, pCM210, pCM211, pCM212, and pCM213 for three technical replicates (blue, red, and yellow lines) of each construct. Vertical black lines indicate time window where steady state was selected. Data shown corresponds to the $OD_{600}$ data shown in Figure S25.

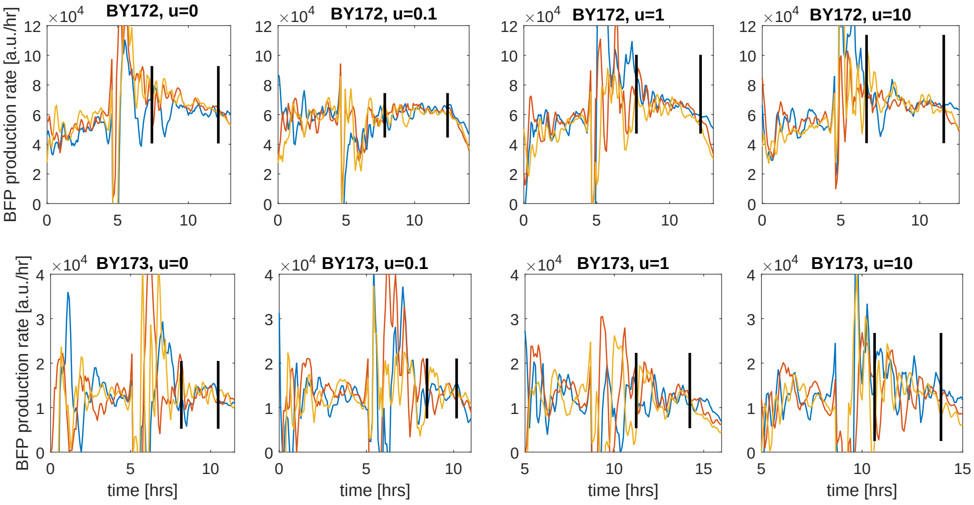


Figure S29: Production rate of BFP per cell for plasmids pCM172 and pCM173 for three technical replicates (blue, red, and yellow lines) of each construct with different levels of induction: 0 nM, 0.1 nM, 1 nM, and 10 nM of DAPG. Vertical black lines indicate time window where steady state was selected. Data shown corresponds to the $OD_{600}$ data shown in Figure S26.

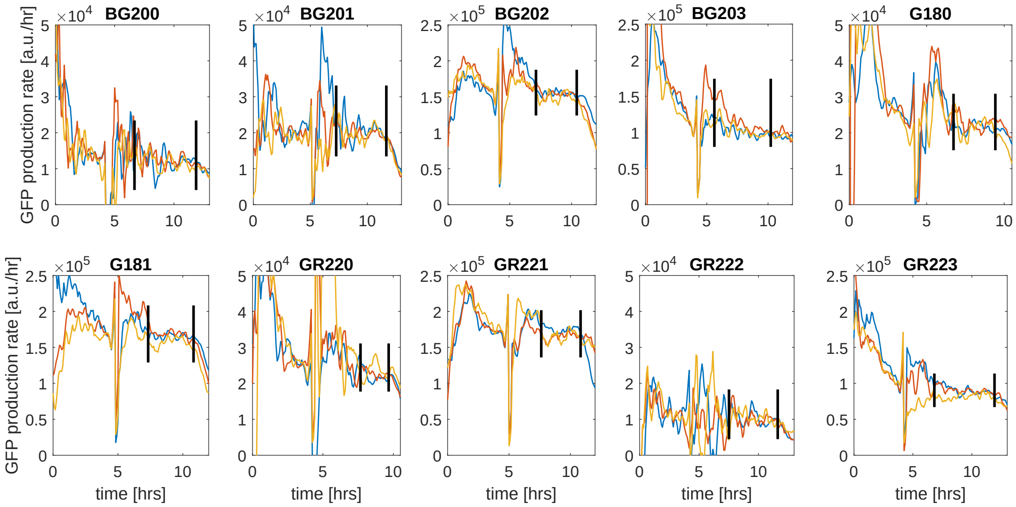


Figure S30: Production rate of GFP per cell for the constitutive plasmids pCM180, pCM181, pCM200, pCM201, pCM202, pCM203, pCM220, pCM221, pCM222, and pCM223 for three technical replicates (blue, red, and yellow lines) of each construct. Vertical black lines indicate time window where steady state was selected. Data shown corresponds to the $OD_{600}$ data shown in Figure S25.

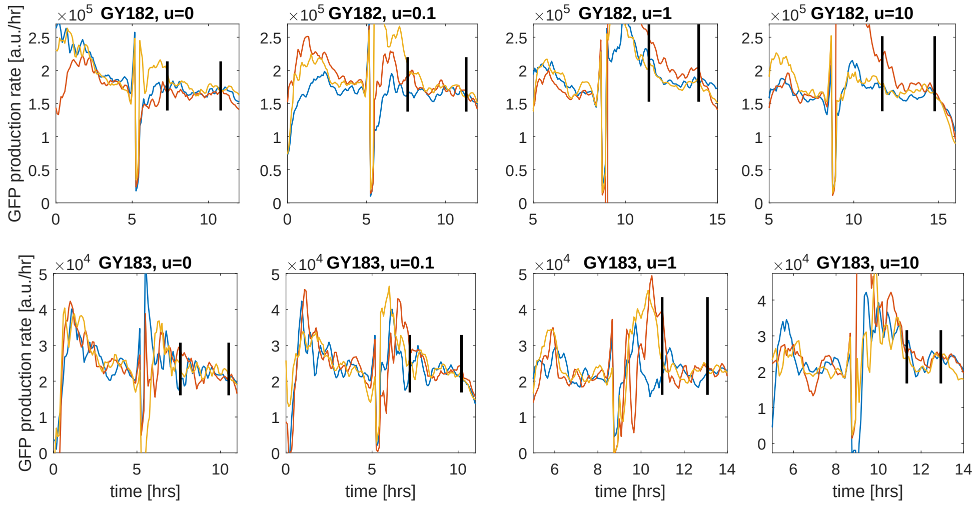


Figure S31: Production rate of GFP per cell for plasmids pCM182 and pCM183 for three technical replicates (blue, red, and yellow lines) of each construct with different levels of induction: 0 nM, 0.1 nM, 1 nM, and 10 nM of DAPG. Vertical black lines indicate time window where steady state was selected. Data shown corresponds to the $OD_{600}$ data shown in Figure S26.

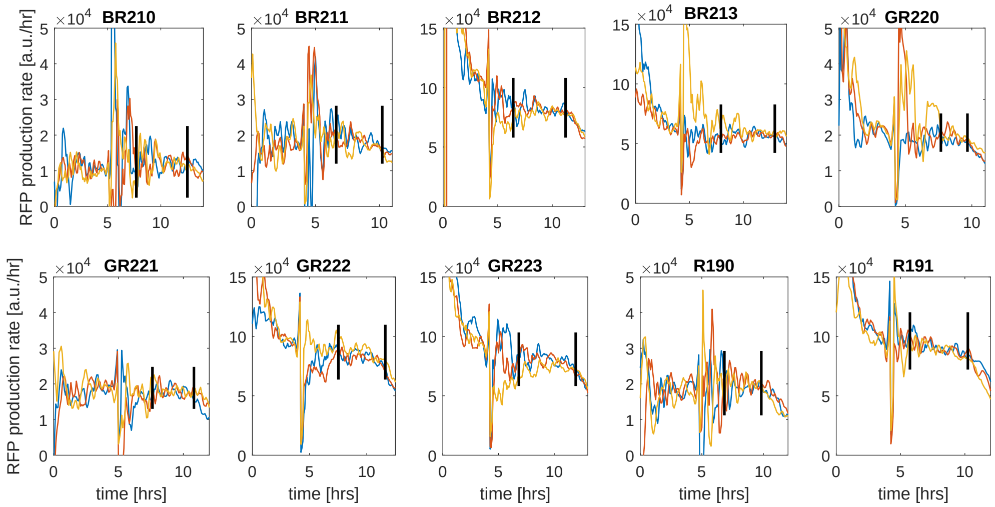


Figure S32: Production rate of RFP per cell for the constitutive plasmids pCM190, pCM191, pCM210, pCM211, pCM212, pCM213, pCM220, pCM221, pCM222, and pCM223 for three technical replicates (blue, red, and yellow lines) of each construct. Vertical black lines indicate time window where steady state was selected. Data shown corresponds to the $OD_{600}$ data shown in Figure S25.

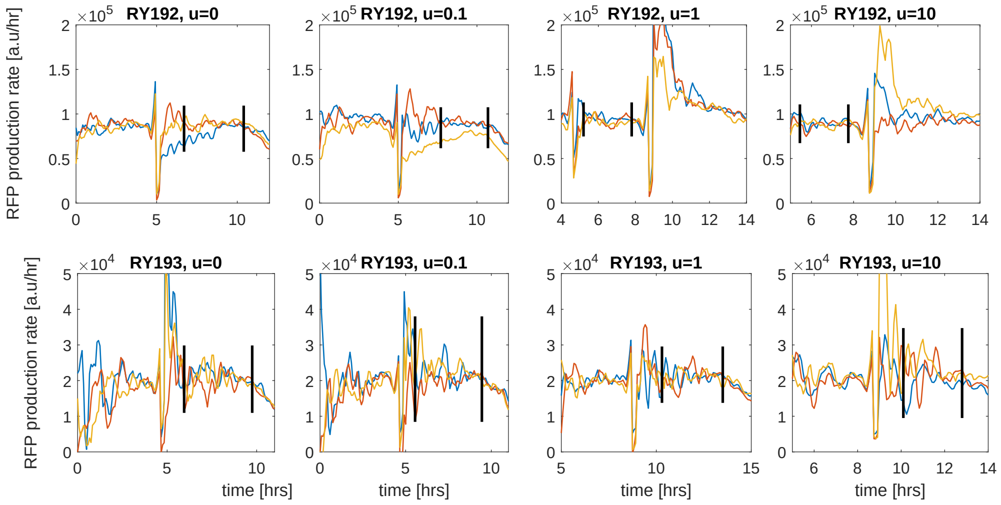


Figure S33: Production rate of RFP per cell for plasmids pCM192 and pCM193 for three technical replicates (blue, red, and yellow lines) of each construct with different levels of induction: 0 nM, 0.1 nM, 1 nM and 10 nM of DAPG. Vertical black lines indicate time window where steady state was selected. Data shown corresponds to the $OD_{600}$ data shown in Figure S27.

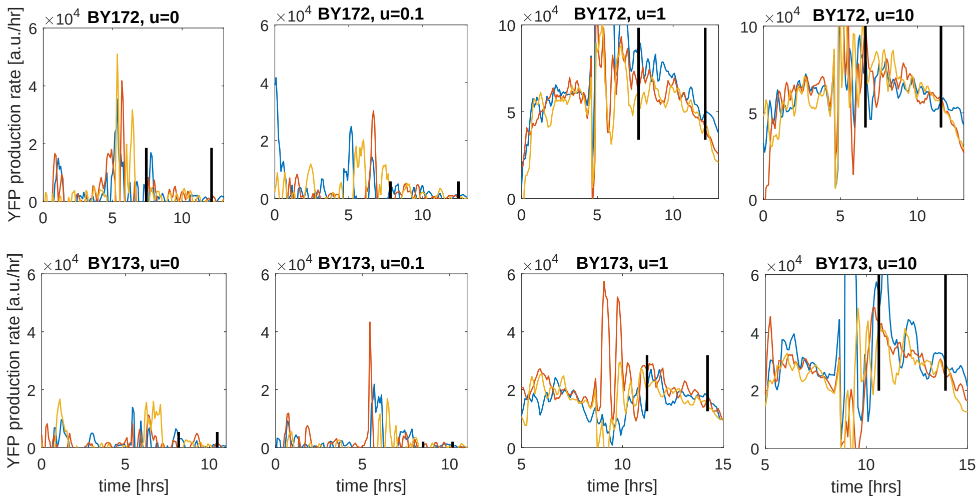


Figure S34: Production rate of YFP per cell for plasmids pCM172 and pCM173 for three technical replicates (blue, red, and yellow lines) of each construct with different levels of induction: 0 nM, 0.1 nM, 1 nM and 10 nM of DAPG. Vertical black lines indicate time window where steady state was selected. Data shown corresponds to the $OD_{600}$ data shown in Figure S26.

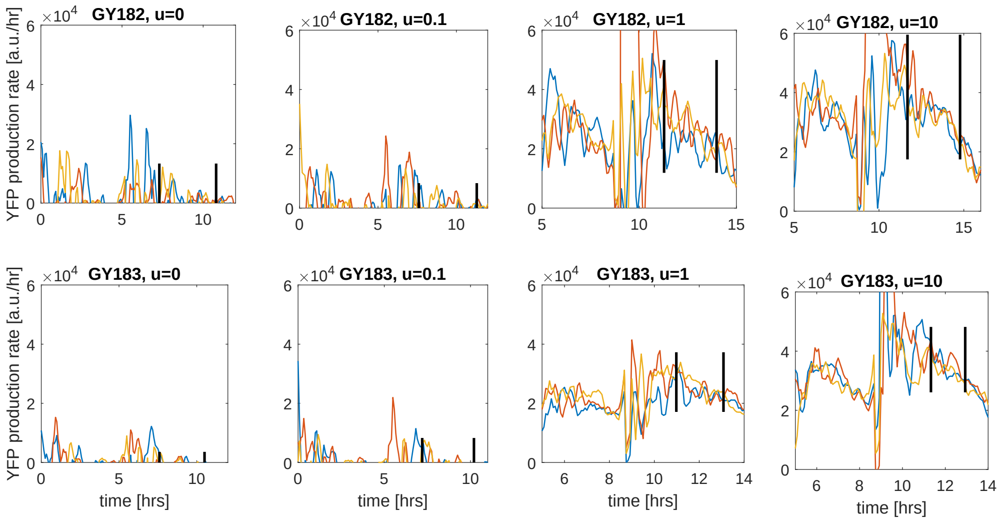


Figure S35: Production rate of YFP per cell for plasmids pCM182 and pCM183 for three technical replicates (blue, red, and yellow lines) of each construct with different levels of induction: 0 nM, 0.1 nM, 1 nM and 10 nM of DAPG. Vertical black lines indicate time window where steady state was selected. Data shown corresponds to the $OD_{600}$ data shown in Figure S26.

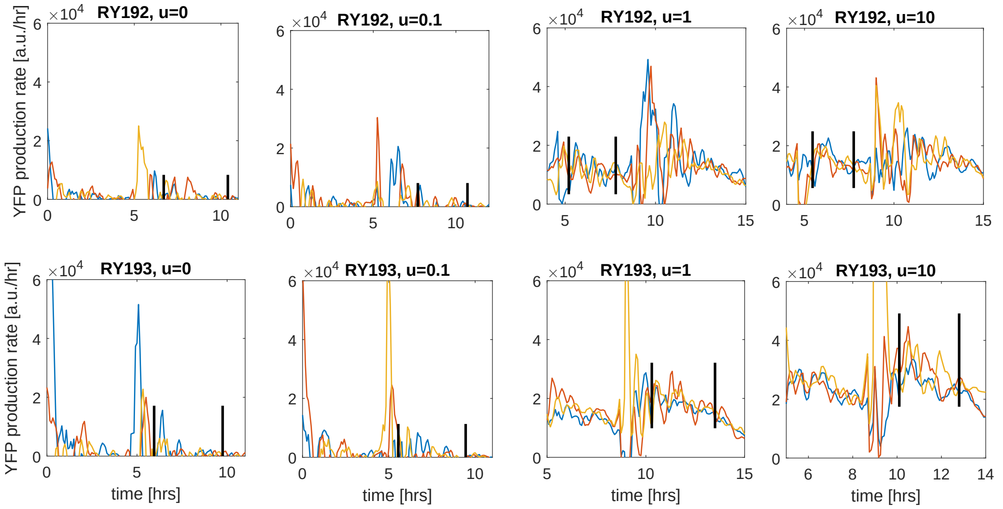


Figure S36: Production rate of YFP per cell for plasmids pCM192 and pCM193 for three technical replicates (blue, red, and yellow lines) of each construct with different levels of induction: 0 nM, 0.1 nM, 1 nM and 10 nM of DAPG. Vertical black lines indicate time window where steady state was selected. Data shown corresponds to the $OD_{600}$ data shown in Figure S27.

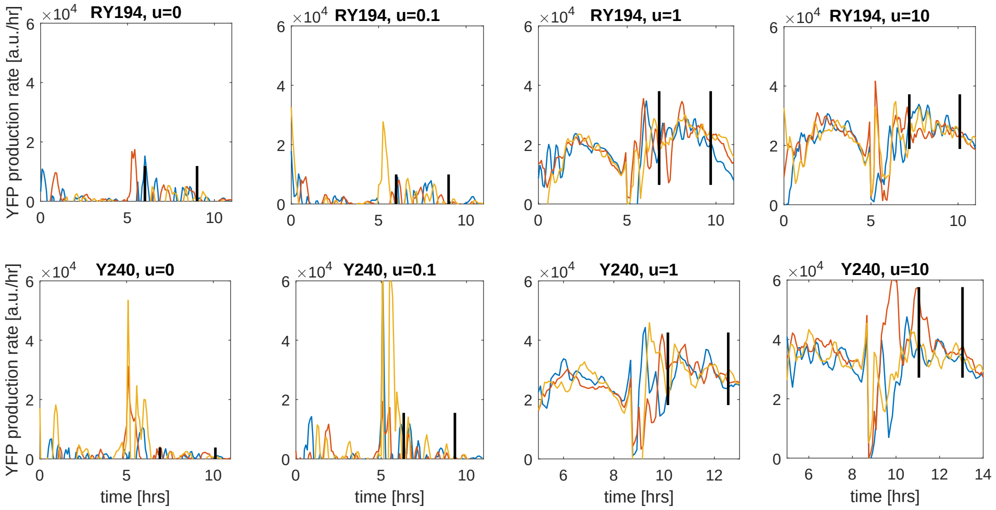


Figure S37: Production rate of YFP per cell for plasmids pCM194 and pCM240 for three technical replicates (blue, red, and yellow lines) of each construct with different levels of induction: 0 nM, 0.1 nM, 1 nM and 10 nM of DAPG. Vertical black lines indicate time window where steady state was selected. Data shown corresponds to the $OD_{600}$ data shown in Figure S27.

Table S10: Table of Plasmid sequences for experimental constructs

| Plasmid | Sequence |
|---|---|
| **pCM170** | TCGTGATGCTTGTCAGGGGGGCGGAGCCTATGGAAAAACGGCTTTGCCGCGGCCCTCTCACTTCCCTGTTAAGTATCTTCCTGGCATCTTCCAGGAAATC<br>TCCGCCCCGTTCGTAAGCCATTTCCGCTCGCCGCAGTCGAACGGGCGCGCCTTGACAGCTAGCTCAGTCCTAGGTATTGTGCTAGCTTACGTCAACCCAC<br>CCTAGGAGAGGATAAGATGAGCGAACTGATCAAAGAGAACATGCACATGAAGCTGTACATGGAGGGTACCGTGGATAATCACCACTTTAAGTGTACTTCT<br>GAGGGCGAGGGTAAGCCGTATGAAGGGACTCAAACGATGCGTATTAAAGTAGTGGAGGGTGGCCCACTGCCGTTTGCTTTCGATATTCTGGCGACGAGCT<br>TTCTGTATGGTAGCAAAACGTTTATAAACCACACTCAGGGCATTCCGGATTTCTTTAAACAAAGCTTTCCGGAAGGTTTTACCTGGGAGCGTGTGACTAC<br>GTATGAAGATGGTGGTGTGTTGACTGCTACTCAAGATACTTCACTGCAGGACGGCTGTCTGATCTATAACGTGAAGATTCGTGGCGTGAACTTTACGAGC<br>AATGGGCCGGTAATGCAAAAAAAAACCCTGGGTTGGGAAGCGTTCACGGAAACTCTGTATCCGGCTGACGGCGGCCTGGAGGGCCGTAACGATATGGCAC<br>TGAAGCTGGTTGGTGGCAGCCACCTGATCGCGAATATCAAAACGACTTATCGCTCTAAAAAACCGGCGAAAAATCTGAAGATGCCGGGTGTTTATTATGT<br>TGACTATCGTCTGGAACGCATTAAAGAAGCGAATAATGAAACTTACGTGGAGCAACACGAGGTTGCAGTGGCGCGCTATTGCGACTTGCCTTCAAAGCTG<br>GGTCACAAACTGAATTAATAAGGAAACACAGAAAAAAGCCCGCACCTGACAGTGCGGGCTTTTTTTTTCGACCAAAGGGACAATGTCCTGAAGAATATTT<br>AAGAAAAAAGCACCCCTCGTCGCCTAGAATTACCTACCGCGGTCGACCATACCTTCGAGACGAACAATAAGGCCGCAAATCGCGGCCTTCGTTATTGATA<br>ACAGCAGGTAAAAAGACGCTTATTATGGAGTTCGACATGGCAATAACGCGACGAATCTACGTCATGACGAGAATAGTATAAACAAAACTGCTGACGGCAG<br>AAGCGTCTCGGTACCAAATTCCAGAAAAGAGACGCTTTCGAGCGTCTTTTTTCGTTTTGGTCCACGCTGATGAATATTCTACGGAATTGCCATAGGCGTT<br>GAACGCTACACGGACGATACGAATTTATGTATAGAGCGTTCAGCCAAAAAACTTAAGACCGCCGGTCTTGTCCACTACCTTGCAGTAATGCGGTGGACAG<br>GATCGGCGGTTTTCTTTTCTCTTCTCAATTCGCGACCCAGGATGAGGTCGCCAAAACTACTAGATTTGGGCGCGCCTTGACGGCTAGCTCAGTCCTAGGT<br>ACAGTGCTAGCTTAATCTTGTCAAACAGTGGCTACGTGAACAGATGCGTAAAGGAGAAGAACTGTTCACTGGAGTTGTCCCAATTCTTGTTGAATTAGAT<br>GGTGATGTTAATGGGCACAAATTTTCTGTGAGTGGAGAGGGTGAAGGTGATGCGACATACGGCAAACTGACCCTTAAATTTATTTGTACTACCGGCAAGC<br>TACCTGTTCCTTGGCCGACACTTGTCACTACTTTCGGTTATGGTGTTCAATGTTTTGCGAGATACCCAGATCATATGAAACAGCATGACTTTTTCAAAAG<br>TGCCATGCCCGAAGGTTATGTACAGGAAAGAACTATATTTTTCAAAGATGACGGGAACTACAAAACACGTGCTGAAGTTAAATTTGAGGGTGATACTCTG<br>GTAAATAGAATCGAATTAAAAGGTATTGATTTTAAAGAAGATGGCAACATTCTTGGTCACAAGCTGGAATACAACTACAACTCACACAATGTTTACATCA<br>TGGCAGACAAACAAAAAAATGGCATCAAAGTTAACTTCAAAATTCGACACAACATTGAAGATGGAAGCGTTCAACTGGCAGACCATTATCAACAGAATAC<br>TCCGATTGGCGATGGCCCTGTACTCTTACCAGACAACCATTACCTGTCCACACAATCTGCCCTTTCGAAAGATCCGAACGAAAAGAGAGATCACATGGTC<br>CTGCTTGAGTTTGTAACAGCTGCTGGGATTACACATGGTATGGATGAGCTATACAAATAATAACCAATTATTGAAGGCCTCCCTAACGGGGGGCCTTTTT<br>TTGTTTCTGGTCTCCCCACTGATAGTGCTAGTGTAGATCACTACTAGAGCCAGGCATCAAATAAAACGAAAGGCTCAGTCGAAAGACTGGGCCTTTCGTT<br>TTATCTGTTGTTTGTCGGTGAACGCTCTCTACTAGAGTCACACTGGCTCACCTTCGGGTGGGCCTTTCTGCGTTTATATACTAGAGGGAATCGTACGGTC<br>GATCCATCTGACTATCGCCTACTTCATTGACAGCTAGCTCAGTCCTAGGTATAATGCTAGCAAATATGTTGTAGAAACAATTGCACGAATGGCTTCCTCC<br>GAAGACGTTATCAAAGAGTTCATGCGTTTCAAAGTTCGTATGGAAGGTTCCGTTAACGGTCACGAGTTCGAAATCGAAGGTGAAGGTGAAGGTCGTCCGT<br>ACGAAGGTACCCAGACCGCTAAACTGAAAGTTACCAAAGGTGGTCCGCTGCCGTTCGCTTGGGACATCCTGTCCCCGCAGTTCCAGTACGGTTCCAAAGC<br>TTACGTTAAACACCCGGCTGACATCCCGGACTACCTGAAACTGTCCTTCCCGGAAGGTTTCAAATGGGAACGTGTTATGAACTTCGAAGACGGTGGTGTT<br>GTTACCGTTACCCAGGACTCCTCCCTGCAAGACGGTGAGTTCATCTACAAAGTTAAACTGCGTGGTACCAACTTCCCGTCCGACGGTCCGGTTATGCAGA<br>AAAAAACCATGGGTTGGGAAGCTTCCACCGAACGTATGTACCCGGAAGACGGTGCTCTGAAAGGTGAAATCAAAATGCGTCTGAAACTGAAAGACGGTGG<br>TCACTACGACGCTGAAGTTAAAACCACCTACATGGCTAAAAAACCGGTTCAGCTGCCGGGTGCTTACAAAACCGACATCAAACTGGACATCACCTCCCAC<br>AACGAAGACTACACCATCGTTGAACAGTACGAACGTGCTGAAGGTCGTCACTCCACCGGTGCTTAATAACGCTGATAGTGCTAGTGTAGATCGCTACTAG<br>AGTACTAGCTCTAGATGCGGCCGCGAATTCCAGAAATCATCCTTAGCGAAAGCTAAGGATTTTTTTTATCTGAAATGCCTCTGTAACAGAGCATTAGCGC<br>AAGGTGATTTTTGTCTTCTTGCGCTAATTTTTTGGCGTCTCCACTGTGCGCGGAACCCCTATTTGTTTATTTTTCTAAATACATTCAAATATGTATCCGC<br>TCATGAGACAATAACCCTGATAAATGCTTCAATAATATTGAAAAAGGAAGAGTATGAGTATTCAACATTTCCGTGTCGCCCTTATTCCCTTTTTTGCGGC<br>ATTTTGCCTTCCTGTTTTTGCTCACCCAGAAACGCTGGTGAAAGTAAAAGATGCTGAAGATCAGTTGGGTGCACGAGTGGGTTACATCGAACTGGATCTC<br>AACAGCGGTAAGATCCTTGAGAGTTTTCGCCCCGAAGAACGTTTTCCAATGATGAGCACTTTTAAAGTTCTGCTATGTGGCGCGGTATTATCCCGTATTG<br>ACGCCGGGCAAGAGCAACTCGGTCGCCGCATACACTATTCTCAGAATGACTTGGTTGAGTACTCACCAGTCACAGAAAAGCATCTTACGGATGGCATGAC<br>AGTAAGAGAATTATGCAGTGCTGCCATAACCATGAGTGATAACACTGCGGCCAACTTACTTCTGACAACGATCGGAGGACCGAAGGAGCTAACCGCTTTT<br>TTGCACAACATGGGGGATCATGTAACTCGCCTTGATCGTTGGGAACCGGAGCTGAATGAAGCCATACCAAACGACGAGCGTGACACCACGATGCCTGTAG<br>CAATGGCAACAACGTTGCGCAAACTATTAACTGGCGAACTACTTACTCTAGCTTCCCGGCAACAATTAATAGACTGGATGGAGGCGGATAAAGTTGCAGG<br>ACCACTTCTGCGCTCGGCCCTTCCGGCTGGCTGGTTTATTGCTGATAAATCTGGAGCCGGTGAGCGTGGGTCTCGCGGTATCATTGCAGCACTGGGGCCA<br>GATGGTAAGCCCTCCCGTATCGTAGTTATCTACACGACGGGGAGTCAGGCAACTATGGATGAACGAAATAGACAGATCGCTGAGATAGGTGCCTCACTGA<br>TTAAGCATTGGTAACTCGAGCTGATCCTTCAAGAGCTCGCTTGGACTCCTGTTGATAGATCCAGTAATGACCTCAGAACTCCATCTGGATTTGTTCAGAA<br>CGCTCGGTTGCCGCCGGGCGTTTTTTATTGGTGAGAATTTACAACTTATATCGTATGGGGCTGACTTCAGGTGCTACATTTGAAGAGATAAATTGCACTG<br>AAATCTAGAAATATTTTATCTGATTAATAAGATGATCTTCTTGAGATCGTTTTGGTCTGCGCGTAATCTCTTGCTCTGAAAACGAAAAAACCGCCTTGCA<br>GGGCGGTTTTTCGAAGGTTCTCTGAGCTACCAACTCTTTGAACCGAGGTAACTGGCTTGGAGGAGCGCAGTCACCAAAACTTGTCCTTTCAGTTTAGCCT<br>TAACCGGCGCATGACTTCAAGACTAACTCCTCTAAATCAATTACCAGTGGCTGCTGCCAGTGGTGCTTTTGCATGTCTTTCCGGGTTGGACTCAAGACGA<br>TAGTTACCGGATAAGGCGCAGCGGTCGGACTGAACGGGGGGTTCGTGCATACAGTCCAGCTTGGAGCGAACTGCCTACCCGGAACTGAGTGTCAGGCGTG<br>GAATGAGACAAACGCGGCCATAACAGCGGAATGACACCGGTAAACCGAAAGGCAGGAACAGGAGAGCGCACGAGGGAGCCGCCAGGGGGAAACGCCTGGT<br>ATCTTTATAGTCCTGTCGGGTTTCGCCACCACTGATTTGAGCGTCAGATT |
| **pCM171** | TCGTGATGCTTGTCAGGGGGGCGGAGCCTATGGAAAAACGGCTTTGCCGCGGCCCTCTCACTTCCCTGTTAAGTATCTTCCTGGCATCTTCCAGGAAATC<br>TCCGCCCCGTTCGTAAGCCATTTCCGCTCGCCGCAGTCGAACGGGCGCGCCTTGACAGCTAGCTCAGTCCTAGGTATTGTGCTAGCTTACGCCGATTCTG<br>CCGGGGGGAATTATAATGAGCGAACTGATCAAAGAGAACATGCACATGAAGCTGTACATGGAGGGTACCGTGGATAATCACCACTTTAAGTGTACTTCTG<br>AGGGCGAGGGTAAGCCGTATGAAGGGACTCAAACGATGCGTATTAAAGTAGTGGAGGGTGGCCCACTGCCGTTTGCTTTCGATATTCTGGCGACGAGCTT<br>TCTGTATGGTAGCAAAACGTTTATAAACCACACTCAGGGCATTCCGGATTTCTTTAAACAAAGCTTTCCGGAAGGTTTTACCTGGGAGCGTGTGACTACG<br>TATGAAGATGGTGGTGTGTTGACTGCTACTCAAGATACTTCACTGCAGGACGGCTGTCTGATCTATAACGTGAAGATTCGTGGCGTGAACTTTACGAGCA<br>ATGGGCCGGTAATGCAAAAAAAAACCCTGGGTTGGGAAGCGTTCACGGAAACTCTGTATCCGGCTGACGGCGGCCTGGAGGGCCGTAACGATATGGCACT<br>GAAGCTGGTTGGTGGCAGCCACCTGATCGCGAATATCAAAACGACTTATCGCTCTAAAAAACCGGCGAAAAATCTGAAGATGCCGGGTGTTTATTATGTT<br>GACTATCGTCTGGAACGCATTAAAGAAGCGAATAATGAAACTTACGTGGAGCAACACGAGGTTGCAGTGGCGCGCTATTGCGACTTGCCTTCAAAGCTGG<br>GTCACAAACTGAATTAATAAGGAAACACAGAAAAAAGCCCGCACCTGACAGTGCGGGCTTTTTTTTTCGACCAAAGGGACAATGTCCTGAAGAATATTTA<br>AGAAAAAAGCACCCCTCGTCGCCTAGAATTACCTACCGCGGTCGACCATACCTTCGAGACGAACAATAAGGCCGCAAATCGCGGCCTTCGTTATTGATAA<br>CAGCAGGTAAAAAGACGCTTATTATGGAGTTCGACATGGCAATAACGCGACGAATCTACGTCATGACGAGAATAGTATAAACAAAACTGCTGACGGCAGA<br>AGCGTCTCGGTACCAAATTCCAGAAAAGAGACGCTTTCGAGCGTCTTTTTTCGTTTTGGTCCACGCTGATGAATATTCTACGGAATTGCCATAGGCGTTG<br>AACGCTACACGGACGATACGAATTTATGTATAGAGCGTTCAGCCAAAAAACTTAAGACCGCCGGTCTTGTCCACTACCTTGCAGTAATGCGGTGGACAGG<br>ATCGGCGGTTTTCTTTTCTCTTCTCAATTCGCGACCCAGGATGAGGTCGCCAAAACTACTAGATTTGGGCGCGCCTTGACGGCTAGCTCAGTCCTAGGTA<br>CAGTGCTAGCTTAATCTTGTCAAACAGTGGCTACGTGAACAGATGCGTAAAGGAGAAGAACTGTTCACTGGAGTTGTCCCAATTCTTGTTGAATTAGATG<br>GTGATGTTAATGGGCACAAATTTTCTGTGAGTGGAGAGGGTGAAGGTGATGCGACATACGGCAAACTGACCCTTAAATTTATTTGTACTACCGGCAAGCT<br>ACCTGTTCCTTGGCCGACACTTGTCACTACTTTCGGTTATGGTGTTCAATGTTTTGCGAGATACCCAGATCATATGAAACAGCATGACTTTTTCAAAAGT<br>GCCATGCCCGAAGGTTATGTACAGGAAAGAACTATATTTTTCAAAGATGACGGGAACTACAAAACACGTGCTGAAGTTAAATTTGAGGGTGATACTCTGG<br>TAAATAGAATCGAATTAAAAGGTATTGATTTTAAAGAAGATGGCAACATTCTTGGTCACAAGCTGGAATACAACTACAACTCACACAATGTTTACATCAT<br>GGCAGACAAACAAAAAAATGGCATCAAAGTTAACTTCAAAATTCGACACAACATTGAAGATGGAAGCGTTCAACTGGCAGACCATTATCAACAGAATACT<br>CCGATTGGCGATGGCCCTGTACTCTTACCAGACAACCATTACCTGTCCACACAATCTGCCCTTTCGAAAGATCCGAACGAAAAGAGAGATCACATGGTCC<br>TGCTTGAGTTTGTAACAGCTGCTGGGATTACACATGGTATGGATGAGCTATACAAATAATAACCAATTATTGAAGGCCTCCCTAACGGGGGGCCTTTTTT<br>TGTTTCTGGTCTCCCCACTGATAGTGCTAGTGTAGATCACTACTAGAGCCAGGCATCAAATAAAACGAAAGGCTCAGTCGAAAGACTGGGCCTTTCGTTT<br>TATCTGTTGTTTGTCGGTGAACGCTCTCTACTAGAGTCACACTGGCTCACCTTCGGGTGGGCCTTTCTGCGTTTATATACTAGAGGGAATCGTACGGTCG |

ATCCATCTGACTATCGCCTACTTCATTGACAGCTAGCTCAGTCCTAGGTATAATGCTAGCAAATATGTTGTAGAAACAATTGCACGAATGGCTTCCTCCG
AAGACGTTATCAAAGAGTTCATGCGTTTCAAAGTTCGTATGGAAGGTTCCGTTAACGGTCACGAGTTCGAAATCGAAGGTGAAGGTGAAGGTCGTCCGTA
CGAAGGTACCCAGACCGCTAAACTGAAAGTTACCAAAGGTGGTCCGCTGCCGTTCGCTTGGGACATCCTGTCCCCGCAGTTCCAGTACGGTTCCAAAGCT
TACGTTAAACACCCGGCTGACATCCCGGACTACCTGAAACTGTCCTTCCCGGAAGGTTTCAAATGGGAACGTGTTATGAACTTCGAAGACGGTGGTGTTG
TTACCGTTACCCAGGACTCCTCCCTGCAAGACGGTGAGTTCATCTACAAAGTTAAACTGCGTGGTACCAACTTCCCGTCCGACGGTCCGGTTATGCAGAA
AAAAACCATGGGTTGGGAAGCTTCCACCGAACGTATGTACCCGGAAGACGGTGCTCTGAAAGGTGAAATCAAAATGCGTCTGAAACTGAAAGACGGTGGT
CACTACGACGCTGAAGTTAAAACCACCTACATGGCTAAAAAACCGGTTCAGCTGCCGGGTGCTTACAAAACCGACATCAAACTGGACATCACCTCCCACA
ACGAAGACTACACCATCGTTGAACAGTACGAACGTGCTGAAGGTCGTCACTCCACCGGTGCTTAATAACGCTGATAGTGCTAGTGTAGATCGCTACTAGA
GTACTAGCTCTAGATGCGGCCGCGAATTCCAGAAATCATCCTTAGCGAAAGCTAAGGATTTTTTTTATCTGAAATGCCTCTGTAACAGAGCATTAGCGCA
AGGTGATTTTTGTCTTCTTGCGCTAATTTTTTGGCGTCTCCACTGTGCGCGGAACCCCTATTTGTTTATTTTTCTAAATACATTCAAATATGTATCCGCT
CATGAGACAATAACCCTGATAAATGCTTCAATAATATTGAAAAAGGAAGAGTATGAGTATTCAACATTTCCGTGTCGCCCTTATTCCCTTTTTTGCGGCA
TTTTGCCTTCCTGTTTTTGCTCACCCAGAAACGCTGGTGAAAGTAAAAGATGCTGAAGATCAGTTGGGTGCACGAGTGGGTTACATCGAACTGGATCTCA
ACAGCGGTAAGATCCTTGAGAGTTTTCGCCCCGAAGAACGTTTTCCAATGATGAGCACTTTTAAAGTTCTGCTATGTGGCGCGGTATTATCCCGTATTGA
CGCCGGGCAAGAGCAACTCGGTCGCCGCATACACTATTCTCAGAATGACTTGGTTGAGTACTCACCAGTCACAGAAAAGCATCTTACGGATGGCATGACA
GTAAGAGAATTATGCAGTGCTGCCATAACCATGAGTGATAACACTGCGGCCAACTTACTTCTGACAACGATCGGAGGACCGAAGGAGCTAACCGCTTTTT
TGCACAACATGGGGGATCATGTAACTCGCCTTGATCGTTGGGAACCGGAGCTGAATGAAGCCATACCAAACGACGAGCGTGACACCACGATGCCTGTAGC
AATGGCAACAACGTTGCGCAAACTATTAACTGGCGAACTACTTACTCTAGCTTCCCGGCAACAATTAATAGACTGGATGGAGGCGGATAAAGTTGCAGGA
CCACTTCTGCGCTCGGCCCTTCCGGCTGGCTGGTTTATTGCTGATAAATCTGGAGCCGGTGAGCGTGGGTCTCGCGGTATCATTGCAGCACTGGGGCCAG
ATGGTAAGCCCTCCCGTATCGTAGTTATCTACACGACGGGGAGTCAGGCAACTATGGATGAACGAAATAGACAGATCGCTGAGATAGGTGCCTCACTGAT
TAAGCATTGGTAACTCGAGCTGATCCTTCAAGAGCTCGCTTGGACTCCTGTTGATAGATCCAGTAATGACCTCAGAACTCCATCTGGATTTGTTCAGAAC
GCTCGGTTGCCGCCGGGCGTTTTTTATTGGTGAGAATTTACAACTTATATCGTATGGGGCTGACTTCAGGTGCTACATTTGAAGAGATAAATTGCACTGA
AATCTAGAAATATTTTATCTGATTAATAAGATGATCTTCTTGAGATCGTTTTGGTCTGCGCGTAATCTCTTGCTCTGAAAACGAAAAAACCGCCTTGCAG
GGCGGTTTTTCGAAGGTTCTCTGAGCTACCAACTCTTTGAACCGAGGTAACTGGCTTGGAGGAGCGCAGTCACCAAAACTTGTCCTTTCAGTTTAGCCTT
AACCGGCGCATGACTTCAAGACTAACTCCTCTAAATCAATTACCAGTGGCTGCTGCCAGTGGTGCTTTTGCATGTCTTTCCGGGTTGGACTCAAGACGAT
AGTTACCGGATAAGGCGCAGCGGTCGGACTGAACGGGGGGTTCGTGCATACAGTCCAGCTTGGAGCGAACTGCCTACCCGGAACTGAGTGTCAGGCGTGG
AATGAGACAAACGCGGCCATAACAGCGGAATGACACCGGTAAACCGAAAGGCAGGAACAGGAGAGCGCACGAGGGAGCCGCCAGGGGGAAACGCCTGGTA
TCTTTATAGTCCTGTCGGGTTTCGCCACCACTGATTTGAGCGTCAGATT

**pCM180**

TCGTGATGCTTGTCAGGGGGGCGGAGCCTATGGAAAAACGGCTTTGCCGCGGCCCTCTCACTTCCCTGTTAAGTATCTTCCTGGCATCTTCCAGGAAATC
TCCGCCCCGTTCGTAAGCCATTTCCGCTCGCCGCAGTCGAACGGGCGCGCCTTGACAGCTAGCTCAGTCCTAGGTATTGTGCTAGCTTACGAGCGATGGG
CTCACTGTTGAGCCTCAATGAGCGAACTGATCAAAGAGAACATGCACATGAAGCTGTACATGGAGGGTACCGTGGATAATCACCACTTTAAGTGTACTTC
TGAGGGCGAGGGTAAGCCGTATGAAGGGACTCAAACGATGCGTATTAAAGTAGTGGAGGGTGGCCCACTGCCGTTTGCTTTCGATATTCTGGCGACGAGC
TTTCTGTATGGTAGCAAAACGTTTATAAACCACACTCAGGGCATTCCGGATTTCTTTAAACAAAGCTTTCCGGAAGGTTTTACCTGGGAGCGTGTGACTA
CGTATGAAGATGGTGGTGTGTTGACTGCTACTCAAGATACTTCACTGCAGGACGGCTGTCTGATCTATAACGTGAAGATTCGTGGCGTGAACTTTACGAG
CAATGGGCCGGTAATGCAAAAAAAAACCCTGGGTTGGGAAGCGTTCACGGAAACTCTGTATCCGGCTGACGGCGGCCTGGAGGGCCGTAACGATATGGCA
CTGAAGCTGGTTGGTGGCAGCCACCTGATCGCGAATATCAAAACGACTTATCGCTCTAAAAAACCGGCGAAAAATCTGAAGATGCCGGGTGTTTATTATG
TTGACTATCGTCTGGAACGCATTAAAGAAGCGAATAATGAAACTTACGTGGAGCAACACGAGGTTGCAGTGGCGCGCTATTGCGACTTGCCTTCAAAGCT
GGGTCACAAACTGAATTAATAAGGAAACACAGAAAAAAGCCCGCACCTGACAGTGCGGGCTTTTTTTTTCGACCAAAGGGACAATGTCCTGAAGAATATT
TAAGAAAAAAGCACCCCTCGTCGCCTAGAATTACCTACCGCGGTCGACCATACCTTCGAGACGAACAATAAGGCCGCAAATCGCGGCCTTCGTTATTGAT
AACAGCAGGTAAAAAGACGCTTATTATGGAGTTCGACATGGCAATAACGCGACGAATCTACGTCATGACGAGAATAGTATAAACAAAACTGCTGACGGCA
GAAGCGTCTCGGTACCAAATTCCAGAAAAGAGACGCTTTCGAGCGTCTTTTTTCGTTTTGGTCCACGCTGATGAATATTCTACGGAATTGCCATAGGCGT
TGAACGCTACACGGACGATACGAATTTATGTATAGAGCGTTCAGCCAAAAAACTTAAGACCGCCGGTCTTGTCCACTACCTTGCAGTAATGCGGTGGACA
GGATCGGCGGTTTTCTTTTCTCTTCTCAATTCGCGACCCAGGATGAGGTCGCCAAAACTACTAGATTTGGGCGCGCCTTGACGGCTAGCTCAGTCCTAGG
TACAGTGCTAGCTTAAT
GCGAGAGGCTGGTTAAGTGGGTGTTG
ATGCGTAAAGGAGAAGAACTGTTCACTGGAGTTGTCCCAATTCTTGTTGAATTAGATGGTGATGTTAATGGGCACAAATTTTCTGTGAGTGGAGAGGGTG
AAGGTGATGCGACATACGGCAAACTGACCCTTAAATTTATTTGTACTACCGGCAAGCTACCTGTTCCTTGGCCGACACTTGTCACTACTTTCGGTTATGG
TGTTCAATGTTTTGCGAGATACCCAGATCATATGAAACAGCATGACTTTTTCAAAAGTGCCATGCCCGAAGGTTATGTACAGGAAAGAACTATATTTTTC
AAAGATGACGGGAACTACAAAACACGTGCTGAAGTTAAATTTGAGGGTGATACTCTGGTAAATAGAATCGAATTAAAAGGTATTGATTTTAAAGAAGATG
GCAACATTCTTGGTCACAAGCTGGAATACAACTACAACTCACACAATGTTTACATCATGGCAGACAAACAAAAAAATGGCATCAAAGTTAACTTCAAAAT
TCGACACAACATTGAAGATGGAAGCGTTCAACTGGCAGACCATTATCAACAGAATACTCCGATTGGCGATGGCCCTGTACTCTTACCAGACAACCATTAC
CTGTCCACACAATCTGCCCTTTCGAAAGATCCGAACGAAAAGAGAGATCACATGGTCCTGCTTGAGTTTGTAACAGCTGCTGGGATTACACATGGTATGG
ATGAGCTATACAAATAATAACCAATTATTGAAGGCCTCCCTAACGGGGGGCCTTTTTTTGTTTCTGGTCTCCCCACTGATAGTGCTAGTGTAGATCACTA
CTAGAGCCAGGCATCAAATAAAACGAAAGGCTCAGTCGAAAGACTGGGCCTTTCGTTTTATCTGTTGTTTGTCGGTGAACGCTCTCTACTAGAGTCACAC
TGGCTCACCTTCGGGTGGGCCTTTCTGCGTTTATATACTAGAGGGAATCGTACGGTCGATCCATCTGACTATCGCCTACTTCATTGACAGCTAGCTCAGT
CCTAGGTATAATGCTAGCAAATATGTTGTAGAAACAATTGCACGAATGGCTTCCTCCGAAGACGTTATCAAAGAGTTCATGCGTTTCAAAGTTCGTATGG
AAGGTTCCGTTAACGGTCACGAGTTCGAAATCGAAGGTGAAGGTGAAGGTCGTCCGTACGAAGGTACCCAGACCGCTAAACTGAAAGTTACCAAAGGTGG
TCCGCTGCCGTTCGCTTGGGACATCCTGTCCCCGCAGTTCCAGTACGGTTCCAAAGCTTACGTTAAACACCCGGCTGACATCCCGGACTACCTGAAACTG
TCCTTCCCGGAAGGTTTCAAATGGGAACGTGTTATGAACTTCGAAGACGGTGGTGTTGTTACCGTTACCCAGGACTCCTCCCTGCAAGACGGTGAGTTCA
TCTACAAAGTTAAACTGCGTGGTACCAACTTCCCGTCCGACGGTCCGGTTATGCAGAAAAAAACCATGGGTTGGGAAGCTTCCACCGAACGTATGTACCC
GGAAGACGGTGCTCTGAAAGGTGAAATCAAAATGCGTCTGAAACTGAAAGACGGTGGTCACTACGACGCTGAAGTTAAAACCACCTACATGGCTAAAAAA
CCGGTTCAGCTGCCGGGTGCTTACAAAACCGACATCAAACTGGACATCACCTCCCACAACGAAGACTACACCATCGTTGAACAGTACGAACGTGCTGAAG
GTCGTCACTCCACCGGTGCTTAATAACGCTGATAGTGCTAGTGTAGATCGCTACTAGAGTACTAGCTCTAGATGCGGCCGCGAATTCCAGAAATCATCCT
TAGCGAAAGCTAAGGATTTTTTTTATCTGAAATGCCTCTGTAACAGAGCATTAGCGCAAGGTGATTTTTGTCTTCTTGCGCTAATTTTTTGGCGTCTCCA
CTGTGCGCGGAACCCCTATTTGTTTATTTTTCTAAATACATTCAAATATGTATCCGCTCATGAGACAATAACCCTGATAAATGCTTCAATAATATTGAAA
AAGGAAGAGTATGAGTATTCAACATTTCCGTGTCGCCCTTATTCCCTTTTTTGCGGCATTTTGCCTTCCTGTTTTTGCTCACCCAGAAACGCTGGTGAAA
GTAAAAGATGCTGAAGATCAGTTGGGTGCACGAGTGGGTTACATCGAACTGGATCTCAACAGCGGTAAGATCCTTGAGAGTTTTCGCCCCGAAGAACGTT
TTCCAATGATGAGCACTTTTAAAGTTCTGCTATGTGGCGCGGTATTATCCCGTATTGACGCCGGGCAAGAGCAACTCGGTCGCCGCATACACTATTCTCA
GAATGACTTGGTTGAGTACTCACCAGTCACAGAAAAGCATCTTACGGATGGCATGACAGTAAGAGAATTATGCAGTGCTGCCATAACCATGAGTGATAAC
ACTGCGGCCAACTTACTTCTGACAACGATCGGAGGACCGAAGGAGCTAACCGCTTTTTTGCACAACATGGGGGATCATGTAACTCGCCTTGATCGTTGGG
AACCGGAGCTGAATGAAGCCATACCAAACGACGAGCGTGACACCACGATGCCTGTAGCAATGGCAACAACGTTGCGCAAACTATTAACTGGCGAACTACT
TACTCTAGCTTCCCGGCAACAATTAATAGACTGGATGGAGGCGGATAAAGTTGCAGGACCACTTCTGCGCTCGGCCCTTCCGGCTGGCTGGTTTATTGCT
GATAAATCTGGAGCCGGTGAGCGTGGGTCTCGCGGTATCATTGCAGCACTGGGGCCAGATGGTAAGCCCTCCCGTATCGTAGTTATCTACACGACGGGGA
GTCAGGCAACTATGGATGAACGAAATAGACAGATCGCTGAGATAGGTGCCTCACTGATTAAGCATTGGTAACTCGAGCTGATCCTTCAAGAGCTCGCTTG
GACTCCTGTTGATAGATCCAGTAATGACCTCAGAACTCCATCTGGATTTGTTCAGAACGCTCGGTTGCCGCCGGGCGTTTTTTATTGGTGAGAATTTACA
ACTTATATCGTATGGGGCTGACTTCAGGTGCTACATTTGAAGAGATAAATTGCACTGAAATCTAGAAATATTTTATCTGATTAATAAGATGATCTTCTTG
AGATCGTTTTGGTCTGCGCGTAATCTCTTGCTCTGAAAACGAAAAAACCGCCTTGCAGGGCGGTTTTTCGAAGGTTCTCTGAGCTACCAACTCTTTGAAC
CGAGGTAACTGGCTTGGAGGAGCGCAGTCACCAAAACTTGTCCTTTCAGTTTAGCCTTAACCGGCGCATGACTTCAAGACTAACTCCTCTAAATCAATTA
CCAGTGGCTGCTGCCAGTGGTGCTTTTGCATGTCTTTCCGGGTTGGACTCAAGACGATAGTTACCGGATAAGGCGCAGCGGTCGGACTGAACGGGGGGTT

CGTGCATACAGTCCAGCTTGGAGCGAACTGCCTACCCGGAACTGAGTGTCAGGCGTGGAATGAGACAAACGCGGCCATAACAGCGGAATGACACCGGTAA
ACCGAAAGGCAGGAACAGGAGAGCGCACGAGGGAGCCGCCAGGGGGAAACGCCTGGTATCTTTATAGTCCTGTCGGGTTTCGCCACCACTGATTTGAGCG
TCAGATT

**pCM181**

TCGTGATGCTTGTCAGGGGGGCGGAGCCTATGGAAAAACGGCTTTGCCGCGGCCCTCTCACTTCCCTGTTAAGTATCTTCCTGGCATCTTCCAGGAAATC
TCCGCCCCGTTCGTAAGCCATTTCCGCTCGCCGCAGTCGAACGGGCGCGCCTTGACAGCTAGCTCAGTCCTAGGTATTGTGCTAGCTTACGAGCGATGGG
CTCACTGTTGAGCCTCAATGAGCGAACTGATCAAAGAGAACATGCACATGAAGCTGTACATGGAGGGTACCGTGGATAATCACCACTTTAAGTGTACTTC
TGAGGGCGAGGGTAAGCCGTATGAAGGGACTCAAACGATGCGTATTAAAGTAGTGGAGGGTGGCCCACTGCCGTTTGCTTTCGATATTCTGGCGACGAGC
TTTCTGTATGGTAGCAAAACGTTTATAAACCACACTCAGGGCATTCCGGATTTCTTTAAACAAAGCTTTCCGGAAGGTTTTACCTGGGAGCGTGTGACTA
CGTATGAAGATGGTGGTGTGTTGACTGCTACTCAAGATACTTCACTGCAGGACGGCTGTCTGATCTATAACGTGAAGATTCGTGGCGTGAACTTTACGAG
CAATGGGCCGGTAATGCAAAAAAAAACCCTGGGTTGGGAAGCGTTCACGGAAACTCTGTATCCGGCTGACGGCGGCCTGGAGGGCCGTAACGATATGGCA
CTGAAGCTGGTTGGTGGCAGCCACCTGATCGCGAATATCAAAACGACTTATCGCTCTAAAAAACCGGCGAAAAATCTGAAGATGCCGGGTGTTTATTATG
TTGACTATCGTCTGGAACGCATTAAAGAAGCGAATAATGAAACTTACGTGGAGCAACACGAGGTTGCAGTGGCGCGCTATTGCGACTTGCCTTCAAAGCT
GGGTCACAAACTGAATTAATAAGGAAACACAGAAAAAAGCCCGCACCTGACAGTGCGGGCTTTTTTTTTCGACCAAAGGGACAATGTCCTGAAGAATATT
TAAGAAAAAAGCACCCCTCGTCGCCTAGAATTACCTACCGCGGTCGACCATACCTTCGAGACGAACAATAAGGCCGCAAATCGCGGCCTTCGTTATTGAT
AACAGCAGGTAAAAAGACGCTTATTATGGAGTTCGACATGGCAATAACGCGACGAATCTACGTCATGACGAGAATAGTATAAACAAAACTGCTGACGGCA
GAAGCGTCTCGGTACCAAATTCCAGAAAAGAGACGCTTTCGAGCGTCTTTTTTCGTTTTGGTCCACGCTGATGAATATTCTACGGAATTGCCATAGGCGT
TGAACGCTACACGGACGATACGAATTTATGTATAGAGCGTTCAGCCAAAAAACTTAAGACCGCCGGTCTTGTCCACTACCTTGCAGTAATGCGGTGGACA
GGATCGGCGGTTTTCTTTTCTCTTCTCAATTCGCGACCCAGGATGAGGTCGCCAAAACTACTAGATTTGGGCGCGCCTTGACGGCTAGCTCAGTCCTAGG
TACAGTGCTAGCTTAATTTCTAATTAGGATAAGGACACCAAGATGCGTAAAGGAGAAGAACTGTTCACTGGAGTTGTCCCAATTCTTGTTGAATTAGATG
GTGATGTTAATGGGCACAAATTTTCTGTGAGTGGAGAGGGTGAAGGTGATGCGACATACGGCAAACTGACCCTTAAATTTATTTGTACTACCGGCAAGCT
ACCTGTTCCTTGGCCGACACTTGTCACTACTTTCGGTTATGGTGTTCAATGTTTTGCGAGATACCCAGATCATATGAAACAGCATGACTTTTTCAAAAGT
GCCATGCCCGAAGGTTATGTACAGGAAAGAACTATATTTTTCAAAGATGACGGGAACTACAAAACACGTGCTGAAGTTAAATTTGAGGGTGATACTCTGG
TAAATAGAATCGAATTAAAAGGTATTGATTTTAAAGAAGATGGCAACATTCTTGGTCACAAGCTGGAATACAACTACAACTCACACAATGTTTACATCAT
GGCAGACAAACAAAAAAATGGCATCAAAGTTAACTTCAAAATTCGACACAACATTGAAGATGGAAGCGTTCAACTGGCAGACCATTATCAACAGAATACT
CCGATTGGCGATGGCCCTGTACTCTTACCAGACAACCATTACCTGTCCACACAATCTGCCCTTTCGAAAGATCCGAACGAAAAGAGAGATCACATGGTCC
TGCTTGAGTTTGTAACAGCTGCTGGGATTACACATGGTATGGATGAGCTATACAAATAATAACCAATTATTGAAGGCCTCCCTAACGGGGGGCCTTTTTT
TGTTTCTGGTCTCCCCACTGATAGTGCTAGTGTAGATCACTACTAGAGCCAGGCATCAAATAAAACGAAAGGCTCAGTCGAAAGACTGGGCCTTTCGTTT
TATCTGTTGTTTGTCGGTGAACGCTCTCTACTAGAGTCACACTGGCTCACCTTCGGGTGGGCCTTTCTGCGTTTATATACTAGAGGGAATCGTACGGTCG
ATCCATCTGACTATCGCCTACTTCATTGACAGCTAGCTCAGTCCTAGGTATAATGCTAGCAAATATGTTGTAGAAACAATTGCACGAATGGCTTCCTCCG
AAGACGTTATCAAAGAGTTCATGCGTTTCAAAGTTCGTATGGAAGGTTCCGTTAACGGTCACGAGTTCGAAATCGAAGGTGAAGGTGAAGGTCGTCCGTA
CGAAGGTACCCAGACCGCTAAACTGAAAGTTACCAAAGGTGGTCCGCTGCCGTTCGCTTGGGACATCCTGTCCCCGCAGTTCCAGTACGGTTCCAAAGCT
TACGTTAAACACCCGGCTGACATCCCGGACTACCTGAAACTGTCCTTCCCGGAAGGTTTCAAATGGGAACGTGTTATGAACTTCGAAGACGGTGGTGTTG
TTACCGTTACCCAGGACTCCTCCCTGCAAGACGGTGAGTTCATCTACAAAGTTAAACTGCGTGGTACCAACTTCCCGTCCGACGGTCCGGTTATGCAGAA
AAAAACCATGGGTTGGGAAGCTTCCACCGAACGTATGTACCCGGAAGACGGTGCTCTGAAAGGTGAAATCAAAATGCGTCTGAAACTGAAAGACGGTGGT
CACTACGACGCTGAAGTTAAAACCACCTACATGGCTAAAAAACCGGTTCAGCTGCCGGGTGCTTACAAAACCGACATCAAACTGGACATCACCTCCCACA
ACGAAGACTACACCATCGTTGAACAGTACGAACGTGCTGAAGGTCGTCACTCCACCGGTGCTTAATAACGCTGATAGTGCTAGTGTAGATCGCTACTAGA
GTACTAGCTCTAGATGCGGCCGCGAATTCCAGAAATCATCCTTAGCGAAAGCTAAGGATTTTTTTTATCTGAAATGCCTCTGTAACAGAGCATTAGCGCA
AGGTGATTTTTGTCTTCTTGCGCTAATTTTTTGGCGTCTCCACTGTGCGCGGAACCCCTATTTGTTTATTTTTCTAAATACATTCAAATATGTATCCGCT
CATGAGACAATAACCCTGATAAATGCTTCAATAATATTGAAAAAGGAAGAGTATGAGTATTCAACATTTCCGTGTCGCCCTTATTCCCTTTTTTGCGGCA
TTTTGCCTTCCTGTTTTTGCTCACCCAGAAACGCTGGTGAAAGTAAAAGATGCTGAAGATCAGTTGGGTGCACGAGTGGGTTACATCGAACTGGATCTCA
ACAGCGGTAAGATCCTTGAGAGTTTTCGCCCCGAAGAACGTTTTCCAATGATGAGCACTTTTAAAGTTCTGCTATGTGGCGCGGTATTATCCCGTATTGA
CGCCGGGCAAGAGCAACTCGGTCGCCGCATACACTATTCTCAGAATGACTTGGTTGAGTACTCACCAGTCACAGAAAAGCATCTTACGGATGGCATGACA
GTAAGAGAATTATGCAGTGCTGCCATAACCATGAGTGATAACACTGCGGCCAACTTACTTCTGACAACGATCGGAGGACCGAAGGAGCTAACCGCTTTTT
TGCACAACATGGGGGATCATGTAACTCGCCTTGATCGTTGGGAACCGGAGCTGAATGAAGCCATACCAAACGACGAGCGTGACACCACGATGCCTGTAGC
AATGGCAACAACGTTGCGCAAACTATTAACTGGCGAACTACTTACTCTAGCTTCCCGGCAACAATTAATAGACTGGATGGAGGCGGATAAAGTTGCAGGA
CCACTTCTGCGCTCGGCCCTTCCGGCTGGCTGGTTTATTGCTGATAAATCTGGAGCCGGTGAGCGTGGGTCTCGCGGTATCATTGCAGCACTGGGGCCAG
ATGGTAAGCCCTCCCGTATCGTAGTTATCTACACGACGGGGAGTCAGGCAACTATGGATGAACGAAATAGACAGATCGCTGAGATAGGTGCCTCACTGAT
TAAGCATTGGTAACTCGAGCTGATCCTTCAAGAGCTCGCTTGGACTCCTGTTGATAGATCCAGTAATGACCTCAGAACTCCATCTGGATTTGTTCAGAAC
GCTCGGTTGCCGCCGGGCGTTTTTTATTGGTGAGAATTTACAACTTATATCGTATGGGGCTGACTTCAGGTGCTACATTTGAAGAGATAAATTGCACTGA
AATCTAGAAATATTTTATCTGATTAATAAGATGATCTTCTTGAGATCGTTTTGGTCTGCGCGTAATCTCTTGCTCTGAAAACGAAAAAACCGCCTTGCAG
GGCGGTTTTTCGAAGGTTCTCTGAGCTACCAACTCTTTGAACCGAGGTAACTGGCTTGGAGGAGCGCAGTCACCAAAACTTGTCCTTTCAGTTTAGCCTT
AACCGGCGCATGACTTCAAGACTAACTCCTCTAAATCAATTACCAGTGGCTGCTGCCAGTGGTGCTTTTGCATGTCTTTCCGGGTTGGACTCAAGACGAT
AGTTACCGGATAAGGCGCAGCGGTCGGACTGAACGGGGGGTTCGTGCATACAGTCCAGCTTGGAGCGAACTGCCTACCCGGAACTGAGTGTCAGGCGTGG
AATGAGACAAACGCGGCCATAACAGCGGAATGACACCGGTAAACCGAAAGGCAGGAACAGGAGAGCGCACGAGGGAGCCGCCAGGGGGAAACGCCTGGTA
TCTTTATAGTCCTGTCGGGTTTCGCCACCACTGATTTGAGCGTCAGATT

**pCM190**

TCGTGATGCTTGTCAGGGGGGCGGAGCCTATGGAAAAACGGCTTTGCCGCGGCCCTCTCACTTCCCTGTTAAGTATCTTCCTGGCATCTTCCAGGAAATC
TCCGCCCCGTTCGTAAGCCATTTCCGCTCGCCGCAGTCGAACGGGCGCGCCTTGACAGCTAGCTCAGTCCTAGGTATTGTGCTAGCTTACGAGCGATGGG
CTCACTGTTGAGCCTCAATGAGCGAACTGATCAAAGAGAACATGCACATGAAGCTGTACATGGAGGGTACCGTGGATAATCACCACTTTAAGTGTACTTC
TGAGGGCGAGGGTAAGCCGTATGAAGGGACTCAAACGATGCGTATTAAAGTAGTGGAGGGTGGCCCACTGCCGTTTGCTTTCGATATTCTGGCGACGAGC
TTTCTGTATGGTAGCAAAACGTTTATAAACCACACTCAGGGCATTCCGGATTTCTTTAAACAAAGCTTTCCGGAAGGTTTTACCTGGGAGCGTGTGACTA
CGTATGAAGATGGTGGTGTGTTGACTGCTACTCAAGATACTTCACTGCAGGACGGCTGTCTGATCTATAACGTGAAGATTCGTGGCGTGAACTTTACGAG
CAATGGGCCGGTAATGCAAAAAAAAACCCTGGGTTGGGAAGCGTTCACGGAAACTCTGTATCCGGCTGACGGCGGCCTGGAGGGCCGTAACGATATGGCA
CTGAAGCTGGTTGGTGGCAGCCACCTGATCGCGAATATCAAAACGACTTATCGCTCTAAAAAACCGGCGAAAAATCTGAAGATGCCGGGTGTTTATTATG
TTGACTATCGTCTGGAACGCATTAAAGAAGCGAATAATGAAACTTACGTGGAGCAACACGAGGTTGCAGTGGCGCGCTATTGCGACTTGCCTTCAAAGCT
GGGTCACAAACTGAATTAATAAGGAAACACAGAAAAAAGCCCGCACCTGACAGTGCGGGCTTTTTTTTTCGACCAAAGGGACAATGTCCTGAAGAATATT
TAAGAAAAAAGCACCCCTCGTCGCCTAGAATTACCTACCGCGGTCGACCATACCTTCGAGACGAACAATAAGGCCGCAAATCGCGGCCTTCGTTATTGAT
AACAGCAGGTAAAAAGACGCTTATTATGGAGTTCGACATGGCAATAACGCGACGAATCTACGTCATGACGAGAATAGTATAAACAAAACTGCTGACGGCA
GAAGCGTCTCGGTACCAAATTCCAGAAAAGAGACGCTTTCGAGCGTCTTTTTTCGTTTTGGTCCACGCTGATGAATATTCTACGGAATTGCCATAGGCGT
TGAACGCTACACGGACGATACGAATTTATGTATAGAGCGTTCAGCCAAAAAACTTAAGACCGCCGGTCTTGTCCACTACCTTGCAGTAATGCGGTGGACA
GGATCGGCGGTTTTCTTTTCTCTTCTCAATTCGCGACCCAGGATGAGGTCGCCAAAACTACTAGATTTGGGCGCGCCTTGACGGCTAGCTCAGTCCTAGG
TACAGTGCTAGCTTAATCTTGTCAAACAGTGGCTACGTGAACAGATGCGTAAAGGAGAAGAACTGTTCACTGGAGTTGTCCCAATTCTTGTTGAATTAGA
TGGTGATGTTAATGGGCACAAATTTTCTGTGAGTGGAGAGGGTGAAGGTGATGCGACATACGGCAAACTGACCCTTAAATTTATTTGTACTACCGGCAAG
CTACCTGTTCCTTGGCCGACACTTGTCACTACTTTCGGTTATGGTGTTCAATGTTTTGCGAGATACCCAGATCATATGAAACAGCATGACTTTTTCAAAA
GTGCCATGCCCGAAGGTTATGTACAGGAAAGAACTATATTTTTCAAAGATGACGGGAACTACAAAACACGTGCTGAAGTTAAATTTGAGGGTGATACTCT
GGTAAATAGAATCGAATTAAAAGGTATTGATTTTAAAGAAGATGGCAACATTCTTGGTCACAAGCTGGAATACAACTACAACTCACACAATGTTTACATC
ATGGCAGACAAACAAAAAAATGGCATCAAAGTTAACTTCAAAATTCGACACAACATTGAAGATGGAAGCGTTCAACTGGCAGACCATTATCAACAGAATA
CTCCGATTGGCGATGGCCCTGTACTCTTACCAGACAACCATTACCTGTCCACACAATCTGCCCTTTCGAAAGATCCGAACGAAAAGAGAGATCACATGGT

CCTGCTTGAGTTTGTAACAGCTGCTGGGATTACACATGGTATGGATGAGCTATACAAATAATAACCAATTATTGAAGGCCTCCCTAACGGGGGGCCTTTT
TTTGTTTCTGGTCTCCCCACTGATAGTGCTAGTGTAGATCACTACTAGAGCCAGGCATCAAATAAAACGAAAGGCTCAGTCGAAAGACTGGGCCTTTCGT
TTTATCTGTTGTTTGTCGGTGAACGCTCTCTACTAGAGTCACACTGGCTCACCTTCGGGTGGGCCTTTCTGCGTTTATATACTAGAGGGAATCGTACGGT
CGATCCATCTGACTATCGCCTACTTCATTGACAGCTAGCTCAGTCCTAGGTATAATGCTAGCTCTAGAGAAAGAGGAGAAATACTAGATGGCTTCCTCCG
AAGACGTTATCAAAGAGTTCATGCGTTTCAAAGTTCGTATGGAAGGTTCCGTTAACGGTCACGAGTTCGAAATCGAAGGTGAAGGTGAAGGTCGTCCGTA
CGAAGGTACCCAGACCGCTAAACTGAAAGTTACCAAAGGTGGTCCGCTGCCGTTCGCTTGGGACATCCTGTCCCCGCAGTTCCAGTACGGTTCCAAAGCT
TACGTTAAACACCCGGCTGACATCCCGGACTACCTGAAACTGTCCTTCCCGGAAGGTTTCAAATGGGAACGTGTTATGAACTTCGAAGACGGTGGTGTTG
TTACCGTTACCCAGGACTCCTCCCTGCAAGACGGTGAGTTCATCTACAAAGTTAAACTGCGTGGTACCAACTTCCCGTCCGACGGTCCGGTTATGCAGAA
AAAAACCATGGGTTGGGAAGCTTCCACCGAACGTATGTACCCGGAAGACGGTGCTCTGAAAGGTGAAATCAAAATGCGTCTGAAACTGAAAGACGGTGGT
CACTACGACGCTGAAGTTAAAACCACCTACATGGCTAAAAAACCGGTTCAGCTGCCGGGTGCTTACAAAACCGACATCAAACTGGACATCACCTCCCACA
ACGAAGACTACACCATCGTTGAACAGTACGAACGTGCTGAAGGTCGTCACTCCACCGGTGCTTAATAACGCTGATAGTGCTAGTGTAGATCGCTACTAGA
GTACTAGCTCTAGATGCGGCCGCGAATTCCAGAAATCATCCTTAGCGAAAGCTAAGGATTTTTTTTATCTGAAATGCCTCTGTAACAGAGCATTAGCGCA
AGGTGATTTTTGTCTTCTTGCGCTAATTTTTTGGCGTCTCCACTGTGCGCGGAACCCCTATTTGTTTATTTTTCTAAATACATTCAAATATGTATCCGCT
CATGAGACAATAACCCTGATAAATGCTTCAATAATATTGAAAAAGGAAGAGTATGAGTATTCAACATTTCCGTGTCGCCCTTATTCCCTTTTTTGCGGCA
TTTTGCCTTCCTGTTTTTGCTCACCCAGAAACGCTGGTGAAAGTAAAAGATGCTGAAGATCAGTTGGGTGCACGAGTGGGTTACATCGAACTGGATCTCA
ACAGCGGTAAGATCCTTGAGAGTTTTCGCCCCGAAGAACGTTTTCCAATGATGAGCACTTTTAAAGTTCTGCTATGTGGCGCGGTATTATCCCGTATTGA
CGCCGGGCAAGAGCAACTCGGTCGCCGCATACACTATTCTCAGAATGACTTGGTTGAGTACTCACCAGTCACAGAAAAGCATCTTACGGATGGCATGACA
GTAAGAGAATTATGCAGTGCTGCCATAACCATGAGTGATAACACTGCGGCCAACTTACTTCTGACAACGATCGGAGGACCGAAGGAGCTAACCGCTTTTT
TGCACAACATGGGGGATCATGTAACTCGCCTTGATCGTTGGGAACCGGAGCTGAATGAAGCCATACCAAACGACGAGCGTGACACCACGATGCCTGTAGC
AATGGCAACAACGTTGCGCAAACTATTAACTGGCGAACTACTTACTCTAGCTTCCCGGCAACAATTAATAGACTGGATGGAGGCGGATAAAGTTGCAGGA
CCACTTCTGCGCTCGGCCCTTCCGGCTGGCTGGTTTATTGCTGATAAATCTGGAGCCGGTGAGCGTGGGTCTCGCGGTATCATTGCAGCACTGGGGCCAG
ATGGTAAGCCCTCCCGTATCGTAGTTATCTACACGACGGGGAGTCAGGCAACTATGGATGAACGAAATAGACAGATCGCTGAGATAGGTGCCTCACTGAT
TAAGCATTGGTAACTCGAGCTGATCCTTCAAGAGCTCGCTTGGACTCCTGTTGATAGATCCAGTAATGACCTCAGAACTCCATCTGGATTTGTTCAGAAC
GCTCGGTTGCCGCCGGGCGTTTTTTATTGGTGAGAATTTACAACTTATATCGTATGGGGCTGACTTCAGGTGCTACATTTGAAGAGATAAATTGCACTGA
AATCTAGAAATATTTTATCTGATTAATAAGATGATCTTCTTGAGATCGTTTTGGTCTGCGCGTAATCTCTTGCTCTGAAAACGAAAAAACCGCCTTGCAG
GGCGGTTTTTCGAAGGTTCTCTGAGCTACCAACTCTTTGAACCGAGGTAACTGGCTTGGAGGAGCGCAGTCACCAAAACTTGTCCTTTCAGTTTAGCCTT
AACCGGCGCATGACTTCAAGACTAACTCCTCTAAATCAATTACCAGTGGCTGCTGCCAGTGGTGCTTTTGCATGTCTTTCCGGGTTGGACTCAAGACGAT
AGTTACCGGATAAGGCGCAGCGGTCGGACTGAACGGGGGGTTCGTGCATACAGTCCAGCTTGGAGCGAACTGCCTACCCGGAACTGAGTGTCAGGCGTGG
AATGAGACAAACGCGGCCATAACAGCGGAATGACACCGGTAAACCGAAAGGCAGGAACAGGAGAGCGCACGAGGGAGCCGCCAGGGGGAAACGCCTGGTA
TCTTTATAGTCCTGTCGGGTTTCGCCACCACTGATTTGAGCGTCAGATT

**pCM191**

TCGTGATGCTTGTCAGGGGGGCGGAGCCTATGGAAAAACGGCTTTGCCGCGGCCCTCTCACTTCCCTGTTAAGTATCTTCCTGGCATCTTCCAGGAAATC
TCCGCCCCGTTCGTAAGCCATTTCCGCTCGCCGCAGTCGAACGGGCGCGCCTTGACAGCTAGCTCAGTCCTAGGTATTGTGCTAGCTTACGAGCGATGGG
CTCACTGTTGAGCCTCAATGAGCGAACTGATCAAAGAGAACATGCACATGAAGCTGTACATGGAGGGTACCGTGGATAATCACCACTTTAAGTGTACTTC
TGAGGGCGAGGGTAAGCCGTATGAAGGGACTCAAACGATGCGTATTAAAGTAGTGGAGGGTGGCCCACTGCCGTTTGCTTTCGATATTCTGGCGACGAGC
TTTCTGTATGGTAGCAAAACGTTTATAAACCACACTCAGGGCATTCCGGATTTCTTTAAACAAAGCTTTCCGGAAGGTTTTACCTGGGAGCGTGTGACTA
CGTATGAAGATGGTGGTGTGTTGACTGCTACTCAAGATACTTCACTGCAGGACGGCTGTCTGATCTATAACGTGAAGATTCGTGGCGTGAACTTTACGAG
CAATGGGCCGGTAATGCAAAAAAAAACCCTGGGTTGGGAAGCGTTCACGGAAACTCTGTATCCGGCTGACGGCGGCCTGGAGGGCCGTAACGATATGGCA
CTGAAGCTGGTTGGTGGCAGCCACCTGATCGCGAATATCAAAACGACTTATCGCTCTAAAAAACCGGCGAAAAATCTGAAGATGCCGGGTGTTTATTATG
TTGACTATCGTCTGGAACGCATTAAAGAAGCGAATAATGAAACTTACGTGGAGCAACACGAGGTTGCAGTGGCGCGCTATTGCGACTTGCCTTCAAAGCT
GGGTCACAAACTGAATTAATAAGGAAACACAGAAAAAAGCCCGCACCTGACAGTGCGGGCTTTTTTTTTCGACCAAAGGGACAATGTCCTGAAGAATATT
TAAGAAAAAAGCACCCCTCGTCGCCTAGAATTACCTACCGCGGTCGACCATACCTTCGAGACGAACAATAAGGCCGCAAATCGCGGCCTTCGTTATTGAT
AACAGCAGGTAAAAAGACGCTTATTATGGAGTTCGACATGGCAATAACGCGACGAATCTACGTCATGACGAGAATAGTATAAACAAAACTGCTGACGGCA
GAAGCGTCTCGGTACCAAATTCCAGAAAAGAGACGCTTTCGAGCGTCTTTTTTCGTTTTGGTCCACGCTGATGAATATTCTACGGAATTGCCATAGGCGT
TGAACGCTACACGGACGATACGAATTTATGTATAGAGCGTTCAGCCAAAAAACTTAAGACCGCCGGTCTTGTCCACTACCTTGCAGTAATGCGGTGGACA
GGATCGGCGGTTTTCTTTTCTCTTCTCAATTCGCGACCCAGGATGAGGTCGCCAAAACTACTAGATTTGGGCGCGCCTTGACGGCTAGCTCAGTCCTAGG
TACAGTGCTAGCTTAATCTTGTCAAACAGTGGCTACGTGAACAGATGCGTAAAGGAGAAGAACTGTTCACTGGAGTTGTCCCAATTCTTGTTGAATTAGA
TGGTGATGTTAATGGGCACAAATTTTCTGTGAGTGGAGAGGGTGAAGGTGATGCGACATACGGCAAACTGACCCTTAAATTTATTTGTACTACCGGCAAG
CTACCTGTTCCTTGGCCGACACTTGTCACTACTTTCGGTTATGGTGTTCAATGTTTTGCGAGATACCCAGATCATATGAAACAGCATGACTTTTTCAAAA
GTGCCATGCCCGAAGGTTATGTACAGGAAAGAACTATATTTTTCAAAGATGACGGGAACTACAAAACACGTGCTGAAGTTAAATTTGAGGGTGATACTCT
GGTAAATAGAATCGAATTAAAAGGTATTGATTTTAAAGAAGATGGCAACATTCTTGGTCACAAGCTGGAATACAACTACAACTCACACAATGTTTACATC
ATGGCAGACAAACAAAAAAATGGCATCAAAGTTAACTTCAAAATTCGACACAACATTGAAGATGGAAGCGTTCAACTGGCAGACCATTATCAACAGAATA
CTCCGATTGGCGATGGCCCTGTACTCTTACCAGACAACCATTACCTGTCCACACAATCTGCCCTTTCGAAAGATCCGAACGAAAAGAGAGATCACATGGT
CCTGCTTGAGTTTGTAACAGCTGCTGGGATTACACATGGTATGGATGAGCTATACAAATAATAACCAATTATTGAAGGCCTCCCTAACGGGGGGCCTTTT
TTTGTTTCTGGTCTCCCCACTGATAGTGCTAGTGTAGATCACTACTAGAGCCAGGCATCAAATAAAACGAAAGGCTCAGTCGAAAGACTGGGCCTTTCGT
TTTATCTGTTGTTTGTCGGTGAACGCTCTCTACTAGAGTCACACTGGCTCACCTTCGGGTGGGCCTTTCTGCGTTTATATACTAGAGGGAATCGTACGGT
CGATCCATCTGACTATCGCCTACTTCATTGACAGCTAGCTCAGTCCTAGGTATAATGCTAGCCTAATTCTACATAGAGTGAGGTAATCTAATGGCTTCCT
CCGAAGACGTTATCAAAGAGTTCATGCGTTTCAAAGTTCGTATGGAAGGTTCCGTTAACGGTCACGAGTTCGAAATCGAAGGTGAAGGTGAAGGTCGTCC
GTACGAAGGTACCCAGACCGCTAAACTGAAAGTTACCAAAGGTGGTCCGCTGCCGTTCGCTTGGGACATCCTGTCCCCGCAGTTCCAGTACGGTTCCAAA
GCTTACGTTAAACACCCGGCTGACATCCCGGACTACCTGAAACTGTCCTTCCCGGAAGGTTTCAAATGGGAACGTGTTATGAACTTCGAAGACGGTGGTG
TTGTTACCGTTACCCAGGACTCCTCCCTGCAAGACGGTGAGTTCATCTACAAAGTTAAACTGCGTGGTACCAACTTCCCGTCCGACGGTCCGGTTATGCA
GAAAAAAACCATGGGTTGGGAAGCTTCCACCGAACGTATGTACCCGGAAGACGGTGCTCTGAAAGGTGAAATCAAAATGCGTCTGAAACTGAAAGACGGT
GGTCACTACGACGCTGAAGTTAAAACCACCTACATGGCTAAAAAACCGGTTCAGCTGCCGGGTGCTTACAAAACCGACATCAAACTGGACATCACCTCCC
ACAACGAAGACTACACCATCGTTGAACAGTACGAACGTGCTGAAGGTCGTCACTCCACCGGTGCTTAATAACGCTGATAGTGCTAGTGTAGATCGCTACT
AGAGTACTAGCTCTAGATGCGGCCGCGAATTCCAGAAATCATCCTTAGCGAAAGCTAAGGATTTTTTTTATCTGAAATGCCTCTGTAACAGAGCATTAGC
GCAAGGTGATTTTTGTCTTCTTGCGCTAATTTTTTGGCGTCTCCACTGTGCGCGGAACCCCTATTTGTTTATTTTTCTAAATACATTCAAATATGTATCC
GCTCATGAGACAATAACCCTGATAAATGCTTCAATAATATTGAAAAAGGAAGAGTATGAGTATTCAACATTTCCGTGTCGCCCTTATTCCCTTTTTTGCG
GCATTTTGCCTTCCTGTTTTTGCTCACCCAGAAACGCTGGTGAAAGTAAAAGATGCTGAAGATCAGTTGGGTGCACGAGTGGGTTACATCGAACTGGATC
TCAACAGCGGTAAGATCCTTGAGAGTTTTCGCCCCGAAGAACGTTTTCCAATGATGAGCACTTTTAAAGTTCTGCTATGTGGCGCGGTATTATCCCGTAT
TGACGCCGGGCAAGAGCAACTCGGTCGCCGCATACACTATTCTCAGAATGACTTGGTTGAGTACTCACCAGTCACAGAAAAGCATCTTACGGATGGCATG
ACAGTAAGAGAATTATGCAGTGCTGCCATAACCATGAGTGATAACACTGCGGCCAACTTACTTCTGACAACGATCGGAGGACCGAAGGAGCTAACCGCTT
TTTTGCACAACATGGGGGATCATGTAACTCGCCTTGATCGTTGGGAACCGGAGCTGAATGAAGCCATACCAAACGACGAGCGTGACACCACGATGCCTGT
AGCAATGGCAACAACGTTGCGCAAACTATTAACTGGCGAACTACTTACTCTAGCTTCCCGGCAACAATTAATAGACTGGATGGAGGCGGATAAAGTTGCA
GGACCACTTCTGCGCTCGGCCCTTCCGGCTGGCTGGTTTATTGCTGATAAATCTGGAGCCGGTGAGCGTGGGTCTCGCGGTATCATTGCAGCACTGGGGC
CAGATGGTAAGCCCTCCCGTATCGTAGTTATCTACACGACGGGGAGTCAGGCAACTATGGATGAACGAAATAGACAGATCGCTGAGATAGGTGCCTCACT
GATTAAGCATTGGTAACTCGAGCTGATCCTTCAAGAGCTCGCTTGGACTCCTGTTGATAGATCCAGTAATGACCTCAGAACTCCATCTGGATTTGTTCAG
AACGCTCGGTTGCCGCCGGGCGTTTTTTATTGGTGAGAATTTACAACTTATATCGTATGGGGCTGACTTCAGGTGCTACATTTGAAGAGATAAATTGCAC
TGAAATCTAGAAATATTTTATCTGATTAATAAGATGATCTTCTTGAGATCGTTTTGGTCTGCGCGTAATCTCTTGCTCTGAAAACGAAAAAACCGCCTTG
CAGGGCGGTTTTTCGAAGGTTCTCTGAGCTACCAACTCTTTGAACCGAGGTAACTGGCTTGGAGGAGCGCAGTCACCAAAACTTGTCCTTTCAGTTTAGC

CTTAACCGGCGCATGACTTCAAGACTAACTCCTCTAAATCAATTACCAGTGGCTGCTGCCAGTGGTGCTTTTGCATGTCTTTCCGGGTTGGACTCAAGAC
GATAGTTACCGGATAAGGCGCAGCGGTCGGACTGAACGGGGGGTTCGTGCATACAGTCCAGCTTGGAGCGAACTGCCTACCCGGAACTGAGTGTCAGGCG
TGGAATGAGACAAACGCGGCCATAACAGCGGAATGACACCGGTAAACCGAAAGGCAGGAACAGGAGAGCGCACGAGGGAGCCGCCAGGGGGAAACGCCTG
GTATCTTTATAGTCCTGTCGGGTTTCGCCACCACTGATTTGAGCGTCAGATT

**pCM200**

TCGTGATGCTTGTCAGGGGGGCGGAGCCTATGGAAAAACGGCTTTGCCGCGGCCCTCTCACTTCCCTGTTAAGTATCTTCCTGGCATCTTCCAGGAAATC
TCCGCCCCGTTCGTAAGCCATTTCCGCTCGCCGCAGTCGAACGGGCGCGCCTTGACAGCTAGCTCAGTCCTAGGTATTGTGCTAGCTTACGTCAACCCAC
CCTAGGAGAGGATAAGATGAGCGAACTGATCAAAGAGAACATGCACATGAAGCTGTACATGGAGGGTACCGTGGATAATCACCACTTTAAGTGTACTTCT
GAGGGCGAGGGTAAGCCGTATGAAGGGACTCAAACGATGCGTATTAAAGTAGTGGAGGGTGGCCCACTGCCGTTTGCTTTCGATATTCTGGCGACGAGCT
TTCTGTATGGTAGCAAAACGTTTATAAACCACACTCAGGGCATTCCGGATTTCTTTAAACAAAGCTTTCCGGAAGGTTTTACCTGGGAGCGTGTGACTAC
GTATGAAGATGGTGGTGTGTTGACTGCTACTCAAGATACTTCACTGCAGGACGGCTGTCTGATCTATAACGTGAAGATTCGTGGCGTGAACTTTACGAGC
AATGGGCCGGTAATGCAAAAAAAAACCCTGGGTTGGGAAGCGTTCACGGAAACTCTGTATCCGGCTGACGGCGGCCTGGAGGGCCGTAACGATATGGCAC
TGAAGCTGGTTGGTGGCAGCCACCTGATCGCGAATATCAAAACGACTTATCGCTCTAAAAAACCGGCGAAAAATCTGAAGATGCCGGGTGTTTATTATGT
TGACTATCGTCTGGAACGCATTAAAGAAGCGAATAATGAAACTTACGTGGAGCAACACGAGGTTGCAGTGGCGCGCTATTGCGACTTGCCTTCAAAGCTG
GGTCACAAACTGAATTAATAAGGAAACACAGAAAAAAGCCCGCACCTGACAGTGCGGGCTTTTTTTTTCGACCAAAGGGACAATGTCCTGAAGAATATTT
AAGAAAAAAGCACCCCTCGTCGCCTAGAATTACCTACCGCGGTCGACCATACCTTCGAGACGAACAATAAGGCCGCAAATCGCGGCCTTCGTTATTGATA
ACAGCAGGTAAAAAGACGCTTATTATGGAGTTCGACATGGCAATAACGCGACGAATCTACGTCATGACGAGAATAGTATAAACAAAACTGCTGACGGCAG
AAGCGTCTCGGTACCAAATTCCAGAAAAGAGACGCTTTCGAGCGTCTTTTTTCGTTTTGGTCCACGCTGATGAATATTCTACGGAATTGCCATAGGCGTT
GAACGCTACACGGACGATACGAATTTATGTATAGAGCGTTCAGCCAAAAAACTTAAGACCGCCGGTCTTGTCCACTACCTTGCAGTAATGCGGTGGACAG
GATCGGCGGTTTTCTTTTCTCTTCTCAATTCGCGACCCAGGATGAGGTCGCCAAAACTACTAGATTTGGGCGCGCCTTGACGGCTAGCTCAGTCCTAGGT
ACAGTGCTAGCTTAATGCGAGAGGCTGGTTAAGTGGGTGTTGATGCGTAAAGGAGAAGAACTGTTCACTGGAGTTGTCCCAATTCTTGTTGAATTAGATG
GTGATGTTAATGGGCACAAATTTTCTGTGAGTGGAGAGGGTGAAGGTGATGCGACATACGGCAAACTGACCCTTAAATTTATTTGTACTACCGGCAAGCT
ACCTGTTCCTTGGCCGACACTTGTCACTACTTTCGGTTATGGTGTTCAATGTTTTGCGAGATACCCAGATCATATGAAACAGCATGACTTTTTCAAAAGT
GCCATGCCCGAAGGTTATGTACAGGAAAGAACTATATTTTTCAAAGATGACGGGAACTACAAAACACGTGCTGAAGTTAAATTTGAGGGTGATACTCTGG
TAAATAGAATCGAATTAAAAGGTATTGATTTTAAAGAAGATGGCAACATTCTTGGTCACAAGCTGGAATACAACTACAACTCACACAATGTTTACATCAT
GGCAGACAAACAAAAAAATGGCATCAAAGTTAACTTCAAAATTCGACACAACATTGAAGATGGAAGCGTTCAACTGGCAGACCATTATCAACAGAATACT
CCGATTGGCGATGGCCCTGTACTCTTACCAGACAACCATTACCTGTCCACACAATCTGCCCTTTCGAAAGATCCGAACGAAAAGAGAGATCACATGGTCC
TGCTTGAGTTTGTAACAGCTGCTGGGATTACACATGGTATGGATGAGCTATACAAATAATAACCAATTATTGAAGGCCTCCCTAACGGGGGGCCTTTTTT
TGTTTCTGGTCTCCCCACTGATAGTGCTAGTGTAGATCACTACTAGAGCCAGGCATCAAATAAAACGAAAGGCTCAGTCGAAAGACTGGGCCTTTCGTTT
TATCTGTTGTTTGTCGGTGAACGCTCTCTACTAGAGTCACACTGGCTCACCTTCGGGTGGGCCTTTCTGCGTTTATATACTAGAGGGAATCGTACGGTCG
ATCCATCTGACTATCGCCTACTTCATTGACAGCTAGCTCAGTCCTAGGTATAATGCTAGCAAATATGTTGTAGAAACAATTGCACGAATGGCTTCCTCCG
AAGACGTTATCAAAGAGTTCATGCGTTTCAAAGTTCGTATGGAAGGTTCCGTTAACGGTCACGAGTTCGAAATCGAAGGTGAAGGTGAAGGTCGTCCGTA
CGAAGGTACCCAGACCGCTAAACTGAAAGTTACCAAAGGTGGTCCGCTGCCGTTCGCTTGGGACATCCTGTCCCCGCAGTTCCAGTACGGTTCCAAAGCT
TACGTTAAACACCCGGCTGACATCCCGGACTACCTGAAACTGTCCTTCCCGGAAGGTTTCAAATGGGAACGTGTTATGAACTTCGAAGACGGTGGTGTTG
TTACCGTTACCCAGGACTCCTCCCTGCAAGACGGTGAGTTCATCTACAAAGTTAAACTGCGTGGTACCAACTTCCCGTCCGACGGTCCGGTTATGCAGAA
AAAAACCATGGGTTGGGAAGCTTCCACCGAACGTATGTACCCGGAAGACGGTGCTCTGAAAGGTGAAATCAAAATGCGTCTGAAACTGAAAGACGGTGGT
CACTACGACGCTGAAGTTAAAACCACCTACATGGCTAAAAAACCGGTTCAGCTGCCGGGTGCTTACAAAACCGACATCAAACTGGACATCACCTCCCACA
ACGAAGACTACACCATCGTTGAACAGTACGAACGTGCTGAAGGTCGTCACTCCACCGGTGCTTAATAACGCTGATAGTGCTAGTGTAGATCGCTACTAGA
GTACTAGCTCTAGATGCGGCCGCGAATTCCAGAAATCATCCTTAGCGAAAGCTAAGGATTTTTTTTATCTGAAATGCCTCTGTAACAGAGCATTAGCGCA
AGGTGATTTTTGTCTTCTTGCGCTAATTTTTTGGCGTCTCCACTGTGCGCGGAACCCCTATTTGTTTATTTTTCTAAATACATTCAAATATGTATCCGCT
CATGAGACAATAACCCTGATAAATGCTTCAATAATATTGAAAAAGGAAGAGTATGAGTATTCAACATTTCCGTGTCGCCCTTATTCCCTTTTTTGCGGCA
TTTTGCCTTCCTGTTTTTGCTCACCCAGAAACGCTGGTGAAAGTAAAAGATGCTGAAGATCAGTTGGGTGCACGAGTGGGTTACATCGAACTGGATCTCA
ACAGCGGTAAGATCCTTGAGAGTTTTCGCCCCGAAGAACGTTTTCCAATGATGAGCACTTTTAAAGTTCTGCTATGTGGCGCGGTATTATCCCGTATTGA
CGCCGGGCAAGAGCAACTCGGTCGCCGCATACACTATTCTCAGAATGACTTGGTTGAGTACTCACCAGTCACAGAAAAGCATCTTACGGATGGCATGACA
GTAAGAGAATTATGCAGTGCTGCCATAACCATGAGTGATAACACTGCGGCCAACTTACTTCTGACAACGATCGGAGGACCGAAGGAGCTAACCGCTTTTT
TGCACAACATGGGGGATCATGTAACTCGCCTTGATCGTTGGGAACCGGAGCTGAATGAAGCCATACCAAACGACGAGCGTGACACCACGATGCCTGTAGC
AATGGCAACAACGTTGCGCAAACTATTAACTGGCGAACTACTTACTCTAGCTTCCCGGCAACAATTAATAGACTGGATGGAGGCGGATAAAGTTGCAGGA
CCACTTCTGCGCTCGGCCCTTCCGGCTGGCTGGTTTATTGCTGATAAATCTGGAGCCGGTGAGCGTGGGTCTCGCGGTATCATTGCAGCACTGGGGCCAG
ATGGTAAGCCCTCCCGTATCGTAGTTATCTACACGACGGGGAGTCAGGCAACTATGGATGAACGAAATAGACAGATCGCTGAGATAGGTGCCTCACTGAT
TAAGCATTGGTAACTCGAGCTGATCCTTCAAGAGCTCGCTTGGACTCCTGTTGATAGATCCAGTAATGACCTCAGAACTCCATCTGGATTTGTTCAGAAC
GCTCGGTTGCCGCCGGGCGTTTTTTATTGGTGAGAATTTACAACTTATATCGTATGGGGCTGACTTCAGGTGCTACATTTGAAGAGATAAATTGCACTGA
AATCTAGAAATATTTTATCTGATTAATAAGATGATCTTCTTGAGATCGTTTTGGTCTGCGCGTAATCTCTTGCTCTGAAAACGAAAAAACCGCCTTGCAG
GGCGGTTTTTCGAAGGTTCTCTGAGCTACCAACTCTTTGAACCGAGGTAACTGGCTTGGAGGAGCGCAGTCACCAAAACTTGTCCTTTCAGTTTAGCCTT
AACCGGCGCATGACTTCAAGACTAACTCCTCTAAATCAATTACCAGTGGCTGCTGCCAGTGGTGCTTTTGCATGTCTTTCCGGGTTGGACTCAAGACGAT
AGTTACCGGATAAGGCGCAGCGGTCGGACTGAACGGGGGGTTCGTGCATACAGTCCAGCTTGGAGCGAACTGCCTACCCGGAACTGAGTGTCAGGCGTGG
AATGAGACAAACGCGGCCATAACAGCGGAATGACACCGGTAAACCGAAAGGCAGGAACAGGAGAGCGCACGAGGGAGCCGCCAGGGGGAAACGCCTGGTA
TCTTTATAGTCCTGTCGGGTTTCGCCACCACTGATTTGAGCGTCAGATT

**pCM201**

TCGTGATGCTTGTCAGGGGGGCGGAGCCTATGGAAAAACGGCTTTGCCGCGGCCCTCTCACTTCCCTGTTAAGTATCTTCCTGGCATCTTCCAGGAAATC
TCCGCCCCGTTCGTAAGCCATTTCCGCTCGCCGCAGTCGAACGGGCGCGCCTTGACAGCTAGCTCAGTCCTAGGTATTGTGCTAGCTTACGCTAATTCTA
CATAGAGTGAGGTAATCTAATGAGCGAACTGATCAAAGAGAACATGCACATGAAGCTGTACATGGAGGGTACCGTGGATAATCACCACTTTAAGTGTACT
TCTGAGGGCGAGGGTAAGCCGTATGAAGGGACTCAAACGATGCGTATTAAAGTAGTGGAGGGTGGCCCACTGCCGTTTGCTTTCGATATTCTGGCGACGA
GCTTTCTGTATGGTAGCAAAACGTTTATAAACCACACTCAGGGCATTCCGGATTTCTTTAAACAAAGCTTTCCGGAAGGTTTTACCTGGGAGCGTGTGAC
TACGTATGAAGATGGTGGTGTGTTGACTGCTACTCAAGATACTTCACTGCAGGACGGCTGTCTGATCTATAACGTGAAGATTCGTGGCGTGAACTTTACG
AGCAATGGGCCGGTAATGCAAAAAAAAACCCTGGGTTGGGAAGCGTTCACGGAAACTCTGTATCCGGCTGACGGCGGCCTGGAGGGCCGTAACGATATGG
CACTGAAGCTGGTTGGTGGCAGCCACCTGATCGCGAATATCAAAACGACTTATCGCTCTAAAAAACCGGCGAAAAATCTGAAGATGCCGGGTGTTTATTA
TGTTGACTATCGTCTGGAACGCATTAAAGAAGCGAATAATGAAACTTACGTGGAGCAACACGAGGTTGCAGTGGCGCGCTATTGCGACTTGCCTTCAAAG
CTGGGTCACAAACTGAATTAATAAGGAAACACAGAAAAAAGCCCGCACCTGACAGTGCGGGCTTTTTTTTTCGACCAAAGGGACAATGTCCTGAAGAATA
TTTAAGAAAAAAGCACCCCTCGTCGCCTAGAATTACCTACCGCGGTCGACCATACCTTCGAGACGAACAATAAGGCCGCAAATCGCGGCCTTCGTTATTG
ATAACAGCAGGTAAAAAGACGCTTATTATGGAGTTCGACATGGCAATAACGCGACGAATCTACGTCATGACGAGAATAGTATAAACAAAACTGCTGACGG
CAGAAGCGTCTCGGTACCAAATTCCAGAAAAGAGACGCTTTCGAGCGTCTTTTTTCGTTTTGGTCCACGCTGATGAATATTCTACGGAATTGCCATAGGC
GTTGAACGCTACACGGACGATACGAATTTATGTATAGAGCGTTCAGCCAAAAAACTTAAGACCGCCGGTCTTGTCCACTACCTTGCAGTAATGCGGTGGA
CAGGATCGGCGGTTTTCTTTTCTCTTCTCAATTCGCGACCCAGGATGAGGTCGCCAAAACTACTAGATTTGGGCGCGCCTTGACGGCTAGCTCAGTCCTA
GGTACAGTGCTAGCTTAATGCGAGAGGCTGGTTAAGTGGGTGTTGATGCGTAAAGGAGAAGAACTGTTCACTGGAGTTGTCCCAATTCTTGTTGAATTAG
ATGGTGATGTTAATGGGCACAAATTTTCTGTGAGTGGAGAGGGTGAAGGTGATGCGACATACGGCAAACTGACCCTTAAATTTATTTGTACTACCGGCAA
GCTACCTGTTCCTTGGCCGACACTTGTCACTACTTTCGGTTATGGTGTTCAATGTTTTGCGAGATACCCAGATCATATGAAACAGCATGACTTTTTCAAA
AGTGCCATGCCCGAAGGTTATGTACAGGAAAGAACTATATTTTTCAAAGATGACGGGAACTACAAAACACGTGCTGAAGTTAAATTTGAGGGTGATACTC
TGGTAAATAGAATCGAATTAAAAGGTATTGATTTTAAAGAAGATGGCAACATTCTTGGTCACAAGCTGGAATACAACTACAACTCACACAATGTTTACAT
CATGGCAGACAAACAAAAAAATGGCATCAAAGTTAACTTCAAAATTCGACACAACATTGAAGATGGAAGCGTTCAACTGGCAGACCATTATCAACAGAAT

ACTCCGATTGGCGATGGCCCTGTACTCTTACCAGACAACCATTACCTGTCCACACAATCTGCCCTTTCGAAAGATCCGAACGAAAAGAGAGATCACATGG
TCCTGCTTGAGTTTGTAACAGCTGCTGGGATTACACATGGTATGGATGAGCTATACAAATAATAACCAATTATTGAAGGCCTCCCTAACGGGGGGCCTTT
TTTTGTTTCTGGTCTCCCCACTGATAGTGCTAGTGTAGATCACTACTAGAGCCAGGCATCAAATAAAACGAAAGGCTCAGTCGAAAGACTGGGCCTTTCG
TTTTATCTGTTGTTTGTCGGTGAACGCTCTCTACTAGAGTCACACTGGCTCACCTTCGGGTGGGCCTTTCTGCGTTTATATACTAGAGGGAATCGTACGG
TCGATCCATCTGACTATCGCCTACTTCATTGACAGCTAGCTCAGTCCTAGGTATAATGCTAGCAAATATGTTGTAGAAACAATTGCACGAATGGCTTCCT
CCGAAGACGTTATCAAAGAGTTCATGCGTTTCAAAGTTCGTATGGAAGGTTCCGTTAACGGTCACGAGTTCGAAATCGAAGGTGAAGGTGAAGGTCGTCC
GTACGAAGGTACCCAGACCGCTAAACTGAAAGTTACCAAAGGTGGTCCGCTGCCGTTCGCTTGGGACATCCTGTCCCCGCAGTTCCAGTACGGTTCCAAA
GCTTACGTTAAACACCCGGCTGACATCCCGGACTACCTGAAACTGTCCTTCCCGGAAGGTTTCAAATGGGAACGTGTTATGAACTTCGAAGACGGTGGTG
TTGTTACCGTTACCCAGGACTCCTCCCTGCAAGACGGTGAGTTCATCTACAAAGTTAAACTGCGTGGTACCAACTTCCCGTCCGACGGTCCGGTTATGCA
GAAAAAAACCATGGGTTGGGAAGCTTCCACCGAACGTATGTACCCGGAAGACGGTGCTCTGAAAGGTGAAATCAAAATGCGTCTGAAACTGAAAGACGGT
GGTCACTACGACGCTGAAGTTAAAACCACCTACATGGCTAAAAAACCGGTTCAGCTGCCGGGTGCTTACAAAACCGACATCAAACTGGACATCACCTCCC
ACAACGAAGACTACACCATCGTTGAACAGTACGAACGTGCTGAAGGTCGTCACTCCACCGGTGCTTAATAACGCTGATAGTGCTAGTGTAGATCGCTACT
AGAGTACTAGCTCTAGATGCGGCCGCGAATTCCAGAAATCATCCTTAGCGAAAGCTAAGGATTTTTTTTATCTGAAATGCCTCTGTAACAGAGCATTAGC
GCAAGGTGATTTTTGTCTTCTTGCGCTAATTTTTTGGCGTCTCCACTGTGCGCGGAACCCCTATTTGTTTATTTTTCTAAATACATTCAAATATGTATCC
GCTCATGAGACAATAACCCTGATAAATGCTTCAATAATATTGAAAAAGGAAGAGTATGAGTATTCAACATTTCCGTGTCGCCCTTATTCCCTTTTTTGCG
GCATTTTGCCTTCCTGTTTTTGCTCACCCAGAAACGCTGGTGAAAGTAAAAGATGCTGAAGATCAGTTGGGTGCACGAGTGGGTTACATCGAACTGGATC
TCAACAGCGGTAAGATCCTTGAGAGTTTTCGCCCCGAAGAACGTTTTCCAATGATGAGCACTTTTAAAGTTCTGCTATGTGGCGCGGTATTATCCCGTAT
TGACGCCGGGCAAGAGCAACTCGGTCGCCGCATACACTATTCTCAGAATGACTTGGTTGAGTACTCACCAGTCACAGAAAAGCATCTTACGGATGGCATG
ACAGTAAGAGAATTATGCAGTGCTGCCATAACCATGAGTGATAACACTGCGGCCAACTTACTTCTGACAACGATCGGAGGACCGAAGGAGCTAACCGCTT
TTTTGCACAACATGGGGGATCATGTAACTCGCCTTGATCGTTGGGAACCGGAGCTGAATGAAGCCATACCAAACGACGAGCGTGACACCACGATGCCTGT
AGCAATGGCAACAACGTTGCGCAAACTATTAACTGGCGAACTACTTACTCTAGCTTCCCGGCAACAATTAATAGACTGGATGGAGGCGGATAAAGTTGCA
GGACCACTTCTGCGCTCGGCCCTTCCGGCTGGCTGGTTTATTGCTGATAAATCTGGAGCCGGTGAGCGTGGGTCTCGCGGTATCATTGCAGCACTGGGGC
CAGATGGTAAGCCCTCCCGTATCGTAGTTATCTACACGACGGGGAGTCAGGCAACTATGGATGAACGAAATAGACAGATCGCTGAGATAGGTGCCTCACT
GATTAAGCATTGGTAACTCGAGCTGATCCTTCAAGAGCTCGCTTGGACTCCTGTTGATAGATCCAGTAATGACCTCAGAACTCCATCTGGATTTGTTCAG
AACGCTCGGTTGCCGCCGGGCGTTTTTTATTGGTGAGAATTTACAACTTATATCGTATGGGGCTGACTTCAGGTGCTACATTTGAAGAGATAAATTGCAC
TGAAATCTAGAAATATTTTATCTGATTAATAAGATGATCTTCTTGAGATCGTTTTGGTCTGCGCGTAATCTCTTGCTCTGAAAACGAAAAAACCGCCTTG
CAGGGCGGTTTTTCGAAGGTTCTCTGAGCTACCAACTCTTTGAACCGAGGTAACTGGCTTGGAGGAGCGCAGTCACCAAAACTTGTCCTTTCAGTTTAGC
CTTAACCGGCGCATGACTTCAAGACTAACTCCTCTAAATCAATTACCAGTGGCTGCTGCCAGTGGTGCTTTTGCATGTCTTTCCGGGTTGGACTCAAGAC
GATAGTTACCGGATAAGGCGCAGCGGTCGGACTGAACGGGGGGTTCGTGCATACAGTCCAGCTTGGAGCGAACTGCCTACCCGGAACTGAGTGTCAGGCG
TGGAATGAGACAAACGCGGCCATAACAGCGGAATGACACCGGTAAACCGAAAGGCAGGAACAGGAGAGCGCACGAGGGAGCCGCCAGGGGGAAACGCCTG
GTATCTTTATAGTCCTGTCGGGTTTCGCCACCACTGATTTGAGCGTCAGATT

**pCM202** TCGTGATGCTTGTCAGGGGGGCGGAGCCTATGGAAAAACGGCTTTGCCGCGGCCCTCTCACTTCCCTGTTAAGTATCTTCCTGGCATCTTCCAGGAAATC
TCCGCCCCGTTCGTAAGCCATTTCCGCTCGCCGCAGTCGAACGGGCGCGCCTTGACAGCTAGCTCAGTCCTAGGTATTGTGCTAGCTTACGCCGATTCTG
CCGGGGGGAATTATAATGAGCGAACTGATCAAAGAGAACATGCACATGAAGCTGTACATGGAGGGTACCGTGGATAATCACCACTTTAAGTGTACTTCTG
AGGGCGAGGGTAAGCCGTATGAAGGGACTCAAACGATGCGTATTAAAGTAGTGGAGGGTGGCCCACTGCCGTTTGCTTTCGATATTCTGGCGACGAGCTT
TCTGTATGGTAGCAAAACGTTTATAAACCACACTCAGGGCATTCCGGATTTCTTTAAACAAAGCTTTCCGGAAGGTTTTACCTGGGAGCGTGTGACTACG
TATGAAGATGGTGGTGTGTTGACTGCTACTCAAGATACTTCACTGCAGGACGGCTGTCTGATCTATAACGTGAAGATTCGTGGCGTGAACTTTACGAGCA
ATGGGCCGGTAATGCAAAAAAAAACCCTGGGTTGGGAAGCGTTCACGGAAACTCTGTATCCGGCTGACGGCGGCCTGGAGGGCCGTAACGATATGGCACT
GAAGCTGGTTGGTGGCAGCCACCTGATCGCGAATATCAAAACGACTTATCGCTCTAAAAAACCGGCGAAAAATCTGAAGATGCCGGGTGTTTATTATGTT
GACTATCGTCTGGAACGCATTAAAGAAGCGAATAATGAAACTTACGTGGAGCAACACGAGGTTGCAGTGGCGCGCTATTGCGACTTGCCTTCAAAGCTGG
GTCACAAACTGAATTAATAAGGAAACACAGAAAAAAGCCCGCACCTGACAGTGCGGGCTTTTTTTTTCGACCAAAGGGACAATGTCCTGAAGAATATTTA
AGAAAAAAGCACCCCTCGTCGCCTAGAATTACCTACCGCGGTCGACCATACCTTCGAGACGAACAATAAGGCCGCAAATCGCGGCCTTCGTTATTGATAA
CAGCAGGTAAAAAGACGCTTATTATGGAGTTCGACATGGCAATAACGCGACGAATCTACGTCATGACGAGAATAGTATAAACAAAACTGCTGACGGCAGA
AGCGTCTCGGTACCAAATTCCAGAAAAGAGACGCTTTCGAGCGTCTTTTTTCGTTTTGGTCCACGCTGATGAATATTCTACGGAATTGCCATAGGCGTTG
AACGCTACACGGACGATACGAATTTATGTATAGAGCGTTCAGCCAAAAAACTTAAGACCGCCGGTCTTGTCCACTACCTTGCAGTAATGCGGTGGACAGG
ATCGGCGGTTTTCTTTTCTCTTCTCAATTCGCGACCCAGGATGAGGTCGCCAAAACTACTAGATTTGGGCGCGCCTTGACGGCTAGCTCAGTCCTAGGTA
CAGTGCTAGCTTAATTTCTAATTAGGATAAGGACACCAAGATGCGTAAAGGAGAAGAACTGTTCACTGGAGTTGTCCCAATTCTTGTTGAATTAGATGGT
GATGTTAATGGGCACAAATTTTCTGTGAGTGGAGAGGGTGAAGGTGATGCGACATACGGCAAACTGACCCTTAAATTTATTTGTACTACCGGCAAGCTAC
CTGTTCCTTGGCCGACACTTGTCACTACTTTCGGTTATGGTGTTCAATGTTTTGCGAGATACCCAGATCATATGAAACAGCATGACTTTTTCAAAAGTGC
CATGCCCGAAGGTTATGTACAGGAAAGAACTATATTTTTCAAAGATGACGGGAACTACAAAACACGTGCTGAAGTTAAATTTGAGGGTGATACTCTGGTA
AATAGAATCGAATTAAAAGGTATTGATTTTAAAGAAGATGGCAACATTCTTGGTCACAAGCTGGAATACAACTACAACTCACACAATGTTTACATCATGG
CAGACAAACAAAAAAATGGCATCAAAGTTAACTTCAAAATTCGACACAACATTGAAGATGGAAGCGTTCAACTGGCAGACCATTATCAACAGAATACTCC
GATTGGCGATGGCCCTGTACTCTTACCAGACAACCATTACCTGTCCACACAATCTGCCCTTTCGAAAGATCCGAACGAAAAGAGAGATCACATGGTCCTG
CTTGAGTTTGTAACAGCTGCTGGGATTACACATGGTATGGATGAGCTATACAAATAATAACCAATTATTGAAGGCCTCCCTAACGGGGGGCCTTTTTTTG
TTTCTGGTCTCCCCACTGATAGTGCTAGTGTAGATCACTACTAGAGCCAGGCATCAAATAAAACGAAAGGCTCAGTCGAAAGACTGGGCCTTTCGTTTTA
TCTGTTGTTTGTCGGTGAACGCTCTCTACTAGAGTCACACTGGCTCACCTTCGGGTGGGCCTTTCTGCGTTTATATACTAGAGGGAATCGTACGGTCGAT
CCATCTGACTATCGCCTACTTCATTGACAGCTAGCTCAGTCCTAGGTATAATGCTAGCAAATATGTTGTAGAAACAATTGCACGAATGGCTTCCTCCGAA
GACGTTATCAAAGAGTTCATGCGTTTCAAAGTTCGTATGGAAGGTTCCGTTAACGGTCACGAGTTCGAAATCGAAGGTGAAGGTGAAGGTCGTCCGTACG
AAGGTACCCAGACCGCTAAACTGAAAGTTACCAAAGGTGGTCCGCTGCCGTTCGCTTGGGACATCCTGTCCCCGCAGTTCCAGTACGGTTCCAAAGCTTA
CGTTAAACACCCGGCTGACATCCCGGACTACCTGAAACTGTCCTTCCCGGAAGGTTTCAAATGGGAACGTGTTATGAACTTCGAAGACGGTGGTGTTGTT
ACCGTTACCCAGGACTCCTCCCTGCAAGACGGTGAGTTCATCTACAAAGTTAAACTGCGTGGTACCAACTTCCCGTCCGACGGTCCGGTTATGCAGAAAA
AAACCATGGGTTGGGAAGCTTCCACCGAACGTATGTACCCGGAAGACGGTGCTCTGAAAGGTGAAATCAAAATGCGTCTGAAACTGAAAGACGGTGGTCA
CTACGACGCTGAAGTTAAAACCACCTACATGGCTAAAAAACCGGTTCAGCTGCCGGGTGCTTACAAAACCGACATCAAACTGGACATCACCTCCCACAAC
GAAGACTACACCATCGTTGAACAGTACGAACGTGCTGAAGGTCGTCACTCCACCGGTGCTTAATAACGCTGATAGTGCTAGTGTAGATCGCTACTAGAGT
ACTAGCTCTAGATGCGGCCGCGAATTCCAGAAATCATCCTTAGCGAAAGCTAAGGATTTTTTTTATCTGAAATGCCTCTGTAACAGAGCATTAGCGCAAG
GTGATTTTTGTCTTCTTGCGCTAATTTTTTGGCGTCTCCACTGTGCGCGGAACCCCTATTTGTTTATTTTTCTAAATACATTCAAATATGTATCCGCTCA
TGAGACAATAACCCTGATAAATGCTTCAATAATATTGAAAAAGGAAGAGTATGAGTATTCAACATTTCCGTGTCGCCCTTATTCCCTTTTTTGCGGCATT
TTGCCTTCCTGTTTTTGCTCACCCAGAAACGCTGGTGAAAGTAAAAGATGCTGAAGATCAGTTGGGTGCACGAGTGGGTTACATCGAACTGGATCTCAAC
AGCGGTAAGATCCTTGAGAGTTTTCGCCCCGAAGAACGTTTTCCAATGATGAGCACTTTTAAAGTTCTGCTATGTGGCGCGGTATTATCCCGTATTGACG
CCGGGCAAGAGCAACTCGGTCGCCGCATACACTATTCTCAGAATGACTTGGTTGAGTACTCACCAGTCACAGAAAAGCATCTTACGGATGGCATGACAGT
AAGAGAATTATGCAGTGCTGCCATAACCATGAGTGATAACACTGCGGCCAACTTACTTCTGACAACGATCGGAGGACCGAAGGAGCTAACCGCTTTTTTG
CACAACATGGGGGATCATGTAACTCGCCTTGATCGTTGGGAACCGGAGCTGAATGAAGCCATACCAAACGACGAGCGTGACACCACGATGCCTGTAGCAA
TGGCAACAACGTTGCGCAAACTATTAACTGGCGAACTACTTACTCTAGCTTCCCGGCAACAATTAATAGACTGGATGGAGGCGGATAAAGTTGCAGGACC
ACTTCTGCGCTCGGCCCTTCCGGCTGGCTGGTTTATTGCTGATAAATCTGGAGCCGGTGAGCGTGGGTCTCGCGGTATCATTGCAGCACTGGGGCCAGAT
GGTAAGCCCTCCCGTATCGTAGTTATCTACACGACGGGGAGTCAGGCAACTATGGATGAACGAAATAGACAGATCGCTGAGATAGGTGCCTCACTGATTA
AGCATTGGTAACTCGAGCTGATCCTTCAAGAGCTCGCTTGGACTCCTGTTGATAGATCCAGTAATGACCTCAGAACTCCATCTGGATTTGTTCAGAACGC
TCGGTTGCCGCCGGGCGTTTTTTATTGGTGAGAATTTACAACTTATATCGTATGGGGCTGACTTCAGGTGCTACATTTGAAGAGATAAATTGCACTGAAA
TCTAGAAATATTTTATCTGATTAATAAGATGATCTTCTTGAGATCGTTTTGGTCTGCGCGTAATCTCTTGCTCTGAAAACGAAAAAACCGCCTTGCAGGG

CGGTTTTTCGAAGGTTCTCTGAGCTACCAACTCTTTGAACCGAGGTAACTGGCTTGGAGGAGCGCAGTCACCAAAACTTGTCCTTTCAGTTTAGCCTTAA
CCGGCGCATGACTTCAAGACTAACTCCTCTAAATCAATTACCAGTGGCTGCTGCCAGTGGTGCTTTTGCATGTCTTTCCGGGTTGGACTCAAGACGATAG
TTACCGGATAAGGCGCAGCGGTCGGACTGAACGGGGGGTTCGTGCATACAGTCCAGCTTGGAGCGAACTGCCTACCCGGAACTGAGTGTCAGGCGTGGAA
TGAGACAAACGCGGCCATAACAGCGGAATGACACCGGTAAACCGAAAGGCAGGAACAGGAGAGCGCACGAGGGAGCCGCCAGGGGGAAACGCCTGGTATC
TTTATAGTCCTGTCGGGTTTCGCCACCACTGATTTGAGCGTCAGATT

**pCM203**

TCGTGATGCTTGTCAGGGGGGCGGAGCCTATGGAAAAACGGCTTTGCCGCGGCCCTCTCACTTCCCTGTTAAGTATCTTCCTGGCATCTTCCAGGAAATC
TCCGCCCCGTTCGTAAGCCATTTCCGCTCGCCGCAGTCGAACGGGCGCGCCTTGACAGCTAGCTCAGTCCTAGGTATTGTGCTAGCTTACGTCAACCCAC
CCTAGGAGAGGATAAGATGAGCGAACTGATCAAAGAGAACATGCACATGAAGCTGTACATGGAGGGTACCGTGGATAATCACCACTTTAAGTGTACTTCT
GAGGGCGAGGGTAAGCCGTATGAAGGGACTCAAACGATGCGTATTAAAGTAGTGGAGGGTGGCCCACTGCCGTTTGCTTTCGATATTCTGGCGACGAGCT
TTCTGTATGGTAGCAAAACGTTTATAAACCACACTCAGGGCATTCCGGATTTCTTTAAACAAAGCTTTCCGGAAGGTTTTACCTGGGAGCGTGTGACTAC
GTATGAAGATGGTGGTGTGTTGACTGCTACTCAAGATACTTCACTGCAGGACGGCTGTCTGATCTATAACGTGAAGATTCGTGGCGTGAACTTTACGAGC
AATGGGCCGGTAATGCAAAAAAAAACCCTGGGTTGGGAAGCGTTCACGGAAACTCTGTATCCGGCTGACGGCGGCCTGGAGGGCCGTAACGATATGGCAC
TGAAGCTGGTTGGTGGCAGCCACCTGATCGCGAATATCAAAACGACTTATCGCTCTAAAAAACCGGCGAAAAATCTGAAGATGCCGGGTGTTTATTATGT
TGACTATCGTCTGGAACGCATTAAAGAAGCGAATAATGAAACTTACGTGGAGCAACACGAGGTTGCAGTGGCGCGCTATTGCGACTTGCCTTCAAAGCTG
GGTCACAAACTGAATTAATAAGGAAACACAGAAAAAAGCCCGCACCTGACAGTGCGGGCTTTTTTTTTCGACCAAAGGGACAATGTCCTGAAGAATATTT
AAGAAAAAAGCACCCCTCGTCGCCTAGAATTACCTACCGCGGTCGACCATACCTTCGAGACGAACAATAAGGCCGCAAATCGCGGCCTTCGTTATTGATA
ACAGCAGGTAAAAAGACGCTTATTATGGAGTTCGACATGGCAATAACGCGACGAATCTACGTCATGACGAGAATAGTATAAACAAAACTGCTGACGGCAG
AAGCGTCTCGGTACCAAATTCCAGAAAAGAGACGCTTTCGAGCGTCTTTTTTCGTTTTGGTCCACGCTGATGAATATTCTACGGAATTGCCATAGGCGTT
GAACGCTACACGGACGATACGAATTTATGTATAGAGCGTTCAGCCAAAAAACTTAAGACCGCCGGTCTTGTCCACTACCTTGCAGTAATGCGGTGGACAG
GATCGGCGGTTTTCTTTTCTCTTCTCAATTCGCGACCCAGGATGAGGTCGCCAAAACTACTAGATTTGGGCGCGCCTTGACGGCTAGCTCAGTCCTAGGT
ACAGTGCTAGCTTAATTTCTAATTAGGATAAGGACACCAAGATGCGTAAAGGAGAAGAACTGTTCACTGGAGTTGTCCCAATTCTTGTTGAATTAGATGG
TGATGTTAATGGGCACAAATTTTCTGTGAGTGGAGAGGGTGAAGGTGATGCGACATACGGCAAACTGACCCTTAAATTTATTTGTACTACCGGCAAGCTA
CCTGTTCCTTGGCCGACACTTGTCACTACTTTCGGTTATGGTGTTCAATGTTTTGCGAGATACCCAGATCATATGAAACAGCATGACTTTTTCAAAAGTG
CCATGCCCGAAGGTTATGTACAGGAAAGAACTATATTTTTCAAAGATGACGGGAACTACAAAACACGTGCTGAAGTTAAATTTGAGGGTGATACTCTGGT
AAATAGAATCGAATTAAAAGGTATTGATTTTAAAGAAGATGGCAACATTCTTGGTCACAAGCTGGAATACAACTACAACTCACACAATGTTTACATCATG
GCAGACAAACAAAAAAATGGCATCAAAGTTAACTTCAAAATTCGACACAACATTGAAGATGGAAGCGTTCAACTGGCAGACCATTATCAACAGAATACTC
CGATTGGCGATGGCCCTGTACTCTTACCAGACAACCATTACCTGTCCACACAATCTGCCCTTTCGAAAGATCCGAACGAAAAGAGAGATCACATGGTCCT
GCTTGAGTTTGTAACAGCTGCTGGGATTACACATGGTATGGATGAGCTATACAAATAATAACCAATTATTGAAGGCCTCCCTAACGGGGGGCCTTTTTTT
GTTTCTGGTCTCCCCACTGATAGTGCTAGTGTAGATCACTACTAGAGCCAGGCATCAAATAAAACGAAAGGCTCAGTCGAAAGACTGGGCCTTTCGTTTT
ATCTGTTGTTTGTCGGTGAACGCTCTCTACTAGAGTCACACTGGCTCACCTTCGGGTGGGCCTTTCTGCGTTTATATACTAGAGGGAATCGTACGGTCGA
TCCATCTGACTATCGCCTACTTCATTGACAGCTAGCTCAGTCCTAGGTATAATGCTAGCAAATATGTTGTAGAAACAATTGCACGAATGGCTTCCTCCGA
AGACGTTATCAAAGAGTTCATGCGTTTCAAAGTTCGTATGGAAGGTTCCGTTAACGGTCACGAGTTCGAAATCGAAGGTGAAGGTGAAGGTCGTCCGTAC
GAAGGTACCCAGACCGCTAAACTGAAAGTTACCAAAGGTGGTCCGCTGCCGTTCGCTTGGGACATCCTGTCCCCGCAGTTCCAGTACGGTTCCAAAGCTT
ACGTTAAACACCCGGCTGACATCCCGGACTACCTGAAACTGTCCTTCCCGGAAGGTTTCAAATGGGAACGTGTTATGAACTTCGAAGACGGTGGTGTTGT
TACCGTTACCCAGGACTCCTCCCTGCAAGACGGTGAGTTCATCTACAAAGTTAAACTGCGTGGTACCAACTTCCCGTCCGACGGTCCGGTTATGCAGAAA
AAAACCATGGGTTGGGAAGCTTCCACCGAACGTATGTACCCGGAAGACGGTGCTCTGAAAGGTGAAATCAAAATGCGTCTGAAACTGAAAGACGGTGGTC
ACTACGACGCTGAAGTTAAAACCACCTACATGGCTAAAAAACCGGTTCAGCTGCCGGGTGCTTACAAAACCGACATCAAACTGGACATCACCTCCCACAA
CGAAGACTACACCATCGTTGAACAGTACGAACGTGCTGAAGGTCGTCACTCCACCGGTGCTTAATAACGCTGATAGTGCTAGTGTAGATCGCTACTAGAG
TACTAGCTCTAGATGCGGCCGCGAATTCCAGAAATCATCCTTAGCGAAAGCTAAGGATTTTTTTTATCTGAAATGCCTCTGTAACAGAGCATTAGCGCAA
GGTGATTTTTGTCTTCTTGCGCTAATTTTTTGGCGTCTCCACTGTGCGCGGAACCCCTATTTGTTTATTTTTCTAAATACATTCAAATATGTATCCGCTC
ATGAGACAATAACCCTGATAAATGCTTCAATAATATTGAAAAAGGAAGAGTATGAGTATTCAACATTTCCGTGTCGCCCTTATTCCCTTTTTTGCGGCAT
TTTGCCTTCCTGTTTTTGCTCACCCAGAAACGCTGGTGAAAGTAAAAGATGCTGAAGATCAGTTGGGTGCACGAGTGGGTTACATCGAACTGGATCTCAA
CAGCGGTAAGATCCTTGAGAGTTTTCGCCCCGAAGAACGTTTTCCAATGATGAGCACTTTTAAAGTTCTGCTATGTGGCGCGGTATTATCCCGTATTGAC
GCCGGGCAAGAGCAACTCGGTCGCCGCATACACTATTCTCAGAATGACTTGGTTGAGTACTCACCAGTCACAGAAAAGCATCTTACGGATGGCATGACAG
TAAGAGAATTATGCAGTGCTGCCATAACCATGAGTGATAACACTGCGGCCAACTTACTTCTGACAACGATCGGAGGACCGAAGGAGCTAACCGCTTTTTT
GCACAACATGGGGGATCATGTAACTCGCCTTGATCGTTGGGAACCGGAGCTGAATGAAGCCATACCAAACGACGAGCGTGACACCACGATGCCTGTAGCA
ATGGCAACAACGTTGCGCAAACTATTAACTGGCGAACTACTTACTCTAGCTTCCCGGCAACAATTAATAGACTGGATGGAGGCGGATAAAGTTGCAGGAC
CACTTCTGCGCTCGGCCCTTCCGGCTGGCTGGTTTATTGCTGATAAATCTGGAGCCGGTGAGCGTGGGTCTCGCGGTATCATTGCAGCACTGGGGCCAGA
TGGTAAGCCCTCCCGTATCGTAGTTATCTACACGACGGGGAGTCAGGCAACTATGGATGAACGAAATAGACAGATCGCTGAGATAGGTGCCTCACTGATT
AAGCATTGGTAACTCGAGCTGATCCTTCAAGAGCTCGCTTGGACTCCTGTTGATAGATCCAGTAATGACCTCAGAACTCCATCTGGATTTGTTCAGAACG
CTCGGTTGCCGCCGGGCGTTTTTTATTGGTGAGAATTTACAACTTATATCGTATGGGGCTGACTTCAGGTGCTACATTTGAAGAGATAAATTGCACTGAA
ATCTAGAAATATTTTATCTGATTAATAAGATGATCTTCTTGAGATCGTTTTGGTCTGCGCGTAATCTCTTGCTCTGAAAACGAAAAAACCGCCTTGCAGG
GCGGTTTTTCGAAGGTTCTCTGAGCTACCAACTCTTTGAACCGAGGTAACTGGCTTGGAGGAGCGCAGTCACCAAAACTTGTCCTTTCAGTTTAGCCTTA
ACCGGCGCATGACTTCAAGACTAACTCCTCTAAATCAATTACCAGTGGCTGCTGCCAGTGGTGCTTTTGCATGTCTTTCCGGGTTGGACTCAAGACGATA
GTTACCGGATAAGGCGCAGCGGTCGGACTGAACGGGGGGTTCGTGCATACAGTCCAGCTTGGAGCGAACTGCCTACCCGGAACTGAGTGTCAGGCGTGGA
ATGAGACAAACGCGGCCATAACAGCGGAATGACACCGGTAAACCGAAAGGCAGGAACAGGAGAGCGCACGAGGGAGCCGCCAGGGGGAAACGCCTGGTAT
CTTTATAGTCCTGTCGGGTTTCGCCACCACTGATTTGAGCGTCAGATT

**pCM210**

TCGTGATGCTTGTCAGGGGGGCGGAGCCTATGGAAAAACGGCTTTGCCGCGGCCCTCTCACTTCCCTGTTAAGTATCTTCCTGGCATCTTCCAGGAAATC
TCCGCCCCGTTCGTAAGCCATTTCCGCTCGCCGCAGTCGAACGGGCGCGCCTTGACAGCTAGCTCAGTCCTAGGTATTGTGCTAGCTTACGTCAACCCAC
CCTAGGAGAGGATAAGATGAGCGAACTGATCAAAGAGAACATGCACATGAAGCTGTACATGGAGGGTACCGTGGATAATCACCACTTTAAGTGTACTTCT
GAGGGCGAGGGTAAGCCGTATGAAGGGACTCAAACGATGCGTATTAAAGTAGTGGAGGGTGGCCCACTGCCGTTTGCTTTCGATATTCTGGCGACGAGCT
TTCTGTATGGTAGCAAAACGTTTATAAACCACACTCAGGGCATTCCGGATTTCTTTAAACAAAGCTTTCCGGAAGGTTTTACCTGGGAGCGTGTGACTAC
GTATGAAGATGGTGGTGTGTTGACTGCTACTCAAGATACTTCACTGCAGGACGGCTGTCTGATCTATAACGTGAAGATTCGTGGCGTGAACTTTACGAGC
AATGGGCCGGTAATGCAAAAAAAAACCCTGGGTTGGGAAGCGTTCACGGAAACTCTGTATCCGGCTGACGGCGGCCTGGAGGGCCGTAACGATATGGCAC
TGAAGCTGGTTGGTGGCAGCCACCTGATCGCGAATATCAAAACGACTTATCGCTCTAAAAAACCGGCGAAAAATCTGAAGATGCCGGGTGTTTATTATGT
TGACTATCGTCTGGAACGCATTAAAGAAGCGAATAATGAAACTTACGTGGAGCAACACGAGGTTGCAGTGGCGCGCTATTGCGACTTGCCTTCAAAGCTG
GGTCACAAACTGAATTAATAAGGAAACACAGAAAAAAGCCCGCACCTGACAGTGCGGGCTTTTTTTTTCGACCAAAGGGACAATGTCCTGAAGAATATTT
AAGAAAAAAGCACCCCTCGTCGCCTAGAATTACCTACCGCGGTCGACCATACCTTCGAGACGAACAATAAGGCCGCAAATCGCGGCCTTCGTTATTGATA
ACAGCAGGTAAAAAGACGCTTATTATGGAGTTCGACATGGCAATAACGCGACGAATCTACGTCATGACGAGAATAGTATAAACAAAACTGCTGACGGCAG
AAGCGTCTCGGTACCAAATTCCAGAAAAGAGACGCTTTCGAGCGTCTTTTTTCGTTTTGGTCCACGCTGATGAATATTCTACGGAATTGCCATAGGCGTT
GAACGCTACACGGACGATACGAATTTATGTATAGAGCGTTCAGCCAAAAAACTTAAGACCGCCGGTCTTGTCCACTACCTTGCAGTAATGCGGTGGACAG
GATCGGCGGTTTTCTTTTCTCTTCTCAATTCGCGACCCAGGATGAGGTCGCCAAAACTACTAGATTTGGGCGCGCCTTGACGGCTAGCTCAGTCCTAGGT
ACAGTGCTAGCTTAATCTTGTCAAACAGTGGCTACGTGAACAGATGCGTAAAGGAGAAGAACTGTTCACTGGAGTTGTCCCAATTCTTGTTGAATTAGAT
GGTGATGTTAATGGGCACAAATTTTCTGTGAGTGGAGAGGGTGAAGGTGATGCGACATACGGCAAACTGACCCTTAAATTTATTTGTACTACCGGCAAGC
TACCTGTTCCTTGGCCGACACTTGTCACTACTTTCGGTTATGGTGTTCAATGTTTTGCGAGATACCCAGATCATATGAAACAGCATGACTTTTTCAAAAG
TGCCATGCCCGAAGGTTATGTACAGGAAAGAACTATATTTTTCAAAGATGACGGGAACTACAAAACACGTGCTGAAGTTAAATTTGAGGGTGATACTCTG
GTAAATAGAATCGAATTAAAAGGTATTGATTTTAAAGAAGATGGCAACATTCTTGGTCACAAGCTGGAATACAACTACAACTCACACAATGTTTACATCA

TGGCAGACAAACAAAAAAATGGCATCAAAGTTAACTTCAAAATTCGACACAACATTGAAGATGGAAGCGTTCAACTGGCAGACCATTATCAACAGAATAC
TCCGATTGGCGATGGCCCTGTACTCTTACCAGACAACCATTACCTGTCCACACAATCTGCCCTTTCGAAAGATCCGAACGAAAAGAGAGATCACATGGTC
CTGCTTGAGTTTGTAACAGCTGCTGGGATTACACATGGTATGGATGAGCTATACAAATAATAACCAATTATTGAAGGCCTCCCTAACGGGGGGCCTTTTT
TTGTTTCTGGTCTCCCCACTGATAGTGCTAGTGTAGATCACTACTAGAGCCAGGCATCAAATAAAACGAAAGGCTCAGTCGAAAGACTGGGCCTTTCGTT
TTATCTGTTGTTTGTCGGTGAACGCTCTCTACTAGAGTCACACTGGCTCACCTTCGGGTGGGCCTTTCTGCGTTTATATACTAGAGGGAATCGTACGGTC
GATCCATCTGACTATCGCCTACTTCATTGACAGCTAGCTCAGTCCTAGGTATAATGCTAGCTCTAGAGAAAGAGGAGAAATACTAGATGGCTTCCTCCGA
AGACGTTATCAAAGAGTTCATGCGTTTCAAAGTTCGTATGGAAGGTTCCGTTAACGGTCACGAGTTCGAAATCGAAGGTGAAGGTGAAGGTCGTCCGTAC
GAAGGTACCCAGACCGCTAAACTGAAAGTTACCAAAGGTGGTCCGCTGCCGTTCGCTTGGGACATCCTGTCCCCGCAGTTCCAGTACGGTTCCAAAGCTT
ACGTTAAACACCCGGCTGACATCCCGGACTACCTGAAACTGTCCTTCCCGGAAGGTTTCAAATGGGAACGTGTTATGAACTTCGAAGACGGTGGTGTTGT
TACCGTTACCCAGGACTCCTCCCTGCAAGACGGTGAGTTCATCTACAAAGTTAAACTGCGTGGTACCAACTTCCCGTCCGACGGTCCGGTTATGCAGAAA
AAAACCATGGGTTGGGAAGCTTCCACCGAACGTATGTACCCGGAAGACGGTGCTCTGAAAGGTGAAATCAAAATGCGTCTGAAACTGAAAGACGGTGGTC
ACTACGACGCTGAAGTTAAAACCACCTACATGGCTAAAAAACCGGTTCAGCTGCCGGGTGCTTACAAAACCGACATCAAACTGGACATCACCTCCCACAA
CGAAGACTACACCATCGTTGAACAGTACGAACGTGCTGAAGGTCGTCACTCCACCGGTGCTTAATAACGCTGATAGTGCTAGTGTAGATCGCTACTAGAG
TACTAGCTCTAGATGCGGCCGCGAATTCCAGAAATCATCCTTAGCGAAAGCTAAGGATTTTTTTTATCTGAAATGCCTCTGTAACAGAGCATTAGCGCAA
GGTGATTTTTGTCTTCTTGCGCTAATTTTTTGGCGTCTCCACTGTGCGCGGAACCCCTATTTGTTTATTTTTCTAAATACATTCAAATATGTATCCGCTC
ATGAGACAATAACCCTGATAAATGCTTCAATAATATTGAAAAAGGAAGAGTATGAGTATTCAACATTTCCGTGTCGCCCTTATTCCCTTTTTTGCGGCAT
TTTGCCTTCCTGTTTTTGCTCACCCAGAAACGCTGGTGAAAGTAAAAGATGCTGAAGATCAGTTGGGTGCACGAGTGGGTTACATCGAACTGGATCTCAA
CAGCGGTAAGATCCTTGAGAGTTTTCGCCCCGAAGAACGTTTTCCAATGATGAGCACTTTTAAAGTTCTGCTATGTGGCGCGGTATTATCCCGTATTGAC
GCCGGGCAAGAGCAACTCGGTCGCCGCATACACTATTCTCAGAATGACTTGGTTGAGTACTCACCAGTCACAGAAAAGCATCTTACGGATGGCATGACAG
TAAGAGAATTATGCAGTGCTGCCATAACCATGAGTGATAACACTGCGGCCAACTTACTTCTGACAACGATCGGAGGACCGAAGGAGCTAACCGCTTTTTT
GCACAACATGGGGGATCATGTAACTCGCCTTGATCGTTGGGAACCGGAGCTGAATGAAGCCATACCAAACGACGAGCGTGACACCACGATGCCTGTAGCA
ATGGCAACAACGTTGCGCAAACTATTAACTGGCGAACTACTTACTCTAGCTTCCCGGCAACAATTAATAGACTGGATGGAGGCGGATAAAGTTGCAGGAC
CACTTCTGCGCTCGGCCCTTCCGGCTGGCTGGTTTATTGCTGATAAATCTGGAGCCGGTGAGCGTGGGTCTCGCGGTATCATTGCAGCACTGGGGCCAGA
TGGTAAGCCCTCCCGTATCGTAGTTATCTACACGACGGGGAGTCAGGCAACTATGGATGAACGAAATAGACAGATCGCTGAGATAGGTGCCTCACTGATT
AAGCATTGGTAACTCGAGCTGATCCTTCAAGAGCTCGCTTGGACTCCTGTTGATAGATCCAGTAATGACCTCAGAACTCCATCTGGATTTGTTCAGAACG
CTCGGTTGCCGCCGGGCGTTTTTTATTGGTGAGAATTTACAACTTATATCGTATGGGGCTGACTTCAGGTGCTACATTTGAAGAGATAAATTGCACTGAA
ATCTAGAAATATTTTATCTGATTAATAAGATGATCTTCTTGAGATCGTTTTGGTCTGCGCGTAATCTCTTGCTCTGAAAACGAAAAAACCGCCTTGCAGG
GCGGTTTTTCGAAGGTTCTCTGAGCTACCAACTCTTTGAACCGAGGTAACTGGCTTGGAGGAGCGCAGTCACCAAAACTTGTCCTTTCAGTTTAGCCTTA
ACCGGCGCATGACTTCAAGACTAACTCCTCTAAATCAATTACCAGTGGCTGCTGCCAGTGGTGCTTTTGCATGTCTTTCCGGGTTGGACTCAAGACGATA
GTTACCGGATAAGGCGCAGCGGTCGGACTGAACGGGGGGTTCGTGCATACAGTCCAGCTTGGAGCGAACTGCCTACCCGGAACTGAGTGTCAGGCGTGGA
ATGAGACAAACGCGGCCATAACAGCGGAATGACACCGGTAAACCGAAAGGCAGGAACAGGAGAGCGCACGAGGGAGCCGCCAGGGGGAAACGCCTGGTAT
CTTTATAGTCCTGTCGGGTTTCGCCACCACTGATTTGAGCGTCAGATT

**pCM211**

TCGTGATGCTTGTCAGGGGGGCGGAGCCTATGGAAAAACGGCTTTGCCGCGGCCCTCTCACTTCCCTGTTAAGTATCTTCCTGGCATCTTCCAGGAAATC
TCCGCCCCGTTCGTAAGCCATTTCCGCTCGCCGCAGTCGAACGGGCGCGCCTTGACAGCTAGCTCAGTCCTAGGTATTGTGCTAGCTTACGCCGATTCTG
CCGGGGGGAATTATAATGAGCGAACTGATCAAAGAGAACATGCACATGAAGCTGTACATGGAGGGTACCGTGGATAATCACCACTTTAAGTGTACTTCTG
AGGGCGAGGGTAAGCCGTATGAAGGGACTCAAACGATGCGTATTAAAGTAGTGGAGGGTGGCCCACTGCCGTTTGCTTTCGATATTCTGGCGACGAGCTT
TCTGTATGGTAGCAAAACGTTTATAAACCACACTCAGGGCATTCCGGATTTCTTTAAACAAAGCTTTCCGGAAGGTTTTACCTGGGAGCGTGTGACTACG
TATGAAGATGGTGGTGTGTTGACTGCTACTCAAGATACTTCACTGCAGGACGGCTGTCTGATCTATAACGTGAAGATTCGTGGCGTGAACTTTACGAGCA
ATGGGCCGGTAATGCAAAAAAAAACCCTGGGTTGGGAAGCGTTCACGGAAACTCTGTATCCGGCTGACGGCGGCCTGGAGGGCCGTAACGATATGGCACT
GAAGCTGGTTGGTGGCAGCCACCTGATCGCGAATATCAAAACGACTTATCGCTCTAAAAAACCGGCGAAAAATCTGAAGATGCCGGGTGTTTATTATGTT
GACTATCGTCTGGAACGCATTAAAGAAGCGAATAATGAAACTTACGTGGAGCAACACGAGGTTGCAGTGGCGCGCTATTGCGACTTGCCTTCAAAGCTGG
GTCACAAACTGAATTAATAAGGAAACACAGAAAAAAGCCCGCACCTGACAGTGCGGGCTTTTTTTTTCGACCAAAGGGACAATGTCCTGAAGAATATTTA
AGAAAAAAGCACCCCTCGTCGCCTAGAATTACCTACCGCGGTCGACCATACCTTCGAGACGAACAATAAGGCCGCAAATCGCGGCCTTCGTTATTGATAA
CAGCAGGTAAAAAGACGCTTATTATGGAGTTCGACATGGCAATAACGCGACGAATCTACGTCATGACGAGAATAGTATAAACAAAACTGCTGACGGCAGA
AGCGTCTCGGTACCAAATTCCAGAAAAGAGACGCTTTCGAGCGTCTTTTTTCGTTTTGGTCCACGCTGATGAATATTCTACGGAATTGCCATAGGCGTTG
AACGCTACACGGACGATACGAATTTATGTATAGAGCGTTCAGCCAAAAAACTTAAGACCGCCGGTCTTGTCCACTACCTTGCAGTAATGCGGTGGACAGG
ATCGGCGGTTTTCTTTTCTCTTCTCAATTCGCGACCCAGGATGAGGTCGCCAAAACTACTAGATTTGGGCGCGCCTTGACGGCTAGCTCAGTCCTAGGTA
CAGTGCTAGCTTAATCTTGTCAAACAGTGGCTACGTGAACAGATGCGTAAAGGAGAAGAACTGTTCACTGGAGTTGTCCCAATTCTTGTTGAATTAGATG
GTGATGTTAATGGGCACAAATTTTCTGTGAGTGGAGAGGGTGAAGGTGATGCGACATACGGCAAACTGACCCTTAAATTTATTTGTACTACCGGCAAGCT
ACCTGTTCCTTGGCCGACACTTGTCACTACTTTCGGTTATGGTGTTCAATGTTTTGCGAGATACCCAGATCATATGAAACAGCATGACTTTTTCAAAAGT
GCCATGCCCGAAGGTTATGTACAGGAAAGAACTATATTTTTCAAAGATGACGGGAACTACAAAACACGTGCTGAAGTTAAATTTGAGGGTGATACTCTGG
TAAATAGAATCGAATTAAAAGGTATTGATTTTAAAGAAGATGGCAACATTCTTGGTCACAAGCTGGAATACAACTACAACTCACACAATGTTTACATCAT
GGCAGACAAACAAAAAAATGGCATCAAAGTTAACTTCAAAATTCGACACAACATTGAAGATGGAAGCGTTCAACTGGCAGACCATTATCAACAGAATACT
CCGATTGGCGATGGCCCTGTACTCTTACCAGACAACCATTACCTGTCCACACAATCTGCCCTTTCGAAAGATCCGAACGAAAAGAGAGATCACATGGTCC
TGCTTGAGTTTGTAACAGCTGCTGGGATTACACATGGTATGGATGAGCTATACAAATAATAACCAATTATTGAAGGCCTCCCTAACGGGGGGCCTTTTT
TGTTTCTGGTCTCCCCACTGATAGTGCTAGTGTAGATCACTACTAGAGCCAGGCATCAAATAAAACGAAAGGCTCAGTCGAAAGACTGGGCCTTTCGTTT
TATCTGTTGTTTGTCGGTGAACGCTCTCTACTAGAGTCACACTGGCTCACCTTCGGGTGGGCCTTTCTGCGTTTATATACTAGAGGGAATCGTACGGTCG
ATCCATCTGACTATCGCCTACTTCATTGACAGCTAGCTCAGTCCTAGGTATAATGCTAGCTCTAGAGAAAGAGGAGAAATACTAGATGGCTTCCTCCGAA
GACGTTATCAAAGAGTTCATGCGTTTCAAAGTTCGTATGGAAGGTTCCGTTAACGGTCACGAGTTCGAAATCGAAGGTGAAGGTGAAGGTCGTCCGTACG
AAGGTACCCAGACCGCTAAACTGAAAGTTACCAAAGGTGGTCCGCTGCCGTTCGCTTGGGACATCCTGTCCCCGCAGTTCCAGTACGGTTCCAAAGCTTA
CGTTAAACACCCGGCTGACATCCCGGACTACCTGAAACTGTCCTTCCCGGAAGGTTTCAAATGGGAACGTGTTATGAACTTCGAAGACGGTGGTGTTGTT
ACCGTTACCCAGGACTCCTCCCTGCAAGACGGTGAGTTCATCTACAAAGTTAAACTGCGTGGTACCAACTTCCCGTCCGACGGTCCGGTTATGCAGAAAA
AAACCATGGGTTGGGAAGCTTCCACCGAACGTATGTACCCGGAAGACGGTGCTCTGAAAGGTGAAATCAAAATGCGTCTGAAACTGAAAGACGGTGGTCA
CTACGACGCTGAAGTTAAAACCACCTACATGGCTAAAAAACCGGTTCAGCTGCCGGGTGCTTACAAAACCGACATCAAACTGGACATCACCTCCCACAAC
GAAGACTACACCATCGTTGAACAGTACGAACGTGCTGAAGGTCGTCACTCCACCGGTGCTTAATAACGCTGATAGTGCTAGTGTAGATCGCTACTAGAGT
ACTAGCTCTAGATGCGGCCGCGAATTCCAGAAATCATCCTTAGCGAAAGCTAAGGATTTTTTTTATCTGAAATGCCTCTGTAACAGAGCATTAGCGCAAG
GTGATTTTTGTCTTCTTGCGCTAATTTTTTGGCGTCTCCACTGTGCGCGGAACCCCTATTTGTTTATTTTTCTAAATACATTCAAATATGTATCCGCTCA
TGAGACAATAACCCTGATAAATGCTTCAATAATATTGAAAAAGGAAGAGTATGAGTATTCAACATTTCCGTGTCGCCCTTATTCCCTTTTTTGCGGCATT
TTGCCTTCCTGTTTTTGCTCACCCAGAAACGCTGGTGAAAGTAAAAGATGCTGAAGATCAGTTGGGTGCACGAGTGGGTTACATCGAACTGGATCTCAAC
AGCGGTAAGATCCTTGAGAGTTTTCGCCCCGAAGAACGTTTTCCAATGATGAGCACTTTTAAAGTTCTGCTATGTGGCGCGGTATTATCCCGTATTGACG
CCGGGCAAGAGCAACTCGGTCGCCGCATACACTATTCTCAGAATGACTTGGTTGAGTACTCACCAGTCACAGAAAAGCATCTTACGGATGGCATGACAGT
AAGAGAATTATGCAGTGCTGCCATAACCATGAGTGATAACACTGCGGCCAACTTACTTCTGACAACGATCGGAGGACCGAAGGAGCTAACCGCTTTTTTG
CACAACATGGGGGATCATGTAACTCGCCTTGATCGTTGGGAACCGGAGCTGAATGAAGCCATACCAAACGACGAGCGTGACACCACGATGCCTGTAGCAA
TGGCAACAACGTTGCGCAAACTATTAACTGGCGAACTACTTACTCTAGCTTCCCGGCAACAATTAATAGACTGGATGGAGGCGGATAAAGTTGCAGGACC
ACTTCTGCGCTCGGCCCTTCCGGCTGGCTGGTTTATTGCTGATAAATCTGGAGCCGGTGAGCGTGGGTCTCGCGGTATCATTGCAGCACTGGGGCCAGAT
GGTAAGCCCTCCCGTATCGTAGTTATCTACACGACGGGGAGTCAGGCAACTATGGATGAACGAAATAGACAGATCGCTGAGATAGGTGCCTCACTGATTA
AGCATTGGTAACTCGAGCTGATCCTTCAAGAGCTCGCTTGGACTCCTGTTGATAGATCCAGTAATGACCTCAGAACTCCATCTGGATTTGTTCAGAACGC
TCGGTTGCCGCCGGGCGTTTTTTATTGGTGAGAATTTACAACTTATATCGTATGGGGCTGACTTCAGGTGCTACATTTGAAGAGATAAATTGCACTGAAA

TCTAGAAATATTTTATCTGATTAATAAGATGATCTTCTTGAGATCGTTTTGGTCTGCGCGTAATCTCTTGCTCTGAAAACGAAAAAACCGCCTTGCAGGG
CGGTTTTTCGAAGGTTCTCTGAGCTACCAACTCTTTGAACCGAGGTAACTGGCTTGGAGGAGCGCAGTCACCAAAACTTGTCCTTTCAGTTTAGCCTTAA
CCGGCGCATGACTTCAAGACTAACTCCTCTAAATCAATTACCAGTGGCTGCTGCCAGTGGTGCTTTTGCATGTCTTTCCGGGTTGGACTCAAGACGATAG
TTACCGGATAAGGCGCAGCGGTCGGACTGAACGGGGGGTTCGTGCATACAGTCCAGCTTGGAGCGAACTGCCTACCCGGAACTGAGTGTCAGGCGTGGAA
TGAGACAAACGCGGCCATAACAGCGGAATGACACCGGTAAACCGAAAGGCAGGAACAGGAGAGCGCACGAGGGAGCCGCCAGGGGGAAACGCCTGGTATC
TTTATAGTCCTGTCGGGTTTCGCCACCACTGATTTGAGCGTCAGATT

**pCM212**

TCGTGATGCTTGTCAGGGGGGCGGAGCCTATGGAAAAACGGCTTTGCCGCGGCCCTCTCACTTCCCTGTTAAGTATCTTCCTGGCATCTTCCAGGAAATC
TCCGCCCCGTTCGTAAGCCATTTCCGCTCGCCGCAGTCGAACGGGCGCGCCTTGACAGCTAGCTCAGTCCTAGGTATTGTGCTAGCTTACGCCGATTCTG
CCGGGGGGAATTATAATGAGCGAACTGATCAAAGAGAACATGCACATGAAGCTGTACATGGAGGGTACCGTGGATAATCACCACTTTAAGTGTACTTCTG
AGGGCGAGGGTAAGCCGTATGAAGGGACTCAAACGATGCGTATTAAAGTAGTGGAGGGTGGCCCACTGCCGTTTGCTTTCGATATTCTGGCGACGAGCTT
TCTGTATGGTAGCAAAACGTTTATAAACCACACTCAGGGCATTCCGGATTTCTTTAAACAAAGCTTTCCGGAAGGTTTTACCTGGGAGCGTGTGACTACG
TATGAAGATGGTGGTGTGTTGACTGCTACTCAAGATACTTCACTGCAGGACGGCTGTCTGATCTATAACGTGAAGATTCGTGGCGTGAACTTTACGAGCA
ATGGGCCGGTAATGCAAAAAAAAACCCTGGGTTGGGAAGCGTTCACGGAAACTCTGTATCCGGCTGACGGCGGCCTGGAGGGCCGTAACGATATGGCACT
GAAGCTGGTTGGTGGCAGCCACCTGATCGCGAATATCAAAACGACTTATCGCTCTAAAAAACCGGCGAAAAATCTGAAGATGCCGGGTGTTTATTATGTT
GACTATCGTCTGGAACGCATTAAAGAAGCGAATAATGAAACTTACGTGGAGCAACACGAGGTTGCAGTGGCGCGCTATTGCGACTTGCCTTCAAAGCTGG
GTCACAAACTGAATTAATAAGGAAACACAGAAAAAAGCCCGCACCTGACAGTGCGGGCTTTTTTTTTCGACCAAAGGGACAATGTCCTGAAGAATATTTA
AGAAAAAAGCACCCCTCGTCGCCTAGAATTACCTACCGCGGTCGACCATACCTTCGAGACGAACAATAAGGCCGCAAATCGCGGCCTTCGTTATTGATAA
CAGCAGGTAAAAAGACGCTTATTATGGAGTTCGACATGGCAATAACGCGACGAATCTACGTCATGACGAGAATAGTATAAACAAAACTGCTGACGGCAGA
AGCGTCTCGGTACCAAATTCCAGAAAAGAGACGCTTTCGAGCGTCTTTTTTCGTTTTGGTCCACGCTGATGAATATTCTACGGAATTGCCATAGGCGTTG
AACGCTACACGGACGATACGAATTTATGTATAGAGCGTTCAGCCAAAAAACTTAAGACCGCCGGTCTTGTCCACTACCTTGCAGTAATGCGGTGGACAGG
ATCGGCGGTTTTCTTTTCTCTTCTCAATTCGCGACCCAGGATGAGGTCGCCAAAACTACTAGATTTGGGCGCGCCTTGACGGCTAGCTCAGTCCTAGGTA
CAGTGCTAGCTTAATCTTGTCAAACAGTGGCTACGTGAACAGATGCGTAAAGGAGAAGAACTGTTCACTGGAGTTGTCCCAATTCTTGTTGAATTAGATG
GTGATGTTAATGGGCACAAATTTTCTGTGAGTGGAGAGGGTGAAGGTGATGCGACATACGGCAAACTGACCCTTAAATTTATTTGTACTACCGGCAAGCT
ACCTGTTCCTTGGCCGACACTTGTCACTACTTTCGGTTATGGTGTTCAATGTTTTGCGAGATACCCAGATCATATGAAACAGCATGACTTTTTCAAAAGT
GCCATGCCCGAAGGTTATGTACAGGAAAGAACTATATTTTTCAAAGATGACGGGAACTACAAAACACGTGCTGAAGTTAAATTTGAGGGTGATACTCTGG
TAAATAGAATCGAATTAAAAGGTATTGATTTTAAAGAAGATGGCAACATTCTTGGTCACAAGCTGGAATACAACTACAACTCACACAATGTTTACATCAT
GGCAGACAAACAAAAAAATGGCATCAAAGTTAACTTCAAAATTCGACACAACATTGAAGATGGAAGCGTTCAACTGGCAGACCATTATCAACAGAATACT
CCGATTGGCGATGGCCCTGTACTCTTACCAGACAACCATTACCTGTCCACACAATCTGCCCTTTCGAAAGATCCGAACGAAAAGAGAGATCACATGGTCC
TGCTTGAGTTTGTAACAGCTGCTGGGATTACACATGGTATGGATGAGCTATACAAATAATAACCAATTATTGAAGGCCTCCCTAACGGGGGGCCTTTTTT
TGTTTCTGGTCTCCCCACTGATAGTGCTAGTGTAGATCACTACTAGAGCCAGGCATCAAATAAAACGAAAGGCTCAGTCGAAAGACTGGGCCTTTCGTTT
TATCTGTTGTTTGTCGGTGAACGCTCTCTACTAGAGTCACACTGGCTCACCTTCGGGTGGGCCTTTCTGCGTTTATATACTAGAGGGAATCGTACGGTCG
ATCCATCTGACTATCGCCTACTTCATTGACAGCTAGCTCAGTCCTAGGTATAATGCTAGCCTAATTCTACATAGAGTGAGGTAATCTAATGGCTTCCTCC
GAAGACGTTATCAAAGAGTTCATGCGTTTCAAAGTTCGTATGGAAGGTTCCGTTAACGGTCACGAGTTCGAAATCGAAGGTGAAGGTGAAGGTCGTCCGT
ACGAAGGTACCCAGACCGCTAAACTGAAAGTTACCAAAGGTGGTCCGCTGCCGTTCGCTTGGGACATCCTGTCCCCGCAGTTCCAGTACGGTTCCAAAGC
TTACGTTAAACACCCGGCTGACATCCCGGACTACCTGAAACTGTCCTTCCCGGAAGGTTTCAAATGGGAACGTGTTATGAACTTCGAAGACGGTGGTGTT
GTTACCGTTACCCAGGACTCCTCCCTGCAAGACGGTGAGTTCATCTACAAAGTTAAACTGCGTGGTACCAACTTCCCGTCCGACGGTCCGGTTATGCAGA
AAAAAACCATGGGTTGGGAAGCTTCCACCGAACGTATGTACCCGGAAGACGGTGCTCTGAAAGGTGAAATCAAAATGCGTCTGAAACTGAAAGACGGTGG
TCACTACGACGCTGAAGTTAAAACCACCTACATGGCTAAAAAACCGGTTCAGCTGCCGGGTGCTTACAAAACCGACATCAAACTGGACATCACCTCCCAC
AACGAAGACTACACCATCGTTGAACAGTACGAACGTGCTGAAGGTCGTCACTCCACCGGTGCTTAATAACGCTGATAGTGCTAGTGTAGATCGCTACTAG
AGTACTAGCTCTAGATGCGGCCGCGAATTCCAGAAATCATCCTTAGCGAAAGCTAAGGATTTTTTTTATCTGAAATGCCTCTGTAACAGAGCATTAGCGC
AAGGTGATTTTTGTCTTCTTGCGCTAATTTTTTGGCGTCTCCACTGTGCGCGGAACCCCTATTTGTTTATTTTTCTAAATACATTCAAATATGTATCCGC
TCATGAGACAATAACCCTGATAAATGCTTCAATAATATTGAAAAAGGAAGAGTATGAGTATTCAACATTTCCGTGTCGCCCTTATTCCCTTTTTTGCGGC
ATTTTGCCTTCCTGTTTTTGCTCACCCAGAAACGCTGGTGAAAGTAAAAGATGCTGAAGATCAGTTGGGTGCACGAGTGGGTTACATCGAACTGGATCTC
AACAGCGGTAAGATCCTTGAGAGTTTTCGCCCCGAAGAACGTTTTCCAATGATGAGCACTTTTAAAGTTCTGCTATGTGGCGCGGTATTATCCCGTATTG
ACGCCGGGCAAGAGCAACTCGGTCGCCGCATACACTATTCTCAGAATGACTTGGTTGAGTACTCACCAGTCACAGAAAAGCATCTTACGGATGGCATGAC
AGTAAGAGAATTATGCAGTGCTGCCATAACCATGAGTGATAACACTGCGGCCAACTTACTTCTGACAACGATCGGAGGACCGAAGGAGCTAACCGCTTTT
TTGCACAACATGGGGGATCATGTAACTCGCCTTGATCGTTGGGAACCGGAGCTGAATGAAGCCATACCAAACGACGAGCGTGACACCACGATGCCTGTAG
CAATGGCAACAACGTTGCGCAAACTATTAACTGGCGAACTACTTACTCTAGCTTCCCGGCAACAATTAATAGACTGGATGGAGGCGGATAAAGTTGCAGG
ACCACTTCTGCGCTCGGCCCTTCCGGCTGGCTGGTTTATTGCTGATAAATCTGGAGCCGGTGAGCGTGGGTCTCGCGGTATCATTGCAGCACTGGGGCCA
GATGGTAAGCCCTCCCGTATCGTAGTTATCTACACGACGGGGAGTCAGGCAACTATGGATGAACGAAATAGACAGATCGCTGAGATAGGTGCCTCACTGA
TTAAGCATTGGTAACTCGAGCTGATCCTTCAAGAGCTCGCTTGGACTCCTGTTGATAGATCCAGTAATGACCTCAGAACTCCATCTGGATTTGTTCAGAA
CGCTCGGTTGCCGCCGGGCGTTTTTTATTGGTGAGAATTTACAACTTATATCGTATGGGGCTGACTTCAGGTGCTACATTTGAAGAGATAAATTGCACTG
AAATCTAGAAATATTTTATCTGATTAATAAGATGATCTTCTTGAGATCGTTTTGGTCTGCGCGTAATCTCTTGCTCTGAAAACGAAAAAACCGCCTTGCA
GGGCGGTTTTTCGAAGGTTCTCTGAGCTACCAACTCTTTGAACCGAGGTAACTGGCTTGGAGGAGCGCAGTCACCAAAACTTGTCCTTTCAGTTTAGCCT
TAACCGGCGCATGACTTCAAGACTAACTCCTCTAAATCAATTACCAGTGGCTGCTGCCAGTGGTGCTTTTGCATGTCTTTCCGGGTTGGACTCAAGACGA
TAGTTACCGGATAAGGCGCAGCGGTCGGACTGAACGGGGGGTTCGTGCATACAGTCCAGCTTGGAGCGAACTGCCTACCCGGAACTGAGTGTCAGGCGTG
GAATGAGACAAACGCGGCCATAACAGCGGAATGACACCGGTAAACCGAAAGGCAGGAACAGGAGAGCGCACGAGGGAGCCGCCAGGGGGAAACGCCTGGT
ATCTTTATAGTCCTGTCGGGTTTCGCCACCACTGATTTGAGCGTCAGATT

**pCM213**

TCGTGATGCTTGTCAGGGGGGCGGAGCCTATGGAAAAACGGCTTTGCCGCGGCCCTCTCACTTCCCTGTTAAGTATCTTCCTGGCATCTTCCAGGAAATC
TCCGCCCCGTTCGTAAGCCATTTCCGCTCGCCGCAGTCGAACGGGCGCGCCTTGACAGCTAGCTCAGTCCTAGGTATTGTGCTAGCTTACGTCAACCCAC
CCTAGGAGAGGATAAGATGAGCGAACTGATCAAAGAGAACATGCACATGAAGCTGTACATGGAGGGTACCGTGGATAATCACCACTTTAAGTGTACTTCT
GAGGGCGAGGGTAAGCCGTATGAAGGGACTCAAACGATGCGTATTAAAGTAGTGGAGGGTGGCCCACTGCCGTTTGCTTTCGATATTCTGGCGACGAGCT
TTCTGTATGGTAGCAAAACGTTTATAAACCACACTCAGGGCATTCCGGATTTCTTTAAACAAAGCTTTCCGGAAGGTTTTACCTGGGAGCGTGTGACTAC
GTATGAAGATGGTGGTGTGTTGACTGCTACTCAAGATACTTCACTGCAGGACGGCTGTCTGATCTATAACGTGAAGATTCGTGGCGTGAACTTTACGAGC
AATGGGCCGGTAATGCAAAAAAAAACCCTGGGTTGGGAAGCGTTCACGGAAACTCTGTATCCGGCTGACGGCGGCCTGGAGGGCCGTAACGATATGGCAC
TGAAGCTGGTTGGTGGCAGCCACCTGATCGCGAATATCAAAACGACTTATCGCTCTAAAAAACCGGCGAAAAATCTGAAGATGCCGGGTGTTTATTATGT
TGACTATCGTCTGGAACGCATTAAAGAAGCGAATAATGAAACTTACGTGGAGCAACACGAGGTTGCAGTGGCGCGCTATTGCGACTTGCCTTCAAAGCTG
GGTCACAAACTGAATTAATAAGGAAACACAGAAAAAAGCCCGCACCTGACAGTGCGGGCTTTTTTTTTCGACCAAAGGGACAATGTCCTGAAGAATATTT
AAGAAAAAAGCACCCCTCGTCGCCTAGAATTACCTACCGCGGTCGACCATACCTTCGAGACGAACAATAAGGCCGCAAATCGCGGCCTTCGTTATTGATA
ACAGCAGGTAAAAAGACGCTTATTATGGAGTTCGACATGGCAATAACGCGACGAATCTACGTCATGACGAGAATAGTATAAACAAAACTGCTGACGGCAG
AAGCGTCTCGGTACCAAATTCCAGAAAAGAGACGCTTTCGAGCGTCTTTTTTCGTTTTGGTCCACGCTGATGAATATTCTACGGAATTGCCATAGGCGTT
GAACGCTACACGGACGATACGAATTTATGTATAGAGCGTTCAGCCAAAAAACTTAAGACCGCCGGTCTTGTCCACTACCTTGCAGTAATGCGGTGGACAG
GATCGGCGGTTTTCTTTTCTCTTCTCAATTCGCGACCCAGGATGAGGTCGCCAAAACTACTAGATTTGGGCGCGCCTTGACGGCTAGCTCAGTCCTAGGT
ACAGTGCTAGCTTAATCTTGTCAAACAGTGGCTACGTGAACAGATGCGTAAAGGAGAAGAACTGTTCACTGGAGTTGTCCCAATTCTTGTTGAATTAGAT
GGTGATGTTAATGGGCACAAATTTTCTGTGAGTGGAGAGGGTGAAGGTGATGCGACATACGGCAAACTGACCCTTAAATTTATTTGTACTACCGGCAAGC
TACCTGTTCCTTGGCCGACACTTGTCACTACTTTCGGTTATGGTGTTCAATGTTTTGCGAGATACCCAGATCATATGAAACAGCATGACTTTTTCAAAAG
TGCCATGCCCGAAGGTTATGTACAGGAAAGAACTATATTTTTCAAAGATGACGGGAACTACAAAACACGTGCTGAAGTTAAATTTGAGGGTGATACTCTG

GTAAATAGAATCGAATTAAAAGGTATTGATTTTAAAGAAGATGGCAACATTCTTGGTCACAAGCTGGAATACAACTACAACTCACACAATGTTTACATCA
TGGCAGACAAACAAAAAAATGGCATCAAAGTTAACTTCAAAATTCGACACAACATTGAAGATGGAAGCGTTCAACTGGCAGACCATTATCAACAGAATAC
TCCGATTGGCGATGGCCCTGTACTCTTACCAGACAACCATTACCTGTCCACACAATCTGCCCTTTCGAAAGATCCGAACGAAAAGAGAGATCACATGGTC
CTGCTTGAGTTTGTAACAGCTGCTGGGATTACACATGGTATGGATGAGCTATACAAATAATAACCAATTATTGAAGGCCTCCCTAACGGGGGGCCTTTTT
TTGTTTCTGGTCTCCCCACTGATAGTGCTAGTGTAGATCACTACTAGAGCCAGGCATCAAATAAAACGAAAGGCTCAGTCGAAAGACTGGGCCTTTCGTT
TTATCTGTTGTTTGTCGGTGAACGCTCTCTACTAGAGTCACACTGGCTCACCTTCGGGTGGGCCTTTCTGCGTTTATATACTAGAGGGAATCGTACGGTC
GATCCATCTGACTATCGCCTACTTCATTGACAGCTAGCTCAGTCCTAGGTATAATGCTAGCCTAATTCTACATAGAGTGAGGTAATCTAATGGCTTCCTC
CGAAGACGTTATCAAAGAGTTCATGCGTTTCAAAGTTCGTATGGAAGGTTCCGTTAACGGTCACGAGTTCGAAATCGAAGGTGAAGGTGAAGGTCGTCCG
TACGAAGGTACCCAGACCGCTAAACTGAAAGTTACCAAAGGTGGTCCGCTGCCGTTCGCTTGGGACATCCTGTCCCCGCAGTTCCAGTACGGTTCCAAAG
CTTACGTTAAACACCCGGCTGACATCCCGGACTACCTGAAACTGTCCTTCCCGGAAGGTTTCAAATGGGAACGTGTTATGAACTTCGAAGACGGTGGTGT
TGTTACCGTTACCCAGGACTCCTCCCTGCAAGACGGTGAGTTCATCTACAAAGTTAAACTGCGTGGTACCAACTTCCCGTCCGACGGTCCGGTTATGCAG
AAAAAAACCATGGGTTGGGAAGCTTCCACCGAACGTATGTACCCGGAAGACGGTGCTCTGAAAGGTGAAATCAAAATGCGTCTGAAACTGAAAGACGGTG
GTCACTACGACGCTGAAGTTAAAACCACCTACATGGCTAAAAAACCGGTTCAGCTGCCGGGTGCTTACAAAACCGACATCAAACTGGACATCACCTCCCA
CAACGAAGACTACACCATCGTTGAACAGTACGAACGTGCTGAAGGTCGTCACTCCACCGGTGCTTAATAACGCTGATAGTGCTAGTGTAGATCGCTACTA
GAGTACTAGCTCTAGATGCGGCCGCGAATTCCAGAAATCATCCTTAGCGAAAGCTAAGGATTTTTTTTATCTGAAATGCCTCTGTAACAGAGCATTAGCG
CAAGGTGATTTTTGTCTTCTTGCGCTAATTTTTTGGCGTCTCCACTGTGCGCGGAACCCCTATTTGTTTATTTTTCTAAATACATTCAAATATGTATCCG
CTCATGAGACAATAACCCTGATAAATGCTTCAATAATATTGAAAAAGGAAGAGTATGAGTATTCAACATTTCCGTGTCGCCCTTATTCCCTTTTTTGCGG
CATTTTGCCTTCCTGTTTTTGCTCACCCAGAAACGCTGGTGAAAGTAAAAGATGCTGAAGATCAGTTGGGTGCACGAGTGGGTTACATCGAACTGGATCT
CAACAGCGGTAAGATCCTTGAGAGTTTTCGCCCCGAAGAACGTTTTCCAATGATGAGCACTTTTAAAGTTCTGCTATGTGGCGCGGTATTATCCCGTATT
GACGCCGGGCAAGAGCAACTCGGTCGCCGCATACACTATTCTCAGAATGACTTGGTTGAGTACTCACCAGTCACAGAAAAGCATCTTACGGATGGCATGA
CAGTAAGAGAATTATGCAGTGCTGCCATAACCATGAGTGATAACACTGCGGCCAACTTACTTCTGACAACGATCGGAGGACCGAAGGAGCTAACCGCTTT
TTTGCACAACATGGGGGATCATGTAACTCGCCTTGATCGTTGGGAACCGGAGCTGAATGAAGCCATACCAAACGACGAGCGTGACACCACGATGCCTGTA
GCAATGGCAACAACGTTGCGCAAACTATTAACTGGCGAACTACTTACTCTAGCTTCCCGGCAACAATTAATAGACTGGATGGAGGCGGATAAAGTTGCAG
GACCACTTCTGCGCTCGGCCCTTCCGGCTGGCTGGTTTATTGCTGATAAATCTGGAGCCGGTGAGCGTGGGTCTCGCGGTATCATTGCAGCACTGGGGCC
AGATGGTAAGCCCTCCCGTATCGTAGTTATCTACACGACGGGGAGTCAGGCAACTATGGATGAACGAAATAGACAGATCGCTGAGATAGGTGCCTCACTG
ATTAAGCATTGGTAACTCGAGCTGATCCTTCAAGAGCTCGCTTGGACTCCTGTTGATAGATCCAGTAATGACCTCAGAACTCCATCTGGATTTGTTCAGA
ACGCTCGGTTGCCGCCGGGCGTTTTTTATTGGTGAGAATTTACAACTTATATCGTATGGGGCTGACTTCAGGTGCTACATTTGAAGAGATAAATTGCACT
GAAATCTAGAAATATTTTATCTGATTAATAAGATGATCTTCTTGAGATCGTTTTGGTCTGCGCGTAATCTCTTGCTCTGAAAACGAAAAAACCGCCTTGC
AGGGCGGTTTTTCGAAGGTTCTCTGAGCTACCAACTCTTTGAACCGAGGTAACTGGCTTGGAGGAGCGCAGTCACCAAAACTTGTCCTTTCAGTTTAGCC
TTAACCGGCGCATGACTTCAAGACTAACTCCTCTAAATCAATTACCAGTGGCTGCTGCCAGTGGTGCTTTTGCATGTCTTTCCGGGTTGGACTCAAGACG
ATAGTTACCGGATAAGGCGCAGCGGTCGGACTGAACGGGGGGTTCGTGCATACAGTCCAGCTTGGAGCGAACTGCCTACCCGGAACTGAGTGTCAGGCGT
GGAATGAGACAAACGCGGCCATAACAGCGGAATGACACCGGTAAACCGAAAGGCAGGAACAGGAGAGCGCACGAGGGAGCCGCCAGGGGGAAACGCCTGG
TATCTTTATAGTCCTGTCGGGTTTCGCCACCACTGATTTGAGCGTCAGATT

**pCM220**

TCGTGATGCTTGTCAGGGGGGCGGAGCCTATGGAAAAACGGCTTTGCCGCGGCCCTCTCACTTCCCTGTTAAGTATCTTCCTGGCATCTTCCAGGAAATC
TCCGCCCCGTTCGTAAGCCATTTCCGCTCGCCGCAGTCGAACGGGCGCGCCTTGACAGCTAGCTCAGTCCTAGGTATTGTGCTAGCTTACGAGCGATGGG
CTCACTGTTGAGCCTCAATGAGCGAACTGATCAAAGAGAACATGCACATGAAGCTGTACATGGAGGGTACCGTGGATAATCACCACTTTAAGTGTACTTC
TGAGGGCGAGGGTAAGCCGTATGAAGGGACTCAAACGATGCGTATTAAAGTAGTGGAGGGTGGCCCACTGCCGTTTGCTTTCGATATTCTGGCGACGAGC
TTTCTGTATGGTAGCAAAACGTTTATAAACCACACTCAGGGCATTCCGGATTTCTTTAAACAAAGCTTTCCGGAAGGTTTTACCTGGGAGCGTGTGACTA
CGTATGAAGATGGTGGTGTGTTGACTGCTACTCAAGATACTTCACTGCAGGACGGCTGTCTGATCTATAACGTGAAGATTCGTGGCGTGAACTTTACGAG
CAATGGGCCGGTAATGCAAAAAAAAACCCTGGGTTGGGAAGCGTTCACGGAAACTCTGTATCCGGCTGACGGCGGCCTGGAGGGCCGTAACGATATGGCA
CTGAAGCTGGTTGGTGGCAGCCACCTGATCGCGAATATCAAAACGACTTATCGCTCTAAAAAACCGGCGAAAAATCTGAAGATGCCGGGTGTTTATTATG
TTGACTATCGTCTGGAACGCATTAAAGAAGCGAATAATGAAACTTACGTGGAGCAACACGAGGTTGCAGTGGCGCGCTATTGCGACTTGCCTTCAAAGCT
GGGTCACAAACTGAATTAATAAGGAAACACAGAAAAAAGCCCGCACCTGACAGTGCGGGCTTTTTTTTTCGACCAAAGGGACAATGTCCTGAAGAATATT
TAAGAAAAAAGCACCCCTCGTCGCCTAGAATTACCTACCGCGGTCGACCATACCTTCGAGACGAACAATAAGGCCGCAAATCGCGGCCTTCGTTATTGAT
AACAGCAGGTAAAAAGACGCTTATTATGGAGTTCGACATGGCAATAACGCGACGAATCTACGTCATGACGAGAATAGTATAAACAAAACTGCTGACGGCA
GAAGCGTCTCGGTACCAAATTCCAGAAAAGAGACGCTTTCGAGCGTCTTTTTTCGTTTTGGTCCACGCTGATGAATATTCTACGGAATTGCCATAGGCGT
TGAACGCTACACGGACGATACGAATTTATGTATAGAGCGTTCAGCCAAAAAACTTAAGACCGCCGGTCTTGTCCACTACCTTGCAGTAATGCGGTGGACA
GGATCGGCGGTTTTCTTTTCTCTTCTCAATTCGCGACCCAGGATGAGGTCGCCAAAACTACTAGATTTGGGCGCGCCTTGACGGCTAGCTCAGTCCTAGG
TACAGTGCTAGCTTAATGCGAGAGGCTGGTTAAGTGGGTGTTGATGCGTAAAGGAGAAGAACTGTTCACTGGAGTTGTCCCAATTCTTGTTGAATTAGAT
GGTGATGTTAATGGGCACAAATTTTCTGTGAGTGGAGAGGGTGAAGGTGATGCGACATACGGCAAACTGACCCTTAAATTTATTTGTACTACCGGCAAGC
TACCTGTTCCTTGGCCGACACTTGTCACTACTTTCGGTTATGGTGTTCAATGTTTTGCGAGATACCCAGATCATATGAAACAGCATGACTTTTTCAAAAG
TGCCATGCCCGAAGGTTATGTACAGGAAAGAACTATATTTTTCAAAGATGACGGGAACTACAAAACACGTGCTGAAGTTAAATTTGAGGGTGATACTCTG
GTAAATAGAATCGAATTAAAAGGTATTGATTTTAAAGAAGATGGCAACATTCTTGGTCACAAGCTGGAATACAACTACAACTCACACAATGTTTACATCA
TGGCAGACAAACAAAAAAATGGCATCAAAGTTAACTTCAAAATTCGACACAACATTGAAGATGGAAGCGTTCAACTGGCAGACCATTATCAACAGAATAC
TCCGATTGGCGATGGCCCTGTACTCTTACCAGACAACCATTACCTGTCCACACAATCTGCCCTTTCGAAAGATCCGAACGAAAAGAGAGATCACATGGTC
CTGCTTGAGTTTGTAACAGCTGCTGGGATTACACATGGTATGGATGAGCTATACAAATAATAACCAATTATTGAAGGCCTCCCTAACGGGGGGCCTTTTT
TTGTTTCTGGTCTCCCCACTGATAGTGCTAGTGTAGATCACTACTAGAGCCAGGCATCAAATAAAACGAAAGGCTCAGTCGAAAGACTGGGCCTTTCGTT
TTATCTGTTGTTTGTCGGTGAACGCTCTCTACTAGAGTCACACTGGCTCACCTTCGGGTGGGCCTTTCTGCGTTTATATACTAGAGGGAATCGTACGGTC
GATCCATCTGACTATCGCCTACTTCATTGACAGCTAGCTCAGTCCTAGGTATAATGCTAGCTCTAGAGAAAGAGGAGAAATACTAGATGGCTTCCTCCGA
AGACGTTATCAAAGAGTTCATGCGTTTCAAAGTTCGTATGGAAGGTTCCGTTAACGGTCACGAGTTCGAAATCGAAGGTGAAGGTGAAGGTCGTCCGTAC
GAAGGTACCCAGACCGCTAAACTGAAAGTTACCAAAGGTGGTCCGCTGCCGTTCGCTTGGGACATCCTGTCCCCGCAGTTCCAGTACGGTTCCAAAGCTT
ACGTTAAACACCCGGCTGACATCCCGGACTACCTGAAACTGTCCTTCCCGGAAGGTTTCAAATGGGAACGTGTTATGAACTTCGAAGACGGTGGTGTTGT
TACCGTTACCCAGGACTCCTCCCTGCAAGACGGTGAGTTCATCTACAAAGTTAAACTGCGTGGTACCAACTTCCCGTCCGACGGTCCGGTTATGCAGAAA
AAAACCATGGGTTGGGAAGCTTCCACCGAACGTATGTACCCGGAAGACGGTGCTCTGAAAGGTGAAATCAAAATGCGTCTGAAACTGAAAGACGGTGGTC
ACTACGACGCTGAAGTTAAAACCACCTACATGGCTAAAAAACCGGTTCAGCTGCCGGGTGCTTACAAAACCGACATCAAACTGGACATCACCTCCCACAA
CGAAGACTACACCATCGTTGAACAGTACGAACGTGCTGAAGGTCGTCACTCCACCGGTGCTTAATAACGCTGATAGTGCTAGTGTAGATCGCTACTAGAG
TACTAGCTCTAGATGCGGCCGCGAATTCCAGAAATCATCCTTAGCGAAAGCTAAGGATTTTTTTTATCTGAAATGCCTCTGTAACAGAGCATTAGCGCAA
GGTGATTTTTGTCTTCTTGCGCTAATTTTTTGGCGTCTCCACTGTGCGCGGAACCCCTATTTGTTTATTTTTCTAAATACATTCAAATATGTATCCGCTC
ATGAGACAATAACCCTGATAAATGCTTCAATAATATTGAAAAAGGAAGAGTATGAGTATTCAACATTTCCGTGTCGCCCTTATTCCCTTTTTTGCGGCAT
TTTGCCTTCCTGTTTTTGCTCACCCAGAAACGCTGGTGAAAGTAAAAGATGCTGAAGATCAGTTGGGTGCACGAGTGGGTTACATCGAACTGGATCTCAA
CAGCGGTAAGATCCTTGAGAGTTTTCGCCCCGAAGAACGTTTTCCAATGATGAGCACTTTTAAAGTTCTGCTATGTGGCGCGGTATTATCCCGTATTGAC
GCCGGGCAAGAGCAACTCGGTCGCCGCATACACTATTCTCAGAATGACTTGGTTGAGTACTCACCAGTCACAGAAAAGCATCTTACGGATGGCATGACAG
TAAGAGAATTATGCAGTGCTGCCATAACCATGAGTGATAACACTGCGGCCAACTTACTTCTGACAACGATCGGAGGACCGAAGGAGCTAACCGCTTTTTT
GCACAACATGGGGGATCATGTAACTCGCCTTGATCGTTGGGAACCGGAGCTGAATGAAGCCATACCAAACGACGAGCGTGACACCACGATGCCTGTAGCA
ATGGCAACAACGTTGCGCAAACTATTAACTGGCGAACTACTTACTCTAGCTTCCCGGCAACAATTAATAGACTGGATGGAGGCGGATAAAGTTGCAGGAC
CACTTCTGCGCTCGGCCCTTCCGGCTGGCTGGTTTATTGCTGATAAATCTGGAGCCGGTGAGCGTGGGTCTCGCGGTATCATTGCAGCACTGGGGCCAGA
TGGTAAGCCCTCCCGTATCGTAGTTATCTACACGACGGGGAGTCAGGCAACTATGGATGAACGAAATAGACAGATCGCTGAGATAGGTGCCTCACTGATT
AAGCATTGGTAACTCGAGCTGATCCTTCAAGAGCTCGCTTGGACTCCTGTTGATAGATCCAGTAATGACCTCAGAACTCCATCTGGATTTGTTCAGAACG

CTCGGTTGCCGCCGGGCGTTTTTTATTGGTGAGAATTTACAACTTATATCGTATGGGGCTGACTTCAGGTGCTACATTTGAAGAGATAAATTGCACTGAA
ATCTAGAAATATTTTATCTGATTAATAAGATGATCTTCTTGAGATCGTTTTGGTCTGCGCGTAATCTCTTGCTCTGAAAACGAAAAAACCGCCTTGCAGG
GCGGTTTTTCGAAGGTTCTCTGAGCTACCAACTCTTTGAACCGAGGTAACTGGCTTGGAGGAGCGCAGTCACCAAAACTTGTCCTTTCAGTTTAGCCTTA
ACCGGCGCATGACTTCAAGACTAACTCCTCTAAATCAATTACCAGTGGCTGCTGCCAGTGGTGCTTTTGCATGTCTTTCCGGGTTGGACTCAAGACGATA
GTTACCGGATAAGGCGCAGCGGTCGGACTGAACGGGGGGTTCGTGCATACAGTCCAGCTTGGAGCGAACTGCCTACCCGGAACTGAGTGTCAGGCGTGGA
ATGAGACAAACGCGGCCATAACAGCGGAATGACACCGGTAAACCGAAAGGCAGGAACAGGAGAGCGCACGAGGGAGCCGCCAGGGGGAAACGCCTGGTAT
CTTTATAGTCCTGTCGGGTTTCGCCACCACTGATTTGAGCGTCAGATT

**pCM221**

TCGTGATGCTTGTCAGGGGGGCGGAGCCTATGGAAAAACGGCTTTGCCGCGGCCCTCTCACTTCCCTGTTAAGTATCTTCCTGGCATCTTCCAGGAAATC
TCCGCCCCGTTCGTAAGCCATTTCCGCTCGCCGCAGTCGAACGGGCGCGCCTTGACAGCTAGCTCAGTCCTAGGTATTGTGCTAGCTTACGAGCGATGGG
CTCACTGTTGAGCCTCAATGAGCGAACTGATCAAAGAGAACATGCACATGAAGCTGTACATGGAGGGTACCGTGGATAATCACCACTTTAAGTGTACTTC
TGAGGGCGAGGGTAAGCCGTATGAAGGGACTCAAACGATGCGTATTAAAGTAGTGGAGGGTGGCCCACTGCCGTTTGCTTTCGATATTCTGGCGACGAGC
TTTCTGTATGGTAGCAAAACGTTTATAAACCACACTCAGGGCATTCCGGATTTCTTTAAACAAAGCTTTCCGGAAGGTTTTACCTGGGAGCGTGTGACTA
CGTATGAAGATGGTGGTGTGTTGACTGCTACTCAAGATACTTCACTGCAGGACGGCTGTCTGATCTATAACGTGAAGATTCGTGGCGTGAACTTTACGAG
CAATGGGCCGGTAATGCAAAAAAAAACCCTGGGTTGGGAAGCGTTCACGGAAACTCTGTATCCGGCTGACGGCGGCCTGGAGGGCCGTAACGATATGGCA
CTGAAGCTGGTTGGTGGCAGCCACCTGATCGCGAATATCAAAACGACTTATCGCTCTAAAAAACCGGCGAAAAATCTGAAGATGCCGGGTGTTTATTATG
TTGACTATCGTCTGGAACGCATTAAAGAAGCGAATAATGAAACTTACGTGGAGCAACACGAGGTTGCAGTGGCGCGCTATTGCGACTTGCCTTCAAAGCT
GGGTCACAAACTGAATTAATAAGGAAACACAGAAAAAAGCCCGCACCTGACAGTGCGGGCTTTTTTTTTCGACCAAAGGGACAATGTCCTGAAGAATATT
TAAGAAAAAAGCACCCCTCGTCGCCTAGAATTACCTACCGCGGTCGACCATACCTTCGAGACGAACAATAAGGCCGCAAATCGCGGCCTTCGTTATTGAT
AACAGCAGGTAAAAAGACGCTTATTATGGAGTTCGACATGGCAATAACGCGACGAATCTACGTCATGACGAGAATAGTATAAACAAAACTGCTGACGGCA
GAAGCGTCTCGGTACCAAATTCCAGAAAAGAGACGCTTTCGAGCGTCTTTTTTCGTTTTGGTCCACGCTGATGAATATTCTACGGAATTGCCATAGGCGT
TGAACGCTACACGGACGATACGAATTTATGTATAGAGCGTTCAGCCAAAAAACTTAAGACCGCCGGTCTTGTCCACTACCTTGCAGTAATGCGGTGGACA
GGATCGGCGGTTTTCTTTTCTCTTCTCAATTCGCGACCCAGGATGAGGTCGCCAAAACTACTAGATTTGGGCGCGCCTTGACGGCTAGCTCAGTCCTAGG
TACAGTGCTAGCTTAATTTCTAATTAGGATAAGGACACCAAGATGCGTAAAGGAGAAGAACTGTTCACTGGAGTTGTCCCAATTCTTGTTGAATTAGATG
GTGATGTTAATGGGCACAAATTTTCTGTGAGTGGAGAGGGTGAAGGTGATGCGACATACGGCAAACTGACCCTTAAATTTATTTGTACTACCGGCAAGCT
ACCTGTTCCTTGGCCGACACTTGTCACTACTTTCGGTTATGGTGTTCAATGTTTTGCGAGATACCCAGATCATATGAAACAGCATGACTTTTTCAAAAGT
GCCATGCCCGAAGGTTATGTACAGGAAAGAACTATATTTTTCAAAGATGACGGGAACTACAAAACACGTGCTGAAGTTAAATTTGAGGGTGATACTCTGG
TAAATAGAATCGAATTAAAAGGTATTGATTTTAAAGAAGATGGCAACATTCTTGGTCACAAGCTGGAATACAACTACAACTCACACAATGTTTACATCAT
GGCAGACAAACAAAAAAATGGCATCAAAGTTAACTTCAAAATTCGACACAACATTGAAGATGGAAGCGTTCAACTGGCAGACCATTATCAACAGAATACT
CCGATTGGCGATGGCCCTGTACTCTTACCAGACAACCATTACCTGTCCACACAATCTGCCCTTTCGAAAGATCCGAACGAAAAGAGAGATCACATGGTCC
TGCTTGAGTTTGTAACAGCTGCTGGGATTACACATGGTATGGATGAGCTATACAAATAATAACCAATTATTGAAGGCCTCCCTAACGGGGGGCCTTTTTT
TGTTTCTGGTCTCCCCACTGATAGTGCTAGTGTAGATCACTACTAGAGCCAGGCATCAAATAAAACGAAAGGCTCAGTCGAAAGACTGGGCCTTTCGTTT
TATCTGTTGTTTGTCGGTGAACGCTCTCTACTAGAGTCACACTGGCTCACCTTCGGGTGGGCCTTTCTGCGTTTATATACTAGAGGGAATCGTACGGTCG
ATCCATCTGACTATCGCCTACTTCATTGACAGCTAGCTCAGTCCTAGGTATAATGCTAGCTCTAGAGAAAGAGGAGAAATACTAGATGGCTTCCTCCGAA
GACGTTATCAAAGAGTTCATGCGTTTCAAAGTTCGTATGGAAGGTTCCGTTAACGGTCACGAGTTCGAAATCGAAGGTGAAGGTGAAGGTCGTCCGTACG
AAGGTACCCAGACCGCTAAACTGAAAGTTACCAAAGGTGGTCCGCTGCCGTTCGCTTGGGACATCCTGTCCCCGCAGTTCCAGTACGGTTCCAAAGCTTA
CGTTAAACACCCGGCTGACATCCCGGACTACCTGAAACTGTCCTTCCCGGAAGGTTTCAAATGGGAACGTGTTATGAACTTCGAAGACGGTGGTGTTGTT
ACCGTTACCCAGGACTCCTCCCTGCAAGACGGTGAGTTCATCTACAAAGTTAAACTGCGTGGTACCAACTTCCCGTCCGACGGTCCGGTTATGCAGAAAA
AAACCATGGGTTGGGAAGCTTCCACCGAACGTATGTACCCGGAAGACGGTGCTCTGAAAGGTGAAATCAAAATGCGTCTGAAACTGAAAGACGGTGGTCA
CTACGACGCTGAAGTTAAAACCACCTACATGGCTAAAAAACCGGTTCAGCTGCCGGGTGCTTACAAAACCGACATCAAACTGGACATCACCTCCCACAAC
GAAGACTACACCATCGTTGAACAGTACGAACGTGCTGAAGGTCGTCACTCCACCGGTGCTTAATAACGCTGATAGTGCTAGTGTAGATCGCTACTAGAGT
ACTAGCTCTAGATGCGGCCGCGAATTCCAGAAATCATCCTTAGCGAAAGCTAAGGATTTTTTTTATCTGAAATGCCTCTGTAACAGAGCATTAGCGCAAG
GTGATTTTTGTCTTCTTGCGCTAATTTTTTGGCGTCTCCACTGTGCGCGGAACCCCTATTTGTTTATTTTTCTAAATACATTCAAATATGTATCCGCTCA
TGAGACAATAACCCTGATAAATGCTTCAATAATATTGAAAAAGGAAGAGTATGAGTATTCAACATTTCCGTGTCGCCCTTATTCCCTTTTTTGCGGCATT
TTGCCTTCCTGTTTTTGCTCACCCAGAAACGCTGGTGAAAGTAAAAGATGCTGAAGATCAGTTGGGTGCACGAGTGGGTTACATCGAACTGGATCTCAAC
AGCGGTAAGATCCTTGAGAGTTTTCGCCCCGAAGAACGTTTTCCAATGATGAGCACTTTTAAAGTTCTGCTATGTGGCGCGGTATTATCCCGTATTGACG
CCGGGCAAGAGCAACTCGGTCGCCGCATACACTATTCTCAGAATGACTTGGTTGAGTACTCACCAGTCACAGAAAAGCATCTTACGGATGGCATGACAGT
AAGAGAATTATGCAGTGCTGCCATAACCATGAGTGATAACACTGCGGCCAACTTACTTCTGACAACGATCGGAGGACCGAAGGAGCTAACCGCTTTTTTG
CACAACATGGGGGATCATGTAACTCGCCTTGATCGTTGGGAACCGGAGCTGAATGAAGCCATACCAAACGACGAGCGTGACACCACGATGCCTGTAGCAA
TGGCAACAACGTTGCGCAAACTATTAACTGGCGAACTACTTACTCTAGCTTCCCGGCAACAATTAATAGACTGGATGGAGGCGGATAAAGTTGCAGGACC
ACTTCTGCGCTCGGCCCTTCCGGCTGGCTGGTTTATTGCTGATAAATCTGGAGCCGGTGAGCGTGGGTCTCGCGGTATCATTGCAGCACTGGGGCCAGAT
GGTAAGCCCTCCCGTATCGTAGTTATCTACACGACGGGGAGTCAGGCAACTATGGATGAACGAAATAGACAGATCGCTGAGATAGGTGCCTCACTGATTA
AGCATTGGTAACTCGAGCTGATCCTTCAAGAGCTCGCTTGGACTCCTGTTGATAGATCCAGTAATGACCTCAGAACTCCATCTGGATTTGTTCAGAACGC
TCGGTTGCCGCCGGGCGTTTTTTATTGGTGAGAATTTACAACTTATATCGTATGGGGCTGACTTCAGGTGCTACATTTGAAGAGATAAATTGCACTGAAA
TCTAGAAATATTTTATCTGATTAATAAGATGATCTTCTTGAGATCGTTTTGGTCTGCGCGTAATCTCTTGCTCTGAAAACGAAAAAACCGCCTTGCAGGG
CGGTTTTTCGAAGGTTCTCTGAGCTACCAACTCTTTGAACCGAGGTAACTGGCTTGGAGGAGCGCAGTCACCAAAACTTGTCCTTTCAGTTTAGCCTTAA
CCGGCGCATGACTTCAAGACTAACTCCTCTAAATCAATTACCAGTGGCTGCTGCCAGTGGTGCTTTTGCATGTCTTTCCGGGTTGGACTCAAGACGATAG
TTACCGGATAAGGCGCAGCGGTCGGACTGAACGGGGGGTTCGTGCATACAGTCCAGCTTGGAGCGAACTGCCTACCCGGAACTGAGTGTCAGGCGTGGAA
TGAGACAAACGCGGCCATAACAGCGGAATGACACCGGTAAACCGAAAGGCAGGAACAGGAGAGCGCACGAGGGAGCCGCCAGGGGGAAACGCCTGGTATC
TTTATAGTCCTGTCGGGTTTCGCCACCACTGATTTGAGCGTCAGATT

**pCM222**

TCGTGATGCTTGTCAGGGGGGCGGAGCCTATGGAAAAACGGCTTTGCCGCGGCCCTCTCACTTCCCTGTTAAGTATCTTCCTGGCATCTTCCAGGAAATC
TCCGCCCCGTTCGTAAGCCATTTCCGCTCGCCGCAGTCGAACGGGCGCGCCTTGACAGCTAGCTCAGTCCTAGGTATTGTGCTAGCTTACGAGCGATGGG
CTCACTGTTGAGCCTCAATGAGCGAACTGATCAAAGAGAACATGCACATGAAGCTGTACATGGAGGGTACCGTGGATAATCACCACTTTAAGTGTACTTC
TGAGGGCGAGGGTAAGCCGTATGAAGGGACTCAAACGATGCGTATTAAAGTAGTGGAGGGTGGCCCACTGCCGTTTGCTTTCGATATTCTGGCGACGAGC
TTTCTGTATGGTAGCAAAACGTTTATAAACCACACTCAGGGCATTCCGGATTTCTTTAAACAAAGCTTTCCGGAAGGTTTTACCTGGGAGCGTGTGACTA
CGTATGAAGATGGTGGTGTGTTGACTGCTACTCAAGATACTTCACTGCAGGACGGCTGTCTGATCTATAACGTGAAGATTCGTGGCGTGAACTTTACGAG
CAATGGGCCGGTAATGCAAAAAAAAACCCTGGGTTGGGAAGCGTTCACGGAAACTCTGTATCCGGCTGACGGCGGCCTGGAGGGCCGTAACGATATGGCA
CTGAAGCTGGTTGGTGGCAGCCACCTGATCGCGAATATCAAAACGACTTATCGCTCTAAAAAACCGGCGAAAAATCTGAAGATGCCGGGTGTTTATTATG
TTGACTATCGTCTGGAACGCATTAAAGAAGCGAATAATGAAACTTACGTGGAGCAACACGAGGTTGCAGTGGCGCGCTATTGCGACTTGCCTTCAAAGCT
GGGTCACAAACTGAATTAATAAGGAAACACAGAAAAAAGCCCGCACCTGACAGTGCGGGCTTTTTTTTTCGACCAAAGGGACAATGTCCTGAAGAATATT
TAAGAAAAAAGCACCCCTCGTCGCCTAGAATTACCTACCGCGGTCGACCATACCTTCGAGACGAACAATAAGGCCGCAAATCGCGGCCTTCGTTATTGAT
AACAGCAGGTAAAAAGACGCTTATTATGGAGTTCGACATGGCAATAACGCGACGAATCTACGTCATGACGAGAATAGTATAAACAAAACTGCTGACGGCA
GAAGCGTCTCGGTACCAAATTCCAGAAAAGAGACGCTTTCGAGCGTCTTTTTTCGTTTTGGTCCACGCTGATGAATATTCTACGGAATTGCCATAGGCGT
TGAACGCTACACGGACGATACGAATTTATGTATAGAGCGTTCAGCCAAAAAACTTAAGACCGCCGGTCTTGTCCACTACCTTGCAGTAATGCGGTGGACA
GGATCGGCGGTTTTCTTTTCTCTTCTCAATTCGCGACCCAGGATGAGGTCGCCAAAACTACTAGATTTGGGCGCGCCTTGACGGCTAGCTCAGTCCTAGG
TACAGTGCTAGCTTAATGCGAGAGGCTGGTTAAGTGGGTGTTGATGCGTAAAGGAGAAGAACTGTTCACTGGAGTTGTCCCAATTCTTGTTGAATTAGAT
GGTGATGTTAATGGGCACAAATTTTCTGTGAGTGGAGAGGGTGAAGGTGATGCGACATACGGCAAACTGACCCTTAAATTTATTTGTACTACCGGCAAGC
TACCTGTTCCTTGGCCGACACTTGTCACTACTTTCGGTTATGGTGTTCAATGTTTTGCGAGATACCCAGATCATATGAAACAGCATGACTTTTTCAAAAG

TGCCATGCCCGAAGGTTATGTACAGGAAAGAACTATATTTTTCAAAGATGACGGGAACTACAAAACACGTGCTGAAGTTAAATTTGAGGGTGATACTCTG
GTAAATAGAATCGAATTAAAAGGTATTGATTTTAAAGAAGATGGCAACATTCTTGGTCACAAGCTGGAATACAACTACAACTCACACAATGTTTACATCA
TGGCAGACAAACAAAAAAATGGCATCAAAGTTAACTTCAAAATTCGACACAACATTGAAGATGGAAGCGTTCAACTGGCAGACCATTATCAACAGAATAC
TCCGATTGGCGATGGCCCTGTACTCTTACCAGACAACCATTACCTGTCCACACAATCTGCCCTTTCGAAAGATCCGAACGAAAAGAGAGATCACATGGTC
CTGCTTGAGTTTGTAACAGCTGCTGGGATTACACATGGTATGGATGAGCTATACAAATAATAACCAATTATTGAAGGCCTCCCTAACGGGGGGCCTTTTT
TTGTTTCTGGTCTCCCCACTGATAGTGCTAGTGTAGATCACTACTAGAGCCAGGCATCAAATAAAACGAAAGGCTCAGTCGAAAGACTGGGCCTTTCGTT
TTATCTGTTGTTTGTCGGTGAACGCTCTCTACTAGAGTCACACTGGCTCACCTTCGGGTGGGCCTTTCTGCGTTTATATACTAGAGGGAATCGTACGGTC
GATCCATCTGACTATCGCCTACTTCATTGACAGCTAGCTCAGTCCTAGGTATAATGCTAGCCTAATTCTACATAGAGTGAGGTAATCTAATGGCTTCCTC
CGAAGACGTTATCAAAGAGTTCATGCGTTTCAAAGTTCGTATGGAAGGTTCCGTTAACGGTCACGAGTTCGAAATCGAAGGTGAAGGTGAAGGTCGTCCG
TACGAAGGTACCCAGACCGCTAAACTGAAAGTTACCAAAGGTGGTCCGCTGCCGTTCGCTTGGGACATCCTGTCCCCGCAGTTCCAGTACGGTTCCAAAG
CTTACGTTAAACACCCGGCTGACATCCCGGACTACCTGAAACTGTCCTTCCCGGAAGGTTTCAAATGGGAACGTGTTATGAACTTCGAAGACGGTGGTGT
TGTTACCGTTACCCAGGACTCCTCCCTGCAAGACGGTGAGTTCATCTACAAAGTTAAACTGCGTGGTACCAACTTCCCGTCCGACGGTCCGGTTATGCAG
AAAAAAACCATGGGTTGGGAAGCTTCCACCGAACGTATGTACCCGGAAGACGGTGCTCTGAAAGGTGAAATCAAAATGCGTCTGAAACTGAAAGACGGTG
GTCACTACGACGCTGAAGTTAAAACCACCTACATGGCTAAAAAACCGGTTCAGCTGCCGGGTGCTTACAAAACCGACATCAAACTGGACATCACCTCCCA
CAACGAAGACTACACCATCGTTGAACAGTACGAACGTGCTGAAGGTCGTCACTCCACCGGTGCTTAATAACGCTGATAGTGCTAGTGTAGATCGCTACTA
GAGTACTAGCTCTAGATGCGGCCGCGAATTCCAGAAATCATCCTTAGCGAAAGCTAAGGATTTTTTTTATCTGAAATGCCTCTGTAACAGAGCATTAGCG
CAAGGTGATTTTTGTCTTCTTGCGCTAATTTTTTGGCGTCTCCACTGTGCGCGGAACCCCTATTTGTTTATTTTTCTAAATACATTCAAATATGTATCCG
CTCATGAGACAATAACCCTGATAAATGCTTCAATAATATTGAAAAAGGAAGAGTATGAGTATTCAACATTTCCGTGTCGCCCTTATTCCCTTTTTTGCGG
CATTTTGCCTTCCTGTTTTTGCTCACCCAGAAACGCTGGTGAAAGTAAAAGATGCTGAAGATCAGTTGGGTGCACGAGTGGGTTACATCGAACTGGATCT
CAACAGCGGTAAGATCCTTGAGAGTTTTCGCCCCGAAGAACGTTTTCCAATGATGAGCACTTTTAAAGTTCTGCTATGTGGCGCGGTATTATCCCGTATT
GACGCCGGGCAAGAGCAACTCGGTCGCCGCATACACTATTCTCAGAATGACTTGGTTGAGTACTCACCAGTCACAGAAAAGCATCTTACGGATGGCATGA
CAGTAAGAGAATTATGCAGTGCTGCCATAACCATGAGTGATAACACTGCGGCCAACTTACTTCTGACAACGATCGGAGGACCGAAGGAGCTAACCGCTTT
TTTGCACAACATGGGGGATCATGTAACTCGCCTTGATCGTTGGGAACCGGAGCTGAATGAAGCCATACCAAACGACGAGCGTGACACCACGATGCCTGTA
GCAATGGCAACAACGTTGCGCAAACTATTAACTGGCGAACTACTTACTCTAGCTTCCCGGCAACAATTAATAGACTGGATGGAGGCGGATAAAGTTGCAG
GACCACTTCTGCGCTCGGCCCTTCCGGCTGGCTGGTTTATTGCTGATAAATCTGGAGCCGGTGAGCGTGGGTCTCGCGGTATCATTGCAGCACTGGGGCC
AGATGGTAAGCCCTCCCGTATCGTAGTTATCTACACGACGGGGAGTCAGGCAACTATGGATGAACGAAATAGACAGATCGCTGAGATAGGTGCCTCACTG
ATTAAGCATTGGTAACTCGAGCTGATCCTTCAAGAGCTCGCTTGGACTCCTGTTGATAGATCCAGTAATGACCTCAGAACTCCATCTGGATTTGTTCAGA
ACGCTCGGTTGCCGCCGGGCGTTTTTTATTGGTGAGAATTTACAACTTATATCGTATGGGGCTGACTTCAGGTGCTACATTTGAAGAGATAAATTGCACT
GAAATCTAGAAATATTTTATCTGATTAATAAGATGATCTTCTTGAGATCGTTTTGGTCTGCGCGTAATCTCTTGCTCTGAAAACGAAAAAACCGCCTTGC
AGGGCGGTTTTTCGAAGGTTCTCTGAGCTACCAACTCTTTGAACCGAGGTAACTGGCTTGGAGGAGCGCAGTCACCAAAACTTGTCCTTTCAGTTTAGCC
TTAACCGGCGCATGACTTCAAGACTAACTCCTCTAAATCAATTACCAGTGGCTGCTGCCAGTGGTGCTTTTGCATGTCTTTCCGGGTTGGACTCAAGACG
ATAGTTACCGGATAAGGCGCAGCGGTCGGACTGAACGGGGGGTTCGTGCATACAGTCCAGCTTGGAGCGAACTGCCTACCCGGAACTGAGTGTCAGGCGT
GGAATGAGACAAACGCGGCCATAACAGCGGAATGACACCGGTAAACCGAAAGGCAGGAACAGGAGAGCGCACGAGGGAGCCGCCAGGGGGAAACGCCTGG
TATCTTTATAGTCCTGTCGGGTTTCGCCACCACTGATTTGAGCGTCAGATT

**pCM223**

TCGTGATGCTTGTCAGGGGGGCGGAGCCTATGGAAAAACGGCTTTGCCGCGGCCCTCTCACTTCCCTGTTAAGTATCTTCCTGGCATCTTCCAGGAAATC
TCCGCCCCGTTCGTAAGCCATTTCCGCTCGCCGCAGTCGAACGGGCGCGCCTTGACAGCTAGCTCAGTCCTAGGTATTGTGCTAGCTTACGAGCGATGGG
CTCACTGTTGAGCCTCAATGAGCGAACTGATCAAAGAGAACATGCACATGAAGCTGTACATGGAGGGTACCGTGGATAATCACCACTTTAAGTGTACTTC
TGAGGGCGAGGGTAAGCCGTATGAAGGGACTCAAACGATGCGTATTAAAGTAGTGGAGGGTGGCCCACTGCCGTTTGCTTTCGATATTCTGGCGACGAGC
TTTCTGTATGGTAGCAAAACGTTTATAAACCACACTCAGGGCATTCCGGATTTCTTTAAACAAAGCTTTCCGGAAGGTTTTACCTGGGAGCGTGTGACTA
CGTATGAAGATGGTGGTGTGTTGACTGCTACTCAAGATACTTCACTGCAGGACGGCTGTCTGATCTATAACGTGAAGATTCGTGGCGTGAACTTTACGAG
CAATGGGCCGGTAATGCAAAAAAAAACCCTGGGTTGGGAAGCGTTCACGGAAACTCTGTATCCGGCTGACGGCGGCCTGGAGGGCCGTAACGATATGGCA
CTGAAGCTGGTTGGTGGCAGCCACCTGATCGCGAATATCAAAACGACTTATCGCTCTAAAAAACCGGCGAAAAATCTGAAGATGCCGGGTGTTTATTATG
TTGACTATCGTCTGGAACGCATTAAAGAAGCGAATAATGAAACTTACGTGGAGCAACACGAGGTTGCAGTGGCGCGCTATTGCGACTTGCCTTCAAAGCT
GGGTCACAAACTGAATTAATAAGGAAACACAGAAAAAAGCCCGCACCTGACAGTGCGGGCTTTTTTTTTCGACCAAAGGGACAATGTCCTGAAGAATATT
TAAGAAAAAAGCACCCCTCGTCGCCTAGAATTACCTACCGCGGTCGACCATACCTTCGAGACGAACAATAAGGCCGCAAATCGCGGCCTTCGTTATTGAT
AACAGCAGGTAAAAAGACGCTTATTATGGAGTTCGACATGGCAATAACGCGACGAATCTACGTCATGACGAGAATAGTATAAACAAAACTGCTGACGGCA
GAAGCGTCTCGGTACCAAATTCCAGAAAAGAGACGCTTTCGAGCGTCTTTTTTCGTTTTGGTCCACGCTGATGAATATTCTACGGAATTGCCATAGGCGT
TGAACGCTACACGGACGATACGAATTTATGTATAGAGCGTTCAGCCAAAAAACTTAAGACCGCCGGTCTTGTCCACTACCTTGCAGTAATGCGGTGGACA
GGATCGGCGGTTTTCTTTTCTCTTCTCAATTCGCGACCCAGGATGAGGTCGCCAAAACTACTAGATTTGGGCGCGCCTTGACGGCTAGCTCAGTCCTAGG
TACAGTGCTAGCTTAATTTCTAATTAGGATAAGGACACCAAGATGCGTAAAGGAGAAGAACTGTTCACTGGAGTTGTCCCAATTCTTGTTGAATTAGATG
GTGATGTTAATGGGCACAAATTTTCTGTGAGTGGAGAGGGTGAAGGTGATGCGACATACGGCAAACTGACCCTTAAATTTATTTGTACTACCGGCAAGCT
ACCTGTTCCTTGGCCGACACTTGTCACTACTTTCGGTTATGGTGTTCAATGTTTTGCGAGATACCCAGATCATATGAAACAGCATGACTTTTTCAAAAGT
GCCATGCCCGAAGGTTATGTACAGGAAAGAACTATATTTTTCAAAGATGACGGGAACTACAAAACACGTGCTGAAGTTAAATTTGAGGGTGATACTCTGG
TAAATAGAATCGAATTAAAAGGTATTGATTTTAAAGAAGATGGCAACATTCTTGGTCACAAGCTGGAATACAACTACAACTCACACAATGTTTACATCAT
GGCAGACAAACAAAAAAATGGCATCAAAGTTAACTTCAAAATTCGACACAACATTGAAGATGGAAGCGTTCAACTGGCAGACCATTATCAACAGAATACT
CCGATTGGCGATGGCCCTGTACTCTTACCAGACAACCATTACCTGTCCACACAATCTGCCCTTTCGAAAGATCCGAACGAAAAGAGAGATCACATGGTCC
TGCTTGAGTTTGTAACAGCTGCTGGGATTACACATGGTATGGATGAGCTATACAAATAATAACCAATTATTGAAGGCCTCCCTAACGGGGGGCCTTTTTT
TGTTTCTGGTCTCCCCACTGATAGTGCTAGTGTAGATCACTACTAGAGCCAGGCATCAAATAAAACGAAAGGCTCAGTCGAAAGACTGGGCCTTTCGTTT
TATCTGTTGTTTGTCGGTGAACGCTCTCTACTAGAGTCACACTGGCTCACCTTCGGGTGGGCCTTTCTGCGTTTATATACTAGAGGGAATCGTACGGTCG
ATCCATCTGACTATCGCCTACTTCATTGACAGCTAGCTCAGTCCTAGGTATAATGCTAGCCTAATTCTACATAGAGTGAGGTAATCTAATGGCTTCCTCC
GAAGACGTTATCAAAGAGTTCATGCGTTTCAAAGTTCGTATGGAAGGTTCCGTTAACGGTCACGAGTTCGAAATCGAAGGTGAAGGTGAAGGTCGTCCGT
ACGAAGGTACCCAGACCGCTAAACTGAAAGTTACCAAAGGTGGTCCGCTGCCGTTCGCTTGGGACATCCTGTCCCCGCAGTTCCAGTACGGTTCCAAAGC
TTACGTTAAACACCCGGCTGACATCCCGGACTACCTGAAACTGTCCTTCCCGGAAGGTTTCAAATGGGAACGTGTTATGAACTTCGAAGACGGTGGTGTT
GTTACCGTTACCCAGGACTCCTCCCTGCAAGACGGTGAGTTCATCTACAAAGTTAAACTGCGTGGTACCAACTTCCCGTCCGACGGTCCGGTTATGCAGA
AAAAAACCATGGGTTGGGAAGCTTCCACCGAACGTATGTACCCGGAAGACGGTGCTCTGAAAGGTGAAATCAAAATGCGTCTGAAACTGAAAGACGGTGG
TCACTACGACGCTGAAGTTAAAACCACCTACATGGCTAAAAAACCGGTTCAGCTGCCGGGTGCTTACAAAACCGACATCAAACTGGACATCACCTCCCAC
AACGAAGACTACACCATCGTTGAACAGTACGAACGTGCTGAAGGTCGTCACTCCACCGGTGCTTAATAACGCTGATAGTGCTAGTGTAGATCGCTACTAG
AGTACTAGCTCTAGATGCGGCCGCGAATTCCAGAAATCATCCTTAGCGAAAGCTAAGGATTTTTTTTATCTGAAATGCCTCTGTAACAGAGCATTAGCGC
AAGGTGATTTTTGTCTTCTTGCGCTAATTTTTTGGCGTCTCCACTGTGCGCGGAACCCCTATTTGTTTATTTTTCTAAATACATTCAAATATGTATCCGC
TCATGAGACAATAACCCTGATAAATGCTTCAATAATATTGAAAAAGGAAGAGTATGAGTATTCAACATTTCCGTGTCGCCCTTATTCCCTTTTTTGCGGC
ATTTTGCCTTCCTGTTTTTGCTCACCCAGAAACGCTGGTGAAAGTAAAAGATGCTGAAGATCAGTTGGGTGCACGAGTGGGTTACATCGAACTGGATCTC
AACAGCGGTAAGATCCTTGAGAGTTTTCGCCCCGAAGAACGTTTTCCAATGATGAGCACTTTTAAAGTTCTGCTATGTGGCGCGGTATTATCCCGTATTG
ACGCCGGGCAAGAGCAACTCGGTCGCCGCATACACTATTCTCAGAATGACTTGGTTGAGTACTCACCAGTCACAGAAAAGCATCTTACGGATGGCATGAC
AGTAAGAGAATTATGCAGTGCTGCCATAACCATGAGTGATAACACTGCGGCCAACTTACTTCTGACAACGATCGGAGGACCGAAGGAGCTAACCGCTTTT
TTGCACAACATGGGGGATCATGTAACTCGCCTTGATCGTTGGGAACCGGAGCTGAATGAAGCCATACCAAACGACGAGCGTGACACCACGATGCCTGTAG
CAATGGCAACAACGTTGCGCAAACTATTAACTGGCGAACTACTTACTCTAGCTTCCCGGCAACAATTAATAGACTGGATGGAGGCGGATAAAGTTGCAGG
ACCACTTCTGCGCTCGGCCCTTCCGGCTGGCTGGTTTATTGCTGATAAATCTGGAGCCGGTGAGCGTGGGTCTCGCGGTATCATTGCAGCACTGGGGCCA
GATGGTAAGCCCTCCCGTATCGTAGTTATCTACACGACGGGGAGTCAGGCAACTATGGATGAACGAAATAGACAGATCGCTGAGATAGGTGCCTCACTGA

TTAAGCATTGGTAACTCGAGCTGATCCTTCAAGAGCTCGCTTGGACTCCTGTTGATAGATCCAGTAATGACCTCAGAACTCCATCTGGATTTGTTCAGAA
CGCTCGGTTGCCGCCGGGCGTTTTTTATTGGTGAGAATTTACAACTTATATCGTATGGGGCTGACTTCAGGTGCTACATTTGAAGAGATAAATTGCACTG
AAATCTAGAAATATTTTATCTGATTAATAAGATGATCTTCTTGAGATCGTTTTGGTCTGCGCGTAATCTCTTGCTCTGAAAACGAAAAAACCGCCTTGCA
GGGCGGTTTTTCGAAGGTTCTCTGAGCTACCAACTCTTTGAACCGAGGTAACTGGCTTGGAGGAGCGCAGTCACCAAAACTTGTCCTTTCAGTTTAGCCT
TAACCGGCGCATGACTTCAAGACTAACTCCTCTAAATCAATTACCAGTGGCTGCTGCCAGTGGTGCTTTTGCATGTCTTTCCGGGTTGGACTCAAGACGA
TAGTTACCGGATAAGGCGCAGCGGTCGGACTGAACGGGGGGTTCGTGCATACAGTCCAGCTTGGAGCGAACTGCCTACCCGGAACTGAGTGTCAGGCGTG
GAATGAGACAAACGCGGCCATAACAGCGGAATGACACCGGTAAACCGAAAGGCAGGAACAGGAGAGCGCACGAGGGAGCCGCCAGGGGGAAACGCCTGGT
ATCTTTATAGTCCTGTCGGGTTTCGCCACCACTGATTTGAGCGTCAGATT

**pCM172**

TCGTGATGCTTGTCAGGGGGGCGGAGCCTATGGAAAAACGGCTTTGCCGCGGCCCTCTCACTTCCCTGTTAAGTATCTTCCTGGCATCTTCCAGGAAATC
TCCGCCCCGTTCGTAAGCCATTTCCGCTCGCCGCAGTCGAACGGGCGCGCCTTGACAGCTAGCTCAGTCCTAGGTATTGTGCTAGCTTACGTCAACCCAC
CCTAGGAGAGGATAAGATGAGCGAACTGATCAAAGAGAACATGCACATGAAGCTGTACATGGAGGGTACCGTGGATAATCACCACTTTAAGTGTACTTCT
GAGGGCGAGGGTAAGCCGTATGAAGGGACTCAAACGATGCGTATTAAAGTAGTGGAGGGTGGCCCACTGCCGTTTGCTTTCGATATTCTGGCGACGAGCT
TTCTGTATGGTAGCAAAACGTTTATAAACCACACTCAGGGCATTCCGGATTTCTTTAAACAAAGCTTTCCGGAAGGTTTTACCTGGGAGCGTGTGACTAC
GTATGAAGATGGTGGTGTGTTGACTGCTACTCAAGATACTTCACTGCAGGACGGCTGTCTGATCTATAACGTGAAGATTCGTGGCGTGAACTTTACGAGC
AATGGGCCGGTAATGCAAAAAAAAACCCTGGGTTGGGAAGCGTTCACGGAAACTCTGTATCCGGCTGACGGCGGCCTGGAGGGCCGTAACGATATGGCAC
TGAAGCTGGTTGGTGGCAGCCACCTGATCGCGAATATCAAAACGACTTATCGCTCTAAAAAACCGGCGAAAAATCTGAAGATGCCGGGTGTTTATTATGT
TGACTATCGTCTGGAACGCATTAAAGAAGCGAATAATGAAACTTACGTGGAGCAACACGAGGTTGCAGTGGCGCGCTATTGCGACTTGCCTTCAAAGCTG
GGTCACAAACTGAATTAATAAGGAAACACAGAAAAAAGCCCGCACCTGACAGTGCGGGCTTTTTTTTTCGACCAAAGGGACAATGTCCTGAAGAATATTT
AAGAAAAAAGCACCCCTCGTCGCCTAGAATTACCTACCGCGGTCGACCATACCTTCGAGACGAACAATAAGGCCGCAAATCGCGGCCTTCGTTATTGATA
ACAGCAGGTAAAAAGACGCTTATTATGGAGTTCGACATGGCAATAACGCGACGAATCTACGTCATGACGAGAATAGTATAAACAAAACTGCTGACGGCAG
AAGCGTCTCGGTACCAAATTCCAGAAAAGAGACGCTTTCGAGCGTCTTTTTTCGTTTTGGTCCACGCTGATGAATATTCTACGGAATTGCCATAGGCGTT
GAACGCTACACGGACGATACGAATTTATGTATAGAGCGTTCAGCCAAAAAACTTAAGACCGCCGGTCTTGTCCACTACCTTGCAGTAATGCGGTGGACAG
GATCGGCGGTTTTCTTTTCTCTTCTCAATTCGCGACCCAGGATGAGGTCGCCAAAACTACTAGATTTGGCTTAACGATCGTTGGCTGCGACGTACGGTGG
AATCTGATTCGTTACCAATTGACATGATACGAAACGTACCGTATCGTTAAGGTAGCTGTCACCGGATGTGCTTTCCGGTCTGATGAGTCCGTGAGGACGA
AACAGCCTCTACAAATAATTTTGTTTAATACTAGAGAAAGAGGGGAAATACTAGATGGTGAGCAAGGGCGAGGAGCTGTTCACCGGGGTGGTGCCCATCC
TGGTCGAGCTGGACGGCGACGTAAACGGCCACAAGTTCAGCGTGTCCGGCGAGGGCGAGGGCGATGCCACCTACGGCAAGCTGACCCTGAAGTTCATCTG
CACCACAGGCAAGCTGCCCGTGCCCTGGCCCACCCTCGTGACCACCTTCGGCTACGGCCTGCAATGCTTCGCCCGCTACCCCGACCACATGAAGCTGCAC
GACTTCTTCAAGTCCGCCATGCCCGAAGGCTACGTCCAGGAGCGCACCATCTTCTTCAAGGACGACGGCAACTACAAGACCCGCGCCGAGGTGAAGTTCG
AGGGCGACACCCTGGTGAACCGCATCGAGCTGAAGGGCATCGACTTCAAGGAGGACGGCAACATCCTGGGGCACAAGCTGGAGTACAACTACAACAGCCA
CAACGTCTATATCATGGCCGACAAGCAGAAGAACGGCATCAAGGTGAACTTCAAGATCCGCCACAACATCGAGGACGGCAGCGTGCAGCTCGCCGACCAC
TACCAGCAGAACACCCCAATCGGCGACGGCCCCGTGCTGCTGCCCGACAACCACTACCTTAGCTACCAGTCCGCCCTGAGCAAAGACCCCAACGAGAAGC
GCGATCACATGGTCCTGCTGGAGTTCGTGACCGCCGCCGGGATCACTCTCGGCATGGACGAGCTGTACAAGTAACTCGGTACCAAATTCCAGAAAAGAGG
CCTCCCGAAAGGGGGGCCTTTTTTCGTTTTGGTCCAATGGCGGCGCGCCATCGAATGGTGCAAAACCTTTCGCGGTATGGCATGATAGCGCCCGGAAGAG
AGTCAATTCATGGGGGTGAATATGGCACGTACCCCGAGCCGTAGCAGCATTGGTAGCCTGCGTAGTCCGCATACCCATAAAGCAATTCTGACCAGCACCA
TTGAAATCCTGAAAGAATGTGGTTATAGCGGTCTGAGCATTGAAAGCGTGGCACGTCGCGCCGGTGCAGGCAAACCGACCATTTATCGTTGGTGGACCAA
CAAAGCAGCACTGATTGCCGAAGTGTATGAAAATGAAATCGAACAGGTACGTAAATTTCCGGATTTGGGTAGCTTTAAAGCCGATCTGGATTTTCTGCTG
CATAATCTGTGGAAAGTTTGGCGTGAAACCATTTGTGGTGAAGCATTTCGTTGTGTTATTGCAGAAGCACAGTTGGACCCTGTAACCCTGACCCAACTGA
AAGATCAGTTTATGGAACGTCGTCGTGAGATACCGAAAAAACTGGTTGAAGATGCCATTAGCAATGGTGAACTGCCGAAAGATATCAATCGTGAACTGCT
GCTGGATATGATTTTTGGTTTTTGTTGGTATCGCCTGCTGACCGAACAGTTGACCGTTGAACAGGATATTGAAGAATTTACCTTCCTGCTGATTAATGGT
GTTTGTCCGGGTACACAGTGTTGATAAGGATCCCTCGGTACCAAAAAAAAAAAAAAAAGACGCTGAAAAGCGTCTTTTTTCGTTTTGGTCCGGCGTCTCCA
CTGTGCGCGGAACCCCTATTTGTTTATTTTTCTAAATACATTCAAATATGTATCCGCTCATGAGACAATAACCCTGATAAATGCTTCAATAATATTGAAA
AAGGAAGAGTATGAGTATTCAACATTTCCGTGTCGCCCTTATTCCCTTTTTTGCGGCATTTTGCCTTCCTGTTTTTGCTCACCCAGAAACGCTGGTGAAA
GTAAAAGATGCTGAAGATCAGTTGGGTGCACGAGTGGGTTACATCGAACTGGATCTCAACAGCGGTAAGATCCTTGAGAGTTTTCGCCCCGAAGAACGTT
TTCCAATGATGAGCACTTTTAAAGTTCTGCTATGTGGCGCGGTATTATCCCGTATTGACGCCGGGCAAGAGCAACTCGGTCGCCGCATACACTATTCTCA
GAATGACTTGGTTGAGTACTCACCAGTCACAGAAAAGCATCTTACGGATGGCATGACAGTAAGAGAATTATGCAGTGCTGCCATAACCATGAGTGATAAC
ACTGCGGCCAACTTACTTCTGACAACGATCGGAGGACCGAAGGAGCTAACCGCTTTTTTGCACAACATGGGGGATCATGTAACTCGCCTTGATCGTTGGG
AACCGGAGCTGAATGAAGCCATACCAAACGACGAGCGTGACACCACGATGCCTGTAGCAATGGCAACAACGTTGCGCAAACTATTAACTGGCGAACTACT
TACTCTAGCTTCCCGGCAACAATTAATAGACTGGATGGAGGCGGATAAAGTTGCAGGACCACTTCTGCGCTCGGCCCTTCCGGCTGGCTGGTTTATTGCT
GATAAATCTGGAGCCGGTGAGCGTGGGTCTCGCGGTATCATTGCAGCACTGGGGCCAGATGGTAAGCCCTCCCGTATCGTAGTTATCTACACGACGGGGA
GTCAGGCAACTATGGATGAACGAAATAGACAGATCGCTGAGATAGGTGCCTCACTGATTAAGCATTGGTAACTCGAGCTGATCCTTCAAGAGCTCGCTTG
GACTCCTGTTGATAGATCCAGTAATGACCTCAGAACTCCATCTGGATTTGTTCAGAACGCTCGGTTGCCGCCGGGCGTTTTTTATTGGTGAGAATTTACA
ACTTATATCGTATGGGGCTGACTTCAGGTGCTACATTTGAAGAGATAAATTGCACTGAAATCTAGAAATATTTTATCTGATTAATAAGATGATCTTCTTG
AGATCGTTTTGGTCTGCGCGTAATCTCTTGCTCTGAAAACGAAAAAACCGCCTTGCAGGGCGGTTTTTCGAAGGTTCTCTGAGCTACCAACTCTTTGAAC
CGAGGTAACTGGCTTGGAGGAGCGCAGTCACCAAAACTTGTCCTTTCAGTTTAGCCTTAACCGGCGCATGACTTCAAGACTAACTCCTCTAAATCAATTA
CCAGTGGCTGCTGCCAGTGGTGCTTTTGCATGTCTTTCCGGGTTGGACTCAAGACGATAGTTACCGGATAAGGCGCAGCGGTCGGACTGAACGGGGGGTT
CGTGCATACAGTCCAGCTTGGAGCGAACTGCCTACCCGGAACTGAGTGTCAGGCGTGGAATGAGACAAACGCGGCCATAACAGCGGAATGACACCGGTAA
ACCGAAAGGCAGGAACAGGAGAGCGCACGAGGGAGCCGCCAGGGGGAAACGCCTGGTATCTTTATAGTCCTGTCGGGTTTCGCCACCACTGATTTGAGCG
TCAGATT

**pCM173**

TCGTGATGCTTGTCAGGGGGGCGGAGCCTATGGAAAAACGGCTTTGCCGCGGCCCTCTCACTTCCCTGTTAAGTATCTTCCTGGCATCTTCCAGGAAATC
TCCGCCCCGTTCGTAAGCCATTTCCGCTCGCCGCAGTCGAACGGGCGCGCCTTGACAGCTAGCTCAGTCCTAGGTATTGTGCTAGCTTACGCCGATTCTG
CCGGGGGGAATTATAATGAGCGAACTGATCAAAGAGAACATGCACATGAAGCTGTACATGGAGGGTACCGTGGATAATCACCACTTTAAGTGTACTTCTG
AGGGCGAGGGTAAGCCGTATGAAGGGACTCAAACGATGCGTATTAAAGTAGTGGAGGGTGGCCCACTGCCGTTTGCTTTCGATATTCTGGCGACGAGCTT
TCTGTATGGTAGCAAAACGTTTATAAACCACACTCAGGGCATTCCGGATTTCTTTAAACAAAGCTTTCCGGAAGGTTTTACCTGGGAGCGTGTGACTACG
TATGAAGATGGTGGTGTGTTGACTGCTACTCAAGATACTTCACTGCAGGACGGCTGTCTGATCTATAACGTGAAGATTCGTGGCGTGAACTTTACGAGCA
ATGGGCCGGTAATGCAAAAAAAAACCCTGGGTTGGGAAGCGTTCACGGAAACTCTGTATCCGGCTGACGGCGGCCTGGAGGGCCGTAACGATATGGCACT
GAAGCTGGTTGGTGGCAGCCACCTGATCGCGAATATCAAAACGACTTATCGCTCTAAAAAACCGGCGAAAAATCTGAAGATGCCGGGTGTTTATTATGTT
GACTATCGTCTGGAACGCATTAAAGAAGCGAATAATGAAACTTACGTGGAGCAACACGAGGTTGCAGTGGCGCGCTATTGCGACTTGCCTTCAAAGCTGG
GTCACAAACTGAATTAATAAGGAAACACAGAAAAAAGCCCGCACCTGACAGTGCGGGCTTTTTTTTTCGACCAAAGGGACAATGTCCTGAAGAATATTTA
AGAAAAAAGCACCCCTCGTCGCCTAGAATTACCTACCGCGGTCGACCATACCTTCGAGACGAACAATAAGGCCGCAAATCGCGGCCTTCGTTATTGATAA
CAGCAGGTAAAAAGACGCTTATTATGGAGTTCGACATGGCAATAACGCGACGAATCTACGTCATGACGAGAATAGTATAAACAAAACTGCTGACGGCAGA
AGCGTCTCGGTACCAAATTCCAGAAAAGAGACGCTTTCGAGCGTCTTTTTTCGTTTTGGTCCACGCTGATGAATATTCTACGGAATTGCCATAGGCGTTG
AACGCTACACGGACGATACGAATTTATGTATAGAGCGTTCAGCCAAAAAACTTAAGACCGCCGGTCTTGTCCACTACCTTGCAGTAATGCGGTGGACAGG
ATCGGCGGTTTTCTTTTCTCTTCTCAATTCGCGACCCAGGATGAGGTCGCCAAAACTACTAGATTTGGCTTAACGATCGTTGGCTGCGACGTACGGTGGA
ATCTGATTCGTTACCAATTGACATGATACGAAACGTACCGTATCGTTAAGGTAGCTGTCACCGGATGTGCTTTCCGGTCTGATGAGTCCGTGAGGACGAA
ACAGCCTCTACAAATAATTTTGTTTAATACTAGAGAAAGAGGGGAAATACTAGATGGTGAGCAAGGGCGAGGAGCTGTTCACCGGGGTGGTGCCCATCCT
GGTCGAGCTGGACGGCGACGTAAACGGCCACAAGTTCAGCGTGTCCGGCGAGGGCGAGGGCGATGCCACCTACGGCAAGCTGACCCTGAAGTTCATCTGC
ACCACAGGCAAGCTGCCCGTGCCCTGGCCCACCCTCGTGACCACCTTCGGCTACGGCCTGCAATGCTTCGCCCGCTACCCCGACCACATGAAGCTGCACG

ACTTCTTCAAGTCCGCCATGCCCGAAGGCTACGTCCAGGAGCGCACCATCTTCTTCAAGGACGACGGCAACTACAAGACCCGCGCCGAGGTGAAGTTCGA
GGGCGACACCCTGGTGAACCGCATCGAGCTGAAGGGCATCGACTTCAAGGAGGACGGCAACATCCTGGGGCACAAGCTGGAGTACAACTACAACAGCCAC
AACGTCTATATCATGGCCGACAAGCAGAAGAACGGCATCAAGGTGAACTTCAAGATCCGCCACAACATCGAGGACGGCAGCGTGCAGCTCGCCGACCACT
ACCAGCAGAACACCCCCAATCGGCGACGGCCCCGTGCTGCTGCCCGACAACCACTACCTTAGCTACCAGTCCGCCCTGAGCAAAGACCCCAACGAGAAGCG
CGATCACATGGTCCTGCTGGAGTTCGTGACCGCCGCCGGGATCACTCTCGGCATGGACGAGCTGTACAAGTAACTCGGTACCAAATTCCAGAAAAGAGGC
CTCCCGAAAGGGGGGCCTTTTTTCGTTTTGGTCCAATGGCGGCGCGCCATCGAATGGTGCAAAACCTTTCGCGGTATGGCATGATAGCGCCCGGAAGAGA
GTCAATTCATGGGGGTGAATATGGCACGTACCCCGAGCCGTAGCAGCATTGGTAGCCTGCGTAGTCCGCATACCCATAAAGCAATTCTGACCAGCACCAT
TGAAATCCTGAAAGAATGTGGTTATAGCGGTCTGAGCATTGAAAGCGTGGCACGTCGCGCCGGTGCAGGCAAACCGACCATTTATCGTTGGTGGACCAAC
AAAGCAGCACTGATTGCCGAAGTGTATGAAAATGAAATCGAACAGGTACGTAAATTTCCGGATTTGGGTAGCTTTAAAGCCGATCTGGATTTTCTGCTGC
ATAATCTGTGGAAAGTTTGGCGTGAAACCATTTGTGGTGAAGCATTTCGTTGTGTTATTGCAGAAGCACAGTTGGACCCTGTAACCCTGACCCAACTGAA
AGATCAGTTTATGGAACGTCGTCGTGAGATACCGAAAAAACTGGTTGAAGATGCCATTAGCAATGGTGAACTGCCGAAAGATATCAATCGTGAACTGCTG
CTGGATATGATTTTTGGTTTTTGTTGGTATCGCCTGCTGACCGAACAGTTGACCGTTGAACAGGATATTGAAGAATTTACCTTCCTGCTGATTAATGGTG
TTTGTCCGGGTACACAGTGTTGATAAGGATCCCTCGGTACCAAAAAAAAAAAAAAAAGACGCTGAAAAGCGTCTTTTTTCGTTTTGGTCCGGCGTCTCCAC
TGTGCGCGGAACCCCTATTTGTTTATTTTTCTAAATACATTCAAATATGTATCCGCTCATGAGACAATAACCCTGATAAATGCTTCAATAATATTGAAAA
AGGAAGAGTATGAGTATTCAACATTTCCGTGTCGCCCTTATTCCCTTTTTTGCGGCATTTTGCCTTCCTGTTTTTGCTCACCCAGAAACGCTGGTGAAAG
TAAAAGATGCTGAAGATCAGTTGGGTGCACGAGTGGGTTACATCGAACTGGATCTCAACAGCGGTAAGATCCTTGAGAGTTTTCGCCCCGAAGAACGTTT
TCCAATGATGAGCACTTTTAAAGTTCTGCTATGTGGCGCGGTATTATCCCGTATTGACGCCGGGCAAGAGCAACTCGGTCGCCGCATACACTATTCTCAG
AATGACTTGGTTGAGTACTCACCAGTCACAGAAAAGCATCTTACGGATGGCATGACAGTAAGAGAATTATGCAGTGCTGCCATAACCATGAGTGATAACA
CTGCGGCCAACTTACTTCTGACAACGATCGGAGGACCGAAGGAGCTAACCGCTTTTTTGCACAACATGGGGGATCATGTAACTCGCCTTGATCGTTGGGA
ACCGGAGCTGAATGAAGCCATACCAAACGACGAGCGTGACACCACGATGCCTGTAGCAATGGCAACAACGTTGCGCAAACTATTAACTGGCGAACTACTT
ACTCTAGCTTCCCGGCAACAATTAATAGACTGGATGGAGGCGGATAAAGTTGCAGGACCACTTCTGCGCTCGGCCCTTCCGGCTGGCTGGTTTATTGCTG
ATAAATCTGGAGCCGGTGAGCGTGGGTCTCGCGGTATCATTGCAGCACTGGGGCCAGATGGTAAGCCCTCCCGTATCGTAGTTATCTACACGACGGGGAG
TCAGGCAACTATGGATGAACGAAATAGACAGATCGCTGAGATAGGTGCCTCACTGATTAAGCATTGGTAACTCGAGCTGATCCTTCAAGAGCTCGCTTGG
ACTCCTGTTGATAGATCCAGTAATGACCTCAGAACTCCATCTGGATTTGTTCAGAACGCTCGGTTGCCGCCGGGCGTTTTTTATTGGTGAGAATTTACAA
CTTATATCGTATGGGGCTGACTTCAGGTGCTACATTTGAAGAGATAAATTGCACTGAAATCTAGAAATATTTTATCTGATTAATAAGATGATCTTCTTGA
GATCGTTTTGGTCTGCGCGTAATCTCTTGCTCTGAAAACGAAAAAACCGCCTTGCAGGGCGGTTTTTCGAAGGTTCTCTGAGCTACCAACTCTTTGAACC
GAGGTAACTGGCTTGGAGGAGCGCAGTCACCAAAACTTGTCCTTTCAGTTTAGCCTTAACCGGCGCATGACTTCAAGACTAACTCCTCTAAATCAATTAC
CAGTGGCTGCTGCCAGTGGTGCTTTTGCATGTCTTTCCGGGTTGGACTCAAGACGATAGTTACCGGATAAGGCGCAGCGGTCGGACTGAACGGGGGGTTC
GTGCATACAGTCCAGCTTGGAGCGAACTGCCTACCCGGAACTGAGTGTCAGGCGTGGAATGAGACAAACGCGGCCATAACAGCGGAATGACACCGGTAAA
CCGAAAGGCAGGAACAGGAGAGCGCACGAGGGAGCCGCCAGGGGGAAACGCCTGGTATCTTTATAGTCCTGTCGGGTTTCGCCACCACTGATTTGAGCGT
CAGATT

**pCM182**

AACGGCTTTGCCGCGGCCCTCTCACTTCCCTGTTAAGTATCTTCCTGGCATCTTCCAGGAAATCTCCGCCCCGTTCGTAAGCCATTTCCGCTCGCCGCAG
TCGAACGTTCGCGACCCAGGATGAGGTCGCCAAAACTACTAGATTTGGGCGCGCCTTGACGGCTAGCTCAGTCCTAGGTACAGTGCTAGCTTAATTTCTA
ATTAGGATAAGGACACCAAGATGCGTAAAGGAGAAGAACTGTTCACTGGAGTTGTCCCAATTCTTGTTGAATTAGATGGTGATGTTAATGGGCACAAATT
TTCTGTGAGTGGAGAGGGTGAAGGTGATGCGACATACGGCAAACTGACCCTTAAATTTATTTGTACTACCGGCAAGCTACCTGTTCCTTGGCCGACACTT
GTCACTACTTTCGGTTATGGTGTTCAATGTTTTGCGAGATACCCAGATCATATGAAACAGCATGACTTTTTCAAAAGTGCCATGCCCGAAGGTTATGTAC
AGGAAAGAACTATATTTTTCAAAGATGACGGGAACTACAAAACACGTGCTGAAGTTAAATTTGAGGGTGATACTCTGGTAAATAGAATCGAATTAAAAGG
TATTGATTTTAAAGAAGATGGCAACATTCTTGGTCACAAGCTGGAATACAACTACAACTCACACAATGTTTACATCATGGCAGACAAACAAAAAAATGGC
ATCAAAGTTAACTTCAAAATTCGACACAACATTGAAGATGGAAGCGTTCAACTGGCAGACCATTATCAACAGAATACTCCGATTGGCGATGGCCCTGTAC
TCTTACCAGACAACCATTACCTGTCCACACAATCTGCCCTTTCGAAAGATCCGAACGAAAAGAGAGATCACATGGTCCTGCTTGAGTTTGTAACAGCTGC
TGGGATTACACATGGTATGGATGAGCTATACAAATAATAACCAATTATTGAAGGCCTCCCTAACGGGGGGCCTTTTTTTGTTTCTGGTCTCCCCACTGAT
AGTGCTAGTGTAGATCACTACTAGAGCCAGGCATCAAATAAAACGAAAGGCTCAGTCGAAAGACTGGGCCTTTCGTTTTATCTGTTGTTTGTCGGTGAAC
GCTCTCTACTAGAGTCACACTGGCTCACCTTCGGGTGGGCCTTTCTGCGTTTATATACTAGAGGGAATCGTACGGTCGATCCATCTGACTATCGCCTACT
TCAGCTTAACGATCGTTGGCTGCGACGTACGGTGGAATCTGATTCGTTACCAATTGACATGATACGAAACGTACCGTATCGTTAAGGTAGCTGTCACCGG
ATGTGCTTTCCGGTCTGATGAGTCCGTGAGGACGAAACAGCCTCTACAAATAATTTTGTTTAATACTAGAGAAAGAGGGGAAATACTAGATGGTGAGCAA
GGGCGAGGAGCTGTTCACCGGGGTGGTGCCCATCCTGGTCGAGCTGGACGGCGACGTAAACGGCCACAAGTTCAGCGTGTCCGGCGAGGGCGAGGGCGAT
GCCACCTACGGCAAGCTGACCCTGAAGTTCATCTGCACCACAGGCAAGCTGCCCGTGCCCTGGCCCACCCTCGTGACCACCTTCGGCTACGGCCTGCAAT
GCTTCGCCCGCTACCCCGACCACATGAAGCTGCACGACTTCTTCAAGTCCGCCATGCCCGAAGGCTACGTCCAGGAGCGCACCATCTTCTTCAAGGACGA
CGGCAACTACAAGACCCGCGCCGAGGTGAAGTTCGAGGGCGACACCCTGGTGAACCGCATCGAGCTGAAGGGCATCGACTTCAAGGAGGACGGCAACATC
CTGGGGCACAAGCTGGAGTACAACTACAACAGCCACAACGTCTATATCATGGCCGACAAGCAGAAGAACGGCATCAAGGTGAACTTCAAGATCCGCCACA
ACATCGAGGACGGCAGCGTGCAGCTCGCCGACCACTACCAGCAGAACACCCCCAATCGGCGACGGCCCCGTGCTGCTGCCCGACAACCACTACCTTAGCTA
CCAGTCCGCCCTGAGCAAAGACCCCAACGAGAAGCGCGATCACATGGTCCTGCTGGAGTTCGTGACCGCCGCCGGGATCACTCTCGGCATGGACGAGCTG
TACAAGTAACTCGGTACCAAATTCCAGAAAAGAGGCCTCCCGAAAGGGGGGCCTTTTTTCGTTTTGGTCCAATGGCGGCGCGCCATCGAATGGTGCAAAA
CCTTTCGCGGTATGGCATGATAGCGCCCGGAAGAGAGTCAATTCATGGGGGTGAATATGGCACGTACCCCGAGCCGTAGCAGCATTGGTAGCCTGCGTAG
TCCGCATACCCATAAAGCAATTCTGACCAGCACCATTGAAATCCTGAAAGAATGTGGTTATAGCGGTCTGAGCATTGAAAGCGTGGCACGTCGCGCCGGT
GCAGGCAAACCGACCATTTATCGTTGGTGGACCAACAAAGCAGCACTGATTGCCGAAGTGTATGAAAATGAAATCGAACAGGTACGTAAATTTCCGGATT
TGGGTAGCTTTAAAGCCGATCTGGATTTTCTGCTGCATAATCTGTGGAAAGTTTGGCGTGAAACCATTTGTGGTGAAGCATTTCGTTGTGTTATTGCAGA
AGCACAGTTGGACCCTGTAACCCTGACCCAACTGAAAGATCAGTTTATGGAACGTCGTCGTGAGATACCGAAAAAACTGGTTGAAGATGCCATTAGCAAT
GGTGAACTGCCGAAAGATATCAATCGTGAACTGCTGCTGGATATGATTTTTGGTTTTTGTTGGTATCGCCTGCTGACCGAACAGTTGACCGTTGAACAGG
ATATTGAAGAATTTACCTTCCTGCTGATTAATGGTGTTTGTCCGGGTACACAGTGTTGATAAGGATCCCTCGGTACCAAAAAAAAAAAAAAAAGACGCTGA
AAAGCGTCTTTTTTCGTTTTGGTCCGGCGTCTCCACTGTGCGCGGAACCCCTATTTGTTTATTTTTCTAAATACATTCAAATATGTATCCGCTCATGAGA
CAATAACCCTGATAAATGCTTCAATAATATTGAAAAAGGAAGAGTATGAGTATTCAACATTTCCGTGTCGCCCTTATTCCCTTTTTTGCGGCATTTTGCC
TTCCTGTTTTTGCTCACCCAGAAACGCTGGTGAAAGTAAAAGATGCTGAAGATCAGTTGGGTGCACGAGTGGGTTACATCGAACTGGATCTCAACAGCGG
TAAGATCCTTGAGAGTTTTCGCCCCGAAGAACGTTTTCCAATGATGAGCACTTTTAAAGTTCTGCTATGTGGCGCGGTATTATCCCGTATTGACGCCGGG
CAAGAGCAACTCGGTCGCCGCATACACTATTCTCAGAATGACTTGGTTGAGTACTCACCAGTCACAGAAAAGCATCTTACGGATGGCATGACAGTAAGAG
AATTATGCAGTGCTGCCATAACCATGAGTGATAACACTGCGGCCAACTTACTTCTGACAACGATCGGAGGACCGAAGGAGCTAACCGCTTTTTTGCACAA
CATGGGGGATCATGTAACTCGCCTTGATCGTTGGGAACCGGAGCTGAATGAAGCCATACCAAACGACGAGCGTGACACCACGATGCCTGTAGCAATGGCA
ACAACGTTGCGCAAACTATTAACTGGCGAACTACTTACTCTAGCTTCCCGGCAACAATTAATAGACTGGATGGAGGCGGATAAAGTTGCAGGACCACTTC
TGCGCTCGGCCCTTCCGGCTGGCTGGTTTATTGCTGATAAATCTGGAGCCGGTGAGCGTGGGTCTCGCGGTATCATTGCAGCACTGGGGCCAGATGGTAA
GCCCTCCCGTATCGTAGTTATCTACACGACGGGGAGTCAGGCAACTATGGATGAACGAAATAGACAGATCGCTGAGATAGGTGCCTCACTGATTAAGCAT
TGGTAACTCGAGCTGATCCTTCAAGAGCTCGCTTGGACTCCTGTTGATAGATCCAGTAATGACCTCAGAACTCCATCTGGATTTGTTCAGAACGCTCGGT
TGCCGCCGGGCGTTTTTTATTGGTGAGAATTTACAACTTATATCGTATGGGGCTGACTTCAGGTGCTACATTTGAAGAGATAAATTGCACTGAAATCTAG
AAATATTTTATCTGATTAATAAGATGATCTTCTTGAGATCGTTTTGGTCTGCGCGTAATCTCTTGCTCTGAAAACGAAAAAACCGCCTTGCAGGGCGGTT
TTTCGAAGGTTCTCTGAGCTACCAACTCTTTGAACCGAGGTAACTGGCTTGGAGGAGCGCAGTCACCAAAACTTGTCCTTTCAGTTTAGCCTTAACCGGC
GCATGACTTCAAGACTAACTCCTCTAAATCAATTACCAGTGGCTGCTGCCAGTGGTGCTTTTGCATGTCTTTCCGGGTTGGACTCAAGACGATAGTTACC
GGATAAGGCGCAGCGGTCGGACTGAACGGGGGGTTCGTGCATACAGTCCAGCTTGGAGCGAACTGCCTACCCGGAACTGAGTGTCAGGCGTGGAATGAGA
CAAACGCGGCCATAACAGCGGAATGACACCGGTAAACCGAAAGGCAGGAACAGGAGAGCGCACGAGGGAGCCGCCAGGGGGAAACGCCTGGTATCTTTAT
AGTCCTGTCGGGTTTCGCCACCACTGATTTGAGCGTCAGATTTCGTGATGCTTGTCAGGGGGGCGGAGCCTATGGAAA

**pCM183**

AACGGCTTTGCCGCGGCCCTCTCACTTCCCTGTTAAGTATCTTCCTGGCATCTTCCAGGAAATCTCCGCCCCGTTCGTAAGCCATTTCCGCTCGCCGCAG
TCGAACGTTCGCGACCCAGGATGAGGTCGCCAAAACTACTAGATTTGGGCGCGCCTTGACGGCTAGCTCAGTCCTAGGTACAGTGCTAGCTTAATGCGAG
AGGCTGGTTAAGTGGGTGTTGATGCGTAAAGGAGAAGAACTGTTCACTGGAGTTGTCCCAATTCTTGTTGAATTAGATGGTGATGTTAATGGGCACAAAT
TTTCTGTGAGTGGAGAGGGTGAAGGTGATGCGACATACGGCAAACTGACCCTTAAATTTATTTGTACTACCGGCAAGCTACCTGTTCCTTGGCCGACACT
TGTCACTACTTTCGGTTATGGTGTTCAATGTTTTGCGAGATACCCAGATCATATGAAACAGCATGACTTTTTCAAAAGTGCCATGCCCGAAGGTTATGTA
CAGGAAAGAACTATATTTTTCAAAGATGACGGGAACTACAAAACACGTGCTGAAGTTAAATTTGAGGGTGATACTCTGGTAAATAGAATCGAATTAAAAG
GTATTGATTTTAAAGAAGATGGCAACATTCTTGGTCACAAGCTGGAATACAACTACAACTCACACAATGTTTACATCATGGCAGACAAACAAAAAAATGG
CATCAAAGTTAACTTCAAAATTCGACACAACATTGAAGATGGAAGCGTTCAACTGGCAGACCATTATCAACAGAATACTCCGATTGGCGATGGCCCTGTA
CTCTTACCAGACAACCATTACCTGTCCACACAATCTGCCCTTTCGAAAGATCCGAACGAAAAGAGAGATCACATGGTCCTGCTTGAGTTTGTAACAGCTG
CTGGGATTACACATGGTATGGATGAGCTATACAAATAATAACCAATTATTGAAGGCCTCCCTAACGGGGGGCCTTTTTTTGTTTCTGGTCTCCCCACTGA
TAGTGCTAGTGTAGATCACTACTAGAGCCAGGCATCAAATAAAACGAAAGGCTCAGTCGAAAGACTGGGCCTTTCGTTTTATCTGTTGTTTGTCGGTGAA
CGCTCTCTACTAGAGTCACACTGGCTCACCTTCGGGTGGGCCTTTCTGCGTTTATATACTAGAGGGAATCGTACGGTCGATCCATCTGACTATCGCCTAC
TTCAGCTTAACGATCGTTGGCTGCGACGTACGGTGGAATCTGATTCGTTACCAATTGACATGATACGAAACGTACCGTATCGTTAAGGTAGCTGTCACCG
GATGTGCTTTCCGGTCTGATGAGTCCGTGAGGACGAAACAGCCTCTACAAATAATTTTGTTTAATACTAGAGAAAGAGGGGAAATACTAGATGGTGAGCA
AGGGCGAGGAGCTGTTCACCGGGGTGGTGCCCATCCTGGTCGAGCTGGACGGCGACGTAAACGGCCACAAGTTCAGCGTGTCCGGCGAGGGCGAGGGCGA
TGCCACCTACGGCAAGCTGACCCTGAAGTTCATCTGCACCACAGGCAAGCTGCCCGTGCCCTGGCCCACCCTCGTGACCACCTTCGGCTACGGCCTGCAA
TGCTTCGCCCGCTACCCCGACCACATGAAGCTGCACGACTTCTTCAAGTCCGCCATGCCCGAAGGCTACGTCCAGGAGCGCACCATCTTCTTCAAGGACG
ACGGCAACTACAAGACCCGCGCCGAGGTGAAGTTCGAGGGCGACACCCTGGTGAACCGCATCGAGCTGAAGGGCATCGACTTCAAGGAGGACGGCAACAT
CCTGGGGCACAAGCTGGAGTACAACTACAACAGCCACAACGTCTATATCATGGCCGACAAGCAGAAGAACGGCATCAAGGTGAACTTCAAGATCCGCCAC
AACATCGAGGACGGCAGCGTGCAGCTCGCCGACCACTACCAGCAGAACACCCCAATCGGCGACGGCCCCGTGCTGCTGCCCGACAACCACTACCTTAGCT
ACCAGTCCGCCCTGAGCAAAGACCCCAACGAGAAGCGCGATCACATGGTCCTGCTGGAGTTCGTGACCGCCGCCGGGATCACTCTCGGCATGGACGAGCT
GTACAAGTAACTCGGTACCAAATTCCAGAAAAGAGGCCTCCCGAAAGGGGGGCCTTTTTTCGTTTTGGTCCAATGGCGGCGCGCCATCGAATGGTGCAAA
ACCTTTCGCGGTATGGCATGATAGCGCCCGGAAGAGAGTCAATTCATGGGGGTGAATATGGCACGTACCCCGAGCCGTAGCAGCATTGGTAGCCTGCGTA
GTCCGCATACCCATAAAGCAATTCTGACCAGCACCATTGAAATCCTGAAAGAATGTGGTTATAGCGGTCTGAGCATTGAAAGCGTGGCACGTCGCGCCGG
TGCAGGCAAACCGACCATTTATCGTTGGTGGACCAACAAAGCAGCACTGATTGCCGAAGTGTATGAAAATGAAATCGAACAGGTACGTAAATTTCCGGAT
TTGGGTAGCTTTAAAGCCGATCTGGATTTTCTGCTGCATAATCTGTGGAAAGTTTGGCGTGAAACCATTTGTGGTGAAGCATTTCGTTGTGTTATTGCAG
AAGCACAGTTGGACCCTGTAACCCTGACCCAACTGAAAGATCAGTTTATGGAACGTCGTCGTGAGATACCGAAAAAACTGGTTGAAGATGCCATTAGCAA
TGGTGAACTGCCGAAAGATATCAATCGTGAACTGCTGCTGGATATGATTTTTGGTTTTTGTTGGTATCGCCTGCTGACCGAACAGTTGACCGTTGAACAG
GATATTGAAGAATTTACCTTCCTGCTGATTAATGGTGTTTGTCCGGGTACACAGTGTTGATAAGGATCCCTCGGTACCAAAAAAAAAAAAAAAAGACGCTG
AAAAGCGTCTTTTTTCGTTTTGGTCCGGCGTCTCCACTGTGCGCGGAACCCCTATTTGTTTATTTTTCTAAATACATTCAAATATGTATCCGCTCATGAG
ACAATAACCCTGATAAATGCTTCAATAATATTGAAAAAGGAAGAGTATGAGTATTCAACATTTCCGTGTCGCCCTTATTCCCTTTTTTGCGGCATTTTGC
CTTCCTGTTTTTGCTCACCCAGAAACGCTGGTGAAAGTAAAAGATGCTGAAGATCAGTTGGGTGCACGAGTGGGTTACATCGAACTGGATCTCAACAGCG
GTAAGATCCTTGAGAGTTTTCGCCCCGAAGAACGTTTTCCAATGATGAGCACTTTTAAAGTTCTGCTATGTGGCGCGGTATTATCCCGTATTGACGCCGG
GCAAGAGCAACTCGGTCGCCGCATACACTATTCTCAGAATGACTTGGTTGAGTACTCACCAGTCACAGAAAAGCATCTTACGGATGGCATGACAGTAAGA
GAATTATGCAGTGCTGCCATAACCATGAGTGATAACACTGCGGCCAACTTACTTCTGACAACGATCGGAGGACCGAAGGAGCTAACCGCTTTTTTGCACA
ACATGGGGGATCATGTAACTCGCCTTGATCGTTGGGAACCGGAGCTGAATGAAGCCATACCAAACGACGAGCGTGACACCACGATGCCTGTAGCAATGGC
AACAACGTTGCGCAAACTATTAACTGGCGAACTACTTACTCTAGCTTCCCGGCAACAATTAATAGACTGGATGGAGGCGGATAAAGTTGCAGGACCACTT
CTGCGCTCGGCCCTTCCGGCTGGCTGGTTTATTGCTGATAAATCTGGAGCCGGTGAGCGTGGGTCTCGCGGTATCATTGCAGCACTGGGGCCAGATGGTA
AGCCCTCCCGTATCGTAGTTATCTACACGACGGGGAGTCAGGCAACTATGGATGAACGAAATAGACAGATCGCTGAGATAGGTGCCTCACTGATTAAGCA
TTGGTAACTCGAGCTGATCCTTCAAGAGCTCGCTTGGACTCCTGTTGATAGATCCAGTAATGACCTCAGAACTCCATCTGGATTTGTTCAGAACGCTCGG
TTGCCGCCGGGCGTTTTTTATTGGTGAGAATTTACAACTTATATCGTATGGGGCTGACTTCAGGTGCTACATTTGAAGAGATAAATTGCACTGAAATCTA
GAAATATTTTATCTGATTAATAAGATGATCTTCTTGAGATCGTTTTGGTCTGCGCGTAATCTCTTGCTCTGAAAACGAAAAAACCGCCTTGCAGGGCGGT
TTTTCGAAGGTTCTCTGAGCTACCAACTCTTTGAACCGAGGTAACTGGCTTGGAGGAGCGCAGTCACCAAAACTTGTCCTTTCAGTTTAGCCTTAACCGG
CGCATGACTTCAAGACTAACTCCTCTAAATCAATTACCAGTGGCTGCTGCCAGTGGTGCTTTTGCATGTCTTTCCGGGTTGGACTCAAGACGATAGTTAC
CGGATAAGGCGCAGCGGTCGGACTGAACGGGGGGTTCGTGCATACAGTCCAGCTTGGAGCGAACTGCCTACCCGGAACTGAGTGTCAGGCGTGGAATGAG
ACAAACGCGGCCATAACAGCGGAATGACACCGGTAAACCGAAAGGCAGGAACAGGAGAGCGCACGAGGGAGCCGCCAGGGGGAAACGCCTGGTATCTTTA
TAGTCCTGTCGGGTTTCGCCACCACTGATTTGAGCGTCAGATTTCGTGATGCTTGTCAGGGGGGCGGAGCCTATGGAAA

**pCM184**

AACGGCTTTGCCGCGGCCCTCTCACTTCCCTGTTAAGTATCTTCCTGGCATCTTCCAGGAAATCTCCGCCCCGTTCGTAAGCCATTTCCGCTCGCCGCAG
TCGAACGTTCGCGACCCAGGATGAGGTCGCCAAAACTACTAGATTTGGGCGCGCCTTGACGGCTAGCTCAGTCCTAGGTACAGTGCTAGCTTAATCTTGT
CAAACAGTGGCTACGTGAACAGATGCGTAAAGGAGAAGAACTGTTCACTGGAGTTGTCCCAATTCTTGTTGAATTAGATGGTGATGTTAATGGGCACAAA
TTTTCTGTGAGTGGAGAGGGTGAAGGTGATGCGACATACGGCAAACTGACCCTTAAATTTATTTGTACTACCGGCAAGCTACCTGTTCCTTGGCCGACAC
TTGTCACTACTTTCGGTTATGGTGTTCAATGTTTTGCGAGATACCCAGATCATATGAAACAGCATGACTTTTTCAAAAGTGCCATGCCCGAAGGTTATGT
ACAGGAAAGAACTATATTTTTCAAAGATGACGGGAACTACAAAACACGTGCTGAAGTTAAATTTGAGGGTGATACTCTGGTAAATAGAATCGAATTAAAA
GGTATTGATTTTAAAGAAGATGGCAACATTCTTGGTCACAAGCTGGAATACAACTACAACTCACACAATGTTTACATCATGGCAGACAAACAAAAAAATG
GCATCAAAGTTAACTTCAAAATTCGACACAACATTGAAGATGGAAGCGTTCAACTGGCAGACCATTATCAACAGAATACTCCGATTGGCGATGGCCCTGT
ACTCTTACCAGACAACCATTACCTGTCCACACAATCTGCCCTTTCGAAAGATCCGAACGAAAAGAGAGATCACATGGTCCTGCTTGAGTTTGTAACAGCT
GCTGGGATTACACATGGTATGGATGAGCTATACAAATAATAACCAATTATTGAAGGCCTCCCTAACGGGGGGCCTTTTTTTGTTTCTGGTCTCCCCACTG
ATAGTGCTAGTGTAGATCACTACTAGAGCCAGGCATCAAATAAAACGAAAGGCTCAGTCGAAAGACTGGGCCTTTCGTTTTATCTGTTGTTTGTCGGTGA
ACGCTCTCTACTAGAGTCACACTGGCTCACCTTCGGGTGGGCCTTTCTGCGTTTATATACTAGAGGGAATCGTACGGTCGATCCATCTGACTATCGCCTA
CTTCAGCTTAACGATCGTTGGCTGCGACGTACGGTGGAATCTGATTCGTTACCAATTGACATGATACGAAACGTACCGTATCGTTAAGGTAGCTGTCACC
GGATGTGCTTTCCGGTCTGATGAGTCCGTGAGGACGAAACAGCCTCTACAAATAATTTTGTTTAATACTAGAGAAAGAGGGGAAATACTAGATGGTGAGC
AAGGGCGAGGAGCTGTTCACCGGGGTGGTGCCCATCCTGGTCGAGCTGGACGGCGACGTAAACGGCCACAAGTTCAGCGTGTCCGGCGAGGGCGAGGGCG
ATGCCACCTACGGCAAGCTGACCCTGAAGTTCATCTGCACCACAGGCAAGCTGCCCGTGCCCTGGCCCACCCTCGTGACCACCTTCGGCTACGGCCTGCA
ATGCTTCGCCCGCTACCCCGACCACATGAAGCTGCACGACTTCTTCAAGTCCGCCATGCCCGAAGGCTACGTCCAGGAGCGCACCATCTTCTTCAAGGAC
GACGGCAACTACAAGACCCGCGCCGAGGTGAAGTTCGAGGGCGACACCCTGGTGAACCGCATCGAGCTGAAGGGCATCGACTTCAAGGAGGACGGCAACA
TCCTGGGGCACAAGCTGGAGTACAACTACAACAGCCACAACGTCTATATCATGGCCGACAAGCAGAAGAACGGCATCAAGGTGAACTTCAAGATCCGCCA
CAACATCGAGGACGGCAGCGTGCAGCTCGCCGACCACTACCAGCAGAACACCCCAATCGGCGACGGCCCCGTGCTGCTGCCCGACAACCACTACCTTAGC
TACCAGTCCGCCCTGAGCAAAGACCCCAACGAGAAGCGCGATCACATGGTCCTGCTGGAGTTCGTGACCGCCGCCGGGATCACTCTCGGCATGGACGAGC
TGTACAAGTAACTCGGTACCAAATTCCAGAAAAGAGGCCTCCCGAAAGGGGGGCCTTTTTTCGTTTTGGTCCAATGGCGGCGCGCCATCGAATGGTGCAA
AACCTTTCGCGGTATGGCATGATAGCGCCCGGAAGAGAGTCAATTCATGGGGGTGAATATGGCACGTACCCCGAGCCGTAGCAGCATTGGTAGCCTGCGT
AGTCCGCATACCCATAAAGCAATTCTGACCAGCACCATTGAAATCCTGAAAGAATGTGGTTATAGCGGTCTGAGCATTGAAAGCGTGGCACGTCGCGCCG
GTGCAGGCAAACCGACCATTTATCGTTGGTGGACCAACAAAGCAGCACTGATTGCCGAAGTGTATGAAAATGAAATCGAACAGGTACGTAAATTTCCGGA
TTTGGGTAGCTTTAAAGCCGATCTGGATTTTCTGCTGCATAATCTGTGGAAAGTTTGGCGTGAAACCATTTGTGGTGAAGCATTTCGTTGTGTTATTGCA
GAAGCACAGTTGGACCCTGTAACCCTGACCCAACTGAAAGATCAGTTTATGGAACGTCGTCGTGAGATACCGAAAAAACTGGTTGAAGATGCCATTAGCA
ATGGTGAACTGCCGAAAGATATCAATCGTGAACTGCTGCTGGATATGATTTTTGGTTTTTGTTGGTATCGCCTGCTGACCGAACAGTTGACCGTTGAACA
GGATATTGAAGAATTTACCTTCCTGCTGATTAATGGTGTTTGTCCGGGTACACAGTGTTGATAAGGATCCCTCGGTACCAAAAAAAAAAAAAAAAGACGCT
GAAAAGCGTCTTTTTTCGTTTTGGTCCGGCGTCTCCACTGTGCGCGGAACCCCTATTTGTTTATTTTTCTAAATACATTCAAATATGTATCCGCTCATGA

GACAATAACCCTGATAAATGCTTCAATAATATTGAAAAAGGAAGAGTATGAGTATTCAACATTTCCGTGTCGCCCTTATTCCCTTTTTTGCGGCATTTTG
CCTTCCTGTTTTTGCTCACCCAGAAACGCTGGTGAAAGTAAAAGATGCTGAAGATCAGTTGGGTGCACGAGTGGGTTACATCGAACTGGATCTCAACAGC
GGTAAGATCCTTGAGAGTTTTCGCCCCGAAGAACGTTTTCCAATGATGAGCACTTTTAAAGTTCTGCTATGTGGCGCGGTATTATCCCGTATTGACGCCG
GGCAAGAGCAACTCGGTCGCCGCATACACTATTCTCAGAATGACTTGGTTGAGTACTCACCAGTCACAGAAAAGCATCTTACGGATGGCATGACAGTAAG
AGAATTATGCAGTGCTGCCATAACCATGAGTGATAACACTGCGGCCAACTTACTTCTGACAACGATCGGAGGACCGAAGGAGCTAACCGCTTTTTTGCAC
AACATGGGGGATCATGTAACTCGCCTTGATCGTTGGGAACCGGAGCTGAATGAAGCCATACCAAACGACGAGCGTGACACCACGATGCCTGTAGCAATGG
CAACAACGTTGCGCAAACTATTAACTGGCGAACTACTTACTCTAGCTTCCCGGCAACAATTAATAGACTGGATGGAGGCGGATAAAGTTGCAGGACCACT
TCTGCGCTCGGCCCTTCCGGCTGGCTGGTTTATTGCTGATAAATCTGGAGCCGGTGAGCGTGGGTCTCGCGGTATCATTGCAGCACTGGGGCCAGATGGT
AAGCCCTCCCGTATCGTAGTTATCTACACGACGGGGAGTCAGGCAACTATGGATGAACGAAATAGACAGATCGCTGAGATAGGTGCCTCACTGATTAAGC
ATTGGTAACTCGAGCTGATCCTTCAAGAGCTCGCTTGGACTCCTGTTGATAGATCCAGTAATGACCTCAGAACTCCATCTGGATTTGTTCAGAACGCTCG
GTTGCCGCCGGGCGTTTTTTATTGGTGAGAATTTACAACTTATATCGTATGGGGCTGACTTCAGGTGCTACATTTGAAGAGATAAATTGCACTGAAATCT
AGAAATATTTTATCTGATTAATAAGATGATCTTCTTGAGATCGTTTTGGTCTGCGCGTAATCTCTTGCTCTGAAAACGAAAAAACCGCCTTGCAGGGCGG
TTTTTCGAAGGTTCTCTGAGCTACCAACTCTTTGAACCGAGGTAACTGGCTTGGAGGAGCGCAGTCACCAAAACTTGTCCTTTCAGTTTAGCCTTAACCG
GCGCATGACTTCAAGACTAACTCCTCTAAATCAATTACCAGTGGCTGCTGCCAGTGGTGCTTTTGCATGTCTTTCCGGGTTGGACTCAAGACGATAGTTA
CCGGATAAGGCGCAGCGGTCGGACTGAACGGGGGGTTCGTGCATACAGTCCAGCTTGGAGCGAACTGCCTACCCGGAACTGAGTGTCAGGCGTGGAATGA
GACAAACGCGGCCATAACAGCGGAATGACACCGGTAAACCGAAAGGCAGGAACAGGAGAGCGCACGAGGGAGCCGCCAGGGGGAAACGCCTGGTATCTTT
ATAGTCCTGTCGGGTTTCGCCACCACTGATTTGAGCGTCAGATTTCGTGATGCTTGTCAGGGGGGCGGAGCCTATGGAAA

**pCM192**

TCGTGATGCTTGTCAGGGGGGCGGAGCCTATGGAAAAACGGCTTTGCCGCGGCCCTCTCACTTCCCTGTTAAGTATCTTCCTGGCATCTTCCAGGAAATC
TCCGCCCCGTTCGTAAGCCATTTCCGCTCGCCGCAGTCGAACGTACTAGAGGGAATCGTACGGTCGATCCATCTGACTATCGCCTACTTCATTGACAGCT
AGCTCAGTCCTAGGTATAATGCTAGCCTAATTCTACATAGAGTGAGGTAATCTAATGGCTTCCTCCGAAGACGTTATCAAAGAGTTCATGCGTTTCAAAG
TTCGTATGGAAGGTTCCGTTAACGGTCACGAGTTCGAAATCGAAGGTGAAGGTGAAGGTCGTCCGTACGAAGGTACCCAGACCGCTAAACTGAAAGTTAC
CAAAGGTGGTCCGCTGCCGTTCGCTTGGGACATCCTGTCCCCGCAGTTCCAGTACGGTTCCAAAGCTTACGTTAAACACCCGGCTGACATCCCGGACTAC
CTGAAACTGTCCTTCCCGGAAGGTTTCAAATGGGAACGTGTTATGAACTTCGAAGACGGTGGTGTTGTTACCGTTACCCAGGACTCCTCCCTGCAAGACG
GTGAGTTCATCTACAAAGTTAAACTGCGTGGTACCAACTTCCCGTCCGACGGTCCGGTTATGCAGAAAAAAACCATGGGTTGGGAAGCTTCCACCGAACG
TATGTACCCGGAAGACGGTGCTCTGAAAGGTGAAATCAAAATGCGTCTGAAACTGAAAGACGGTGGTCACTACGACGCTGAAGTTAAAACCACCTACATG
GCTAAAAAACCGGTTCAGCTGCCGGGTGCTTACAAAACCGACATCAAACTGGACATCACCTCCCACAACGAAGACTACACCATCGTTGAACAGTACGAAC
GTGCTGAAGGTCGTCACTCCACCGGTGCTTAATAACGCTGATAGTGCTAGTGTAGATCGCTACTAGAGTACTAGCTCTAGATGCGGCCGCGAATTCCAGA
AATCATCCTTAGCGAAAGCTAAGGATTTTTTTTATCTGAAATGCCTCTGTAACAGAGCATTAGCGCAAGGTGATTTTTGTCTTCTTGCGCTAATTTTTTG
GTCCAACTTAGGCACCAATTATTGAAGGCCTCCCTAACGGGGGGCCTTTTTTTGTTTCTGGTCTCCCGCTTAACGATCGTTGGCTGCGACGTACGGTGGA
ATCTGATTCGTTACCAATTGACATGATACGAAACGTACCGTATCGTTAAGGTAGCTGTCACCGGATGTGCTTTCCGGTCTGATGAGTCCGTGAGGACGAA
ACAGCCTCTACAAATAATTTTGTTTAATACTAGAGAAAGAGGGGAAATACTAGATGGTGAGCAAGGGCGAGGAGCTGTTCACCGGGGTGGTGCCCATCCT
GGTCGAGCTGGACGGCGACGTAAACGGCCACAAGTTCAGCGTGTCCGGCGAGGGCGAGGGCGATGCCACCTACGGCAAGCTGACCCTGAAGTTCATCTGC
ACCACAGGCAAGCTGCCCGTGCCCTGGCCCACCCTCGTGACCACCTTCGGCTACGGCCTGCAATGCTTCGCCCGCTACCCCGACCACATGAAGCTGCACG
ACTTCTTCAAGTCCGCCATGCCCGAAGGCTACGTCCAGGAGCGCACCATCTTCTTCAAGGACGACGGCAACTACAAGACCCGCGCCGAGGTGAAGTTCGA
GGGCGACACCCTGGTGAACCGCATCGAGCTGAAGGGCATCGACTTCAAGGAGGACGGCAACATCCTGGGGCACAAGCTGGAGTACAACTACAACAGCCAC
AACGTCTATATCATGGCCGACAAGCAGAAGAACGGCATCAAGGTGAACTTCAAGATCCGCCACAACATCGAGGACGGCAGCGTGCAGCTCGCCGACCACT
ACCAGCAGAACACCCCCAATCGGCGACGGCCCCGTGCTGCTGCCCGACAACCACTACCTTAGCTACCAGTCCGCCCTGAGCAAAGACCCCAACGAGAAGCG
CGATCACATGGTCCTGCTGGAGTTCGTGACCGCCGCCGGGATCACTCTCGGCATGGACGAGCTGTACAAGTAACTCGGTACCAAATTCCAGAAAAGAGGC
CTCCCGAAAGGGGGGCCTTTTTTCGTTTTGGTCCAATGGCGGCGCGCCATCGAATGGTGCAAAACCTTTCGCGGTATGGCATGATAGCGCCCGGAAGAGA
GTCAATTCATGGGGGTGAATATGGCACGTACCCCGAGCCGTAGCAGCATTGGTAGCCTGCGTAGTCCGCATACCCATAAAGCAATTCTGACCAGCACCAT
TGAAATCCTGAAAGAATGTGGTTATAGCGGTCTGAGCATTGAAAGCGTGGCACGTCGCGCCGGTGCAGGCAAACCGACCATTTATCGTTGGTGGACCAAC
AAAGCAGCACTGATTGCCGAAGTGTATGAAAATGAAATCGAACAGGTACGTAAATTTCCGGATTTGGGTAGCTTTAAAGCCGATCTGGATTTTCTGCTGC
ATAATCTGTGGAAAGTTTGGCGTGAAACCATTTGTGGTGAAGCATTTCGTTGTGTTATTGCAGAAGCACAGTTGGACCCTGTAACCCTGACCCAACTGAA
AGATCAGTTTATGGAACGTCGTCGTGAGATACCGAAAAAACTGGTTGAAGATGCCATTAGCAATGGTGAACTGCCGAAAGATATCAATCGTGAACTGCTG
CTGGATATGATTTTTGGTTTTTGTTGGTATCGCCTGCTGACCGAACAGTTGACCGTTGAACAGGATATTGAAGAATTTACCTTCCTGCTGATTAATGGTG
TTTGTCCGGGTACACAGTGTTGATAAGGATCCCTCGGTACCAAAAAAAAAAAAAAAAGACGCTGAAAAGCGTCTTTTTTCGTTTTGGTCCGGCGTCTCCAC
TGTGCGCGGAACCCCTATTTGTTTATTTTTCTAAATACATTCAAATATGTATCCGCTCATGAGACAATAACCCTGATAAATGCTTCAATAATATTGAAAA
AGGAAGAGTATGAGTATTCAACATTTCCGTGTCGCCCTTATTCCCTTTTTTGCGGCATTTTGCCTTCCTGTTTTTGCTCACCCAGAAACGCTGGTGAAAG
TAAAAGATGCTGAAGATCAGTTGGGTGCACGAGTGGGTTACATCGAACTGGATCTCAACAGCGGTAAGATCCTTGAGAGTTTTCGCCCCGAAGAACGTTT
TCCAATGATGAGCACTTTTAAAGTTCTGCTATGTGGCGCGGTATTATCCCGTATTGACGCCGGGCAAGAGCAACTCGGTCGCCGCATACACTATTCTCAG
AATGACTTGGTTGAGTACTCACCAGTCACAGAAAAGCATCTTACGGATGGCATGACAGTAAGAGAATTATGCAGTGCTGCCATAACCATGAGTGATAACA
CTGCGGCCAACTTACTTCTGACAACGATCGGAGGACCGAAGGAGCTAACCGCTTTTTTGCACAACATGGGGGATCATGTAACTCGCCTTGATCGTTGGGA
ACCGGAGCTGAATGAAGCCATACCAAACGACGAGCGTGACACCACGATGCCTGTAGCAATGGCAACAACGTTGCGCAAACTATTAACTGGCGAACTACTT
ACTCTAGCTTCCCGGCAACAATTAATAGACTGGATGGAGGCGGATAAAGTTGCAGGACCACTTCTGCGCTCGGCCCTTCCGGCTGGCTGGTTTATTGCTG
ATAAATCTGGAGCCGGTGAGCGTGGGTCTCGCGGTATCATTGCAGCACTGGGGCCAGATGGTAAGCCCTCCCGTATCGTAGTTATCTACACGACGGGGAG
TCAGGCAACTATGGATGAACGAAATAGACAGATCGCTGAGATAGGTGCCTCACTGATTAAGCATTGGTAACTCGAGCTGATCCTTCAAGAGCTCGCTTGG
ACTCCTGTTGATAGATCCAGTAATGACCTCAGAACTCCATCTGGATTTGTTCAGAACGCTCGGTTGCCGCCGGGCGTTTTTTATTGGTGAGAATTTACAA
CTTATATCGTATGGGGCTGACTTCAGGTGCTACATTTGAAGAGATAAATTGCACTGAAATCTAGAAATATTTTATCTGATTAATAAGATGATCTTCTTGA
GATCGTTTTGGTCTGCGCGTAATCTCTTGCTCTGAAAACGAAAAAACCGCCTTGCAGGGCGGTTTTTCGAAGGTTCTCTGAGCTACCAACTCTTTGAACC
GAGGTAACTGGCTTGGAGGAGCGCAGTCACCAAAACTTGTCCTTTCAGTTTAGCCTTAACCGGCGCATGACTTCAAGACTAACTCCTCTAAATCAATTAC
CAGTGGCTGCTGCCAGTGGTGCTTTTGCATGTCTTTCCGGGTTGGACTCAAGACGATAGTTACCGGATAAGGCGCAGCGGTCGGACTGAACGGGGGGTTC
GTGCATACAGTCCAGCTTGGAGCGAACTGCCTACCCGGAACTGAGTGTCAGGCGTGGAATGAGACAAACGCGGCCATAACAGCGGAATGACACCGGTAAA
CCGAAAGGCAGGAACAGGAGAGCGCACGAGGGAGCCGCCAGGGGGAAACGCCTGGTATCTTTATAGTCCTGTCGGGTTTCGCCACCACTGATTTGAGCGT
CAGATT

**pCM193**

TCGTGATGCTTGTCAGGGGGGCGGAGCCTATGGAAAAACGGCTTTGCCGCGGCCCTCTCACTTCCCTGTTAAGTATCTTCCTGGCATCTTCCAGGAAATC
TCCGCCCCGTTCGTAAGCCATTTCCGCTCGCCGCAGTCGAACGTACTAGAGGGAATCGTACGGTCGATCCATCTGACTATCGCCTACTTCATTGACAGCT
AGCTCAGTCCTAGGTATAATGCTAGCTCTAGAGAAAGAGGAGAAATACTAGATGGCTTCCTCCGAAGACGTTATCAAAGAGTTCATGCGTTTCAAAGTTC
GTATGGAAGGTTCCGTTAACGGTCACGAGTTCGAAATCGAAGGTGAAGGTGAAGGTCGTCCGTACGAAGGTACCCAGACCGCTAAACTGAAAGTTACCAA
AGGTGGTCCGCTGCCGTTCGCTTGGGACATCCTGTCCCCGCAGTTCCAGTACGGTTCCAAAGCTTACGTTAAACACCCGGCTGACATCCCGGACTACCTG
AAACTGTCCTTCCCGGAAGGTTTCAAATGGGAACGTGTTATGAACTTCGAAGACGGTGGTGTTGTTACCGTTACCCAGGACTCCTCCCTGCAAGACGGTG
AGTTCATCTACAAAGTTAAACTGCGTGGTACCAACTTCCCGTCCGACGGTCCGGTTATGCAGAAAAAAACCATGGGTTGGGAAGCTTCCACCGAACGTAT
GTACCCGGAAGACGGTGCTCTGAAAGGTGAAATCAAAATGCGTCTGAAACTGAAAGACGGTGGTCACTACGACGCTGAAGTTAAAACCACCTACATGGCT
AAAAAACCGGTTCAGCTGCCGGGTGCTTACAAAACCGACATCAAACTGGACATCACCTCCCACAACGAAGACTACACCATCGTTGAACAGTACGAACGTG
CTGAAGGTCGTCACTCCACCGGTGCTTAATAACGCTGATAGTGCTAGTGTAGATCGCTACTAGAGTACTAGCTCTAGATGCGGCCGCGAATTCCAGAAAT
CATCCTTAGCGAAAGCTAAGGATTTTTTTTATCTGAAATGCCTCTGTAACAGAGCATTAGCGCAAGGTGATTTTTGTCTTCTTGCGCTAATTTTTTGGTC
CAACTTAGGCACCAATTATTGAAGGCCTCCCTAACGGGGGGCCTTTTTTTGTTTCTGGTCTCCCGCTTAACGATCGTTGGCTGCGACGTACGGTGGAATC
TGATTCGTTACCAATTGACATGATACGAAACGTACCGTATCGTTAAGGTAGCTGTCACCGGATGTGCTTTCCGGTCTGATGAGTCCGTGAGGACGAAACA

GCCTCTACAAATAATTTTGTTTAATACTAGAGAAAGAGGGGAAATACTAGATGGTGAGCAAGGGCGAGGAGCTGTTCACCGGGGTGGTGCCCATCCTGGT
CGAGCTGGACGGCGACGTAAACGGCCACAAGTTCAGCGTGTCCGGCGAGGGCGAGGGCGATGCCACCTACGGCAAGCTGACCCTGAAGTTCATCTGCACC
ACAGGCAAGCTGCCCGTGCCCTGGCCCACCCTCGTGACCACCTTCGGCTACGGCCTGCAATGCTTCGCCCGCTACCCCGACCACATGAAGCTGCACGACT
TCTTCAAGTCCGCCATGCCCGAAGGCTACGTCCAGGAGCGCACCATCTTCTTCAAGGACGACGGCAACTACAAGACCCGCGCCGAGGTGAAGTTCGAGGG
CGACACCCTGGTGAACCGCATCGAGCTGAAGGGCATCGACTTCAAGGAGGACGGCAACATCCTGGGGCACAAGCTGGAGTACAACTACAACAGCCACAAC
GTCTATATCATGGCCGACAAGCAGAAGAACGGCATCAAGGTGAACTTCAAGATCCGCCACAACATCGAGGACGGCAGCGTGCAGCTCGCCGACCACTACC
AGCAGAACACCCCAATCGGCGACGGCCCCGTGCTGCTGCCCGACAACCACTACCTTAGCTACCAGTCCGCCCTGAGCAAAGACCCCAACGAGAAGCGCGA
TCACATGGTCCTGCTGGAGTTCGTGACCGCCGCCGGGATCACTCTCGGCATGGACGAGCTGTACAAGTAACTCGGTACCAAATTCCAGAAAAGAGGCCTC
CCGAAAGGGGGGCCTTTTTTCGTTTTGGTCCAATGGCGGCGCGCCATCGAATGGTGCAAAACCTTTCGCGGTATGGCATGATAGCGCCCGGAAGAGAGTC
AATTCATGGGGGTGAATATGGCACGTACCCCGAGCCGTAGCAGCATTGGTAGCCTGCGTAGTCCGCATACCCATAAAGCAATTCTGACCAGCACCATTGA
AATCCTGAAAGAATGTGGTTATAGCGGTCTGAGCATTGAAAGCGTGGCACGTCGCGCCGGTGCAGGCAAACCGACCATTTATCGTTGGTGGACCAACAAA
GCAGCACTGATTGCCGAAGTGTATGAAAATGAAATCGAACAGGTACGTAAATTTCCGGATTTGGGTAGCTTTAAAGCCGATCTGGATTTTCTGCTGCATA
ATCTGTGGAAAGTTTGGCGTGAAACCATTTGTGGTGAAGCATTTCGTTGTGTTATTGCAGAAGCACAGTTGGACCCTGTAACCCTGACCCAACTGAAAGA
TCAGTTTATGGAACGTCGTCGTGAGATACCGAAAAAACTGGTTGAAGATGCCATTAGCAATGGTGAACTGCCGAAAGATATCAATCGTGAACTGCTGCTG
GATATGATTTTTGGTTTTTGTTGGTATCGCCTGCTGACCGAACAGTTGACCGTTGAACAGGATATTGAAGAATTTACCTTCCTGCTGATTAATGGTGTTT
GTCCGGGTACACAGTGTTGATAAGGATCCCTCGGTACCAAAAAAAAAAAAAAAAGACGCTGAAAAGCGTCTTTTTTCGTTTTGGTCCGGCGTCTCCACTGT
GCGCGGAACCCCTATTTGTTTATTTTTCTAAATACATTCAAATATGTATCCGCTCATGAGACAATAACCCTGATAAATGCTTCAATAATATTGAAAAAGG
AAGAGTATGAGTATTCAACATTTCCGTGTCGCCCTTATTCCCTTTTTTGCGGCATTTTGCCTTCCTGTTTTTGCTCACCCAGAAACGCTGGTGAAAGTAA
AAGATGCTGAAGATCAGTTGGGTGCACGAGTGGGTTACATCGAACTGGATCTCAACAGCGGTAAGATCCTTGAGAGTTTTCGCCCCGAAGAACGTTTTCC
AATGATGAGCACTTTTAAAGTTCTGCTATGTGGCGCGGTATTATCCCGTATTGACGCCGGGCAAGAGCAACTCGGTCGCCGCATACACTATTCTCAGAAT
GACTTGGTTGAGTACTCACCAGTCACAGAAAAGCATCTTACGGATGGCATGACAGTAAGAGAATTATGCAGTGCTGCCATAACCATGAGTGATAACACTG
CGGCCAACTTACTTCTGACAACGATCGGAGGACCGAAGGAGCTAACCGCTTTTTTGCACAACATGGGGGATCATGTAACTCGCCTTGATCGTTGGGAACC
GGAGCTGAATGAAGCCATACCAAACGACGAGCGTGACACCACGATGCCTGTAGCAATGGCAACAACGTTGCGCAAACTATTAACTGGCGAACTACTTACT
CTAGCTTCCCGGCAACAATTAATAGACTGGATGGAGGCGGATAAAGTTGCAGGACCACTTCTGCGCTCGGCCCTTCCGGCTGGCTGGTTTATTGCTGATA
AATCTGGAGCCGGTGAGCGTGGGTCTCGCGGTATCATTGCAGCACTGGGGCCAGATGGTAAGCCCTCCCGTATCGTAGTTATCTACACGACGGGGAGTCA
GGCAACTATGGATGAACGAAATAGACAGATCGCTGAGATAGGTGCCTCACTGATTAAGCATTGGTAACTCGAGCTGATCCTTCAAGAGCTCGCTTGGACT
CCTGTTGATAGATCCAGTAATGACCTCAGAACTCCATCTGGATTTGTTCAGAACGCTCGGTTGCCGCCGGGCGTTTTTTATTGGTGAGAATTTACAACTT
ATATCGTATGGGGCTGACTTCAGGTGCTACATTTGAAGAGATAAATTGCACTGAAATCTAGAAATATTTTATCTGATTAATAAGATGATCTTCTTGAGAT
CGTTTTGGTCTGCGCGTAATCTCTTGCTCTGAAAACGAAAAAACCGCCTTGCAGGGCGGTTTTTCGAAGGTTCTCTGAGCTACCAACTCTTTGAACCGAG
GTAACTGGCTTGGAGGAGCGCAGTCACCAAAACTTGTCCTTTCAGTTTAGCCTTAACCGGCGCATGACTTCAAGACTAACTCCTCTAAATCAATTACCAG
TGGCTGCTGCCAGTGGTGCTTTTGCATGTCTTTCCGGGTTGGACTCAAGACGATAGTTACCGGATAAGGCGCAGCGGTCGGACTGAACGGGGGGTTCGTG
CATACAGTCCAGCTTGGAGCGAACTGCCTACCCGGAACTGAGTGTCAGGCGTGGAATGAGACAAACGCGGCCATAACAGCGGAATGACACCGGTAAACCG
AAAGGCAGGAACAGGAGAGCGCACGAGGGAGCCGCCAGGGGGAAACGCCTGGTATCTTTATAGTCCTGTCGGGTTTCGCCACCACTGATTTGAGCGTCAG
ATT

**pCM194**

TCGTGATGCTTGTCAGGGGGGCGGAGCCTATGGAAAAACGGCTTTGCCGCGGCCCTCTCACTTCCCTGTTAAGTATCTTCCTGGCATCTTCCAGGAAATC
TCCGCCCCGTTCGTAAGCCATTTCCGCTCGCCGCAGTCGAACGTACTAGAGGGAATCGTACGGTCGATCCATCTGACTATCGCCTACTTCATTGACAGCT
AGCTCAGTCCTAGGTATAATGCTAGCAAATATGTTGTAGAAACAATTGCACGAATGGCTTCCTCCGAAGACGTTATCAAAGAGTTCATGCGTTTCAAAGT
TCGTATGGAAGGTTCCGTTAACGGTCACGAGTTCGAAATCGAAGGTGAAGGTGAAGGTCGTCCGTACGAAGGTACCCAGACCGCTAAACTGAAAGTTACC
AAAGGTGGTCCGCTGCCGTTCGCTTGGGACATCCTGTCCCCGCAGTTCCAGTACGGTTCCAAAGCTTACGTTAAACACCCGGCTGACATCCCGGACTACC
TGAAACTGTCCTTCCCGGAAGGTTTCAAATGGGAACGTGTTATGAACTTCGAAGACGGTGGTGTTGTTACCGTTACCCAGGACTCCTCCCTGCAAGACGG
TGAGTTCATCTACAAAGTTAAACTGCGTGGTACCAACTTCCCGTCCGACGGTCCGGTTATGCAGAAAAAAACCATGGGTTGGGAAGCTTCCACCGAACGT
ATGTACCCGGAAGACGGTGCTCTGAAAGGTGAAATCAAAATGCGTCTGAAACTGAAAGACGGTGGTCACTACGACGCTGAAGTTAAAACCACCTACATGG
CTAAAAAACCGGTTCAGCTGCCGGGTGCTTACAAAACCGACATCAAACTGGACATCACCTCCCACAACGAAGACTACACCATCGTTGAACAGTACGAACG
TGCTGAAGGTCGTCACTCCACCGGTGCTTAATAACGCTGATAGTGCTAGTGTAGATCGCTACTAGAGTACTAGCTCTAGATGCGGCCGCGAATTCCAGAA
ATCATCCTTAGCGAAAGCTAAGGATTTTTTTTATCTGAAATGCCTCTGTAACAGAGCATTAGCGCAAGGTGATTTTTGTCTTCTTGCGCTAATTTTTTGG
TCCAACTTAGGCACCAATTATTGAAGGCCTCCCTAACGGGGGGCCTTTTTTTGTTTCTGGTCTCCCGCTTAACGATCGTTGGCTGCGACGTACGGTGGAA
TCTGATTCGTTACCAATTGACATGATACGAAACGTACCGTATCGTTAAGGTAGCTGTCACCGGATGTGCTTTCCGGTCTGATGAGTCCGTGAGGACGAAA
CAGCCTCTACAAATAATTTTGTTTAATACTAGAGAAAGAGGGGAAATACTAGATGGTGAGCAAGGGCGAGGAGCTGTTCACCGGGGTGGTGCCCATCCTG
GTCGAGCTGGACGGCGACGTAAACGGCCACAAGTTCAGCGTGTCCGGCGAGGGCGAGGGCGATGCCACCTACGGCAAGCTGACCCTGAAGTTCATCTGCA
CCACAGGCAAGCTGCCCGTGCCCTGGCCCACCCTCGTGACCACCTTCGGCTACGGCCTGCAATGCTTCGCCCGCTACCCCGACCACATGAAGCTGCACGA
CTTCTTCAAGTCCGCCATGCCCGAAGGCTACGTCCAGGAGCGCACCATCTTCTTCAAGGACGACGGCAACTACAAGACCCGCGCCGAGGTGAAGTTCGAG
GGCGACACCCTGGTGAACCGCATCGAGCTGAAGGGCATCGACTTCAAGGAGGACGGCAACATCCTGGGGCACAAGCTGGAGTACAACTACAACAGCCACA
ACGTCTATATCATGGCCGACAAGCAGAAGAACGGCATCAAGGTGAACTTCAAGATCCGCCACAACATCGAGGACGGCAGCGTGCAGCTCGCCGACCACTA
CCAGCAGAACACCCCAATCGGCGACGGCCCCGTGCTGCTGCCCGACAACCACTACCTTAGCTACCAGTCCGCCCTGAGCAAAGACCCCAACGAGAAGCGC
GATCACATGGTCCTGCTGGAGTTCGTGACCGCCGCCGGGATCACTCTCGGCATGGACGAGCTGTACAAGTAACTCGGTACCAAATTCCAGAAAAGAGGCC
TCCCGAAAGGGGGGCCTTTTTTCGTTTTGGTCCAATGGCGGCGCGCCATCGAATGGTGCAAAACCTTTCGCGGTATGGCATGATAGCGCCCGGAAGAGAG
TCAATTCATGGGGGTGAATATGGCACGTACCCCGAGCCGTAGCAGCATTGGTAGCCTGCGTAGTCCGCATACCCATAAAGCAATTCTGACCAGCACCATT
GAAATCCTGAAAGAATGTGGTTATAGCGGTCTGAGCATTGAAAGCGTGGCACGTCGCGCCGGTGCAGGCAAACCGACCATTTATCGTTGGTGGACCAACA
AAGCAGCACTGATTGCCGAAGTGTATGAAAATGAAATCGAACAGGTACGTAAATTTCCGGATTTGGGTAGCTTTAAAGCCGATCTGGATTTTCTGCTGCA
TAATCTGTGGAAAGTTTGGCGTGAAACCATTTGTGGTGAAGCATTTCGTTGTGTTATTGCAGAAGCACAGTTGGACCCTGTAACCCTGACCCAACTGAAA
GATCAGTTTATGGAACGTCGTCGTGAGATACCGAAAAAACTGGTTGAAGATGCCATTAGCAATGGTGAACTGCCGAAAGATATCAATCGTGAACTGCTGC
TGGATATGATTTTTGGTTTTTGTTGGTATCGCCTGCTGACCGAACAGTTGACCGTTGAACAGGATATTGAAGAATTTACCTTCCTGCTGATTAATGGTGT
TTGTCCGGGTACACAGTGTTGATAAGGATCCCTCGGTACCAAAAAAAAAAAAAAAAGACGCTGAAAAGCGTCTTTTTTCGTTTTGGTCCGGCGTCTCCACT
GTGCGCGGAACCCCTATTTGTTTATTTTTCTAAATACATTCAAATATGTATCCGCTCATGAGACAATAACCCTGATAAATGCTTCAATAATATTGAAAAA
GGAAGAGTATGAGTATTCAACATTTCCGTGTCGCCCTTATTCCCTTTTTTGCGGCATTTTGCCTTCCTGTTTTTGCTCACCCAGAAACGCTGGTGAAAGT
AAAAGATGCTGAAGATCAGTTGGGTGCACGAGTGGGTTACATCGAACTGGATCTCAACAGCGGTAAGATCCTTGAGAGTTTTCGCCCCGAAGAACGTTTT
CCAATGATGAGCACTTTTAAAGTTCTGCTATGTGGCGCGGTATTATCCCGTATTGACGCCGGGCAAGAGCAACTCGGTCGCCGCATACACTATTCTCAGA
ATGACTTGGTTGAGTACTCACCAGTCACAGAAAAGCATCTTACGGATGGCATGACAGTAAGAGAATTATGCAGTGCTGCCATAACCATGAGTGATAACAC
TGCGGCCAACTTACTTCTGACAACGATCGGAGGACCGAAGGAGCTAACCGCTTTTTTGCACAACATGGGGGATCATGTAACTCGCCTTGATCGTTGGGAA
CCGGAGCTGAATGAAGCCATACCAAACGACGAGCGTGACACCACGATGCCTGTAGCAATGGCAACAACGTTGCGCAAACTATTAACTGGCGAACTACTTA
CTCTAGCTTCCCGGCAACAATTAATAGACTGGATGGAGGCGGATAAAGTTGCAGGACCACTTCTGCGCTCGGCCCTTCCGGCTGGCTGGTTTATTGCTGA
TAAATCTGGAGCCGGTGAGCGTGGGTCTCGCGGTATCATTGCAGCACTGGGGCCAGATGGTAAGCCCTCCCGTATCGTAGTTATCTACACGACGGGGAGT
CAGGCAACTATGGATGAACGAAATAGACAGATCGCTGAGATAGGTGCCTCACTGATTAAGCATTGGTAACTCGAGCTGATCCTTCAAGAGCTCGCTTGGA
CTCCTGTTGATAGATCCAGTAATGACCTCAGAACTCCATCTGGATTTGTTCAGAACGCTCGGTTGCCGCCGGGCGTTTTTTATTGGTGAGAATTTACAAC
TTATATCGTATGGGGCTGACTTCAGGTGCTACATTTGAAGAGATAAATTGCACTGAAATCTAGAAATATTTTATCTGATTAATAAGATGATCTTCTTGAG
ATCGTTTTGGTCTGCGCGTAATCTCTTGCTCTGAAAACGAAAAAACCGCCTTGCAGGGCGGTTTTTCGAAGGTTCTCTGAGCTACCAACTCTTTGAACCG
AGGTAACTGGCTTGGAGGAGCGCAGTCACCAAAACTTGTCCTTTCAGTTTAGCCTTAACCGGCGCATGACTTCAAGACTAACTCCTCTAAATCAATTACC
AGTGGCTGCTGCCAGTGGTGCTTTTGCATGTCTTTCCGGGTTGGACTCAAGACGATAGTTACCGGATAAGGCGCAGCGGTCGGACTGAACGGGGGGTTCG

TGCATACAGTCCAGCTTGGAGCGAACTGCCTACCCGGAACTGAGTGTCAGGCGTGGAATGAGACAAACGCGGCCATAACAGCGGAATGACACCGGTAAAC
CGAAAGGCAGGAACAGGAGAGCGCACGAGGGAGCCGCCAGGGGGAAACGCCTGGTATCTTTATAGTCCTGTCGGGTTTCGCCACCACTGATTTGAGCGTC
AGATT

**pCM240**

TCGTGATGCTTGTCAGGGGGGCGGAGCCTATGGAAAAACGGCTTTGCCGCGGCCCTCTCACTTCCCTGTTAAGTATCTTCCTGGCATCTTCCAGGAAATC
TCCGCCCCGTTCGTAAGCCATTTCCGCTCGCCGCAGTCGAACGGCTTAACGATCGTTGGCTGCGACGTACGGTGGAATCTGATTCGTTACCAATTGACAT
GATACGAAACGTACCGTATCGTTAAGGTAGCTGTCACCGGATGTGCTTTCCGGTCTGATGAGTCCGTGAGGACGAAACAGCCTCTACAAATAATTTTGTT
TAATACTAGAGAAAGAGGGGAAATACTAGATGGTGAGCAAGGGCGAGGAGCTGTTCACCGGGGTGGTGCCCATCCTGGTCGAGCTGGACGGCGACGTAAA
CGGCCACAAGTTCAGCGTGTCCGGCGAGGGCGAGGGCGATGCCACCTACGGCAAGCTGACCCTGAAGTTCATCTGCACCACAGGCAAGCTGCCCGTGCCC
TGGCCCACCCTCGTGACCACCTTCGGCTACGGCCTGCAATGCTTCGCCCGCTACCCCGACCACATGAAGCTGCACGACTTCTTCAAGTCCGCCATGCCCG
AAGGCTACGTCCAGGAGCGCACCATCTTCTTCAAGGACGACGGCAACTACAAGACCCGCGCCGAGGTGAAGTTCGAGGGCGACACCCTGGTGAACCGCAT
CGAGCTGAAGGGCATCGACTTCAAGGAGGACGGCAACATCCTGGGGCACAAGCTGGAGTACAACTACAACAGCCACAACGTCTATATCATGGCCGACAAG
CAGAAGAACGGCATCAAGGTGAACTTCAAGATCCGCCACAACATCGAGGACGGCAGCGTGCAGCTCGCCGACCACTACCAGCAGAACACCCCAATCGGCG
ACGGCCCCGTGCTGCTGCCCGACAACCACTACCTTAGCTACCAGTCCGCCCTGAGCAAAGACCCCAACGAGAAGCGCGATCACATGGTCCTGCTGGAGTT
CGTGACCGCCGCCGGGATCACTCTCGGCATGGACGAGCTGTACAAGTAACTCGGTACCAAATTCCAGAAAAGAGGCCTCCCGAAAGGGGGGCCTTTTTTC
GTTTTGGTCCAATGGCGGCGCGCCATCGAATGGTGCAAAACCTTTCGCGGTATGGCATGATAGCGCCCGGAAGAGAGTCAATTCATGGGGGTGAATATGG
CACGTACCCCGAGCCGTAGCAGCATTGGTAGCCTGCGTAGTCCGCATACCCATAAAGCAATTCTGACCAGCACCATTGAAATCCTGAAAGAATGTGGTTA
TAGCGGTCTGAGCATTGAAAGCGTGGCACGTCGCGCCGGTGCAGGCAAACCGACCATTTATCGTTGGTGGACCAACAAAGCAGCACTGATTGCCGAAGTG
TATGAAAATGAAATCGAACAGGTACGTAAATTTCCGGATTTGGGTAGCTTTAAAGCCGATCTGGATTTTCTGCTGCATAATCTGTGGAAAGTTTGGCGTG
AAACCATTTGTGGTGAAGCATTTCGTTGTGTTATTGCAGAAGCACAGTTGGACCCTGTAACCCTGACCCAACTGAAAGATCAGTTTATGGAACGTCGTCG
TGAGATACCGAAAAAACTGGTTGAAGATGCCATTAGCAATGGTGAACTGCCGAAAGATATCAATCGTGAACTGCTGCTGGATATGATTTTTGGTTTTTGT
TGGTATCGCCTGCTGACCGAACAGTTGACCGTTGAACAGGATATTGAAGAATTTACCTTCCTGCTGATTAATGGTGTTTGTCCGGGTACACAGTGTTGAT
AAGGATCCCTCGGTACCAAAAAAAAAAAAAAAAAGACGCTGAAAAGCGTCTTTTTTCGTTTTGGTCCGGCGTCTCCACTGTGCGCGGAACCCCTATTTGTTT
ATTTTTCTAAATACATTCAAATATGTATCCGCTCATGAGACAATAACCCTGATAAATGCTTCAATAATATTGAAAAAGGAAGAGTATGAGTATTCAACAT
TTCCGTGTCGCCCTTATTCCCTTTTTTGCGGCATTTTGCCTTCCTGTTTTTGCTCACCCAGAAACGCTGGTGAAAGTAAAAGATGCTGAAGATCAGTTGG
GTGCACGAGTGGGTTACATCGAACTGGATCTCAACAGCGGTAAGATCCTTGAGAGTTTTCGCCCCGAAGAACGTTTTCCAATGATGAGCACTTTTAAAGT
TCTGCTATGTGGCGCGGTATTATCCCGTATTGACGCCGGGCAAGAGCAACTCGGTCGCCGCATACACTATTCTCAGAATGACTTGGTTGAGTACTCACCA
GTCACAGAAAAGCATCTTACGGATGGCATGACAGTAAGAGAATTATGCAGTGCTGCCATAACCATGAGTGATAACACTGCGGCCAACTTACTTCTGACAA
CGATCGGAGGACCGAAGGAGCTAACCGCTTTTTTGCACAACATGGGGGATCATGTAACTCGCCTTGATCGTTGGGAACCGGAGCTGAATGAAGCCATACC
AAACGACGAGCGTGACACCACGATGCCTGTAGCAATGGCAACAACGTTGCGCAAACTATTAACTGGCGAACTACTTACTCTAGCTTCCCGGCAACAATTA
ATAGACTGGATGGAGGCGGATAAAGTTGCAGGACCACTTCTGCGCTCGGCCCTTCCGGCTGGCTGGTTTATTGCTGATAAATCTGGAGCCGGTGAGCGTG
GGTCTCGCGGTATCATTGCAGCACTGGGGCCAGATGGTAAGCCCTCCCGTATCGTAGTTATCTACACGACGGGGAGTCAGGCAACTATGGATGAACGAAA
TAGACAGATCGCTGAGATAGGTGCCTCACTGATTAAGCATTGGTAACTCGAGCTGATCCTTCAAGAGCTCGCTTGGACTCCTGTTGATAGATCCAGTAAT
GACCTCAGAACTCCATCTGGATTTGTTCAGAACGCTCGGTTGCCGCCGGGCGTTTTTTATTGGTGAGAATTTACAACTTATATCGTATGGGGCTGACTTC
AGGTGCTACATTTGAAGAGATAAATTGCACTGAAATCTAGAAATATTTTATCTGATTAATAAGATGATCTTCTTGAGATCGTTTTGGTCTGCGCGTAATC
TCTTGCTCTGAAAACGAAAAAACCGCCTTGCAGGGCGGTTTTTCGAAGGTTCTCTGAGCTACCAACTCTTTGAACCGAGGTAACTGGCTTGGAGGAGCGC
AGTCACCAAAACTTGTCCTTTCAGTTTAGCCTTAACCGGCGCATGACTTCAAGACTAACTCCTCTAAATCAATTACCAGTGGCTGCTGCCAGTGGTGCTT
TTGCATGTCTTTCCGGGTTGGACTCAAGACGATAGTTACCGGATAAGGCGCAGCGGTCGGACTGAACGGGGGGTTCGTGCATACAGTCCAGCTTGGAGCG
AACTGCCTACCCGGAACTGAGTGTCAGGCGTGGAATGAGACAAACGCGGCCATAACAGCGGAATGACACCGGTAAACCGAAAGGCAGGAACAGGAGAGCG
CACGAGGGAGCCGCCAGGGGGAAACGCCTGGTATCTTTATAGTCCTGTCGGGTTTCGCCACCACTGATTTGAGCGTCAGATT

Gold indicates promoters, green indicates ribosome binding sites, red indicates coding sequences, blue indicates terminators, purple indicates ribozyme insulators, and black indicates the plasmid backbone and spacers.
